# OmicOS: A Comprehensive Omics Ecosystem Infrastructure and Agent System for the AI Era

**DOI:** 10.64898/2026.06.11.731775

**Authors:** Zehua Zeng, Xu Meng, Lei Hu, Chen Li, Peng Liu, Yu Shi, Xuejiao Ma, Lianchong Gao, Xuehai Wang, Zhi Luo, Yawen Zheng, Jieshen Xian, Ziheng Lin, Haoliang Zhu, Zhaorui Jiang, Sheng Mao, Yifan Lu, Wenzhuo Tang, Qiangwei Peng, Yuqing Ma, Liping Zhou, Cencan Xing, Xuegong Zhang, Yuanyan Xiong, Hongwu Du

**Affiliations:** School of Chemistry and Biological Engineering, University of Science and Technology Beijing, Beijing 100083, China; Daxing Research Institute, University of Science and Technology Beijing, Beijing 100083, China; Department of Genetics, Stanford University, Stanford, CA 94305, USA; Department of Biochemistry, Key Laboratory of Gene Engineering of the Ministry of Education, School of Life Sciences, Sun Yat-sen University, Guangzhou 510275, China; College of Informatics, Hubei Key Laboratory of Agricultural Bioinformatics, Huazhong Agricultural University, Wuhan 430070, China; School of Life Sciences, Westlake University, Hangzhou 310030, China; MOE Key Laboratory of Bioinformatics, Bioinformatics Division of BNRIST (Beijing National Research Center for Information Science and Technology), Department of Automation, Tsinghua University, Beijing 100084, China; School of Life Sciences, Tsinghua University, Beijing 100084, China; Department of Surgery of Traditional Chinese Medicine, China-Japan Friendship Hospital, Beijing 100029, China; Key Laboratory of Systems Biomedicine (Ministry of Education), Shanghai Center for Systems Biomedicine, Shanghai Jiao Tong University, Shanghai 200240, China; Department of Physiology and Pharmacology, Karolinska Institutet, Stockholm SE-171 77, Sweden; School of Public Health, Guangzhou Medical University, Guangzhou 511436, China; School of Basic Medical Sciences, Capital Medical University, Beijing 100069, China; School of the First Clinical Medical Sciences, Wenzhou Medical University, Wenzhou 325000, China; School of Environment and Energy, Peking University Shenzhen Graduate School, Shenzhen 518055, China; Electronic Information School, Wuhan University, Wuhan 430072, China; Department of Statistics and Probability, Michigan State University, East Lansing, MI 48824, USA; School of Mathematical Sciences, Peking University, Beijing 100871, China; Center for Precision Medicine and Healthcare, Tsinghua-Berkeley Shenzhen Institute, Shenzhen 518055, China; Institute of Biopharmaceutical and Health Engineering, Tsinghua Shenzhen International Graduate School, Shenzhen 518055, China; Basic Science and Engineering Initiative, Betty Irene Moore Children’s Heart Center, Lucile Packard Children’s Hospital Stanford, Palo Alto, CA 94304, USA; Department of Artificial Intelligence, School of Engineering, Westlake University, Hangzhou 310030, China; Nurturing Center of Jiangsu Province for State Laboratory of AI Imaging and Interventional Radiology, Department of Radiology, Zhongda Hospital, School of Medicine, Southeast University, Nanjing 210009, China

## Abstract

Biology has accumulated a vast ecosystem of omics methods, but much of this ecosystem remains built for expert humans rather than scientific agents. Methods are scattered across Python packages, R/Bioconductor and CRAN workflows, command-line tools, incompatible data containers and implicit object states, making even routine analyses difficult for an AI system to choose, execute and verify reliably. Here we introduce OmicOS, a comprehensive omics ecosystem infrastructure and agent system that turns OmicVerse V2, an open-source omics community, into an executable foundation for agentic biology. OmicVerse V2 provides the community substrate: scalable AnnDataOOM-compatible rust backends, agent-friendly Python algorithms for single-cell, spatial, bulk and multi-omics analysis, interfaces to single-cell foundation models, and Python-native reconstructions of historically R-centred Bioconductor/CRAN-style workflows. OmicOS makes this substrate actionable by registering analytical functions as state-aware capability contracts, allowing agents to inspect live data objects, select valid methods, execute controlled workflows and record provenance. The result is not a fixed pipeline, but a programmable omics environment in which agents compose real analyses from verified community methods rather than inventing tools. Across external and purpose-built benchmarks, OmicOS ranked first among the evaluated systems, reaching 81.2% on BiomniBench. Adding OmicVerse to a minimal agent improved task completion by up to 34.2 percentage points with qwen-3.6-35b, and controlled ablations showed that the gains came from registry-grounded execution rather than from larger models, documentation retrieval or unrestricted tool exposure. The same infrastructure scaled to atlas-sized data, reproduced R-centred workflows in Python and converted external pathology software into agent-usable skills. In a discovery task starting from a whole-body spatial map and the term “Alzheimer’s disease”, OmicOS composed a non-canonical workflow that integrated spatial expression, genetic association, eQTL and colocalization evidence to nominate a colon epithelial risk axis centred on PICALM, CD2AP and CR1. Together, OmicVerse and OmicOS define an open foundation for AI-era omics, showing how a community of biological methods can be transformed into a reliable, extensible and agent-operable system for discovery.

**Highlight:**

- **OmicVerse 2.0** consolidates 694 methods spanning 11 omics domains into agent-callable high-level APIs.
- **RebuildR** automatically reconstructs and evolves R/Bioconductor methods as Python-native implementations under output-equivalence gates.
- **OmicOS** establishes a state-of-the-art omics agent harness, ranking first on general omics benchmarks across models and substantially improving the analytical capability of local open-source models.
- Compositional use of ecosystem modules nominates a colon epithelial axis associated with Alzheimer’s disease risk.
- External algorithm packages supporting automatic iterative evolution can be integrated into the OmicOS ecosystem.

## Main

Modern biomedical discovery has become multi-modal, spanning genomics, epigenomics, transcriptomics, proteomics, metabolomics, microbiome profiling, spatial measurements and single-cell atlases^1–5^. The computational response has been a powerful but fragmented ecosystem—Bioconductor^6^, Scanpy^7^ and AnnData^8^, scverse^9^, MuData/MUON^10^, Squidpy/SpatialData^11,12^ and OmicVerse^13^—with domain methods for differential expression, enrichment, microbiome and other assays scattered across incompatible languages, package conventions, data containers and dependency systems^14–16^, a fragmentation now compounded by single-cell and spatial foundation models with their own interfaces^17–19^. These workflows are intrinsically stateful: whether a method is applicable depends on whether raw counts, normalized matrices, highly variable genes, embeddings, neighbourhood graphs, batch covariates, spatial coordinates or prior annotations already exist on the object, and a bare function name conveys none of this, so the same ecosystem that empowers human experts is what makes the field hard to automate^7,10,20,21^. Scale compounds the problem: the dominant in-memory data model loads entire objects into RAM, so atlas-scale data increasingly exceeds what a single workstation can hold, making scalable yet API-compatible storage a prerequisite for any agent expected to operate on real datasets^109^

Large language model agents have changed how computational tasks are approached, coupling reasoning with action selection, tool use, environment interaction and iterative correction^22–24^. Multi-agent and software-engineering systems further show that realized capability depends strongly on the interface between the model and its execution environment, because repository navigation, file editing, testing and error feedback determine whether a model can complete a real task^25–27^. This insight has reached biomedicine through “AI scientist” systems that generate hypotheses, design experiments, write and run code, and draft manuscripts^28–32^, and recent benchmarks make one pattern explicit: realistic agent performance is inseparable from the dataset inspection, method selection, code execution and result interpretation that the environment must support^33–35^. A central bottleneck for AI-era omics is therefore not only whether a model can reason about biology, but whether it can act reliably inside a validated, state-aware execution space—suggesting that omics-agent capability should be treated as a property of a model–harness–ecosystem system rather than of the language model alone.

Existing infrastructure provides important pieces of this stack but does not yet jointly expose them as a bounded, state-aware action space over evolving omics objects. Workflow engines such as Snakemake^36,37^, Nextflow^36^ and Galaxy^38^ deliver strong provenance and scalability, yet were not designed as live biological action spaces for autonomous agents. Semantic standards and registries—FAIR^39^, EDAM^40^ and bio.tools^41^—make tools discoverable, but metadata alone cannot guarantee that a chosen method runs with valid arguments, satisfied prerequisites and recoverable failures. Two barriers are specific to omics. First, many canonical statistical pipelines remain in R and Bioconductor while modern agentic workflows are Python-centred, leaving a structural R/Python divide^6,7,14^. Second, naive reimplementation is unsafe, because biologically equivalent outputs can differ by cluster-label permutations, embedding rotations, branch orientations or pseudotime scaling, so correctness cannot be judged by byte-level identity^42–45^.

Here we introduce OmicOS (https://omicos.cn/) and OmicVerse V2 (https://github.com/omicverse), OmicOS is an agent-facing operating layer that exposes a reconstructed omics ecosystem as a state-aware, executable action space rather than asking agents to assemble one-off scripts from scattered documentation. OmicOS is organized as three layers. At the base, AnnDataOOM and MuDataOOM are Rust-engineered out-of-memory backends that hold only the working set in RAM while the remainder stays on disk; on a personal laptop they read and visualize a variable from a million-cell dataset in about one second, at speed parity with in-memory AnnData, while remaining fully API-compatible with AnnData and MuData. On this backend, OmicVerse v2 reconstructs and unifies methods across major omics domains—single-cell, spatial, bulk, multi-omics, foundation-model, molecular, epigenomic, immune-repertoire, proteomic, microbiome and metabolomic analysis, and rewrites the reconstructed omics preprocessing methods in full as GPU-accelerated implementations on PyTorch and Apple MLX backends (**Fig. 1a**). OmicVerse-RebuildR brings R-centred pipelines into Python by treating each R reference implementation as an executable specification, accepting a port only after output-class-aware equivalence is verified and then optimizing it under correctness-preserving constraints^42–44^. A complementary pipeline extends the same discipline to third-party Python tools beyond the native ecosystem: rather than wrapping them ad hoc, it iteratively distils a package into a portable agent-and-skill bundle through a reflection–edit–validate optimization loop^46^, admitting it only after rubric-based quality, API-grounding and cross-modality-fabrication gates pass. We illustrate this by porting LazySlide^47^ for direct haematoxylin-and-eosin (H&E) whole-slide analysis, where the resulting skill builds H&E-based prognostic survival readouts and chains into OmicVerse’s H&E gene-expression-prediction module to generate pseudo-spatial profiles from histology alone. At the top, OmicOS lifts these methods into explicit capability contracts that declare required and produced object states, parameter schemas, dependencies and side effects, coupled to a live runtime that validates prerequisites, updates biological state, records execution lineage and localizes failures.

**Figure 1.**
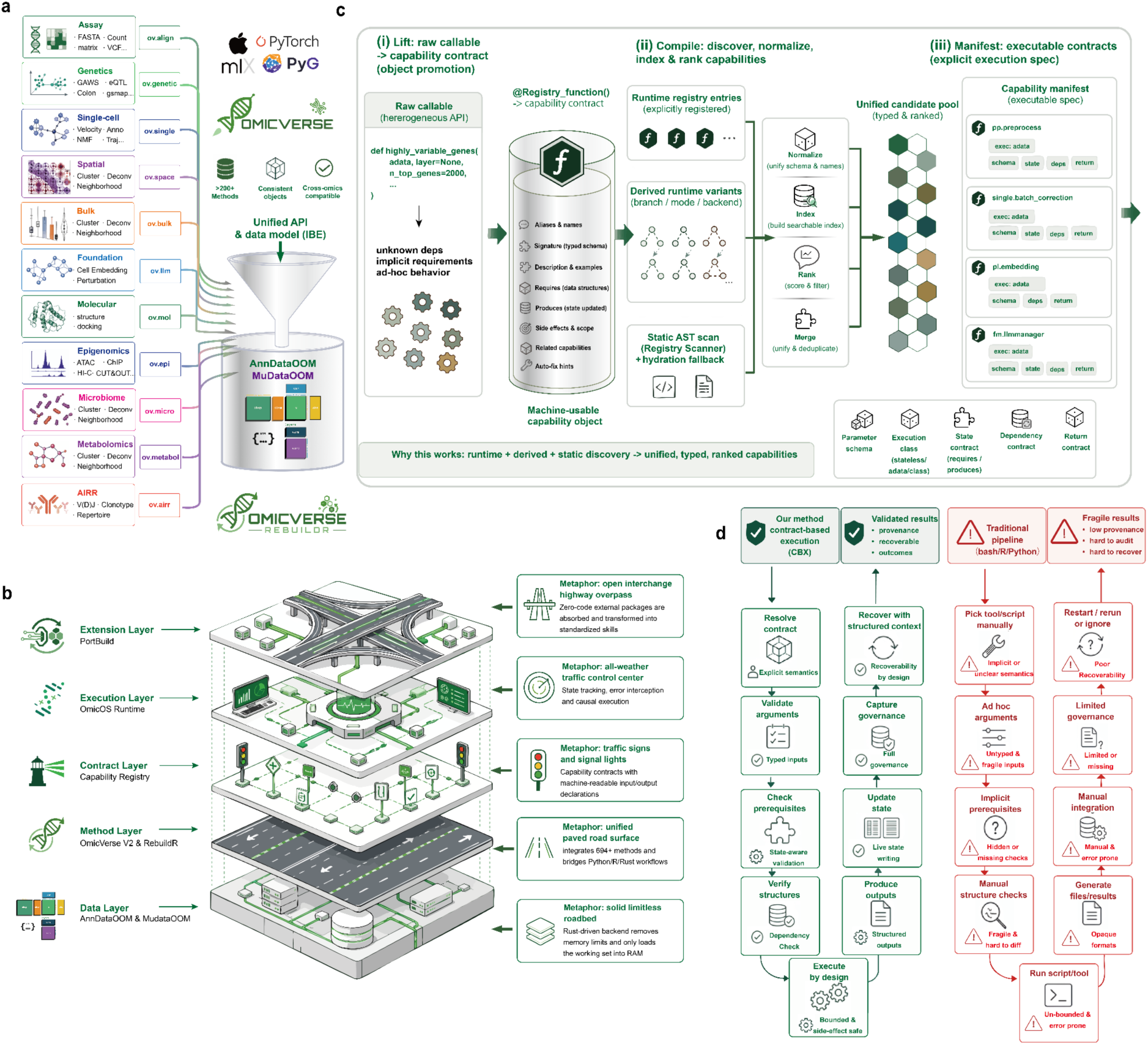
| OmicOS establishes an agent-friendly reconstruction ecosystem for reproducible multi-omics analysis. **a,** OmicOS is built on OmicVerse, a unified Python-native omics ecosystem that consolidates end-to-end pipelines across major data modalities—assay processing, preprocessing, statistical genetics, bulk and single-cell transcriptomics, spatial transcriptomics, single-cell foundation models, molecular analysis, epigenomics, immune-repertoire (AIRR), microbiome and metabolomics—exposed through consistent high-level ov.* APIs and a shared AnnData/MuData data model. A Rust-engineered out-of-memory backend (AnnDataOOM/MuDataOOM) and GPU/accelerator support (PyTorch, MLX, PyG) underlie the unified interface, and many historically R-centred workflows are reconstructed as Python-accessible implementations through OmicVerse-RebuildR (Fig. 3). **b,** Layered design of the OmicOS execution stack. AnnDataOOM/MuDataOOM provides scalable on-disk data storage; OmicVerse v2 and RebuildR unify Python and reconstructed R workflows; the capability registry converts methods into machine-readable contracts; the OmicOS runtime manages state, provenance and recoverable execution; and PortBuild imports external packages as standardized agent skills. **c,** Registered scientific functions are compiled into agent-executable capability contracts in three stages—lift, compile and manifest. Each raw callable is promoted into a structured, inspectable object encoding aliases and biological intent, accepted inputs, required object states (requires), parameters, dependencies and prerequisites, produced fields (produces) and an error policy (auto_fix). Runtime registration, static abstract-syntax-tree scanning and metadata normalization discover, deduplicate and rank these capabilities into a bounded, typed analytical action space for agent planning, validation and execution. **d,** Schematic comparison of OmicOS contract-based execution (left) and conventional script-based analysis (right). Script-based pipelines require manual resolution of package-specific syntax, dependency conflicts, object conversions, prerequisite checks and opaque failures, whereas the contract-grounded layer resolves, type-checks, validates, executes, traces and repairs each step within a reproducible runtime, yielding typed and recoverable failure states rather than untyped crashes

This design allowed us to evaluate agentic omics as a model–harness pair. Across two external task suites (BixBench-Verified-50 and BiomniBench) and a purpose-built benchmark (OmicBench), controlled ablations on a deliberately minimal agent showed that adding the execution layer—rather than enlarging the model—substantially improved task completion, with the largest gains in smaller or local models, narrowing the gap to closed-source frontier systems on omics tasks. Reconstruction proved faithful as well as practical, preserving benchmark-level behaviour while improving runtime over R references, and the composability of the ecosystem allowed OmicOS to assemble non-canonical workflows—demonstrated, as a proof-of-concept rather than a primary disease claim, by a whole-body spatial-genetic analysis that integrates spatially resolved genetic-risk mapping^48^, single-cell reference mapping, and human eQTL and colocalization evidence^49,50^ to nominate a colon epithelial axis of Alzheimer’s-disease genetic risk. Together, these results support a single conclusion: reliable agentic omics requires a state-aware execution substrate—from a scalable data backend through reconstruction-verified methods to contract-grounded execution—not merely a stronger language model.

## Results

### OmicOS converts a comprehensive omics ecosystem into an agent-executable analysis substrate

We built OmicOS on OmicVerse to give agents a single, unified substrate for omics analysis rather than a scatter of incompatible tools. Across 14 modules, OmicVerse exposes 694 analysis methods under one ov.* namespace and a shared AnnData/MuData data model, spanning assay processing, preprocessing, single-cell, spatial, bulk, foundation-model, molecular, epigenomic, microbiome, metabolomic and multi-omics analysis (**Fig. 1a**; **Supplementary Table 1**). Module sizes range from a large single-cell module (ov.single, 151 methods) and immune-repertoire module (ov.airr, 102) to a compact molecular one (ov.mol, 12), and more than 21 single-cell foundation models are interfaced, six with local runtimes (scGPT, Geneformer, scFoundation, CellPLM, UCE and scMulan). To avoid leaving canonical statistics stranded in R, we folded historically R-centred pipelines into the same Python-accessible infrastructure as 58 reconstructed R/Bioconductor/CRAN packages, each accepted against an output-class-aware equivalence criterion (38 with quantified parity metrics) and reachable through the same high-level interfaces (**Supplementary Table 2**).

A function name alone rarely tells an agent how to use it: it does not reveal whether the input must contain raw counts, normalized expression, highly variable genes, a neighbour graph, batch labels, spatial coordinates, embeddings or prior annotations. ov.single.scanpy_cellanno_from_dict, for instance, requires that Leiden clustering has already run and written a cluster column, a precondition absent from its signature (**Fig. 1b**). To make such requirements explicit, OmicOS lifts each callable into a machine-readable capability object encoding aliases, biological intent, a parameter schema, accepted inputs, required object states, dependency constraints, side effects and return structure, so that the hidden precondition becomes an explicit prerequisites = {“functions”: [“leiden”]} field and the produced annotation column is declared for downstream steps (**Fig. 1b**; **Supplementary Table 3**). Three mechanisms populate this registry—runtime registration through an import-time decorator, static abstract-syntax-tree scanning of unimported source, and metadata hydration that merges the two—and rank capabilities for a free-text task (**Fig. 1b**; **Supplementary Note 1**). Auditing the public API showed how breadth and execution-readiness relate: of the 694 exposed methods, 231 were promoted to capability objects across nine categories; all 231 are discoverable by intent, with a mean of 4.3 aliases each (including multilingual aliases such as 质控 and 手动注释) and runnable examples, while 45.9% (∼106 functions) additionally declare a full requires/produces/prerequisites contract, concentrated in the preprocessing and single-cell layers (**Supplementary Note 1**). The resulting manifest gives an agent a bounded action space of type- and state-aware operations rather than free-form code.

OmicOS connects these contracts to a live AnnData/MuData-centred runtime so that multi-step analyses proceed as traceable state transitions. Because downstream steps depend on whether counts have been normalized, embeddings computed, neighbour graphs constructed, batches recorded, cell types annotated or modalities aligned, OmicOS holds this evolving state in shared objects spanning layers, observations, variables, embeddings, graphs, modality views, intermediate artifacts and provenance records (**Fig. 1c**). A workflow planner, state executor, capability grounder and verifier–curator operate over this state, and because each capability declares both its required input state and its produced output state, the runtime checks prerequisites, updates the object, records execution lineage and recovers from failures with structured context (**Fig. 1c**). A complete single-cell trace—raw counts → QC → normalization and HVG selection → scaling → PCA → neighbour graph → Leiden clustering → UMAP → annotation—shows each step’s produced fields (for example, ov.pp.pca writing obsm[’X_pca’]) satisfying the next step’s required fields, with every transition appended to a provenance log (**Supplementary Table 4**). Because state and lineage rather than code are shared, the same capability runs identically from chat, notebooks, command-line interfaces, graphical workspaces or programmatic API calls (**Fig. 1c**).

We next asked how this changes failure. A code-generating agent that emits free-form scripts receives failures mainly as untyped tracebacks, so a missing precomputed graph, a mistyped keyword, a broken native library or a malformed object all arrive in the same form (**Fig. 1d**). OmicOS instead decomposes execution into semantic resolution, argument validation, prerequisite checking, dependency verification, state update, structured-output generation and provenance capture, attributing a failure to one of five classes—missing state, invalid arguments, unavailable dependency, incompatible object structure or failed execution—before or at execution (**Fig. 1d**; **Supplementary Note 2**). Across seven analysis tasks spanning these classes, a baseline code-generating agent averaged 26.3 turns and 4.7 distinct errors per task and failed outright on two—a Dask-orchestration crash in a hand-rolled SCENIC pipeline and a use_rep=’spatial’ mismatch that surfaced in scikit-learn as a “0 feature(s)” error—whereas running the same operations through registered capabilities (for example ov.single.SCENIC and ov.fm.run) typed each crash and passed all seven (**Supplementary Table 5**). The benefit is partly entangled with using a maintained API: in one task both agents hit the same scVelo TypeError and used the same workaround, so the contract supplied typed context rather than an automatic fix. This focused comparison shows contracts converting opaque crashes into typed failure states, not a powered claim about general reliability.

Together, the ecosystem, reconstruction, contract and runtime layers make OmicOS an infrastructure for reproducible multi-omics analysis. The reconstruction layer brings R-centred pipelines into Python as 43 reconstructed workflows assembled from the ported packages, 28 of them wired into the public ov.* API and accelerated over their R references from roughly four-fold to ∼280-fold in the most favourable Rust-backed case (NMF; **Supplementary Table 6**); the contract layer declares what each method requires, produces and modifies; and the runtime preserves state, artifacts, provenance and recovery context across steps. To test whether this substrate translates into agent performance, we next evaluated OmicOS on existing biomedical-agent benchmarks and in controlled with- and without-OmicVerse comparisons using the same minimal agent.

### OmicOS achieves state-of-the-art omics-agent performance and enables local models to approach frontier systems

To benchmark OmicOS against existing systems, we evaluated it on BixBench-Verified-50, an external verified benchmark of heterogeneous biomedical data-analysis tasks. OmicOS reached 90.0% Pass@1 (45/50), the highest of the evaluated systems and ahead of the published biomedical agent Biomni Lab (88.7%), Edison Analysis (78.0%), Claude Code (65.3%) and the OpenAI Agents SDK (61.3%; **Fig. 2a**). To assess how those answers were reached, a fixed LLM judge scored each best-per-question trajectory on five process dimensions, and OmicOS maintained a high mean of 94.8, ranging from 93.0 on transcriptomic tasks (14 tasks, all passed) to 96.7 on epigenomic and functional tasks (**Fig. 2b**; **Extended Data Fig. 1a**; **Supplementary Table 7**). Because this process score is decoupled from the binary outcome—and a trajectory audit showed that answer-fidelity scores can slightly overstate correctness—we treat it as complementary to Pass@1. The 50 questions resolved into 15 analysis nodes spanning four of OmicOS’s eight capability families—general omics analysis (32 tasks), phylogenomics (14), variant analysis (4) and the orchestration layer—so the result reflects broad analytic competence rather than mastery of a single pipeline (**Extended Data Fig. 1b**; **Supplementary Table 8**).

**Figure 2.**
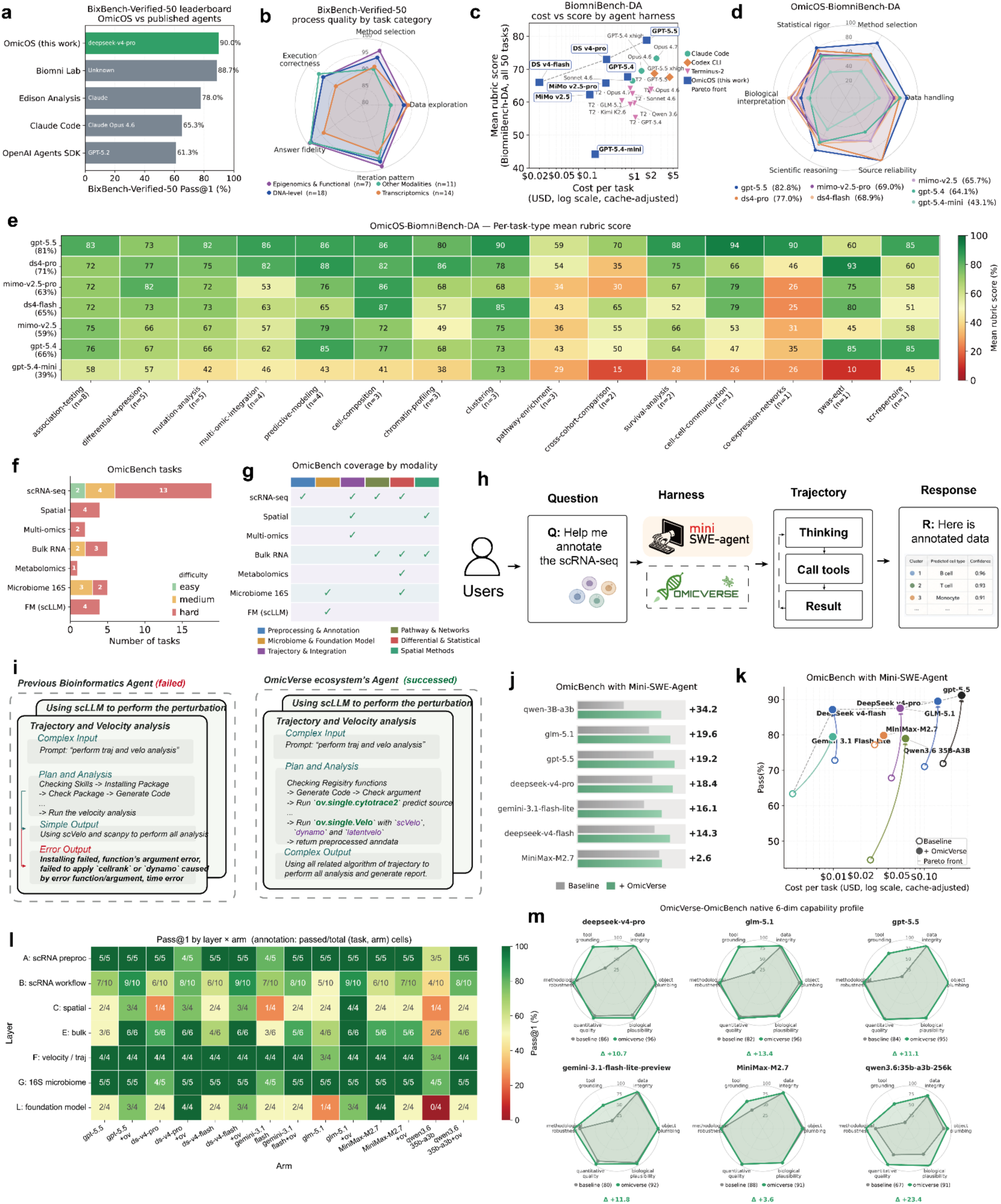
| OmicOS improves accuracy, cost efficiency and robustness of agentic biomedical data analysis. **a,** BixBench-Verified-50 leaderboard comparing OmicOS with published biomedical agents and general-purpose coding-agent systems. Bars show Pass@1 performance, with OmicOS achieving the highest score among the evaluated systems. **b,** Process-quality analysis of BixBench-Verified-50 tasks stratified by task category. Radar plots summarize rubric scores for model selection, data exploration, iteration pattern, answer fidelity and execution correctness across transcriptomic, DNA-level, epigenomic and other biomedical data-analysis tasks. **c,** Cost–performance analysis on BiomniBench-DA. Each point represents an agent harness–model combination, with mean rubric score plotted against cache-adjusted cost per task on a logarithmic scale. OmicOS-enabled agents shift the Pareto frontier towards higher analytical quality at lower or comparable cost. **d,** Capability profile of OmicOS-BiomniBench-DA across six evaluation dimensions: statistical rigor, method selection, data handling, source reliability, scientific reasoning and biological interpretation. Radar plots show performance differences across model backbones when equipped with the OmicOS execution harness. **e,** Per-task-type heat map of OmicOS-BiomniBench-DA performance. Rows indicate model backbones and columns indicate biomedical task types, including association testing, differential expression, mutation analysis, multi-omics integration, predictive modelling, cell composition, chromatin profiling, clustering, pathway enrichment, cross-cohort comparison, survival analysis, cell–cell communication, co-expression networks, genome-wide association studies and T-cell receptor repertoire analysis. Color denotes the mean rubric score. **f,** Composition of the OmicBench task suite by modality and difficulty. Tasks span scRNA-seq, spatial transcriptomics, multi-omics, bulk RNA-seq, metabolomics, 16S microbiome analysis and single-cell foundation-model workflows, with difficulty labels assigned to each task. **g,** Coverage matrix of OmicBench across analytical capability classes. Check marks indicate whether each modality includes tasks involving preprocessing and annotation, microbiome or foundation-model analysis, trajectory and integration, pathway and network analysis, differential and statistical analysis, or spatial methods. **h,** Minimal-agent evaluation design using Mini-SWE-Agent. Mini-SWE-Agent provides a deliberately simple agent loop consisting of task reasoning, tool calling and result generation. This lightweight harness is used as a controlled testbed to determine whether adding OmicVerse capabilities improves an agent’s ability to answer omics-analysis questions, rather than attributing gains to a more complex agent architecture. **i,** Representative failure and recovery example for trajectory and velocity analysis. A baseline bioinformatics agent fails when directly generating package-specific code with invalid assumptions or arguments, whereas the OmicVerse-enabled agent succeeds by grounding the request in registered functions, checking arguments, selecting appropriate trajectory and velocity tools, and returning processed AnnData objects. **j,** Model-level improvement on OmicBench after adding OmicVerse to the same Mini-SWE-Agent framework. Bars compare baseline Mini-SWE-Agent performance with OmicVerse-augmented performance across multiple model backbones, with numerical labels indicating the absolute gain. **k,** Cost–accuracy trade-off for OmicBench under the Mini-SWE-Agent harness. Points connect baseline and OmicVerse-augmented runs for the same model, showing that OmicVerse improves Pass@1 across a broad cost range without requiring a more complex agent design. **l,** Layer-by-arm Pass@1 heat map for OmicBench. Rows correspond to benchmark layers, including scRNA-seq preprocessing, scRNA-seq workflows, spatial analysis, bulk RNA-seq, trajectory and velocity analysis, 16S microbiome analysis and foundation-model tasks. Columns compare model arms with or without OmicVerse under the same minimal-agent setting. Each cell reports the number of passed tasks over the total number of evaluated tasks. **m,** Native six-dimensional capability profiles for OmicVerse-OmicBench. Radar plots compare baseline and OmicVerse-augmented agents across tool grounding, data integrity, object plumbing, biological plausibility, quantitative quality and methodological robustness. Green traces indicate OmicVerse-enabled runs, showing consistent improvements in agentic omics-analysis capability across diverse model families.

**Extended Data Fig. 1.**
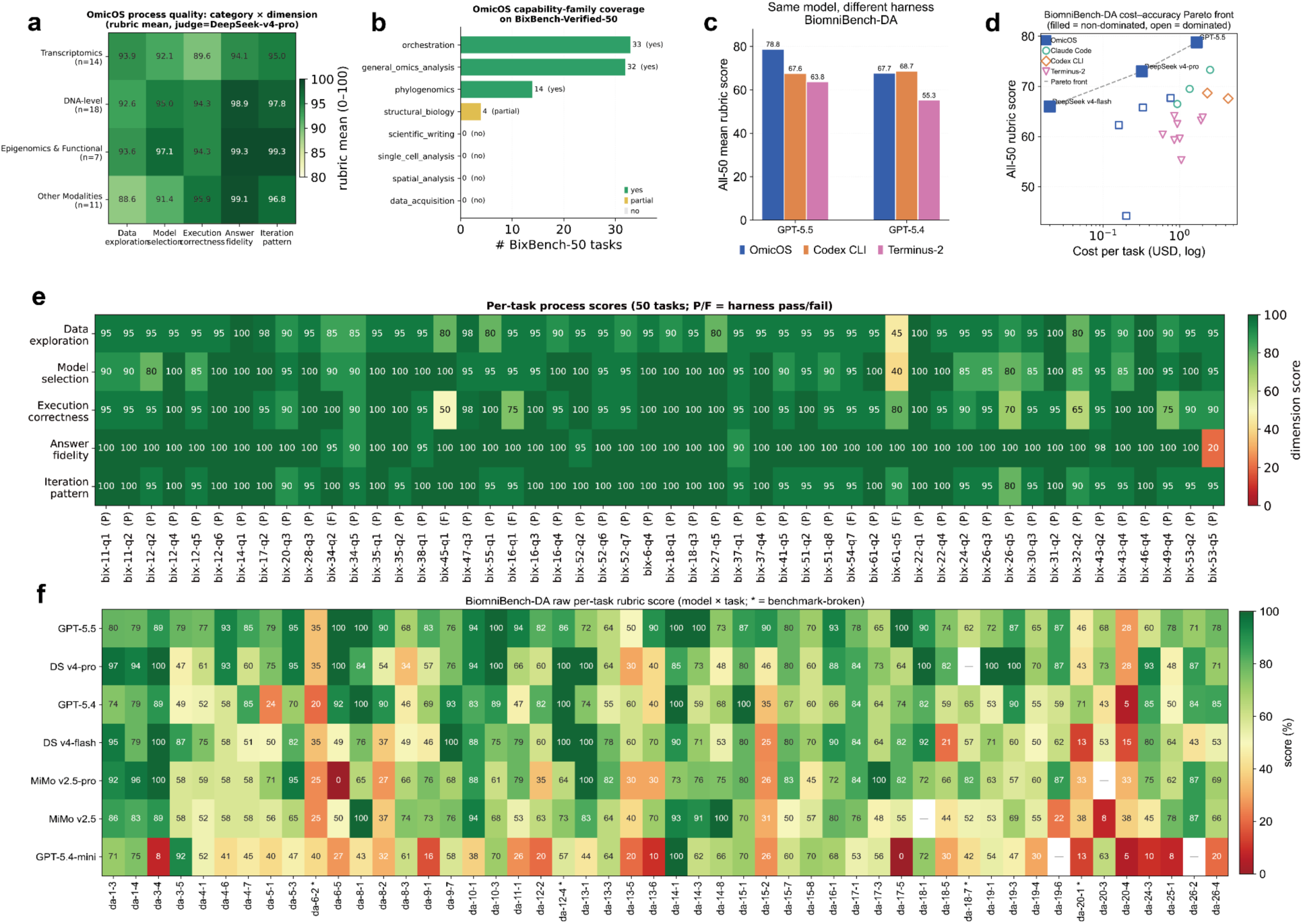
| Extended process-quality and raw benchmark analyses for OmicOS on BixBench-Verified-50 and BiomniBench-DA. **a,** Process-quality scores of OmicOS on BixBench-Verified-50 stratified by task category and evaluation dimension. Rows indicate transcriptomics, DNA-level analysis, epigenomics and functional analysis, and other modalities; columns indicate data exploration, model selection, execution correctness, answer fidelity and iteration pattern. Values denote mean rubric scores on a 0–100 scale. **b,** Capability-family coverage of OmicOS across BixBench-Verified-50 tasks. Bars show the number of benchmark tasks covered by each capability family, with coverage status indicated as complete, partial or absent. OmicOS covered major orchestration, general omics analysis and phylogenomics tasks, with partial coverage for structural biology and no coverage for several categories not represented in the current execution ecosystem. **c,** Same-model cross-harness comparison on BiomniBench-DA. Mean all-50-task rubric scores are shown for GPT-5.5 and GPT-5.4 when executed through OmicOS, Codex CLI or Terminus-2. The comparison illustrates that realized biomedical-analysis performance depends on the agent harness as well as the underlying language model. **d,** Cost–accuracy Pareto analysis on BiomniBench-DA. Each point represents an agent-harness and model combination, with mean all-50-task rubric score plotted against cache-adjusted cost per task on a logarithmic scale. Filled points indicate non-dominated Pareto-front configurations, and open points indicate dominated configurations. **e,** Per-task process-quality heat map for BixBench-Verified-50. Rows correspond to the five process-quality dimensions, and columns correspond to individual benchmark tasks. Column labels indicate whether the OmicOS harness passed or failed each task. Color denotes the dimension-level rubric score. **f,** Raw per-task BiomniBench-DA rubric scores across OmicOS model backbones. Rows indicate model backbones and columns indicate individual BiomniBench-DA tasks. Color denotes the raw task-level rubric score. Asterisks mark benchmark-broken tasks that were excluded from corrected capability-profile analyses in the main figure.

**Extended Data Figure 2.**
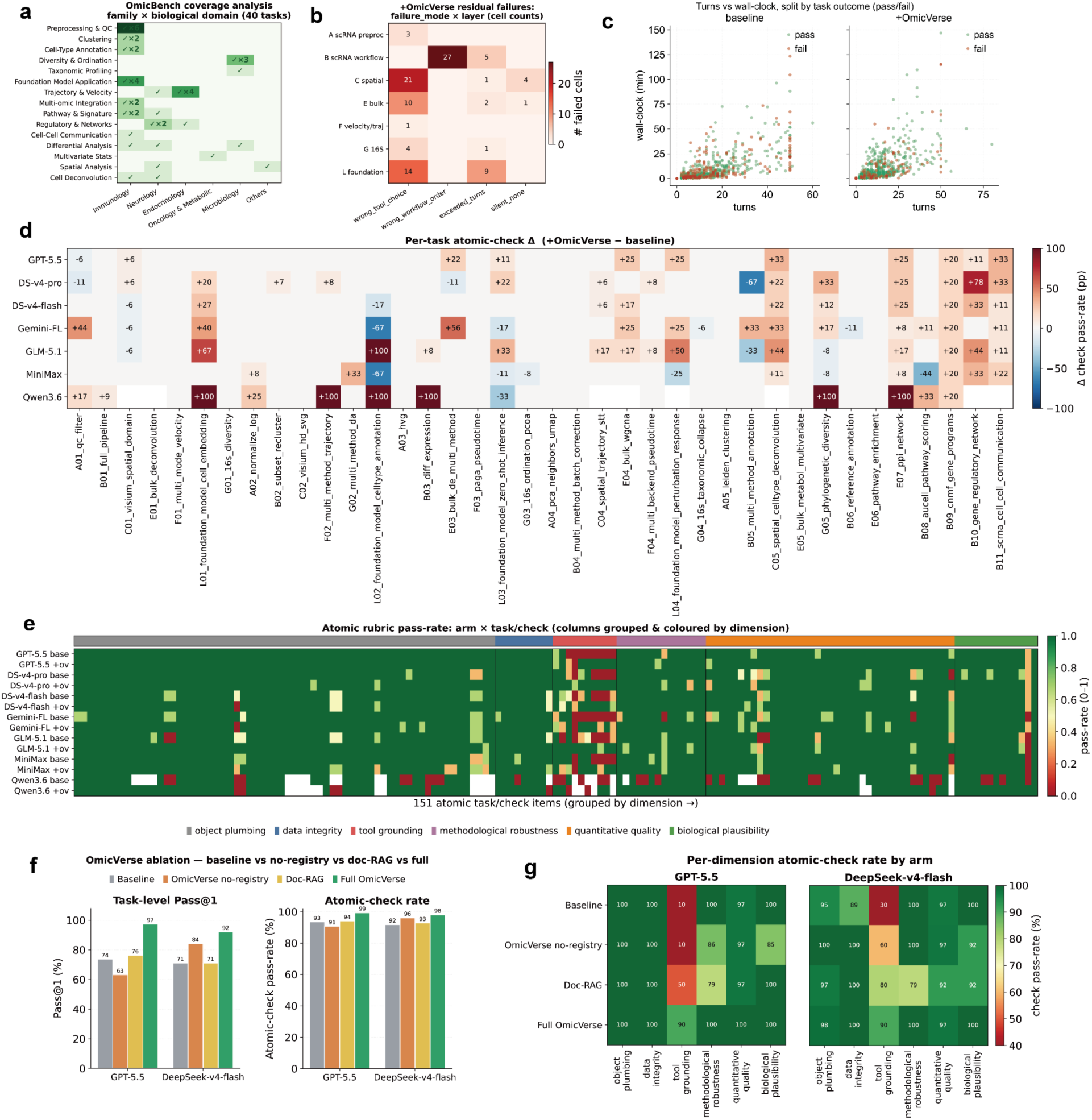
| OmicBench benchmark structure and the mechanism of the OmicVerse gain under a minimal agent. (All panels use the deliberately minimal Mini-SWE-Agent harness; “+OmicVerse” adds the registry-grounded OmicVerse execution layer while holding the agent loop fixed. An arm denotes a model evaluated under a given condition. Each task is graded by a set of atomic rubric checks, which are grouped into six native capability dimensions: object plumbing, data integrity, tool grounding, methodological robustness, quantitative quality and biological plausibility. Pass@1 counts a task as passed only when all of its checks pass; the atomic-check pass-rate is the fraction of individual rubric items passed, and is therefore always higher than Pass@1 because a failing task still passes most of its checks.) **(a)** OmicBench coverage. Each of the 40 tasks is mapped to one fine-grained analysis family (rows; 15 families) and one biological/disease domain (columns); cells give the number of tasks (✓, or ✓×n). The benchmark spans many analysis types but is concentrated in immunology-flavoured single-cell tasks. **(b)** Residual failures under +OmicVerse. For tasks that still failed after augmentation (majority of seeds), the number of failed cells by failure mode (columns) × OmicBench layer (rows). Residual failures concentrate in the spatial (C) and foundation-model (L) layers and in wrong-tool / wrong-workflow-order modes rather than runtime crashes. **(c)** Agent efficiency. Per-(task, seed) number of turns (LLM calls) versus wall-clock time, split by task outcome (green = pass, red = fail), for the baseline (left) and +OmicVerse (right) arms pooled across the seven model families. Adding OmicVerse does not inflate the turn or time budget. **(d)** Per-task, per-model change in atomic-check pass-rate from baseline to +OmicVerse (Δ, percentage points); rows = seven model families, columns = the 38 evaluated tasks. Warm/red = +OmicVerse raises the check rate, cool/blue = lowers it. Gains are broadly distributed across tasks and models rather than confined to a few easy items. **(e)** Full atomic rubric matrix. Pass-rate of every one of the 151 atomic task/check items (columns, grouped and colour-coded by capability dimension; see top strip and key) across all 14 arms (rows; each model as baseline and +OmicVerse). The +OmicVerse rows are greener overall, with the largest shift in the tool-grounding block, showing that a single task-level pass is composed of different combinations of dimension-specific checks and that OmicVerse most improves whether the correct domain tool is actually invoked. **(f, g)** OmicVerse ablation isolating three mechanisms — documentation access (Doc-RAG), named-tool access without the structured registry (OmicVerse no-registry) and full registry-grounded execution (Full OmicVerse) — versus baseline, for GPT-5.5 and DeepSeek-v4-flash on a common seed over 38 tasks. (f) Task-level Pass@1 (left) and atomic-check pass-rate (right) by arm: Full OmicVerse is best on both metrics for both models (Pass@1 73.7→97.4% for GPT-5.5 and 71.1→92.1% for DeepSeek-v4-flash), beating Doc-RAG (76.3% and 71.1%) and no-registry (63.2% and 84.2%); for GPT-5.5 the no-registry arm even falls below baseline, indicating that exposing tool names without the registry/contract layer can mislead the model. (g) Per-dimension atomic-check rate by arm for the two models; Full OmicVerse’s advantage is concentrated in tool grounding and the methodological and quantitative dimensions. Together, panels (f) and (g) attribute the OmicBench gain to the contract/registry execution layer rather than to additional documentation text or tool names alone.

To separate the contribution of the harness from that of the model, we placed OmicOS in the BiomniBench-DA cost–performance space alongside the general-purpose agents Claude Code, Codex CLI and Terminus-2. OmicOS with GPT-5.5 attained the highest all-50-task mean rubric score, 78.8 at a cache-adjusted $1.67 per task, above Claude Code with Opus 4.7 (73.3 at $2.50), Codex CLI with GPT-5.5 (67.6 at $4.30) and Terminus-2 with GPT-5.5 (63.8 at $2.00; **Fig. 2c**; **Supplementary Table 9**). The same model behaved very differently across harnesses—GPT-5.5 scored 78.8 under OmicOS but 67.6 under Codex CLI and 63.8 under Terminus-2, an 11–15-point swing at equal or lower cost (**Extended Data Fig. 1c**)—and OmicOS also reached competitive low-cost points, with DeepSeek-v4-pro at 73.0 ($0.32 per task) and DeepSeek-v4-flash at 66.0 (∼$0.02 per task; **Fig. 2c**; **Extended Data Fig. 1d**). Scores are computed on the same 50 tasks for every system, but because the external rows were transcribed from a published chart rather than re-measured under one tool stack and cache convention, we read the cost axis as indicative. The consistent same-model swing nonetheless indicates that realized biomedical-analysis capability depends on the agent harness, not the model alone.

To profile model capability within the OmicOS harness, we excluded four documented benchmark-broken tasks (da-12-4, da-18-7, da-20-1, da-6-2) identically from every model after inspecting each task’s instruction, rubric and grade record (**Fig. 2d**; **Supplementary Note 3**); this raised every model’s score by only 0.2–1.5 points and left the ranking unchanged (Supplementary Table 10). On the corrected 46-task set, GPT-5.5 scored highest at 82.8%, followed by DeepSeek-v4-pro (77.0%), mimo-v2.5-pro (69.0%), DeepSeek-v4-flash (68.9%), mimo-v2.5 (65.7%), GPT-5.4 (64.1%) and GPT-5.4-mini (43.1%; Fig. 2d; Supplementary Table 10). GPT-5.5 led on source reliability (100.0), method selection (87.6), data handling (84.4), scientific reasoning (83.2) and statistical rigor (79.9) but fell to 61.3 on biological interpretation—a panel-wide weak axis on which DeepSeek-v4-flash (78.0) and mimo-v2.5-pro (75.7) scored higher, suggesting that domain interpretation is not closed by scaling the strongest general model (**Fig. 2d**; **Supplementary Table 11**). A per-task-type heat map showed GPT-5.5 strong across association testing (83), mutation analysis (82), multi-omics integration (86), predictive modelling (86), cell composition (86), clustering (90), survival analysis (88), cell–cell communication (94), co-expression networks (90) and TCR repertoire (85) but weaker on pathway enrichment (59) and GWAS (60), while other models showed distinct profiles—DeepSeek-v4-pro strong on GWAS (93) yet weak on cross-cohort comparison (35) and co-expression networks (46), and GPT-5.4-mini low on cross-cohort comparison (15), GWAS (10), cell–cell communication (26) and co-expression networks (26; **Fig. 2e**). Several task types comprise only one or two tasks and should be read as single runs (**Extended Data Fig. 1f**; **Supplementary Table 12**).

The cross-harness comparison is indicative rather than controlled, so to isolate the effect of the execution layer we constructed OmicBench and evaluated it with Mini-SWE-Agent, a deliberately minimal harness in which only the availability of OmicVerse is varied (**Fig. 2h**). OmicBench contains 40 tasks across seven modalities—scRNA-seq (19), spatial transcriptomics (4), multi-omics (2), bulk RNA-seq (5), metabolomics (1), 16S microbiome (5) and single-cell foundation-model workflows (4)—and is deliberately hard, with 29 hard, 9 medium and 2 easy tasks (Fig. 2f; Supplementary Table 13). Each prompt states only the scientific goal and never names an ov.* API, and grading is programmatic and package-agnostic: checks assert biological and numerical criteria (clustering ARI against known labels, marker overlap against an oracle, multi-method consensus) and accept equivalent outputs from any library, with tool-grounding checks crediting third-party packages such as cellphonedb, liana, scGPT and Geneformer, so a baseline agent can pass with Scanpy, Bioconductor or hand-written code (**Supplementary Table 13**). The tasks span six capability classes—preprocessing and annotation (10), microbiome and foundation-model analysis (8), trajectory and integration (7), pathway and network analysis (7), spatial methods (4) and differential and statistical analysis (4)—and 15 fine-grained families (**Fig. 2g; Extended Data Fig. 2a**; **Supplementary Table 14**). In a representative trajectory-and-velocity task, the baseline agent failed after generating package-specific code with invalid assumptions, whereas the OmicVerse-enabled agent grounded the request in registered functions, checked arguments and returned a processed AnnData object (**Fig. 2i**).

Adding OmicVerse to this fixed harness raised Pass@1 for every model family, and most for the weaker ones. Across the 38-task evaluation subset (the two layer-D multi-omics tasks were not run), Qwen3.6-35B-A3B improved most, from 44.7% to 78.9% (+34.2 percentage points), followed by GLM-5.1 (67.9%→87.5%, +19.6), GPT-5.5 (71.9%→91.2%, +19.2), DeepSeek-v4-pro (71.1%→89.5%, +18.4), Gemini-3.1-flash-lite (63.4%→79.5%, +16.1), DeepSeek-v4-flash (72.8%→87.1%, +14.3) and MiniMax-M2.7 (77.2%→79.8%, +2.6; **Fig. 2j**; **Supplementary Table 15**); six models ran at three seeds and the locally served Qwen arm at one. The gains were distributed across layers rather than one task family—scRNA-seq preprocessing and workflows, bulk RNA-seq, trajectory and velocity, 16S microbiome and foundation-model tasks—with GLM-5.1 rising from 25/38 to 36/38 task passes and Qwen3.6-35B-A3B from 17/38 to 30/38 by majority-seed vote, although the near-saturated MiniMax-M2.7 dropped slightly (31→30; **Fig. 2l**; **Supplementary Table 16**). Of the 35 residual failures after augmentation, spatial and foundation-model tasks remained hardest and the dominant cause was model reasoning (15/35)—the agent failing to invoke or orchestrate the specialised tool rather than crashing—so OmicVerse improves execution without removing every model- or modality-specific limit (**Fig. 2l**; **Supplementary Table 17**).

Plotted against cost, adding OmicVerse moved nearly all arms upward at similar per-task cost and pushed several open-weights configurations toward the Pareto frontier (**Fig. 2k**; **Supplementary Table 18**). After augmentation the open-weights arms DeepSeek-v4-pro (89.5% Pass@1), GLM-5.1 (87.5%) and DeepSeek-v4-flash (87.1%) approached the strongest closed-source arm, GPT-5.5 + OmicVerse (91.2%), and the locally served Qwen3.6-35B-A3B reached 78.9% (**Fig. 2j,k**); here closed-source denotes API-only models (GPT-5.5, Gemini), open-weights models with downloadable weights (DeepSeek, GLM, MiniMax, Qwen), and only Qwen was served locally. Decomposing the same grading signal into six native capability dimensions—used as a diagnostic of where the gain falls, not as an independent benchmark—showed the same pattern: OmicVerse lifted DeepSeek-v4-pro from 85.7 to 96.4 (+10.7), GLM-5.1 from 82.3 to 95.7 (+13.4), GPT-5.5 from 84.0 to 95.2 (+11.1), Gemini-3.1-flash-lite from 79.9 to 91.7 (+11.8), MiniMax-M2.7 from 87.8 to 91.4 (+3.6) and Qwen3.6-35B-A3B from 67.2 to 90.6 (+23.4; **Fig. 2m**; **Supplementary Table 19**). The uplift was largest for the weakest baseline (Qwen3.6-35B-A3B) and smallest for the strongest (MiniMax-M2.7), concentrated in tool grounding and methodological robustness, and did not inflate the per-task turn or wall-clock budget (**Extended Data Fig. 2c,e**).

To show that this gain comes from the execution layer rather than from extra documentation or tool names, we ran a four-arm ablation on the same harness—baseline, full OmicVerse, OmicVerse without the registry (tools named but without the structured registry and contract layer), and a documentation-RAG variant—for GPT-5.5 and DeepSeek-v4-flash on a common seed over the 38 tasks (Extended Data Fig. 2f,g; Supplementary Table 20). Full OmicVerse was best on both Pass@1 and atomic-check pass-rate for both models: GPT-5.5 rose from 73.7% (baseline) to 76.3% (doc-RAG) to 97.4% (full), while the no-registry arm fell to 63.2%, below baseline; DeepSeek-v4-flash rose from 71.1% (baseline and doc-RAG) and 84.2% (no-registry) to 92.1% (full), with the same ordering at the atomic-check level (GPT-5.5 99.3% versus ≤94.0% for the partial arms; DeepSeek-v4-flash 98.0% versus ≤96.0%). Because neither documentation text nor un-grounded tool names recovered more than a fraction of the gain—and naming tools without grounding even pushed GPT-5.5 below baseline—this single-seed ablation attributes the gain to the contract and registry execution layer.

Across BixBench-Verified-50, BiomniBench-DA and OmicBench, OmicOS improved omics-agent performance through ecosystem-grounded execution. It ranked first on both external benchmarks—90.0% Pass@1 on BixBench-Verified-50 and the top all-50-task rubric score (78.8) on BiomniBench-DA, where the same model scored very differently across harnesses—and in the controlled Mini-SWE-Agent setting, adding OmicVerse raised Pass@1 by up to 34.2 percentage points and broadened native capability profiles across model families, with the ablation tracing the gain to the registry-grounded execution layer rather than to documentation or tool names. Together these results indicate that OmicOS turns the OmicVerse ecosystem into transferable capability for agentic biomedical data analysis, rather than acting as a stronger prompt or a larger model.

### Reconstructing complete omics-analysis pipelines establishes the software infrastructure required for agentic biological discovery

Many canonical omics pipelines—differential analysis, trajectory inference, repertoire and network analysis, eQTL and TWAS analysis, microbiome analysis and statistical modelling—were written in R or Bioconductor, while modern analysis runs on Python objects such as AnnData and MuData. To close this gap we built OmicVerse-RebuildR, which ports complete R-centred workflows into Python while preserving reference behaviour, and paired it with an upper design layer that unifies the ported algorithms behind a small set of shared, optimized interfaces (**Fig. 3a–d**). The reconstruction catalogue now holds 62 packages (59 pure-Python and 3 Rust), 56 released on PyPI and 45 wired into a public ov.* entry point. Reconstructed-R ports and native-Python tools share one dispatch pattern over a single namespace, agree on a fixed object-state contract, and read and write one AnnData/MuData object model (the IBE substrate), so that across nine dispatchers a ported and a native tool are called indistinguishably—py-Slingshot and Palantir, for example, both run through ov.single.TrajInfer (**Supplementary Table 22**). Each port follows a fixed protocol in which the R source is the executable specification, parity is class-aware, and reconstruction is held to identity rather than metric optimization, run through six stages from duplicate-checking discovery to release (**Supplementary Note 4**).

**Figure 3.**
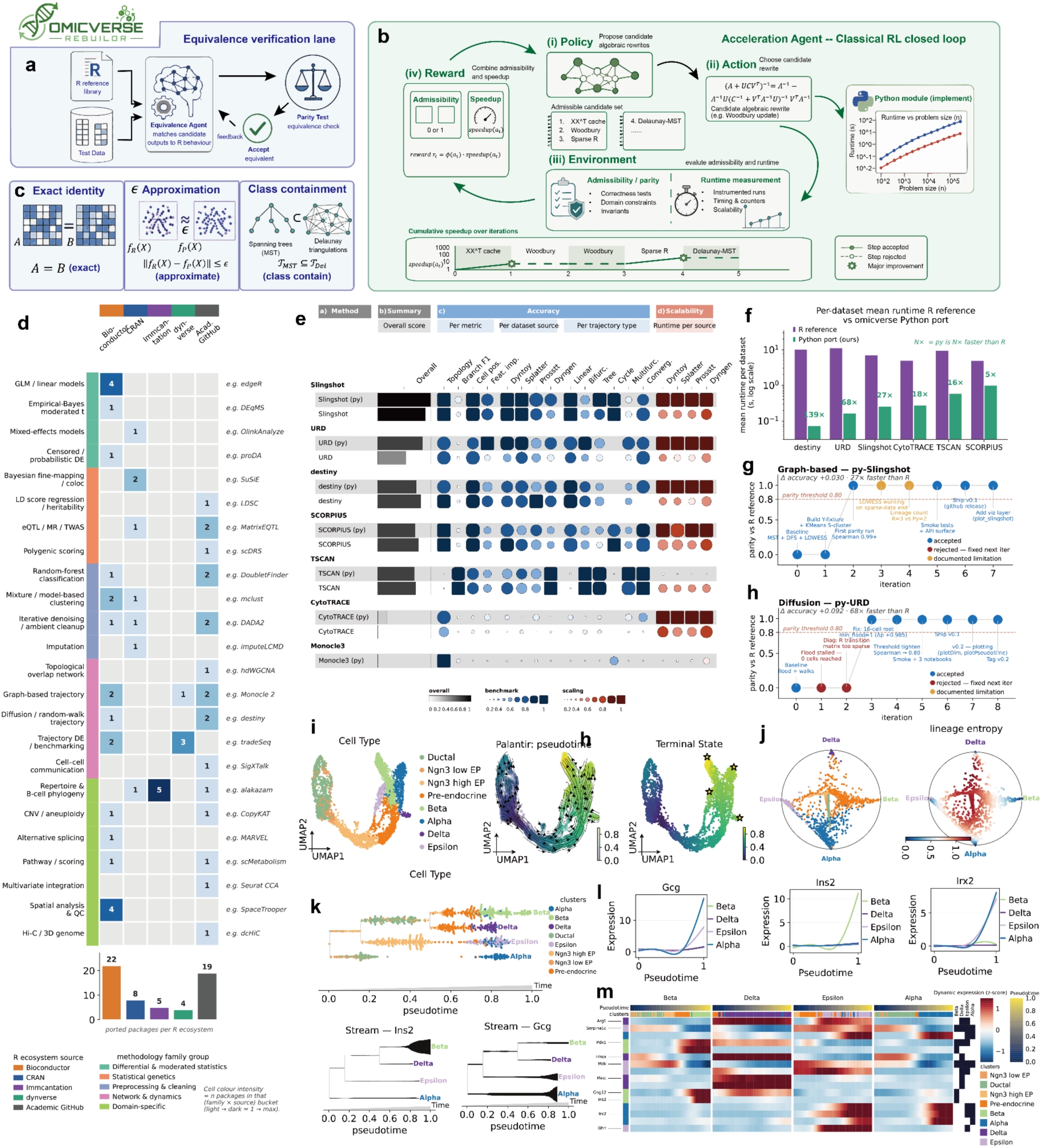
| Self-evolving reconstruction of the R omics ecosystem enables a unified pseudotime-fate analysis framework. **a,** Equivalence-verification lane of OmicVerse-RebuildR. For each target R package, the R reference and shared fixtures serve as the executable specification; an equivalence agent iteratively reconstructs a candidate Python implementation and compares it against R under pre-registered, class-aware parity tests, accepting a port only when the equivalence criteria are met. **b,** Self-evolving acceleration loop. After a parity-cleared baseline port, an acceleration agent runs a closed loop of policy (propose algebraic/algorithmic rewrites), action (apply a rewrite), environment (evaluate admissibility and runtime) and reward (admissibility + measured speedup), preserving equivalence to R throughout. **c,** Admissibility criteria: rewrites are accepted only under exact identity, bounded ε-approximation or class-containment guarantees. **d,** Reconstruction landscape by methodology family (rows) and source ecosystem (columns: Bioconductor, CRAN, immunogenomics, dynverse, academic GitHub); numbers give reconstructed targets per category, representative methods at right, and the bar plot the packages per ecosystem. **e,** Benchmark overview of reconstructed trajectory/pseudotime methods versus their R references across topology, branch assignment, pseudotime consistency and runtime, over multiple datasets and trajectory types. **f,** Per-dataset runtime (mean per dataset, log scale) for R references versus OmicVerse Python ports of representative trajectory methods. **g,** Iterative evolution of py-Slingshot: parity to R and cumulative acceleration across accepted and rejected rewrites, with annotated implementation updates. **h,** Iterative evolution of py-URD: parity and speed across the self-evolving optimization. **i,** Unified pseudotime-fate analysis with PseudotimeFate, the upper-level design of omicverse.single.Traj. Using a sparse PCCA+ strategy in place of the heavier CellRank/pyGPCCA stack, it couples pseudotime ordering, terminal-state inference and fate-probability estimation in one framework. The pancreatic endocrine example shows cell-type annotation, pseudotime and terminal states in a shared embedding. **j,** Circular fate embeddings summarizing branch bias toward terminal endocrine states and lineage entropy. **k,** Lineage-resolved pseudotime and gene dynamics; representative markers (Gcg, Ins2, Irx2) show fate-specific expression along pseudotime. **m,** Stream plots of branch-specific expression trends and a pseudotime-ordered dynamic heat map of transcriptional programs across terminal endocrine lineages.

**Extended Data Fig. 3.**
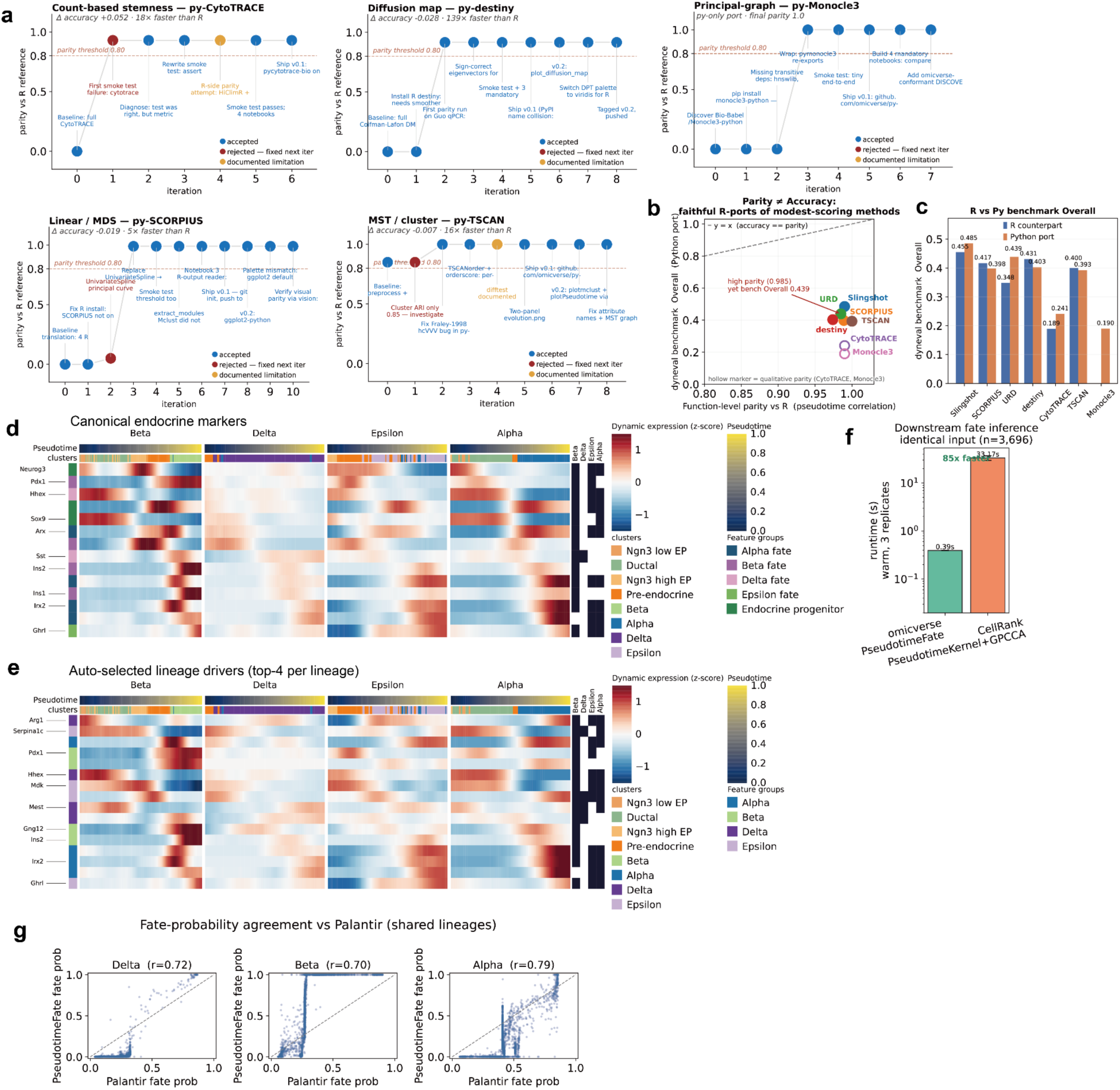
| Auditable reconstruction of trajectory ports and lineage-resolved downstream analysis with PseudotimeFate. **a,** Per-port reconstruction-evolution traces for representative trajectory-inference ports not shown in the main figure. Each panel plots parity with the R reference against reconstruction iteration for one Python port: count-based stemness (**py-CytoTRACE**), diffusion map (**py-destiny**), principal-graph inference (**py-Monocle3**), linear/MDS trajectory inference (**py-SCORPIUS**) and minimum-spanning-tree/cluster-based trajectory inference (**py-TSCAN**). Blue markers indicate accepted iterations, red markers indicate rejected iterations that were fixed in the next step, and orange markers indicate documented limitations. Dashed horizontal lines denote the pre-registered parity gate of 0.80. Text annotations summarize the major implementation changes, debugging steps or limitations recorded during reconstruction. Italic subtitles report the benchmark accuracy change relative to the R reference and the runtime speed-up. These traces show that reconstruction failures, such as spline degeneracy in **py-SCORPIUS** or rejected parity steps in **py-CytoTRACE** and **py-TSCAN**, were explicitly documented and resolved while maintaining parity above the predefined threshold. **b,** Relationship between function-level parity to the R reference and trajectory-benchmark accuracy for reconstructed Python trajectory ports. The x axis shows parity with the R implementation, and the y axis shows dynbenchmark-style overall accuracy. The dashed line indicates equality between the two quantities. High parity and high benchmark accuracy represent distinct properties: parity measures fidelity of the Python port to the R reference, whereas benchmark accuracy measures the trajectory-recovery performance of the underlying method. For example, **py-URD** achieved high R-reference parity but modest benchmark accuracy, indicating faithful reconstruction of a method whose benchmark behaviour is itself limited. Hollow markers denote ports with qualitative parity only. **c,** Paired R-versus-Python benchmark accuracy for representative trajectory methods. Bars show dynbenchmark-style overall scores for R references and reconstructed Python ports. Python ports reproduced the benchmark behaviour of their R counterparts and, in several cases, matched or exceeded the R reference, including **py-Slingshot** (0.485 versus 0.455), **py-URD** (0.439 versus 0.348) and **py-CytoTRACE** (0.241 versus 0.189). Other ports showed comparable scores, including **py-TSCAN** (0.393 versus 0.400), **py-SCORPIUS** (0.398 versus 0.417) and **py-destiny** (0.403 versus 0.431). **d,** Dynamic expression heat map of canonical endocrine markers along pseudotime, split by the four mature lineages recovered by PseudotimeFate: Beta, Delta, Epsilon and Alpha. The analysis was performed on a pancreatic endocrine dataset containing 3,696 cells and 3,000 highly variable genes. Rows show fitted, z-scored expression trends, and columns represent pseudotime-ordered cells or bins within each fate branch. Top annotations indicate pseudotime, and side annotations indicate endocrine feature groups and cell-state clusters. Canonical markers are organized by expected fate programs, including Alpha markers (**Gcg**, **Arx**, **Irx1**, **Irx2**), Beta markers (**Ins1**, **Ins2**, **Pdx1**, **Pax4**), Delta markers (**Sst**, **Hhex**), the Epsilon marker **Ghrl**, and endocrine progenitor markers (**Sox9**, **Neurog3**, **Fev**). **e,** Dynamic expression heat map of automatically selected lineage drivers. The top four genes selected for each lineage by PseudotimeFate are shown across the same pseudotime-ordered fate branches as in **d**. The unbiased driver set recovers branch-specific expression programs that are consistent with the curated endocrine marker patterns. **f,** Runtime comparison of downstream fate inference on identical input data. OmicVerse **PseudotimeFate** and a comparable CellRank-style **PseudotimeKernel + GPCCA** workflow were run on the same AnnData object, Palantir pseudotime and k-nearest-neighbour graph with 3,696 cells. Bars show mean runtime over three warm replicates on a log scale. PseudotimeFate completed in 0.39 ± 0.00 s, whereas the CellRank-style workflow required 33.17 ± 3.66 s, corresponding to an approximately 85-fold speed-up. **g,** Agreement between PseudotimeFate and Palantir fate probabilities for shared terminal lineages. Each point represents one cell, with Palantir fate probability on the x axis and PseudotimeFate fate probability on the y axis. Pearson correlations are shown for Delta (r = 0.72), Beta (r = 0.70) and Alpha (r = 0.79). The concordance indicates that PseudotimeFate recovers fate-probability landscapes consistent with an established pseudotime-fate framework despite using a distinct macrostate inference procedure.

RebuildR accepts a port only after class-aware equivalence verification, treating the R implementation as the executable specification. For each package, the R reference and shared fixtures define the expected behaviour, and an equivalence agent translates that behaviour into a candidate Python package whose outputs are compared against R under parity tests registered before any code is written (**Fig. 3a**). We matched each test to the class of output it checks (**Supplementary Table 23**): deterministic matrices to bit-equivalence (max|Δ| < 1×10⁻¹⁰; e.g. py-matrixeqtl β max|diff| < 2×10⁻¹⁵, py-coloc posteriors < 3×10⁻¹⁶), test statistics to rank and top-hit overlap, embeddings to rotation-invariant subspace agreement (Procrustes ≥ 0.80; py-destiny 0.916), clusterings to label-permutation-invariant agreement (ARI ≥ 0.90), pseudotime to rank correlation (Spearman ≥ 0.80, usually ≥ 0.99; py-Slingshot ≥ 0.99, py-URD 0.985), and tree topologies, discrete sets and stochastic outputs to structure-, set-or distribution-level criteria (**Fig. 3a,c**). This separation lets a failure be read correctly: a failed deterministic-kernel test is a bug, whereas a failed invariance-class test is a genuine scientific disagreement rather than a relabeling artifact, although the guarantee is fixture-level rather than a proof over all inputs.

Once a port is equivalent, RebuildR optimizes it through a self-evolving acceleration loop that cannot break equivalence. The acceleration agent runs a closed loop over policy, action, environment and reward: it proposes a rewrite, applies it, measures runtime and parity, and accepts it only if it is faster and still clears the equivalence gate (**Fig. 3b**). Every accepted rewrite carries an admissibility proof of one of three kinds—exact mathematical identity, bounded ε-approximation or class-containment guarantee (**Fig. 3c**)—which keeps optimization from drifting into a different algorithm. Across the seven packages with full telemetry, 34 iterations gave 27 accepted and 4 rejected rewrites (2 for dropping parity, 2 for no wall-clock gain), with exact-identity rewrites in all seven and bounded-ε rewrites in four; one rewrite that wrongly assumed k-nearest-neighbour contiguity dropped parity from 0.984 to 0.753 and was rejected at the gate (**Supplementary Table 24**). Logged speedups reach 102-fold here and extend to roughly 280-fold (rust-NMF) and 192-fold (py-inferCNV) across the broader catalogue, though structured policy–action–reward telemetry is so far published for 7 of the ∼62 packages.

RebuildR has grown beyond any single method family. Its landscape spans five source ecosystems—Bioconductor, CRAN, immunogenomics resources, dynverse and academic GitHub—and comprises 58 reconstructed or targeted components (22 Bioconductor, 8 CRAN, 5 immunogenomics, 4 dynverse, 19 academic GitHub) across 24 methodology families, from generalized linear and empirical-Bayes models through Bayesian fine-mapping, eQTL, TWAS and polygenic scoring, model-based clustering, ambient-RNA correction and imputation, co-expression and regulatory networks, graph- and diffusion-based trajectory inference, B-cell-receptor analysis, copy-number inference, alternative splicing, pathway scoring and multivariate integration (**Fig. 3d**; **Supplementary Table 25**). Because breadth is not completeness, we report five non-nested maturity statuses for these components (**Supplementary Table 26**): all are ported, 46 carry a quantified parity metric (12 only a stated one), 19 have a benchmarked acceleration, 43 are integrated into a public ov.* entry point, and 40 are agent-executable out of the box through the 788-function OmicVerse registry that ov.Agent and the MCP server call.

We used trajectory inference as a stress test, because it combines diverse algorithmic structures with non-trivial output invariances. We benchmarked 13 methods (six R references and seven Python ports) on 14 synthetic datasets from four simulators (Dyntoy, Splatter, PROSSTT, dyngen) over six topology classes and 300–600 cells, scoring 182 method-by-dataset cells with dynbenchmark-style metrics through the Python dyneval port (**Fig. 3e**; **Supplementary Tables 27, 28**; **Supplementary Note 5**). The Python ports matched or exceeded their R counterparts in three of six paired comparisons—py-Slingshot 0.485 versus 0.455, py-URD 0.439 versus 0.348, py-CytoTRACE 0.241 versus 0.189—while R stayed ahead for destiny (0.431 versus 0.403), SCORPIUS (0.417 versus 0.398) and TSCAN (0.400 versus 0.393), with Monocle3 available only as a Python port (0.190; **Fig. 3e**; **Supplementary Table 28**). These dyneval scores are low in absolute terms on deliberately hard synthetic topologies and measure benchmark-level trajectory agreement, a different axis from the function-level parity the ports are actually gated on (pseudotime Spearman ≥ 0.99, embedding Procrustes ≥ 0.80); on that benchmark axis the ports were comparable to their R references.

The ports were also markedly faster. Every Python trajectory port beat its R reference, with a median speedup of 22.6-fold and method-specific speedups of 139-fold for destiny, 68-fold for URD, 27-fold for Slingshot, 18-fold for CytoTRACE, 16-fold for TSCAN and 5-fold for SCORPIUS, at modest peak memory (**Fig. 3f**; **Extended Data Fig. 3a**; **Supplementary Table 29**). The gains came from different sources: destiny’s 139-fold speedup is algorithmic, replacing a dense full eigendecomposition with a sparse truncated top-k solve so that the advantage grows with dataset size, whereas URD’s 68-fold speedup is an implementation effect from vectorizing an interpreted R loop and would shrink if R ran in-process (**Supplementary Note 6**). Because the benchmarks used small synthetic datasets (300–600 cells) and Monocle3 has no R baseline, these factors are implementation-and input-dependent rather than an intrinsic language-level claim.

RebuildR keeps an auditable record of both successes and failures. Fourteen packages carry a published evolution log totalling 106 iterations, of which 8 were rejected and 6 noted as known limitations, all 14 ending above their parity threshold (**Supplementary Table 30**). For py-Slingshot, the port reached a higher benchmark score than R (0.485 versus 0.455) at a 27-fold speedup over eight iterations with no rejections and two documented limitations, such as R emitting three lineages to the port’s two without affecting pseudotime parity (**Fig. 3g**). For py-URD, two failed iterations first recorded that the R-derived transition matrix was too sparse for a single-root flood, after which the accepted fix—a 16-cell root cluster with a relaxed minimum flooded-cell criterion—raised parity from 0 to 0.985 in one step, giving 0.439 versus 0.348 and a 68-fold speedup (**Fig. 3h**; **Extended Data Fig. 3a**). These logs currently cover 14 of the ∼62 packages.

Reconstruction supplies the algorithms; the upper design layer makes them interoperable. This layer re-designs and optimizes unified APIs so that heterogeneous backends share one interface and, where the algorithm allows, one optimized downstream engine, and PseudotimeFate—the upper-level design behind omicverse.single.Traj—is one example. It treats pseudotime as a common interface: once any backend writes a pseudotime vector into adata.obs, the same workflow proceeds—pseudotime-biased k-nearest-neighbour graph, directed transition matrix, macrostate inference and fate probabilities—whether the pseudotime came from Palantir, Slingshot, SCORPIUS, destiny, URD, Monocle 3 or scTour (**Fig. 3i**). To keep this light, PseudotimeFate uses a sparse PCCA+ strategy (a sparse ARPACK eigensolver plus a deterministic inner-simplex assignment) in pure NumPy/SciPy instead of the heavier CellRank/pyGPCCA Schur-decomposition stack; the approximation is exact for reversible transition matrices and valid for the near-reversible pseudotime-biased graph it is built for, but not for strongly non-reversible chains, where real-Schur GPCCA remains appropriate (**Supplementary Note 7**). On a pancreatic endocrine differentiation dataset (3,696 cells, 3,000 highly variable genes), it identified four mature endocrine terminal macrostates—Beta, Delta, Epsilon and Alpha—at high self-residency (**Fig. 3i; Supplementary Table 31**). Under a standardized comparison with identical input, hardware and warm-up, PseudotimeFate finished in 0.392 s against 33.17 s for the matched CellRank PseudotimeKernel-plus-GPCCA workflow (84.6-fold faster) and recovered all four terminal states at precision and recall of 1.0, versus CellRank’s default of three (**Extended Data Fig. 3f**; **Supplementary Note 8**). This is a lightweight, backend-agnostic fate layer rather than a CellRank replacement: the gap is environment-specific (the install lacked petsc/slepc, with which CellRank’s sparse Krylov–Schur path narrows it to a few seconds), the sub-second figure is the warm cost (a cold call is ∼12 s), and CellRank’s default terminal-state count is tunable.

Through this single interface, PseudotimeFate recovered the expected endocrine fate structure and supported lineage-resolved analysis. UMAP placed annotated cell states, Palantir pseudotime with streamlines and inferred terminal states in one embedding (**Fig. 3i**), and a circular fate projection separated the Alpha, Beta, Delta and Epsilon fates while gathering high-entropy intermediate states at the centre (**Fig. 3j**). Lineage-driver analysis recovered the canonical endocrine markers—Gcg for Alpha (r = 0.593), Pdx1 and Ins2 for Beta (0.498 and 0.491), Sst for Delta (0.658) and Ghrl for Epsilon (0.523), all top drivers at adjusted P below the numerical reporting threshold (**Fig. 3k**; **Supplementary Table 32**)—and dynamic-trend, stream and dynamic-heatmap views summarized branch-specific programs across the four lineages (**Fig. 3k,m**; **Extended Data Fig. 3d,e**). As a demonstration dataset its value is reproduction rather than discovery: terminal-state recovery reached precision and recall of 1.0, and fate probabilities agreed moderately with Palantir on shared lineages (Delta r = 0.72, Beta r = 0.70, Alpha r = 0.79; **Extended Data Fig. 3f,g**; **Supplementary Table 32**).

RebuildR and the unified API layer thus form a two-layer strategy: reconstruct the canonical method ecosystem, then expose its outputs through shared, optimized interfaces. The lower layer turns R-centred pipelines into parity-tested, accelerated and documented Python packages across five ecosystems and 24 methodology families, and the trajectory benchmark shows the ports staying comparable to R while running a median of 22.6-fold faster; the upper layer harmonizes reconstructed and native backends behind shared entry points over one object model (**Supplementary Table 22**), with PseudotimeFate as one example. Acceleration and evolution telemetry currently cover subsets of the catalogue, the PseudotimeFate speedup is environment-dependent, and the pancreas analysis is a demonstration—gaps we expect to close as the loop is run with logging across the full catalogue. This reconstruction connects the ecosystem of Fig. 1 to the benchmark gains of Fig. 2, giving agents the equivalence-gated algorithmic infrastructure that composable biological analysis needs.

### OmicOS composes ecosystem-scale analytical modules to discover biological insights beyond conventional pipelines

OmicOS treats a dataset and a scientific question as a strategy-design problem rather than a fixed pipeline. Its strategy composer clarifies the objective, gates the input through an exploratory-data-analysis check of shape, type and pairing, surveys the available OmicVerse functions and skills, enumerates analytical primitives, generates several candidate workflows, self-audits them for circularity, feasibility, fabricated tool names, compute fit and biological yield, ranks them, and emits a human-readable strategy, a machine-readable plan and a delegation plan (**Fig. 4a**). To test whether the agent could build a workflow from almost nothing, we gave it a deliberately underspecified task: a whole-body mouse spatial transcriptomic dataset and the term “Alzheimer’s disease”, with no analysis plan, no named methods and no assembled reference. The agent designed and executed the workflow itself; the human operator only chose among agent-proposed options and gave corrective feedback at decision points. The full trajectory—preprocessing, gsMap, age-resolved CMap, eQTL/colocalization, sensitivity and cross-species notebooks—is publicly available.

**Figure 4.**
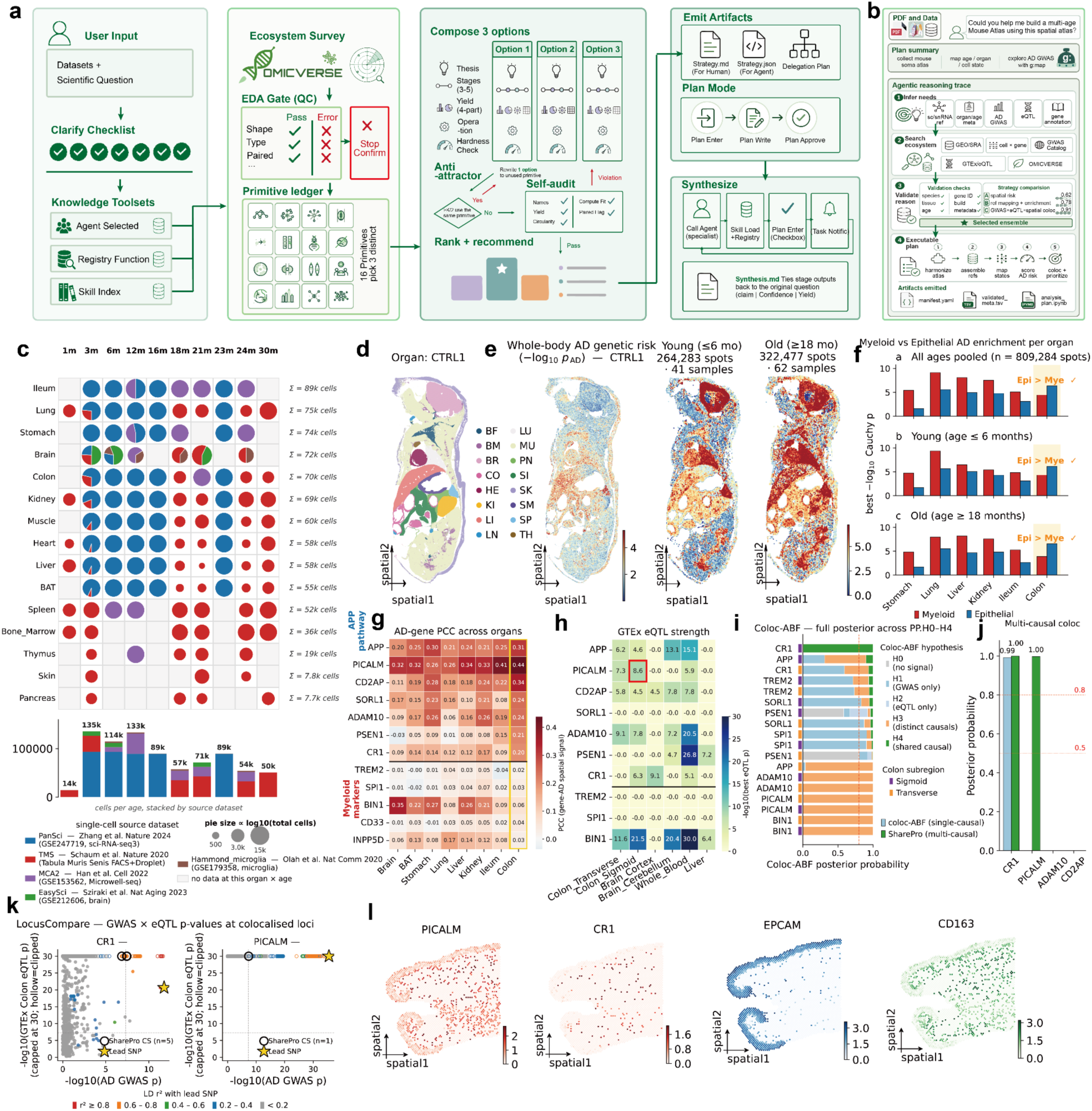
| OmicOS composes a multi-stage spatial-genetic discovery workflow to uncover colon-associated Alzheimer’s disease risk. **a,** OmicOS strategy-composition workflow. OmicOS clarifies the analytical objective, inspects the input data, surveys the OmicVerse ecosystem, registry functions and skill index to enumerate analytical primitives, applies exploratory-data-analysis gating, composes multiple candidate strategies with anti-attractor and self-audit checks, ranks them, and emits a human-readable strategy, a machine-readable plan and a delegation plan; specialist agents then execute the selected workflow stage by stage. **b,** OmicOS-generated plan for the whole-body mouse spatial dataset: collection of a mouse single-cell reference, cell-type mapping onto spots with CMap, and Alzheimer’s disease (AD) genetic-risk mapping with gsMap. **c,** Agent-assembled multi-age mouse single-cell reference. Public single-cell datasets across organs (rows) and ages were collected and harmonized; dot size, cell number; colour, dataset source. **d,** Whole-body spatial annotation of the control mouse section, showing spatial domains for major organs. **e,** Spatial AD genetic-risk scores (gsMap) for the control sample and for young and old age groups across the whole body. **f,** Myeloid versus epithelial contributions to AD genetic-risk enrichment per organ, for all ages pooled, young and old; the colon shows epithelial-dominant enrichment. **g,** Organ-level association between AD risk genes (APP, PICALM, CD2AP, SORL1, ADAM10, PSEN1, CR1, TREM2, SPI1, BIN1, CD33, INPP5D) and cell-type-enriched spatial programs (heat map). **h,** GTEx eQTL strength for the candidate genes across colon, brain, whole blood and liver (heat map). **i,** Colocalization posterior probabilities across AD GWAS and colon eQTL signals (no signal, GWAS-only, eQTL-only, distinct causal, shared causal) under single- and multi-causal models. **j,** Multi-causal (SharePro) shared-causal posterior probabilities for CR1, PICALM, ADAM10 and CD2AP. **k,** LocusCompare plots of AD GWAS versus colon eQTL association for CR1 and PICALM, with lead variants and credible-set signals. **l,** Spatial expression of PICALM and CR1 with the epithelial marker EPCAM and the myeloid marker CD163 in human colon Visium data.

**Extended Data Fig. 4.**
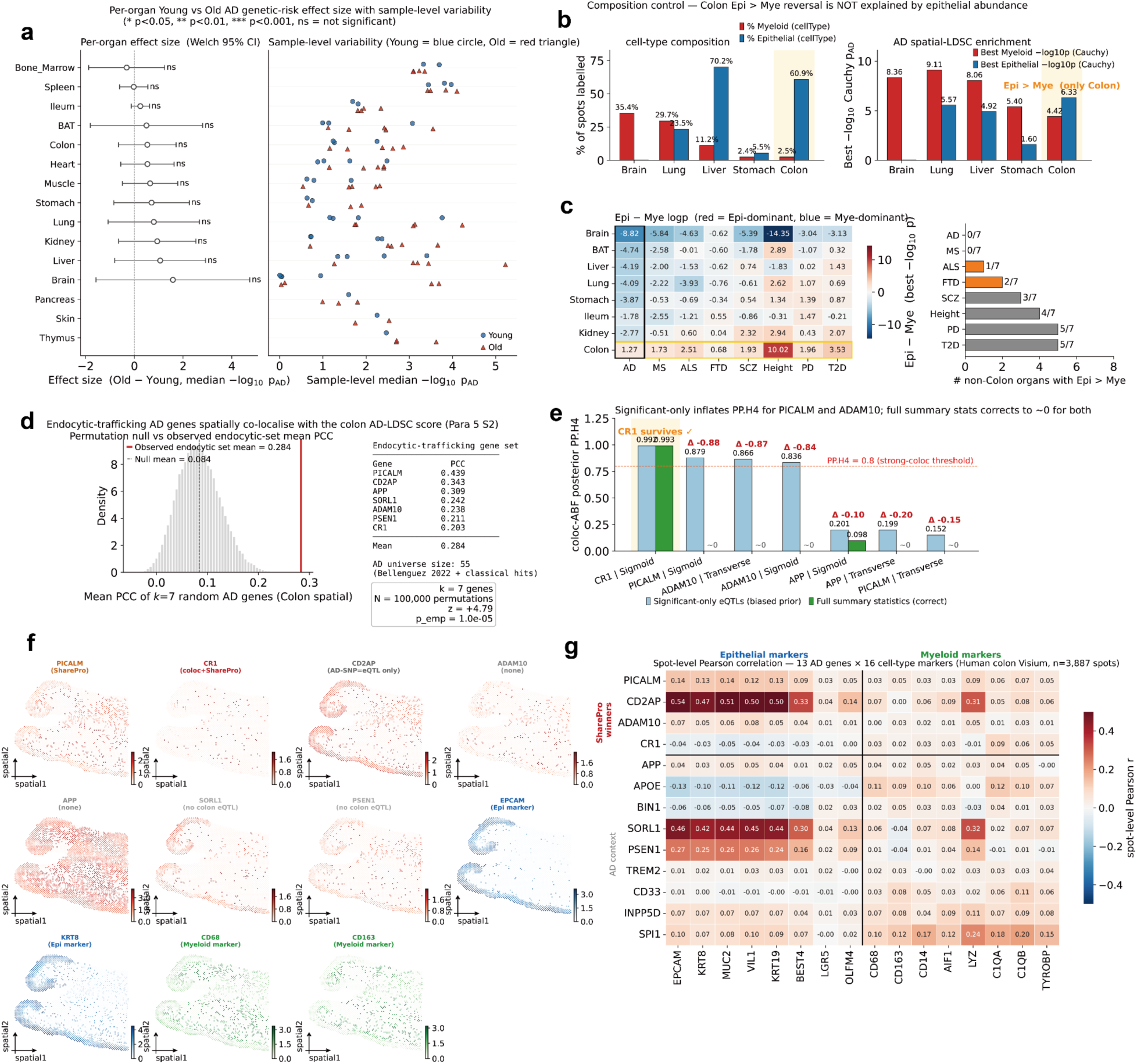
| Sensitivity analyses and cross-dataset validation of the colon epithelial Alzheimer’s disease genetic-risk signal. **a,** Age-associated changes in organ-level Alzheimer’s disease genetic-risk scores. Left, per-organ effect sizes comparing old versus young samples, shown as median differences in −log₁₀ P_AD with Welch 95% confidence intervals; significance levels are indicated above the intervals. Right, sample-level variability for each organ (young, blue circles; old, red triangles). **b,** Cell-composition control for the colon epithelial-over-myeloid reversal. Left, fraction of spots labelled myeloid or epithelial across representative organs. Right, best myeloid and best epithelial Alzheimer’s disease spatial-LDSC enrichment scores (−log₁₀ Cauchy-combined P). The colon is the only displayed organ in which epithelial enrichment exceeds myeloid enrichment, indicating the reversal is not explained by epithelial abundance alone. **c,** Multi-trait comparison of epithelial-versus-myeloid enrichment across organs. Left, heat map of the difference between epithelial and myeloid enrichment scores across traits and organs (red, epithelial-dominant; blue, myeloid-dominant). Right, number of non-colon organs in which epithelial enrichment exceeds myeloid enrichment per trait. **d,** Permutation analysis of endocytic-trafficking Alzheimer’s disease genes in colon spatial genetic-risk maps. The null distribution shows the mean Pearson correlation from randomly sampled seven-gene sets; the observed endocytic-trafficking set exceeds the null. The table lists the candidate genes and their individual spatial correlation coefficients. **e,** Single-causal colocalization using significant-only eQTLs versus full eQTL summary statistics. Bars show coloc-ABF posterior probability for a shared causal variant (PP.H4). Significant-only filtering inflated PP.H4 for several loci, including PICALM and ADAM10, whereas full-summary colocalization corrected these to near zero; CR1 remained strongly supported. **f,** Spatial expression maps for prioritized Alzheimer’s disease genes and cell-type markers in human colon Visium data. Candidate genes are annotated by genetic-support category (SharePro-supported PICALM; coloc- and SharePro-supported CR1; AD SNP–eQTL-supported CD2AP; and genes without colocalization or colon eQTL support). Epithelial markers (EPCAM, KRT8) and myeloid markers (CD68, CD163) provide spatial references. **g,** Spot-level spatial correlation between Alzheimer’s disease candidate genes (rows) and epithelial or myeloid marker genes (columns) in human colon Visium data. Values are Pearson correlation coefficients across 3,887 spots.

**Extended Data Fig. 5.**
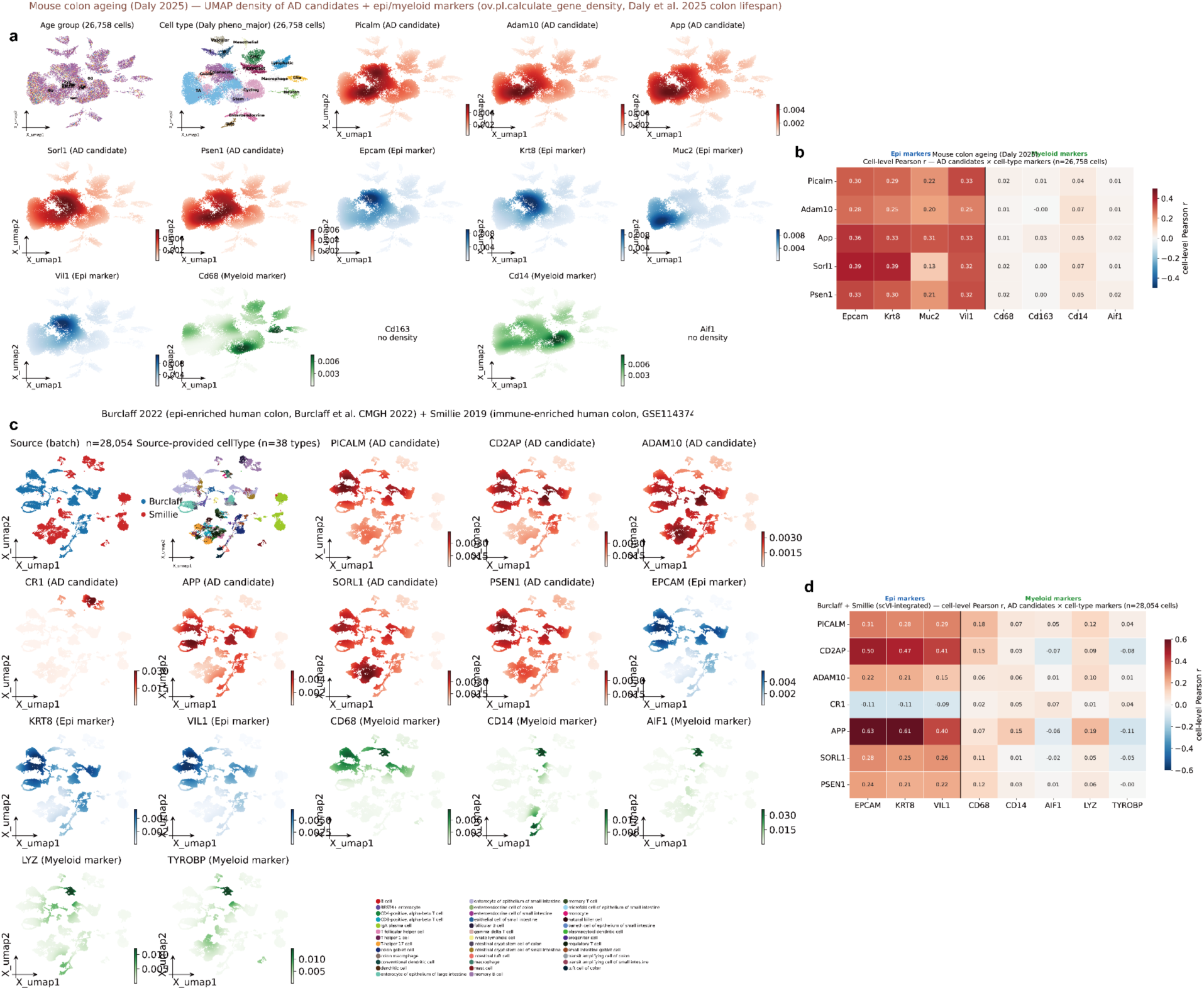
| Single-cell validation of epithelial association for colon Alzheimer’s disease candidate genes. **a,** Gene-expression density maps in a mouse colon ageing single-cell dataset. UMAP embeddings are shown for 26,758 colon cells from the Daly et al. 2025 lifespan dataset, coloured by age group, major cell type, Alzheimer’s disease candidate genes and epithelial or myeloid marker genes. Candidate genes include **Picalm**, **Adam10**, **App**, **Sorl1** and **Psen1**; epithelial markers include **Epcam**, **Krt8**, **Muc2** and **Vil1**; and myeloid markers include **Cd68**, **Cd14**, **Cd163** and **Aif1**. Density patterns show that several candidate genes are enriched in epithelial regions of the manifold rather than in myeloid regions. **b,** Cell-level correlation between Alzheimer’s disease candidate genes and epithelial or myeloid marker genes in the mouse colon ageing dataset. Values denote Pearson correlation coefficients across 26,758 cells. Candidate genes show stronger correlations with epithelial markers than with myeloid markers, supporting an epithelial-associated expression program in mouse colon ageing. **c,** Gene-expression density maps in an integrated human colon single-cell reference combining epithelial-enriched Burclaff et al. 2022 data and immune-enriched Smillie et al. 2019 data. UMAP embeddings are shown for 28,054 cells, coloured by dataset source, source-provided cell type, Alzheimer’s disease candidate genes and epithelial or myeloid markers. Candidate genes include **PICALM**, **CD2AP**, **ADAM10**, **CR1**, **APP**, **SORL1** and **PSEN1**; epithelial markers include **EPCAM**, **KRT8** and **VIL1**; and myeloid markers include **CD68**, **CD14**, **AIF1**, **LYZ** and **TYROBP**. **d,** Cell-level correlation between Alzheimer’s disease candidate genes and epithelial or myeloid marker genes in the integrated human colon reference. Values denote Pearson correlation coefficients across 28,054 cells. Several candidate genes, including **PICALM**, **CD2AP**, **APP**, **SORL1** and **PSEN1**, show stronger association with epithelial markers than with myeloid markers, whereas **CR1** shows weaker epithelial association. Together with the spatial analyses, these single-cell references support a colon epithelial component of the Alzheimer’s disease candidate-gene program.

From the spatial dataset and the disease term alone, OmicOS recognized that the question was addressable by composing two OmicVerse modules—gsMap, to project Alzheimer’s disease GWAS signals onto spatial coordinates by spatial linkage-disequilibrium-score regression, and CMap, to map single-cell reference information onto spatial spots—and that it first needed reference data it did not have (**Fig. 4b**). The agent therefore searched public repositories, downloaded and preprocessed five single-cell datasets, and assembled them across age groups into a reference of 809,284 cells across 126 organ–age combinations spanning 1–30 months: PanSci (GSE247719; 454,620 cells), Tabula Muris Senis, Mouse Cell Atlas 2.0 (GSE153562), EasySci (GSE212606) and a brain-microglia dataset (GSE179358), each projected onto a 6-week whole-body Array-seq spatial reference (**Fig. 4c**; **Supplementary Table 33**). That the agent retrieved and built this reference itself, rather than receiving a pre-made atlas, is what let the analysis separate tissue location, cell-type abundance, ageing and organ-specific gene programs that the spatial data alone could not distinguish.

With the reference in place, gsMap produced per-spot Alzheimer’s disease genetic-risk scores across the anatomically annotated control section (**Fig. 4d,e**). After age-resolved CMap projection, the agent built young and old whole-body risk maps from 264,283 young spots (41 samples) and 322,477 old spots (62 samples); the old-reference map carried a higher median per-spot risk (−log₁₀P 2.12 versus 1.19; **Fig. 4e**; **Supplementary Table 34**; **Extended Data Fig. 4a**). Because per-spot summaries are not independent within a sample, we read this age contrast at the organ and sample level. These maps located where disease-associated risk fell and set up a cell-type-resolved analysis.

Across most organs, genetic-risk enrichment was strongest in tissue-resident myeloid or macrophage-like annotations, consistent with the microglial architecture of Alzheimer’s disease genetics. The colon broke this pattern: epithelial enrichment exceeded myeloid enrichment across all ages (6.33 versus 4.42), in young samples (6.13 versus 4.31) and in old samples (6.55 versus 3.91; Fig. 4f). A composition control argued against an abundance artifact—other non-myeloid-rich organs stayed myeloid-dominant, and the stomach did not reproduce the reversal at a comparable myeloid fraction (**Extended Data Fig. 4b**). To place the colon signal in context, the agent repeated the epithelial-versus-myeloid comparison across eight GWAS traits—Alzheimer’s disease (AD), Parkinson’s disease (PD), amyotrophic lateral sclerosis (ALS), multiple sclerosis (MS), frontotemporal dementia (FD), type 2 diabetes (T2D), schizophrenia (SCZ) and height—and scored, for each trait, how many non-colon organs showed epithelial-over-myeloid enrichment (**Extended Data Fig. 4c**). Because a colon epithelial skew was not unique to Alzheimer’s disease and the colon holds few myeloid cells, the Alzheimer’s-disease-relevant pattern is the conjunction of broad myeloid dominance, the colon exception and the specific risk genes driving the colon program rather than colon epithelial enrichment alone. The agent carried the colon epithelial signal forward to gene-level analysis.

To resolve the signal to genes, OmicOS scored gene-level spatial specificity for Alzheimer’s disease risk genes across organs, highlighting colon-associated patterns for APP, PICALM, CD2AP, SORL1, ADAM10, PSEN1 and CR1 (**Fig. 4g**). In the colon, a seven-gene endocytic-trafficking set co-localized with the colon risk score more strongly than random gene sets (mean Pearson r = 0.284; z = 4.79; empirical P = 1×10⁻⁵ over 100,000 permutations; **Extended Data Fig. 4d**). Because APP and ADAM10 later failed shared-causal colocalization, the agent relabelled this set an endocytic-trafficking and epithelial-barrier candidate program rather than an APP-processing pathway, correcting a label its own figure-reading step had first produced. Candidates were thus prioritized by the intersection of spatial specificity, gene membership, epithelial enrichment and genetic support.

To seek human genetic support, the agent queried GTEx v8 eQTLs and found colon regulatory signals for several candidates: PICALM had 93 colon eQTL variants and no detectable bulk-brain eQTL, CD2AP had 200 colon eQTL variants including an Alzheimer’s disease lead SNP that was itself a colon eQTL, and CR1 had 24 colon eQTL variants overlapping an Alzheimer’s disease SNP (**Fig. 4h**; **Supplementary Table 35**). Single-causal colocalization (coloc-ABF) on full summary statistics supported CR1 as a shared-causal colon signal (PP.H4 = 0.993, colon sigmoid) but placed PICALM on a distinct signal (PP.H3 = 0.999); restricting to significant eQTLs inflated PP.H4 for PICALM and ADAM10, which full-summary colocalization corrected (**Fig. 4i,k**; **Extended Data Fig. 4e**). Multi-causal SharePro then recovered a shared PICALM signal (0.998) and supported CR1 (0.999) but not ADAM10 (0.0005; **Fig. 4j,k**). The genetic evidence is therefore tiered: CR1 has single- and multi-causal support; PICALM has multi-causal support but a distinct single-causal signal; CD2AP has an Alzheimer’s disease SNP–colon eQTL overlap with multi-causal analysis pending; and ADAM10 is SMR-significant but rejected by HEIDI as a linkage artifact (**Supplementary Table 36**). No gene passed simultaneous coloc-ABF, SMR and HEIDI support, and the GTEx colon eQTLs are from bulk tissue.

To corroborate localization, the agent turned to human colon Visium data, where PICALM and CR1 co-localized spatially with the epithelial marker EPCAM and the myeloid marker CD163 (**Fig. 4l**). Across a marker grid of 13 Alzheimer’s disease genes against 16 cell-type markers over 3,887 spots, the epithelial-associated candidates tracked epithelial markers more closely than the classical myeloid genes TREM2, CD33 and SPI1 (**Extended Data Fig. 4g**), and the same pattern held in Human Gut Cell Atlas and mouse colon ageing data (**Extended Data Fig. 5**). This is spatial-expression corroboration, not genetic validation; neither the mouse maps nor the human bulk eQTLs establish the colon epithelium as a causal tissue.

The workflow ended on a hypothesis: a subset of Alzheimer’s disease risk may act through colon epithelial endocytic-trafficking and barrier programs rather than myeloid biology alone, with the exception driven by PICALM, CD2AP and CR1. PICALM is a clathrin-assembly protein with colon regulatory evidence and multi-causal colocalization support; CD2AP is tied to junctional biology and carried an Alzheimer’s disease SNP–colon eQTL overlap; and CR1 had the strongest single-causal colocalization. We present this as a mechanistic model rather than a demonstrated pathway, and the natural next step is functional testing—perturbing PICALM or CD2AP in colon epithelial organoids or barrier models and measuring tight-junction trafficking, permeability and inflammatory response under risk-allele or expression perturbation.

From a minimal input,one spatial dataset and a disease term, OmicOS autonomously identified the applicable modules (gsMap and CMap), retrieved and assembled the reference data they needed, and composed a workflow spanning whole-body spatial transcriptomics, genetic-risk projection, age-resolved single-cell mapping, cell-type and gene-level scoring, GTEx eQTL analysis, single- and multi-causal colocalization, and human spatial-expression corroboration, with the human operator limited to error-correcting judgment. The workflow surfaced a colon epithelial exception to the otherwise myeloid-dominant Alzheimer’s disease risk architecture and nominated CR1, PICALM and CD2AP as colon-supported candidates. A conventional single-track pipeline would not have produced this composition, which extends the benchmark results of Fig. 2 into autonomous hypothesis generation.

### OmicOS-PortBuild converts external pathology packages into agent-usable skills for H&E-to-spatial-omics discovery

OmicVerse provides broad native coverage of omics workflows, but many useful biomedical tools live in external Python packages with their own tutorials, source code, API conventions and notebooks, which a scientific agent cannot use directly: the agent must first learn which APIs are stable, which functions correspond to which biological task, what inputs they need and how their outputs should be read. To make such packages agent-usable without rewriting them into OmicVerse, we built OmicOS-PortBuild, which ingests a package’s tutorials, source repository, API documentation and notebooks and partitions them into capability domains such as whole-slide image analysis, spatial omics, gene-expression analysis, cell segmentation and cross-modality analysis (**Fig. 5a**). As a test case, we asked PortBuild to convert LazySlide, an external H&E pathology-analysis package not adapted to OmicVerse, into OmicOS-compatible agents and skills, so that an external method could become agent-callable through the same interfaces as a native module. (**Supplementary Table 37**)

**Figure 5.**
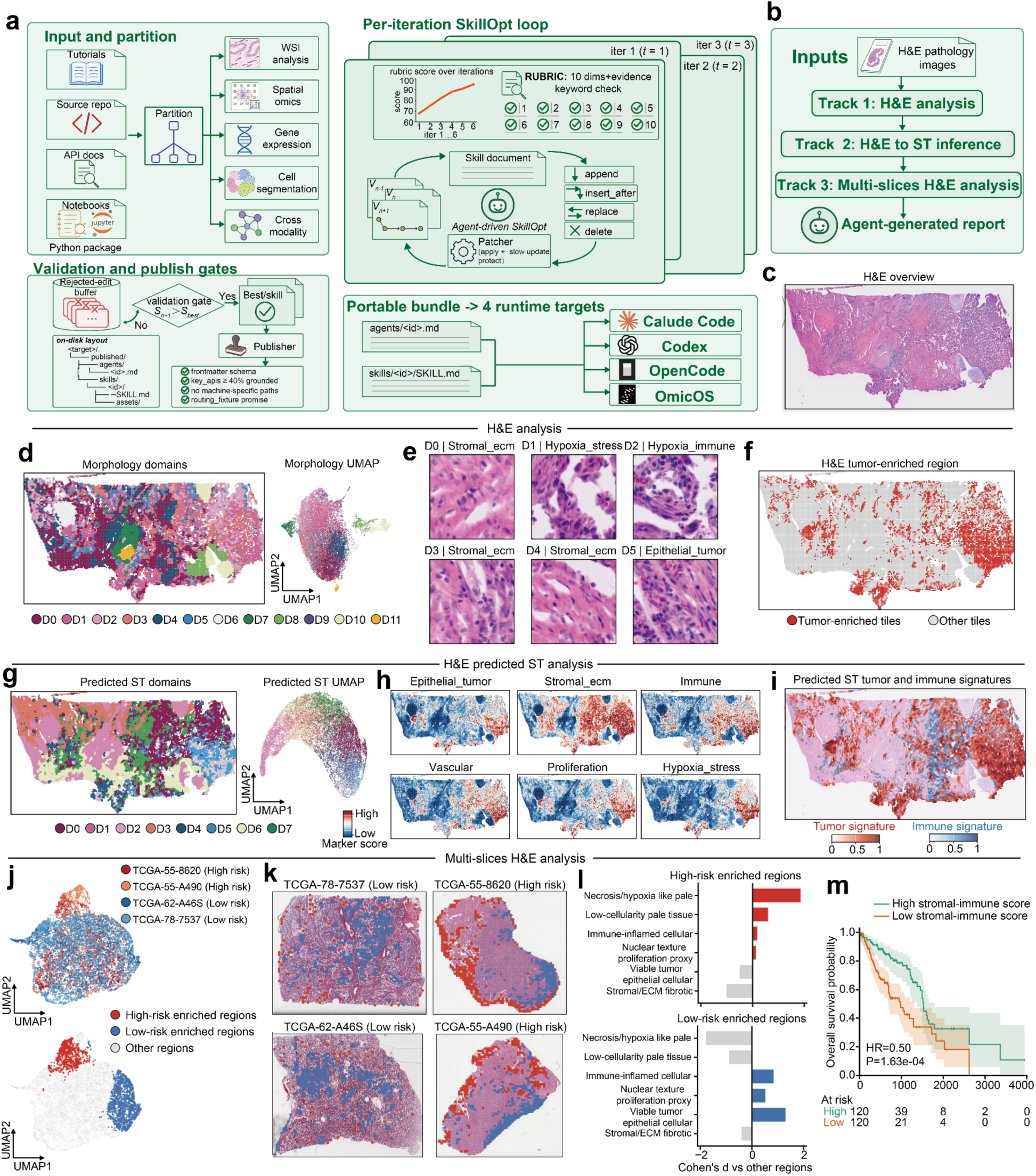
| OmicOS-PortBuild converts an external H&E-analysis package into agent-usable skills for H&E-to-spatial-omics discovery. **a,** Overview of the OmicOS-PortBuild pipeline. External Python packages that are not originally adapted to OmicOS can be converted into portable agent skills by ingesting tutorials, source repositories, API documentation and notebooks. The package is first partitioned into independent capability domains, such as whole-slide image analysis, spatial omics, gene-expression analysis, cell segmentation and cross-modality analysis. Each partition is transformed into an OmicOS-compatible skill through an iterative port-building loop, in which candidate skill documents are refined through protected edit operations, scored by rubric-based checks and accepted only when they improve validation performance. Candidate skills then pass validation and publishing gates, including schema checks, grounded key APIs, machine-path linting and routing-fixture compatibility, before being packaged as runtime-agnostic bundles for Claude Code, Codex, OpenCode or OmicOS. This enables external packages such as LazySlide to become part of the OmicOS execution ecosystem without requiring prior manual adaptation in OmicVerse. **b,** Agent-composed pathology workflow enabled by LazySlide-derived OmicOS-PortBuild skills. Given H&E pathology images as input, OmicOS organizes the analysis into three tracks: direct H&E morphology analysis, H&E-to-spatial-transcriptomics inference and combined downstream analysis, producing an agent-generated report. **c,** Overview of the input H&E-stained whole-slide image used for downstream morphology and inferred spatial-omics analysis. **d,** Morphology-domain analysis from H&E image tiles. Left, spatial distribution of morphology domains across the tissue section. Right, UMAP representation of morphology tiles, coloured by the same domain assignments. **e,** Representative H&E image patches from selected morphology domains. Domains capture distinct histopathological patterns, including stromal extracellular-matrix regions, hypoxia- or stress-associated morphology, stromal-immune regions and epithelial-tumor regions. **f,** Spatial map of H&E-derived cancer-enriched regions. Red points indicate tiles classified as cancer-enriched, whereas grey points indicate other tissue tiles. **g,** Predicted spatial-transcriptomics domains inferred from H&E images. Left, spatial distribution of predicted ST domains across the section. Right, UMAP representation of predicted ST features, with marker-score intensity shown as a continuous annotation. **h,** Predicted spatial-transcriptomics signature maps for major tissue programs. Marker programme scores are shown for epithelial tumor, stromal extracellular matrix, immune, vascular, proliferation and hypoxia/stress signatures. **i,** Overlay of predicted tumor and immune signatures across the H&E section. Epithelial-tumor and immune programme scores were robustly normalized across tiles, and tiles above the 65th percentile for each programme are shown in red or blue, respectively. **j,** Multi-section H&E morphology embedding across high- and low-risk samples. Top, UMAP coloured by sample identity and clinical risk group. Bottom, regions enriched in high-risk or low-risk samples are highlighted on the shared morphology embedding. **k,** Spatial localization of risk-enriched morphology regions across four H&E sections. Red marks high-risk-enriched regions, and blue marks low-risk-enriched regions, shown on representative high- and low-risk samples. **l,** Histopathological features enriched in high-risk and low-risk regions. Bars show Cohen’s d effect sizes comparing each risk-enriched region class with other tissue regions. High-risk-enriched regions are associated with necrosis/hypoxia-like pale morphology and low-cellularity pale tissue, whereas low-risk-enriched regions show comparatively stronger viable-tumor, epithelial-cellular and immune-inflamed morphology. **m,** Kaplan–Meier analysis of overall survival stratified by predicted stromal–immune score. Patients with higher stromal–immune scores show improved survival compared with those with lower scores, with hazard ratio and log-rank P value indicated. Shaded regions denote confidence intervals, and the table reports the number of patients at risk over time.

**Extended Data Fig. 6.**
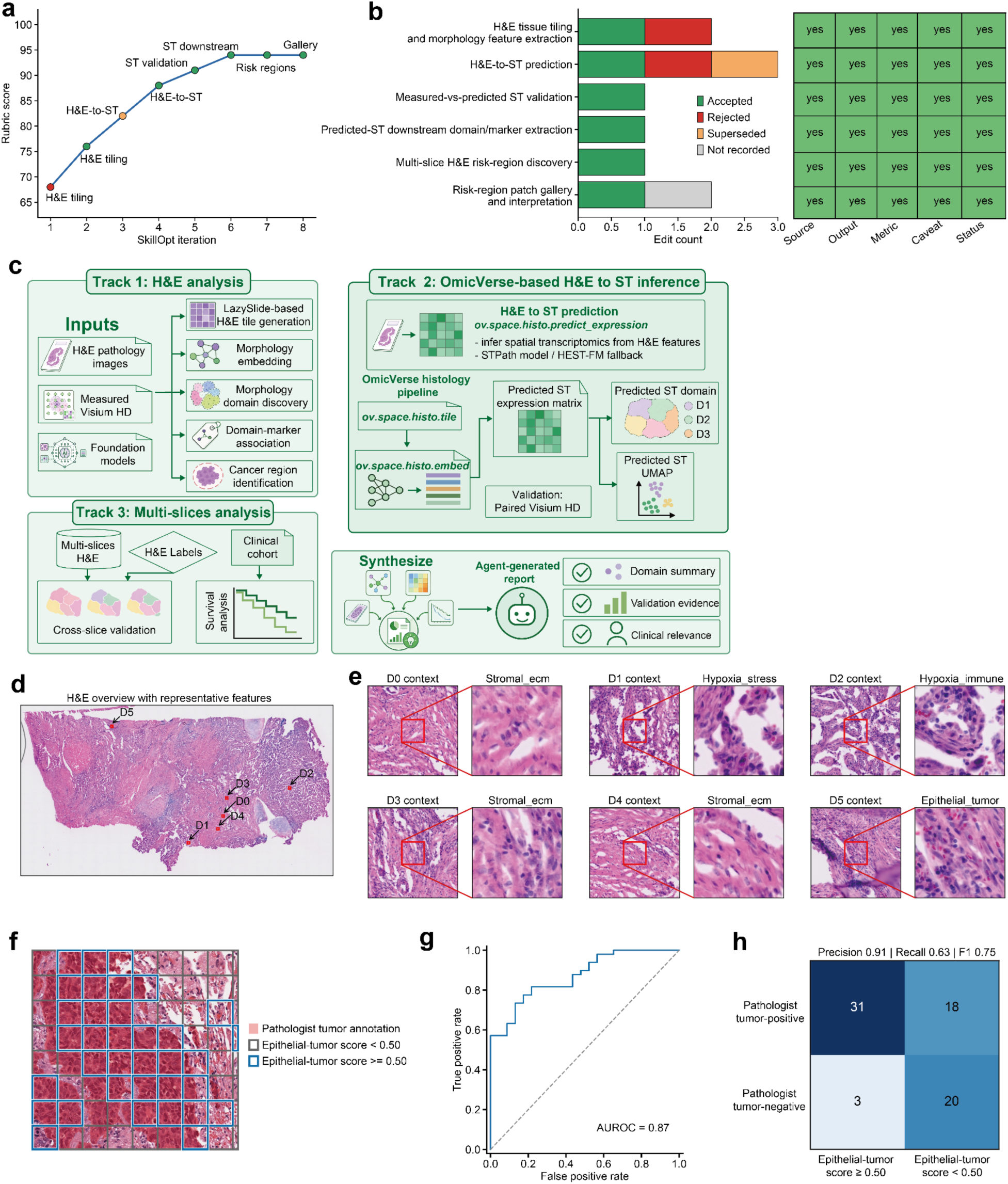
| OmicOS-PortBuild-derived skills optimization, representative H&E regions and external tumor-tile benchmark for the OmicOS pathology workflow. **a,** Detailed schematic of the OmicOS-PortBuild-derived H&E-to-spatial-transcriptomics analysis workflow. The generated skill is organized into three analysis tracks. Track 1 performs H&E morphology analysis using H&E pathology images, measured Visium HD data and foundation-model-derived image features to generate H&E tiles, morphology embeddings, morphology domains, domain-marker associations and cancer-region annotations. Track 2 performs OmicVerse-based H&E-to-spatial-transcriptomics inference by tiling H&E images, extracting histology embeddings, predicting spatial gene-expression profiles and deriving predicted ST domains and UMAP representations, with paired Visium HD data used for validation. Track 3 performs multi-slice H&E analysis by integrating multi-slice H&E images, H&E-derived labels and clinical cohort information to support cross-slice validation and survival analysis. Outputs from the three tracks are synthesized into an agent-generated report summarizing domain composition, validation evidence and clinical relevance. **b,** Per-iteration SkillOpt rubric score trace for LazySlide-derived skill development. The x-axis shows SkillOpt iterations, and the annotated points mark successive workflow capabilities including H&E tiling, H&E-to-ST inference, ST validation, ST downstream analysis, risk-region discovery and patch-gallery generation. **c,** SkillOpt edit history and final validation matrix. Left, horizontal stacked bars summarize accepted, rejected, superseded and unrecorded edits for each derived workflow capability. Right, the validation matrix records whether each capability retained source, output, metric, caveat and status fields in the final report. **d,** Whole-slide H&E overview with selected representative morphology regions. The annotated D0–D5 regions indicate representative H&E-derived morphology contexts selected for downstream high-resolution patch inspection and interpretation. **e,** Representative high-resolution H&E patches from selected morphology contexts. For each selected region, the left image shows the local H&E context and the right image shows the magnified patch used for morphology interpretation. The selected patches are associated with distinct H&E-derived morphology programs, including stromal extracellular-matrix, hypoxia/stress, hypoxia/immune and epithelial-tumor-associated patterns. **f,** External IGNITE Non Small Cell Lung Cancer (NSCLC) tumor-annotation benchmark overlay. The annotated ROI from the IGNITE NSCLC dataset was tiled and scored using the OmicOS H&E tumor-enrichment rule. Pathologist tumor annotation is shown in red, OmicOS-classified tumor-positive tiles are outlined in blue and OmicOS-classified tumor-negative tiles are outlined in grey. **g,** Receiver operating characteristic (ROC) curve for tile-level tumor classification on the IGNITE annotated ROI. The curve evaluates separation of pathologist tumor-positive and tumor-negative tiles. h, Tile-level confusion matrix for the IGNITE benchmark. Counts compare OmicOS tumor-positive and tumor-negative tile calls with pathologist-derived ground-truth tumor-positive and tumor-negative tile labels, with summary classification metrics shown above the matrix.

**Extended Data Fig. 7.**
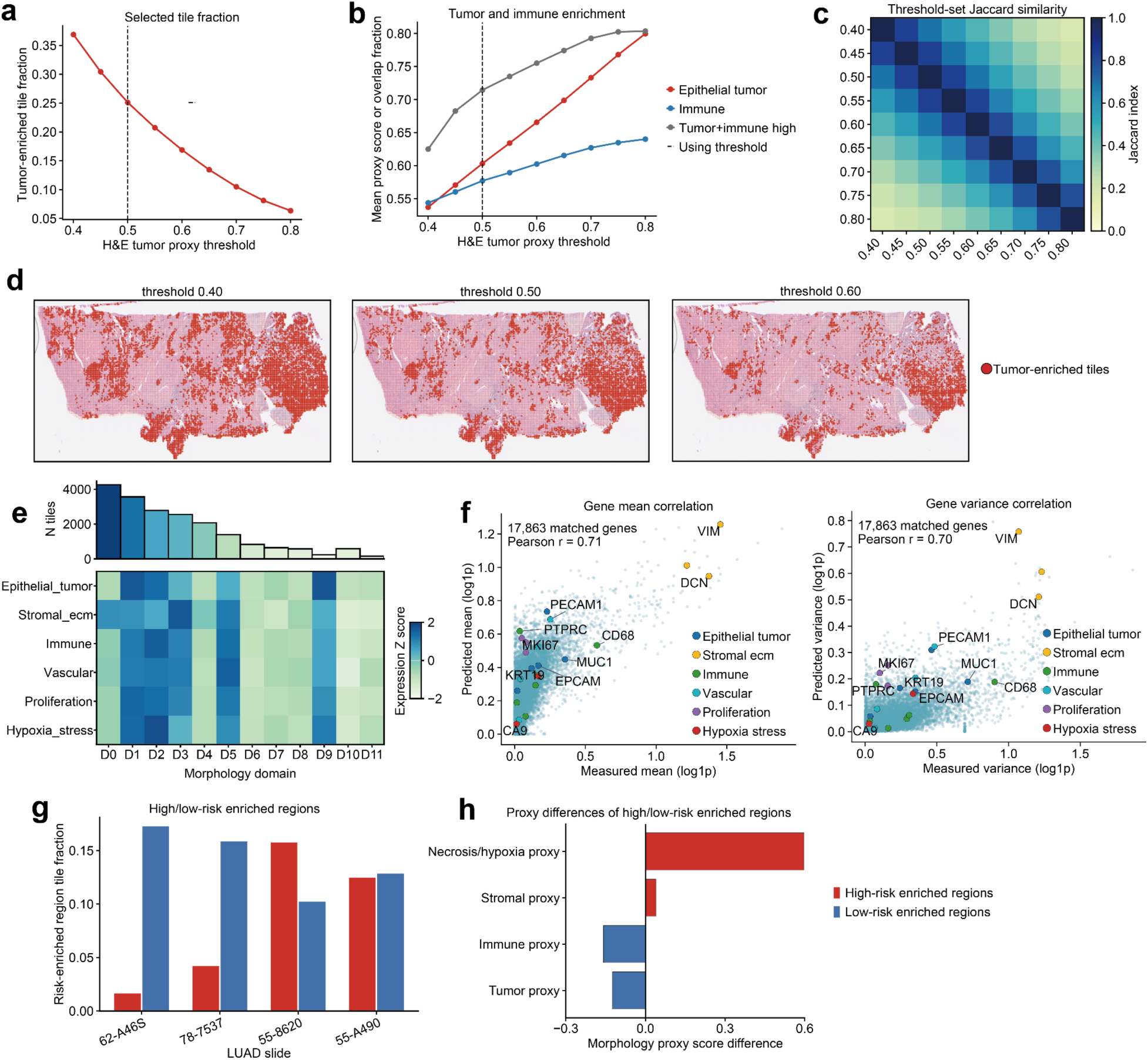
| Threshold sensitivity, H&E-to-ST validation and multi-slice risk-region quantification. **a,** Fraction of H&E tiles classified as tumor-enriched across tumor-enrichment score thresholds. The dashed vertical line marks the main operating threshold used for the corresponding Fig. 5 tumor-enriched region map. **b,** tumor and immune signature enrichment across H&E tumor-proxy thresholds. Lines show the mean epithelial-tumor score, mean immune score and tumor-plus-immune-high overlap fraction among selected tiles, with the main operating threshold indicated by the dashed vertical line. **c,** Jaccard similarity between tumor-enriched tile sets obtained at different H&E tumor-proxy thresholds. The matrix compares threshold-defined tile sets, with higher values indicating greater overlap between selected tile sets. **d,** Spatial maps of tumor-enriched tiles at selected H&E tumor-proxy thresholds. Red points indicate tiles selected as tumor-enriched at each threshold over the same H&E tissue section. **e,** Summary of predicted spatial-transcriptomics domains inferred from H&E images. The bar plot shows the abundance of H&E-predicted ST domains. The heatmap shows marker-program enrichment calculated from the paired Visium HD measurements for the corresponding predicted ST domains, indicating domain-specific epithelial, stromal, immune, vascular, proliferation and hypoxia/stress-associated transcriptional patterns. **f,** Gene-level validation of H&E-predicted spatial expression using measured Visium HD data. Left, each point represents one matched gene, comparing the mean measured expression with the mean H&E-predicted expression across spatial locations. Right, each point represents one matched gene, comparing the measured expression variance with the predicted expression variance across spatial locations. Selected marker genes associated with epithelial tumor, stromal extracellular matrix, immune, vascular, proliferation and hypoxia/stress programs are highlighted, and Pearson correlation coefficients are indicated in each plot. **g,** Fraction of tiles assigned to high-risk-enriched and low-risk-enriched morphology regions across LUAD H&E sections. Bars report the risk-enriched region tile fraction for each slide. **h,** Morphology proxy score differences between high-risk-enriched and low-risk-enriched regions. Bars show mean proxy score differences for tumor, immune, stromal and necrosis/hypoxia proxies, with high-risk-enriched regions showing higher necrosis/hypoxia proxy scores and low-risk-enriched regions showing higher tumor and immune proxy scores in this comparison.

PortBuild refines each capability partition into a skill document through an iterative, auditable loop rather than a one-shot wrapper. For each partition, an agent-driven SkillOpt procedure proposes candidate edits—append, insert-after, replace or delete—applied through a protected patcher that preserves the edit history, and scores each candidate against a ten-dimension rubric with an evidence-keyword check (**Fig. 5a**). Only edits that improve the current best score are accepted, failed edits are kept in a rejected-edit buffer rather than overwritten, and the rubric score rises across iterations (**Fig. 5a**). Before publication, each skill must clear hard gates—frontmatter-schema validation, grounding of at least 40% of its key APIs against real source, removal of machine-specific paths and routing-fixture compatibility—after which it is packaged into a runtime-agnostic bundle of agent definitions, skill documents and assets that deploys to Claude Code, Codex, OpenCode or OmicOS (**Fig. 5a**). The integration is therefore a reproducible port-building procedure, leaving each skill inspectable and portable across execution environments. (**Extended Data Fig. 6a-b**)

Using the LazySlide-derived skills, OmicOS composed an H&E pathology workflow in three tracks (**Fig. 5b; Extended Data Fig. 6c**). Given H&E whole-slide images, the first track quantified morphology directly from the stained image, the second inferred spatial-transcriptomic programs from image features, and the third combined morphology-derived and predicted spatial-omics features across sections for clinical interpretation, ending in an agent-generated report. Because LazySlide had been converted into agent-usable skills, OmicOS could run these pathology and spatial-omics analyses even though the package was never natively designed for the OmicOS ecosystem, bringing H&E images into the same agentic framework used for omics analysis. (**Supplementary Table 38**)

The first track recovered spatially organized morphology from the whole-slide image (**Fig. 5c**). Tile-level analysis partitioned the section into twelve morphology domains and embedded the tiles in a morphology UMAP that showed both spatially localized and manifold-level structure (**Fig. 5d**), and representative patches gave interpretable domain identities—stromal extracellular-matrix, hypoxia- or stress-associated, hypoxia-immune and epithelial-tumor morphology (**Fig. 5e; Extended Data Fig. 6d-e**). Mapping cancer-enriched tiles back onto the section showed that tumor-associated morphology occupied spatially restricted territories rather than spreading uniformly across the tissue (**Fig. 5f; Extended Data Fig. 6f-h and Extended Data Fig. 7a-d**).The second track turned morphology into predicted molecular programs. From the H&E image alone, OmicOS inferred eight predicted spatial-transcriptomic domains and a predicted ST UMAP (**Fig. 5g**) and recovered marker-score maps for major tissue programs—epithelial tumor, stromal extracellular matrix, immune, vascular, proliferation and hypoxia/stress (**Fig. 5h**). Overlaying the predicted tumor and immune signatures revealed spatial heterogeneity, with tumor-dominated regions, immune-enriched regions and zones where the two programs were adjacent or overlapping (**Fig. 5i**). These programs are inferred rather than directly measured, so the maps generate molecular hypotheses from histology where matched spatial-transcriptomic data are unavailable rather than replacing such measurements. (**Extended Data Fig. 7e-f**)

The third track extended the analysis across four H&E sections from high- and low-risk patients. OmicOS embedded tiles from all sections in a shared morphology space and coloured it by sample and clinical risk group; high-risk- and low-risk-enriched regions separated within this shared embedding, indicating that risk-associated tissue patterns recurred across sections rather than being confined to one image (**Fig. 5j**). Projecting these regions back onto the four sections (TCGA-55-8620 and TCGA-55-A490, high risk; TCGA-62-A46S and TCGA-78-7537, low risk) showed that high-risk- and low-risk-enriched regions occupied distinct spatial territories within each section (**Fig. 5k**). (**Supplementary Table 39**)

The risk-enriched regions differed in histopathology. Relative to other tissue, high-risk-enriched regions were most strongly associated with necrosis- or hypoxia-like pale morphology and low-cellularity pale tissue (positive Cohen’s d), whereas low-risk-enriched regions were associated with viable tumor and immune-inflamed cellular morphology (**Fig. 5l**). These contrasts indicate that the H&E-derived morphology features capture clinically meaningful variation in tissue state—hypoxia, cellularity, stromal organization and immune contexture—though the feature labels are inferred from image-derived domains and would benefit from expert pathology review. (**Extended Data Fig. 7g-h**; **Supplementary Table 40.**)

Finally, we tested whether an H&E-derived feature carried prognostic information by stratifying patients on a predicted stromal–immune score from the morphology analysis. Patients with a high stromal–immune score had better overall survival than those with a low score (hazard ratio 0.50; log-rank P = 1.63 × 10⁻⁴; 120 patients per group at baseline; **Fig. 5m**). This single-cohort, univariate association would need confirmation with multivariable Cox models adjusting for stage, age and treatment, and validation in an independent cohort, but it shows that the PortBuild-enabled workflow can produce a clinically meaningful readout rather than image-domain features alone. (**Supplementary Table 41**)

Together, these analyses show that OmicOS is not closed to its native ecosystem. PortBuild converted the external LazySlide pathology package into portable skills through capability partitioning, rubric-guided iterative refinement, validation gates and runtime-agnostic publishing, and those skills let OmicOS compose a pathology workflow spanning morphology-domain analysis, H&E-to-spatial-transcriptomics inference, tumor–immune signature mapping, multi-section risk-region analysis and survival association. The workflow recovered interpretable morphology domains, predicted molecular tissue programs, risk-associated histological regions and a stromal–immune score associated with overall survival. Fig. 5 thus extends OmicOS from a reconstructed omics ecosystem to an extensible agent platform that can absorb external tools and apply them to pathology-to-spatial-omics discovery.

## Discussion

Here we show that reliable agentic omics requires a state-aware execution substrate, rather than a language model that is simply larger or prompted with more context. OmicOS provides such a substrate by linking the physical layer of omics data to an agent-facing execution layer. At its base, AnnDataOOM and MuDataOOM preserve compatibility with AnnData and MuData while replacing the in-memory assumption with a Rust-backed out-of-memory implementation, allowing large atlas-scale objects to remain on local disk while only the active working set is brought into memory^9,10,51^. Above this storage layer, OmicOS reconstructs and unifies a broad OmicVerse method ecosystem, exposes functions through a machine-readable registry, and couples them to capability contracts that declare the biological state a method requires, the state it produces, its parameters, dependencies and side effects^13,52^. This changes the role of an omics method in an AI system. Rather than being an isolated callable selected from documentation, each method becomes a validated transition between biological object states: raw counts are transformed into normalized matrices, matrices into embeddings and graphs, and graphs into clusters, trajectories, cell-type annotations, genetic-risk maps or spatial programs. The conceptual advance is therefore a shift from tool use to scientific execution. Workflow engines have established provenance and scale, and omics libraries have established powerful analytical methods; OmicOS connects these ideas into a live, state-aware biological execution space in which a language model can search, rank, validate and execute operations while preserving lineage and recovering from typed failures^39,53,54^. In this sense, the central contribution of OmicOS is not a single algorithm or benchmark result, but a bounded substrate that converts model reasoning into reproducible omics action.

Omics provides an unusually stringent test for scientific agents because the challenge is not only to interpret biological concepts, but to execute valid operations inside stateful, heterogeneous and resource-constrained data systems^21^. A single analysis can span raw sequencing inputs, gene identifiers, batch covariates, embeddings, neighbourhood graphs, spatial coordinates, reference atlases, GWAS and eQTL evidence, image-derived features and foundation-model outputs. Despite this conceptual accessibility, conventional agents remain operationally fragile because method applicability in omics is often encoded in object state rather than in the function signature^20^. A cell-annotation step may require clusters, a trajectory method may require a neighbour graph, a velocity model may require spliced and unspliced layers, and a spatial workflow may require matched coordinates and tissue metadata; when these states are absent, free-form scripts fail as missing prerequisites, invalid arguments, dependency errors or malformed objects. Scale adds a second constraint: atlas-scale single-cell and spatial datasets increasingly exceed ordinary memory budgets, even though an agent must still inspect, subset, visualize and update them interactively^55^. AnnDataOOM and MuDataOOM address this constraint by preserving the familiar AnnData/MuData programming model while changing the memory model beneath it, keeping data on local disk and bringing only the working set into memory through a Rust-backed out-of-memory implementation^51^. Thus, the current limits of biomedical agents follow naturally from the omics setting: general-purpose agents have learned to reason, call tools and edit code, but their realized capability depends on the harness that converts reasoning into valid execution^22,56,57^. The same language model can therefore appear competent or brittle depending on whether its harness supplies a validated biological action space.

The central novelty of OmicOS is that it builds a five-layer, interdependent infrastructure for AI-era omics, in which each layer supplies the state, methods, contracts, execution semantics and extensibility required by the next. The first layer is the scalable data substrate: AnnDataOOM and MuDataOOM preserve the AnnData/MuData programming model while replacing the assumption that atlas-scale objects must fit in memory. The second layer is the reconstructed method substrate: OmicVerse and OmicVerse-RebuildR bring Python-native workflows, historically R-centred Bioconductor and CRAN methods, and domain-specific omics tools into a unified execution ecosystem, treating reference implementations as executable specifications and admitting ports only through output-class-aware equivalence tests^6,58^. The third layer is the registry and contract layer: raw callables are lifted, compiled, normalized, indexed, ranked and emitted as capability manifests, so methods are no longer passive documentation entries but typed operations over live biological state. The fourth layer is the agentic execution layer: OmicOS evaluates omics agents as model–harness–ecosystem systems, and the benchmark ablations show that registry-grounded execution, rather than documentation retrieval or ungrounded tool exposure, accounts for the gain. The fifth layer is the extensibility and discovery layer: PortBuild can convert external packages, tutorials, API documentation and notebooks into runtime-portable skills, while the spatial Alzheimer’s disease workflow shows that the same substrate can compose genetic-risk mapping, single-cell references, eQTL analysis, colocalization and spatial validation into a hypothesis-generating analysis. These layers are iterative rather than additive: scalable data access enables method execution, reconstructed methods define the action space, contracts make actions valid, the runtime turns valid actions into traceable workflows, and extensibility feeds new tools and discoveries back into the ecosystem. In this architecture, OmicOS is not a collection of wrappers but an operating infrastructure for scientific execution in which omics data, methods, agents and provenance co-evolve within a single inspectable substrate.

This distinction explains why OmicOS cannot be reduced to either expert-operated scripts or static wrappers. Expert users can, of course, assemble individual analyses from Scanpy, Bioconductor, spatial-omics, genetic-association and pathology tools^6,7^; what is missing is a shared live state in which each operation declares what it consumes, what it produces and how subsequent operations inherit that result. Static wrappers also expose tools, but exposure alone is not execution validity: in the OmicBench ablation, providing named OmicVerse tools without registry grounding did not reproduce full-system performance and, for GPT-5.5, reduced Pass@1 below baseline. OmicOS instead preserves continuity across layers, linking out-of-memory atlas handling, preprocessing, trajectory inference, reconstructed R methods, foundation-model calls, spatial genetic mapping, eQTL and colocalization analysis, H&E morphology analysis, H&E-to-spatial-transcriptomic inference and survival readouts within one provenance-preserving substrate. This continuity allows hypotheses to propagate across tool boundaries without being manually reassembled after each package, notebook or runtime transition. The goal is therefore not to automate away scientific judgement, but to make human–AI collaboration auditable: agents can propose and execute traceable workflows, while investigators inspect object state, provenance, assumptions and typed failure classes instead of reverse-engineering free-form scripts.

Several limitations should be considered when interpreting these results. First, the main constraint of the registry is not whether OmicVerse functions can be made addressable, but whether their metadata remain synchronized with evolving APIs, dependencies, side effects and biological prerequisites. As the ecosystem expands, capability contracts will require continuous auditing so that agent-executable metadata continue to reflect both the underlying code and current scientific practice. Second, although the benchmark evidence spans external and internal suites and the ablations support registry-grounded execution as the active mechanism, the evaluation remains incomplete: some task families contain few examples, external cost–performance comparisons were not fully remeasured under a single infrastructure, and residual failures persist in spatial and foundation-model workflows. Future benchmarks should therefore emphasize harder multimodal, longitudinal and perturbational tasks, together with standardized cost, runtime and leakage controls. Third, equivalence-gated reconstruction provides a practical software guarantee rather than a proof over all biological inputs. RebuildR compares Python ports against R references using fixtures and output-class-aware parity tests, and its acceleration loop rejects rewrites that break equivalence; nevertheless, broader coverage of edge cases, stochastic outputs, unusual object structures and atlas-scale datasets will be needed as the catalogue grows. Fourth, the discovery workflows should be interpreted as hypothesis-generating rather than mechanistic proof. The Alzheimer’s disease colon epithelial signal is supported by spatial genetic-risk mapping, organ- and cell-type analysis, GTEx eQTL evidence, colocalization and human spatial-expression corroboration, but bulk-tissue eQTLs, cross-species mapping and colocalization assumptions cannot establish tissue causality; perturbation in colon epithelial organoids or barrier models will be required to test mechanism. Fifth, the H&E workflow is clinically suggestive but not yet a validated pathology model. The stromal–immune survival association is single-cohort and univariate, predicted spatial-transcriptomic programs require calibration against measured spatial omics, and morphology labels require pathologist review and independent validation. Finally, increased autonomy creates governance requirements. OmicOS improves provenance, inspection and recoverability, but scientific agents that search public resources, execute workflows and emit biological hypotheses will still require safeguards against benchmark leakage, hallucinated interpretation, privacy risks, unsupported clinical claims and dual-use misuse^59^.

Looking forward, OmicOS points to a general playbook for AI-era omics: major analytical operations should be exposed not as code fragments, but as registry-grounded state transitions with explicit assumptions, dependencies, outputs, provenance and validation checks. A key next step is uncertainty-aware execution, in which method selection, object-state validity, statistical inference, spatial mapping, colocalization and biological interpretation carry calibrated confidence so that agents can distinguish well-supported conclusions from weak extrapolations. OmicOS also provides a natural interface to biological foundation models and AI virtual-cell efforts: foundation models can learn representations and predict cellular behaviour, whereas a state-aware execution substrate is needed to connect those predictions to real datasets, verified tools, perturbation workflows and interpretable outputs^60,61^. Prospective evaluation will be essential, moving benchmarks from retrospective task completion to settings in which agents design analyses before outcomes are known, select validation experiments, handle held-out perturbations and revise hypotheses when evidence contradicts their initial interpretation. More broadly, OmicOS suggests a shift from AI systems that write omics code to AI systems that operate within validated omics infrastructure. In this view, the central object is not a single model, workflow engine or package, but an executable ecosystem in which models, methods, data objects and provenance are coupled. Such infrastructure is needed for AI-era omics to move from compelling demonstrations towards reliable biological discovery.

## Material and Methods

### Design principles and overview

OmicOS was designed around a single principle: realised omics-agent capability is a property of a model–harness–ecosystem system rather than of the language model alone, so the analytical action space exposed to an agent must be bounded, type- and state-aware, and reconstruction-verified. OmicOS is organised as five interdependent layers, each supplying the state, methods, contracts, execution semantics or extensibility required by the next: (i) a scalable out-of-memory data substrate (AnnDataOOM/MuDataOOM); (ii) a reconstructed method substrate (OmicVerse v2 together with OmicVerse-RebuildR and OmicOS-PortBuild); (iii) a machine-readable capability registry and contract layer; (iv) a live runtime that grounds, validates, executes and traces operations; and (v) an extensibility and discovery layer. Throughout, the AnnData and MuData objects are the shared physical state on which all operations read and write^7–10^. Unless stated otherwise, single-cell preprocessing followed the canonical OmicVerse chain — quality control → normalisation and log transformation → highly variable gene (HVG) selection → scaling → principal component analysis (PCA) → neighbourhood graph → Leiden clustering → uniform manifold approximation and projection (UMAP) → annotation — with each step’s produced object state satisfying the next step’s required state. All software versions, parameter defaults and dataset accessions are given in the relevant subsection below, and a consolidated list of methods-only references continues the main reference numbering from^62^.

### Out-of-memory AnnData and MuData backend

AnnDataOOM (anndataoom v0.1.8) is a Rust-engineered, disk-backed store that is API-compatible with anndata.AnnData^8^ and registered as an abstract subclass so that isinstance(obj, AnnData) resolves to True. The expression matrix X is held on disk in HDF5 (.h5ad) through the vendored scverse/anndata-rs engine (revision aa15c8d, the same I/O layer used by SnapATAC2^62^), exposed to Python as a PyO3 extension; only the working set is materialised in RAM. Dense (float32/float64), CSR and CSC layouts are supported, and sparsity is preserved through normalisation and the log transform (densification occurs only where algorithmically required, e.g. z-scoring and PCA). Zarr is not yet implemented.

The backend represents adata.X as a lazy chain of operator nodes — a backed-array root performing real HDF5 I/O, with subset, transform (size-factor normalisation plus a log1p flag) and scale (per-gene mean μ, standard deviation σ, optional clip) nodes layered above it. Reads pull data downward through the chain in chunks (default chunk_size, 1,000 rows) and transformed chunks flow upward, fusing all pending transforms in a single streaming pass so that peak working-set memory is chunk_size × n_vars, independent of n_obs. Stacked subsets are flattened to a single node so subsetting is O(1). Per-cell total-count normalisation is applied as a left multiplication by a sparse diagonal of inverse size factors,

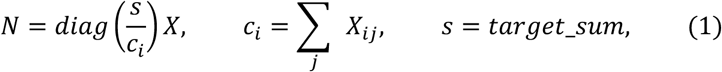

where target_sum, when unset, defaults to the median of non-zero per-cell counts, followed by the sparsity-preserving transform 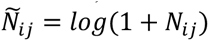 (2). Per-gene mean and variance are computed in one chunked pass using a sparse-native *E*[*X*^2^] − *E*[*X*]^2^ identity within each chunk and Welford’s numerically stable accumulation across chunks in float64^63^ returning sample variance (relative error floor ≈ 7 × 10⁻⁸). Scaling stores only the per-gene μ and σ vectors plus an optional clip (default max_value, 10).

PCA over the lazy matrix uses randomised singular value decomposition^64^. When the post-HVG scaled block fits a configurable threshold (default 16 GB) it is materialised once — slicing the ≈2,000 HVG columns off the sparse parent *before* densification (valid because normalisation is per-row and log1p is element-wise) so that only a chunk × n_HVG block is ever dense — and factorised by randomized_svd; otherwise an implicit-centering path rewrites every matrix–vector product through

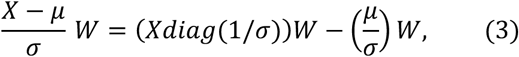

so that no dense n_obs × n_vars buffer is allocated, using a hand-rolled Halko loop with n_power_iters (default 4) and n_oversamples (default 10). The first 50 components were retained (n_pcs, 50). Top components were numerically identical to standard PCA (top-PC |cos| = 1.0; max abs difference ≈ 10⁻⁷). MuDataOOM (mudataoom) is a strict subclass of mudata.MuData^10^ whose per-modality matrices are AnnDataOOM objects; read_h5mu(…, backed=“r”) walks /mod/* and builds one virtual .h5ad per modality via HDF5 external links into the source .h5mu, so X is streamed without copying. Downstream graph and embedding steps optionally offload to GPU through PyTorch (CUDA/MPS) and PyTorch Geometric, and PCA for in-memory objects dispatches to a PyTorch solver; the out-of-memory preprocessing operators are pure-CPU and mode-invariant.

Benchmarks used seven Tabula Sapiens^65^ slices from CZ CELLxGENE Discover[66] (collection e5f58829-1a66-40b5-a624-9046778e74f5) on a shared 60,606-gene panel, ranging from 5,000 to 1,053,033 cells (the 1M object formed by concatenating the stromal, epithelial and immune compartments). Relative to dense in-memory AnnData, peak resident memory fell from ≈120 GB (out-of-memory failure) to ≈0.7–5.0 GB at 1.05 × 10⁶ cells (≈17–170× across datasets); the 1M-cell object completed the full pipeline within the reported wall-clock at single-node memory. Runs were performed under a 256 GB per-process cap (CPU node) with the GPU-mixed comparison on an NVIDIA H100.

### The OmicVerse v2 analysis ecosystem

OmicOS is built on OmicVerse^13^ (omicverse v2.2.1rc1), which exposes 694 analysis methods across 14 modules under one ov.* namespace over a shared AnnData/MuData model: ov.single (151 methods), ov.airr (102), ov.metabol (64), ov.genetics (59), ov.space (48), ov.micro (41), ov.external (39), ov.llm/ov.fm (38), ov.epi (36), ov.alignment (32), ov.pp (31), ov.bulk (25), ov.es (16) and ov.mol (12). The data model is AnnData (extended to MuData for multimodal data), with pandas.DataFrame retained where appropriate (ov.bulk, ov.genetics). Fixed slot conventions are enforced across backends (pseudotime in adata.obs, embeddings in adata.obsm, result matrices in obsm/layers, method metadata in adata.uns). More than 21 single-cell foundation models are interfaced through a model-factory dispatcher, six with local runtimes (scGPT^18^, Geneformer^61^, scFoundation^17^, CellPLM^17,66^, UCE and scMulan^67^), each writing an obsm[’X_<MODEL>’] embedding. GPU-accelerated preprocessing is provided for two backends selected at runtime — PyTorch (CUDA/MPS/ROCm/XPU) and Apple MLX (mlx.core, device mps) with a scikit-learn fallback. Representative defaults used throughout were ov.pp.preprocess (mode=’shiftlog|pearson’, n_HVGs=2000), ov.pp.scale (max_value=10), ov.pp.pca (n_pcs=50, layer=’scaled’), ov.pp.neighbors (n_neighbors=15, use_rep=’X_pca’) and ov.pp.leiden (resolution=1.0); batch integration used Harmony^68^ and gene-regulatory-network inference used GRNBoost2^69^.

### Capability registry and machine-readable contracts

Each raw OmicVerse callable is lifted into a machine-readable *capability object* by an import-time decorator (@register_function) that validates metadata and introspects the live signature and docstring. A capability object encodes a nine-field contract: aliases; category and description; a parameter schema (signature with defaults); requires (the input object state that must be present, with keys drawn from the allowlist {layers, obsm, obsp, uns, var, obs, varm}); prerequisites (hard functions and soft optional_functions that must or should have run first); dependency constraints; produces (the object fields written, using the same allowlist); the declared return structure; and an error policy auto_fix ∈ {none, auto, escalate}. Registration requires at least one alias, a non-empty category and description, a legal auto_fix, and allowlist-valid contract keys. Each entry is indexed under its full name, short name and every alias for constant-time resolution. Virtual entries are derived automatically from the abstract syntax tree of the registered callable: every public method of a registered class becomes its own entry, and a simple string-dispatch if/elif branch (e.g. if method==“harmony”) becomes one synthetic entry per branch (e.g. batch_correction[method=harmony]), gated by an allowlist of capability-bearing parameters (method, backend, mode, source, task, model, …).

Three discovery mechanisms populate the registry: (i) runtime registration through the import-time decorator; (ii) static abstract-syntax-tree scanning of unimported source, which literal-evaluates decorator arguments with no import side effects and is cached and invalidated by file modification time; and (iii) metadata hydration that merges the two, unioning list and dict fields with deduplication, back-filling empty scalars and letting the runtime source win on conflict. Auditing the public API, 231 callables were promoted to capability objects across nine categories, with a mean of 4.3 aliases each (median 5; range 2–12), including multilingual aliases such as 质控 and 手动注释; 100% carry discovery metadata and at least one worked example, and 45.9% (≈106 functions) additionally declare a full requires/produces/prerequisites/auto_fix contract, concentrated in the preprocessing and single-cell layers. The broader @register_function registry of 788 functions is the surface that ov.Agent and the Model Context Protocol (MCP) server expose, with the MCP tool manifest built directly from the registry. Metadata completeness, but not runtime correctness, was summarised by an agent-friendliness score (AFS) defined as a weighted sum of four 0–1 sub-scores — discoverability D, contract clarity C, signature clarity S and behaviour B:

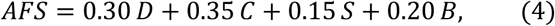

with a library-wide mean of 0.61 (median 0.58).

### Capability discovery, ranking and the agent action space

For a free-text task, candidate capabilities are ranked by a seven-component structured key — exact-name match (4.0 for a full/registry-name match, 3.0 for a short-name or alias match), alias overlap, name overlap, a structural kind preference (parent function > class method > dispatch branch), a weak score over description/docstring/examples, a contract-text overlap and a source bias (runtime > static) — collapsed to a scalar

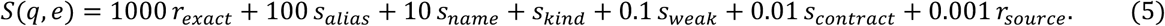

Queries are tokenised over mixed Latin and Chinese (CJK) character ranges so that multilingual aliases match, and a fuzzy fallback (cutoff 0.6) resolves near-misses. The top entries are rendered into a compact agent block (name, signature, description, hard prerequisites, requires/produces maps and an example); prerequisites and contracts are surfaced for planning and are not used to silently filter or gate execution. The resulting manifest gives an agent a bounded action space of type- and state-aware operations rather than free-form code^52^.

### The OmicOS runtime and contract-localized failure recovery

The OmicOS runtime (omicos-core v0.2.0) is a Rust daemon that hosts a shared IPython kernel, exposes an HTTP/server-sent-events contract and presents chat, command-line, notebook, graphical-workspace and programmatic surfaces over one live analytical state. Each chat turn runs a single tool loop (iterate_turn): the runtime estimates tokens and auto-compacts before the model call when above a threshold (default ≈70% of the model context window, at most twice per turn), streams the model response, and — when tool calls are returned — records the assistant message and tool-use blocks into an append-only trajectory store, dispatches each tool through one shared executor, appends the results and continues until a tool-free answer or the iteration cap (max_tool_iterations, default 100). Specialist delegation (call_agent) spawns a child turn with a fresh history and a new execution context, bounded by an agent-depth cap (MAX_AGENT_DEPTH, 6) with ancestor-cycle detection; an optional output schema makes the child reply with validated JSON, which is the runtime’s structured-output path. Provenance is captured as an append-only trajectory (system prompt once, user and assistant messages, full tool results, compaction summaries and sidechain forks) together with per-call usage records.

Because each capability declares both its required input state and its produced output state, execution decomposes against the contract into semantic resolution (alias → entry) → argument/parameter-schema validation → prerequisite checking → dependency and state verification → state update → structured-output generation → provenance capture, so that each step’s produces satisfies the next step’s requires (a complete raw-counts → QC → normalisation/HVG → scaling → PCA → neighbours → Leiden → UMAP → annotation trace is given in Supplementary Table 4). A failure can therefore be attributed, before or at the point of execution, to one of five classes: (1) **missing state** (an unsatisfied requires contract); (2) **invalid arguments** (a parameter-schema violation); (3) **unavailable dependency** (a backend that cannot load); (4) **incompatible object structure** (nominally valid arguments but an object layout that violates a structural expectation, e.g. use_rep=’spatial’ resolving to a zero-feature array); and (5) **failed execution** (a genuine runtime or logic fault). Recovery is structured per class — substitute an interchangeable backend, reposition a data slot to satisfy a structural contract, or delegate orchestration to a validated wrapper — rather than editing the line a traceback points at. In a focused seven-task comparison spanning these classes, a baseline code-generating agent averaged 26.3 turns and 4.7 distinct errors per task and failed outright on two, whereas running the same operations through registered capabilities typed each crash and passed all seven; this comparison is illustrative (n = 7, non-random) rather than a powered reliability claim, and in one task both agents hit the same library TypeError, so the contract supplied typed context rather than an automatic fix (**Supplementary Note 2**; **Supplementary Table 5**).

### Strategy composition for autonomous workflow design

For open-ended tasks, a strategy-composition agent designs a cross-modality, multi-stage plan and never executes the pipeline itself. It (i) clarifies the objective through up-front questions on datasets/modalities/pairing, scientific question, prior object state, novelty budget, compute budget, deliverable and constraints; (ii) gates the input through a single exploratory-data-analysis cell that checks shape, annotation state, coordinate/batch keys, gene-identifier format and cross-modality pairing; (iii) surveys the ecosystem in parallel over the agent roster, the registry and the skill index; (iv) enumerates analytical primitives; (v) generates two or three candidate workflows (never one, never four or more), each with named real agent/skill/function identifiers, a novelty label, explicit risks and a four-part expected yield (metric, numerical threshold/comparison, comparison set, and statistical unit-of-inference plus test family); (vi) self-audits for circularity, executability, fabricated tool names, compute-budget fit and primitive diversity; (vii) ranks and recommends one; and (viii) emits a human-readable strategy, a machine-readable plan and a stage-by-stage delegation plan. The agent body was tuned over iterations v0→v3 against three OmicVerse-bundled datasets (snapshot, lineage and tumor-microenvironment), scored by an independent judge on a five-axis rubric (ecosystem grounding, yield specificity, novelty, risk honesty and executability; each 0–10); the selected iteration raised the mean from 31/50 to 43/50 and reduced fabricated-tool-name hallucinations from a mean of 6 per run to 0 by injecting the authoritative catalogue at runtime.

### OmicVerse-RebuildR: reference-grounded R-to-Python reconstruction

OmicVerse-RebuildR is a fixed six-stage protocol that ports R/Bioconductor packages to pure-Python py-<PKG> mirrors validated by an executable, class-aware numerical-parity gate. The R reference is treated as the executable specification — run on a fixed fixture every iteration — and is never reverse-engineered from publications. The objective is fixture-level, class-aware equivalence,

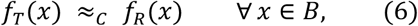

where B is the canonical fixture set and ≈_C is the parity relation for algorithm class C; this is an empirical guarantee at fixture level, not a proof over the full input domain. The stages are: (0) discovery, with a duplicate and upstream-dependency audit committed before any algorithmic code; (1) shape templating, copying only directory layout and the test rig from a class-matched prior port; (2) dual side-by-side conda environments for the Python target and the R reference; (3) a two-agent inner loop (equivalence then acceleration); (4) validation against the pre-registered gate; and (5) release to PyPI and GitHub, after which the port becomes the seed template for the next, lowering the marginal cost of each subsequent port. Coverage required at least 95% of exported (NAMESPACE) functions plus every transitive dependency. The catalogue comprises 62 packages (59 pure-Python and 3 Rust), 56 released on PyPI and 45 wired into a public ov.* entry point, spanning five source ecosystems (22 Bioconductor, 8 CRAN, 5 immunogenomics, 4 dynverse, 19 academic GitHub) and 24 methodology families; maturity is reported as five non-nested statuses (**Supplementary Tables 21, 25, 26**).

### Class-aware parity verification

Parity is matched to the class of output it checks (Supplementary Table 23). Eight algorithm classes each fix one metric and one pre-registered, read-only threshold, implemented over canonical library functions: deterministic numerical outputs are gated by element-wise tolerance (np.allclose, absolute tolerance from 1 × 10⁻¹³ to a hard ceiling of 1 × 10⁻⁶) plus Pearson agreement (e.g. py-matrixeqtl β max|Δ| < 2 × 10⁻¹⁵; py-coloc posteriors < 3 × 10⁻¹⁶); stochastic outputs by a two-sample Kolmogorov–Smirnov test (p ≥ 0.05) or a Wasserstein-1 bound; clusterings by the label-permutation-invariant adjusted Rand index (ARI),

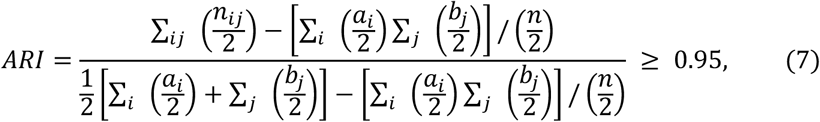

where *n_ij_* = |*U_i_* ∩ *V_j_*|[73]; embeddings by rotation–invariant Procrustes similarity,

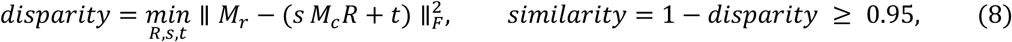

(reported per-port gates ranged down to ≥ 0.80, e.g. py-destiny eigenvector Procrustes 0.916); ranked outputs by a top-50 Jaccard (≥ 0.80) and Spearman rank correlation (≥ 0.9); ordinal/pseudotime outputs by Pearson correlation (≥ 0.99, with values within 1 × 10⁻¹² of unity treated as exact); classifications by F1 (≥ 0.95); and statistical-inference outputs by Spearman on −log₁₀ p (≥ 0.90) plus a top-50 Jaccard (≥ 0.7). The Spearman coefficient used throughout is Pearson correlation on the ranks of the two vectors. Every output of a multi-output port declares its own gate and all must pass; no per-fixture tuning or multiple-comparison relaxation is permitted, and the random seed is fixed at 42 in both reference and candidate. Because parity is class-aware, a failed deterministic-kernel test signals a genuine bug whereas a failed invariance-class test signals a scientific disagreement rather than a relabelling artefact.

### Acceleration under admissibility constraints

Once a port clears the equivalence gate, it is optimised by a self-evolving acceleration loop that cannot break equivalence. The loop is a verifier-guided, test-time search (an MDP-like closed loop) in which the policy is the in-context language model (no weight update), the action is one rewrite drawn from an algebraic/algorithmic playbook, the environment is the parity test plus a stopwatch on the canonical fixture, and the reward is

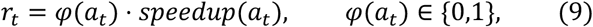

where φ is an admissibility indicator. A rewrite is merged only if it clears the parity gate and improves wall-clock by more than 1.05× (else rolled back), up to a maximum of 20 iterations. Wall-clock was measured as one discarded warm-up run plus three measured runs (mean ± s.d.; if the coefficient of variation exceeded 10%, five runs with the median) under time.perf_counter() with BLAS threads pinned to 8. Every committed rewrite carries an admissibility proof of one of three kinds: (E) an exact algebraic identity (bit-equivalent, e.g. caching *X*^T^*X*, the Woodbury identity (*I* + *λU*𝛬*U*^T^)^-1^ = *I* − *λU*(𝛬^-1^ + *λU*^T^*U*)^-1^*U*^T^, the Sherman–Morrison update or vectorisation); (B) a bounded ε-approximation with a closed-form perturbation bound, e.g. for sparse soft-assignment row truncation ∥ *W*_*new*_ − *W*_*old*_ ∥_*F*_≤ *k nK*𝜀 (10); or (C) a class-containment theorem (e.g. Euclidean minimum spanning tree ⊆ Delaunay triangulation). Across the seven packages with full telemetry, 34 acceleration iterations yielded 27 accepted and 4 rejected rewrites (2 for dropping parity, 2 for no wall-clock gain; 0 inadmissible), with exact-identity rewrites in all seven and bounded-ε rewrites in four; one rewrite that wrongly assumed k-nearest-neighbour contiguity dropped parity from 0.984 to 0.753 and was rejected at the gate. Logged speedups reached 102× (py-SpaceTrooper) within the telemetry set and extend to ≈280× (rust-NMF) and ≈192× (py-inferCNV) across the broader catalogue (**Supplementary Table 24**). Because R references were executed as a subprocess while Python ports ran in-process, on small fixtures the fixed per-call R-invocation cost is a non-trivial fraction of R wall-clock, so headline speedups blend durable compute-level gains with fixed invocation overhead, and are input- and implementation-dependent rather than a language-level claim.

### OmicOS-PortBuild: external-package skill porting

OmicOS-PortBuild extends the same discipline to third-party Python packages that are not natively part of OmicVerse, distilling each into a portable agent-and-skill bundle through a reflection–edit–validate optimisation loop^46^. A package’s tutorials, source repository, API documentation and notebooks are ingested and partitioned into capability domains (e.g. whole-slide image analysis, spatial omics, gene-expression analysis, cell segmentation and cross-modality analysis). For each partition, an agent-driven SkillOpt procedure proposes candidate edits — append, insert-after, replace or delete — applied through a protected patcher that preserves the full edit history, and scores each candidate against a ten-dimension rubric (each 1–5, normalised to 100) with an evidence-keyword check; only edits that improve the current best score are accepted, and rejected edits are retained in a buffer rather than overwritten. The optimiser was run with four epochs, a rollout batch of 40, a minibatch of 8, an edit budget of 4 and a quality pass-threshold of 75 (hard-gate minimum 3/5). Before publication, each skill must clear hard gates — frontmatter-schema validation, grounding of at least 40% of its key APIs against real source, removal of machine-specific paths and routing-fixture compatibility — after which it is packaged into a runtime-agnostic bundle of agent definitions, skill documents and assets deployable to Claude Code, Codex, OpenCode or OmicOS. As a demonstration, PortBuild converted LazySlide^47^ (≥ v0.9) into six published skills under one pathology agent, leaving each skill inspectable and portable across execution environments.

### PseudotimeFate: backend-agnostic macrostate and fate inference

PseudotimeFate (the upper-level design behind omicverse.single.Traj) treats pseudotime as a common interface: once any backend writes a pseudotime vector into adata.obs, the same four-stage workflow proceeds — a pseudotime-biased k-nearest-neighbour graph; an adaptive-Gaussian, row-normalised directed transition matrix P; macrostate inference; and fate-probability computation — whether the pseudotime came from Palantir^70^, Slingshot^43^, SCORPIUS^42^, destiny^71^, URD^72^, Monocle 3^73^ or scTour^74^. To remain lightweight, PseudotimeFate retains the geometry of Perron-cluster cluster analysis (PCCA+)^75^ but replaces the two expensive pieces of the generalised PCCA+ (GPCCA) stack^44^ with sparse, deterministic surrogates implemented in pure NumPy/SciPy. The sorted real-Schur basis is replaced by the top-K eigenvectors of *P*^T^ obtained by a sparse ARPACK Krylov solver, with complex-conjugate pairs split into real and imaginary parts and Gram–Schmidt orthonormalised; soft memberships are assigned by the inner-simplex algorithm without crispness optimisation,

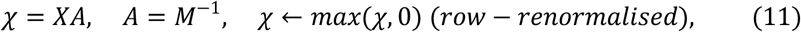

where M are the simplex-vertex rows. Hard one-hot memberships Z give the Galerkin coarse-grained transition matrix and residency,

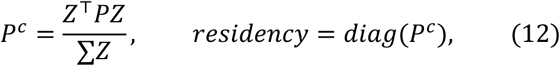

matching pyGPCCA’s coarse-grained operator; terminal macrostates are those with mean-pseudotime rank above a late-pseudotime quantile and residency above a threshold, excluding the initial state. Fate is the absorption probability into each terminal cell-set, solved by Neumann-series power iteration of

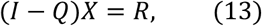

with Q the transient-to-transient block and R aggregated over each terminal set, together with a per-cell lineage entropy − ∑*_k_* 𝜒*_k_log*𝜒*_k_* (14). The eigenvector-for-Schur substitution is exact for reversible P and a bounded approximation otherwise; because the pseudotime-biased k-nearest-neighbour graph is near-reversible, the surrogate spans essentially the same metastable subspace as the true Schur basis, whereas for strongly non-reversible chains the rigorous real-Schur GPCCA remains appropriate (Supplementary Note 7). On a pancreatic endocrine differentiation dataset (3,696 cells, 3,000 HVGs) PseudotimeFate recovered four mature terminal macrostates (Beta, Delta, Epsilon, Alpha) at high self-residency, recovered canonical lineage drivers by per-gene correlation with fate probability (Gcg for Alpha, r = 0.593; Sst for Delta, r = 0.658; Pdx1/Ins2 for Beta; Ghrl for Epsilon), and under a standardised comparison with identical input, hardware and warm-up completed in 0.392 s versus 33.17 s for the matched CellRank PseudotimeKernel-plus-GPCCA workflow (84.6×) at terminal-state precision and recall of 1.0 (**Supplementary Notes 7, 8**). The speedup is environment-specific (the comparison environment lacked petsc/slepc) and the sub-second figure is the warm cost.

### Trajectory-inference benchmark and metrics

Reconstructed trajectory methods were benchmarked against gold-standard synthetic trajectories using the dynbenchmark metric set^76^ re-implemented as the pure-Python py-dyneval port. Thirteen methods (six R references and seven Python ports) were scored on 14 synthetic datasets from four simulators (Dyntoy, Splatter^77^, PROSSTT^78^ and dyngen^79^) spanning six topology classes at 300–600 cells, giving 182 method-by-dataset cells. To prevent leakage, predicted branchings were taken only from each adapter’s own output (no fallback k-means with the gold branch count), R and Python adapters read one shared on-disk fixture, and the root cell was chosen by an identical data-driven rule (never the gold root). Four families were reported, each in [0,1] with negative correlations clamped to 0. Topology used the Hamming–Ipsen–Mikhailov distance,

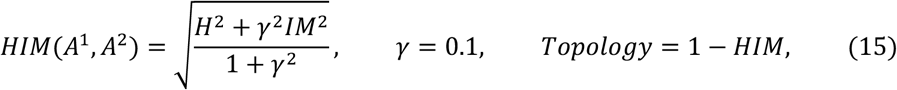

with H the normalised upper-triangular L1 difference of the milestone adjacencies and IM the L2 distance of their Laplacian spectral densities. Branch assignment used the harmonic mean of Jaccard-based recovery and relevance,

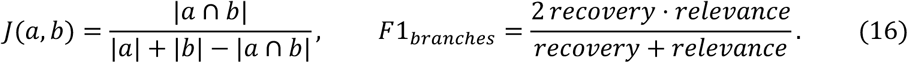

Cell positions used the Spearman correlation of the flattened gold and predicted geodesic-distance matrices over a shared waypoint set, clamped to be non-negative (17). Features used a gold-importance-weighted Pearson correlation of random-forest (50 trees, depth 8) Gini importances regressing pseudotime onto expression, with weights *w_g_* = *max*(*a_g_*, 0) (18). The reported overall score for each method was the arithmetic mean of the four families; per-family values are reported individually so that the canonical geometric-mean aggregation can be recomputed (**Supplementary Note 5**; **Supplementary Tables 27, 28**).

### BixBench-Verified-50 evaluation

BixBench-Verified-50 is the 50-question expert-verified subset of BixBench^33^ (HuggingFace phylobio/BixBench-Verified-50, spanning 33 data capsules); the per-task verification was performed by the dataset authors. Each task is a short-answer bioinformatics question answered against a per-question data capsule whose held-out expert notebook is never staged into the agent workspace. For each task, a fresh workspace was staged, the OmicOS runtime was launched against it, the question was sent over the streaming chat endpoint with the full OmicOS tool surface (Python execution, notebook and file tools, shell and registry lookup), and the final line FINAL ANSWER: … was extracted and graded by the dataset’s own verifier (exact-string, numeric-range or fixed-LLM verifier). The agent backbone was GPT-5.5 (1,000,000-token context, max_tool_iterations=100, full permissions) and one specialist or generalist agent was assigned per question by first-match routing. Pass@1 is binary per task,

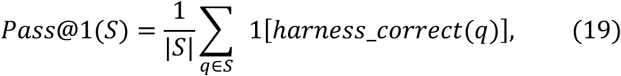

reaching 45/50 (90.0%). Four documented, reversible grader deviations from strict BixBench semantics (numeric tolerance for pure-number answers, percent↔fraction tolerance, range scanning over all emitted numbers, and a rounding-tolerant LLM verifier) make the score non-identical to the public leaderboard; external system scores were transcribed from the published leaderboard and were not re-measured. To assess process quality independently of the binary outcome, each best-per-question trajectory was scored by a fixed DeepSeek-v4-pro judge (temperature 0, anchored 0–100 calibration) on five process dimensions — data exploration, method/model selection, execution correctness, answer fidelity and iteration pattern — with 95% confidence intervals from a percentile bootstrap (10,000 resamples, seed 20260530). A 13-of-50 raw-trajectory audit grounded four of the five dimensions in every audited case and identified answer fidelity as a mild over-estimate of final-answer correctness (**Supplementary Tables 7, 8**).

### BiomniBench-DA evaluation

BiomniBench-DA^35^ (HuggingFace phylobio/BiomniBench-DA) comprises 50 process-level biomedical data-analysis tasks; unlike answer-only grading it scores the full analytical trajectory, with the agent writing a markdown narrative (trace.md) and a final answer.txt. Each task carries a 100-point rubric split across criteria with discrete levels (A full, B partial, C zero); a fixed DeepSeek-v4-pro judge selected one level per criterion, levels were mapped to points programmatically, and the per-cell score was the clamped point total divided by 100, with a cell counted correct at score ≥ 0.7:

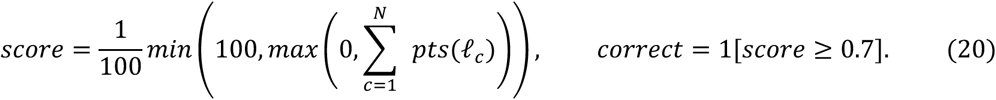

Tasks were driven through OmicOS’s vertical-specialist router (21 specialists; the generalist deliberately omitted) at max_tool_iterations=150. Every rubric criterion was classified into one of six published capability dimensions — data handling, method selection, statistical rigor, source reliability, scientific reasoning and biological interpretation — by a documented keyword-priority heuristic (auditable per criterion), and the per-dimension percentage was the point-weighted earned/maximum ratio, with the overall capability score the unweighted mean of the six. Because a BiomniBench task is a long agent loop that resends a growing context (1–2 × 10⁶ cumulative input tokens; measured cache hit ≈94–96%), two cost figures were reported per model: a naive cost and a cache-adjusted cost in which only positive per-call input increments were billed as fresh and the remainder at the provider’s cache-read price,

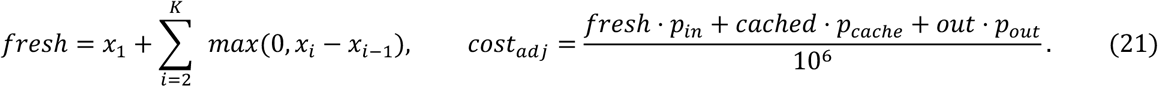

The cache-adjusted figure is therefore a best-case lower bound, and external harness costs (Claude Code, Codex CLI, Terminus-2) were transcribed from a published chart under an assumed cache-adjusted convention; we therefore read the cost axis as indicative rather than re-measured. Seven model backbones were evaluated as single fresh runs against a held-constant roster. Four tasks (da-12-4, da-18-7, da-20-1, da-6-2) were excluded as benchmark-broken — their gold answer or rubric cannot be satisfied without a methodological or factual error (e.g. da-12-4 rests on retracted tumor-microbiome data and a marginal raw-count p-value) — applying the same fixed exclusion symmetrically to every model’s denominator to give a corrected 46-task set; removing exactly these four reproduced the reported per-model scores to the decimal, and the excluded cells did not advantage the leading model (**Supplementary Note 3**; **Supplementary Tables 9–12**).

### OmicBench construction and evaluation

OmicBench is a purpose-built suite of 40 omics-analysis tasks across seven modalities (scRNA-seq 19, spatial transcriptomics 4, multi-omics 2, bulk RNA-seq 5, metabolomics 1, 16S microbiome 5 and single-cell foundation-model workflows 4), with 29 hard, 9 medium and 2 easy tasks mapping to six capability classes and 15 fine-grained families. Each prompt states only the scientific goal and never names an ov.* API. Grading is fully programmatic and package-agnostic: each task declares a set of atomic checks (151 atomic checks across the 40 tasks) that assert biological and numerical criteria and accept equivalent outputs from any library. A task score is the fraction of its checks that pass, and a task passes only when all of its checks pass,

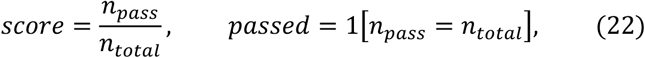

so that Pass@1 over (task, seed) cells is the stricter signal and the atomic-check pass-rate the finer one. Representative checks include clustering ARI against known labels (equation (7); e.g. ari_vs_celltype ≥ 0.30), the mean top-N Jaccard of recovered markers against an oracle,

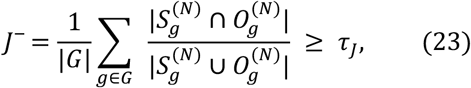

multi-method consensus (minimum pairwise Spearman of pseudotime or agreement of deconvolution fractions), scIB-style silhouette improvements, and negation-aware tool-grounding checks that scan trajectory tool outputs for canonical upstream-library markers (e.g. cellphonedb, scGPT, Geneformer, PyWGCNA) and credit a hit only when it is not within ±200 characters of an unavailability message. Checks were grouped into six native capability dimensions (object plumbing, data integrity, tool grounding, methodological robustness, quantitative quality and biological plausibility). Canonical marker panels and ligand–receptor references were hard-coded from PanglaoDB, CellMarker 2.0, CellChatDB v2 and developmental-pancreas references^80^.

### OmicVerse ablation on OmicBench

To isolate the effect of the execution layer, OmicBench was evaluated under Mini-SWE-Agent, a deliberately minimal “reason plus one shell command per turn” harness^26^ driven through LiteLLM, in which the model, agent loop, tool surface, data and environment were held identical between arms and only the availability of OmicVerse was varied (a prior-knowledge ablation; both arms could import omicverse). Across a 38-task evaluation subset, six models were run at three seeds and a locally served Qwen3.6-35B-A3B model at one seed (1,411 graded cells in total); adding OmicVerse raised Pass@1 for every model family, most for the weakest. A four-arm ablation (baseline, full OmicVerse, OmicVerse without the registry, and a documentation-retrieval-augmented-generation control over 613 OmicVerse docstrings embedded with a sentence-transformer^81^) was run for GPT-5.5 and DeepSeek-v4-flash on a common seed: the full registry-grounded arm was best on both Pass@1 and atomic-check pass-rate, the documentation control approximated baseline, and naming tools without the registry fell below baseline for GPT-5.5 — attributing the gain to the contract and registry execution layer rather than to documentation text or tool names (**Supplementary Tables 13–20**).

### Whole-body spatial-genetic Alzheimer’s disease workflow

From a single whole-body mouse spatial-transcriptomics dataset and the term “Alzheimer’s disease”, with no analysis plan or named methods, the OmicOS agent autonomously composed and executed the workflow; the human operator only chose among agent-proposed options and corrected errors. The agent recognised that the question was addressable by composing two OmicVerse modules — gsMap^48^, which projects Alzheimer’s-disease genome-wide association study (GWAS) signal onto spatial coordinates by spatial linkage-disequilibrium-score regression (spatial S-LDSC)^82^, and CMap^83^, which maps single-cell reference information onto spatial spots — and assembled the reference data they required. Five public single-cell datasets were retrieved and harmonised across age groups into a reference of 809,284 cells spanning 126 organ–age combinations (1–30 months): PanSci (GEO GSE247719, 454,620 cells)^84^, Tabula Muris Senis^85^, Mouse Cell Atlas 2.0 (GSE153562)^86^, EasySci (GSE212606)^87^ and a brain-microglia dataset (GSE179358)^88^, each projected by CMap onto a 6-week whole-body Array-seq spatial reference (GSE248904)^89^. gsMap was run per (organ × age × source) projection in quick mode with the Alzheimer’s-disease GWAS of Bellenguez et al^90^. (GWAS Catalog GCST90027158) and a mouse→human homologue mapping, producing a per-spot S-LDSC χ² statistic whose expectation, given per-annotation effects 𝜏*_c_* and LD scores ℓ(*j*, *c*), is

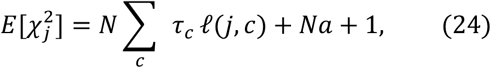

and a per-annotation enrichment from a Cauchy (ACAT) combination[85] of per-spot p-values,

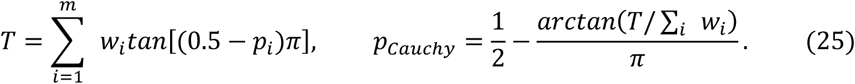

Per-spot risk was summarised as −*log*_10_*p_AD_*; age-resolved young and old whole-body maps were built from 264,283 young spots (41 samples) and 322,477 old spots (62 samples), with a higher median per-spot risk in the old map (2.12 versus 1.19), read at organ and sample level because per-spot summaries are not independent within a sample. Across most organs genetic-risk enrichment was strongest in tissue-resident myeloid annotations, with the colon the exception (epithelial > myeloid across all ages, 6.33 versus 4.42, in −log₁₀ Cauchy-combined P); a cell-composition control and an eight-trait comparison (Alzheimer’s disease, Parkinson’s disease, amyotrophic lateral sclerosis, multiple sclerosis, frontotemporal dementia, type 2 diabetes, schizophrenia and height) established that the Alzheimer’s-relevant pattern is the conjunction of broad myeloid dominance, the colon exception and the specific colon risk genes. A seven-gene endocytic-trafficking set co-localised with the colon risk score more strongly than random seven-gene sets (mean Pearson r = 0.284; z = 4.79; empirical P = 1 × 10⁻⁵ over 100,000 permutations).

Human genetic support was sought from GTEx v8[49] colon (Transverse and Sigmoid) expression quantitative trait loci (eQTLs): PICALM had 93 colon eQTL variants and no detectable bulk-brain eQTL, CD2AP had 200 (including an Alzheimer’s-disease lead SNP that was itself a colon eQTL) and CR1 had 24 overlapping an Alzheimer’s-disease SNP. Single-causal colocalisation (coloc-ABF)[50] on full summary statistics computed the posterior over five hypotheses from per-SNP Wakefield approximate Bayes factors[86],

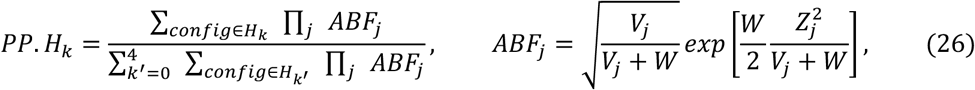

supporting CR1 as a shared-causal colon signal (PP.H4 = 0.993, colon sigmoid) but placing PICALM on a distinct signal (PP.H3 = 0.999); restricting to significant-only eQTLs inflated PP.H4 for PICALM and ADAM10, which full-summary colocalisation corrected. Multi-causal SharePro^91^ recovered a shared PICALM signal (0.998) and supported CR1 (0.999) but not ADAM10 (0.0005), and summary-data-based Mendelian randomisation with the HEIDI test^92^ flagged ADAM10 as a linkage artefact; no gene passed simultaneous coloc-ABF, SMR and HEIDI support. Localisation was corroborated in human colon Visium data (3,887 spots), where PICALM and CR1 co-localised with the epithelial marker EPCAM and the myeloid marker CD163. All GTEx colon eQTLs are from bulk tissue and the analysis is hypothesis-generating; neither the mouse maps nor the human bulk eQTLs establish the colon epithelium as a causal tissue (**Supplementary Tables 33–36**; **Extended Data Figs. 4, 5**).

### Histopathology analysis with LazySlide and PortBuild

Using the PortBuild-derived LazySlide^47^ skills, OmicOS composed a haematoxylin-and-eosin (H&E) pathology workflow in three tracks. The H&E analysis used the public 10x Genomics Visium HD Human Lung Cancer Fixed Frozen H&E image and matched Visium HD spatial-transcriptomics data. The 10x Visium HD H&E image was represented by 19,659 retained tissue tile centres. Around each centre, Prov-GigaPath morphology features were extracted from image patches at physical footprints of 16, 32 and 64 µm, corresponding to 58, 117 and 234 pixels at 0.274 µm per pixel. The displayed morphology-domain analysis used the 32-µm feature table. Morphology domains were identified from Prov-GigaPath^93^ features extracted from H&E tiles and the resulting smoothed labels comprised 12 domains. Tiles with H&E tumor-enrichment scores ≥ 0.50 were classified as tumor-enriched tiles.

An external benchmark used the publicly available IGNITE NSCLC H&E tissue-compartment segmentation data^94^. The annotated lung adenocarcinoma ROI patient22_he_roi1 was cropped to the pathologist-annotated area and divided into a nominal 128-pixel tile grid, yielding 72 valid tiles. For each tile, the ground-truth tumor fraction was calculated from the pathologist annotation mask, and tiles with at least 10% tumor-annotated pixels were labelled tumor-positive. OmicOS assigned each tile an H&E epithelial-tumor proxy score from tile-level H&E image features, and tiles with scores ≥ 0.50 were classified as tumor-positive (Supplementary Note 9). Precision, recall and F1 score were calculated from the tile-level labels, and AUROC was calculated from the continuous OmicOS-calculated epithelial-tumor proxy score.

For the H&E-to-spatial-transcriptomics analysis, predicted expression profiles for 38,982 genes were generated for 19,659 H&E tiles at a 32-µm spatial scale using the OmicVerse histology workflow and STPath zero-shot generative foundation model^95^. Predicted spatial-transcriptomics programme scores were calculated from this predicted expression matrix using predefined marker-gene sets. For each marker gene, predicted expression values across tiles were robustly normalized to the 1st–99th percentile range and clipped to [0,1]. For each tile, a programme score was calculated as the mean normalized expression of the available marker genes in that programme. The marker sets were: epithelial tumor (EPCAM, KRT19, KRT8, KRT18, MUC1 and SOX2), stromal ECM (COL1A1, COL1A2, DCN, LUM and VIM), immune (PTPRC, CD3D, CD3E, CD68, LST1 and MS4A1), vascular (PECAM1, VWF, KDR and EMCN), proliferation (MKI67, TOP2A and PCNA), and hypoxia/stress (VEGFA, CA9 and HIF1A). Predicted ST domains were then derived by clustering the six tile-level programme scores, with weighted spatial coordinates included during clustering, followed by neighbourhood smoothing, yielding eight inferred domains (D0-D7). The tile-level programme scores were mapped to H&E tile coordinates to generate spatial programme maps.

The multi-section LUAD analysis used four TCGA-LUAD diagnostic H&E sections, comprising two high-risk sections (TCGA-55-8620 and TCGA-55-A490) and two low-risk sections (TCGA-62-A46S and TCGA-78-7537). Tissue regions were segmented in downsampled slide thumbnails with the longer image dimension set to 2,048 pixels and tiled on a 32-pixel grid at thumbnail resolution. Tiles with at least 25% tissue were retained, yielding 9,298 H&E thumbnail tiles. For each tile, thumbnail-level H&E colour and texture features were extracted and converted by robust normalization and fixed linear weighting into descriptive tumor, immune, stromal and necrosis/hypoxia morphology proxy scores for morphology interpretation (Supplementary Note 9). For the risk-region analysis, morphology programmes were computed from z-scored H&E morphology proxy features and compared separately for high-risk- and low-risk-enriched regions against all other retained tiles using Cohen’s d,

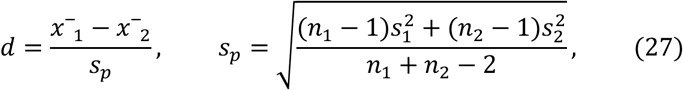

Here, *x*^−^_1_ and *x*^−^_2_ denote the mean programme scores in the target risk-enriched region and in all other retained tiles, respectively; *n*_1_ and *n*_2_ denote the corresponding numbers of tiles; and *n*^2^_1_ and *n*^2^_2_ denote the corresponding sample variances used to calculate the pooled standard deviation.

We used the TCGA-LUAD cohort to evaluate the association between an H&E-derived stromal-immune score and overall survival. Diagnostic H&E sections with matched overall-survival and clinical metadata were processed to compute patient-level H&E stromal-immune scores (Supplementary Note 9), resulting in 240 TCGA-LUAD patients. Patients were dichotomized at the cohort median H&E stromal-immune score and assigned to H&E stromal-immune high group and H&E stromal-immune low group. Overall survival was estimated using the Kaplan–Meier estimator,

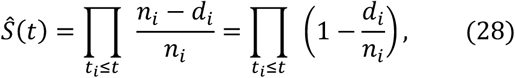

where d_i_ is the number of events at time t_i_ and n_i_ is the number of patients at risk immediately before t_i_. Survival curves were compared using a two-sided log-rank test. The displayed hazard ratio was estimated using a penalized univariable Cox proportional-hazards model with the median-defined H&E stromal-immune high-group indicator as the covariate.

### Statistics and reproducibility

No statistical method was used to predetermine sample size; the benchmark task counts are fixed by the source suites (BixBench-Verified-50, 50 tasks; BiomniBench-DA, 50 tasks, corrected to 46; OmicBench, 40 tasks) and the reconstruction and trajectory analyses are deterministic given a fixed seed. Random seeds were pinned throughout: 42 for reconstruction parity and acceleration, 0 (and 0–2 for multi-seed benchmark runs) for stochastic preprocessing and evaluation, and 20260530 for bootstrap resampling. Benchmark-broken BiomniBench-DA tasks (da-12-4, da-18-7, da-20-1, da-6-2) were excluded by a single fixed rule applied symmetrically to every model before any score was computed, with the rationale documented per task (Supplementary Note 3); no other data were excluded. Bootstrap confidence intervals used the percentile method with 10,000 resamples. Correlations are Pearson unless Spearman is stated; permutation P values are empirical over the stated number of permutations; colocalisation, SMR/HEIDI and survival statistics are reported with the exact posteriors, effect sizes and P values in the corresponding figures and tables. The experiments were not randomised and the investigators were not blinded, as all grading was either programmatic and deterministic or performed by a fixed automated judge; the with- and without-OmicVerse ablation held the model, agent loop, tools, data and environment identical between arms, varying only the execution layer. Wall-clock comparisons used one discarded warm-up plus three measured replicates (median of five when the coefficient of variation exceeded 10%) with BLAS threads pinned. Cross-harness cost comparisons that include rows transcribed from published charts are treated as indicative rather than re-measured under a single tool stack.

## Data availability

The benchmark task suites are available as the gated HuggingFace datasets phylobio/BixBench-Verified-50(https://huggingface.co/datasets/phylobio/BixBench-Verified-50) and phylobio/BiomniBench-DA(https://huggingface.co/datasets/phylobio/BiomniBench-DA); OmicBench tasks, rubrics and fixtures are available at omicverse/OmicBench (https://huggingface.co/datasets/omicverse/OmicBench). The out-of-memory benchmark used Tabula Sapiens data from CZ CELLxGENE Discover (collection e5f58829-1a66-40b5-a624-9046778e74f5). The Alzheimer’s-disease workflow used public datasets under GEO/GWAS-Catalog accessions PanSci^84^ GSE247719, Mouse Cell Atlas 2.0^86^ GSE153562, EasySci^87^ GSE212606, brain microglia^88^ GSE179358, whole-body Array-seq^89^ GSE248904, Tabula Muris Senis^84^ (figshare), the Bellenguez et al. Alzheimer’s-disease GWAS GCST90027158^90^, and GTEx v8 colon eQTLs; additional colon validation datasets are listed in **Supplementary Table 33**. Pancreatic-endocrine and other demonstration datasets are distributed with OmicVerse. The H&E image and matched spatial-transcriptomics data used for the H&E-to-spatial-omics analyses were obtained from the public 10x Genomics Visium HD Spatial Gene Expression Library, Human Lung Cancer (Fixed Frozen) dataset. TCGA-LUAD diagnostic H&E whole-slide images and clinical metadata used for the multi-section LUAD and survival analyses were obtained from the NCI Genomic Data Commons Data Portal/API under project ‘TCGA-LUAD’. The data supporting the external pathologist-annotation benchmark in Extended Data Fig. 6 are publicly available in the IGNITE NSCLC data toolkit on Zenodo(https://doi.org/10.5281/zenodo.15674785). Source data are provided with this paper.

## Code availability

The source code for OmicVerse is available at https://github.com/omicverse/omicverse. The out-of-memory backends AnnDataOOM and MuDataOOM are available at https://github.com/omicverse/anndata-oom and https://github.com/omicverse/mudata-oom, respectively. OmicVerse-RebuildR is available at https://github.com/omicverse/omicverse-rebuildr, and OmicOS-PortBuild is available at https://github.com/omicverse/omicos-portbuild. Reconstructed py-* and rust-* packages are distributed through the OmicVerse GitHub organization (https://github.com/omicverse/) and, where applicable, through PyPI. The OmicOS runtime, including omicos-core, is available from the corresponding Primordecode page (https://omicos.cn/). The benchmark reproduction trajectories and evaluation records for OmicBench, BixBench and BiomniBench are available at https://github.com/omicverse/OmicVerse-OmicBench, https://github.com/omicverse/OmicOS-BixBench and https://github.com/omicverse/OmicOS-BiomniBench, respectively. The H&E-to-spatial-omics reproduction is available at https://github.com/omicverse/OmicOS-HE_Reproducibility. Per-package reconstruction reports, parity manifests and acceleration-iteration logs are provided with the documented reconstructed ports. Repository-specific licences, installation instructions and version information are provided in the corresponding repositories.

## Acknowledgements

We first thank all members of the 112 Laboratory at the University of Science and Technology Beijing for their support of this project. We are also grateful to all members of the Qiu Lab at Stanford University for their valuable suggestions and discussions during the development of this work. We thank Jianming Zeng, founder of Bioinformatics Skill Tree, for his early promotion of the OmicVerse ecosystem. We also thank all OmicVerse users who have submitted issues, pull requests and feedback, which helped improve the ecosystem over time. We also thank Xiaotong Wu from the Xie Lab at Tsinghua University for helpful guidance on epigenomics.

## Author Contributions

Z.Z. conceived, founded and led the OmicVerse open-source community, and was responsible for the overall construction, integration and maintenance of the OmicOS ecosystem. X. Meng served as an active maintainer of OmicVerse and contributed to the development of pseudotime and cell–cell communication modules. L.H. developed algorithms for the spatial multi-omics component, contributed to the validation of the OmicOS external-module PortBuild framework and performed H&E-related experiments. C.L. contributed to the self-evolving components of the OmicVerse ecosystem and helped develop the agentic concepts underlying OmicOS. P.L. and Y.S. contributed to iterative testing of the OmicOS ecosystem. X. Ma performed practical testing of OmicOS and contributed to external pathology-slide validation experiments. L.G. contributed to iterative testing of OmicOS and developed Skill and Agent templates. X.W. developed the OmicVerse Agent components. Z. Luo developed and maintained bulk-transcriptomics algorithms in OmicVerse. Y.Z. contributed to the development and testing of bulk-transcriptomics algorithms in OmicVerse. J.X. developed and maintained copy-number-variation algorithms in OmicVerse. Z. Lin contributed to the reconstruction of spatial-transcriptomics algorithms and the development of single-cell metabolomics algorithms in OmicVerse. H.Z. designed logos for the reconstructed OmicVerse algorithm packages. Z.J. contributed to the reconstruction and development of metagenomics algorithms. S.M. contributed to the reconstruction and development of Hi-C algorithms. Y.L. advised on the design of spatial-transcriptomics algorithms in OmicVerse and contributed to OmicOS testing. W.T. advised on the design of single-cell foundation-model algorithms in OmicVerse and contributed to OmicOS testing. Q.P. advised on the design of transcriptional-dynamics algorithms in OmicVerse and contributed to OmicOS testing. Y.M. contributed to OmicOS testing. L.Z. performed validation testing of OmicOS in hydrogel-related applications and helped organize the analytical logic. C.X. performed validation testing of OmicOS in Alzheimer’s disease-related applications and helped organize the analytical logic. X.Z. contributed to the design of the OmicOS ecosystem and revised the manuscript. Y.X. contributed to the conceptualization of the OmicVerse ecosystem, supervised experimental validation of OmicOS and revised the manuscript. H.D. contributed to the conceptualization of the OmicVerse ecosystem and supervised the application of OmicOS in biomedical research. Z.Z. wrote the manuscript with input and revisions from all authors.

