## Supplementary Notes 1-8 and Tables 1-36 for "OmicOS: A Comprehensive Omics Ecosystem Infrastructure and Agent System for the AI Era"

—-------------------------------------------------

### OmicOS: A Comprehensive Omics Ecosystem Infrastructure and Agent System for the AI Era

Zehua Zeng[1,2,3,4,21†,✉], Xu Meng[5†], Lei Hu[6,22†], Chen Li[3,7†], Peng Liu[7], Yu Shi[8], Xuejiao Ma[9], Lianchong Gao[10,23], Xuehai Wang[11], Zhi Luo[1,2], Yawen Zheng[1,2], Jieshen Xian[12], Ziheng Lin[13], Haoliang Zhu[14], Zhaorui Jiang[15], Sheng Mao[6], Yifan Lu[3,4,16], Wenzhuo Tang[17], Qiangwei Peng[18], Yuqing Ma[19,20], Liping Zhou[1], Cencan Xing[1,2], Xuegong Zhang[7,✉], Yuanyan Xiong[4,✉] , Hongwu Du[1,2,✉]

*[1]School of Chemistry and Biological Engineering, University of Science and Technology Beijing, Beijing 100083, China*
*[2]Daxing Research Institute, University of Science and Technology Beijing, Beijing 100083, China*
*[3]Department of Genetics, Stanford University, Stanford, CA 94305, USA*
*[4]Department of Biochemistry, Key Laboratory of Gene Engineering of the Ministry of Education, School of Life Sciences, Sun Yat-sen University, Guangzhou 510275, China*
*[5]College of Informatics, Hubei Key Laboratory of Agricultural Bioinformatics, Huazhong Agricultural University, Wuhan 430070, China*
*[6]School of Life Sciences, Westlake University, Hangzhou 310030, China*
*[7]MOE Key Laboratory of Bioinformatics, Bioinformatics Division of BNRIST (Beijing National Research Center for Information Science and Technology), Department of Automation, Tsinghua University, Beijing 100084, China*
*[8]School of Life Sciences, Tsinghua University, Beijing 100084, China*
*[9]Department of Surgery of Traditional Chinese Medicine, China-Japan Friendship Hospital, Beijing 100029, China*
*[10]Key Laboratory of Systems Biomedicine (Ministry of Education), Shanghai Center for Systems Biomedicine, Shanghai Jiao Tong University, Shanghai 200240, China*
*[11]Department of Physiology and Pharmacology, Karolinska Institutet, Stockholm SE-171 77, Sweden*
*[12]School of Public Health, Guangzhou Medical University, Guangzhou 511436, China*
*[13]School of Basic Medical Sciences, Capital Medical University, Beijing 100069, China*
*[14]School of the First Clinical Medical Sciences, Wenzhou Medical University, Wenzhou 325000, China*
*[15]School of Environment and Energy, Peking University Shenzhen Graduate School, Shenzhen 518055, China*
*[16]Electronic Information School, Wuhan University, Wuhan 430072, China*
*[17]Department of Statistics and Probability, Michigan State University, East Lansing, MI 48824, USA*
*[18]School of Mathematical Sciences, Peking University, Beijing 100871, China*
*[19]Center for Precision Medicine and Healthcare, Tsinghua-Berkeley Shenzhen Institute, Shenzhen 518055, China*
*[20]Institute of Biopharmaceutical and Health Engineering, Tsinghua Shenzhen International Graduate School, Shenzhen 518055, China*
*[21]Basic Science and Engineering Initiative, Betty Irene Moore Children's Heart Center, Lucile Packard Children's Hospital Stanford, Palo Alto, CA 94304, USA*
*[22]Department of Artificial Intelligence, School of Engineering, Westlake University, Hangzhou 310030, China*
*[23]Nurturing Center of Jiangsu Province for State Laboratory of AI Imaging and Interventional Radiology, Department of Radiology, Zhongda Hospital, School of Medicine, Southeast University, Nanjing 210009, China*

*[†]These authors contributed equally to this work.*

### Supplementary Notes

#### *Supplementary Note 1 | A function registry that lifts raw omics callables into machine-readable capability objects*

##### 1. A worked example of the problem

Consider a real OmicVerse function for manual cell-type annotation, whose Python signature is roughly

```
scanpy_cellanno_from_dict(adata, anno_dict, clustertype='leiden', ...)
```

To an autonomous LLM agent that sees only this callable, three facts that determine whether the call succeeds are invisible:

1. **It cannot be run first.** The function maps *cluster IDs* to cell types, so the data must already have been clustered — concretely, `leiden` must have run and written a `leiden` column. Nothing in the signature says this. An agent that calls it on a freshly loaded matrix gets a meaningless or failing result. 2. **The agent may not find it.** A user request such as “给我的细胞手动注释” (“manually annotate my cells”) shares no tokens with the identifier `scanpy_cellanno_from_dict`; signature or keyword search misses it. 3. **The agent cannot chain it.** It writes a new `obs` column of cell-type labels — the input that a downstream plot or differential-expression step needs — but the signature does not declare that this is what it produces.

This is the general failure mode behind S1: a bare callable, even with a docstring, does not tell an agent what the function is *for* in biological terms, what data state it expects (raw counts? normalized expression? a neighbour graph? batch labels? a prior clustering?), what must run before it, or what it leaves behind. Two functions with identical signatures can differ completely in all of these.

OmicOS solves this by lifting every raw callable into a **capability object** — a structured, machine-readable contract carried alongside the function. For the annotation function above, the contract is literally:

```
aliases       = ["字典注释", "scanpy_cellanno_from_dict", "manual_annotation",
                 "手动注释", "字典映射注释"]
category      = "single"
description   = "Manual cell type annotation from cluster-to-celltype dictionary mapping"
prerequisites = {"functions": ["leiden"]}          # HARD: leiden must run first
requires      = {"obs": []}                         # reads a user-named cluster column
produces      = {"obs": []}                         # writes a new {anno}_celltype column
auto_fix      = "none"
examples      = ["cluster2annotation = {'0':'T cell','1':'B cell', ...}",
                 "ov.single.scanpy_cellanno_from_dict(adata, anno_dict=...,"
                 "                                     clustertype='leiden')"]
related       = ["single.pySCSA", "single.get_celltype_marker", "utils.embedding"]
```

Now the same three facts are explicit and machine-readable: the Chinese alias 手动注释 resolves the user’s request to this function; `prerequisites={"functions":["leiden"]}` tells the agent to cluster first; and `produces` declares the new annotation column so a downstream step can consume it.

##### 2. What the contract encodes (S2), through real entries

Every registered function becomes one canonical *entry* (a dictionary) indexed under its full name, short name and **every** alias, so any of them resolves to the same object in constant time. The contract has nine fields; we illustrate each with two further real entries — `ov.pp.qc` (quality control) and `ov.pp.identify_robust_genes` (robust-gene/HVG initialization):

```
# ov.pp.qc
aliases     = ["质控", "qc", "quality_control", "质量控制"]
category    = "preprocessing"
description = "Perform comprehensive quality control on single-cell data ..."
requires    = {}                                   # accepts a raw AnnData
produces    = {"obs": ["n_genes","n_counts","pct_counts_mt"], "var": ["mt","n_cells"]}

# ov.pp.identify_robust_genes
aliases       = ["稳健基因","robust_genes","identify_robust_genes","基因过滤","expressed_genes"]
category      = "preprocessing"
prerequisites = {"optional_functions": ["qc","filter_cells","filter_genes"]}   # SOFT
produces      = {"var": ["n_cells","percent_cells","robust","highly_variable_features"]}
```

| # | Field | What it encodes | In the examples above |
|---|---|---|---|
| 1 | **aliases** | every name a user might use — abbreviations, full phrases, **multilingual** | `qc` is also 质控/质量控制; matched by exact, fuzzy (cutoff 0.6) and token overlap |
| 2 | **category + description** | biological intent: a browsable bucket + a one-line purpose | `category="preprocessing"`, "comprehensive quality control …" |
| 3 | **parameter schema** | the signature + each parameter with its default, captured by introspection | e.g. `percent_cells=0.05` |
| 4 | **requires** | accepted inputs / required data state, as `key → constraints` | `qc.requires={}` → takes a raw object; keys come from an allowlist (`shape`, `attributes`, `layers`, `obsm`, `obs`, `var`, …) |
| 5 | **prerequisites** | required object states / which functions must run first | annotation needs `{"functions":["leiden"]}` (hard); robust-genes uses `{"optional_functions":[...]}` (soft) |
| 6 | **dependency constraints** | the same two-tier `prerequisites` — hard vs optional — lets the agent order steps | hard = must precede; optional = may precede |
| 7 | **produces** | side effects: the new state/attributes/metrics/files created | qc writes `obs:[n_genes,n_counts,pct_counts_mt]`, `var:[mt,n_cells]` |
| 8 | **return structure** | declared inside `produces` (`fields/attributes/shape`) for chaining | robust-genes adds `var:[robust, highly_variable_features]` consumed by HVG selection |
| 9 | **auto_fix** | error policy: none / auto / `escalate` | `"none"` = the agent decides on failure |

Each entry also stores the callable itself, the dotted `full_name`, the `module`, the extracted docstring, the `examples`, the `related` cross-references, and a `source` tag (how it was discovered). The contract-key allowlist is domain-agnostic by default and can be overridden per library, so the same machinery works outside omics.

A companion **agent-friendliness score (AFS)** audits how completely an entry is filled in (a metadata-quality measure, not a correctness measure). It is a weighted sum of four 0–1 sub-scores: **Discoverability** D (0.30; ≥3 aliases, a multilingual alias, a category, a ≥40-character description), **Contract clarity** C (0.35; a non-empty `requires`/`produces`, a declared `auto_fix`, and declared prerequisites), **Signature clarity** S (0.15; defaulted keyword args + a dispatch hint), and **Behavioural** B (0.20; ≥1 example + ≥1 related). `ov.pp.qc`, for instance, scores full marks on D and B (rich multilingual aliases, worked examples) and partial on C (it declares `produces` but no prerequisites).

#### 3. How capabilities are discovered and ranked (S3)

Three mechanisms turn raw callables into ranked, executable capabilities.

**Runtime registration** is the canonical path: the `@register_function` decorator fires at import, validates the metadata (a non-empty alias/category/description are mandatory; `auto_fix` must be legal; `prerequisites` keys must be `functions/optional_functions`; `requires/produces` keys must be in the allowlist), introspects the live signature and docstring, and indexes the entry under its name and all aliases. It also auto-derives **virtual entries** by reading the function's syntax tree: a public method on a registered class becomes its own entry, and an `if method=="harmony" / elif method=="scvi"` dispatch becomes one synthetic entry per branch (e.g. `batch_correction[method=harmony]`), so an agent can target a specific algorithm rather than a monolithic function.

**Static source-code scanning** covers code that has not been imported (and foreign libraries with no decorator). A scanner walks the source tree, parses each file's syntax tree, and recovers the decorator's arguments by literal evaluation — without import-time side effects — synthesizing an entry (public name, module, a signature from argument names, docstring) and the same method/branch expansion. Results are cached on disk and invalidated by file modification time.

**Metadata hydration** merges the two: when a function appears in both, the runtime entry (which holds the real callable and resolved docstring) wins; internal names are normalized to public ones; class–method relations are resolved; missing scalar fields are back-filled from whichever source has them; and list fields (aliases, examples, related) are deduplicated.

**Ranking for a task.** A free-text task is scored against every candidate by a seven-part key — exact-name match (4.0 full / 3.0 short or alias), alias overlap, name overlap, a structural "kind" preference (parent function > method > branch), a weak score over description/docstring/examples, a contract-text overlap, and a source bias (runtime > static). Non-matches are dropped; the rest are sorted and the top entries are rendered into a compact agent block: name, signature, description, full docstring, hard prerequisites, the `requires/produces` maps, and an example. Crucially, prerequisites and contracts are *surfaced for planning*, not used to silently filter — the agent decides ordering from them.

#### 4. Capability-discovery statistics

Auditing the public OmicVerse API over all registered functions:

| Metric | Value |
|---|---|
| Raw callables promoted to capability objects (contracts) | **231** |
| Capability categories | 9 (preprocessing, single, space, bulk, plotting, pl, utils, trajectory, biocontext) |
| Aliases per function | mean **4.3**, median 5 (range 2–12) |
| Carry discovery metadata (aliases + category + description) | **100 %** |
| Carry ≥1 worked example + related links | **100 %** |
| Carry an explicit input/output/dependency contract (`requires/produces/prerequisites/auto_fix`) | **45.9 %** |
| Carry most contract fields | **35.1 %** |
| Carry signature dispatch hints | **29.0 %** |

| | |
|---|---|
| Library-wide AFS | mean **0.61**, median 0.58 (p10 0.50, p90 0.81) |

The pattern is consistent with the worked examples: **discovery and grounding metadata are universal** — all 231 functions are findable by intent (a mean of 4.3 names each, including Chinese aliases like 质控 and 手动注释) and carry runnable examples — which is exactly what lets a weaker model locate and call the right tool. The heavier machine-checkable **execution contract** (explicit `requires/produces/prerequisites`, as in the `scanpy_cellanno_from_dict → leiden` dependency) is present on a substantial minority (~36–46 %) and concentrates in the preprocessing and single-cell layers — an actively expanding annotation frontier rather than a finished layer.

#### Caveats

- AFS measures **metadata completeness, not correctness**: a function declaring `produces={"obs":["pct_counts_mt"]}` is not verified at runtime to actually write that column.
- Statistics are over the **231 registered public functions** (those promoted to capability objects); the count of raw callables the scanner visits *before* promotion is not separately recorded, so we report promoted = 231.
- `requires/prerequisites/produces` are surfaced to the agent for planning but do **not** gate ranking or execution at search time; satisfying them is left to the agent's plan.
- Some contracts use intentionally empty value lists (e.g. `requires={"obs":[]}` for the annotation function, where the exact cluster column is user-named at call time); these declare *which slot* is read/written without pinning the specific key.
- Branch-variant discovery parses simple `if/elif` string dispatch; nested or complex conditionals may not be captured.
- Static-scan timings (~4–5 s cold scan of ~1,000 files) are approximate, not benchmarked.

### *Supplementary Note 2 | A localizable failure taxonomy (baseline agents vs OmicOS)*

#### Motivation

A code-generating agent that drives an omics analysis by emitting free-form Python scripts has exactly one failure signal: the interpreter raises, the cell crashes, and a traceback comes back. Every distinct underlying problem — a missing precomputed graph, a mistyped keyword, a broken native library, a malformed `AnnData` slot, a transient logic bug — arrives in the same shape, an opaque crash at the bottom of a stack trace. The agent cannot tell *which kind* of thing went wrong without re-reasoning over the whole script, so it tends to thrash: it edits the line the traceback points at, re-runs, hits the next manifestation, and repeats. In our comparison set the baseline averaged 26.3 turns and 4.7 distinct errors per task, with two of seven tasks (B10, C04) never recovering.

OmicOS executes against a registry of *capabilities*, each of which declares (i) its required input state, (ii) a parameter schema, and (iii) the state it produces. Because execution is decomposed against these declared contracts, a failure can be attributed to a specific class *before or at the point of execution* rather than discovered as a generic crash. We organize the attributable failures into five classes and ground each in observed trajectories.

#### The five classes

**1. Missing state.** The capability's *required-input* contract is unsatisfied — a precomputed object the step depends on is absent. In **A05_leiden_clustering**, `sc.tl.leiden` requires a neighbor graph in adata.obsp['connectivities']/['distances'] and adata.uns['neighbors']. Here neighbors *were* precomputed, so neither agent tripped this class — but had they not been, the baseline would have surfaced a bare KeyError: 'connectivities' indistinguishable from a typo, whereas a requires: neighbors contract element makes the same failure attributable up front.

**2. Invalid arguments.** The call reaches a real function with arguments that violate its parameter schema. In **B05_multi_method_annotation**, the baseline passed mode='best_match' to celltypist.annotate, which expects the literal 'best match' (space, not underscore), yielding ValueError: Unrecognized 'mode' value. In **F01_multi_mode_velocity**, the baseline passed n_top_genes=2000 to scVelo's filter_and_normalize, which forwards to normalize_per_cell() and raised TypeError: normalize_per_cell() got an unexpected keyword argument 'n_top_genes'. Both are pure parameter-schema violations: an enum constraint on mode, or a documented "n_top_genes not supported; subset HVGs separately," would localize them before runtime.

**3. Unavailable dependency.** The required software backend cannot be loaded at all. In **L01_foundation_model_cell_embedding**, importing scgpt.tokenizer.gene_tokenizer pulled in torchtext, which failed with OSError: libtorchtext.so: undefined symbol: _ZN5torch3jit17parseSchemaOrName... — a torch/torchtext ABI mismatch. This is not a logic error and no amount of script editing fixes it. OmicOS's ov.fm.run declares multiple interchangeable backends (scgpt, geneformer, cellplm), so the unavailable-dependency class is attributable and the system programmatically selects a working backend instead of trial-and-error.

**4. Incompatible object structure.** Arguments are nominally valid but the object's internal layout violates the consuming step's structural expectations. In **C04_spatial_trajectory_stt**, the baseline called sc.pp.neighbors(..., use_rep='spatial'), but scanpy's use_rep lookup contract expects an X_*-prefixed obsm slot or a layers key, not a bare 'spatial'. The lookup resolved to an empty array and surfaced deep inside sklearn as ValueError: Found array with 0 feature(s) (shape=(3460, 0)) — a structural mismatch wearing the costume of a numeric error. OmicOS resolved it by copying coordinates into obsm['X_spatial'] to satisfy the schema contract.

**5. Failed execution.** A genuine runtime/logic failure inside otherwise-valid, available, well-typed code. In **B10_gene_regulatory_network**, the baseline's hand-rolled SCENIC/arboreto pipeline raised TypeError: Must supply at least one delayed object from inside dask_expr from_delayed() — an empty Dask graph produced by the orchestration, not by a bad argument. After 50 turns and 12 errors it never recovered; OmicOS used the registered ov.single.SCENIC wrapper, which configures Dask and partitioning internally. The f-string SyntaxErrors in **E04_bulk_wgcna** and **A05** are milder members of this class — incidental logic faults that a wrapper with declared produces outputs sidesteps.

#### Why the contract makes failures localizable

The recurring pattern is that the *same crash* belongs to a definite class once the capability's contract exists. C04's 0 feature(s) error is sklearn's, but the cause is a use_rep structural contract violation; F01's TypeError is scVelo's, but the cause is a parameter-schema violation. Without contracts the agent only sees the downstream library's error and edits blindly. With required-state / parameter-schema / produced-state declarations, OmicOS attributes the fault to a class and recovers with structured context — substitute a backend (L01), reposition a slot (C04), delegate orchestration to a validated wrapper (B10, E04) — rather than thrashing. Across the set, every OmicOS trajectory passed; the two outright baseline failures (B10, C04) were precisely the cases whose root cause was buried deepest below the surface traceback.

#### *Supplementary Note 3 | Rationale for excluding the four benchmark-broken BiomniBench-DA tasks*

Supplementary Note 2 documents a benchmark-calibration decision underlying the six-dimensional capability radar in Fig. 2d (data handling, method selection, statistical rigor, biological interpretation, scientific reasoning, source reliability). The radar is computed over a corrected 46-task capability set: four of the 50 BiomniBench-DA tasks (da-12-4, da-18-7, da-20-1, da-6-2) were excluded because they are benchmark-broken — items whose gold answer or grading rubric is invalid on primary-source inspection of the task instruction, rubric, grade record, and raw agent trajectory, independent of any model's behavior. In every case the agent performed standard, defensible, instruction-faithful science, but the rubric awards full credit only for a specific number, statistical test, or selection filter that is either (i) absent from or contradicted by the task instruction, (ii) statistically wrong for the question posed, (iii) scientifically incorrect or internally inconsistent, or (iv) derived from retracted/contaminated source data. The general principle for exclusion is narrow: we remove a cell only when the benchmark's own gold/rubric cannot be satisfied without committing a methodological or factual error, not when an agent simply scored poorly. The published 46-task capability set (Fig. 2d) excludes exactly these four benchmark-broken items; this four-task exclusion reproduces the reported per-model scores to the decimal. Two other tasks (da-19-1, da-8-3) carry their own failure-case write-ups for unrelated benchmark issues but are RETAINED in the 46-task capability set and are outside the scope of this note.

##### da-12-4 — Gold answer rests on retracted source data and a marginal raw-count p-value; the agent's correct conclusion is penalized for a p-value that differs from the broken benchmark value.

- **Original question:** From lung cancer patients with matched tumor-sample microbiome and RNA-seq data, is Kocuria significantly associated with poor prognosis, defined as HR > 1 and p-value < 0.05 in univariate Cox proportional hazards analysis (TCGA-LUAD)?
- **Failure mode:** The agent built the correct matched cohort (457 patients, 173 events), ran per-taxon univariate Cox models on raw aggregated counts, applied the instructed HR > 1 AND p < 0.05 rule, and reported Kocuria with HR = 1.0114, 95% CI 1.0004-1.0226, p = 0.0428, concluding 'Yes, significantly associated with poor prognosis' — exactly the gold conclusion. Criterion 4 was nonetheless docked from A (27/27) to B (13/27) solely because the reported p = 0.0428 is ~1.8x larger than the rubric's expected p ~= 0.0234, dropping the cell to 0.86 even though the grader notes 'conclusion remains correct and well-supported.' The penalty is a numeric mismatch against a fragile benchmark value, not a reasoning error.
- **Why the benchmark item is invalid:** The benchmark item is invalid on three converging grounds. (1) The rubric's expected value is exact and brittle: Criterion 4 level A requires 'HR ~= 1.0124 and p ~= 0.0234 (within tolerance)', a value obtainable only under one untransformed raw-count encoding; log-relative, log-raw, and binary-presence encodings all give p > 0.05, and the raw-count result dies under BH-FDR across 230 taxa (q = 0.76). (2) The provenance is retracted: the TCGA tumor-microbiome data (Poore et al. 2020, Nature) was retracted in 2024 (Gihawi et al.) for batch/contamination/human-read errors, so the methodology behind p ~= 0.0234 is no longer defensible. (3) Kocuria is a recognized low-biomass sequencing contaminant (89% zeros, 51/457 nonzero) with no peer-reviewed link to LUAD prognosis. The gold answer is literally correct only under an under-specified recipe (raw count, nominal p, no FDR) built on retracted data, so the cell cannot fairly discriminate model capability.

##### da-18-7 — Rubric demands a one-sided Fisher's exact test for an explicitly two-directional question, penalizing the statistically correct two-sided test.

- **Original question:** Does the genomic data show ESR1 mutations and MAPK pathway alterations as mutually exclusive or co-occurring in post-hormonal therapy HR+/HER2- breast tumors (MSK-IMPACT cohort)?
- **Failure mode:** The agent constructed a justified 705-sample post-treatment HR+/HER2- cohort, identified 131 ESR1-mutant samples (all top variants within the LBD residue range 300-550), defined MAPK alteration over a documented 32-gene RTK/RAS/MAPK set with role-aware CNAs, built the 2x2 table, and ran a two-sided Fisher's exact test (OR = 0.76, p = 0.17), correctly concluding the two are neither significantly exclusive nor co-occurring. The rubric docked 15 points across Criteria 2, 3, and 4 — most damagingly Criterion 4, which awards A only for a 'one-sided Fisher's exact test.'
- **Why the benchmark item is invalid:** Criterion 4 penalizes the statistically correct test for the question posed. The instruction (line 7) asks whether the alterations are 'mutually exclusive OR co-occurring' — a bidirectional association question with no prior directional hypothesis, for which two-sided Fisher's exact is the standard and correct choice; one-sided would presuppose testing only for exclusivity. The rubric's level-A text ('performs a one-sided Fisher's exact test to evaluate mutual exclusivity') directly contradicts the instruction's bidirectional framing. Criteria 2 and 3 are phantom penalties for un-communicated specifics (an exact ESR1 amino-acid range and a particular MAPK gene list): the agent's observed ESR1 variants already lie within residues 300-550, and its 32-gene MAPK set documents and contains every rubric gene, so formal restriction would not change the counts or the conclusion. The cell rewards a backward statistical choice and undocumented selection rules rather than scientific quality.

##### da-20-1 — Rubric pins parameter values and a 'most-similar pair' answer that are absent from the instruction, contradicted by a statistical tie, and justified by a biologically incorrect marker rationale.

- **Original question:** Do the four primary cell types show distinct or overlapping baseline transcriptional signatures, and which cell types are most similar to each other (DRUG-seq bulk RNA-seq)?
- **Failure mode:** The agent ran a clean unsupervised pipeline (576 DMSO baseline samples, logCPM, top-5,000 HVGs, 20-component PCA, k-means k=4) and recovered all four cell types essentially perfectly (ARI = 0.991, purity = 0.997), then ranked pair similarity by three metrics and reported SkMM-Fibroblast as most similar (with AoSMC-SkMM a statistical near-tie, <1% apart). It lost points on Criterion 1 (C, used all three DMSO concentrations rather than 0.0625% alone), Criterion 2 (5,000 vs ~10,000 genes), Criterion 3 (20 vs 50-100 components), and Criterion 5 (C, reported SkMM-Fibroblast instead of the mandated AoSMC-SkMM), scoring 0.46 despite flawless clustering.
- **Why the benchmark item is invalid:** The benchmark item fails on three points. (1) Inverted requirement: Criterion 1 demands DMSO at 'percent_volume_dmso == 0.0625', but the instruction says only 'DMSO controls' with no concentration constraint — using all three balanced vehicle levels is scientifically sound and the 0.0625% restriction appears nowhere in the brief. (2) Over-determined tie with a wrong rationale: Criterion 5 requires AoSMC-SkMM as the answer 'due to their shared mesenchymal/contractile lineage' evidenced by ACTA2 and TAGLN, but the data ranks SkMM-Fibroblast and AoSMC-SkMM within <1% on all three metrics (an unacknowledged tie), and ACTA2/TAGLN are canonical smooth-muscle and fibroblast markers, not skeletal-myoblast markers (skeletal myoblasts are marked by PAX7/MYOD1/MYOG/MYF5). The rubric's biological justification is internally inconsistent and factually incorrect. (3) Criteria 2

and 3 pin exact parameter values (~10,000 genes, 50-100 SVD components) that appear nowhere in the instruction, which only asks for a 'standard DRUG-seq pipeline.' A perfect clustering is graded as a near-failure on undocumented, partly incorrect requirements.

##### da-6-2 — Rubric imposes a hidden, internally incoherent all-four-timepoint significance filter that contradicts the temporal-dynamics question and discards 86% of responsive genes.

- **Original question:** How does gene expression in vastus lateralis skeletal muscle (SKM-VL) dynamically change over an 8-week endurance-training course in males and females, and what are the predominant sex-specific or shared temporal expression patterns (MoTrPAC)?
- **Failure mode:** The agent filtered to the SKM-VL transcriptomics slice, applied BH-FDR per timepoint, built a 3-state (Up/Down/none) directional encoding per timepoint, classified genes as shared/female-only/male-only mutually exclusively, ranked patterns, and reported quantitative temporal dynamics plus named example genes and biology. Criterion 2 was scored C (0/20) for not requiring significance at ALL four timepoints, and the failure cascaded to Criterion 4 (B instead of A, -10), pushing the cell to 0.35 despite standard, higher-resolution temporal-genomics methodology.
- **Why the benchmark item is invalid:** The item fails on four axes. (1) Rubric-instruction mismatch: Criterion 2 demands significance at all four timepoints (1w/2w/4w/8w) per sex, a completeness requirement absent from the instruction, which asks how expression 'dynamically changes' and for 'temporal patterns' — language that explicitly embraces onset, transient, and late-response genes. (2) Internal incoherence: the criterion cites 'training_q < 0.05' as the metric, but training_q is a single per-gene FDR constant identical across all four timepoints (verified: all 1,164 gene-sex pairs share one training_q), so the all-4 filter is not even computable from the named metric. (3) The filter contradicts the task: it retains only 161/1,164 (13.8%) of responsive gene-sex pairs, discarding 86% that embody the very temporal dynamics the question asks about. (4) Factual slip: Criterion 1 conflates SKM-VL (vastus lateralis) with 'gastrocnemius', two distinct tissues, corroborating a hastily written rubric. Exceeding ~70/100 requires applying the hidden, incoherent filter.

#### Symmetric exclusion across models

The four excluded task IDs form a single fixed set (da-12-4, da-18-7, da-20-1, da-6-2), applied as one fixed exclusion set and removed identically from the per-dimension score denominator of every one of the seven models (gpt-5.5, ds4-pro, gpt-5.4, ds4-flash, mimo-v2.5-pro, mimo-v2.5, gpt-5.4-mini). The exclusion is model-independent: each of the four is a real, fully graded cell for all seven models (28/28 grade.json records present), and the same four cells are dropped from all models before any dimension is computed. The decision is not selectively favorable to the top model. The per-model raw scores on these four cells vary widely and do not advantage the leader — e.g. da-18-7 ranges from 0.00 (ds4-pro) to 0.82 (mimo-v2.5-pro); da-12-4 is 0.86 for the top model gpt-5.5 but a perfect 1.00 for ds4-pro, gpt-5.4, and ds4-flash, so removing it actually withdraws cells on which three competitors outscored the leader; da-20-1 ranges 0.13-0.71 and da-6-2 ranges 0.20-0.40 across models. The cut is therefore a fixed correction applied uniformly to the denominator, not a per-model cherry-pick.

#### Caveats

- Grading is performed by an LLM judge against a fixed text rubric (tests/llm_judge.py + tests/rubric.txt per task); the invalidity arguments concern the rubric/gold definitions themselves, which are deterministic artifacts, not run-to-run judge variance.

- Trajectory-level evidence (what the agent actually computed, e.g. cohort sizes, encodings, reported statistics) was audited from a single canonical run per task (the OmicOS gpt-5.5 vertical_agent_selector run); the rubric and instruction defects, however, are properties of the benchmark item and hold independently of which model is graded.
- This is a documented, pre-registered benchmark-calibration decision encoded in the radar pipeline (FAIL_CASES in bench_radar.py, with one auditable failure-case write-up per task under docs/failure-cases/), not a silent or post-hoc drop; the four cells remain in the raw run records and can be re-included to reproduce the uncorrected 50-task figure.
- Exclusion was applied only to items where the gold/rubric cannot be satisfied without a methodological or factual error (wrong test, undocumented or self-contradictory filter, biologically incorrect rationale, or retracted source data); tasks where the agent merely scored low under a valid rubric were retained.
- The published 46-task set excludes exactly the four broken-rubric items addressed here; removing only these four reproduces the Fig. 2d per-model scores exactly. Two additional documented tasks (da-19-1 rubric-vs-instruction mismatch; da-8-3 missing classification data) are retained in the capability set, so the scientific-validity exclusion set for Fig. 2d is exactly four.

### *Supplementary Note 4 | OmicVerse-RebuildR community protocol*

#### 1. Overview — a community-scale reconstruction framework

OmicVerse-RebuildR is, in the words of its own `README.md`, "a fixed, reproducible protocol for porting R / Bioconductor packages to pure-Python `py-<pkg>` standalones, with cryptographic-grade numerical parity against the R reference." It exists so that a distributed community of contributors (human or LLM agent) can rebuild canonical R/Bioconductor methods as pure-Python mirrors that are demonstrably *identical* to the original — not merely "good enough" — and publish them under a shared GitHub organization (`github.com/omicverse` by default) and on PyPI.

Three foundational principles, quoted from the repo, frame everything below:

1. **The R source is the executable spec.** No reverse-engineering from papers. The agent runs the R reference on a fixed input fixture and compares its draft to that output every iteration.
2. **Parity is class-aware.** "Same output" means different things for embeddings (rotation-invariant), clustering (label-permutation-invariant), or pseudotime (correlation-invariant). The protocol pre-registers which numerical metric applies to which output and locks the threshold *before any agent code is written*.
3. **Reconstruction is not metric optimization.** The algorithm is never tuned to "look better" — it is tuned to be **identical** to R, then searched for speed under provably-equivalent algebraic rewrites.

**The agent pipeline stages.** `PROTOCOL.md` formalizes the objective as `f_T(x) ≈_C f_R(x)` for all `x ∈ B`, where `B` is the canonical fixture set and `≈_C` is the parity relation for algorithm class `C`. This is explicitly *fixture-level equivalence, not a proof of semantic equivalence over the full input domain*. The protocol runs in six sequential stages (numbered Stage 0–5 in `PROTOCOL.md`; Phase 0–5 in `CHECKLIST.md`):

| Stage | Name | What happens | Key artifacts |
|---|---|---|---|
| **0 / 0.5** | **Discovery** | Check whether the target is already ported; identify which upstream R deps have py-mirrors under the org. STOP if | `DISCOVERY.md` (committed before any algorithmic code) |

| | | | |
|---|---|---|---|
| | | duplicate; reuse deps as `pyproject.toml` deps if found. | |
| 1 | **Shape template** | Copy directory layout + test scaffold from a prior port matching the algorithm class. Do **NOT** copy algorithmic code or thresholds. | `py-<pkg>` layout from `templates/` |
| 2 | **Dual environments** | Provision a Python target env and an R reference env side-by-side; both see the same fixture files. | conda envs `rebuild-py`, `rebuild-r` |
| 3 | **Two-agent inner loop** | (3a) Equivalence Agent: translate R → Python, iterate until the pre-registered class-aware parity gate clears. (3b) Acceleration Agent: verifier-guided search over algebraic rewrites for speed, each carrying an admissibility proof. | `ITERATION_LOG.md`; per-iteration branches |
| 4 | **Validate** | Re-confirm the gate at the committed threshold (never tightened or loosened). Produce all mandatory deliverables. | `CHECKLIST.md`, four notebooks, `AUDIT.md` |
| 5 | **Release** | Ship wheel to PyPI; publish `github.com/<org>/py-<pkg>`; complete `RECONSTRUCTION_REPORT.md` + four notebooks; vendor into OmicVerse; add as seed template. | PyPI + GitHub release; `RECONSTRUCTION_REPORT.md` |

**Always-first invariant:** Phase 0.5 (Discovery) is non-skippable. If discovery is skipped the protocol fails — at risk of re-implementing upstream work that already exists.

**No deferred items in Phase 4:** all artefacts are mandatory. The repo records that TSCAN-v0.1 deferred Notebooks 1+2 and shipped without them; v0.2 added them retroactively; v0.3 added Notebook 3; v7 added Notebook 4. The protocol now blocks all four from being skipped.

The eight algorithm classes and their pre-registered gates, the three acceleration admissibility-proof classes, and the engine scripts (`engine/loop.py`, `engine/parity_metrics.py`, `engine/benchmark.py`, `engine/discover_omicverse_deps.py`, `engine/r_function_audit.py`, `engine/plot_evolution.py`, `engine/smoke_test.py`) are the runnable substrate that contributors fill in; the engine "is the structure the LLM coding agent fills in — not an autonomous orchestrator."

---

#### 2. Contribution rules

A new port is proposed, built, and added by following `SETUP.md` then `PROTOCOL.md` / `CHECKLIST.md` verbatim. The repo's hard rules (`PROTOCOL.md` "Critical Hard Rules") govern what is and is not allowed.

**Who/what is eligible (from README "When to Use RebuildR").** Use RebuildR when the target is an R / Bioconductor package with clear numerical output (vector, matrix, table, cluster IDs, p-values); a canonical input fixture can be constructed small enough that the R reference runs end-to-end in < 1 min; the upstream license is permissive or copyleft you can match; and you can commit roughly 1–5 working days. Do **not** use it when the "R package" is closed-source or paper-only (no executable spec means no parity oracle); when the algorithm depends critically on R-only C++ extensions or S4-heavy Bioconductor classes; when you want a Python algorithm *better* than R (the kit refuses to widen the gate — fork after the port lands); or for GPU-only kernels with no CPU reference.

**The executable-R-as-spec rule.** The R source is the spec. Every Equivalence-Agent iteration runs the R reference driver on the fixture and compares the Python candidate against that JSON output under the pre-registered class-C metric. Contributors never reverse-engineer from a paper, and never tune the algorithm to improve a metric — the reference output is the ceiling, not the floor (per `EVALUATION.md`: "Never beat R metric (beating it means the algorithm has drifted)").

**Setup / environment prerequisites (`SETUP.md`).**

- Prerequisites: Python ≥ 3.9, R ≥ 4.0, `conda/mamba`, `gh` CLI, `git`, Jupyter. (`discover_omicverse_deps.py` calls `gh repo list <org>`; release helpers call `gh repo create` / `gh release create`.)
- Two side-by-side conda envs: a Python target env `rebuild-py` (`pip install -r omicverse-rebuildr/requirements.txt`) and an R reference env `rebuild-r` (`r-base=4.3` + Bioconductor + the upstream R package per port).
- Tell the kit where the envs live by exporting `PYTHON_TEST_ENV` and `R_TEST_ENV` (look up paths via `conda info --envs`). Default fallbacks in `engine/loop.py` and `templates/test_exact_match.template.py` are literal `/path/to/your/<X>/env` placeholders; if not exported, the agent exits with a clear error.
- Authenticate `gh` (`gh auth login`) — needed for Discovery (Phase 0.5) and Release (Phase 5). Optionally authenticate `twine` via `~/.pypirc` for PyPI; if you are not a maintainer, Phase 5 `twine upload` is rejected and you ship the wheel + tarball as GitHub release assets instead.
- Smoke-test the kit: `python -m engine.smoke_test`, expecting `[smoke] OK — all engine modules import and 8 parity metrics compute correctly.`
- Run the duplicate check before anything else: `python -m engine.discover_omicverse_deps --check <YourTargetRPackage>`. If it reports "No existing omicverse port found", proceed with `PROTOCOL.md`.

**Org override.** The kit refers to `github.com/omicverse` throughout. For a different org, find-replace `s/omicverse/<your-org>/g` across docs; engine scripts read the org from `engine/discover_omicverse_deps.py::ORG_NAME`, and ports publish under `github.com/<org>/py-<pkg>` (override with `REBUILDR_ORG`).

**The ten hard rules (verbatim from `PROTOCOL.md`).**

4. **Discovery must complete first** — if the target is already ported, the protocol ends immediately.
5. **`DISCOVERY.md` committed before algorithmic code** — documents the duplicate check and upstream dep decisions.
6. **Parity threshold pre-registered in `manifest.yaml` before Step 3 begins** — read-only during all subsequent work.
7. **No threshold widening** — if the gate fails after Acceleration, roll back to the last clearing commit; never adjust the threshold.
8. **Equivalence Agent exits only when the pre-registered gate clears** — do not tighten/loosen to fit candidate output.
9. **Acceleration runs only after the Equivalence gate passes** — never attempt rewrites before parity is achieved.
10. **All four notebooks mandatory at release** — missing any one blocks publication.
11. **Every committed rewrite must cite admissibility evidence** — inline comment or docstring required.
12. **Bounded approximations must derive a perturbation bound** — recorded in `MATH.md` or docstring, not handwaved.
13. **Package layout copied, code not copied** — only repository structure and test rig come from a template, never algorithms.

**Discovery dependency-action mapping (`DISCOVERY.md`).** For each upstream dep, the contributor records a decision: a direct mirror of the *target* → **stop, reuse existing repo**; a mirror of an `Imports:` dep → add to `pyproject.toml::dependencies` (hard dep); a mirror of a `Suggests:` dep → add as optional dep; a `LinkingTo:` dep → usually a Rust port, use as hard dep; description-substring-only match → lower confidence, read before depending; no match → native Python equivalent (scipy/sklearn/statsmodels/networkx) or a roadmap TODO. When publishing a new port whose repo name does not follow the auto-mirror rule (R::<X> → omicverse/py-<X>, py-<X>R, rust-<X>), add it to ALIAS_MAP. The repo list is cached in engine/omicverse_repos.cache.json for 24 h (override with --refresh).

---

#### 3. Port template & required repo layout

Every port copies its scaffolding (layout + test rig only, never algorithmic code) from templates/ and a class-matched seed port. The canonical directory structure from TEMPLATE.md:

```
py-<pkg>/
├── README.md                  # User-facing: Install / Quickstart / Function-map / Reproducing-R /
Citation / License
├── LICENSE                    # Match upstream R package license (GPL-3 if upstream GPL≥2; MIT if
MIT/BSD/Apache)
├── pyproject.toml             # Build + deps + metadata; version pinned to 0.1.0
├── .gitignore                 # Standard Python + ignore data/raw, <pkg>-ref/, dist/
│
├── <pkg_modulename>/          # The actual Python package (lower snake_case)
│   ├── __init__.py            # Public API surface (class + function exports)
│   ├── core.py                # Main class / orchestrator (entry point)
│   ├── <algorithm_a>.py       # One file per algorithmic step
│   ├── <algorithm_b>.py
│   ├── preprocessing.py
│   └── utils.py
│
├── tests/
│   ├── __init__.py
│   ├── conftest.py            # pytest fixtures (load reference outputs); session-level caching
│   ├── test_smoke.py          # "does it run at all" — fastest test
│   ├── test_exact_match.py    # THE parity gate against the R reference (asserts metric ≥ threshold)
│   ├── test_<feature>.py      # Per-feature / per-function parity tests, locked as gates clear
│   ├── parity_metrics.py      # Vendored copy of engine/parity_metrics.py
│   ├── _run_candidate.py      # Candidate runner → produces data/candidate_output.json
│   └── r_reference_driver.R   # R reference runner (pins seed, dumps reference_output.json)
│
├── examples/                  # MANDATORY: all 4 notebooks below, pre-executed with outputs committed
│   ├── compare_R_vs_Python.ipynb        # Notebook 1 — pipeline-level parity (6-section schema)
│   ├── tutorial_<dataset>.ipynb         # Notebook 2 — Python-only function tutorial (7 sections)
│   ├── function_by_function_R_parity.ipynb  # Notebook 3 — R⇄Python per-function parameter dictionary
│   ├── evolution.ipynb                  # Notebook 4 — per-iteration narrative (one ## Iteration N header each)
│   ├── r_per_function_dump.R            # R driver dumping per-function outputs (for Notebook 3)
│   ├── evolution.png                    # Two-panel acceleration plot (auto-rendered by engine.plot_evolution)
│   ├── r_driver_<dataset>.R             # R driver for Notebook 1 (if not already in tests/)
│   ├── *.executed.ipynb                 # Pre-executed copies for GitHub preview (optional)
│   └── data/                            # Small fixture copies (or .gitignore'd if large)
│
├── data/                      # Reference fixtures
│   ├── manifest.yaml          # Pre-registered parity gate — committed BEFORE agent work, read-only after
│   ├── fixture_<dataset>.h5ad # Canonical fixture (cells×genes AnnData, raw counts in .X)
│   ├── reference_output.json  # Cached R reference output
│   └── candidate_output.json  # Cached Python candidate output
│
├── <pkg>-ref/                 # Cloned upstream R source (gitignored)
│   ├── R/                     # R/*.R (audited by engine.r_function_audit)
│   ├── DESCRIPTION            # Parsed by discover_omicverse_deps for dep audit
│   ├── NAMESPACE              # Exported-function list (drives §2 coverage audit)
│   └── R_README_original.md
│
├── dist/                      # Build artefacts (gitignored)
│
├── MATH.md                    # Acceleration Agent's derivations (only if class B/C; every (B) bound here)
├── ITERATION_LOG.md           # Per-iteration YAML log (Phase 3 only)
```

```
├── RECONSTRUCTION_REPORT.md        # Structured "is the port done?" audit, 8 sections (Phase 4)
├── DISCOVERY.md                    # Phase 0.5 upstream dep audit (committed before algorithmic code)
├── CHECKLIST.md                    # Per-port progress tracker (copied from omicverse-rebuildr)
└── PROTOCOL_NOTES.md               # (optional) port-specific deviations
```

**Naming conventions (`TEMPLATE.md`).** GitHub repo py-<PkgName> (matches R casing, e.g. py-DoubletFinder); PyPI distribution py<pkgname> lowercase (pydoubletfinder); Python module <pkgname> snake_case; public class <PkgName> PascalCase; public functions snake_case (paramSweep → param_sweep).

**`data/manifest.yaml` schema (from `engine/manifest.template.yaml`).** This is the pre-registered gate; committed before the Step 3 agent loop and read-only after. Keys:

```
package: py-<PkgName>
upstream:
  name: <UpstreamR>                 # e.g. DoubletFinder, Slingshot, TSCAN
  version: "1.2.3"                  # pinned upstream version
  source: <CRAN | Bioconductor | GitHub>
  url: https://github.com/<user>/<repo>

algorithm_class: ordinal | clustering | embedding | deterministic | stochastic |
                 ranked | classification | inference |
                 deterministic-strict | deterministic-bounded   # pick ONE (PARITY_TAXONOMY.md)

parity_threshold: <float>           # from PARITY_TAXONOMY.md defaults (0.99 ordinal, 0.95 clustering, 1e-8
deterministic)
parity_rtol: 1e-8                   # optional, for multi-order-of-magnitude outputs
parity_atol: 1e-10

fixture:
  path: data/fixture_<dataset>.h5ad  # cells×genes AnnData, raw counts in .X
  source: <public URL or scanpy.datasets.<name>()>
  description: "<one line>"
  expected_shape: [<n_cells>, <n_genes>]

reference_command: tests/r_reference_driver.R
seed: 42                            # pinned immediately before the algorithm call in BOTH ref and candidate

outputs:                            # supports multi-output algorithms; ALL must pass
  - name: <output_name>
    type: "1d vector" | "matrix" | "cluster labels" | ...
    location_reference: "$.FieldName"          # JSON path in reference_output.json
    location_candidate: "adata.obs['field']"   # how to find it in candidate AnnData
    metric: <pearson | ari | procrustes | max_abs_err | ...>
    threshold: <float>

held_out_fixture:                   # optional, for hidden-fixture scoring
  path: null
  description: null

acceleration:
  enabled: true                     # false for class-A translation-only ports
  max_iterations: 20
  must_pass_gate: true              # rewrites that break the gate are rolled back

audit_class: null                   # A | B | C — filled AFTER the port is done
created_by: <agent or human handle>
created_at: <YYYY-MM-DD>
```

**`pyproject.toml` conventions (`templates/pyproject.template.toml`).** name = "py<pkgname>" (lowercase, no hyphens); version = "0.1.0" (pinned until first release); requires-python = ">=3.9"; license = { file = "LICENSE" }; core dependencies = ["numpy>=1.23", "scipy>=1.10", "pandas>=1.5", "anndata>=0.9", "scikit-learn>=1.2", "matplotlib>=3.6", "tqdm"]; [project.optional-dependencies] dev =

["pytest>=7", "pytest-cov", "ruff"] and scanpy = ["scanpy>=1.9"]; [project.urls] point Homepage/Repository/Issues at github.com/omicverse/py-<PkgName>; [tool.setuptools.packages.find] excludes tests*, examples*, <pkgname>-ref*; [tool.pytest.ini_options] sets testpaths = ["tests"].

**R reference driver pattern (`tests/r_reference_driver.R`).** Invoked under $R_TEST_ENV as Rscript r_reference_driver.R <fixture.{h5ad,rds}> <output.json>; loads the fixture, runs set.seed(42) **before** the algorithm call, runs <UpstreamR>::<entry_point>(...) with pinned defaults, flattens the result to a JSON list whose keys match manifest.yaml::outputs[*].location_reference, and writes it with write_json(..., auto_unbox = TRUE, digits = NA, na = "null").

**Parity gate test (`tests/test_exact_match.py`).** Loads manifest.yaml; runs the R driver under $R_TEST_ENV → reference_output.json; runs _run_candidate.py under $PYTHON_TEST_ENV → candidate_output.json; for each outputs[] extracts location_reference/location_candidate, computes the class-aware metric, asserts metric >= threshold, prints each sub-gate pass/fail, fails the build if **any** gate fails, and **does not permit threshold loosening**.

**`RECONSTRUCTION_REPORT.md` — 8 mandatory sections** (RECONSTRUCTION_REPORT.template.md): §1 Identity (package, upstream version, class, threshold + final parity, audit class A/B/C, LOC, speedup, memory gain); §2 R-function coverage audit (§2.1 exported from NAMESPACE, §2.2 internal helpers, §2.3 coverage summary requiring ≥ **95% exported + every transitive dependency ported**, §2.4 deliberately skipped, §2.5 dependencies reused from omicverse with LOC saved); §3 Parity evidence (§3.1 per-output, §3.2 per-fixture wall-clock + speedup, §3.3 reproducible reference command); §4 Acceleration evidence (§4.1 two-panel evolution.png, §4.2 accepted rewrites, §4.3 rejected rewrites); §5 Code-quality audit (all ✅, non-skippable); §6 Known limitations; §7 Integration into omicverse; §8 Sign-off (author, date, active time, audit class).

**`ITERATION_LOG.md` schema (`ITERATION_LOG.template.md`).** One YAML block per Acceleration step (baseline iter: 0 plus every attempt, accepted or rejected), parsed by engine/plot_evolution.py. Per-iteration keys include iter, status (ACCEPT | REJECT_GATE | REJECT_SLOW | REJECT_INADMISSIBLE), action, playbook_section, admissibility (exact | bounded | containment), admissibility_evidence, perturbation_bound (B-only), wall_clock_mean_s / wall_clock_stddev_s / wall_clock_runs_s (warmup-excluded) / warmup_run_s, speedup_vs_previous, speedup_vs_baseline, parity_metric, parity_delta_vs_baseline, parity_passes, and math_reason_for_dip (rendered next to the dip on Plot 2).

---

#### 4. Review checklist

The full pass/fail review checklist, grouped by category, drawn from CHECKLIST.md, PARITY_TAXONOMY.md, EVALUATION.md, ACCELERATION_PLAYBOOK.md, and NOTEBOOKS.md. Every box must be ticked; "deferred" is not a valid status.

##### 4.1 Pre-registration & Discovery

- [ ] Canonical R entry point and output type identified (Phase 0).
- [ ] Algorithm class looked up in PARITY_TAXONOMY.md; one class chosen per output.
- [ ] Parity threshold set from PARITY_TAXONOMY.md defaults; canonical fixture + seed: 42 chosen.
- [ ] data/manifest.yaml committed with algorithm_class, parity_threshold, fixture_path, reference_command, seed, outputs[] — **read-only thereafter**.
- [ ] python -m engine.discover_omicverse_deps --check <PkgName> run; no sister port exists (else STOP and open an issue).
- [ ] Upstream cloned into <pkg>-ref/; --description <pkg>-ref/DESCRIPTION --output DISCOVERY.md run; each R dep mapped to hard dep / optional dep / native-Python replacement.
- [ ] DISCOVERY.md committed **before** any algorithmic code.

#### 4.2 Parity gates — matched to output class (`PARITY_TAXONOMY.md`)

The pre-registered gate must clear on all required fixtures at the committed threshold; no per-fixture tuning, no Bonferroni-style correction — one class, one threshold, fixed before agent work.

- [ ] **Class 1 — Deterministic numerical**: element-wise max_abs_err < atol AND Pearson. Sub-tiers deterministic-strict (1e-13), deterministic-standard/deterministic (1e-8), deterministic-bounded (1e-6). **Hard ceiling: any threshold > 1e-6 is rejected by `is_pass()`** — switch to ordinal or embedding instead.
- [ ] **Class 2 — Stochastic numerical**: KS p-value ≥ 0.05 **or** Wasserstein-1 ≤ 1% of scale.
- [ ] **Class 3 — Combinatorial clustering**: label-invariant **ARI ≥ 0.95** (NMI / Fowlkes–Mallows secondary).
- [ ] **Class 4 — Continuous embedding**: rotation-invariant **Procrustes ≥ 0.95** (similarity = 1 - disparity).
- [ ] **Class 5 — Ranked output**: **top-50 Jaccard ≥ 0.8** and Spearman ≥ 0.9.
- [ ] **Class 6 — Ordinal / pseudotime**: **Pearson ≥ 0.99** (is_pass subtracts 1e-12; ≥ 1 - 1e-12 treated as exact); Spearman secondary.
- [ ] **Class 7 — Classification**: agreement / **F1 ≥ 0.95** (preserve original label names; do not relabel to 0/1).
- [ ] **Class 8 — Statistical inference**: **Spearman ≥ 0.90 on −log10 p** and top-50 Jaccard ≥ 0.7.
- [ ] If the entry point returns multiple outputs of different classes, one gate per output in manifest.yaml::outputs[] and **all** pass.
- [ ] For deterministic-bounded, MATH.md derives Σ bound ≤ atol. Metrics imported from engine/parity_metrics.py (vendored into tests/parity_metrics.py), not redefined.

#### 4.3 Tests / CI

- [ ] Per-function parity tests locked as tests/test_<function>_parity.py as each leaf clears (dependency order; leaves first, then orchestrator).
- [ ] tests/test_smoke.py runs (import + instantiate + don't crash).
- [ ] tests/test_exact_match.py clears the pre-registered gate on all required fixtures; threshold never loosened.
- [ ] pytest -q green; pip install -e . works.
- [ ] Smoke / benchmark notebook in examples/ runs end-to-end on the public fixture.

#### 4.4 Acceleration — admissibility proofs (`ACCELERATION_PLAYBOOK.md`, `EVALUATION.md`)

Runs **only after** the Equivalence gate passes. Every committed rewrite must carry one of three proofs, cited inline in code (e.g. # acceleration: §3.1 MST ⊆ Delaunay (C)):

- [ ] **(E) Exact identity** — bit-equivalent by algebraic identity (e.g. cache X^T X outside loop §1.1; Woodbury §1.2; Schur complement; memoization; vectorization).
- [ ] **(B) Bounded ε-approximation** — error bounded by a closed-form expression in MATH.md, not handwaved (e.g. sparse soft-assignment row truncation §2.1 with ‖W_new − W_old‖_F ≤ κ·n·K·ε; top-K kNN truncation §2.2, only when the downstream consumer is local-neighbourhood-bound).
- [ ] **(C) Class-containment theorem** — a known theorem guarantees identical output (e.g. Euclidean MST ⊆ Delaunay §3.1; kNN ⊆ relative-neighbourhood graph §3.2).
- [ ] Baseline benchmark recorded as iter: 0 (1 warmup + 3 measured runs; time.perf_counter(); BLAS threads pinned OMP/OPENBLAS/MKL_NUM_THREADS=8; if stddev > 10% of mean, run 5× and report median).
- [ ] A rewrite is merged only if **speedup > 1.05×** and the gate still clears; otherwise rolled back. Every attempt (accept + reject) appended as a YAML block to ITERATION_LOG.md.
- [ ] Every accuracy dip captioned with a math reason (a dip 1.000 → 0.998 with no (B) justification is a bug, not an optimization). Reference output is the ceiling — never beat R.

- [ ] python -m engine.plot_evolution --port-dir . renders examples/evolution.png; time is monotone-non-increasing on accepted iters, accuracy flat at ≈1.0 with annotated dips only where (B) fired.

##### 4.5 Docs / artifacts — four mandatory notebooks + report (`NOTEBOOKS.md`)

All four notebooks pre-executed with outputs committed; re-executed before release via jupyter nbconvert --to notebook --execute examples/*.ipynb --output {}. **If any one is missing, the port is NOT released — no "deferred" exception.**

- [ ] examples/compare_R_vs_Python.ipynb — 6-section schema (Setup / R reference run / Python candidate run / per-output parity per outputs[] / wall-clock comparison / Verdict), executed end-to-end in a fresh kernel, every parity sub-gate clears.
- [ ] examples/tutorial_<dataset>.ipynb — 7 sections, one subsection per public function from RECONSTRUCTION_REPORT §2.2, class-API mirror + ≥ 3 pitfalls + next-steps.
- [ ] examples/function_by_function_R_parity.ipynb — one subsection per public R function with a parameter table (R name / Py name / type / default / range / description for **every** parameter), R one-liner (markdown) + Python call + numerical comparison + sub-verdict, plus an aggregate verdict table.
- [ ] examples/r_per_function_dump.R — dumps per-function R outputs to JSON for Notebook 3.
- [ ] examples/evolution.ipynb — one ## Iteration N — <title> header per iteration (**minimum 2**: ## Iteration 0 — Baseline + ≥ 1 follow-up, even for class A), each with ≥ 3-sentence markdown narrative AND a code cell producing a subplot, final cell renders the 2-panel evolution.png. Header count is itself a quality signal.
- [ ] README.md complete: Install / Quickstart / Function-map table / Reproducing-R / Citation / License.
- [ ] MATH.md lists every (B) bounded-approximation rewrite with its perturbation bound (if class B/C).
- [ ] RECONSTRUCTION_REPORT.md filled: engine.r_function_audit → AUDIT.md pasted into §2; §1 / §3 / §4 / §6 / §8 complete; evolution.png embedded in §4.1; §5 code-quality all ✅; signed §8 with date + active time.

##### 4.6 Release & OmicVerse integration

- [ ] Version pinned to 0.1.0; license chosen per TEMPLATE.md decision matrix (MIT, or upstream license whichever more restrictive).
- [ ] Repo pushed to github.com/omicverse/py-<PkgName>; PyPI release workflow triggered.
- [ ] Port added to examples/ROADMAP_TRAJ.md as ✅ done; added to TEMPLATE.md seed-template table if it sets a new template for its class.
- [ ] Vendored into omicverse/external/<pkgname>/ (or omicverse/single/_<pkgname>.py for a small port).
- [ ] Exposed at top-level omicverse.<subpackage>.<PkgName> (matching existing patterns, cf. ov.single.Monocle); tutorial added to omicverse-guide.
- [ ] Audit retrospective written: class A (translation-only) / B (minor optimisation) / C (major algorithmic change with proof); which playbook entries fired and at what speedup; any new rewrite to add to the playbook.

---

#### 5. Release criteria

A port is **accepted/validated** (Stage 4, PROTOCOL.md) iff:

- All required fixtures pass the class-C gate at the pre-registered threshold (read-only).
- pip install . succeeds in a fresh conda env.
- pytest -q is green.
- The smoke notebook in examples/ runs end-to-end on the public fixture.

If the gate fails after Acceleration rewrites, the contributor rolls back to the last commit that cleared it; the gate is **never widened**.

A validated port is **released** (Stage 5) only when every one of the following ships:

- A fully-filled RECONSTRUCTION_REPORT.md including: the R-function coverage audit (python -m engine.r_function_audit --r-source <pkg>-ref --py-package <pkgname>), per-fixture parity values, the two-plot evolution figure (python -m engine.plot_evolution --port-dir .), and accepted-vs-rejected rewrites with admissibility evidence.
- All four mandatory notebooks under examples/, pre-executed: compare_R_vs_Python.ipynb, tutorial_<dataset>.ipynb, function_by_function_R_parity.ipynb, evolution.ipynb (one ## Iteration N header per iteration). "All four are mandatory — if any missing, the port is not released."
- omicverse/py-<pkg> repository created; pyproject.toml version pinned to 0.1.0; published to PyPI under MIT or the upstream R package's license (match upstream when more restrictive).
- The §5 code-quality audit all ✅ and non-skippable: pip install -e . in a fresh env, pytest -q green, all four notebooks executed with outputs committed, r_per_function_dump.R present, evolution.png rendered, README.md complete, MATH.md with bounds (if B/C), ITERATION_LOG.md complete, DISCOVERY.md committed, license compatible, version 0.1.0, GitHub repo created under the org, PyPI release passed CI.

After release the port is **integrated and made agent-callable**: vendored into omicverse/external/<pkgname>/ (or omicverse/single/_<pkgname>.py), exposed as omicverse.<subpackage>.<PkgName>, added to omicverse-guide, ticked off on examples/ROADMAP_TRAJ.md, and **added to `TEMPLATE.md` as a candidate seed template** so the next port can copy its layout. The engine loop (python -m engine.loop --port-dir <path> --phase equivalence | acceleration | release) exposes the same equivalence_phase / acceleration_phase / parity_diff functions the agent calls from inside its tool-use loop, so a released port is reproducible and re-runnable.

---

#### 6. How it scales (community)

**Self-seeding template economy.** Each finished port becomes the seed template for the next: layout, test rig, and packaging are copied (never algorithmic code), so the marginal cost of each port falls. EXAMPLE_WALKTHROUGH.md records that a first-time TSCAN port took ~3–5 working days for the Equivalence phase + ~1 day for Phase 4–5, but "subsequent trajectory ports inherit Mclust resolution and py-monocle2-style packaging, so the next 5 ports should take ~2 days each on average." Ports are also reusable as *dependencies*: if Port B depends on Port A's output, A's py-mirror is listed as a hard or optional dep in B's pyproject.toml (e.g. py-TSCAN reuses py-mclustR).

**Class-matched seed templates** (PROTOCOL.md / TEMPLATE.md): Ordinal/trajectory → py-monocle2; Classification → py-DoubletFinder; Clustering → py-mclustR; Statistical inference → py-miloR; Continuous embedding → py-CCA; Deterministic numerical → rust-bandnorm.

**Shipped ports (as of protocol v7):** ✅ py-monocle2 (2026-04, class C, 102× speedup via MST$\subseteq$Delaunay + Woodbury + $X^\top X$ cache), ✅ py-mclustR (2026-05, class A, fixed a Fraley 1998 hcVVV bug), ✅ py-TSCAN (2026-05, class A, ~28× speedup, full discovery + py-mclustR reuse). Already in OmicVerse (do **not** re-port): Slingshot, Palantir, VIA, CytoTRACE 2, CellFateGenie, STT, scdiffusion. Already-Python (do not re-port): PAGA, scVelo, CellRank, scFates, Wishbone, Wanderlust, elPiGraph.

**Community roadmap (`examples/ROADMAP_TRAJ.md`), ranked by citation density × benchmark importance:**

- **Tier 1:** tradeSeq (~152 cites/yr, inference, seed py-miloR), URD (~123/yr, ordinal, py-monocle2), TSCAN (~55/yr, ordinal), destiny/DPT (~34/yr, ordinal).
- **Tier 2:** SCORPIUS (~17/yr, ordinal), condiments (~19/yr, inference).

- **Tier 3:** SLICER, psupertime, Totem, Mpath, PhenoPath, ouija, CellTrails, reCAT, FateID, RaceID/StemID.

Suggested execution order: `py-TSCAN` (test the loop on a simple port) → `py-tradeSeq` (highest-ROI; needed by every benchmark) → `py-destiny` → `py-URD` → `py-SCORPIUS` → `py-condiments`. Each port "should follow PROTOCOL.md verbatim and tick through CHECKLIST.md."

**Engine automation underpinning scale.** The `engine/` package makes the protocol runnable and the review mechanical: `discover_omicverse_deps.py` (duplicate + dep audit, 24 h cached in `omicverse_repos.cache.json`), `parity_metrics.py` (the 8 class metrics + `compute_parity` / `is_pass` dispatcher with the deterministic hard ceiling enforced via `ValueError`), `benchmark.py` (`time_callable`: warmup-excluded 3-run mean, auto-extending to 5 runs + median when CV > 10%), `r_function_audit.py` (parses `NAMESPACE` + `R/*.R` → `AUDIT.md` coverage), `plot_evolution.py` (renders the two-panel figure from `ITERATION_LOG.md`), `loop.py` (the runnable two-agent loop the agent fills in — `Manifest`, `TrialResult`, `RewriteCandidate`, `equivalence_phase`, `acceleration_phase`, `parity_diff`), and `smoke_test.py` (30-second sanity check that all engine modules import and the 8 metrics compute). `smoke_test.py` deliberately does **not** test gh auth, conda env paths, R installation, or internet access — those are the contributor's setup responsibility.

**Long-term maintenance posture.** The repo is explicit about its boundaries: the kit pushes nowhere automatically (Phase 5's `gh repo create` and `twine upload` are user-driven); a depended-on port must itself be parity-validated before reuse (check its `RECONSTRUCTION_REPORT.md` — a port with `xfail` vs R is not safe to depend on); and depend on a port's public API only, never re-export internals, to avoid coupling to layout that may shift between releases. The protocol's own version history (v1 baseline → v3 forced Discovery → v4–v5 mandatory notebooks → v6 portability/`REBUILDR_ORG/SETUP.md` → v7 `evolution.ipynb` forcing function) shows the governance model: each audit of shipped ports that surfaces a skipped step becomes a new non-skippable forcing function in the protocol.

### *Supplementary Note 5 | dynbenchmark / dyneval trajectory metrics — definitions and formulas*

The trajectory benchmark (Fig. 3e) scores each (method × dataset) prediction against the gold-standard trajectory using the dynbenchmark metric set (Saelens et al., *Nat. Biotechnol.* 2019), implemented in the pure-Python **pydyneval** port (`py-dyneval`). Four metric families are reported — **Topology**, **Branch assignment**, **Cell positions** and **Features** — each a scalar in `[0, 1]` where `1` is a perfect match. This note gives the exact mathematical definition of every metric **as implemented**, grounded in the port source (`pydyneval/metric_*.py`, `pydynwrap/geodesic.py`, and the benchmark adapter `run_benchmark_v2.py`). Negative correlations are clamped to `0` throughout. The no-leak audit and root-selection rule are documented in the companion note `Supplementary_Note_dyneval_metrics.md`.

#### 0. Shared objects

- **Milestone network** `G = (V, E)`: a (directed) graph whose vertices `V` are milestones (branch endpoints / branch points) and whose edges `E` carry a `length`. Gold and predicted trajectories each have one. The adjacency matrix `A` has `A_{ij} = length(i→j)` (symmetrised for the topology metrics).
- **Milestone percentages**: each cell's position is a convex combination of the milestones of the edge it lies on; this places cells continuously on `G`.
- **Geodesic distance** `d_G(i, j)`: the shortest-path distance **along the trajectory graph** between the on-graph positions of cells `i` and `j` (`pydynwrap.calculate_geodesic_distances`, a 1:1 port of `dynwrap::calculate_geodesic_distances`). To bound cost it is evaluated between **waypoint cells** (≤ 100 sampled cells), giving a `n_waypoints × n_cells` matrix `D`.

The four reported scores map to dyneval metric IDs as follows (`run_benchmark_v2.py`, lines 352–358):

| Reported metric | dyneval metric ID | family |
|---|---|---|

| | | |
|---|---|---|
| **Topology** | `him` | milestone-graph distance |
| **Branch assignment** | `F1_branches` | cell-grouping agreement |
| **Cell positions** | `correlation` | geodesic-distance agreement |
| **Features** | `featureimp_wcor` | feature-importance agreement |

---

#### 1. Topology — `him` (Hamming–Ipsen–Mikhailov)

`pydyneval/metric_him.py::calculate_him` ports `dyneval::calculate_him` (R `netdist::netdist(·, "HIM")`).
Let $A^1$, $A^2$ be the symmetric adjacencies of the gold and predicted milestone graphs on a **shared node set**, each normalised $A \leftarrow A / \Sigma_{ij} A_{ij}$.

**Hamming component** (normalised L1 over the upper triangle):

$$H(A^1, A^2) = \left( \Sigma_{i<j} |A^1_{ij} - A^2_{ij}| \right) / \max\left( \Sigma_{i<j}|A^1_{ij}| + \Sigma_{i<j}|A^2_{ij}|, \varepsilon \right)$$

**Ipsen–Mikhailov component** (spectral distance of the graph Laplacians). For graph Laplacian $L = D - A$ (with degree matrix $D = \mathrm{diag}(\Sigma_j A_{ij})$), let $\{w_k\}$ be its non-negative eigenvalues and define vibrational frequencies $\omega_k = \sqrt{w_k}$. The spectral density is a sum of Lorentzians of width $\gamma$:

$$\rho(\omega) = (1/Z) \cdot \Sigma_k \; (\gamma/\pi) / \left( (\omega - \omega_k)^2 + \gamma^2 \right), \quad Z \text{ chosen so } \int \rho(\omega)\, d\omega = 1$$

and the Ipsen–Mikhailov distance is the L2 distance of the two densities (numerically integrated on a grid $\omega \in [0, 1.2\cdot\omega_{max}]$, 200 points):

$$IM(A^1, A^2) = \mathrm{sqrt}\left( \int ( \rho_1(\omega) - \rho_2(\omega) )^2 \, d\omega \right)$$

**HIM** combines the two with mixing constant $\gamma = 0.1$:

$$HIM(A^1, A^2) = \mathrm{sqrt}\left( ( H^2 + \gamma^2\cdot IM^2 ) / ( 1 + \gamma^2 ) \right)$$

**Reported score:** $\text{Topology} = 1 - HIM \in [0, 1]$ (returns `0` if either graph has no edges).

**Benchmark milestone-graph construction** (`run_benchmark_v2.py::_topology_him`, applied identically to gold and prediction so no method is advantaged): within each branch `b`, cells are binned by pseudotime into $K = 4$ milestones `b_m0 … b_m3` via `K+1` evenly spaced edges; an undirected edge is added between the milestones of **consecutive-pseudotime** cells in the same branch, and the first milestones `b_m0` of all branches are connected (shared origin). HIM is then computed on these adjacencies.

##### Topology extras (full dyneval set)

- **`edge_flip`** (`metric_flip.py`): Jaccard of the undirected edge sets, $\text{edge\_flip} = |E_1 \cap E_2| / |E_1 \cup E_2|$. (The `pydynbenchmark.metrics_extra.edge_flip` variant instead returns `1 - (#differing upper-triangle entries)/\binom{n}{2}`.)
- **`isomorphic`** (`metric_isomorphic.py`): `isomorphic = 1` if the two milestone graphs are isomorphic (`networkx.is_isomorphic`), else `0`.

---

#### 2. Branch assignment — `F1_branches` (and `F1_milestones`)

`pydyneval/metric_mapping.py`. Cells are grouped: for **branches**, by the trajectory edge they lie on (`group_onto_trajectory_edges`); for **milestones**, by the nearest milestone (`group_onto_nearest_milestones`). Let `g_data` be the gold grouping and `g_pred` the predicted grouping of the same cells. For every (gold group `a`, predicted group `b`) the **Jaccard overlap** is

$$J(a, b) = |a \cap b| / ( |a| + |b| - |a \cap b| )$$

($|a \cap b|$ = number of cells in both). Two one-sided scores follow:

```
recovery  = mean over gold groups a of  max_b J(a, b)       (how well each gold branch is recovered)
relevance = mean over pred groups b of  max_a J(a, b)       (how relevant each predicted branch is)
```

and the metric is their **harmonic mean**:

```
F1_branches = 2 · recovery · relevance / ( recovery + relevance )      (0 if either is 0)
```

**`F1_milestones`** is identical but with the milestone grouping. (The coarser `pydynbenchmark.metrics.f1_branches` helper instead aligns predicted to gold labels by **Hungarian assignment** on the confusion matrix and reports macro-averaged `sklearn.f1_score`; the reported benchmark uses the Jaccard `F1_branches` above.)

---

#### 3. Cell positions — `correlation`

`pydyneval/metric_correlation.py`. Build the gold and predicted geodesic-distance matrices over the shared waypoint set, `D_gold`, `D_pred` (`Inf → 1e308`), flatten them to vectors, and take the **Spearman rank correlation**, clamped to `[0, 1]`:

```
Cell_positions = max( 0, Spearman_ρ( vec(D_gold), vec(D_pred) ) )
```

where `Spearman_ρ(x, y) = Pearson( rank(x), rank(y) )`. Returns `0` if either vector is constant. This rewards predictions that **preserve cell–cell distances along the trajectory**, and is invariant to a monotone reparameterisation of pseudotime (which is why pseudotime parity uses rank/correlation).

---

#### 4. Features — `featureimp_wcor` (and `featureimp_cor`)

`pydyneval/metric_featureimp.py` ports `dynfeature::calculate_overall_feature_importance`. For a trajectory, fit a `RandomForestRegressor` (`n_estimators=50, max_depth=8, random_state=42`) regressing the per-cell **pseudotime** onto the gene-expression matrix, and read each gene's Gini importance. Do this for the gold trajectory (`a_g = imp_data`) and the predicted trajectory (`b_g = imp_pred`) over the set of common genes `g`.

**Unweighted (`featureimp_cor`):** Pearson correlation of the two importance vectors,

```
featureimp_cor = max( 0, Pearson(a, b) )
```

**Weighted (`featureimp_wcor`, the reported Features score):** importance-weighted Pearson with weights `w_g = max(a_g, 0)` (the gold importances, so genes that matter to the true trajectory dominate):

```
μ_a = Σ_g w_g a_g / Σ_g w_g ,            μ_b = Σ_g w_g b_g / Σ_g w_g
cov   = Σ_g w_g (a_g − μ_a)(b_g − μ_b) / Σ_g w_g
var_a = Σ_g w_g (a_g − μ_a)²            / Σ_g w_g ,   var_b analogous
featureimp_wcor = max( 0, cov / √(var_a · var_b) )
```

(The `pydynbenchmark.metrics.wcor_features` helper is a lighter alternative: pick the top-`N` genes by gold-pseudotime correlation and score `mean_j |Pearson(gene_j, pred_pt)|`.)

---

#### 5. Aggregation to the reported "Overall"

For each (method, dataset) the benchmark stores the four metrics `{Topology, Branch_assignment, Cell_positions, Features}`. The per-method scores and the **Overall** are the **arithmetic means** (`trajectory_benchmark.ipynb`, cell 4):

```
per_metric[method, m] = mean over datasets of metric m
Overall[method]       = (1/4) · Σ_{m ∈ {Topology, Branch_assignment, Cell_positions, Features}}
per_metric[method, m]
```

> **Note (honest caveat).** Canonical dynbenchmark aggregates metrics by the **geometric** mean (`pydynbenchmark.metrics.calculate_metrics` implements `overall = (∏_m (metric_m + 1e-6))^{1/k}`), which penalises any single near-zero metric more harshly. The Fig. 3e benchmark reports the **arithmetic** mean of the four families (the values in Supplementary Table 4_8); the two differ for methods that score very low on one family. The per-metric values are reported individually so either aggregation can be recomputed.

#### 6. Two implementations in the codebase

- **`pydyneval`** (`metric_him`, `metric_correlation`, `metric_mapping`, `metric_featureimp`, `metric_flip`, `metric_isomorphic`) — the faithful 1:1 dyneval port; its `calculate_metrics(metrics= ["correlation","him","F1_branches","featureimp_wcor"])` produces the four **reported** scores.
- **`pydynbenchmark.metrics` / `metrics_extra`** — lighter benchmark helpers (`him_distance`, `f1_branches` via Hungarian + macro-F1, `cor_dist` on a pseudotime-derived distance proxy, `wcor_features`, `f1_milestones`, `edge_flip`, `isomorphic_topology`, `pseudotime_spearman`) used for auxiliary/topology computations. Where the two differ, this note states the reported (pydyneval) definition first and the helper variant in parentheses.

##### Metric summary

| Metric (reported) | dyneval ID | Range | Formula (one line) | Invariance |
|---|---|---|---|---|
| Topology | him | [0,1] | $1 - \sqrt{(H^2 + \gamma^2 IM^2)/(1+\gamma^2)}$, $\gamma$=0.1 | graph relabeling |
| Branch assignment | F1_branches | [0,1] | harmonic mean of Jaccard recovery & relevance | branch relabeling |
| Cell positions | correlation | [0,1] | max(0, Spearman(vec D_gold, vec D_pred)) | monotone pseudotime rescaling |
| Features | featureimp_wcor | [0,1] | gold-importance-weighted Pearson of RF importances | feature ranking |
| (extra) edge_flip | edge_flip | [0,1] | | $E_1 \cap E_2$ |
| (extra) isomorphic | isomorphic | {0,1} | $1[G_1 \equiv G_2]$ | full graph relabeling |
| (extra) F1_milestones | F1_milestones | [0,1] | as F1_branches on milestone grouping | milestone relabeling |
| (extra) featureimp_cor | featureimp_cor | [0,1] | max(0, Pearson(imp_gold, imp_pred)) | feature ranking |

#### *Supplementary Note 6 | Method-specific optimization explanation for the large-speedup trajectory ports (destiny, URD)*

**Scope.** Two trajectory ports carry order-of-magnitude speedups over their R references on the benchmark — **py-destiny (~139×)** and **py-URD (~68×)**. Because a single "speedup" number can hide whether a port simply runs the same algorithm faster or has quietly become a *different* method, this note decomposes each speedup into five sources — (i) Python implementation choices, (ii) sparse linear algebra, (iii) caching/precomputation, (iv) algorithmic rewrites, and (v) reduced language/runtime overhead — and states the admissibility class of every algorithmic change. Attributions are grounded in the port source code (`pydestiny/diffmap.py`, `pydestiny/dpt.py`, `pyurd/flood.py`, `pyurd/random_walk.py`) and the R reference behaviour.

##### Benchmark-invocation caveat (read first)

The R references are executed as a **subprocess**: `conda run -p /scratch/users/steorra/env/CMAP Rscript r_methods_driver.R …` (R 4.4.3, CMAP env), with a netcdf `LD_LIBRARY_PATH` prepended. The Python ports run **in-process** in the omicdev env. On the benchmark's deliberately small datasets (300–600 cells, 100–150 genes), the **fixed per-call R-invocation cost** — conda-environment activation + Rscript interpreter startup + package/S4 load + shared-library resolution, on the order of seconds — is a *non-trivial fraction* of total R wall-clock. The headline speedups therefore blend (a) a durable **compute-level** speedup (sparse eigensolver, vectorized flood) with (b) a **fixed invocation overhead** that would shrink in relative terms on larger datasets. The compute-level component is the scientifically meaningful one; the per-dataset detail in Supplementary Table 4_9 (speedup rising on the smallest datasets, e.g. ~900× on the 300-cell prosstt_bifurcation for destiny) is the signature of this fixed-overhead contribution.

##### py-destiny — ~139× (diffusion map + DPT)

**R reference behaviour.** R `destiny` (Angerer et al. 2016) builds a **dense N×N** Gaussian kernel, density-normalises the full matrix, and computes a **full eigendecomposition** of the symmetrised transition matrix, inside the R interpreter with S4-object overhead.

**Attribution.**

- **Python implementation choices — major.** Full pipeline (cdist, local-σ, kernel, normalization) in NumPy/SciPy C bindings, no Python loops; broadcasting for $K=\exp(-D^2/\sigma)$
- **Sparse linear algebra — dominant.** kNN truncation of the kernel (top-k per cell via np.partition mask) → sparse P; scipy.sparse.linalg.eigsh for only the top n_eigs=5 eigenpairs, vs R destiny's dense N×N kernel + full eigendecomposition ($O(N^3) \rightarrow O(N \cdot k^2 \cdot \text{iters})$)
- **Caching / precomputation — minor.** Pairwise distance matrix D computed once and reused for σ-estimation and kernel build
- **Algorithmic rewrites — moderate.** Local-σ (mean of n_local NN distances) + kNN sparsification of the kernel; exact/bounded — eigenvalues match to ~ARPACK rounding, eigenvector Procrustes 0.916, DPT Pearson 0.974
- **Reduced language/runtime overhead — moderate.** Runs in-process vs R via `conda run -p CMAP Rscript` subprocess: conda-env activation + Rscript startup + netcdf lib load + S4 dispatch (~seconds fixed cost)

**Admissibility.** The kNN-sparsified kernel and local-σ are exact/bounded approximations, not a different algorithm: eigenvalues match R to ARPACK rounding, eigenvector subspace Procrustes = 0.916 (gate 0.80), DPT Pearson = 0.974 (gate 0.85) — all parity gates cleared. The dominant lever (sparse truncated eigensolver computing only the top-5 eigenpairs of a sparse kernel, versus a dense full eigendecomposition) is the canonical "same statistical target, lower complexity" rewrite.

#### py-URD — ~68× (flood-pseudotime random walks)

**R reference behaviour.** R `URD` (Farrell et al. 2018) `floodPseudotime` runs many stochastic random-walk simulations from the root over the diffusion-map transition matrix, with **per-cell, per-step loops in interpreted R** — the dominant cost.

**Attribution.**

- **Python implementation choices — dominant.** floodPseudotimeCalc inner step fully vectorized: tm_sub=tm[ix_(unvisited,visited)] slice, visit_prob=1−exp(Σ log(1−p)) over all unvisited cells in one reduce, rng.binomial(1,visit_prob) batch-samples all cells — replaces R's nested per-cell random-walk loops (the dominant R cost)
- **Sparse linear algebra — major.** Transition matrix from pydestiny is sparse CSR (~k≈20 nnz/row); sparse slicing avoids dense N×N copies (but normalization/sub-matrix materialization caps the gain)
- **Caching / precomputation — moderate.** Reuses the pydestiny diffusion-map transitions/eigvecs/eigvals on the URD object instead of recomputing the DM
- **Algorithmic rewrites — minor.** n=30 stochastic floods (empirically sufficient; Spearman 0.985) — not a correctness-breaking shortcut; root-cluster fix (16 cells) from evolution.ipynb iter 3 improved accuracy, not speed
- **Reduced language/runtime overhead — minor.** Same `conda run -p CMAP Rscript` fixed cost as destiny; smaller relative share because the flood loop dominates R wall-clock

**Admissibility.** Class-C (stochastic / distributional equivalence): both implementations run n=30 stochastic floods with *different* RNG streams (R Mersenne-Twister vs NumPy PCG64), so per-cell outputs cannot be bit-identical by construction; equivalence is established on the aggregated pseudotime (Spearman 0.985 vs R). Crucially, the speedup comes from **vectorising the existing stochastic BFS** (batch Bernoulli sampling over a sliced transition sub-matrix), **not** from reducing the simulation count or substituting a deterministic surrogate — the statistical target is unchanged.

##### Contrast: where the speedup is NOT algorithmic

- **py-destiny's dominant lever is genuinely algorithmic-complexity** (dense full eigendecomposition → sparse truncated top-k), so its gain is durable and grows with N.
- **py-URD's dominant lever is implementation** (vectorisation of an interpreted loop); the algorithm is unchanged, so its gain reflects Python-vs-R execution efficiency plus the fixed invocation overhead, and would partially compress if R were called in-process via rpy2 rather than as a `conda run` subprocess.
- For both, **caching** (reusing the shared pydestiny diffusion map) is a real but secondary integration win, and **reduced runtime overhead** is inflated on small datasets by the conda/Rscript subprocess cost.

#### Attribution matrix (companion CSV/XLSX)

| Method | Python impl. | Sparse linalg | Caching | Algorithmic rewrite | Runtime overhead |
|---|---|---|---|---|---|
| py-destiny (~139×) | major | dominant | minor | moderate | moderate |
| py-URD (~68×) | dominant | major | moderate | minor | minor |

#### Other large-speedup methods (brief)

- **Slingshot (27×), TSCAN (16×), CytoTRACE (18×):** dominated by NumPy vectorisation + reduced R/subprocess overhead; modest sparse-graph/MST reuse. TSCAN's measured replicate CV (11.7%) and the documented cold-cache anomaly indicate part of its R/Py gap is invocation-overhead-sensitive.
- **SCORPIUS (5×):** the smallest gap and the highest Python replicate variance (rep-CV ~44%) — its principal-curve/MDS core is already NumPy-vectorised in both ecosystems, leaving mostly language overhead, hence the small and noisy speedup.

#### Reproducibility note

Speedups are from Supplementary Table 4_9 (Python ports re-measured with warm-up + 3 replicates + `tracemalloc` peak memory in omicdev; R references single-run via `conda run -p CMAP Rscript`). A subprocess-overhead-controlled re-run (R via rpy2 in-process, or `/usr/bin/time -v` accounting of the Rscript startup separately from compute) would isolate the durable compute-level speedup from the fixed invocation cost; this is the recommended follow-up for a definitive per-cause magnitude split.

#### *Supplementary Note 7 | Mathematical comparison: GPCCA vs PCCA+ vs the PseudotimeFate sparse approximation*

`ov.single.PseudotimeFate` (the upper-level design behind `omicverse.single.Traj`) unifies downstream fate analysis across pseudotime backends. Once any backend writes a pseudotime vector into `adata.obs[<key>]`, PseudotimeFate runs one backend-agnostic four-stage workflow: **(1)** pseudotime-biased kNN graph, **(2)** adaptive-Gaussian directed transition matrix `P` (row-normalised), **(3)** macrostate inference, and **(4)** fate-probability computation. Stage 3 is where it departs from CellRank's GPCCA stack. This note formalises the macrostate/fate mathematics and states exactly which approximations the implementation makes and why they are acceptable for a pseudotime-biased transition matrix. All claims are grounded in `omicverse/single/_pseudotime_fate.py`.

##### Common goal

All three methods take a row-stochastic transition matrix $P \in \mathbb{R}^{n\times n}$ over cells and seek `K` **metastable macrostates** whose self-transition (residency) is high, designate the late, high-residency ones as **terminal states**, and compute each transient cell's **absorption (fate) probability** into each terminal set. They couple terminal-state inference and fate estimation through a coarse-grained (Galerkin) transition matrix. The methods differ only in *how the metastable subspace is extracted and how soft memberships are assigned*.

##### GPCCA (Generalized PCCA+; pyGPCCA, CellRank)

GPCCA computes a **sorted real Schur decomposition** $P = QRQ^T$ and keeps the leading `K` Schur vectors `X = Q[:, :K]` spanning the dominant invariant subspace. Because it uses the *real Schur form* rather than eigenvectors, it is valid for **non-reversible** `P` (complex eigenvalues are kept in 2×2 real blocks). It then runs **PCCA+** on `X`: the inner-simplex algorithm (ISA) gives an initial rotation `A`, which is then **optimised** (Gauss–Newton on a crispness functional) subject to $\chi = X\cdot A \geq 0$ and row-sums = 1, maximising membership crispness. Terminal states are high-residency macrostates of the coarse-grained $P_c = x^T P x$; fate = absorption into those sets. Dominant cost: the sorted Schur decomposition plus the iterative crispness optimisation.

##### Classic PCCA+ (Röblitz & Weber 2013)

Identical to GPCCA except the metastable subspace is spanned by the **top-`K` dominant eigenvectors** of `P` rather than Schur vectors. This requires `P` to be **reversible** (real spectrum clustered near 1); for non-reversible chains the eigenvectors become complex and the simplex structure breaks. The ISA-plus-crispness-optimisation membership step is the same.

##### Implemented sparse approximation (PseudotimeFate)

PseudotimeFate keeps the PCCA+ *geometry* but replaces the two expensive pieces with sparse, deterministic surrogates (functions `_top_schur_vectors`, `_indexsearch`, `_pcca_plus_approx`, `_coarse_grain`, `_solve_fate_probabilities`):

1. **Schur basis → sparse top-K eigenvectors of $P^T$.** `spla.eigs(Pᵀ, k=K, which='LM')` (ARPACK Krylov) returns the leading `K` eigenvectors; complex-conjugate pairs are split into their real and imaginary parts, then Gram–Schmidt (QR) orthonormalised — a real basis for the metastable subspace. This avoids the dense `O(n³) scipy.linalg.schur` while keeping the macrostate signal. It is **exact for reversible P** and a **bounded approximation** otherwise; the pseudotime-biased kNN graph is near-reversible, so the surrogate spans essentially the same subspace as the true Schur basis.
2. **PCCA+ membership → bare ISA, no crispness optimisation.** `_indexsearch` greedily selects `K` simplex vertices (vertex 1 = max-‖·‖ row; vertex k = row furthest from the affine hull of the previous k−1, via running orthogonal projection, `O(nK²)`). The affine map `A = M⁻¹` (M = vertex rows) gives χ = X·A, clipped at 0 and row-renormalised. The implementation **deliberately skips** the `O(K³)` crispness-optimisation rotation that GPCCA/PCCA+ apply on top. Because terminal states are taken from the **argmax** of χ (not its crispness) and absorption is aggregated over **whole macrostate cell-sets**, the omitted optimisation changes soft membership *values* but not the discrete terminal-state *set* — the source notes terminal-state agreement comparable to GPCCA on the pancreas fixture.
3. **Coarse-graining and terminal selection.** Hard one-hot memberships `Z` give the Galerkin `Pᶜ = (ZᵀPZ)/ΣZ` (matches pyGPCCA's `coarse_T`); `residency = diag(Pᶜ)`. Terminal macrostates = those with mean-pseudotime rank ≥ `late_pt_quantile` **and** residency ≥ `residency_threshold`, excluding the initial state and deduped by dominant cluster.
4. **Fate = absorption probabilities.** `(I - Q)X = R` (Q = transient-to-transient block, R column-aggregated over each terminal cell-set) solved by Neumann-series power iteration — the same absorption target as CellRank/Palantir — yielding a smooth fate landscape plus per-cell `lineage_entropy = -Σ x log x`.

##### The three approximations, and why they hold here

- **Eigenvectors of $P^T$ instead of a sorted real Schur form.** Rigorous GPCCA uses Schur to handle strong non-reversibility; the bias-kNN transition matrix is nearly reversible, so the leading eigenvectors span the same metastable subspace (bounded approximation, exact in the reversible limit).

- **No crispness optimisation.** Affects soft-membership sharpness only; the discrete terminal-state set (argmax + residency/pseudotime gating) and the set-aggregated absorption are invariant to it.
- **Power-iteration absorption instead of coarse-grained fate.** Identical conceptual target (absorption into terminal sets); numerically equivalent.

##### Complexity and dependencies

GPCCA's dominant cost is a sorted real Schur decomposition (dense `O(n³)` or Krylov-Schur via SLEPc) plus iterative crispness optimisation, and it pulls in pyGPCCA (and CellRank's kernel stack). PseudotimeFate runs sparse ARPACK (`O(n·K·iters)`) + ISA (`O(n·K²)`) + a sparse absorption solve, in **pure numpy/scipy.sparse with no pyGPCCA, no jax, no CellRank** — the engineering trade that lets the same fate workflow sit on top of *any* pseudotime backend at low cost.

##### Comparison table

| Aspect | GPCCA (CellRank / pyGPCCA) | Classic PCCA+ (Röblitz–Weber 2013) | Implemented sparse approx. (PseudotimeFate) |
|---|---|---|---|
| Problem solved | Metastable/terminal macrostates + fate (absorption) probabilities from a row-stochastic transition matrix P | same | same |
| Spectral decomposition | Sorted real Schur decomposition of P (top-K invariant subspace) | Top-K dominant eigenvectors of P | Top-K right eigenvectors of $P^T$ via sparse ARPACK (spla.eigs, k=K, 'LM'), Gram–Schmidt; complex pairs split into (real, imag) |
| Reversibility requirement | None — real Schur handles non-reversible P | Requires reversible P / real spectrum near 1 | Exact for reversible P; bounded approximation for mildly non-reversible (bias-kNN is near-reversible) |

| Aspect | GPCCA (CellRank / pyGPCCA) | Classic PCCA+ (Röblitz–Weber 2013) | Implemented sparse approx. (PseudotimeFate) |
|---|---|---|---|
| Soft membership (χ) | ISA initial guess + crispness optimization of rotation A (Gauss–Newton) | ISA + crispness optimization of A | ISA only (_indexsearch): A = $M^{-1}$, χ = X·A, clip<0, renormalize — NO crispness optimization |
| Determinism | Optimization may use restarts | Optimization may use restarts | Fully deterministic, single pass, no restart |
| Coarse-graining | Galerkin P_c = $\chi^T P \chi$ (normalized) | Galerkin | Galerkin $P^c$ = $(Z^T P Z)/\Sigma Z$ on hard one-hot Z (matches pyGPCCA coarse_T) |
| Terminal-state selection | High self-residency macrostates from P_c | — | late-pseudotime (≥ quantile) AND residency ≥ threshold, deduped by dominant cluster |
| Fate probabilities | Absorption into terminal macrostate sets | Absorption | (I−Q)X = R via Neumann-series power iteration; R column-aggregated over each terminal cell-set; + lineage entropy |
| Dominant cost | Sorted real Schur (dense $O(n^3)$ or Krylov-Schur) + iterative A-optimization | $O(n^3)$ eig + A-optimization | Sparse ARPACK O(n·K·iters) + ISA $O(n \cdot K^2)$ + sparse solve — no dense $O(n^3)$, no pyGPCCA/jax |
| Implementation | pyGPCCA + CellRank (Schur via SLEPc/scipy; optimization) | Roeblitz & Weber 2013 | omicverse single/_pseudotime_fate.py — pure numpy/scipy.sparse |

##### Caveat

The eigenvector-for-Schur substitution is a **bounded approximation that degrades for strongly non-reversible transition matrices** — if a backend or biasing scheme produces a markedly non-reversible `P` (large directed cycles), the leading `Pᵀ` eigenvectors can become complex-dominated and the real/imag split no longer cleanly spans the metastable subspace. For such cases the rigorous real-Schur GPCCA remains the correct choice; PseudotimeFate's regime of validity is the near-reversible, pseudotime-biased kNN graph it is built for.

#### *Supplementary Note 8 | PseudotimeFate vs CellRank GPCCA — standardized runtime & terminal-state comparison*

S5 states that PseudotimeFate completes in under one second on the pancreas dataset whereas the comparable CellRank pseudotime-kernel + GPCCA workflow takes tens of seconds to minutes. This benchmark makes that claim reproducible: both estimators receive **identical input** (the same `AnnData`, the same Palantir pseudotime column, the same kNN graph) on the **same hardware and dependency stack**, are timed with the **same warm-up + replicate policy**, and are evaluated on the **same terminal-state ground truth**. The executed notebook is `trajectory_compare.ipynb` (all cells run, outputs retained).

##### Result

| Method | Runtime (s, warm, 3 rep) | Terminal states | Precision | Recall | Macrostate backend |
|---|---|---|---|---|---|
| ov.single.PseudotimeFate | 0.392 ± 0.004 | 4 (Alpha, Beta, Delta, Epsilon) | 1.00 | 1.00 | pure numpy/scipy; ARPACK eigs($P^T$) + ISA |
| CellRank PseudotimeKernel + GPCCA | 33.17 ± 3.66 | 8 → 5 clusters (Beta, Delta, Epsilon + Ductal, Ngn3) | 0.60 | 0.75 | dense brandts Schur (petsc/slepc absent) + pyGPCCA |

**`ov.single.PseudotimeFate` is 84.6× faster** (0.392 s vs 33.17 s) **and recovers the terminal states more cleanly** — all four mature endocrine fates (Alpha, Beta, Delta, Epsilon) at precision = recall = 1.0, versus CellRank's out-of-the-box `predict_terminal_states`, which recovers three (misses Alpha) and additionally flags early Ductal / Ngn3-high macrostates as terminal (precision 0.60, recall 0.75).

##### Standardization (what makes this a fair comparison)

- **Identical input.** Both methods read the same `adata` (3,696 cells × 3,000 HVGs), the same `palantir_pseudotime`, and the same `adata.obsp['connectivities']` kNN graph. Neither recomputes the embedding, neighbours, or pseudotime.
- **Identical macrostate budget.** Both are asked for `n=10` macrostates with `clusters` as the cluster annotation.
- **Warm-up + replicates.** One untimed warm-up call per method is discarded (it excludes the one-off JIT/ARPACK warm-up for PseudotimeFate and any first-call caching for CellRank), then 3 timed replicates (`time.perf_counter`) give mean ± SD.
- **Same hardware & deps.** Intel Xeon Platinum 8462Y+ (64 cores), Python 3.10.20, omicverse 2.2.1rc1, cellrank 2.0.7, pygpcca 1.0.4, scipy 1.15.3, numpy 2.2.6 — one conda environment (omicdev).
- **Same terminal-state evaluation.** Recovered/missed/spurious are scored against the biological ground truth {Alpha, Beta, Delta, Epsilon}; CellRank's `<cluster>_<k>` terminal labels are mapped to clusters by stripping the numeric suffix.

##### Why the gap exists

CellRank's cost is dominated by the **real Schur decomposition** inside GPCCA. In this environment `petsc4py`/`slepc4py` are absent, so pyGPCCA falls back to the dense `brandts` method (it densifies the 3,696×3,696 transition matrix), which is the bulk of the 33 s. PseudotimeFate replaces that step with a **sparse ARPACK `eigs(Pᵀ, k=10)`** surrogate of the Schur basis plus the deterministic inner-simplex (ISA) membership assignment — `O(n·K²)`, sub-second — and computes fate by a sparse absorption solve (see the accompanying Supplementary Note on GPCCA vs PCCA+).

##### Caveats (so the claim is not overstated)

- **Environment-dependent magnitude.** The 84.6× gap is specific to the common *no-petsc/slepc* install. With `petsc4py`+`slepc4py` present, CellRank uses a sparse Krylov–Schur method and the absolute CellRank time drops substantially; the gap would narrow (though PseudotimeFate's lighter eigs+ISA path would still be faster). The figure reports tens of seconds here; on a petsc-enabled machine CellRank is closer to a few seconds.
- **Out-of-the-box terminal states.** CellRank's precision/recall reflect its **default** `predict_terminal_states()` (automatic state count via the eigengap/stability). Manually fixing the number of terminal states to 4 or applying stability filtering would improve CellRank's terminal-state recovery; this benchmark deliberately compares default pipelines, not hand-tuned ones.
- **Cold-start.** PseudotimeFate's *first* (cold) call is ~12 s (JIT/ARPACK/scipy warm-up); the sub-second number is the warm, steady-state cost relevant to an agent that calls it repeatedly. CellRank's cost is compute-bound (dense Schur), so warm-up does not reduce it.

- **Single dataset.** This is the pancreas fixture only (n=3,696); the per-dataset trajectory-method benchmark (Supplementary Tables 4_7–4_9) covers the broader runtime picture.

##### Reproduce

Run `trajectory_compare.ipynb` in the omicdev kernel: load `data/pancreas_with_palantir.h5ad`, then the two timed blocks and the runtime/terminal-state cells. Total execution ≈ 2–3 min (dominated by the 4 CellRank GPCCA runs). The runtime bar chart is written to `figures/trajectory_compare_runtime.png`.

.

### Supplementary Tables

#### *Supplementary Table 1*

OmicVerse module-by-pipeline coverage, method counts, representative complete workflows, and input/output object types. Rows follow the Fig. 1a / S4 narrative order (assay processing → preprocessing → single-cell → spatial → bulk → foundation-model → molecular → epigenomic → microbiome → metabolomics), followed by cross-cutting / support modules.

##### Table 1a | Main table

| Namespace | Modality | Package | # Methods | Input objects | Output objects | Representative complete workflow | Key pipeline stages |
|---|---|---|---|---|---|---|---|
| ov.alignment | Assay processing (FASTQ/FASTA/BAM alignment & quantification) | omicverse/alignment/ | 32 | FASTQ (single/paired), FASTA (transcriptome/genome/ASV), SRA accessions, BAM, GTF/GFF, SINTAX/RDP/SILVA refs | AnnData (sparse CSR counts), FASTQ (filtered), BAM, count matrices (TSV/HDF5/MTX), Newick tree, aligned FASTA, taxonomy/OTU/ASV tables | SRA → FASTQ → QC (fastp) → STAR alignment → featureCounts quantification (bulk_rnaseq_pipeline) | Data acquisition, QC/preprocessing, alignment, quantification, ASV inference, taxonomy, count-matrix generation, phylogenetics |
| ov.pp | Preprocessing | omicverse/pp/ | 31 | AnnData, numpy.ndarray, scipy.sparse CSR | AnnData (in-place embeddings/clusters/QC), numpy.ndarray, boolean filter masks | QC → filter_cells → filter_genes → preprocess → scale → pca → neighbors → leiden → umap | QC, preprocessing, normalization, HVG selection, scaling, regression, dimensionality reduction, neighborhood, clustering, ambient-RNA removal, doublet detection |
| ov.single | Single-cell (scRNA-seq, scATAC-seq, scProteomics/EVs) | omicverse/single/ | 151 | AnnData, MuData, pandas.DataFrame, h5ad, FASTQ (sub-workflows), cell-gene count matrices | AnnData (annotated), obsm embeddings, obs metadata, uns analysis results, DataFrame, figures | Spliced/unspliced → Velo.filter_genes() → Velo.cal_velocity() (dynamo/scvelo/regvelo) → velocity_embedding → cellrank_fate | QC, preprocessing, dim. reduction, clustering, annotation, trajectory, RNA velocity, gene regulation, cell-cell communication, pathway, metabolic, CNV, multi-omics, perturbation, EV proteomics |
| ov.space | Spatial | omicverse/space/ | 48 | AnnData (spatial), WSI images (.svs/.ndpi), H&E histology, spatial coords (obsm['spatial']), expression matrices | AnnData (annotated), obsm embeddings, cell-type proportion matrices, var gene rankings, tissue-zone activations, SVG flags, neighborhood graphs | QC + spatial_neighbors → svg (PROST/pearsonr) → pySTAGATE cluster → visualize spatial domains | QC, preprocessing, spatial-graph construction, clustering, SVG detection, deconvolution, integration, trajectory, tissue-zoning, histology embedding |
| ov.bulk | Bulk RNA-seq | omicverse/bulk/ | 25 | pandas.DataFrame (counts/ranked | DataFrame (DEG/enrichment/W | pyDEG(raw_counts) → | QC, normalization, DE analysis, gene-module |

| | | | | | | | |
|---|---|---|---|---|---|---|---|
| | | | | genes), AnnData, gene-symbol lists, file paths/organism codes | GCNA modules), AnnData (in-place corrected layers), matplotlib figures, NetworkX graph, pandas Series | drop_duplicates_index → normalize → deg_analysis → foldchange_set → plot_volcano | detection, enrichment, network analysis, survival analysis, deconvolution, annotation |
| ov.llm (ov.fm) | Single-cell foundation models | omicverse/llm/ | 38 | AnnData (.X expression), model-checkpoint path, optional .obs batch/cell-type labels | AnnData (.obsm['X_<model>']), numpy.ndarray embeddings, dict (predictions / fine-tuning / integration results) | end_to_end_scgpt_annotation(reference, query, model_path) → fine-tune on reference + predict on query | Annotation, embedding, integration, fine-tuning, perturbation prediction |
| ov.mol | Molecular | omicverse/mol/ | 12 | gene symbol, UniProt accession, PDB ID, AA sequence, SMILES, drug name, RDKit Mol, molecular files, MolStructure | MolStructure, py3Dmol.view, matplotlib PAE axes, pandas.DataFrame (pockets/drugs), DockingResult, dict (druggability), PDB path | fetch_structure('EGFR') → pockets(s) → dock(s,'gefitinib',pocket=1) → view_docking(s,result) | Structure acquisition, visualization, confidence assessment, pocket detection, druggability scoring, drug lookup, ligand prep, docking, validation |
| ov.epi (scattered; no top-level namespace) | Epigenomics (ATAC/regulatory) | omicverse/single/_atac.py + external modules | 36 | AnnData (ATAC-seq), pandas.DataFrame (peak/region tables), genomic regions (chr:start-end), variant IDs (rsid/cCRE) | AnnData (merged ATAC), pandas.DataFrame (TF sites/cCRE/regulatory variants), dict (regulatory metadata) | genomic region → CCREClient.region_to_ccre_screen → cCRE ID + RegulomeDBClient.query_variant → regulatory annotation | QC, preprocessing, feature selection, accessibility quantification, TF annotation, regulatory-element annotation, integration |
| ov.micro | Microbiome (16S/amplicon metagenomics) | omicverse/micro/ | 41 | AnnData (samples × ASVs/OTUs, int counts), Newick tree, pandas.DataFrame, paired AnnData tables | AnnData, pandas.DataFrame (distance/diversity/DA), numpy.ndarray (ordination), embedding/cooccurrence tables | QC → rarefy → Alpha → Beta → Ordinate (PCoA) → scanpy.pl | QC/filtering, normalization, alpha diversity, beta diversity, ordination, differential abundance, phylogenetics, meta-analysis, multi-omics integration |
| ov.metabol | Metabolomics & Lipidomics | omicverse/metabol/ | 64 | AnnData (samples × metabolites), CSV/TSV peak tables (MetaboAnalyst/wide), LC-MS feature tables (m/z + RT) | AnnData (transformed/normalized), pandas.DataFrame (stats/VIP/ROC), PLSDA/ASCA/BiomarkerPanel results, matplotlib axes | read_metaboanalyst → pyMetabo.impute(qrilc) → normalize(pqn) → transform(log) → differential(welch_t) → opls_da(1) | I/O, QC, batch/drift correction, imputation, normalization, transformation, univariate, multivariate, multi-factor designs, biomarker discovery, correlation, MOFA, pathway enrichment, lipidomics, mass annotation, visualization |
| ov.es | Gene-set / enrichment scoring | omicverse/es/ | 16 | AnnData, pandas.DataFrame, numpy.ndarray | DataFrame (cells × signature scores), DataFrame (p-values), AnnData.obsm keys | decouple(adata, signatures, methods=['aucell','gsea','ulm'], cons=True) → dict with consensus | Enrichment scoring, pathway activity, transcriptional regulation, consensus integration, enrichment testing |
| ov.genetics | Statistical genetics / GWAS | omicverse/genetics/ | 59 | AnnData (samples×SNPs, cells×genes, genes×samples), PLINK (.bed/.bim/.fam), VCF, GWAS sumstats (TSV/CSV), DataFrames/arrays, Z-scores, LD matrices, TWAS models, LD-score filesets | AnnData (QC'd genotype), DataFrame (association/clumping/QC), Series (LD $r^2$), backend objects (SuSiE/ColocABF/MRInput/Hsq), GWASStudy, PC tuples, figures | gwas_qc(adata) → gwas_association(geno, pheno) → finemap(z, R, method='susie_rss') → get_credible_sets | I/O, QC, association, fine-mapping, locus definition, colocalization, Mendelian randomization, single-cell disease relevance, heritability, genetic correlation, TWAS, visualization, synthetic data, spatial GWAS |
| ov.airr | Immune repertoire (AIRR, TCR/BCR) | omicverse/airr/ | 102 | AnnData (per-cell TCR/BCR), AIRR-format TSV, 10x Cell Ranger outputs, pandas.DataFrame (contigs), pyimmunarch.ImmunData (bulk) | AnnData (single-cell), pandas.DataFrame (metrics), numpy.ndarray (distances/coords), matplotlib.Axes, annotated DataFrames (clones/SHM/phylo) | read_10x_vdj → chain_qc → define_clonotypes → clonal_expansion | I/O, QC, clonotype definition, repertoire metrics, visualization, bulk analysis, B-cell/Ig analysis, TCR specificity, TCR-GEX integration |
| ov.external | Integrated third-party tools (multi-domain) | omicverse/external/ | 39 | AnnData, MuData, pandas.DataFrame, numpy.ndarray, GTF/GFF, expression matrices, protein IDs, SMILES | AnnData (.obsm/.varm/.uns results), pandas.DataFrame, networkx.Graph, trajectory/pseudotime objects, factor matrices, figures, .obs annotations | scanorama.correct() → harmony.run_harmony() for batch correction & integration | QC, preprocessing, batch correction, integration, normalization, feature selection, dim. reduction, clustering, trajectory, annotation, differential testing, |

| | | | | | | | |
|---|---|---|---|---|---|---|---|
| | | | | | | | network analysis, spatial, visualization |

#### Table 1b | Modality × pipeline-stage coverage matrix

| Module | Assay/IO | Preproc/QC | Clustering/Embedding | Annotation | Trajectory/Dynamics | Differential/Statistics | Integration/Batch | Deconvolution | Enrichment/Pathway | Foundation-model | Visualization |
|---|---|---|---|---|---|---|---|---|---|---|---|
| ov.alignment | ✓ | ✓ | | ✓ | | | | | | | |
| ov.pp | | ✓ | ✓ | | | | | | | | ✓ |
| ov.single | ✓ | ✓ | ✓ | ✓ | ✓ | ✓ | ✓ | ✓ | ✓ | | ✓ |
| ov.space | ✓ | ✓ | ✓ | ✓ | ✓ | ✓ | ✓ | ✓ | | | ✓ |
| ov.bulk | ✓ | ✓ | | ✓ | | ✓ | ✓ | ✓ | ✓ | | ✓ |
| ov.llm (ov.fm) | | ✓ | ✓ | ✓ | | | ✓ | | | ✓ | |
| ov.mol | ✓ | | | | | | | | | | ✓ |
| ov.epi | ✓ | ✓ | | ✓ | ✓ | | ✓ | | | | |
| ov.micro | ✓ | ✓ | ✓ | ✓ | | ✓ | ✓ | | | | ✓ |
| ov.metabol | ✓ | ✓ | ✓ | ✓ | | ✓ | ✓ | | ✓ | | ✓ |
| ov.es | | | | | | ✓ | | | ✓ | | |
| ov.genetics | ✓ | ✓ | | ✓ | | ✓ | | | ✓ | | ✓ |
| ov.airr | ✓ | ✓ | ✓ | ✓ | | ✓ | ✓ | | | | ✓ |
| ov.external | ✓ | ✓ | ✓ | ✓ | ✓ | ✓ | ✓ | | ✓ | | ✓ |

#### Summary statistics

- **Total methods across all modules:** 694
- **Number of modules surveyed:** 14
- **Core S4-narrative modules (10):** ov.alignment, ov.pp, ov.single, ov.space, ov.bulk, ov.llm/ov.fm, ov.mol, ov.epi, ov.micro, ov.metabol — 478 methods combined
- **Cross-cutting / support modules (4):** ov.es, ov.genetics, ov.airr, ov.external — 216 methods combined
- **Largest modules by method count:** ov.single (151), ov.airr (102), ov.metabol (64), ov.genetics (59), ov.space (48)
- **Smallest module:** ov.mol (12)
- **Integrated third-party tools (via ov.external):** 39 publicly exported tools/packages (~30 additional internal/experimental tools), spanning 29+ biological databases via datacollect
- **Foundation models accessible (ov.llm):** 21+ single-cell foundation models (6 fully integrated with local runtimes — scGPT, Geneformer, scFoundation, CellPLM, UCE, scMulan; 15+ registered as stubs pending upstream runtime integration)
- **Primary shared data container:** AnnData (with MuData for multi-omics; pandas.DataFrame for tabular/statistical-genetics and bulk inputs)

### *Supplementary Table 2 | R/Bioconductor-to-Python reconstruction map*

Pure-Python (and Rust) re-implementations of R / Bioconductor / CRAN tools authored under github.com/omicverse, with original source ecosystem, numerical parity status, availability within the OmicVerse package, and the importing ov.* function(s). Rows grouped by analysis domain.

| Reconstructed package | Original R tool | Source ecosystem | Domain | Parity status | Availability in OmicVerse | Importing ov function(s) | GitHub |
|---|---|---|---|---|---|---|---|
| **Bulk RNA-seq differential expression & expression** | | | | | | | |
| py-edgeR | edgeR (Robinson, McCarthy & Smyth, Bioinformatics 2010) | Bioconductor | bulk RNA-seq differential expression (negative-binomial GLM / quasi-likelihood F-test for count data) | stated parity / unquantified ('R-parity tested against Bioconductor edgeR'; no numeric metric) | vendored-external | ov.bulk.pyDEG.deg_analysis(method='edger') | github.com/omicverse/py-edgeR |
| py-limma | limma (Ritchie et al., Nucleic Acids Research 2015; | Bioconductor | bulk DE — linear models & empirical-Bayes | stated parity / unquantified (faithful port of limma core; | vendored-external | ov.bulk.pyDEG (limma/limma-voom backend) | github.com/omicverse/py-limma |

| | | | | | | | |
|---|---|---|---|---|---|---|---|
| | eBayes from Smyth 2004) | | moderated differential expression (voom/lmFit/eBayes/ topTable) | powers omicverse edgeR/limma-voom and pydeqms backends; no verbatim numeric metric) | | | |
| py-mfuzz | Mfuzz (Futschik & Carlisle 2005; Kumar & Futschik 2007); fuzzy c-means core ported from e1071 cmeans | Bioconductor | bulk/time-series gene expression — fuzzy c-means soft clustering of temporal trajectories | membership Pearson r = 1.0, centres r = 1.0, hard-assignment ARI ~0.99; fill_NA(knn) bit-exact (max abs diff ~1e-15); Dmin curve Pearson r = 1.0 vs R (random init caveat) | pip-dependency | ov.bulk.temporal_clusters | github.com/omicverse/py-mfuzz |
| **Single-cell trajectory inference** | | | | | | | |
| py-monocle2 | Monocle 2 (Qiu et al., Nature Methods 2017) | Bioconductor | single-cell trajectory inference (DDRTree pseudotime) | Pseudotime correlation with R Monocle 2 >= 0.99 on every benchmark dataset; 30-100x faster than R | vendored-external | ov.single.Monocle | github.com/omicverse/py-monocle2 |
| py-Monocle3 | monocle3 (Cao et al., Nature 2019; Trapnell et al., Nat Biotechnol 2014) | Bioconductor | single-cell trajectory inference (principal-graph pseudotime) | stated parity / unquantified (wrapper re-exporting Bio-Babel/Monocle3-python; parity report deferred to upstream Bio-Babel, no numeric metric in this repo) | pip-dependency | ov.single.TrajInfer | github.com/omicverse/py-Monocle3 |
| py-Slingshot | slingshot (Street et al., BMC Genomics 2018) | Bioconductor | trajectory — cell-lineage (MST/principal-curve) + pseudotime inference for scRNA-seq | pseudotime Spearman 0.99+ vs R Slingshot on a Y-shaped fixture (matched lineages) | vendored-external | ov.single.TrajInfer(method='slingshot') | github.com/omicverse/py-Slingshot |
| py-TSCAN | TSCAN (Ji & Ji, Nucleic Acids Research 2016) | Bioconductor | trajectory — mclust+MST+spline pseudotime reconstruction for scRNA-seq | Pearson = 1.000000 vs R pseudotime in verification mode (when Mclust clusters match) | pip-dependency | ov.single.TrajInfer (inference(method='TSCAN')) | github.com/omicverse/py-TSCAN |
| py-SCORPIUS | SCORPIUS (Cannoodt et al., bioRxiv 2016) | R-dynverse (rcannood/SCORPIUS) | trajectory — linear trajectory inference for scRNA-seq via MDS + Hastie-Stuetzle principal curves | "Pseudotime Pearson = 0.989 vs R SCORPIUS"; "Low-dim space Procrustes = 0.999 vs R" | pip-dependency | ov.single.TrajInfer(method='scorpius') | github.com/omicverse/py-SCORPIUS |
| py-destiny | destiny (Angerer et al., Bioinformatics 2016; theislab/destiny) | Bioconductor | trajectory inference — diffusion maps + diffusion pseudotime (DPT) for scRNA-seq | Eigenvectors Procrustes 0.916 (thr 0.80); Eigenvalues Pearson 0.967 (thr 0.95); DPT Pearson 0.974 (thr 0.85) vs R | both | ov.single Traj method='destiny'; ov.single Traj method='urd' | github.com/omicverse/py-destiny |
| py-URD | URD (Farrell et al., Science 2018; farrellja/URD) | R (other) | trajectory — branching pseudotime (flood-based) + random-walk lineage attribution for scRNA-seq | "Pseudotime parity vs R URD on Guo qPCR data (428 cells): Spearman 0.985, Pearson 0.992" | pip-dependency | ov.single.TrajInfer (method='urd') | github.com/omicverse/py-URD |
| py-CytoTRACE | CytoTRACE (Gulati et al., Science 2020) | R (other) | trajectory — gene-count-based stemness / developmental-potential pseudotime for scRNA-seq | stated parity / unquantified (function map marks ported functions as ported, no numerical metric in README) | pip-dependency | ov.single.TrajInfer (method='cytotrace') | github.com/omicverse/py-CytoTRACE |
| py-tradeSeq | tradeSeq (Van den Berge et al., Nature Communications 11:1201, 2020) | Bioconductor | trajectory — trajectory-based differential expression for scRNA-seq via NB-GAM (associationTest, startVsEndTest, patternTest) | associationTest Spearman = 0.84 on -log10 p (top-50 Jaccard 0.67); startVsEndTest Spearman 0.45 (approximate) | standalone-only | — (not wired into omicverse) | github.com/omicverse/py-tradeSeq |
| py-condiments | condiments (Roux de Bezieux et al., Nature Communications 2024) | R (other) | trajectory — differential trajectory analysis across conditions in scRNA-seq | imbalance_score Pearson = 1.0000 vs R (topologyTest simplified/approximate in v0.1) | standalone-only | — (not wired into omicverse) | github.com/omicverse/py-condiments |
| py-dynwrap | dynwrap (dynverse; Saelens et al. 2019) | R/dynverse | trajectory inference data wrapper / benchmarking (geodesic distances, milestone networks for dyneval/dynbenchmark) | byte-equivalent geodesic distances (max\|Δ\| = 0.0 on canonical fixture) | standalone-only | — (not wired into omicverse) | github.com/omicverse/py-dynwrap |
| py-dyneval | dyneval (Saelens et al., Nat Biotechnol 2019) | R/dynverse | trajectory inference benchmark metrics | R parity on canonical 3-branch/30-cell fixture: correlation 0.9812 R vs 0.9813 Py (\|Δ\|0.0001); him/edge_flip/isomor | standalone-only | — (not wired into omicverse) | github.com/omicverse/py-dyneval |

| | | | | | | | |
|---|---|---|---|---|---|---|---|
| | | | | phic/F1_branches \|Δ\|=0.0; F1_milestones 0.6667 both; featureimp_cor 0.9622 vs 0.9005 (\|Δ\|0.0617); featureimp_wcor 0.9664 vs 0.9548 (\|Δ\|0.0115). 8/14 dyneval metric IDs ported in v0.1 | | | |
| py-dynbenchmark | dynbenchmark (Saelens et al., Nat Biotechnol 2019) | R/dynverse | trajectory inference benchmarking (dyneval metrics + synthetic generators + harness) | stated parity / unquantified ('Python implementation of dyneval metrics + synthetic generators + benchmark harness'; no numeric metric) | standalone-only | — (not wired into omicverse) | github.com/omicverse/py-dynbenchmark |
| **Single-cell integration, networks, factorization & metacells** | | | | | | | |
| py-cca | Seurat RunCCA (Stuart, Butler, Hoffman, Hafemeister et al., Cell 2019) | R (other) | single-cell batch integration (canonical correlation analysis) | Numerical parity with Seurat::RunCCA across 9 (size x num_cc) configs: singular values match to ~1e-7, subspaces match to ~1e-3 (rotation within near-degenerate eigenspaces only source of difference) | pip-dependency (undeclared optional runtime import; not in pyproject) | ov.single.batch_correction (methods='cca') | github.com/omicverse/py-cca |
| py-Augur | Augur (Skinnider et al., Nature Communications 2021) | R (other) | single-cell cell-type prioritization (AUC-based ranking of perturbation-responsive cell types) | AUC ranking perfectly preserved (Spearman rho = 1.0), Pearson r = 0.9999 on benchmark datasets | standalone-only | — (not wired into omicverse) | github.com/omicverse/py-Augur |
| py-hdWGCNA | hdWGCNA (Morabito et al., Cell Reports Methods 2023) | R (other) | single-cell co-expression network analysis (WGCNA on scRNA-seq) | "SFT R^2, kME, and hMEs Pearson r = 1.0000 on benchmark datasets" | standalone-only | — (not wired into omicverse) | github.com/omicverse/py-hdWGCNA |
| py-SigXTalk | SigXTalk — deep-learning (HGNN) pathway-crosstalk framework (no first-author/year in README) | R (other) | single-cell — cell-cell communication / signaling pathway crosstalk (PRS) inference via hypergraph neural network | Pearson Correlation: 0.9912 (Python vs R, PBMC3k dataset) | standalone-only | — (not wired into omicverse) | github.com/omicverse/py-SigXTalk |
| rust-NMF | NMF (Gaujoux & Seoighe 2010; algorithms brunet/lee/offset/nsNMF/snmf-r/snmf-l, plus HALS/E-HALS/dnmf) | CRAN | single-cell gene-program / matrix factorization (NMF) for scRNA-seq; dimensionality reduction & module identification | bit-equivalent to R NMF within f64 round-off (max\|ΔW\| < 1e-12) for brunet/lee/offset/nsNMF/snmf-r/snmf-l; ~76x R 'lee' at 16 threads, up to ~218-280x with HALS+NNDSVD on pbmc8k; ARI 0.89 vs cell types (lee+NNDSVD) | pip-dependency (optional 'nmf-rs', not declared in pyproject) | ov.single.NMF | github.com/omicverse/rust-NMF |
| supercell | SuperCell (Bilous et al., BMC Bioinformatics 2022) | R (other) | single-cell metacell construction / coarse-graining | stated parity / unquantified | vendored-external | ov.single.MetaCell | —, external-only |
| scTenifold | scTenifoldNet / scTenifoldKnk (Osorio et al., 2020/2022) | R (other) | single-cell gene-regulatory network comparison & virtual knockout (perturbation) | unknown | vendored-external | ov.single.scTenifoldKnk | —, external-only |
| **Single-cell metabolism** | | | | | | | |
| py-scmetabolism | scMetabolism (Wu et al., Cancer Discovery 2021) | R (other) | single-cell metabolism — metabolic pathway-activity scoring (AUCell/VISION/ssGSEA/GSVA over KEGG/REACTOME) | "Correlation with R scMetabolism >= 0.99 for most methods"; also "mathematically equivalent" and "3-45x faster than R" | vendored-external | ov.single.Metabolism(method='scmetabolism') | github.com/omicverse/py-scmetabolism |
| **Single-cell differential abundance & splicing** | | | | | | | |
| py-miloR | Milo / miloR (Dann et al., Nature Biotechnology 2022) | Bioconductor | single-cell differential abundance testing on kNN neighbourhoods | stated parity / unquantified ('Same API as R miloR'; TMM normalisation and QL-F GLM said to match miloR's edgeR step, no numeric correlation quoted) | vendored-external | ov.single.Milo | github.com/omicverse/py-miloR |
| py-MARVEL | MARVEL (Wen et al., NAR Genomics Bioinformatics 2023) | R (other) | single-cell alternative splicing (PSI quantification, | Min Pearson r 0.9999961 (variable_splicing), most artifacts Pearson | standalone-only | — (not wired into omicverse) | github.com/omicverse/py-MARVEL |

| | | | | | | | |
|---|---|---|---|---|---|---|---|
| | | | differential splicing, isoform switching) | r ~1.0 with max abs diff 6.66e-16 to 8.91e-4 (reference fixtures / external replay benchmarks vs R MARVEL) | | | |
| **Doublet detection & ambient-RNA QC (single-cell)** | | | | | | | |
| py-scDblFinder | scDblFinder (Germain et al., F1000Research 2022) | Bioconductor | doublet detection (classifier-based; cxds → kNN features → xgboost iterative scoring → thresholding) for scRNA-seq | stated parity / unquantified ('mirrors R one-to-one'; per-primitive + end-to-end smoke tests, no numeric metric) | pip-dependency | ov.pp.qc (doublets_method='scdblfinder'); ov.pp.scdblfinder | github.com/omicverse/py-scDblFinder |
| py-DoubletFinder | DoubletFinder (McGinnis et al., Cell Systems 2019) | R (other) | doublet detection (scRNA-seq) | bit-for-bit reproducibility against R reference when fed matching PCA embeddings + artificial-doublet cell pairs; pANN/BCreal/classification bit-for-bit | both (in-tree wrapper vendored; underlying port undeclared optional pip dep) | ov.pp.qc; ov.pp.qc_cpu_gpu_mixed | github.com/omicverse/py-DoubletFinder |
| py-decontx | DecontX / decontX (Yang et al., Genome Biology 2020; celda Bioconductor) | Bioconductor | single-cell RNA-seq ambient/cross-contaminating RNA decontamination | contamination fraction, decontaminated matrix and per-cell theta agree at Pearson r > 0.99 vs Bioconductor reference | both | ov.pp.ambient.remove_ambient(method='decontx'); ov.pp.ambient.estimate_contamination(method='decontx') | github.com/omicverse/py-decontx |
| py-soupx | SoupX (Young & Behjati, GigaScience 2020, 9(12):giaa151) | R (other) | single-cell QC — ambient ('soup') RNA contamination removal from droplet scRNA-seq | bit-for-bit reproducibility against the R reference on the deterministic kernels — qualitative, no numeric coefficient | pip-dependency | ov.pp.remove_ambient (method='soupx'); ov.pp.estimate_contamination | github.com/omicverse/py-soupx |
| py-fastcar | FastCAR (Gocht/Berg et al., BMC Genomics 2023) | R (other) | single-cell ambient-RNA removal / QC (deterministic per-gene ambient-RNA correction for droplet scRNA-seq) | bit-for-bit identical to R FastCAR (deterministic, no RNG) | pip-dependency | ov.pp.ambient.remove_ambient(method='fastcar') | github.com/omicverse/py-fastcar |
| py-sccdc | scCDC (Wang et al., Genome Biology 2024) | R (other) | single-cell ambient-RNA decontamination (entropy-based gene-specific GCG detection and correction for scRNA-seq/snRNA-seq) | bit-for-bit reproducibility vs R reference for the deterministic core — per-gene/per-cluster entropy and corrected count matrix match scCDC exactly | pip-dependency | ov.pp.ambient.remove_ambient (method='sccdc') | github.com/omicverse/py-sccdc |
| **Single-cell CNV inference** | | | | | | | |
| py-CopyKAT | CopyKAT (Gao et al., Nat Biotechnol 2021) | R (other) | single-cell CNV — aneuploid/diploid classification and bin-level CNA calling from scRNA-seq | 17-patient 5-cancer benchmark: median py-vs-R ARI 0.988, median 23.4x faster on 8 cores | pip-dependency (lazy import; not a hard pyproject dep) | ov.single.CNV | github.com/omicverse/py-CopyKAT |
| py-inferCNV | inferCNV (Broad Institute / Tickle, Tirosh et al.) | Broad/R | single-cell CNV inference from scRNA-seq | Spearman rho = 1.0000 on continuous post-Phase-1 CNV matrix (3 10x patients), 0.9998 on smart-seq2 fixture; Pearson 0.993-0.999; i3 HMM Jaccard 1.000, i6 Jaccard 0.979; several steps bit-exact (max_diff < 1e-10) | standalone-only (optional runtime import only; not pip dep, not vendored) | ov.single.CNV | github.com/omicverse/py-inferCNV |
| **Immune repertoire (AIRR-seq)** | | | | | | | |
| py-alakazam | alakazam (Gupta et al., Bioinformatics 2015) | R/Immcantation | immune repertoire (AIRR-seq: Hill diversity, gene usage, clonal abundance, CDR3 AA properties, BCR lineage) | stated parity / unquantified — 'aims for numerical parity with alakazam 1.4.3' (qualitative, no numeric metric) | pip-dependency | ov.airr.hill_diversity; ov.airr.aa_properties; ov.airr.clonal_abundance; ov.airr.bcr_gene_usage | github.com/omicverse/py-alakazam |
| py-shazam | shazam (Gupta, Vander Heiden et al., Bioinformatics 2015; CRAN shazam 1.3.2) | R/Immcantation (CRAN) | immune repertoire — B-cell somatic hypermutation (SHM) targeting & BASELINe antigen-driven selection analysis | stated parity / unquantified — numerical parity with shazam 1.3.2 is the design goal, no numeric metric | pip-dependency | ov.airr.distance_threshold; ov.airr.clonal_clustering; ov.airr.mutation_analysis; ov.airr.shm_targeting; ov.airr.baseline_selection | github.com/omicverse/py-shazam |

| | | | | | | | |
|---|---|---|---|---|---|---|---|
| py-scoper | scoper (Immcantation, Kleinstein Lab; CRAN scoper) | R/Immcantation (CRAN) | immune repertoire — B-cell clonal clustering/partitioning of AIRR-seq BCR data (identical / hierarchical / spectral) | stated parity / unquantified (methods re-implemented 'faithfully'; no numeric metric) | pip-dependency | ov.airr.clonal_clustering | github.com/omicverse/py-scoper |
| py-tigger | tigger (Immcantation / Kleinstein Lab; tigger 1.1.3) | R/Immcantation | immune repertoire (AIRR-Seq) — immunoglobulin novel-allele discovery & V genotyping | novel-allele table, genotype membership and reassigned calls match R exactly; Bayesian likelihood agrees to relative difference below 1e-13 | pip-dependency | ov.airr.find_novel_alleles; ov.airr.infer_genotype | github.com/omicverse/py-tigger |
| py-dowser | dowser (Hoehn, Pybus & Kleinstein, PLoS Comput. Biol. 2022) | R/Immcantation | immune repertoire / B-cell receptor phylogenetics (AIRR-seq) | Validated vs dowser 2.4.1 on ExampleAirr: createGermlines/makeSkyline bit-exact; formatClones per-clone counts & germline lengths exact; getTrees max-parsimony score exact (topology may differ for ambiguous clones, RF small); getDivergence close; correlationTest correlation & slope match, permutation p within Monte-Carlo tolerance | pip-dependency | ov.airr.reconstruct_germlines; ov.airr.lineage_trees; ov.airr.lineage_tests | github.com/omicverse/py-dowser |
| py-immunarch | immunarch 0.10.3 (Nazarov & ImmunoMind team) | CRAN | immune repertoire / bulk TCR-BCR (AIRR-seq) analytics: diversity, clonality, overlap, gene usage, public clonotypes | "bit-exact vs immunarch (max rel diff ~1e-13)" | pip-dependency | ov.airr.repertoire_diversity; ov.airr.repertoire_overlap_bulk; ov.airr.gene_usage_bulk; ov.airr.clonality; ov.airr.kmer_analysis; ov.airr.public_clonotypes; ov.airr.track_clonotypes; ov.airr.load_example_immdata | github.com/omicverse/py-immunarch |
| py-gliph | turboGliph 0.99.2 (R wrapper of GLIPH, Glanville et al. Nature 2017; GLIPH2, Huang et al. Nat. Biotechnol. 2020) | R (other) | immune repertoire / TCR CDR3 specificity-group clustering | "deterministic parts bit-exact vs turboGliph" (qualitative; no single numeric metric) | pip-dependency | ov.airr.specificity_groups | github.com/omicverse/py-gliph |
| **Proteomics / mass-spec** | | | | | | | |
| py-MSstats | MSstats (Choi et al., Bioinformatics 2014) | Bioconductor | mass-spec proteomics (DDA/DIA): summarization, group comparison, sample-size design | Bit-for-bit vs R MSstats 4.14.2: TMP LogIntensities 5e-14, log2FC 8e-15, linear summarization 9e-14, designSampleSize numSample exact; Pearson r=1.0 on every output; 2.5x-3.9x faster | pip-dependency | ov.protein.summarize; ov.protein.sample_size; ov.protein.contrast_matrix | github.com/omicverse/py-MSstats |
| py-DEqMS | DEqMS (Zhu et al., Molecular & Cellular Proteomics 2020) | Bioconductor | proteomics differential expression (peptide-count-aware moderated t-tests for mass-spec) | limma lmFit (coefs/sigma) and eBayes (df.prior/s2.prior/t/p) bit-for-bit; DEqMS sca.t/sca.P Pearson r=1.0, 100% top-500 DE gene agreement; 4.8x faster than R on 5000x12 benchmark | both | ov.protein.de(method='deqms') | github.com/omicverse/py-DEqMS |
| py-proDA | proDA (Ahlmann-Eltze & Anders 2020) | Bioconductor | proteomics (MNAR probabilistic differential expression for label-free mass-spec; censored-ML dropout model) | Wald t_statistic and pval agree with R proDA at Pearson r > 0.99 on the synthetic 300x12 benchmark; invprobit dropout curve matched bit-exactly to R proDA::invprobit | pip-dependency | ov.protein.de (method='proda') | github.com/omicverse/py-proDA |
| py-imputeLCMD | imputeLCMD (Lazar et al., J Proteome Res 2016) | Bioconductor | proteomics — left-censored missing-value imputation for label-free LC-MS/MS | "Bit-for-bit reproduction" for MinDet/ZERO; KS-parity for MinProb/QRILC; KNN/SVD Pearson r > 0.99 vs R; MLE Pearson r ~ 0.98 vs R | pip-dependency | ov.protein.pp.impute | github.com/omicverse/py-imputeLCMD |

| | | | | | | | |
|---|---|---|---|---|---|---|---|
| py-OlinkAnalyze | OlinkAnalyze 3.8.2 (Olink Proteomics AB) | CRAN | proteomics (Olink NPX QC, bridge/subset normalization, per-protein differential expression) | Pearson r > 0.99 (often = 1.0) on per-protein test statistics and p-values for t-test, Wilcoxon, LMM, ANOVA and Kruskal-Wallis vs OlinkAnalyze 3.8.2 | pip-dependency (declared but not actually imported/wired; Olink LMM path reimplemented inline with statsmodels) | — (declared dep; no runtime import) | github.com/omicverse/py-OlinkAnalyze |
| **Metabolomics / lipidomics** | | | | | | | |
| py-lipidr | lipidr (Mohamed, Molendijk & Hill, J. Proteome Res. 2020, 19(7):2890-2897) | Bioconductor | lipidomics — normalization, differential analysis, Lipid Set Enrichment Analysis (LSEA) | stated parity / unquantified (faithful port of lipidr 2.20.0 computational core; reuses pygoslin + python-limma; no verbatim numeric metric) | pip-dependency | ov.metabol.read_skyline; ov.metabol.summarize_transitions; ov.metabol.normalize_pqn; ov.metabol.normalize_istd; ov.metabol.de_lipids; ov.metabol.lsea; ov.metabol.lipid_mva | github.com/omicverse/py-lipidr |
| **Spatial transcriptomics & spatial QC** | | | | | | | |
| py-spanorm | SpaNorm (Bhuva DD, Salim A, Mohamed A; bhuvad/SpaNorm) | Bioconductor | spatial transcriptomics normalisation (spatially-aware GLM normalisation + SVG calling + GLM-PCA) | Real-data correlation 0.999964 (10x Visium SP1); per-gene mean correlation 0.999574; synthetic-data correlation 0.974; R function coverage 25/25 | standalone-only | — (not wired into omicverse) | github.com/omicverse/py-spanorm |
| py-SpaceTrooper | SpaceTrooper (Righelli, Banzi, Marchionni et al., 2024; drighelli/SpaceTrooper) | Bioconductor | spatial QC (image-based spatial transcriptomics: CosMx/MERFISH/Xenium per-cell QC + GLM QC score) | QC_score Pearson r 0.989 (gate >=0.90); intermediate metrics machine-precision (log2SignalDensity max abs err 4.88e-15, Area_um 5.68e-14, total 0.0); outlier labels exact match | standalone-only | — (not wired into omicverse) | github.com/omicverse/py-SpaceTrooper |
| py-spotsweeper | SpotSweeper (MicTott/SpotSweeper, v1.5.0) | Bioconductor | spatial QC (spatially-aware outlier/artifact detection: local variance, MAD local outliers, artifact + Visium outlier flagging) | localVariance max abs err 2.67e-6 (gate 1e-5); localOutliers F1 1.0; findArtifacts ARI 1.0; flagVisiumOutliers exact match 1.0 | standalone-only | — (not wired into omicverse) | github.com/omicverse/py-spotsweeper |
| py-Statial | Statial (Ameen, Iyengar, Qin, Ghazanfar, Patrick, 2022; SydneyBioX/Statial) | Bioconductor | spatial proteomics / single-cell spatial (Kontextual L-function, distances, abundances, state changes) | get_distances max abs err 1.33e-11; get_abundances max abs err 0.0; Kontextual L-function max abs err 7.11e-15; Kontextual value relative error 5.14% (gate 10%, gap from cKDTree vs spatstat closepairs) | standalone-only | — (not wired into omicverse) | github.com/omicverse/py-Statial |
| **Genetics / GWAS** | | | | | | | |
| py-coloc | coloc (Giambartolomei et al., PLoS Genet. 2014; Wallace 2020/2021; Wakefield 2009) — coloc 5.2.3 | CRAN | GWAS/genetics — Bayesian colocalisation of two genetic traits (Approximate Bayes Factors / SuSiE) | posterior probabilities match R package to machine precision (< 1e-4 absolute, typically < 1e-15) | pip-dependency | ov.genetics.colocalize; ov.genetics.finemap_abf; ov.genetics.coloc_sensitivity | github.com/omicverse/py-coloc |
| py-susie | susieR (Wang, Sarkar, Carbonetto & Stephens, JRSS-B 2020; Zou et al., PLoS Genetics 2022) | CRAN | GWAS/genetics — Bayesian statistical fine-mapping (SuSiE / susie_rss credible sets, PIPs) | stated parity / unquantified (faithful port of susieR 0.14.2; no single verbatim numeric metric) | pip-dependency | ov.genetics.finemap; ov.genetics.get_credible_sets; ov.genetics.get_pip; ov.genetics.finemap_plot | github.com/omicverse/py-susie |
| py-ldsc | LDSC / LD Score Regression (Bulik-Sullivan et al. 2015; Finucane et al. 2015, 2018) | Broad/R | GWAS/genetics — LD score regression, SNP heritability, genetic correlation, partitioned h2, LDSC-SEG | Qualitative: 'matches the original numerically' — exact block jackknife and deterministic weighted regressions agree with ldsc 'to floating-point tolerance' (no single verbatim correlation number) | pip-dependency | ov.genetics.heritability; ov.genetics.genetic_correlation; ov.genetics.partitioned_heritability; ov.genetics.ldsc_cell_type; ov.genetics.munge_sumstats | github.com/omicverse/py-ldsc |
| py-matrixeqtl | MatrixEQTL (Shabalin, Bioinformatics 2012, 28(10):1353-1358) | CRAN | GWAS/genetics — fast cis/trans eQTL mapping | Bit-for-bit vs upstream MatrixEQTL 2.3 ('reproduces the upstream results bit-for-bit'; per-quantity Pearson r vs R table with max \|diff\|) | pip-dependency | ov.genetics.eqtl_map | github.com/omicverse/py-matrixeqtl |

| | | | | | | | |
|---|---|---|---|---|---|---|---|
| py-twas | MetaXcan / S-PrediXcan / PrediXcan (Barbeira et al., Nature Communications 2018; hakyimlab/MetaXcan) | R (other) — see caveat (original MetaXcan is Python) | GWAS/genetics — transcriptome-wide association study (TWAS): S-PrediXcan, S-MultiXcan, PrediXcan | "numerically identical to the original MetaXcan (verified to machine precision against a live MetaXcan run)" | pip-dependency | ov.genetics.twas; ov.genetics.load_twas_model; ov.genetics.make_twas_covariance; ov.genetics.build_twas_model; ov.genetics.twas_manhattan | github.com/omicverse/py-twas |
| py-twosamplemr | MendelianRandomization (Yavorska & Burgess, IJE 2017) + TwoSampleMR (Hemani et al., eLife 2018) | CRAN | GWAS/genetics — two-sample Mendelian randomization (IVW/Egger/median/mode/maxlik/cML, harmonisation, diagnostics) | "reproduce MendelianRandomization 0.10.0 bit-for-bit (relative difference < 1e-5); bootstrap-based estimators agree to within Monte-Carlo error"; mr_ivw/mr_egger/mr_maxlik/mr_divw/mr_conmix "bit-exact", mr_median/mr_mbe estimate exact + bootstrap SE approximate | pip-dependency | ov.genetics.mendelian_randomization; ov.genetics.harmonize; ov.genetics.mr_heterogeneity; ov.genetics.mr_pleiotropy; ov.genetics.mr_steiger; ov.genetics.mr_scatter; ov.genetics.mr_forest | github.com/omicverse/py-twosamplemr |
| py-scdrs | scDRS (Zhang, Hou et al. & Price, Nature Genetics 2022, 54:1572-1580) | R (other) / Python-original (Price lab; GWAS-genetics) | GWAS/genetics — single-cell disease relevance scoring (cell-level association of GWAS gene sets) | "reproduces the original scdrs package's control gene sets, scores and p-values bit-for-bit on the bundled toy data" (bit-identical, qualitative; no numeric coefficient) | pip-dependency | ov.genetics.disease_relevance_score; ov.genetics.score_downstream; ov.genetics.scdrs_celltype_plot | github.com/omicverse/py-scdrs |
| **Hi-C / 3D genome & epigenome** | | | | | | | |
| py-dcHiC | dcHiC (ay-lab/dcHiC) | R (other) | Hi-C differential compartment analysis (A/B compartment calling + differential compartments) | compartment_pc1 Pearson \|r\| 1.0000 (max \|Δ\|=1.6e-14, bit-equivalent to dcHiC C++ kernel); differential_padj Spearman 0.981, top-50 Jaccard 0.961 | standalone-only | — (not wired into omicverse) | github.com/omicverse/py-dcHiC |
| rust-bandnorm | BandNorm (Shen, Zheng, Keleş 2022; sshen82/BandNorm) | R (other) | single-cell Hi-C — scGAD scoring + band-normalisation for cell-type discovery/embedding | "bit-equivalent (within f64 ε) to the R reference"; max \|Rust − R\| < 1e-10 on raw counts (integer-exact), < 1e-6 on Z-scored GAD; ~24× (10-30×) faster | standalone-only | — (not wired into omicverse) | github.com/omicverse/rust-bandnorm |
| rust-scHiCluster | scHiCluster (Zhou et al., PNAS 2019) — Gaussian convolution + RWR imputation | R (other) | single-cell Hi-C contact-matrix imputation (RWR + convolution) | bit-equivalent within float-32 epsilon: max\|E_rust - E_scipy\| = 8.94e-8, Pearson 1.000000, nnz match exactly on real Chang chr1; ~9.6x faster on long chromosomes, ~2.7x end-to-end; test_parity.py 11/11 pass (max-rel-error < 1e-4) | standalone-only | — (not wired into omicverse) | github.com/omicverse/rust-scHiCluster |
| **Amplicon / microbiome** | | | | | | | |
| py-dada2 | DADA2 (Callahan et al., Nature Methods 2016) | Bioconductor | amplicon microbiome — exact amplicon sequence variant (ASV) inference from amplicon sequencing | ASV-level identity matches R DADA2 on the canonical MiSeq SOP test data (qualitative; no numeric metric) | pip-dependency (lazy import; not a hard pyproject dep) | ov.alignment.dada2_pipeline; ov.alignment.dada2.filter_and_trim; ov.alignment.dada2.learn_errors; ov.alignment.dada2.denoise; ov.alignment.dada2.merge_pairs; ov.alignment.dada2.make_seqtab; ov.alignment.dada2.remove_chimeras | github.com/omicverse/py-dada2 |
| **Clustering / utilities** | | | | | | | |
| py-mclustR | mclust (Scrucca, Fop, Murphy, Raftery 2016) | CRAN | Gaussian mixture model clustering (BIC model selection, 14 covariance parameterizations) for single-cell | 12 of 14 models bitwise/machine-epsilon parity vs CRAN mclust 6.1.1 (~1e-15); overall 224 records Pearson r 0.9935, loglik r 0.9997; outliers EVE r 0.9751 / VVE r 0.9351 | standalone-only (optional code path; user must pip install pymclustR) | ov.utils.cluster (method='pymclustR') | github.com/omicverse/py-mclustR |

| | | | | | | | |
|---|---|---|---|---|---|---|---|
| py-funkyheatmap | funkyheatmap 0.5.2 (R; used for Saelens et al. 2019 dynbenchmark figures) | R/dynverse | visualization — dynbenchmark-style data-frame heatmaps (funky rectangles/circles/bars/pies) for benchmark/multi-metric tables | max \|Δ\| = 0.0 on xmin/xmax/ymin/ymax/corner_size/color_value (parity gate ≤1e-6 on canonical fixture) | standalone-only (wired via undeclared optional pip install) | ov.pl.funky_heatmap | github.com/omicverse/py-funkyheatmap |

#### Coverage summary

- Total reconstructed packages (deduplicated): 60 rows of collected data → 58 unique packages (the two both-availability AIRR/QC entries and vendored duplicates were already single rows; no cross-batch duplicates remained after merge).
- By source ecosystem:

  - Bioconductor: 21 (py-edgeR, py-limma, py-mfuzz, py-monocle2, py-Monocle3, py-Slingshot, py-TSCAN, py-destiny, py-tradeSeq, py-miloR, py-scDblFinder, py-decontx, py-MSstats, py-DEqMS, py-proDA, py-imputeLCMD, py-lipidr, py-spanorm, py-SpaceTrooper, py-spotsweeper, py-Statial, py-dada2) — 22 listed; note py-dada2 brings the Bioconductor total to 22.
  - CRAN: 9 (py-coloc, py-susie, py-matrixeqtl, py-twosamplemr, py-immunarch, py-mclustR, py-OlinkAnalyze, rust-NMF, py-shazam[1]/py-scoper[1] — see Immcantation note)
  - R/Immcantation: 5 (py-alakazam, py-shazam, py-scoper, py-tigger, py-dowser)
  - R/dynverse: 5 (py-dynwrap, py-dyneval, py-dynbenchmark, py-SCORPIUS, py-funkyheatmap)
  - Broad/R: 2 (py-ldsc, py-inferCNV)
  - R (other) / unlabelled: 16 (py-Augur, py-cca, py-condiments, py-CopyKAT, py-CytoTRACE, py-DoubletFinder, py-fastcar, py-gliph, py-hdWGCNA, py-MARVEL, py-sccdc, py-scmetabolism, py-SigXTalk, py-soupx, py-twas, py-URD, supercell, scTenifold, rust-bandnorm, rust-scHiCluster)

- Integration into OmicVerse:

  - Wired in (pip-dependency + vendored-external + both, including optional/lazy runtime imports that reach a public ov.* function): 37
  - Standalone-only (no ov.* import path): 21 — py-Augur, py-condiments, py-dynbenchmark, py-dyneval, py-dynwrap, py-hdWGCNA, py-inferCNV, py-MARVEL, py-SigXTalk, py-spanorm, py-SpaceTrooper, py-spotsweeper, py-Statial, py-tradeSeq, py-dcHiC, rust-bandnorm, rust-scHiCluster, py-mclustR, py-funkyheatmap, plus py-cca and rust-NMF have wrapper code but ship as undeclared optional installs.
  - Vendored in external/ (or in-tree): py-edgeR, py-limma, py-monocle2, py-Slingshot, py-miloR, py-scmetabolism, supercell, scTenifold.

- Parity:

  - Quantified parity metric (numeric correlation / abs-error / ARI / Pearson / Spearman / Procrustes): ~38 packages.
  - Stated/qualitative parity ("bit-for-bit", "faithful port", "aims for parity") with no single numeric metric: ~19 (py-alakazam, py-shazam, py-scoper, py-edgeR, py-limma, py-Monocle3, py-miloR, py-scDblFinder, py-CytoTRACE, py-dada2, py-lipidr, py-susie, py-dynbenchmark, py-soupx, py-fastcar, py-sccdc, py-gliph, py-immunarch[1], supercell).
  - Unknown: 1 (scTenifold).

- Implementation language:

  - Pure-Python: 55
  - Rust (PyO3/maturin extensions): 3 (rust-NMF, rust-bandnorm, rust-scHiCluster).

#### *Supplementary Table 3 | Representative OmicVerse function contracts*

Fourteen core OmicVerse capability objects, extracted verbatim from their registered contracts: aliases, required AnnData/MuData fields and preconditions, parameter schema, produced fields, and the failure modes the contract makes inspectable before execution. Each step's produced fields satisfy the next step's required fields, defining a bounded, ordered analytical action space.

| Function | Analysis role | Category | Aliases (incl. multilingual) | Required fields / preconditions | Parameter schema (key args) | Produced fields | auto_fix | Failure modes guarded by contract |
|---|---|---|---|---|---|---|---|---|
| ov.pp.qc | Quality control | preprocessing | 质控, qc, quality_control, 质量控制 | raw object (no precondition) | (positional input only) | obs: n_genes, n_counts, pct_counts_mt; var: mt, n_cells | none | hallucinated package/argument or wrong layer/dtype on free invocation |
| ov.pp.preprocess | Normalization + HVG (end-to-end) | preprocessing | 预处理, preprocess, preprocessing, 数据预处理, HVG (+3) | recommended: qc | mode='shiftlog\|pearson', target_sum=…, n_HVGs=2000, organism='human', … (+3) | layers: counts; var: highly_variable_features, means, variances, residual_variances | none | hallucinated package/argument or wrong layer/dtype on free invocation |
| ov.pp.highly_variable_features | HVG selection | preprocessing | 高变基因, highly_variable_features, hvg, hvf, 高变特征筛选 | var: robust \| must-run-first: identify_robust_genes \| recommended: log1p, qc | flavor='pegasus', n_top=2000, span=0.02, min_disp=0.5, … (+5) | var: hvf_rank, highly_variable_features | none | invalid object transition — called before identify_robust_genes (required prior step missing); missing required input state (var: robust); hallucinated package/argument or wrong layer/dtype on free invocation |
| ov.pp.scale | Scaling | preprocessing | 标准化, scale, scaling, 标准化处理 | recommended: normalize, qc | max_value=10, layers_add='scaled', to_sparse=False, use_implicit_centering=False | layers: scaled | none | hallucinated package/argument or wrong layer/dtype on free invocation |
| ov.pp.pca | PCA | preprocessing | 主成分分析, pca, PCA, 降维 | layers: scaled \| must-run-first: scale \| recommended: qc, preprocess | n_pcs=50, layer='scaled', inplace=True | obsm: X_pca; varm: PCs; uns: pca | escalate | invalid object transition — called before scale (required prior step missing); missing required input state (layers: scaled); hallucinated package/argument or wrong layer/dtype on free invocation |
| ov.pp.neighbors | kNN graph | preprocessing | 计算邻居, neighbors, knn, 邻居图 | obsm: X_pca \| recommended: pca | n_neighbors=15, knn=True, random_state=0, method='umap', … (+8) | obsp: distances, connectivities; uns: neighbors | auto | missing required input state (obsm: X_pca); hallucinated package/argument or wrong layer/dtype on free invocation |
| ov.pp.leiden | Clustering | preprocessing | 莱顿聚类, leiden, clustering, 聚类 | uns: neighbors; obsp: connectivities \| must-run-first: neighbors \| recommended: pca, umap | resolution=1.0, random_state=0, key_added='leiden', local_iterations=100, … (+3) | obs: leiden | auto | invalid object transition — called before neighbors (required prior step missing); missing required input state (uns: neighbors; obsp: connectivities); hallucinated package/argument or wrong layer/dtype on free invocation |
| ov.pp.umap | UMAP embedding | preprocessing | umap, UMAP, 非线性降维 | uns: neighbors; obsp: connectivities, distances \| must-run-first: neighbors \| recommended: pca | (positional input only) | obsm: X_umap | auto | invalid object transition — called before neighbors (required prior step missing); missing required input state (uns: neighbors; obsp: connectivities, distances); hallucinated |

| | | | | | | | | |
|---|---|---|---|---|---|---|---|---|
| | | | | | | | | package/argument or wrong layer/dtype on free invocation |
| ov.single.scanpy_cellanno_from_dict | Cell-type annotation | single | 字典注释, scanpy_cellanno_from_dict, manual_annotation, 手动注释, 字典映射注释 | must-run-first: leiden | anno_name='major', clustertype='leiden', … (+1) | obs | none | invalid object transition — called before leiden (required prior step missing); hallucinated package/argument or wrong layer/dtype on free invocation |
| ov.single.batch_correction | Batch integration | single | 批次校正, batch_correction, batch_correct, 数据整合, 去批次效应 | recommended: preprocess, scale, pca | use_rep='scaled\|original\|X_pca', methods='harmony', n_pcs=50, … (+1) | obsm | none | hallucinated package/argument or wrong layer/dtype on free invocation |
| ov.single.gsva | Gene-set / pathway scoring | enrichment | gsva, GSVA, 基因集变异分析 | var: gene symbols matching signature keys \| recommended: preprocess | signatures | obsm: score_gsva | none | missing required input state (var: gene symbols matching signature keys); hallucinated package/argument or wrong layer/dtype on free invocation |
| ov.single.grn | Gene-regulatory network | single | grn, GRN, infer_grn, GRNBoost2 prior, GENIE3 prior (+2) | layers: expression layer or X; var: gene names \| recommended: single.SCENIC | method='grnboost2', log=True, seed=42, n_workers=2, … (+6) | — | none | missing required input state (layers: expression layer or X; var: gene names); hallucinated package/argument or wrong layer/dtype on free invocation |
| ov.space.create_communication_anndata | Spatial cell–cell communication | space | 通信AnnData, create_communication_anndata, cellchat格式转换, commot汇总, 细胞通信汇总 | obsp: commot-*; obs: clustering_column | n_permutations=100, … (+1) | layers: means, pvalues; obs: sender, receiver, cell_type_pair; var: interacting_pair, classification | none | missing required input state (obsp: commot-*; obs: clustering_column); hallucinated package/argument or wrong layer/dtype on free invocation |
| ov.single.chain_qc | Immune-repertoire (AIRR) QC | airr | chain_qc, airr_chain_qc, 免疫链质控, 链配对质控 | obs: VJ_1_junction_aa, VDJ_1_junction_aa | (positional input only) | obs: chain_pairing, receptor_subtype | none | missing required input state (obs: VJ_1_junction_aa, VDJ_1_junction_aa); hallucinated package/argument or wrong layer/dtype on free invocation |

#### Notes

- Each row is a real OmicVerse capability object extracted verbatim from its @register_function contract in source (aliases, category, requires, prerequisites, produces, auto_fix) plus the introspected parameter signature; 14 functions chosen to span the core single-cell analysis chain and several modalities.
- 'Required fields / preconditions' combines the contract's `requires` (object state the function reads: obs/var/obsm/obsp/uns/layers keys) with `prerequisites` (must-run-first = hard dependency functions; recommended = optional). Together these define the valid object transition into the function.
- 'Produced fields' is the contract's `produces` (the obs/var/obsm/uns/layers keys the function writes), which downstream capabilities can then require — e.g. ov.pp.pca produces obsm:X_pca, which ov.pp.neighbors and ov.pp.mde require.
- 'Failure modes guarded by contract' are DERIVED from the same contract: they are the invalid transitions the contract makes inspectable before execution (calling a function before its required upstream step, or on an object lacking the required field/layer), plus the free-call errors (hallucinated package/argument, wrong layer/dtype) that selecting from the bounded manifest prevents. They are not drawn from a separate failure log.
- The chain qc → preprocess → highly_variable_features → scale → pca → neighbors → leiden/umap is fully encoded: each step's `produces` satisfies the next step's `requires`, so the registry defines a bounded, ordered action space rather than free-form code (S4).

### *Supplementary Table 4 | A complete state-transition trace over the live biological object*

One end-to-end scRNA-seq pipeline executed through OmicOS, showing how the AnnData object and its provenance evolve. Each row is a registered capability; the prerequisite check and added fields come from its contract, the BEFORE state is the cumulative object, and the provenance entry mirrors the runtime execution trace. Each step's added fields satisfy the next step's prerequisites.

| Step | Capability | Operation | Prerequisite check (requires / must-run-first) | AnnData state — BEFORE | Fields added (produces) | Provenance / lineage entry |
|---|---|---|---|---|---|---|
| 1 | ov.pp.qc | QC | raw object (none) | X (raw counts), obs:_barcodes, var:_gene_names | obs:n_genes, obs:n_counts, obs:pct_counts_mt, var:mt, var:n_cells | step 1: ov.pp.qc(tresh={mito_perc:0.2, nUMIs:500, detected_genes:250}) → wrote [obs:n_genes, obs:n_counts, obs:pct_counts_mt, var:mt, var:n_cells]; prereq_ok=∅ |
| 2 | ov.pp.preprocess | Normalization + HVG | raw object (none) | X (raw counts), obs:_barcodes, var:_gene_names, obs:n_genes, obs:n_counts, obs:pct_counts_mt, var:mt, var:n_cells | layers:counts, var:highly_variable_features, var:means, var:variances, var:residual_variances | step 2: ov.pp.preprocess(mode='shiftlog\|pearson', n_HVGs=2000) → wrote [layers:counts, var:highly_variable_features, var:means, var:variances, var:residual_variances]; prereq_ok=∅ |
| 3 | ov.pp.scale | Scaling | raw object (none) | X (raw counts), obs:_barcodes, var:_gene_names, obs:n_genes, obs:n_counts, obs:pct_counts_mt, var:mt, var:n_cells, layers:counts, var:highly_variable_features, var:means, var:variances, var:residual_variances | layers:scaled | step 3: ov.pp.scale(max_value=10) → wrote [layers:scaled]; prereq_ok=∅ |
| 4 | ov.pp.pca | Linear embedding (PCA) | ran: scale \| fields: layers:scaled | X (raw counts), obs:_barcodes, var:_gene_names, obs:n_genes, obs:n_counts, obs:pct_counts_mt, var:mt, var:n_cells, layers:counts, var:highly_variable_features, var:means, var:variances, var:residual_variances, layers:scaled | obsm:X_pca, varm:PCs, uns:pca | step 4: ov.pp.pca(n_pcs=50, layer='scaled') → wrote [obsm:X_pca, varm:PCs, uns:pca]; prereq_ok=✓ |
| 5 | ov.pp.neighbors | kNN graph | fields: obsm:X_pca | X (raw counts), obs:_barcodes, var:_gene_names, obs:n_genes, obs:n_counts, obs:pct_counts_mt, var:mt, var:n_cells, layers:counts, var:highly_variable_features, var:means, var:variances, var:residual_variances, layers:scaled, obsm:X_pca, varm:PCs, uns:pca | obsp:distances, obsp:connectivities, uns:neighbors | step 5: ov.pp.neighbors(n_neighbors=15, use_rep='X_pca') → wrote [obsp:distances, obsp:connectivities, uns:neighbors]; prereq_ok=✓ |
| 6 | ov.pp.leiden | Clustering | ran: neighbors \| fields: uns:neighbors, obsp:connectivities | X (raw counts), obs:_barcodes, var:_gene_names, obs:n_genes, obs:n_counts, obs:pct_counts_mt, var:mt, var:n_cells, layers:counts, var:highly_variable_features, var:means, var:variances, var:residual_variances, layers:scaled, obsm:X_pca, varm:PCs, uns:pca, obsp:distances, | obs:leiden | step 6: ov.pp.leiden(resolution=1.0) → wrote [obs:leiden]; prereq_ok=✓ |

| | | | | | | |
|---|---|---|---|---|---|---|
| | | | | obsp:connectivities, uns:neighbors | | |
| 7 | ov.pp.umap | UMAP | ran: neighbors \| fields: uns:neighbors, obsp:connectivities, obsp:distances | X (raw counts), obs:_barcodes, var:_gene_names, obs:n_genes, obs:n_counts, obs:pct_counts_mt, var:mt, var:n_cells, layers:counts, var:highly_variable_features, var:means, var:variances, var:residual_variances, layers:scaled, obsm:X_pca, varm:PCs, uns:pca, obsp:distances, obsp:connectivities, uns:neighbors, obs:leiden | obsm:X_umap | step 7: ov.pp.umap(min_dist=0.5) → wrote [obsm:X_umap]; prereq_ok=✓ |
| 8 | ov.single.scanpy_cellanno_from_dict | Cell-type annotation | ran: leiden | X (raw counts), obs:_barcodes, var:_gene_names, obs:n_genes, obs:n_counts, obs:pct_counts_mt, var:mt, var:n_cells, layers:counts, var:highly_variable_features, var:means, var:variances, var:residual_variances, layers:scaled, obsm:X_pca, varm:PCs, uns:pca, obsp:distances, obsp:connectivities, uns:neighbors, obs:leiden, obsm:X_umap | obs:{anno}_celltype | step 8: ov.single.scanpy_cellanno_from_dict(anno_dict={'0':'T cell',...}, clustertype='leiden') → wrote [obs:{anno}_celltype]; prereq_ok=✓ |

#### Notes

- A single complete scRNA-seq trace: raw counts → QC → normalization+HVG → scaling → PCA → kNN graph → Leiden clustering → UMAP → cell-type annotation. Each step is a real registered OmicVerse capability; its 'Prerequisite check' and 'Fields added' columns are taken verbatim from the function's @register_function contract.
- 'AnnData state — BEFORE' is the cumulative biological object entering the step: the raw object (X counts, obs barcodes, var gene names) plus the union of all fields written by prior steps. 'Fields added' is the step's declared `produces`. Because each step's produces satisfies the next step's requires, prerequisites can be checked before execution and the object is updated in a validated, ordered sequence (S4).
- The 'Provenance / lineage entry' mirrors the record the runtime writes to its execution trace/session history (the invoked capability, key parameters, and the fields written), so each transition is auditable and a failed step can be retried with structured context (which prerequisite was unmet, which fields were expected).
- The same validated capability is invoked identically regardless of surface (chat, notebook, CLI, GUI workspace or programmatic API); only the live AnnData state and this lineage record are shared across surfaces (S3).
- Parameter values shown are representative defaults for illustration; the contract stores the full parameter schema. The annotation step writes a user-named obs column in place, shown here as obs:{anno}_celltype.

### *Supplementary Table 5 | Failure taxonomy: baseline code-generating agent vs OmicOS*

Table S4. Per-task failure localization, baseline code-generating agent vs. OmicOS. For each of seven omics analysis tasks (model in parentheses) we report the baseline's approach, the verbatim error it surfaced, the number of turns and distinct errors before resolution, the baseline outcome, the assigned failure class (one of: missing state, invalid arguments, unavailable dependency, incompatible object structure, failed execution), the OmicOS outcome, and the capability-contract element (required-input state, parameter schema, or produced state) that renders the failure attributable to its class before or at execution. Baseline crashes are opaque tracebacks from downstream libraries (e.g., sklearn's "0 feature(s)" for C04, scVelo's TypeError for F01); the contract column shows how the same crash maps to a specific, recoverable failure class.

| Task | Model | Localizable failure class | Baseline approach (free-form code) | Baseline error (opaque crash) | Baseline turns / errors | Baseline outcome | OmicOS approach (grounded capability) | OmicOS outcome | Attributable by contract element |
|---|---|---|---|---|---|---|---|---|---|
| B05_multi_method_annotation | DeepSeek-v4-flash | invalid arguments | CellTypist reference-based annotation (celltypist.annotate) with 'Healthy_COVID19_PBMC.pkl' model + manual marker-based gene scoring (scanpy.tl.score_genes with PBMC marker gene l… | ValueError: 🛑 Unrecognized `mode` value, should be one of `'best match'` or `'prob match'` — baseline passed mode='best_match' (underscore) but API e… | 33 / 4 | passed | Grounded ov.single.Annotation manager with registry lookup (ov.utils.registry_lookup('cell type annotation')) → discovered registered ov.single.Annotation(adata) capability → used… | passed | The baseline crash would be attributable upfront via **parameter schema in requires/produces** — celltypist.annotate docstring should specify that `mode` parameter accepts exact string literals 'best… |
| F01_multi_mode_velocity | gpt-5.5 | invalid arguments | Attempted direct scVelo preprocessing using scv.pp.filter_and_normalize(adata, min_shared_counts=20, n_top_genes=2000, flavor='seurat') followed by scv.pp.moments() and two veloci… | TypeError: normalize_per_cell() got an unexpected keyword argument 'n_top_genes' | 6 / 4 | passed | Turn 1: Registry lookup (ov.utils.registry_lookup('rna velocity')) discovered ov.single.Velo() unified class with explicit documentation separating filter_genes(min_shared_counts=… | passed | Parameter schema contract: scvelo 0.3.4's filter_and_normalize() should document that n_top_genes parameter is NOT supported and that gene subsetting must be done separately via a dedicated HVG-selec… |
| L01_foundation_model_cell_embedding | gpt-5.5 | unavailable dependency | Direct import and invocation of scgpt package (installed in conda environment) without intermediate registry/wrapper lookup; attempted to import scgpt module to inspect its API an… | OSError: /scratch/users/steorra/env/omicdev/lib/python3.10/site-packages/torchtext/lib/libtorchtext.so: undefined symbol: _ZN5torch3jit17parseSchemaO… | 7 / 2 | passed | Used ov.utils.registry_lookup to discover the unified foundation-model wrapper ov.fm.run (which abstracts over multiple backends: scgpt, geneformer, cellplm, etc.) and its signatu… | passed | A contract element documenting ov.fm.run's checkpoint_dir parameter and its expected behavior when models are pre-downloaded locally (e.g. in a task-specific lookup table of default checkpoint locati… |
| C04_spatial_trajectory_stt | DeepSeek-v4-flash | incompatible object structure | Call `sc.pp.neighbors(adata, n_neighbors=15, n_pcs=0, use_rep='spatial', key_added='spatial_neighbors')` directly on spatial coordinates in obsm['spatial'] to build a kNN graph, t… | ValueError: Found array with 0 feature(s) (shape=(3460, 0)) while a minimum of 1 is required by KNeighborsTransformer | 62 / 7 | failed | Copy spatial coordinates from obsm['spatial'] into the standard representation slot obsm['X_spatial'], then call sc.pp.neighbors(adata, use_rep='X_spatial', n_pcs=None) with the p… | passed | The obsm['spatial'] representation key does not conform to scanpy's use_rep lookup contract—use_rep expects either a named obsm slot starting with 'X_' or a layers key, not a bare 'spatial' key. The … |
| A05_leiden_clustering | DeepSeek-v4-flash | failed execution | Direct use of scanpy sc.tl.leiden() at three resolutions (0.4, 0.8, 1.2), evaluation via silhouette_score on PCA embeddings (X_pca), selection of best resolution, storage of leide… | SyntaxError: f-string expression part cannot include a backslash (line 28: f-string with escaped quote); KeyError: 'leiden_r0.4' due to kernel state … | 6 / 2 | passed | Registry-aware approach: called ov.utils.registry_lookup() and skill_lookup() for task discovery (though final code still used sc.tl.leiden); scanned 5 resolutions (0.4-2.0 vs bas… | passed | A contract element 'prerequisite: neighbors graph in adata.obp[connectivities/distances] and adata.uns[neighbors]' would have prevented the confusion signaled by the hint 'missing state (neighbor gra… |
| B10_gene_regulatory_network | DeepSeek-v4-flash | failed execution | Direct use of arboreto.algo.grnboost2() and attempted pyscenic imports (pyscenic.grn.grnboost2, pyscenic.prune) to build a manual SCENIC pipeline with Dask-based inference, motif … | TypeError: Must supply at least one delayed object (from arboreto/core.py:450, dask_expr/io/_delayed.py:118 in create_graph() when calling from_delay… | 50 / 12 | failed | Used registry_lookup() to discover ov.single.SCENIC class registered in OmicVerse, then used the grounded ov.single.SCENIC API with cal_grn() and cal_regulons() methods, passing p… | passed | The SCENIC class should have a "prerequisites" contract element specifying that grnboost2-based GRN inference requires either a configured dask client or explicit client_or_address parameter, and tha… |
| E04_bulk_wgcna | DeepSeek-v4-flash | failed execution | Manual WGCNA implementation: loaded 192×55448 expression matrix, selected top 2000 genes by variance, computed Pearson correlations, soft-thresholded to adjacency | SyntaxError in f-string formatting (rapidly fixed) | 20 / 2 | passed | OmicVerse-grounded: called ov.bulk.pyWGCNA() wrapper (species='mus musculus', networkType='signed hybrid') on top 1000 MAD-selected genes, ran preprocess() → | passed | The baseline encountered SyntaxError in f-string formatting, a failed execution issue. A parameter schema element (e.g., 'produces: module_trait_corr DataFrame') combined with |

| | | | | | | | | | |
|---|---|---|---|---|---|---|---|---|---|
| | | | matrix, computed… | | | | calculate_soft_thresh… | | pre-execution type chec… |

#### Summary

Across 7 compared tasks the baseline agent averaged 26.3 turns and 4.7 distinct errors per task and failed outright on 2/7 (B10, C04), whereas OmicOS passed 7/7; the localizable failure classes observed were failed execution (3: B10, A05, E04), invalid arguments (2: B05, F01), incompatible object structure (1: C04), and unavailable dependency (1: L01).

- Small, non-random sample: 7 tasks across two models (deepseek_v4_flash, gpt-5.5), not a controlled benchmark; mean turns/errors are descriptive, not statistically powered, and turn counts conflate agents with different stopping behaviors.
- The five-class taxonomy is presented in full, but only 4 classes are instantiated by real failures in this set. 'Missing state' did not actually occur in any compared task — in A05 the neighbor graph was precomputed, so that class is discussed counterfactually ('would have failed if...') rather than observed.
- Outcome is confounded with approach choice, not contracts in isolation. OmicOS trajectories also chose grounded wrappers (e.g., ov.single.SCENIC, ov.fm.run) over hand-rolled pipelines; the benefit attributed to declared contracts is entangled with simply using a maintained, validated API. F01 shows this clearly — OmicOS hit the identical n_top_genes TypeError and converged on the same manual workaround as the baseline, so the contract provided context, not an automatic fix.
- Baseline 'failed' vs 'passed' is coarse: 5/7 baselines eventually passed, several after many turns and incidental bugs (e.g., f-string SyntaxErrors in E04/A05) that are not really about the contract framework and inflate the apparent gap.
- Failure-class assignment is a post-hoc labeling by us of single observed errors; some tasks surfaced multiple distinct errors (B05 had 4, F01 had 4) and were assigned to one dominant class, which simplifies genuinely mixed trajectories.
- The localizability claims describe contract elements that would make a failure attributable; they are largely a design argument about the registry schema rather than a measured ablation showing the contract alone changes the outcome.
- Errors are quoted as reported in the trajectory logs and reflect the specific library versions present (e.g., scVelo 0.3.4, the torch/torchtext build in the omicdev env); some failures are version-specific and would not reproduce under other dependency pins.

### *Supplementary Table 6 | Reconstructed R workflows in OmicVerse*

Complete analytical workflows reconstructed from R/Bioconductor/CRAN, with original R package(s), the Python reconstruction package(s), the explicit parity criterion (acceptance gate), acceleration status vs the R original, and the OmicVerse integration point. Several workflows bundle multiple ported packages. Rows grouped by analysis domain.

| Reconstructed workflow | Original R package(s) | Source ecosystem | Python package(s) | Parity criterion | Acceleration status | OmicVerse integration point |
|---|---|---|---|---|---|---|
| **Bulk RNA-seq & transcriptomics** | | | | | | |
| Bulk RNA-seq differential expression (limma-voom / edgeR) | edgeR (Robinson et al. 2010); limma (Ritchie et al. 2015; eBayes Smyth 2004) | Bioconductor | py-edgeR + py-limma | edgeR: stated R-parity tested against Bioconductor edgeR (NB-GLM / QL-F-test), no numeric gate. limma: stated parity / unquantified (faithful pure-Python port of limma core, no verbatim metric) | parity-focused, not benchmarked (both pure-Python/numpy/scipy, no rpy2/R) | ov.bulk.pyDEG.deg_analysis(method='edger'); ov.bulk.pyDEG (limma/limma-voom backend) |
| Time-series / temporal soft clustering | Mfuzz (Futschik & Carlisle 2005; Kumar & Futschik 2007); e1071 cmeans | Bioconductor | py-mfuzz | standardise/mestimate bit-exact (rel-diff ~1e-15), fill_NA(knn) bit-exact (max abs diff ~1e-15), mfuzz membership Pearson r=1.0 + centres r=1.0 + hard-assignment ARI ~0.99, Dmin Pearson r=1.0 (deterministic kernels asserted bit-exact; fuzzy c-means via agreement, no single numeric gate) | parity-focused (pure-Python/numpy: numpy/scipy/pandas) | ov.bulk.temporal_clusters |
| **Proteomics** | | | | | | |
| Mass-spec proteomics DDA/DIA quantitative workflow | MSstats (Choi et al. 2014); DEqMS (Zhu et al. 2020); proDA (Ahlmann-Eltze & Anders 2020); imputeLCMD (Lazar et al. 2016); limma core (Smyth 2004) | Bioconductor | py-MSstats + py-DEqMS + py-proDA + py-imputeLCMD (+py-limma) | MSstats: bit-for-bit vs MSstats 4.14.2 (TMP LogIntensities max\|Δ\|=5e-14, log2FC 8e-15; Pearson r=1.0; LMM-fallback SE ≤5%, pval r≥0.99). DEqMS: | accelerated-vs-R: MSstats 2.5–3.9x (groupComparison 6.20x); DEqMS 4.78x (88x incl. Rscript startup); proDA ~equal inner-loop / ~2.7x end-to-end; | ov.protein.summarize / ov.protein.sample_size / ov.protein.contrast_matrix ; ov.protein.de(method='deqms'\|'proda'); ov.protein.pp.impute |

| | | | | | | |
|---|---|---|---|---|---|---|
| | | | | limma lmFit bit-exact (atol 1e-9), eBayes t/p atol 1e-4, sca.t/sca.P 5% rtol & r=1.0, top-N 100% agreement. proDA: invprobit bit-exact (atol 1e-12), coef r>0.999, Wald t/pval r>0.99, top-DE >95%. imputeLCMD: MinDet/ZERO np.allclose atol 1e-12, MinProb/QRILC KS p>0.01, KNN/SVD r>0.99, MLE r~0.98, model.Selector 100% flag agreement | imputeLCMD parity-focused. All pure-Python/numpy/scipy/statsmodels, no rpy2 | |
| Olink NPX proteomics QC & differential expression | OlinkAnalyze 3.8.2 (Olink Proteomics AB) | CRAN | py-OlinkAnalyze | Explicit gates vs OlinkAnalyze 3.8.2: t-test estimate atol 1e-8 & statistic/p r>0.99; Wilcoxon \|r\|>0.99; LMM r>0.95; ANOVA F/p r=1.0; Kruskal r=1.0; bridge_selector 100% sample-set overlap; LOD flags >95% agreement | parity-focused (pure-Python: scipy/statsmodels) | standalone repo (not yet wired into ov.*) |
| **Lipidomics / metabolomics** | | | | | | |
| Lipidomics normalization, DE & LSEA | lipidr 2.20.0 (Mohamed et al. 2020) | Bioconductor | py-lipidr | Per-step Pearson r vs lipidr 2.20.0 on A1_data.csv: normalize_pqn r=1.000, normalize_istd r=0.997, de_analysis logFC & p r=1.000, annotate_lipids class agreement 0.99, lsea ES r=0.95 & p r=0.91 ('within target tolerance', no explicit numeric gate) | accelerated-vs-R, ~8x (pure-Python/numpy/scipy; reuses pygoslin + python-limma) | ov.metabol.read_skyline / summarize_transitions / normalize_pqn / normalize_istd / de_lipids / lsea / lipid_mva |
| Single-cell metabolic pathway scoring | scMetabolism (Wu et al. 2021) | R (other) | py-scmetabolism | Correlation with R scMetabolism ≥ 0.99 for most methods (achieved VISION 0.9988, ssGSEA 1.0000, GSVA 0.9870, AUCell 0.9327) | accelerated-vs-R, 3–45x (VISION 45.7x, GSVA 43.5x, AUCell 3.5x, ssGSEA 3.0x); pure-Python/numpy, no rpy2/R | ov.single.Metabolism(method='scmetabolism') |
| **Single-cell QC & preprocessing** | | | | | | |
| Ambient/contaminating-RNA decontamination | SoupX (Young & Behjati 2020); DecontX (Yang et al. 2020); FastCAR (Gocht/Berg et al. 2023); scCDC (Wang et al. 2024) | R (other) / Bioconductor | py-soupx + py-decontx + py-fastcar + py-sccdc | soupx: bit-for-bit on deterministic kernels (estimateSoup ~1e-16, quickMarkers ~1e-16, adjustCounts fixed-rho max abs diff 0 / cell-level ~1e-13, autoEstCont rho exact). decontx: contamination fraction / decontaminated matrix / per-cell theta Pearson r>0.99 vs Bioconductor decontX. fastcar: bit-for-bit identical to R FastCAR (deterministic, gMax/corrected matrix/threshold table element-for-element). sccdc: per-gene/per-cluster Shannon entropy bit-exact, GCG list identical, corrected matrix bit-exact, contamination ratio bit-exact, entropy divergence r>0.99 | parity-focused (all pure-Python/numpy/scipy, no rpy2/R) | ov.pp.ambient.remove_ambient(method='soupx'\|'decontx'\|'fastcar'\|'sccdc'); ov.pp.estimate_contamination |
| Doublet detection | DoubletFinder (McGinnis et al. 2019); scDblFinder (Germain et al. 2022) | R (other) / Bioconductor | py-DoubletFinder + py-scDblFinder | DoubletFinder: bit-for-bit vs R when fed matching PCA embeddings + artificial-doublet pairs (pANN/BCreal/classification, max\|Δ\|=0 on deterministic inputs). scDblFinder: classification overlap gate ≥70% (observed 97.0%/96.2%), score Spearman gate ≥0.2 (observed 0.30); deterministic features (libsizes/cxds/kNN/initial score) atol 1e-12 (xgboost classifier | parity-focused (pure-Python NumPy/SciPy/xgboost) | ov.pp.qc (doublets_method='scdblfinder'); ov.pp.scdblfinder; ov.pp.qc_cpu_gpu_mixed |

| | | | | | | |
|---|---|---|---|---|---|---|
| | | | | intentionally not bit-aligned) | | |
| **Single-cell integration & coarse-graining** | | | | | | |
| Batch integration via canonical correlation analysis | Seurat RunCCA (Stuart, Butler, Hoffman, Hafemeister et al. 2019) | R (other) | py-cca | Tolerance gates vs Seurat::RunCCA (9 size×num_cc configs): per-component singular-value rel error <1e-5; per-component embedding correlation >0.999; Frobenius projector distance <5e-3; 18 parity tests pass | parity-focused (pure-Python/numpy; scipy.sparse.linalg.svds ARPACK for irlba, no rpy2/Rcpp) | ov.single.batch_correction (methods='cca') |
| Metacell construction / coarse-graining | SuperCell (Bilous et al. 2022) | R (other) | supercell | stated parity / unquantified — vendored pure-Python re-implementation of SuperCell core (kNN graph + walktrap), no numeric gate in source | parity-focused, not benchmarked (pure-Python: igraph + sklearn/numpy) | ov.single.MetaCell |
| Gene-program / matrix factorization (NMF) | NMF (Gaujoux & Seoighe 2010) — brunet/lee/offset/nsNMF /snmf-r/snmf-l + HALS/E-HALS/dnmf | CRAN | rust-NMF | bit-equivalence max\|ΔW\| <1e-12 vs R NMF (f64 round-off) for the six R algorithms (hals/ehals/dnmf custom, not bit-eq); biological ARI 0.89 vs cell types (lee+NNDSVD) | rust-accelerated, ~76x (lee, bit-eq) up to ~218–280x (hals/dnmf+NNDSVD), 16 threads — Rust (PyO3/maturin + rayon + matrixmultiply); pbmc8k benchmark | ov.single.NMF |
| Single-cell co-expression network analysis (WGCNA) | hdWGCNA (Morabito et al. 2023) | R (other) | py-hdWGCNA | SFT R^2, kME, hMEs Pearson r=1.0000 vs R hdWGCNA; gates hMEs r 0.9999, MEs 0.9999, kME 1.0000, module count 7=7, assignment agreement 1.0 (no explicit numeric threshold stated) | accelerated-vs-R, 3.1x total (44.62s→14.43s); per-step up to 17.4x (TestSoftPowers); pure-Python/numpy (AnnData-native; MiniBatchKMeans, ThreadPool Harmony, ball-tree KNN, fast SVD) | standalone repo (not yet wired into ov.*) |
| **Single-cell CNV** | | | | | | |
| Copy-number variation inference | CopyKAT (Gao et al. 2021); inferCNV (Tickle, Tirosh et al., Broad) | R (other) / Broad | py-CopyKAT + py-inferCNV | CopyKAT: median py-vs-R classification ARI 0.988 (kappa 0.994) on 17-patient 5-cancer sweep; preprocessing bit-exact ≤1e-12, bin aggregation + Kalman smoother bit-exact ≤1e-10 (GMM/clustering/tree-cut approximate). inferCNV: Spearman rho=1.0000 on continuous post-Phase-1 matrix (10x), 0.9998 smart-seq2, Pearson 0.993–0.999; per-step bit-exact <1e-10 (7/8 Phase 1); HMM Jaccard i3 1.000, i6 0.979 (gates i3≥0.99, i6≥0.96) | accelerated-vs-R: CopyKAT median 23.4x on 8 cores (up to 82.4x); inferCNV 25–192x. pure-Python NumPy/SciPy/Numba (+numba JIT for inferCNV) | ov.single.CNV |
| **Single-cell differential abundance** | | | | | | |
| Differential abundance testing on kNN neighbourhoods | Milo / miloR (Dann et al. 2022) | Bioconductor | py-miloR | stated parity / unquantified — README "Same API as R miloR"; TMM normalisation (calcNormFactors) + statsmodels QL-F NB GLM "match miloR's edgeR step"; no numeric tolerance gate | parity-focused, not benchmarked (pure-Python/numpy: NumPy+SciPy+statsmodels QL-F GLM; no rpy2/edgeR) | ov.single.Milo |
| **Single-cell trajectory inference** | | | | | | |
| scRNA-seq trajectory inference (TrajInfer family) | Monocle 3 (Cao et al. 2019); CytoTRACE (Gulati et al. 2020); Slingshot (Street et al. 2018); SCORPIUS (Cannoodt et al. 2016); URD (Farrell et al. 2018); destiny/DPT (Angerer et al. 2016) | Bioconductor / R-dynverse / R (other) | py-Monocle3 + py-CytoTRACE + py-Slingshot + py-SCORPIUS + py-URD + py-destiny | Monocle3: stated parity / unquantified (thin wrapper re-exporting Bio-Babel/Monocle3-python, defers R-parity upstream). CytoTRACE: stated parity / unquantified (functions marked ported, no metric). Slingshot: pseudotime Spearman ≥0.99 vs R (Y-shaped fixture). SCORPIUS: pseudotime Pearson ≥0.95 (achieved 0.989), low-dim Procrustes ≥0.85 (0.999). URD: pseudotime Spearman 0.985 / Pearson 0.992 (threshold 0.80) on Guo qPCR 428 cells. destiny: | mostly parity-focused (all pure-Python/numpy/scipy/sklearn); Monocle3 not benchmarked | ov.single.TrajInfer(method='monocle3'\|'cytotrace'\|'slingshot'\|'scorpius'\|'urd'\|'destiny') |

| | | | | | | |
|---|---|---|---|---|---|---|
| | | | | eigenvectors Procrustes ≥0.80 (0.916), eigenvalues Pearson ≥0.95 (0.967), DPT Pearson ≥0.85 (0.974) | | |
| DDRTree trajectory inference | Monocle 2 (Qiu et al. 2017) | Bioconductor | py-monocle2 | Pseudotime correlation with R Monocle 2 ≥ 0.99 on every benchmark dataset (achieved 0.9900 on pancreas); 'exact' mode targets bitwise parity (Delaunay MST bit-identical; eigsh PCA matches irlba top-k to 1e-5 rel) | accelerated-vs-R, 30–100x (102x on pancreas 3,696 cells, 'fast' mode); pure-Python/numpy (fast DDRTree, Delaunay-MST, eigsh PCA, no C++/rpy2) | ov.single.Monocle |
| mclust+MST+spline pseudotime | TSCAN (Ji & Ji 2016) | Bioconductor | py-TSCAN | Pearson = 1.000000 vs R pseudotime in verification mode (Mclust clusters match); pre-registered gate in tests/test_exact_match.py; steps bit-identical to R TSCAN | accelerated-vs-R, ~28x; pure-Python/numpy (NumPy/SciPy/pymclust R) | ov.single.TrajInfer(method='TSCAN') |
| Trajectory-based differential expression (NB-GAM) | tradeSeq (Van den Berge et al. 2020) | Bioconductor | py-tradeSeq | associationTest Spearman on -log10 p ≥ 0.70 (explicit threshold; measured 0.84, top-50 Jaccard 0.67); startVsEndTest Spearman 0.45 (approximate, no threshold) | accelerated-vs-R, 5x faster than R fitGAM; pure-Python (statsmodels.gam vs R mgcv) | standalone repo (not yet wired into ov.*) |
| Differential trajectory analysis across conditions | condiments (Roux de Bezieux et al. 2024) | R (other) | py-condiments | imbalance_score Pearson = 1.0000 vs R on canonical toy fixture (no explicit gate); progressionTest agrees on significance; topologyTest simplified/approximate in v0.1 | parity-focused (pure-Python/numpy) | standalone repo (not yet wired into ov.*) |
| Cell-type prioritization (perturbation response) | Augur (Skinnider et al. 2021) | R (other) | py-Augur | Pearson r (AUC) 0.9999 ≥ 0.95 gate (PASS); Spearman rho (ranking) 1.0000 = 1.0 gate (PASS); ranking CellTypeC>B>A preserved | accelerated-vs-R, 3.8x (227.8s→59.8s on calculate_auc); pure-Python (statsmodels lowess, custom _FastRandomForest, n_jobs=1) | standalone repo (not yet wired into ov.*) |
| Trajectory-inference benchmarking (dynverse) | dynwrap, dyneval, dynbenchmark (Saelens et al. 2019) | R/dynverse | py-dynwrap + py-dyneval + py-dynbenchmark | dynwrap: byte-equivalent geodesic distances, max\|Δ\|=0.0 on 3-branch/30-cell fixture. dyneval: correlation 0.9812(R)/0.9813(Py) \|Δ\|0.0001; him/edge_flip/isomorphic/F1_branches \|Δ\|=0.0; featureimp_cor 0.9622 vs 0.9005; 8/14 metric IDs ported (no explicit gate; featureimp differs from RF RNG). dynbenchmark: stated parity / unquantified (one-line repo description, no gate) | dynwrap & dyneval parity-focused; dynbenchmark not-benchmarked (all pure-Python/numpy: NumPy/SciPy/NetworkX/sklearn) | standalone repo (not yet wired into ov.*) |
| Benchmark-table visualization | funkyheatmap 0.5.2 (R; dynbenchmark figures) | R/dynverse | py-funkyheatmap | Parity gate max\|Δ\| over funkyrect_data/circle_data geometry ≤1e-6 on canonical fixture; achieved max\|Δ\|=0.0 on xmin/xmax/ymin/ymax/corner_size/color_value (bit-for-bit) | parity-focused (pure-Python/numpy: numpy/pandas/matplotlib, hand-rolled cubic-Bezier rounded rects, no R/ggplot/rpy2) | ov.pl.funky_heatmap |
| **Single-cell clustering & misc** | | | | | | |
| Gaussian mixture model clustering | mclust (Scrucca, Fop, Murphy, Raftery 2016) | CRAN | py-mclustR | Per-model table vs mclust 6.1.1: 12/14 models bitwise/machine-epsilon parity (loglik rel-err ~1e-15, z/mean/sigma corr 1.0000, classification 1.0000); overall 224 records z corr 0.9935, loglik 0.9997; outliers EVE z corr 0.9751 / VVE 0.9351 (no single explicit threshold) | parity-focused (pure-Python/numpy) | ov.utils.cluster(method='pymclustR') |
| Single-cell alternative splicing | MARVEL (Wen et al. 2023) | R (other) | py-MARVEL | min Pearson r 0.9999961 (variable_splicing); most artifacts r~1.0, max abs diff 6.66e-16 to 8.91e-4 vs R MARVEL (no explicit | accelerated-vs-R, 3.23x (plate full replay), 2.96x (droplet); pure-Python/numpy | standalone repo (not yet wired into ov.*) |

| | | | | | | |
|---|---|---|---|---|---|---|
| | | | | gate; reference fixtures + external replay) | | |
| Cell-cell signaling crosstalk inference | SigXTalk (HGNN framework) | R (other) | py-SigXTalk | Pearson Correlation 0.9912 (Python vs R, PBMC3k) — achieved, no explicit gate | parity-focused; GPU/torch (HGNN deep-learning, device cpu/cuda) | standalone repo (not yet wired into ov.*) |
| Gene-regulatory network comparison & virtual knockout | scTenifoldNet / scTenifoldKnk (Osorio et al. 2020/2022) | R (other) | scTenifold | unknown / unquantified — vendored pure-Python port (PCNet networks, manifold alignment, d_regulation, virtual KO); no parity gate or vs-R metric | parity-focused, not benchmarked (pure-Python/numpy; randomized-SVD network construction) | ov.single.scTenifoldKnk |
| **Spatial transcriptomics** | | | | | | |
| Spatial normalization (spatially-aware GLM) | SpaNorm (Bhuva, Salim, Mohamed) | Bioconductor | py-spanorm | Real-data correlation 0.999964 on 10x Visium SP1 (100 genes × 2940 spots); per-gene mean correlation 0.999574; synthetic 0.974; R function coverage 25/25 (no explicit gate) | accelerated-vs-R, 16.6x faster; pure-Python/numpy (IRLS NB-GLM) | standalone repo (not yet wired into ov.*) |
| Image-based spatial QC (cell-level GLM score) | SpaceTrooper (Righelli, Banzi, Marchionni et al. 2024) | Bioconductor | py-SpaceTrooper | QC_score Pearson r $\geq$ 0.90 gate (achieved 0.989); intermediate metrics max abs err <1e-8 (log2SignalDensity 4.88e-15, Area_um 5.68e-14, total 0.0); outlier labels exact match | accelerated-vs-R, 102x; pure-Python/numpy (sklearn LogisticRegression replaces glmnet) | standalone repo (not yet wired into ov.*) |
| Spatial outlier/artifact QC | SpotSweeper (MicTott, v1.5.0) | Bioconductor | py-spotsweeper | localVariance gate max abs err <1e-5 (achieved 2.67e-6); localOutliers F1=1.0; findArtifacts ARI=1.0; flagVisiumOutliers exact match 1.0 | accelerated-vs-R (factor not stated); pure-Python/numpy (kNN + Huber robust regression, PCA + k-means) | standalone repo (not yet wired into ov.*) |
| Spatial proteomics cell-state / spatial association | Statial (Ameen, Iyengar, Qin, Ghazanfar, Patrick 2022) | Bioconductor | py-Statial | get_distances gate <1e-8 (achieved 1.33e-11); get_abundances <1e-8 (0.0); Kontextual L-function <1e-8 (7.11e-15); Kontextual value rel err <10% (achieved 5.14%) | accelerated-vs-R, 13.6x total (get_distances 16.9x, get_abundances 12.4x, Kontextual 2.9x); pure-Python (scipy cKDTree, scikit-learn) | standalone repo (not yet wired into ov.*) |
| **Microbiome / amplicon** | | | | | | |
| Amplicon sequence-variant (ASV) inference | DADA2 (Callahan et al. 2016) | Bioconductor | py-dada2 | ASV-level identity matches R DADA2 on canonical MiSeq SOP test data (qualitative; no numeric metric or gate) | parity-focused (pure-Python, no rpy2/R) | ov.alignment.dada2_pipeline (+ filter_and_trim / learn_errors / denoise / merge_pairs / make_seqtab / remove_chimeras) |
| **Immune repertoire (AIRR-seq)** | | | | | | |
| B-cell AIRR / BCR analysis (Immcantation) | alakazam (Gupta et al. 2015); shazam (Gupta, Vander Heiden et al. 2015); scoper; tigger; dowser (Hoehn et al. 2022) | R/Immcantation | py-alakazam + py-shazam + py-scoper + py-tigger + py-dowser | alakazam: deterministic functions bit-exact / rel-diff <1e-6 vs alakazam 1.4.3, bootstrap (estimateAbundance/alphaDiversity) Pearson r>0.999, lineage identical-length tree 31/49. shazam: observedMutations/distToNearest/calcTargetingDistance bit-exact (max abs diff 0), substitution matrix & expectedMutations rel-diff <1e-15, findThreshold <1e-8, BASELINe sigma <1e-13, baselineCI <1e-8. scoper: ARI >0.95 + clone count within a few percent (identicalClones 1.000, hierarchical 1.000, spectral ~0.98–0.99). tigger: novel-allele table/genotype/calls match R exactly, Bayesian likelihood rel-diff <1e-13. dowser: createGermlines/makeSkyline bit-exact, getTrees max-parsimony exact (RF small), correlationTest match (permutation p within MC tolerance) | parity-focused (all pure-Python/numpy: numpy/scipy/pandas/biopython, no rpy2/PHYLIP/phylo binaries) | ov.airr.hill_diversity / aa_properties / clonal_abundance / bcr_gene_usage; ov.airr.mutation_analysis / shm_targeting / baseline_selection / distance_threshold; ov.airr.clonal_clustering; ov.airr.find_novel_alleles / infer_genotype; ov.airr.reconstruct_germlines / lineage_trees / lineage_tests |
| Bulk & TCR repertoire analytics | immunarch 0.10.3 (Nazarov & ImmunoMind); | CRAN / R (other) | py-immunarch + py-gliph | immunarch: rel-diff <1e-6 for every function family vs immunarch 0.10.3; | immunarch accelerated-vs-R ~60x (incl. Rscript startup); | ov.airr.repertoire_diversity / repertoire_overlap_bulk / |

| | | | | | | |
|---|---|---|---|---|---|---|
| | turboGliph 0.99.2 (GLIPH/GLIPH2; Glanville et al. 2017; Huang et al. 2020) | | | achieved bit-exact, max rel diff ~1e-13. gliph: deterministic parts bit-exact vs turboGliph (find_motifs counts, GLIPH2 hypergeometric p/fold-change/motifs, global structures, edge list, clusters, de_novo PWMs); RNG-dependent resampling scores correlation >0.95 | gliph parity-focused. Both pure-Python/numpy (numpy/scipy/pandas/networkx/sklearn/matplotlib) | gene_usage_bulk / clonality / kmer_analysis / public_clonotypes / track_clonotypes; ov.airr.specificity_groups |
| **Epigenome / Hi-C** | | | | | | |
| Single-cell Hi-C band-normalization & scGAD | BandNorm (Shen, Zheng, Keleş 2022) | R (other) | rust-bandnorm | max\|Rust − R\| <1e-10 on raw counts (integer-exact), <1e-6 on Z-scored GAD; bit-equivalent within f64 epsilon | rust-accelerated, ~24x (10–30x); Rust (PyO3/maturin); scGADExample 100 cells × ~1.3M contact pairs | standalone repo (not yet wired into ov.*) |
| Single-cell Hi-C contact-matrix imputation | scHiCluster (Zhou et al. 2019) | R (other) | rust-scHiCluster | bit-equivalence within float-32 epsilon: max-rel-error <1e-4 (11/11 pass); achieved max\|E_rust−E_scipy\|=8.94e-8, Pearson r=1.000000, nnz match exactly on real Chang chr1 | rust-accelerated, ~9.6x on long chromosomes, ~2.7x end-to-end per cell (+3.1x multiprocess); Rust (PyO3/maturin), steps 2–9 in Rust, cooler/HDF5 I/O in Python | standalone repo (not yet wired into ov.*) |
| Bulk Hi-C differential compartment analysis | dcHiC (ay-lab/dcHiC) | R (other) | py-dcHiC | compartment_pc1 Pearson \|r\| ≥0.99 (achieved 1.0000, max\|Δ\|=1.6e-14, bit-equivalent to dcHiC C++ kernel); differential_padj Spearman(-log10p) ≥0.90 (achieved 0.981), top-50 Jaccard ≥0.7 (achieved 0.961) | parity-focused (pure-Python/numpy: NumPy/SciPy/scikit-learn) | standalone repo (not yet wired into ov.*) |
| **Statistical genetics / GWAS** | | | | | | |
| Bayesian colocalization | coloc 5.2.3 (Giambartolomei et al. 2014; Wallace 2020/2021; Wakefield 2009) | CRAN | py-coloc | vs coloc 5.2.3 on coloc_test_data (500 SNPs): coloc_abf PP.H0..H4 max abs diff <3e-16 & SNP.PP.H4 r=1; finemap_abf SNP.PP r=1; coloc_susie <4e-16; coloc_signals <4e-16; sensitivity grid <1e-6 (README: posterior probs match to machine precision, <1e-4, typically <1e-15) | parity-focused (pure-Python/numpy: numpy/scipy/pandas/matplotlib, no R/rpy2) | ov.genetics.colocalize / finemap_abf / coloc_sensitivity |
| Bayesian statistical fine-mapping | susieR 0.14.2 (Wang et al. 2020; Zou et al. 2022) | CRAN | py-susie | PIP Pearson r=1.000 (max abs diff ~1e-11 susie, ~1e-9 susie_rss); per-effect alpha ~1e-11; ELBO diff ~1e-12; credible sets identical vs susieR 0.14.2 (no explicit pass/fail threshold; tests/test_r_parity.py is the gate) | parity-focused (pure-Python/numpy: numpy/scipy/pandas) | ov.genetics.finemap / get_credible_sets / get_pip / finemap_plot |
| LD score regression / heritability | LDSC (Bulik-Sullivan et al. 2015; Finucane et al. 2015, 2018) | Broad/R | py-ldsc | deterministic h2/intercept/rg/jackknife-SE reproduce to rtol=1e-6 on simulate_test; exact-equality identities hold; statistical unbiasedness checks (README: matches numerically, agrees to floating-point tolerance) | parity-focused (pure-Python/numpy/scipy, no rpy2) | ov.genetics.heritability / genetic_correlation / partitioned_heritability / ldsc_cell_type / munge_sumstats |
| cis/trans eQTL mapping | MatrixEQTL 2.3 (Shabalin 2012) | CRAN | py-matrixeqtl | Bit-for-bit vs MatrixEQTL 2.3: per-quantity Pearson r=1.0000000, beta max\|diff\|<2e-15, t/F <1.1e-11, p <2e-14, FDR <3e-14; cis/trans split counts match exactly | parity-focused, not benchmarked (pure-Python/numpy: numpy/scipy/pandas; raw timing only, no R-vs-Py factor) | ov.genetics.eqtl_map |
| Transcriptome-wide association (TWAS) | MetaXcan / S-PrediXcan / PrediXcan (Barbeira et al. 2018) | R (other) | py-twas | S-PrediXcan zscore/effect_size/pvalue/var_g bit-identical to MetaXcan (max abs diff ~4e-16, z r=1.0); S-MultiXcan ~1e-15; PrediXcan OLS/logistic ~1e-9 vs statsmodels (cell-by-cell agreement, | parity-focused (pure-Python/numpy: numpy/scipy/pandas/sqlite3, no rpy2/compiled) | ov.genetics.twas / load_twas_model / make_twas_covariance / build_twas_model / twas_manhattan |

| | | | | | | |
|---|---|---|---|---|---|---|
| | | | | tolerance-style bit-identity, no ≥ gate) | | |
| Two-sample Mendelian randomization | MendelianRandomization 0.10.0 (Yavorska & Burgess 2017); TwoSampleMR (Hemani et al. 2018) | CRAN | py-twosamplemr | Closed-form estimators reproduce MendelianRandomization 0.10.0 bit-for-bit (rel-diff <1e-5); bootstrap estimators within Monte-Carlo error (bootstrap SE within ~10%); per-method rel diffs 2e-16 to 6e-12 | parity-focused (pure-Python/numpy: numpy/scipy/pandas/matplotlib, no rpy2; line-for-line port of R nmmin C optimiser) | ov.genetics.mendelian_randomization / harmonize / mr_heterogeneity / mr_pleiotropy / mr_steiger / mr_scatter / mr_forest |
| Single-cell disease-relevance scoring | scDRS (Zhang, Hou et al. & Price 2022) | R (other) | py-scdrs | tests/test_parity.py asserts agreement of preprocess (gene stats, mean-variance bins), score_cell (raw/norm score, pval, mc_pval, zscore) and downstream group/gene/correlation stats vs upstream scdrs with identical random_seed on toy data; bit-exact (gate = bit-for-bit, no numeric threshold value) | parity-focused, not benchmarked (pure-Python/numpy: numpy/scipy/pandas/anndata/scanpy, no R/rpy2) | ov.genetics.disease_relevance_score / score_downstream / scdrs_celltype_plot |

#### Summary

- 43 reconstructed workflows assembled from all 64 ported packages (10 multi-package bundles — bulk DE, proteomics DDA/DIA, ambient-RNA, doublet QC, CNV, TrajInfer family, Immcantation B-cell AIRR, bulk/TCR repertoire, dynverse benchmarking — covering ~33 packages; the remaining rows are single-package workflows).
- Acceleration posture: 15 workflows are accelerated-vs-R or Rust-accelerated; ~14 are parity-focused (comparable, not benchmarked for speed); 3 contain at least one explicitly not-benchmarked port (TrajInfer/Monocle3, dynverse/dynbenchmark, MatrixEQTL, scDRS). Acceleration leaders span ~3.8x (Augur) to ~280x (NMF HALS) and ~192x (inferCNV).
- Rust-backed: 3 workflows — NMF (~76–280x), single-cell Hi-C BandNorm (~24x), single-cell Hi-C scHiCluster (~9.6x) — all PyO3/maturin.
- OmicVerse integration: 28 workflows are wired into the public ov. API (bulk DE, temporal clustering, proteomics, lipidomics, scMetabolism, ambient removal, doublet QC, CCA integration, MetaCell, NMF, CNV, Milo differential abundance, the TrajInfer family + Monocle2 + TSCAN, scTenifoldKnk, mclust, DADA2, both AIRR bundles, funky_heatmap, and all six genetics workflows); 15 are standalone repos not yet wired into ov. (Olink, hdWGCNA, tradeSeq, condiments, Augur, dynverse, MARVEL, SigXTalk, the four spatial ports, all three Hi-C ports).
- Parity criteria range from strict bit-equivalence / machine-precision gates (MatrixEQTL, coloc, susie, TWAS, MR, DoubletFinder, FastCAR, soupx, shazam, tigger, immunarch, MSstats, dcHiC compartments, dynwrap, funkyheatmap, NMF, Hi-C ports) through moderate correlation/ARI thresholds (Augur r≥0.95, scoper ARI>0.95, scMetabolism r≥0.99, tradeSeq Spearman≥0.70, SpaceTrooper r≥0.90, destiny Procrustes≥0.80) to qualitative "stated parity / unquantified" ports with no numeric gate (edgeR, limma, lipidr, CytoTRACE, Monocle3, miloR, SuperCell, scDblFinder narrative, dynbenchmark, scTenifold).

#### Notes

- Multi-package bundles: four workflows fuse the most packages — proteomics DDA/DIA (ov.protein: MSstats+DEqMS+proDA+imputeLCMD+limma), ambient-RNA removal (ov.pp.ambient: soupx+decontx+fastcar+sccdc), the Immcantation B-cell AIRR stack (ov.airr: alakazam+shazam+scoper+tigger+dowser), and the TrajInfer family (ov.single.TrajInfer: Monocle3+CytoTRACE+Slingshot+SCORPIUS+URD+destiny). Bulk DE (edgeR+limma), doublet QC (DoubletFinder+scDblFinder), bulk/TCR repertoire (immunarch+gliph), and dynverse benchmarking (dynwrap+dyneval+dynbenchmark) are smaller bundles.
- Acceleration leaders: Rust ports dominate (NMF ~76–280x; BandNorm ~24x; scHiCluster ~9.6x). Among pure-Python ports, inferCNV (25–192x), Monocle2 (30–100x), SpaceTrooper (102x), immunarch (~60x), CopyKAT (median 23.4x, up to 82.4x), scMetabolism (3–45x), TSCAN (~28x), spanorm (16.6x), and Statial (13.6x) are the strongest.
- Strict bit-equivalence gates (exact-match / machine-precision): MatrixEQTL, coloc, susie, TWAS, MR closed-form estimators, DoubletFinder, FastCAR, soupx kernels, shazam deterministic functions, tigger, immunarch, MSstats, DEqMS lmFit/eBayes, dcHiC compartment PC1, dynwrap geodesics, funkyheatmap geometry, scDRS, and all three Rust ports.
- Loose correlation/threshold gates (RNG or approximation-bound): Augur (r≥0.95), scoper (ARI>0.95), tradeSeq (Spearman≥0.70), destiny (Procrustes≥0.80/Pearson≥0.85–0.95), SpaceTrooper (r≥0.90), URD (≥0.80), scDblFinder (overlap≥70%, score Spearman≥0.2), and scMetabolism (r≥0.99 for most methods).

- Standalone-only workflows pending ov.* integration: OlinkAnalyze, hdWGCNA, tradeSeq, condiments, Augur, MARVEL, SigXTalk, the dynverse trio, all four spatial ports (SpaNorm, SpaceTrooper, SpotSweeper, Statial), and all three Hi-C ports (BandNorm, scHiCluster, dcHiC) — each validated against its R reference but not yet exposed through the public API.

### *Supplementary Table 7 | BixBench process-quality evaluation by task category*

Process-quality scores for OmicOS on BixBench-Verified-50 (50 questions). Every best-per-question trajectory was graded by a DeepSeek-v4-pro judge on five 0-100 process dimensions (model selection, data exploration, iteration pattern, answer fidelity, execution correctness). Table 5a gives per-category means with 95% bootstrap confidence intervals; Table 5b lists every task ID with its judge scores; the judge protocol, raw-trajectory validation, and caveats follow.

#### Table 5a | Per-category rubric means (mean [95% CI])

| Category | n tasks | Pass@1 | Data exploration | Model selection | Execution correctness | Answer fidelity | Iteration pattern | Category mean (5 dims) |
|---|---|---|---|---|---|---|---|---|
| Transcriptomics | 14 | 14/14 (100%) | 93.9 [91.1, 96.4] | 92.1 [88.2, 95.7] | 89.6 [83.2, 94.6] | 94.1 [82.6, 100.0] | 95.0 [92.1, 97.1] | 93.0 [89.5, 96.0] |
| DNA-level | 18 | 16/18 (89%) | 92.6 [89.7, 95.1] | 95.0 [92.2, 97.5] | 94.3 [88.4, 98.1] | 98.9 [97.5, 100.0] | 97.8 [96.4, 99.2] | 95.7 [94.0, 97.2] |
| Epigenomics & Functional | 7 | 6/7 (86%) | 93.6 [92.1, 95.0] | 97.1 [95.7, 99.3] | 94.3 [87.1, 98.6] | 99.3 [97.9, 100.0] | 99.3 [97.9, 100.0] | 96.7 [95.0, 98.0] |
| Other Modalities | 11 | 9/11 (82%) | 88.6 [79.1, 95.0] | 91.4 [80.0, 98.2] | 95.9 [92.3, 98.6] | 99.1 [97.3, 100.0] | 96.8 [95.0, 98.6] | 94.4 [89.3, 97.5] |
| **All categories (overall)** | **50** | **45/50 (90%)** | **92.2 [89.6, 94.2]** | **93.7 [90.7, 96.0]** | **93.4 [90.4, 95.9]** | **97.7 [94.2, 99.7]** | **97.0 [95.8, 98.0]** | **94.8 [93.1, 96.2]** |

#### Table 5b | Per-task judge scores

| Task ID | Category bucket | Raw category tags | Agent | Harness Pass@1 | Data exploration | Model selection | Execution correctness | Answer fidelity | Iteration pattern | Task mean |
|---|---|---|---|---|---|---|---|---|---|---|
| bix-22-q1 | Transcriptomics | RNA-seq; Sequence Analysis; Transcriptomics | omicverse_omni | pass | 100 | 100 | 100 | 100 | 100 | 100.0 |
| bix-22-q4 | Transcriptomics | RNA-seq; Sequence Analysis; Transcriptomics | omicverse_omni | pass | 95 | 100 | 95 | 100 | 100 | 98.0 |
| bix-24-q2 | Transcriptomics | Differential Expression Analysis; RNA-seq; Sequence Analysis; Transcriptomics | bulk_rna_analyst | pass | 95 | 85 | 90 | 100 | 95 | 93.0 |
| bix-26-q3 | Transcriptomics | Differential Expression Analysis; RNA-seq; Sequence Analysis; Transcriptomics | bulk_rna_analyst | pass | 95 | 85 | 95 | 100 | 95 | 94.0 |
| bix-26-q5 | Transcriptomics | Differential Expression Analysis; RNA-seq; Sequence Analysis; Transcriptomics | bulk_rna_analyst | pass | 90 | 80 | 70 | 100 | 80 | 84.0 |
| bix-30-q3 | Transcriptomics | Differential Expression Analysis; Transcriptomics | bulk_rna_analyst | pass | 95 | 85 | 95 | 100 | 95 | 94.0 |
| bix-31-q2 | Transcriptomics | Differential Expression Analysis; RNA-seq; | bulk_rna_analyst | pass | 100 | 100 | 95 | 100 | 100 | 99.0 |

| | | | | | | | | | | |
|---|---|---|---|---|---|---|---|---|---|---|
| | | Transcriptomics | | | | | | | | |
| bix-32-q2 | Transcriptomics | Differential Expression Analysis; RNA-seq; Sequence Analysis; Transcriptomics | bulk_rna_analyst | pass | 80 | 85 | 65 | 100 | 90 | 84.0 |
| bix-43-q2 | Transcriptomics | Differential Expression Analysis; RNA-seq; Transcriptomics | bulk_rna_analyst | pass | 95 | 95 | 95 | 98 | 95 | 95.6 |
| bix-43-q4 | Transcriptomics | Differential Expression Analysis; RNA-seq; Transcriptomics | bulk_rna_analyst | pass | 90 | 85 | 100 | 100 | 95 | 94.0 |
| bix-46-q4 | Transcriptomics | Differential Expression Analysis; RNA-seq; Sequence Analysis; Transcriptomics | bulk_rna_analyst | pass | 100 | 100 | 100 | 100 | 100 | 100.0 |
| bix-49-q4 | Transcriptomics | Differential Expression Analysis; RNA-seq; Transcriptomics | bulk_rna_analyst | pass | 90 | 95 | 75 | 100 | 95 | 91.0 |
| bix-53-q2 | Transcriptomics | Differential Expression Analysis; RNA-seq; Sequence Analysis; Transcriptomics | bulk_rna_analyst | pass | 95 | 100 | 90 | 100 | 95 | 96.0 |
| bix-53-q5 | Transcriptomics | Differential Expression Analysis; RNA-seq; Sequence Analysis; Transcriptomics | bulk_rna_analyst | pass | 95 | 95 | 90 | 20 | 95 | 79.0 |
| bix-11-q1 | DNA-level | Genomics; Phylogenetics and Evolutionary Analysis | omicverse_omni | pass | 95 | 90 | 95 | 100 | 100 | 96.0 |
| bix-11-q2 | DNA-level | Genomics; Phylogenetics and Evolutionary Analysis | omicverse_omni | pass | 95 | 90 | 95 | 100 | 100 | 96.0 |
| bix-12-q2 | DNA-level | Genomics; Phylogenetics and Evolutionary Analysis; Whole Genome Sequencing (WGS) | omicverse_omni | pass | 95 | 80 | 95 | 100 | 95 | 93.0 |
| bix-12-q4 | DNA-level | Genomics; Phylogenetics and Evolutionary Analysis; Whole Genome Sequencing (WGS) | omicverse_omni | pass | 95 | 100 | 100 | 100 | 100 | 99.0 |
| bix-12-q5 | DNA-level | Genomics; Phylogenetics and Evolutionary Analysis; Whole Genome Sequencing (WGS) | omicverse_omni | pass | 95 | 85 | 95 | 100 | 95 | 94.0 |
| bix-12-q6 | DNA-level | Genomics; Phylogenetics and Evolutionary | omicverse_omni | pass | 95 | 100 | 100 | 100 | 100 | 99.0 |

| | | | | | | | | | | |
|---|---|---|---|---|---|---|---|---|---|---|
| | | Analysis; Whole Genome Sequencing (WGS) | | | | | | | | |
| bix-14-q1 | DNA-level | Genomic Variant Analysis; Genomics | omicverse_omni | pass | 100 | 100 | 100 | 100 | 100 | 100.0 |
| bix-17-q2 | DNA-level | Genomic Variant Analysis; Genomics | omicverse_omni | pass | 98 | 100 | 95 | 100 | 100 | 98.6 |
| bix-20-q3 | DNA-level | Genomic Variant Analysis; Genomics | omicverse_omni | pass | 90 | 95 | 90 | 100 | 90 | 93.0 |
| bix-28-q3 | DNA-level | Genomics; Phylogenetics and Evolutionary Analysis; Whole Genome Sequencing (WGS) | omicverse_omni | pass | 95 | 100 | 100 | 100 | 95 | 98.0 |
| bix-34-q2 | DNA-level | Genomics; Phylogenetics and Evolutionary Analysis; Whole Genome Sequencing (WGS) | omicverse_omni | fail | 85 | 90 | 100 | 95 | 100 | 94.0 |
| bix-34-q5 | DNA-level | Genomics; Phylogenetics and Evolutionary Analysis; Whole Genome Sequencing (WGS) | omicverse_omni | pass | 85 | 90 | 90 | 90 | 95 | 90.0 |
| bix-35-q1 | DNA-level | Genomics; Phylogenetics and Evolutionary Analysis; Whole Genome Sequencing (WGS) | omicverse_omni | pass | 95 | 100 | 100 | 100 | 100 | 99.0 |
| bix-35-q2 | DNA-level | Genomics; Phylogenetics and Evolutionary Analysis; Whole Genome Sequencing (WGS) | omicverse_omni | pass | 95 | 100 | 100 | 100 | 95 | 98.0 |
| bix-38-q1 | DNA-level | Genomics; Phylogenetics and Evolutionary Analysis; Whole Genome Sequencing (WGS) | omicverse_omni | pass | 95 | 100 | 95 | 95 | 100 | 97.0 |
| bix-45-q1 | DNA-level | Genomics; Phylogenetics; Whole Genome Sequencing (WGS) | omicverse_omni | fail | 80 | 100 | 50 | 100 | 100 | 86.0 |
| bix-47-q3 | DNA-level | Genomic Variant Analysis; Genomics | omicverse_omni | pass | 98 | 95 | 98 | 100 | 100 | 98.2 |
| bix-55-q1 | DNA-level | Genomics; Phylogenetics and Evolutionary Analysis; Whole Genome Sequencing (WGS) | omicverse_omni | pass | 80 | 95 | 100 | 100 | 95 | 94.0 |
| bix-16-q1 | Epigenomics & Functional | Functional Genomics; Transcriptomics | omicverse_omni | fail | 95 | 95 | 75 | 100 | 95 | 92.0 |
| bix-16-q3 | Epigenomics & Functional | Functional Genomics; Transcriptomics | omicverse_omni | pass | 95 | 95 | 100 | 100 | 100 | 98.0 |

| | | | | | | | | | | |
|---|---|---|---|---|---|---|---|---|---|---|
| bix-16-q4 | Epigenomics & Functional | Functional Genomics; Transcriptomics | omicverse_omni | pass | 90 | 100 | 95 | 100 | 100 | 97.0 |
| bix-52-q2 | Epigenomics & Functional | Epigenomics | omicverse_omni | pass | 95 | 100 | 100 | 95 | 100 | 98.0 |
| bix-52-q6 | Epigenomics & Functional | Epigenomics | omicverse_omni | pass | 95 | 95 | 95 | 100 | 100 | 97.0 |
| bix-52-q7 | Epigenomics & Functional | Epigenomics | omicverse_omni | pass | 90 | 95 | 95 | 100 | 100 | 96.0 |
| bix-6-q4 | Epigenomics & Functional | Functional Genomics | omicverse_omni | pass | 95 | 100 | 100 | 100 | 100 | 99.0 |
| bix-18-q1 | Other Modalities | Imaging | omicverse_omni | pass | 90 | 100 | 100 | 100 | 100 | 98.0 |
| bix-18-q3 | Other Modalities | Imaging | omicverse_omni | pass | 95 | 100 | 100 | 100 | 100 | 99.0 |
| bix-27-q5 | Other Modalities | Machine Learning and Artificial Intelligence; RNA-seq; Transcriptomics | bulk_rna_analyst | pass | 80 | 100 | 100 | 100 | 100 | 96.0 |
| bix-37-q1 | Other Modalities | Differential Expression Analysis; Proteomics; RNA-seq | bulk_rna_analyst | pass | 95 | 95 | 95 | 90 | 95 | 94.0 |
| bix-37-q4 | Other Modalities | Differential Expression Analysis; Proteomics; RNA-seq | bulk_rna_analyst | pass | 95 | 100 | 100 | 100 | 100 | 99.0 |
| bix-41-q5 | Other Modalities | Imaging | omicverse_omni | pass | 95 | 90 | 95 | 100 | 95 | 95.0 |
| bix-51-q2 | Other Modalities | Other | omicverse_omni | pass | 95 | 95 | 100 | 100 | 95 | 97.0 |
| bix-51-q8 | Other Modalities | Other | omicverse_omni | pass | 95 | 90 | 95 | 100 | 95 | 95.0 |
| bix-54-q7 | Other Modalities | Imaging | omicverse_omni | fail | 95 | 95 | 95 | 100 | 95 | 96.0 |
| bix-61-q2 | Other Modalities | Antimicrobial Resistance; Genomics; SNP Analysis; Whole Genome Sequencing (WGS) | omicverse_omni | pass | 95 | 100 | 95 | 100 | 100 | 98.0 |
| bix-61-q5 | Other Modalities | Antimicrobial Resistance; Genomics; SNP Analysis; Whole Genome Sequencing (WGS) | omicverse_omni | fail | 45 | 40 | 80 | 100 | 90 | 71.0 |

### Judge protocol

- Judge model: DeepSeek-v4-pro (deepseek-v4-pro), called via the OpenAI-compatible chat-completions API at temperature 0.0, max_tokens 4096, 3-retry exponential backoff.
- Unit of grading: one trajectory per BixBench-Verified-50 question (50 trajectories). When a question had multiple graded attempts, the BEST cell was selected with pass-priority (a harness-correct attempt outranks an incorrect one; ties broken by most recent mtime), matching the eval-report dedup convention.
- Judge input: a faithful compact render of the agent's sse.log — the user task, then every tool call with its name, arguments AND tool OUTPUT (the step messages' tool_calls[].result), interleaved with assistant text; head+tail truncated to ~80k chars only for very long runs. The judge additionally received the agent's final answer, the ground-truth ideal, and whether BixBench's own binary grader passed.
- Scoring: five 0-100 continuous process dimensions, defined for short-answer bioinformatics trajectories (distinct from the 6-dimension BiomniBench-DA narrative framework). Calibration anchors given to the judge: 100 = expert-level, 70 = competent with notable gaps, 40 = substantial deficiencies, 0 = unaddressed / fundamentally wrong.
- The dataset's native pass/fail is graded independently; the judge scores PROCESS quality only (an agent can be correct with weak process, or wrong with a clean attempt).
- Dimension definitions — Data exploration: did the agent inspect file structure / columns / shape / missing values before computing? Model selection (method_selection): was the chosen library / tool / statistical test canonical for the question's intent (e.g. DESeq2 vs t-test, PhyKIT vs custom code)? Execution correctness: did code run cleanly with intermediate values consistent

with the data (no silent off-by-one / unit errors)? Answer fidelity: does the final answer match the requested format (precision, unit, scientific notation, ordering, separator)? Iteration pattern: on error, did the agent diagnose-then-fix, or thrash / hallucinate?

- Scores are cached in bixbench_radar_grades_v2.jsonl; grader source is _grade_bixbench_trajectories_v2.py. Total judge spend ~$0.96.

#### Raw-trajectory validation

- Transcriptomics: Grounded (audited bix-53-q5, bix-22-q1, bix-30-q3). All five dimensions trace to concrete evidence; exact-match answers (bix-22-q1 'CD14', bix-30-q3 '0:0') are correctly rewarded, and the one low cell — bix-53-q5 answer fidelity = 20 — correctly penalizes a real format error (agent reported 10.0 instead of the ideal fraction 0.1).
- DNA-level: Mostly grounded (audited bix-34-q2, bix-45-q1, bix-14-q1, bix-38-q1). Data exploration, model selection, execution correctness and iteration pattern are well-supported by file-listing and tool-invocation evidence in the raw logs; bix-45-q1 answer fidelity = 100 is over-scored given the agent's answer (1.52e-56) differs from the ideal (7.70e-54) by ~100x (it ran PhyKIT rcv on all alignments rather than only shared orthologs).
- Epigenomics & Functional: Mostly grounded (audited all sampled bix-16-q1, bix-6-q4, bix-16-q4). Four process dimensions are well-supported, but bix-16-q1 answer fidelity = 100 is ungrounded: the agent reported CCND1 (computationally consistent with its own code) while the ideal answer is CDKN1A — a wrong-answer mismatch the judge did not catch.
- Other Modalities: Mostly grounded (audited bix-54-q7, bix-61-q5, bix-18-q3). Method, execution and iteration scores hold (including a verified R-path recovery in bix-54-q7), and bix-61-q5's low model-selection critique — no variant-quality filtering before computing Ts/Tv — is confirmed in the code; answer fidelity again downplays documented misses (bix-54-q7 ~3% outside the ideal range, bix-61-q5 4.5% off).
- Overall: direct inspection of raw trajectories supports the S3 claim that OmicOS maintained high process quality across all four categories. In all 13 audited trajectories the four process dimensions (data exploration, model selection, execution correctness, iteration pattern) were consistently traceable to concrete log evidence — file listings, canonical tool selection, error-free or successfully-recovered execution, and linear non-redundant iteration — and no process dimension was judged systematically inflated. The one consistent weakness is answer fidelity, which in a minority of cases rewards a well-formatted or plausible answer without fully penalizing a mismatch against the ideal (over-scored cells: bix-45-q1, bix-16-q1); the four process dimensions are therefore well-grounded, while answer fidelity should be read as a mild over-estimate of true final-answer correctness.

#### Caveats

- Single LLM judge: every score comes from one model (DeepSeek-v4-pro) at temperature 0.0 with no second grader or human adjudication, so scores are reproducible but judge-specific biases are not averaged out and no inter-rater agreement is reported.
- LLM-as-judge reliability: a raw-trajectory audit found the answer-fidelity dimension occasionally scores on format or plausibility rather than correctness against the ideal — bix-45-q1 scored 100 despite a ~100x magnitude error, and bix-16-q1 scored 100 despite the agent reporting CCND1 when the ideal is CDKN1A — so high answer-fidelity scores can overstate true answer correctness. The other four process dimensions were grounded in every audited case.
- Process scores are deliberately decoupled from the binary outcome: Transcriptomics is 100% Pass@1 yet has a lower category mean than Epigenomics & Functional (86% Pass@1), because answer-fidelity formatting penalties (percentage-vs-fraction) lower process scores without failing the harness, and vice-versa.
- Per-category n is small and uneven (7-18 tasks); 95% CIs are percentile bootstrap (10,000 resamples, seed 20260530) over the per-question scores, which respects the [0,100] bound rather than the normal approximation that would clip above 100. Category-level comparisons are low-power and sensitive to individual trajectories (e.g. Epigenomics & Functional has only 7 tasks); wide CIs (Transcriptomics answer fidelity, Other-Modalities model selection) reflect single low-scoring tasks, not a broad deficiency.
- Categories are collapsed from BixBench's 17 overlapping tags into 4 mutually-exclusive buckets by first-match-wins priority (Epigenomics/Functional > Other modalities > Transcriptomics > DNA-level); a task carrying multiple tags is assigned to exactly one bucket, so the per-bucket counts (14/18/7/11) partition the 50 questions.
- Best-cell-per-question deduplication with pass-priority means each question's score reflects its single best attempt, not the agent's average or worst case, which can flatter aggregates.
- Scores are bounded at 100 (continuous 0-100), so distributions are ceiling-censored for strong dimensions and means/CIs near the top of the scale are compressed, not symmetric.
- The raw-trajectory validation is a spot-check: only 13 of 50 graded trajectories were re-read against raw logs (DNA-level 4/18, Other Modalities 3/11, Transcriptomics 3/14, Epigenomics 3/7), not an exhaustive re-grade.
- The judge input was a ~80k-char head+tail render of each sse.log, and the dataset's binary pass/fail was provided alongside the answer; for very long trajectories the middle may be unseen, and the binary signal may anchor answer-fidelity scores.

#### *Supplementary Table 8 | BixBench-Verified-50 task taxonomy mapped to OmicOS capability families*

Each of the 50 BixBench-Verified-50 tasks was classified into a fine-grained analysis taxonomy node (from its question text and trajectory) and mapped to the OmicOS capability family / specialist agent whose toolset is its natural home. Table 6a is the per-node mapping; Table 6b is the coverage matrix over all eight OmicOS capability families.

##### Table 6a | Task taxonomy → OmicOS capability family

| BixBench taxonomy node | BixBench raw tags | n tasks | Task IDs | OmicOS capability family | Natural-home agent | Representative tools exercised | Agent run in eval |
|---|---|---|---|---|---|---|---|
| Bulk RNA-seq differential expression (DESeq2/pydeseq2, LFC shrinkage, covariates/batch) | Bulk RNA-seq differential expression - DEG count quantification with LFC shrinkage; Bulk RNA-seq differential expression with covariates - sex-stratified analysis; Bulk RNA-seq differential expression with batch correction and log-fold-change shrinkage; Bulk RNA-seq differential expression - gene-level log2 fold change extraction; miRNA differential expression - multiple testing correction comparison | 5 | bix-49-q4,bix-31-q2,bix-53-q2,bix-46-q4,bix-30-q3 | general_omics_analysis | bulk_rna_analyst | pydeseq2; pandas; scipy.stats; statsmodels | bulk_rna_analyst |
| DEG + pathway/GO enrichment (gseapy: GO BP, Reactome, KEGG, WikiPathways) | Bulk RNA-seq differential expression + GO Biological Process enrichment; Bulk RNA-seq differential expression + Reactome pathway enrichment with odds ratio extraction; KEGG pathway over-representation analysis - consensus pathways across multiple strains; KEGG pathway enrichment - condition-specific enrichment discovery; WikiPathways enrichment - disease/phenotype-specific pathway analysis; Pathway enrichment - overlap ratio computation between DEGs and pathway members; KEGG pathway | 7 | bix-24-q2,bix-43-q2,bix-32-q2,bix-26-q5,bix-53-q5,bix-43-q4,bix-26-q3 | general_omics_analysis | bulk_rna_analyst | gseapy; pydeseq2; pandas; mygene; scipy.stats | bulk_rna_analyst |

| | | | | | | | |
|---|---|---|---|---|---|---|---|
| | enrichment - gene-level pathway membership quantification | | | | | | |
| Phylogenomic tree-shape & evolutionary-rate statistics (PhyKIT: evo_rate, treeness, RCV, long_branch_score, tree_len) | Evolutionary rate comparison across orthologs (Mann-Whitney U statistic); Gene-specific evolutionary rate (PhyKIT evo_rate single gene); Tree-shape statistics (treeness/RCV relative treeness comparison); Tree-shape statistics (treeness/RCV threshold filtering); Tree-shape statistics (long branch scores/edge statistics); Tree-shape statistics (relative compositional variation - RCV Mann-Whitney U); Tree-shape statistics (tree length fold-change) | 7 | bix-35-q2,bix-35-q1,bix-11-q1,bix-11-q2,bix-28-q3,bix-45-q1,bix-38-q1 | phylogenomics | phylogenomics_analyst | phykit (evo_rate, treeness, rcv, long_branch_score, tree_len); scipy.stats.mannwhitneyu; pandas | omicverse_omni |
| Alignment quality metrics (parsimony-informative sites) | Alignment quality metrics (parsimony informative sites percentage); Alignment quality metrics (parsimony informative sites Mann-Whitney U); Alignment quality metrics (parsimony informative sites extremum); Alignment quality metrics (parsimony informative site count Mann-Whitney U) | 4 | bix-12-q2,bix-12-q4,bix-12-q5,bix-12-q6 | phylogenomics | phylogenomics_analyst | parsimony informative site detection; scipy.stats.mannwhitneyu; pandas | omicverse_omni |
| Phylogenetic distance metrics (patristic distances) | Phylogenetic distance metrics (patristic distances); Phylogenetic distance metrics (patristic distances single gene) | 2 | bix-34-q5,bix-34-q2 | phylogenomics | phylogenomics_analyst | newick tree parsing; patristic distance calculation | omicverse_omni |
| Ortholog completeness (BUSCO single-copy orthologs across proteomes) | Ortholog completeness (BUSCO single-copy orthologs across proteomes) | 1 | bix-55-q1 | phylogenomics | phylogenomics_analyst | BUSCO parsing (eukaryota_odb10) | omicverse_omni |
| Germline/somatic variant classification & counting (VAF filtering, | Germline variant classification (VAF-based filtering and annotation); Germline variant | 4 | bix-14-q1,bix-17-q2,bix-47-q3,bix-20-q3 | structural_biology | variant_analyst | pandas; VAF thresholding; variant location/annotation filtering; ClinVar classification | omicverse_omni |

| | | | | | | | |
|---|---|---|---|---|---|---|---|
| ClinVar/coding annotation) | counting (somatic CHIP variants by VAF and location); Variant counting (gene-level variant burden in individual); Variant classification (ClinVar pathogenicity annotation) | | | | | | |
| Expression-essentiality / CRISPR-screen correlation (Spearman, FDR) | Expression-essentiality correlation (Spearman rank); Expression-essentiality correlation with multiple testing correction; Expression-essentiality correlation (extreme negative); Functional screening assay (CRISPR) correlation | 4 | bix-16-q3,bix-16-q4,bix-16-q1,bix-6-q4 | general_omics_analysis | cancer_dependency_analyst | scipy.stats.spearmanr; statsmodels.stats.multitest.multipletests; pandas; numpy | omicverse_omni |
| Colony morphometrics from imaging assays (area, circularity, regression modeling) | Colony morphometrics - mean circularity measurement; Colony morphometrics - phenotype matching across strain mixtures; Colony morphometrics - percent reduction in area; Non-linear regression with spline and polynomial models - optimal prediction | 4 | bix-18-q1,bix-41-q5,bix-18-q3,bix-54-q7 | general_omics_analysis | tabular_genomics_analyst | pandas; R; lm; spline; quadratic/cubic polynomial models | omicverse_omni |
| Gene length vs expression correlation (Pearson, cell-type-specific) | Gene expression-genomic feature correlation - gene length vs expression association; Gene expression-genomic feature correlation - cell-type-specific length-expression association | 2 | bix-22-q4,bix-22-q1 | general_omics_analysis | bulk_rna_analyst | scipy.stats (Pearson); pandas | omicverse_omni |
| Methylation/epigenomic density & filtering (CpG density per chromosome) | Methylation/epigenomic density calculation; Methylation filtering/data preprocessing; Methylation/epigenomic density by chromosome | 3 | bix-52-q2,bix-52-q7,bix-52-q6 | general_omics_analysis | bulk_epigenomics_analyst | pandas; numpy | omicverse_omni |
| Clinical covariate regression (logistic regression, AIC, coefficients) | Logistic regression coefficient estimation; Logistic regression model selection via AIC | 2 | bix-51-q8,bix-51-q2 | general_omics_analysis | tabular_genomics_analyst | statsmodels.api; sklearn.linear_model.LogisticRegression; numpy; pandas | omicverse_omni |
| Proteomics differential abundance (protein fold | Log2 fold change calculation - proteomics differential abundance; Fold | 2 | bix-37-q4,bix-37-q1 | general_omics_analysis | bulk_rna_analyst | pandas | bulk_rna_analyst |

| | | | | | | | |
|---|---|---|---|---|---|---|---|
| change tumor vs normal) | change calculation - proteomics protein abundance | | | | | | |
| WGS read alignment & variant quality metrics (coverage depth, Ts/Tv ratio) | Whole genome sequencing coverage depth - read alignment statistics; Transition/transversion (Ts/Tv) ratio - SNP type classification | 2 | bix-61-q2,bix-61-q5 | general_omics_analysis | tabular_genomics_analyst | bwa-mem; samtools; vcf parsing; R; lm | omicverse_omni |
| Expression-matrix dimensionality reduction (PCA variance explained) | Dimensionality reduction via PCA - variance explained | 1 | bix-27-q5 | general_omics_analysis | bulk_rna_analyst | sklearn.decomposition.PCA; numpy; pandas | bulk_rna_analyst |

#### Table 6b | OmicOS capability-family coverage

| OmicOS capability family | Example agents | Exercised by BixBench-50 | n BixBench tasks | Note |
|---|---|---|---|---|
| data_acquisition | GEO-everything | no | 0 | BixBench tasks ship with the data capsule already provided; no GEO/SRA search, metadata fetch, or FASTQ download is required. The one WGS task (bix-61-q2) maps reads from supplied FASTQ but does not acquire data from public archives. |
| general_omics_analysis | bulk_rna_analyst; tabular_genomics_analyst; cancer_dependency_analyst; bulk_epigenomics_analyst; proteomics_analyst_pro; statistical_genetics_analyst | yes | 32 | Dominant family: bulk RNA-seq DEG + enrichment, proteomics fold change, methylation density, expression-essentiality/CRISPR-screen correlation, tabular clinical regression, WGS metrics, and colony morphometrics. |
| orchestration | omicverse_omni; analysis_strategist; vertical_agent_selector | yes | 33 | omicverse_omni (the orchestration generalist) actually executed 33 of the 50 tasks as the catch-all runner, including all phylogenomics, variant, epigenomics, correlation, and tabular tasks; counted here as the executing agent rather than the natural-home family. |
| phylogenomics | phylogenomics_analyst | yes | 14 | Strongly exercised: PhyKIT-based tree-shape/evo-rate stats, parsimony-informative-site alignment quality, patristic distances, and BUSCO ortholog completeness. Note: ran under omicverse_omni in the eval, not the phylogenomics_analyst specialist. |
| scientific_writing | scientific_writer; literature_pro; clinical_translator_pro; paper_critic; humanize | no | 0 | BixBench-Verified-50 asks for numeric/categorical analytic answers, not manuscript drafting, literature search, clinical translation, or peer review; |

| | | | | |
|---|---|---|---|---|
| | | | | no writing/literature task is present. |
| single_cell_analysis | single_cell_preprocessor; single_cell_annotator_pro; single_cell_downstream_analyst_pro; single_cell_trajectory_pro; cell_cell_communication_pro | no | 0 | No scRNA-seq/scATAC single-cell data in this 50-set. Bulk immune-cell-type tasks (bix-22) use bulk expression matrices grouped by cell type, not single-cell objects. |
| spatial_analysis | spatial_omics_orchestrator; he_to_st_predictor | no | 0 | No Visium/Xenium/CosMx spatial transcriptomics or H&E imaging data in this 50-set. |
| structural_biology | variant_analyst; structural_biologist; antibody_engineer; binder_designer; primer_design_assistant | partial | 4 | Exercised only via the variant_analyst arm: VAF/ClinVar germline-somatic variant classification and counting (bix-14/17/20/47). No protein-structure prediction, binder/antibody design, docking, or primer design is present. |

#### Interpretation (S4)

- BixBench-Verified-50 is analytically heterogeneous: the 50 tasks collapse into 15 distinct BixBench taxonomy nodes spanning 4 of the 8 OmicOS capability families (general_omics_analysis = 32 tasks, phylogenomics = 14, structural_biology/variant = 4, plus orchestration as the execution layer; with single_cell_analysis, spatial_analysis, data_acquisition, and scientific_writing absent because the corresponding scRNA/Visium/archive-download/writing data types are not in this 50-set). Within these families the tasks touch at least 8 natural-home specialist agents (bulk_rna_analyst, tabular_genomics_analyst, cancer_dependency_analyst, bulk_epigenomics_analyst, phylogenomics_analyst, variant_analyst, plus proteomics-style bulk work routed through bulk_rna_analyst), and exercise a wide concrete toolset (pydeseq2, gseapy, mygene, phykit, scipy.stats Mann-Whitney/Spearman/Pearson, statsmodels, sklearn PCA/LogisticRegression, bwa-mem/samtools, R lm/spline, BUSCO and patristic-distance parsing). Because the 90% leaderboard result is achieved across this breadth of analysis types, statistical methods, and data modalities, it reflects general omics-agent execution competence rather than mastery of a single optimized pipeline family.

#### Caveats

- The orchestration row (33 tasks) counts the agent that actually ran tasks in the eval (omicverse_omni as catch-all), which overlaps with the natural-home-family rows; it is not additive with the 50-task total. The taxonomy_rows partition (which sums to 50) is keyed on natural-home capability family, not the executing agent.
- In the eval, 33 of 50 tasks were executed by the omicverse_omni orchestration generalist rather than their natural-home specialist; only the bulk_rna_analyst arm (DEG/enrichment/proteomics/PCA = 17 tasks) ran on its native specialist. The natural-home-agent column therefore reflects intended routing, not realized routing.
- Proteomics fold-change tasks (bix-37) and PCA (bix-27) are listed under bulk_rna_analyst because that is the agent that ran them and their natural home in the catalog; proteomics_analyst_pro exists as a dedicated family member but was not the assigned runner.
- Two tasks carry harness_pass=false in the source records (bix-34-q2 patristic distance, bix-45-q1 RCV Mann-Whitney U); they are retained in the taxonomy counts as classified tasks regardless of pass/fail.
- Tool lists are taken verbatim from the per-task omicverse_tools_exercised records; no tools were inferred or added beyond what the records state.

### *Supplementary Table 9 | BiomniBench-DA harness x model cost-performance table*

Every system plotted in the BiomniBench-DA cost-performance space (Fig. 2c): agent harness, underlying model, inference mode, tool access, all-50-task mean rubric score and cache-adjusted cost per task. OmicOS rows are recomputed from the real run logs; external rows are transcribed from the published comparison chart. Full reproducibility columns (naive cost, measured cache hit, per-row pricing) are in the spreadsheet.

| Harness | Model | Inference mode | Tool access | All-50 rubric score | Cost/task (USD) | Cost basis |
|---|---|---|---|---|---|---|
| OmicOS (this work) | GPT-5.5 | Temperature unset (provider default); no forced reasoning-effort override; 1M context; max_tool_iterations=150 | Full OmicOS toolset via vertical_agent_selector -> vertical specialist agent: file_manager, run_python_code (Python kernel), shell, omicverse_lookup (function registry), notebook, plan/think/task/skill/memory; permission_mode=full, allow_shell, allow_file_write | **78.8** | 1.67 | cache-adjusted (this work, scripts/bench_cost.py) |
| OmicOS (this work) | DeepSeek v4-pro | Temperature unset (provider default); no forced reasoning-effort override; 1M context; max_tool_iterations=150 | Full OmicOS toolset via vertical_agent_selector -> vertical specialist agent: file_manager, run_python_code (Python kernel), shell, omicverse_lookup (function registry), notebook, plan/think/task/skill/memory; permission_mode=full, allow_shell, allow_file_write | 73.0 | 0.32 | cache-adjusted (this work, scripts/bench_cost.py) |
| OmicOS (this work) | GPT-5.4 | Temperature unset (provider default); no forced reasoning-effort override; 1M context; max_tool_iterations=150 | Full OmicOS toolset via vertical_agent_selector -> vertical specialist agent: file_manager, run_python_code (Python kernel), shell, omicverse_lookup (function registry), notebook, plan/think/task/skill/memory; permission_mode=full, allow_shell, allow_file_write | 67.7 | 0.76 | cache-adjusted (this work, scripts/bench_cost.py) |
| OmicOS (this work) | DeepSeek v4-flash | Temperature unset (provider default); no forced reasoning-effort override; 1M context; max_tool_iterations=150 | Full OmicOS toolset via vertical_agent_selector -> vertical specialist agent: file_manager, run_python_code (Python kernel), shell, omicverse_lookup (function registry), notebook, plan/think/task/skill/memory; permission_mode=full, allow_shell, allow_file_write | 66.0 | 0.02 | cache-adjusted (this work, scripts/bench_cost.py) |
| OmicOS (this work) | MiMo v2.5-pro | Temperature unset (provider default); no forced reasoning-effort override; 1M context; max_tool_iterations=150 | Full OmicOS toolset via vertical_agent_selector -> vertical specialist agent: file_manager, run_python_code (Python kernel), shell, omicverse_lookup (function registry), notebook, plan/think/task/skill/memory; permission_mode=full | 65.8 | 0.33 | cache-adjusted (this work, scripts/bench_cost.py) |

| | | | | | | |
|---|---|---|---|---|---|---|
| | | | , allow_shell, allow_file_write | | | |
| OmicOS (this work) | MiMo v2.5 | Temperature unset (provider default); no forced reasoning-effort override; 1M context; max_tool_iterations=150 | Full OmicOS toolset via vertical_agent_selector -> vertical specialist agent: file_manager, run_python_code (Python kernel), shell, omicverse_lookup (function registry), notebook, plan/think/task/skill/memory; permission_mode=full, allow_shell, allow_file_write | 62.3 | 0.16 | cache-adjusted (this work, scripts/bench_cost.py) |
| OmicOS (this work) | GPT-5.4-mini | Temperature unset (provider default); no forced reasoning-effort override; 1M context; max_tool_iterations=150 | Full OmicOS toolset via vertical_agent_selector -> vertical specialist agent: file_manager, run_python_code (Python kernel), shell, omicverse_lookup (function registry), notebook, plan/think/task/skill/memory; permission_mode=full, allow_shell, allow_file_write | 44.2 | 0.2 | cache-adjusted (this work, scripts/bench_cost.py) |
| Claude Code | Claude Opus 4.7 | max thinking | General-purpose agentic coding CLI (bash, file read/edit/write); no OmicVerse function registry | 73.3 | 2.5 | as published (harness-reported, assumed cache-adjusted) |
| Claude Code | Claude Opus 4.6 | max thinking | General-purpose agentic coding CLI (bash, file read/edit/write); no OmicVerse function registry | 69.5 | 1.35 | as published (harness-reported, assumed cache-adjusted) |
| Claude Code | Claude Sonnet 4.6 | max thinking | General-purpose agentic coding CLI (bash, file read/edit/write); no OmicVerse function registry | 66.5 | 0.93 | as published (harness-reported, assumed cache-adjusted) |
| Codex CLI | GPT-5.4 | xhigh reasoning | OpenAI terminal coding agent (sandboxed shell, apply_patch); no OmicVerse function registry | 68.7 | 2.3 | as published (harness-reported, assumed cache-adjusted) |
| Codex CLI | GPT-5.5 | xhigh reasoning | OpenAI terminal coding agent (sandboxed shell, apply_patch); no OmicVerse function registry | 67.6 | 4.3 | as published (harness-reported, assumed cache-adjusted) |
| Terminus-2 | Claude Opus 4.7 | max thinking | Terminal-Bench agent harness (terminal/tmux tool loop); no OmicVerse function registry | 64.1 | 0.85 | as published (harness-reported, assumed cache-adjusted) |
| Terminus-2 | Claude Sonnet 4.6 | max thinking | Terminal-Bench agent harness (terminal/tmux tool loop); no OmicVerse function registry | 62.5 | 0.93 | as published (harness-reported, assumed cache-adjusted) |
| Terminus-2 | Claude Opus 4.6 | max thinking | Terminal-Bench agent harness (terminal/tmux tool loop); no OmicVerse function registry | 63.2 | 1.9 | as published (harness-reported, assumed cache-adjusted) |

| | | | | | | |
|---|---|---|---|---|---|---|
| Terminus-2 | GPT-5.5 | default | Terminal-Bench agent harness (terminal/tmux tool loop); no OmicVerse function registry | 63.8 | 2.0 | as published (harness-reported, assumed cache-adjusted) |
| Terminus-2 | GLM-5.1 | default | Terminal-Bench agent harness (terminal/tmux tool loop); no OmicVerse function registry | 60.4 | 0.6 | as published (harness-reported, assumed cache-adjusted) |
| Terminus-2 | Qwen 3.6 | default | Terminal-Bench agent harness (terminal/tmux tool loop); no OmicVerse function registry | 59.6 | 1.0 | as published (harness-reported, assumed cache-adjusted) |
| Terminus-2 | Kimi K2.6 | default | Terminal-Bench agent harness (terminal/tmux tool loop); no OmicVerse function registry | 59.3 | 0.85 | as published (harness-reported, assumed cache-adjusted) |
| Terminus-2 | GPT-5.4 | default | Terminal-Bench agent harness (terminal/tmux tool loop); no OmicVerse function registry | 55.3 | 1.05 | as published (harness-reported, assumed cache-adjusted) |

#### Interpretation (S2)

- Among the plotted systems OmicOS with GPT-5.5 attains the highest all-50-task mean rubric score, 78.8 at a cache-adjusted $1.67/task, dominating Claude Code + Opus 4.7 (73.3 at $2.50), Codex CLI + GPT-5.5 xhigh (67.6 at $4.30) and Terminus-2 + GPT-5.5 (63.8 at $2.00). Holding the model fixed, the harness alone moves realized capability substantially: GPT-5.5 scores 78.8 under OmicOS but 67.6 (Codex CLI) and 63.8 (Terminus-2) under general-purpose harnesses — a +11 to +15 point swing at equal or lower cost — so realized biomedical-analysis capability depends strongly on the agent harness, not only on the underlying language model.

#### Cache-adjustment assumptions

- OmicOS cost is cache-adjusted and recomputed here from each run's per-call usage log (scripts/bench_cost.py). A BiomniBench-DA task is a long agent loop that resends the growing conversation every call, so cumulative input reaches 1-2M tokens/task; ~94-96% of that is an unchanged cached prefix (measured per model, see 'Measured cache hit').
- Because the usage log records only total input_tokens (cached+uncached merged), the uncached share is ESTIMATED: per cell the per-call input counts are sorted by time and only the positive increments are treated as fresh (uncached) input; the remainder is billed at the provider's cache-read price. This assumes a warm cache, making the cache-adjusted figure a best-case lower bound; the true cost sits between it and the naive cost (also tabulated).
- Cache-read pricing assumptions: OpenAI/codex models = 10% of input price (90%-off cache reads); DeepSeek cache-hit ~ near-zero; MiMo/MiniMax publishes no cache price, so 10% is assumed. List input/cache/output prices per 1M tokens are given per row (as of 2026-05). DeepSeek v4-pro used list price even though a 75%-off promo ran to 2026-05-31.
- External-harness costs (Claude Code, Codex CLI, Terminus-2) are transcribed verbatim from a published BiomniBench-DA cost/score comparison chart; their cost basis is each harness's own reported cost, assumed to be cache-adjusted real cost, and is NOT recomputed here. They are therefore not guaranteed to use an identical cache-adjustment convention to the OmicOS rows.

#### Caveats

- Score axis = mean rubric score over all 50 BiomniBench-DA tasks with infra-failure cells (no_output/error/serve_failed/judge_unavailable) excluded; the 6 documented failure-case tasks ARE included, for parity with the external chart. OmicOS rubric grading uses a fixed DeepSeek-v4-pro judge held constant across the campaign.
- OmicOS rows are single fresh runs per model against one held-constant agent/skill roster (vertical_agent_selector routing; omicverse_omni generalist deliberately excluded), not best-of-N. Two backends graded 49/50 and one 48/50 after infra exclusions (see 'n tasks scored').
- Cost and score are not measured under an identical environment for OmicOS vs the external harnesses (different tool stacks, sandboxes and cache conventions); the comparison is indicative of harness-level differences, not a controlled single-variable ablation.
- Inference-mode labels for external systems ('max thinking', 'xhigh reasoning') are as published on the source chart; OmicOS leaves temperature and reasoning effort at each provider's default.

#### *Supplementary Table 10 | BiomniBench-DA corrected 46-task vs original all-50 capability scores*

Per model: the original all-50 scores and the corrected 46-task capability scores side by side, with the six-dimensional breakdown on the corrected set. The corrected set removes the four documented benchmark-broken tasks (Supplementary Note 2). All numbers recomputed from the run logs.

| Model | Backend / provider | Original all-50 task-mean (%) | Original all-50 six-dim (%) | Corrected 46-task six-dim (%) | Δ (corrected – all-50 six-dim) | n (all-50, infra-excl) | n (46-task, infra-excl) | Data handling | Method selection | Statistical rigor | Biological interpretation | Scientific reasoning | Source reliability |
|---|---|---|---|---|---|---|---|---|---|---|---|---|---|
| GPT-5.5 | codex (OpenAI) | 78.8 | 81.3 | **82.8** | 1.5 | 50 | 46 | 84.4 | 87.6 | 79.9 | 61.3 | 83.2 | 100.0 |
| DeepSeek v4-pro | deepseek | 73.0 | 76.5 | 77.0 | 0.5 | 49 | 46 | 74.9 | 69.4 | 75.3 | 68.9 | 73.3 | 100.0 |
| MiMo v2.5-pro | custom_openai (MiniMax) | 65.8 | 68.4 | 69.0 | 0.6 | 49 | 45 | 67.2 | 66.7 | 69.0 | 75.7 | 66.7 | 68.6 |
| DeepSeek v4-flash | deepseek | 66.0 | 68.0 | 68.9 | 0.8 | 50 | 46 | 71.0 | 60.6 | 62.7 | 78.0 | 72.3 | 68.6 |
| MiMo v2.5 | custom_openai (MiniMax) | 62.3 | 64.8 | 65.7 | 0.9 | 49 | 45 | 62.5 | 66.7 | 61.9 | 67.9 | 66.7 | 68.6 |
| GPT-5.4 | codex (OpenAI) | 67.7 | 63.9 | 64.1 | 0.2 | 50 | 46 | 75.2 | 68.1 | 66.8 | 49.4 | 70.8 | 54.3 |
| GPT-5.4-mini | codex (OpenAI) | 44.2 | 42.7 | 43.1 | 0.3 | 48 | 44 | 46.3 | 42.9 | 41.4 | 52.0 | 50.1 | 25.7 |

##### Method

- Corrected capability set = the 50 BiomniBench-DA tasks minus the four documented benchmark-broken tasks {da-12-4, da-18-7, da-20-1, da-6-2} = 46 tasks; infra-failure cells (no_output/error/serve_failed/judge_unavailable) are additionally excluded per model, so the scored n is 44-46 per model (see 'n (46-task)').
- Six-dimensional scoring (scripts/bench_radar.py): each rubric criterion is mapped to one of six BiomniBench-DA dimensions (data handling, method selection, statistical rigor, biological interpretation, scientific reasoning, source reliability) by a keyword-priority heuristic; a dimension score = sum of earned points / sum of A-level (max) points over every (cell, criterion) pair in that dimension; the overall capability score = mean of the six dimension percentages. This is a point-weighted score and differs from the simple per-task rubric mean used for the cost-performance axis (Fig. 2c / Supplementary Table 7).
- 'Original all-50 task-mean' is the simple mean of per-task rubric scores over all 50 tasks (infra-excluded) — the Fig. 2c y-axis. 'Original all-50 six-dim' applies the identical six-dimensional scoring to all 50 tasks with NO task exclusion, isolating the effect of removing the four broken tasks (the Δ column). 'Corrected 46-task six-dim' is the published Fig. 2d score.
- Excluding the four broken tasks raises every model's six-dim score only modestly (Δ = +0.2 to +1.5 points) and does not change the top rank (GPT-5.5), confirming the correction is not what drives the leaderboard ordering.

#### *Supplementary Table 11 | Per-model, per-dimension values underlying the BiomniBench-DA capability radar (Fig. 2d)*

The six-dimensional capability scores plotted in Fig. 2d, with the rubric points they are computed from. Corrected 46-task set; reproduces the radar polygons exactly. The biological-interpretation column (shaded) is the panel-wide weak axis.

##### Table 11a | Radar values (per-dimension %)

| Model | Data handling | Method selection | Statistical rigor | Biological interpretation | Scientific reasoning | Source reliability | Overall (mean of 6) | n tasks |
|---|---|---|---|---|---|---|---|---|
| GPT-5.5 | 84.4 | 87.6 | 79.9 | 61.3 | 83.2 | 100.0 | **82.8** | 46 |

| | | | | | | | | |
|---|---|---|---|---|---|---|---|---|
| DeepSeek v4-pro | 74.9 | 69.4 | 75.3 | 68.9 | 73.3 | 100.0 | 77.0 | 46 |
| MiMo v2.5-pro | 67.2 | 66.7 | 69.0 | 75.7 | 66.7 | 68.6 | 69.0 | 45 |
| DeepSeek v4-flash | 71.0 | 60.6 | 62.7 | 78.0 | 72.3 | 68.6 | 68.9 | 46 |
| MiMo v2.5 | 62.5 | 66.7 | 61.9 | 67.9 | 66.7 | 68.6 | 65.7 | 45 |
| GPT-5.4 | 75.2 | 68.1 | 66.8 | 49.4 | 70.8 | 54.3 | 64.1 | 46 |
| GPT-5.4-mini | 46.3 | 42.9 | 41.4 | 52.0 | 50.1 | 25.7 | 43.1 | 44 |

#### Table 11b | Underlying rubric points (earned / max A-points)

| Model | Data handling | Method selection | Statistical rigor | Biological interpretation | Scientific reasoning | Source reliability |
|---|---|---|---|---|---|---|
| GPT-5.5 | 1099/1302 | 418/477 | 1491/1866 | 268/437 | 402/483 | 35/35 |
| DeepSeek v4-pro | 975/1302 | 331/477 | 1405/1866 | 301/437 | 354/483 | 35/35 |
| MiMo v2.5-pro | 818/1217 | 318/477 | 1218/1766 | 331/437 | 312/468 | 24/35 |
| DeepSeek v4-flash | 924/1302 | 289/477 | 1170/1866 | 341/437 | 349/483 | 24/35 |
| MiMo v2.5 | 785/1257 | 318/477 | 1134/1831 | 283/417 | 322/483 | 24/35 |
| GPT-5.4 | 979/1302 | 325/477 | 1246/1866 | 216/437 | 342/483 | 19/35 |
| GPT-5.4-mini | 577/1245 | 198/462 | 727/1758 | 222/427 | 237/473 | 9/35 |

#### Table 11c | Dimension key

| Dimension | What it aggregates | Total A-points (max, gpt-5.5 denom) |
|---|---|---|
| Data handling | Data loading/selection/subsetting, structure & annotation, baseline filtering, table construction | 1302 |
| Method selection | Methodology / model / normalization / QC / network / design-matrix choices and their appropriateness | 477 |
| Statistical rigor | Test choice, significance/FDR, metric computation, ranking, association/correlation, quantitative reporting accuracy | 1866 |
| Biological interpretation | Biological meaning/annotation/mechanism, phenotype interpretation, immunological/biological insight | 437 |
| Scientific reasoning | Translational implications, assumptions/limitations, candidate prioritisation, discussion & argument | 483 |
| Source reliability | Citations, references, source grounding, retraction awareness | 35 |

#### Interpretation (S3)

- GPT-5.5 leads on five of six dimensions — source reliability 100.0, method selection 87.6, data handling 84.4, scientific reasoning 83.2 and statistical rigor 79.9 — but drops to 61.3 on biological interpretation, its weakest axis and ~18 points below its next-lowest dimension. The same dip appears across the panel (GPT-5.4 49.4; even the DeepSeek/MiMo models, which trail overall, match or beat GPT-5.5 on biological interpretation: ds4-flash 78.0, mimo-v2.5-pro 75.7), indicating that domain-specific biological interpretation is a shared weakness not closed by scaling the strongest general model.

#### Method

- Radar value (percentage) for a (model, dimension) = sum of earned rubric points / sum of A-level (max) points, over every (task, criterion) pair whose criterion maps to that dimension, on the corrected 46-task set (50 tasks minus the four benchmark-broken tasks da-12-4, da-18-7, da-20-1, da-6-2; infra-failure cells additionally excluded per model). Overall = mean of the six dimension percentages. This reproduces the Fig. 2d radar polygons exactly (scripts/bench_radar.py).
- Each rubric criterion is assigned to one of the six dimensions by a documented keyword-priority heuristic on the criterion title+description (classify_dim in bench_radar.py); the full per-criterion assignment is auditable in reports/omicos_dim_map.csv.
- Dimensions carry very different rubric weight (Table 9c): statistical rigor draws the most A-points (~1.9k) and source reliability the fewest (~35, from few citation criteria), so source reliability saturates at 100% for several models on a small denominator and should not be read as equally information-rich as the high-load dimensions.
- Per-model denominators differ slightly because infra-failure cells are dropped per model (n = 44-46), so a dimension's max A-points is not identical across rows.

#### Caveats

- Scores are from a single fixed-judge (DeepSeek-v4-pro) rubric grading; the dimension axis is a post-hoc keyword classification of criteria, not a label shipped with the benchmark, so a few criteria could be argued into an adjacent dimension (audit via reports/omicos_dim_map.csv).

- OmicOS rows are single fresh runs per model against one held-constant agent/skill roster, not best-of-N; two-to-six cells per model are infra-excluded (n = 44-46).
- Source reliability rests on a small A-point denominator (~35), so its percentage is high-variance and a 100% there reflects a handful of citation criteria, not breadth comparable to the other axes.

#### *Supplementary Table 12 | BiomniBench-DA task ledger and per-task-type heat-map matrix*

All 50 BiomniBench-DA tasks with task type, included/excluded status, usable-cell and infra-failure counts, and per-model rubric score (10a); and the per-model x per-task-type mean matrix plotted in Fig. 2e (10b). Excluded rows shaded. Recomputed from grade records.

##### Table 12a | All 50 tasks

| Task ID | Task type | Status | Usable cells | Infra-fail cells | GPT-5.5 | DeepSeek v4-pro | MiMo v2.5-pro | DeepSeek v4-flash | MiMo v2.5 | GPT-5.4 | GPT-5.4-mini |
|---|---|---|---|---|---|---|---|---|---|---|---|
| da-1-4 | association-testing | included | 7 | 0 | 79.0 | 94.0 | 96.0 | 79.0 | 83.0 | 79.0 | 75.0 |
| da-8-1 | association-testing | included | 7 | 0 | 100.0 | 84.0 | 65.0 | 76.0 | 100.0 | 100.0 | 43.0 |
| da-8-2 | association-testing | included | 7 | 0 | 90.0 | 54.0 | 27.0 | 37.0 | 37.0 | 90.0 | 32.0 |
| da-9-7 | association-testing | included | 7 | 0 | 76.0 | 76.0 | 68.0 | 100.0 | 76.0 | 93.0 | 58.0 |
| da-13-3 | association-testing | included | 7 | 0 | 64.0 | 64.0 | 82.0 | 78.0 | 64.0 | 55.0 | 64.0 |
| da-14-3 | association-testing | included | 7 | 0 | 100.0 | 73.0 | 76.0 | 71.0 | 91.0 | 59.0 | 62.0 |
| da-14-8 | association-testing | included | 7 | 0 | 73.0 | 48.0 | 75.0 | 53.0 | 100.0 | 68.0 | 66.0 |
| da-15-7 | association-testing | included | 7 | 0 | 80.0 | 80.0 | 83.0 | 80.0 | 50.0 | 67.0 | 60.0 |
| da-8-3 | differential-expression | included | 7 | 0 | 68.0 | 34.0 | 66.0 | 49.0 | 74.0 | 46.0 | 61.0 |
| da-13-1 | differential-expression | included | 7 | 0 | 72.0 | 100.0 | 100.0 | 100.0 | 83.0 | 74.0 | 44.0 |
| da-15-1 | differential-expression | included | 7 | 0 | 87.0 | 80.0 | 80.0 | 80.0 | 70.0 | 100.0 | 70.0 |
| da-17-3 | differential-expression | included | 7 | 0 | 65.0 | 73.0 | 100.0 | 64.0 | 48.0 | 64.0 | 56.0 |
| da-19-1 | differential-expression | included | 7 | 0 | 72.0 | 100.0 | 63.0 | 71.0 | 53.0 | 53.0 | 54.0 |
| da-3-4 | mutation-analysis | included | 7 | 0 | 89.0 | 100.0 | 100.0 | 100.0 | 89.0 | 89.0 | 8.0 |
| da-3-5 | mutation-analysis | included | 7 | 0 | 79.0 | 47.0 | 58.0 | 87.0 | 58.0 | 49.0 | 92.0 |
| da-18-1 | mutation-analysis | included | 6 | 1 | 90.0 | 100.0 | 72.0 | 92.0 | infra | 82.0 | 72.0 |
| da-18-5 | mutation-analysis | included | 7 | 0 | 74.0 | 82.0 | 66.0 | 21.0 | 44.0 | 59.0 | 30.0 |
| da-25-1 | mutation-analysis | included | 7 | 0 | 78.0 | 48.0 | 62.0 | 64.0 | 78.0 | 50.0 | 8.0 |
| da-5-1 | multi-omic-integration | included | 7 | 0 | 79.0 | 75.0 | 71.0 | 50.0 | 56.0 | 24.0 | 40.0 |
| da-5-3 | multi-omic-integration | included | 7 | 0 | 95.0 | 95.0 | 95.0 | 82.0 | 65.0 | 70.0 | 47.0 |
| da-6-5 | multi-omic-integration | included | 7 | 0 | 100.0 | 100.0 | 0.0 | 49.0 | 50.0 | 92.0 | 27.0 |
| da-15-8 | multi-omic-integration | included | 7 | 0 | 70.0 | 60.0 | 45.0 | 70.0 | 57.0 | 60.0 | 70.0 |
| da-10-1 | predictive-modeling | included | 7 | 0 | 94.0 | 94.0 | 88.0 | 88.0 | 94.0 | 83.0 | 38.0 |
| da-10-3 | predictive-modeling | included | 7 | 0 | 100.0 | 100.0 | 61.0 | 75.0 | 68.0 | 89.0 | 70.0 |
| da-26-2 | predictive-modeling | included | 6 | 1 | 71.0 | 87.0 | 87.0 | 43.0 | 87.0 | 84.0 | infra |
| da-26-4 | predictive-modeling | included | 7 | 0 | 78.0 | 71.0 | 69.0 | 53.0 | 66.0 | 85.0 | 20.0 |
| da-1-3 | cell-composition | included | 7 | 0 | 80.0 | 97.0 | 92.0 | 95.0 | 86.0 | 74.0 | 71.0 |

| | | | | | | | | | | | |
|---|---|---|---|---|---|---|---|---|---|---|---|
| da-17-1 | cell-composition | included | 7 | 0 | 78.0 | 84.0 | 84.0 | 84.0 | 76.0 | 84.0 | 53.0 |
| da-17-5 | cell-composition | included | 7 | 0 | 100.0 | 64.0 | 82.0 | 82.0 | 55.0 | 74.0 | 0.0 |
| da-19-3 | chromatin-profiling | included | 7 | 0 | 87.0 | 100.0 | 57.0 | 60.0 | 69.0 | 90.0 | 47.0 |
| da-19-4 | chromatin-profiling | included | 7 | 0 | 65.0 | 70.0 | 60.0 | 50.0 | 55.0 | 55.0 | 30.0 |
| da-19-6 | chromatin-profiling | included | 6 | 1 | 87.0 | 87.0 | 87.0 | 62.0 | 22.0 | 59.0 | infra |
| da-4-1 | clustering | included | 7 | 0 | 77.0 | 61.0 | 59.0 | 75.0 | 52.0 | 52.0 | 52.0 |
| da-14-1 | clustering | included | 7 | 0 | 100.0 | 85.0 | 73.0 | 90.0 | 93.0 | 100.0 | 100.0 |
| da-16-1 | clustering | included | 7 | 0 | 93.0 | 88.0 | 72.0 | 90.0 | 80.0 | 66.0 | 68.0 |
| da-12-2 | pathway-enrichment | included | 7 | 0 | 82.0 | 60.0 | 35.0 | 60.0 | 63.0 | 82.0 | 20.0 |
| da-20-3 | pathway-enrichment | included | 6 | 1 | 68.0 | 73.0 | infra | 53.0 | 8.0 | 43.0 | 63.0 |
| da-20-4 | pathway-enrichment | included | 7 | 0 | 28.0 | 28.0 | 33.0 | 15.0 | 38.0 | 5.0 | 5.0 |
| da-13-5 | cross-cohort-comparison | included | 7 | 0 | 50.0 | 30.0 | 30.0 | 60.0 | 40.0 | 60.0 | 20.0 |
| da-13-6 | cross-cohort-comparison | included | 7 | 0 | 90.0 | 40.0 | 30.0 | 70.0 | 70.0 | 40.0 | 10.0 |
| da-4-6 | survival-analysis | included | 7 | 0 | 93.0 | 93.0 | 58.0 | 58.0 | 58.0 | 58.0 | 41.0 |
| da-9-1 | survival-analysis | included | 7 | 0 | 83.0 | 57.0 | 76.0 | 46.0 | 73.0 | 69.0 | 16.0 |
| da-11-1 | cell-cell-communication | included | 7 | 0 | 94.0 | 66.0 | 79.0 | 79.0 | 53.0 | 47.0 | 26.0 |
| da-15-2 | co-expression-networks | included | 7 | 0 | 90.0 | 46.0 | 26.0 | 25.0 | 31.0 | 35.0 | 26.0 |
| da-24-3 | gwas-eqtl | included | 7 | 0 | 60.0 | 93.0 | 75.0 | 80.0 | 45.0 | 85.0 | 10.0 |
| da-4-7 | tcr-repertoire | included | 7 | 0 | 85.0 | 60.0 | 58.0 | 51.0 | 58.0 | 85.0 | 45.0 |
| da-18-7 | mutation-analysis | EXCLUDED | 6 | 1 | 62.0 | infra | 82.0 | 57.0 | 52.0 | 65.0 | 42.0 |
| da-20-1 | clustering | EXCLUDED | 7 | 0 | 46.0 | 43.0 | 33.0 | 13.0 | 38.0 | 71.0 | 13.0 |
| da-12-4 | survival-analysis | EXCLUDED | 7 | 0 | 86.0 | 100.0 | 64.0 | 100.0 | 73.0 | 100.0 | 57.0 |
| da-6-2 | longitudinal-analysis | EXCLUDED | 7 | 0 | 35.0 | 35.0 | 25.0 | 35.0 | 25.0 | 20.0 | 40.0 |

#### Table 12b | Per-model × per-task-type mean (%)

| Model | association-testing (n=8) | differential-expression (n=5) | mutation-analysis (n=5) | multi-omic-integration (n=4) | predictive-modeling (n=4) | cell-composition (n=3) | chromatin-profiling (n=3) | clustering (n=3) | pathway-enrichment (n=3) | cross-cohort-comparison (n=2) | survival-analysis (n=2) | cell-cell-communication (n=1) | co-expression-networks (n=1) | gwas-eqtl (n=1) | tcr-repertoire (n=1) | Row mean (over types) |
|---|---|---|---|---|---|---|---|---|---|---|---|---|---|---|---|---|
| GPT-5.5 | 82.8 | 72.8 | 82.0 | 86.0 | 85.8 | 86.0 | 79.7 | 90.0 | 59.3 | 70.0 | 88.0 | 94.0 | 90.0 | 60.0 | 85.0 | 80.8 |
| DeepSeek v4-pro | 71.6 | 77.4 | 75.4 | 82.5 | 88.0 | 81.7 | 85.7 | 78.0 | 53.7 | 35.0 | 75.0 | 66.0 | 46.0 | 93.0 | 60.0 | 71.3 |
| MiMo v2.5-pro | 71.5 | 81.8 | 71.6 | 52.8 | 76.2 | 86.0 | 68.0 | 68.0 | 34.0 | 30.0 | 67.0 | 79.0 | 26.0 | 75.0 | 58.0 | 63.0 |
| DeepSeek v4-flash | 71.8 | 72.8 | 72.8 | 62.8 | 64.8 | 87.0 | 57.3 | 85.0 | 42.7 | 65.0 | 52.0 | 79.0 | 25.0 | 80.0 | 51.0 | 64.6 |
| MiMo v2.5 | 75.1 | 65.6 | 67.2 | 57.0 | 78.8 | 72.3 | 48.7 | 75.0 | 36.3 | 55.0 | 65.5 | 53.0 | 31.0 | 45.0 | 58.0 | 58.9 |
| GPT-5.4 | 76.4 | 67.4 | 65.8 | 61.5 | 85.2 | 77.3 | 68.0 | 72.7 | 43.3 | 50.0 | 63.5 | 47.0 | 35.0 | 85.0 | 85.0 | 65.5 |
| GPT-5.4-mini | 57.5 | 57.0 | 42.0 | 46.0 | 42.7 | 41.3 | 38.5 | 73.3 | 29.3 | 15.0 | 28.5 | 26.0 | 26.0 | 10.0 | 45.0 | 38.5 |

#### Method

- Task type = each task's [metadata].task_type from data/biomnibench-da/<task>/task.toml. Per-model score = the task's grade.json rubric score x100; a cell is treated as a infra-failure (and excluded from any mean) when grade_mode is one of no_output/error/judge_unavailable/serve_failed. No agent trajectories are read — scores come from the already-written grade records.

- Heat-map cell (Table 10b) = mean per-model rubric score over the INCLUDED tasks of that task type, infra cells dropped. The four benchmark-broken tasks (da-12-4, da-18-7, da-20-1, da-6-2; Supplementary Note 2) are marked EXCLUDED in 10a and omitted from 10b, so heat-map values reflect model capability rather than rubric-vs-question defects (S6). This reproduces the Fig. 2e cells exactly.
- The 50 tasks span 16 distinct task types; the heat map shows 15 columns because the only 'longitudinal-analysis' task (da-6-2) is one of the four excluded items, so that type has no included task. Excluded-task types: da-12-4 survival-analysis, da-18-7 mutation-analysis, da-20-1 clustering, da-6-2 longitudinal-analysis (each type still appears in 10b via its remaining included tasks, except longitudinal-analysis).
- Row mean (over types) in 10b is the unweighted mean of a model's task-type cells (the Fig. 2e row label); it differs from the all-50 task-mean (Suppl. Table 7/8) and the 46-task six-dim capability score (Suppl. Table 8/9) because it averages per-type means, not per-task scores.

#### Interpretation (S4)

GPT-5.5 holds high per-task-type means across association testing (83), mutation analysis (82), multi-omics integration (86), predictive modelling (86), cell composition (86), clustering (90), survival analysis (88), cell-cell communication (94), co-expression networks (90) and TCR repertoire (85), but is weaker on pathway enrichment (59) and GWAS (60) — small-n, methodology-sensitive types.

#### Interpretation (S5)

Task-type profiles differ by model: DeepSeek-v4-pro is strong on GWAS (93) yet weak on cross-cohort comparison (35) and co-expression networks (46); GPT-5.4-mini collapses on cross-cohort comparison (15), GWAS (10), cell-cell communication (26) and co-expression networks (26). Several of these are single-task (n=1) or two-task (n=2) types, so a cell is one-to-two graded runs.

#### Interpretation (S6)

Because the four benchmark-broken tasks are excluded symmetrically from every model before the per-type means are taken (Supplementary Note 2), the heat map is intended to reflect model capability rather than rubric-versus-question mismatch.

#### Caveats

- Many task types are small (10 of 15 shown types have n<=3; cell-cell communication, co-expression networks, GWAS-eQTL and TCR repertoire are n=1), so those heat-map cells are a single graded run and are high-variance — read them as anecdotes, not stable estimates.
- Single fixed-judge (DeepSeek-v4-pro) rubric grading; single fresh run per model (not best-of-N). Infra-failure cells are excluded per model, so a few per-type means rest on <7 models (see Usable cells / Infra-fail cells in 10a).
- Excluded tasks are retained in 10a (marked EXCLUDED with their per-model scores shown) for transparency; they are only removed from the 10b per-type means.

### *Supplementary Table 13 | Complete OmicBench task list (40 tasks)*

All 40 OmicBench tasks with task ID, data modality, analytical layer, difficulty, input fixture, expected output and programmatic grading checks. Grouped by modality. Difficulty shaded (hard/medium/easy).

| Task ID | Modality | Layer | Difficulty | Fixture | Expected output | Grading checks (n) | Grading checks |
|---|---|---|---|---|---|---|---|
| A01_qc_filter | scRNA-seq | A — scRNA preprocessing floor (PBMC) | medium | pbmc8k_raw.h5ad | QC-filtered AnnData with obs columns pct_counts_mt, pct_counts_ribo, pct_counts_hb (QC metrics); obs boolean columns with 2+ doublet score columns (method names containing 'doublet_score'); obs boolean column doublet_consensus; ~5500-7700 cells post-QC+doublet removal; ~12000-20939 genes remaining. | 6 | 1. mt_obs: obs column pct_counts_mt or alias mito_perc present; 2. ribo_obs: obs column pct_counts_ribo or alias ribo_perc/pct_ribo present; 3. cells_after_qc: cell count in [5500, 7700] after QC+doublet removal; 4. genes_filtered: gene count in [12000, 20939] after filtering; 5. mt_values_sane: pct_counts_mt values in [0, 100] range with |

| | | | | | | | |
|---|---|---|---|---|---|---|---|
| | | | | | | | variance (not all identical); 6. multi_doublet_methods: ≥2 doublet score columns detected, pairwise Spearman correlation ≥0.1, doublet_consensus obs column present |
| A02_normalize_log | scRNA-seq | A — scRNA preprocessing floor (PBMC) | easy | A01_qc_oracle.h5ad | Normalized AnnData with raw integer counts preserved in layer named 'counts' or 'raw_counts'; X matrix containing log1p-transformed normalized values in range [0, 15] (non-integer); per-cell sum of expm1(X) equals 10000±5%. | 4 | 1. counts_layer: layer 'counts' or 'raw_counts' present; 2. x_is_normalized: X values in [0, 15] with non-integer dtype (confirms log-transform); 3. per_cell_sum: per-cell expm1(X) sum ≈10000 (tolerance ±5%); 4. counts_int: layer counts/raw_counts contains integer-like values (no floating-point conversions) |
| A03_hvg | scRNA-seq | A — scRNA preprocessing floor (PBMC) | easy | A02_normalize_oracle.h5ad | AnnData with var boolean column 'highly_variable' or 'highly_variable_features' marking 2000±50 genes; all genes retained (not subset); dataset shape unchanged from input. | 3 | 1. hvg_present: var column 'highly_variable' or 'highly_variable_features' present; 2. hvg_count: exactly 2000±50 genes marked True in HVG column; 3. hvg_includes_canonical_markers: ≥5 canonical PBMC markers (CD3D, CD3E, CD4, CD8A, CD8B, CD79A, CD79B, MS4A1, CD14, LYZ, S100A8, S100A9, FCGR3A, NKG7, GNLY, GZMB, GZMK, CST3, FCER1A, IL7R, CCR7) appear in HVG-marked genes |
| A04_pca_neighbors_umap | scRNA-seq | A — scRNA preprocessing floor (PBMC) | medium | A03_hvg_oracle.h5ad | AnnData with obsm key X_pca (PCA on HVGs, 50 components); obsm key X_umap (2D UMAP, ~7710 cells × 2); uns key 'neighbors' containing neighborhood graph (15 neighbors in PCA space). | 5 | 1. pca: obsm key 'X_pca' present; 2. neighbors: uns key 'neighbors' present (neighborhood graph); 3. umap: obsm key 'X_umap' present; 4. umap_shape: X_umap has shape ~7710×2 (±50 cells tolerance); 5. umap_preserves_celltype: silhouette score of X_umap conditioned on cell_type obs ≥0.1 |
| A05_leiden_clustering | scRNA-seq | A — scRNA preprocessing floor (PBMC) | hard | A04_pca_neighbors_umap_oracle.h5ad | AnnData with obs column 'leiden' or 'clusters' containing final cluster labels (5-30 distinct clusters); obs columns matching pattern 'leiden_r*' / 'res*' / 'resolution_*' from ≥3 resolution sweeps; uns entry 'leiden_resolution' or 'selected_resolution' recording chosen resolution value. | 5 | 1. leiden_present: obs column 'leiden' or 'clusters' present; 2. cluster_count: final clustering has 5-30 unique cluster labels; 3. resolution_recorded: uns key 'leiden_resolution' or 'selected_resolution' or 'resolution_chosen' present with numeric value; 4. multi_resolution_stability: ≥3 obs columns matching leiden_r*/res*/resolution_* pattern detected, pairwise ARI between resolutions ≥0.4 (proves multi-resolution sweep with stable clustering); 5. ari_vs_celltype: ARI of |

| | | | | | | | |
|---|---|---|---|---|---|---|---|
| | | | | | | | leiden obs vs ground-truth cell_type ≥0.3 |
| B01_full_pipeline | scRNA-seq | B — scRNA-seq downstream workflows | hard | pbmc8k_raw.h5ad | Complete PBMC preprocessing-to-clustering pipeline producing: obs columns pct_counts_mt/pct_counts_ribo/doublet_consensus (QC), leiden/clusters for 5-25 clusters, layer counts for raw integers, var column highly_variable for ~2000 HVGs, obsm keys X_pca/X_umap, uns leiden_resolution; ~5500-8000 cells post-QC. ARI vs cell_type ground-truth >= 0.25. | 11 | 11 checks: (1) mt_obs: obs has pct_counts_mt or mito_perc; (2) ribo_obs: obs has pct_counts_ribo/ribo_perc/pct_ribo; (3) leiden_present: obs has leiden or clusters; (4) counts_layer: layer has counts or raw_counts (integer); (5) hvg: var has highly_variable or highly_variable_features (~2000 count); (6) pca: obsm has X_pca; (7) umap: obsm has X_umap; (8) cluster_count: 5 <= nunique(leiden/clusters) <= 25; (9) cells_after_full_qc: shape[0] in [5500,8000]; (10) resolution_chosen: uns has leiden_resolution/selected_resolution/resolution_chosen; (11) ari_vs_celltype: clustering_ari(leiden vs cell_type) >= 0.25 |
| B02_subset_recluster | scRNA-seq | B — scRNA-seq downstream workflows | hard | A05_leiden_oracle.h5ad | T-cell subset with new sub-clustering: obs column t_subcluster/subcluster/leiden_sub/leiden with 3-12 subclusters; subset size 1000-1600 cells; all cells contain 'T cell' substring in cell_type obs; subclusters split by CD4 vs CD8A expression (>=1 CD4-dominant and >=1 CD8A-dominant). | 5 | 5 checks: (1) subset_size: shape[0] in [1000,1600]; (2) only_t_cells: all obs[cell_type] values contain substring 'T cell'; (3) tsubcluster_present: obs has t_subcluster/subcluster/leiden_sub/leiden; (4) subcluster_count: 3 <= nunique(t_subcluster) <= 12; (5) cd4_cd8_split: subcluster_marker_split(CD4 vs CD8A) — requires >=1 CD4-dominant cluster and >=1 CD8A-dominant cluster |
| B03_diff_expression | scRNA-seq | B — scRNA-seq downstream workflows | hard | A05_leiden_oracle.h5ad | Cluster marker genes via >=2 distinct DE methods (e.g., Wilcoxon + t-test + COSG) with results stored in separate uns entries; standard uns[rank_genes_groups] populated with 'names' and 'pvals_adj' subkeys; top-20 genes per cluster show >=0.3 Jaccard overlap across methods; >=3 clusters have canonical PBMC markers in top-10 genes. | 4 | 4 checks: (1) rank_present: uns has rank_genes_groups; (2) rank_substructure: uns[rank_genes_groups] dict has 'names' and 'pvals_adj' subkeys; (3) multi_method_de_consistency: >=2 DE method results in uns with >=0.3 mean Jaccard overlap in top-20 genes per cluster; (4) top_markers_canonical_per_cluster: >=3 clusters have >=1 canonical PBMC marker (from CD3D/CD8A/CD79A/MS4A1/CD14/LYZ/NKG7/GNLY/etc.) in top-10 genes |
| B04_multi_method_batch_correction | scRNA-seq | B — scRNA-seq downstream workflows | hard | neurips2021_batch_all.h5ad | Batch correction via >=2 distinct methods (e.g., Harmony + Combat/Scanorama) stored in obsm/layer keys matching pattern (harmony\|combat\|scanorama\|scvi\|mnn\|bbknn\|corrected); | 4 | 4 checks: (1) batch_obs: obs has batch column; (2) batch_separation_reduced: batch-silhouette(corrected embedding) - batch-silhouette(X_pca) >= 0.02 drop (at |

<table>
<tr><th></th><th></th><th></th><th></th><th></th><th></th><th></th><th></th></tr>
<tr><td></td><td></td><td></td><td></td><td></td><td>obs[batch] column preserved; corrected embeddings show batch-silhouette drop >=0.02 from X_pca baseline, cell-type silhouette >=50% preserved, kNN-batch-mixing LISI score >=0.60.</td><td></td><td>least one corrected key); (3) celltype_silhouette_preserved: silhouette(corrected | cell_type) >= 0.5 * silhouette(X_pca | cell_type); (4) kbet_lisi_quantitative: kNN-batch-mixing LISI score >=0.60 on corrected embedding (15-NN mixing of different batch neighbors, normalized)</td></tr>
<tr><td>B05_multi_method_annotation</td><td>scRNA-seq</td><td>B — scRNA-seq downstream workflows</td><td>hard</td><td>A05_leiden_oracle.h5ad</td><td>Cell-type annotations via >=2 distinct methods with obs columns matching pattern cell_type_<method>/annotation_<method> (e.g., cell_type_cosg, annotation_scsa); family-normalized pairwise agreement >=60% across methods (normalizes CD4 vs T-cell to same family).</td><td>1</td><td>1 check: multi_annotation: >=2 obs columns matching (cell_type|celltype|annotation)_[a-zA-Z0-9_]+ (excluding bare cell_type, predicted_celltype, ground_truth columns); family-normalized pairwise agreement >=0.60 (CD4 T and CD8 T both map to t_cell family, etc.)</td></tr>
<tr><td>B06_reference_annotation</td><td>scRNA-seq</td><td>B — scRNA-seq downstream workflows</td><td>medium</td><td>pbmc8k_raw.h5ad</td><td>Reference-based label transfer via 70/30 split: obs column cell_type_predicted/predicted_celltype/celltype_pred/cell_type_transfer populated on query half (non-empty predictions); obs[cell_type] retained on reference half; 3-50 unique predicted labels (PBMC cell-type diversity); reconstructed single AnnData with both halves.</td><td>3</td><td>3 checks: (1) predicted_obs: obs has cell_type_predicted/predicted_celltype/celltype_pred/cell_type_transfer; (2) ref_celltype_present: obs has cell_type (ground-truth on reference half); (3) predicted_coverage: 3 <= nunique(cell_type_predicted) <= 50 (diverse predictions, not collapsed to single type)</td></tr>
<tr><td>B08_aucell_pathway_scoring</td><td>scRNA-seq</td><td>B — scRNA-seq downstream workflows</td><td>medium</td><td>pbmc8k_raw.h5ad</td><td>Per-cell gene-set activity scores in obs columns matching ^(aucell|score|signature|auc)_; >=2 scores for T-cell and B-cell families; T-cell score peaks in T-cell family (z-gap >=0.3), B-cell score peaks in B-cell family (z-gap >=0.3).</td><td>3</td><td>3 checks: (1) score_columns_regex: obs has >=1 column matching ^(aucell|score|signature|auc)_; (2) score_count_matching: >=2 obs columns match ^(aucell|score|signature|auc)_ pattern; (3) tb_celltype_specificity: >=2 scores are cell-type specific with (mean_family - mean_non-family) / std >= 0.3 (T-cell signature peaks in T-cell family, B-cell signature peaks in B-cell family)</td></tr>
<tr><td>B09_cnmf_gene_programs</td><td>scRNA-seq</td><td>B — scRNA-seq downstream workflows</td><td>hard</td><td>pbmc8k_raw.h5ad</td><td>Consensus NMF via real cNMF or omicverse.single.pycnmf: obsm key X_cnmf/cnmf_usages/gep_usages (n_cells x n_programs); uns key containing 'cnmf' and 'programs'/'spectra'/'gene_loadings'; obs column cnmf_dominant_program/dominant_program with 4-25 unique programs; real tool invocation evidence required (no hand-rolled aliasing).</td><td>5</td><td>5 checks: (1) cnmf_obsm: obsm has key matching X_cnmf|cnmf_usages|gep_usages; (2) cnmf_programs_uns: uns has key matching cnmf.*program|cnmf.*spectra|cnmf.*gene_loading|gep_program; (3) dominant_program: obs has cnmf_dominant_program or dominant_program; (4) n_programs: 4 <= nunique(cnmf_dominant_program) <= 25; (5) cnmf_real_call:</td></tr>
</table>

<table>
<tr><td></td><td></td><td></td><td></td><td></td><td></td><td></td><td>tool_output_evidence matching [cNMF]/[import cnmf]/ov.single.pycnmf/.factorize()/.consensus()/.load_results() patterns in tool messages</td></tr>
<tr><td>B10_gene_regulatory_network</td><td>scRNA-seq</td><td>B — scRNA-seq downstream workflows</td><td>hard</td><td>pancreas_endo_raw.h5ad</td><td>Gene regulatory network via real SCENIC/scenicplus or equivalent: obsm key matching regulon|tf_module|^X_auc|auc_ (n_cells x n_regulons); uns key matching regulon|network|grn with edge list or per-module gene membership; >=3 distinct TF modules identified.</td><td>3</td><td>3 checks: (1) regulon_obsm: obsm has key matching regulon|tf_module|^X_auc|auc_; (2) regulon_uns: uns has key matching regulon|network|grn; (3) n_regulons: obsm[regulon key].shape[1] >= 3 (at least 3 TF modules)</td></tr>
<tr><td>B11_scrna_cell_cell_communication</td><td>scRNA-seq</td><td>B — scRNA-seq downstream workflows</td><td>hard</td><td>pbmc8k_raw.h5ad</td><td>Cell-cell communication (CCC) via real library (cellphonedb/liana/commot): uns key matching ccc|lr_pairs|cell_communication|cellphonedb|liana|interactions with DataFrame containing sender/receiver cell-types, ligand/receptor genes, and significance scores (p-value/FDR); >=10 interactions; real tool invocation evidence required.</td><td>3</td><td>3 checks: (1) ccc_table: uns has key matching ccc|lr_pairs|cell_communication|cellphonedb|liana|interactions with sender/receiver/ligand/receptor columns; (2) celltype_obs: obs has cell_type/celltype/predicted_celltype; (3) ccc_real_call: tool_output_evidence matching cellphonedb/liana/commot/ov.single patterns in tool messages (no hand-rolled permutation tests)</td></tr>
<tr><td>F01_multi_mode_velocity</td><td>scRNA-seq</td><td>F — Trajectory & RNA velocity</td><td>hard</td><td>pancreas_endo_raw.h5ad</td><td>Two velocity-estimation modes (e.g., stochastic vs. dynamical, scvelo vs. graphvelo) with velocity matrices in layers velocity and velocity_alt; UMAP-projected velocity vectors in obsm key velocity_umap; und mode names recorded in uns key velocity_modes. Agent must subset to ~2000 HVGs before compute-heavy steps.</td><td>4</td><td>1. velocity_layer: at least one layer matching regex ^(velocity|velo)(_[A-Za-z0-9]+)?$ present; 2. velocity_umap_obsm: obsm key matching regex ^(velocity|velo)(_[A-Za-z0-9]+)?_umap$ present; 3. modes_directionally_agree: mean per-cell cosine similarity between first two velocity layers ≥0.05 (detects directional agreement, excludes random/fabricated modes); 4. velocity_flows_from_root: velocity vectors in UMAP space point outward from Ductal progenitor cluster with mean outward cosine ≥0.04 (validates biological plausibility of developmental flow).</td></tr>
<tr><td>F02_multi_method_trajectory</td><td>scRNA-seq</td><td>F — Trajectory & RNA velocity</td><td>hard</td><td>pancreas_endo_raw.h5ad</td><td>Two distinct pseudotime estimates from different algorithms (obs columns containing 'pseudotime') both anchored at Ductal root; ≥2 terminal/branch states identified (via uns key containing 'terminal'/'branch'/'fate'/'macrostate' or obs categorical 'terminal_state');</td><td>4</td><td>1. two_pseudotime_obs: ≥2 distinct pseudotime obs columns present; 2. pseudotime_anchored_at_root: median pseudotime in Ductal cluster is ≥10% of overall range below median in non-Ductal clusters for all pseudotime methods; 3. branch_detection: ≥2 terminal/branch</td></tr>
</table>

| | | | | | | | |
|---|---|---|---|---|---|---|---|
| | | | | | trajectory method names recorded in uns key trajectory_methods. | | states detected (via uns regex match or obs categorical count); 4. pseudotime_methods_agree: pairwise Spearman correlation between all pseudotime columns ≥0.4 (minimum pair ≥0.4, confirming trajectory signal is consistent across methods). |
| F03_paga_pseudotime | scRNA-seq | F — Trajectory & RNA velocity | hard | pancreas_endo_raw.h5ad | PAGA cluster connectivity stored in uns['paga'] with 'connectivities' subkey; DPT pseudotime in obs column (dpt_pseudotime or pseudotime alias); uns['neighbors'] containing neighborhood graph; obs['clusters'] containing cluster labels if not present. | 4 | 1. neighbors: uns key 'neighbors' present (graph structure); 2. paga: uns key 'paga' present; 3. paga_connectivities: uns['paga'] contains 'connectivities' subkey (cluster connectivity matrix); 4. pseudotime_obs: obs column matching alias group [dpt_pseudotime, pseudotime] present (DPT pseudotime values). |
| F04_multi_backend_pseudotime | scRNA-seq | F — Trajectory & RNA velocity | hard | pancreas_endo_raw.h5ad | ≥2 distinct pseudotime obs columns (names ending in '_pseudotime', '_dpt', or starting with 'pseudotime_') from different trajectory-inference backends, both anchored at Ductal root cluster; pairwise Spearman ≥0.4 between methods confirming convergence on real developmental signal. | 4 | 1. clusters_obs: obs['clusters'] present (cluster labels); 2. two_pseudotime_obs: ≥2 obs columns matching regex 'pseudotime$\|_dpt$\|^pseudotime_' (multiple methods); 3. pseudotime_anchored_at_root: median pseudotime in Ductal ≥10% of range below non-Ductal medians for all pseudotime columns; 4. pseudotime_methods_agree: pairwise Spearman ≥0.4 between all pseudotime columns (minimum pair ≥0.4, confirming each new method agrees with others). |
| C01_visium_spatial_domain | Spatial | C — Spatial transcriptomics (Visium/Visium-HD) | hard | visium_dlpfc_151676 | Spatial domain clustering on DLPFC Visium data (151676) with discrete spatial-aware clusters in obs column (4-12 domains), spatial coordinates in obsm key 'spatial', and ground-truth cortical-layer labels loaded into obs 'ground_truth_layer' for evaluation. | 6 | spatial_obsm: obsm['spatial'] key must exist with 2D coordinates; ground_truth_loaded: obs['ground_truth_layer'] column required to contain manual cortical-layer labels; spatial_domain: obs column (alias clusters/leiden/spatial_clusters) contains discrete cluster assignments; n_domains: unique values in spatial_domain ∈ [4,12] capturing ~7 layers + white matter; spatial_silhouette: silhouette score of spatial_domain clusters computed on obsm['spatial'] coordinates ≥ 0.05 (real spatial methods yield ~0.10-0.30; random/expression-only ~0); ari_vs_ground_truth: Adjusted Rand Index of predicted spatial_domain vs ground_truth_layer ≥ |

| | | | | | | | |
|---|---|---|---|---|---|---|---|
| | | | | | | | 0.20 (spatial-aware methods on DLPFC achieve this; expression-only clustering fails) |
| C02_visium_hd_svg | Spatial | C — Spatial transcriptomics (Visium/Visium-HD) | hard | visium_hd_8um | Spatially-variable genes (SVGs) identified on Visium HD colon cancer tissue (8μm binning) with spatial coordinates in obsm['spatial'] and ≥100 SVGs marked as True in a boolean var column (aliases: space_variable_features/spatially_variable/svg/spatial_variable_genes). | 3 | spatial_obsm: obsm['spatial'] key present with 2D bin-level coordinates; svg_marker: var column matching aliases (space_variable_features/spatially_variable/svg/spatial_variable_genes) exists as boolean; svg_count: number of True values in the svg marker column ≥ 100 (real Visium HD SVG analyses identify hundreds of genes; threshold catches near-empty results) |
| C04_spatial_trajectory_stt | Spatial | C — Spatial transcriptomics (Visium/Visium-HD) | hard | visium_dlpfc_151676 | Spatially-aware trajectory inference on DLPFC Visium respecting cortical layer structure with spatial coordinates in obsm['spatial'], continuous per-spot pseudotime in obs (aliases: spatial_pseudotime/stt_pseudotime/pseudotime_spatial/pseudotime), discrete spatial domains in obs (4-12 unique values; aliases: region/spatial_domain/domain/clusters), and a transition matrix or tensor in uns matching pattern 'transition\|tensor\|stt\|trajectory' capturing inter-domain state transitions. | 6 | spatial_obsm: obsm['spatial'] key present; pseudotime_obs: obs column matching aliases contains numeric per-spot pseudotime values; spatial_domain_obs: obs column matching aliases contains discrete domain labels; domain_unique_count: unique values in spatial_domain ∈ [4,12] representing ~7 layers + white matter; transition_uns: uns key matching regex pattern (transition\|tensor\|stt\|trajectory) holds per-domain transition statistics; stt_real_call: trajectory tool output from minisweagent_trajectory.json must contain evidence of real STT/spatial-transition-tensor library invocation (patterns: ov.space.STT, import stt, [STT] markers) — hand-rolled stats with labeled dominions not acceptable |
| C05_spatial_celltype_deconvolution | Spatial | C — Spatial transcriptomics (Visium/Visium-HD) | hard | spatial_lymph_node | Cell-type fractions deconvolved from lymph-node spatial transcriptomics using scRNA reference, with spatial coordinates in obsm['spatial'], per-spot fractions for ≥5 cell types in obs columns or uns table (pattern: frac_\|_fraction\|cell_type_frac) where values sum to ~1.0 across ≥5 distinct cell-type columns. | 3 | spatial_obsm: obsm['spatial'] key present with 2D spot coordinates; fractions_present: obs columns or uns DataFrame matching pattern (frac_\|_fraction\|cell_type_frac) exist with ≥5 columns representing distinct cell types; per-sample/spot fractions sum to 1.0 ± 0.1 tolerance; min_n_celltypes: ≥5 cell-type columns with nonzero values (not all-zero fractions); deconv_real_call: tool output evidence must show invocation of real deconvolution library (patterns: cell2location.run_colocation, |

| | | | | | | | |
|---|---|---|---|---|---|---|---|
| | | | | | | | cell2location.cluster, cell2location.models, Tangram, CARD, DestVI, ov.space.Deconvolution) — manually-computed marker-fraction overlays not acceptable |
| D01_multiome_joint_embedding | Multi-omics | D — Multi-omics integration (multiome) | hard | multiome_pbmc | AnnData with paired 10x Multiome RNA+ATAC data; joint embedding in obsm key X_mofa/X_glue/X_multivi/X_wnn/X_joint (≥5 factors), single-modality embeddings in obsm X_pca (RNA) and X_atac (ATAC), and cell-type annotations in obs (cell_type or cell_type_pred) with ≥5 distinct types. | 6 | joint_obsm: obsm key from [X_mofa, X_glue, X_multivi, X_wnn, X_joint] required; single_modality_present: obsm keys [X_pca or X_rna] required; celltype_obs: obs keys [cell_type, celltype, or cell_type_pred] required; joint_factors: obsm[X_mofa] must have ≥5 dimensions; factor_variance_concentrated: top-5 factors explain ≥30% of total variance; joint_outperforms_single: joint embedding cell-type silhouette exceeds best single modality by ≥0.02 |
| D02_peak_gene_linking | Multi-omics | D — Multi-omics integration (multiome) | hard | multiome_pbmc | AnnData with peak-to-gene linkage table in uns key peak_gene_links/peak_to_gene/gene_peak/regulatory_links; table includes columns for peak ID, gene symbol, correlation/score, and genomic distance; minimum ≥500 links with ≥30% within ±10 kb of TSS (promoter-proximal). | 2 | links_uns: uns key from [peak_gene_links, peak_to_gene, gene_peak, regulatory_links] required; links_count_and_promoter_frac: ≥500 peak-gene links required AND ≥30% of links fall within ±10 kb of TSS |
| E01_bulk_deconvolution | Bulk RNA | E — Bulk RNA-seq & metabolomics | hard | bulk_dg | Bulk RNA-seq with per-sample cell-type fractions (≥3 cell types, sums ≈1) stored as obs columns or uns DataFrame from ≥2 distinct deconvolution methods (CIBERSORTx, BayesPrism, MuSiC, Bisque, SCDC, or similar). Cross-method per-cell-type fraction Pearson correlation ≥0.4 demonstrates method agreement. | 2 | fractions_present: per-sample cell-type fractions sum to ≈1 with ≥3 nonzero cell types (matches pattern frac_\|_fraction\|cell_type_frac in obs, or in a DataFrame in uns); multi_method_agreement: ≥2 deconv methods with per-cell-type fraction Pearson correlation ≥0.4 (tables identified by pattern fractions\|deconv\|frac_ in uns keys, shape=(n_samples, n_celltypes)) |
| E03_bulk_de_multi_method | Bulk RNA | E — Bulk RNA-seq & metabolomics | hard | dg_bulk_normal.h5ad | Bulk RNA-seq with rank_genes_groups uns entry containing 'names' and 'pvals_adj' subkeys; ≥2 distinct DE method result tables (uns keys matching pattern rank_genes\|de_\|wilcoxon\|ttest\|deseq\|limma\|pydeg\|edger) with per-comparison top-30 gene Jaccard overlap ≥0.15 (accommodates small-n Wilcoxon saturation; DESeq2+limma+t-test typically clear 0.30). | 3 | rank_present: uns['rank_genes_groups'] exists; rank_substructure: uns['rank_genes_groups'] contains dict subkeys 'names' and 'pvals_adj'; multi_method_de_consistency: ≥2 DE method tables extracted, mean pairwise top-30 gene Jaccard ≥0.15 (per-cluster Jaccard for single-cell rank_genes_groups, or bulk DataFrame Jaccard) |
| E04_bulk_wgcna | Bulk RNA | E — Bulk RNA-seq & metabolomics | hard | wgcna_5xfad | WGCNA co-expression analysis on ~2000 HVG bulk RNA-seq (192 samples × genes): var column | 4 | module_assignment: var column exists matching pattern module_color\|wgcna_module\|module\|modu |

| | | | | | | | |
|---|---|---|---|---|---|---|---|
| | | | | | with ≥3 distinct modules (names like 'turquoise', 'blue', 'brown', or module IDs); uns DataFrame keyed 'module_trait_corr' (or variants) with modules×traits correlation matrix (including Genotype and Age traits). | | les\|module_id\|module_label; n_modules: var module column has ≥3, ≤50 unique values; module_trait_table: uns key exists matching pattern module_trait; wgcna_real_call: trajectory contains evidence of real PyWGCNA/omicverse invocation (e.g. PyWGCNA, soft threshold power, TOM matrix, cutreeDynamic, goodSamplesGenes, mergeCloseModules markers) |
| E06_pathway_enrichment | Bulk RNA | E — Bulk RNA-seq & metabolomics | medium | dg_bulk_normal.h5ad | Bulk RNA-seq with rank_genes_groups DE result; uns DataFrame (keyed with pattern enrich\|gsea) containing ≥2 distinct databases/sources and ≥1 pathway row with adjusted p-value ≤0.10 (FDR/padj/qvalue column required). | 2 | rank_present: uns['rank_genes_groups'] exists; enrichment_table: ≥1 uns key matches pattern enrich\|gsea (grader checks for 2+ database sources and ≥1 row with FDR ≤0.10 via pathways_enrichment_quality check, though exact check name not in task spec) |
| E07_ppi_network | Bulk RNA | E — Bulk RNA-seq & metabolomics | medium | dg_bulk_normal.h5ad | Bulk RNA-seq with rank_genes_groups DE result on top-30 genes; uns entries for: (1) PPI interaction table (DataFrame with source/protein_a, target/protein_b, confidence/score columns), containing ≥5 edges; (2) hub/degree summary table with per-gene centrality metrics. | 4 | rank_present: uns['rank_genes_groups'] exists; ppi_table: ≥1 uns key matches pattern ppi\|interaction\|network (must be DataFrame with ≥5 rows and source/target/confidence columns); hub_summary: ≥1 uns key matches pattern hub\|degree\|central (per-gene centrality summary); ppi_real_call: trajectory contains evidence of real STRINGdb/PPI invocation (e.g., stringdb import, ov.bulk.string_interaction, requests to string-db.org, [STRING] marker) |
| E05_bulk_metabol_multivariate | Metabolomics | E — Bulk RNA-seq & metabolomics | hard | metabol_cachexia.csv | Metabolomics AnnData (77 patients × 63 metabolites) with obs['group'] (or alias 'class') containing exactly 2 distinct values (cachexic/control); ≥2 distinct multivariate embeddings in obsm (e.g., X_pca + X_plsda); X_plsda silhouette score on group separation ≥0.15 (accommodates small-sample metabolomics baseline ~0.15). | 5 | group_obs: obs column present matching group\|class\|Muscle loss\|muscle_loss; two_distinct_groups: obs[group] column has exactly 2 unique values; multivariate_obsm: ≥1 obsm key matches regex X_(pca\|plsda\|opls.?da\|pls.?da\|pls); two_distinct_methods: ≥2 obsm keys match regex X_(pca\|plsda\|opls.?da\|pls.?da\|pls\|tsne\|umap\|fa); supervised_separation: obsm['X_plsda'] silhouette score by group ≥0.15 |
| G01_16s_diversity | Microbiome 16S | G — 16S microbiome analysis | medium | mothur_sop_16s.h5ad | Per-sample alpha diversity metrics (shannon, observed_otus, simpson) in obs; beta-diversity distance matrix | 4 | alpha_present: one or more of shannon/observed_otus/observed_asvs/simpson present and non-zero in obs; beta_present: |

| | | | | | | | |
|---|---|---|---|---|---|---|---|
| | | | | | (Bray-Curtis, UniFrac, or other) in obsp or uns; statistical test results (Kruskal-Wallis/Mann-Whitney for group alpha-diversity and PERMANOVA/ANOSIM for group beta-diversity separation) in uns dict entries. | | pairwise distance matrix at key matching beta_distance/bray_curtis/unifrac/weighted_unifrac in uns or obsp; alpha_group_test: group-level Shannon test result stored in uns with key matching pattern 'alpha_test\|kruskal\|mann_whitney\|group_diff', with numeric pvalue field; beta_permanova: PERMANOVA/ANOSIM test stored in uns with key matching pattern 'permanova\|anosim\|beta_test', pvalue ≤ 0.05 |
| G02_multi_method_da | Microbiome 16S | G — 16S microbiome analysis | hard | mothur_sop_16s.h5ad | Two or more differential-abundance (DA) result tables stored as uns DataFrames, each containing per-taxon p-values/FDR and log-fold-change statistics (Wilcoxon, DESeq2, ANCOM-BC, LEfSe, ALDEx2, or similar frameworks); method names identifiable in the uns key names (e.g. da_wilcoxon, da_deseq2, da_ancombc); top-ranked taxa should partially overlap across methods (Jaccard ≥ 0.20 on top-30 taxa). | 1 | multi_method_da_results: ≥2 DA result tables detected in uns matching pattern 'da_\|differential_abundance\|wilcox\|deseq\|ancombc\|aldex\|lefse'; each has ≥5 rows (taxa); pairwise Jaccard of top-30 significant taxa ≥ 0.20 across methods (captures agreement on real signal without requiring identical rankings) |
| G03_16s_ordination_pcoa | Microbiome 16S | G — 16S microbiome analysis | medium | mothur_sop_16s.h5ad | Bray-Curtis distance matrix in obsp key containing 'braycurtis' or 'beta_distance'; per-sample 2D-or-3D PCoA coordinates in obsm key containing 'pcoa'; per-axis variance-explained array in uns key containing both 'pcoa' and 'var' (e.g., [0.32, 0.18, 0.09]); PERMANOVA or ANOSIM test result (pvalue ≤ 0.05) in uns key matching pattern 'permanova\|anosim\|ordination_test' with fields pvalue and statistic. | 4 | beta_dist_present: pairwise distance matrix present in uns or obsp at key matching 'braycurtis\|bray_curtis\|beta_distance\|unifrac\|weighted_unifrac'; pcoa_obsm: at least one obsm key matching regex 'pcoa\|PCoA' present; pcoa_variance: at least one uns key matching regex 'pcoa.*var\|var.*pcoa\|variance_explained\|expl' present; group_separation: PERMANOVA/ANOSIM test in uns with key matching 'permanova\|anosim\|ordination_test\|beta_test', pvalue ≤ 0.05 |
| G04_16s_taxonomic_collapse | Microbiome 16S | G — 16S microbiome analysis | medium | mothur_sop_16s.h5ad | Genus-level and phylum-level abundance tables (collapsed sums of ASV counts) in uns DataFrames; ASV-level Shannon diversity in obs column 'shannon'; genus-level Shannon diversity in obs column matching regex 'shannon.*genus\|genus.*shannon'; for ≥60% of samples, ASV-level Shannon > genus-level Shannon (information collapses upon aggregation); top-5 most abundant genera | 6 | genus_table: uns key matching regex '(abundance\|collaps(e\|ed)\|table\|counts).*genus\|genus.*(abundance\|collaps\|table\|counts)' present; phylum_table: uns key matching regex '(abundance\|collaps(e\|ed)\|table\|counts).*phylum\|phylum.*(abundance\|collaps\|table\|counts)' present; asv_shannon: obs column 'shannon' present; genus_shannon: obs column matching regex |

<table>
<tr><th></th><th></th><th></th><th></th><th></th><th></th><th></th><th></th></tr>
<tr><td></td><td></td><td></td><td></td><td></td><td>stored as uns array matching regex 'top.*genus|top.*genera|top_taxa'.</td><td></td><td>'shannon.*genus|genus.*shannon' present; shannon_ordering: obs['shannon'] > obs[genus_shannon_col] in ≥60% of samples (min_fraction 0.6, comparison 'a_gt_b'); top_genera: uns key matching regex 'top.*genus|top.*genera|top_taxa' present</td></tr>
<tr><td>G05_phylogenetic_diversity</td><td>Microbiome 16S</td><td>G — 16S microbiome analysis</td><td>hard</td><td>mothur_sop_16s.h5ad</td><td>Phylogenetic tree in Newick format stored in uns key containing 'tree|phylogen|newick'; per-sample Faith's phylogenetic diversity in obs column 'faith_pd' (or 'pd_whole_tree' or 'phylogenetic_diversity'); UniFrac distance matrix (weighted or unweighted) in obsp key matching 'unifrac|weighted_unifrac|unweighted_unifrac'; PERMANOVA or ANOSIM test on UniFrac distance results in uns with key matching regex '(unifrac|tree).*(permanova|anosim|test)|(permanova|anosim).*(unifrac|tree)'.</td><td>4</td><td>tree_newick: uns key matching regex 'tree|phylogen|newick' present; faith_pd_obs: obs column matching any of 'faith_pd|pd_whole_tree|phylogenetic_diversity' present; unifrac_distance: UniFrac distance matrix in uns or obsp key matching any of 'unifrac|weighted_unifrac|unweighted_unifrac' with shape [n_samples, n_samples]; unifrac_test: uns key matching regex '(unifrac|tree).*(permanova|anosim|test)|(permanova|anosim).*(unifrac|tree)' present</td></tr>
<tr><td>L01_foundation_model_cell_embedding</td><td>FM (scLLM)</td><td>L — Single-cell foundation models (scLLM)</td><td>hard</td><td>pbmc8k_raw.h5ad</td><td>obsm key starting with X_ and identifying the FM backend (X_scgpt, X_geneformer, X_uce, X_cellplm, X_scfoundation, X_tabula, or similar); shape (n_cells, n_embed_dims) with n_embed_dims >= 16. Neighbors graph recomputed on FM embedding under uns['neighbors']. Cell-type silhouette on FM embedding >= 0.05.</td><td>5</td><td>4; fm_embedding_obsm: obsm key must match regex for foundation models (scgpt, geneformer, scfoundation, uce, cellplm, tabula, etc.); embedding_dim_at_least: embedding has >=16 dimensions; neighbors_on_fm: neighbors graph present in uns; celltype_silhouette_fm: silhouette(FM_obsm | cell_type) >= 0.05 to validate structure preservation; fm_real_call: trajectory must contain evidence of real FM invocation (bracketed [scGPT]/[Geneformer] markers, SCLLMManager calls, or import statements from real libraries)</td></tr>
<tr><td>L02_foundation_model_celltype_annotation</td><td>FM (scLLM)</td><td>L — Single-cell foundation models (scLLM)</td><td>hard</td><td>pbmc8k_raw.h5ad</td><td>obs column cell_type_<model> (or <model>_celltype variant) with fine-tuned cell-type predictions covering >=95% of cells. obsm key X_<model> with embedding used for annotation. Evidence of real fine-tuning: per-epoch metrics (train_acc, val_acc, epoch N/M progress bars) in trajectory.</td><td>1</td><td>1; fine_tune_evidence: trajectory must contain per-epoch training metrics (train_acc=/val_acc=/Fine-tuning: progress bar) from real omicverse SCLLMManager or upstream library (scGPT/Geneformer/scFoundation/UCE/CellPLM/Tabula), with bracketed [ModelName] prefixes or function signatures proving actual training loop execution</td></tr>
<tr><td>L03_foundation_model_zero_shot_inference</td><td>FM (scLLM)</td><td>L — Single-cell foundation models (scLLM)</td><td>hard</td><td>pbmc8k_raw.h5ad</td><td>Layer or obsm key marked with FM identifier holding inferred output (pattern:</td><td>3</td><td>3; fm_inference_artifact: must have layer or obsm key matching (scgpt|geneformer|scf</td></tr>
</table>

<table>
<tr><th></th><th></th><th></th><th></th><th></th><th></th><th></th><th></th></tr>
<tr><td></td><td></td><td></td><td></td><td></td><td>layers['<model>_imputed'/'_denoised'] or obsm['X_<model>_corrected']). uns dict with FM model name and inference_summary/metrics keys. Deliverable is either gene-expression imputation/denoising (held-out gene prediction) or batch-effect-corrected cell representation.</td><td></td><td>oundation|uce|cellplm|tabula|fm|foundation).*(imputed|denoised|corrected|inferred|reconstructed); fm_inference_summary: must have uns key matching (scgpt|geneformer|scfoundation|uce|cellplm|tabula|fm|foundation).*(inference|metrics|summary) or exact model-name key; fm_real_call: trajectory must show real FM inference invocation (bracketed [ModelName] + (Loading|Loaded|Embedding|Predicting|Inference|Imputation|Denoising), or calls to get_embeddings/embed_data/predict_celltypes/EmbExtractor/Classifier, or imports from real FM libraries)</td></tr>
<tr><td>L04_foundation_model_perturbation_response</td><td>FM (scLLM)</td><td>L — Single-cell foundation models (scLLM)</td><td>hard</td><td>pbmc8k_raw.h5ad</td><td>Layer key containing 'perturb' or '*_pred/*_predicted' with predicted post-perturbation expression matrix. uns key containing 'perturbation' and FM model name, with at minimum: target-gene name, model identifier, summary of top-N up/down-regulated genes. obs column containing 'perturb_response' or '*_response/*_pred_magnitude' with per-cell response magnitude (norm of perturbed-minus-observed difference).</td><td>4</td><td>4; perturb_layer: must have layer matching regex (perturb|_pred$|_predicted$); perturb_uns: must have uns key matching regex (perturb); perturb_response_obs: must have obs column matching regex (perturb|_response$|_pred_magnitude); perturbation_real_call: trajectory must show real perturbation-prediction API call (bracketed [FM_Name] + (Perturbation|Predicting|Inference|Loaded), or calls to predict_perturbation/InSilicoPerturber/[scFoundation] Perturb, or real FM library invocations)</td></tr>
</table>

#### Summary

Modality: scRNA-seq 19, Spatial 4, Multi-omics 2, Bulk RNA 5, Metabolomics 1, Microbiome 16S 5, FM (scLLM) 4 (=40).
Difficulty: 29 hard, 9 medium, 2 easy.

#### Notes

- OmicBench = 40 omics-analysis tasks across seven data modalities (scRNA-seq 19, spatial transcriptomics 4, multi-omics 2, bulk RNA-seq 5, metabolomics 1, 16S microbiome 5, single-cell foundation-model workflows 4) and eight analytical layers (A-G, L). Difficulty: 29 hard, 9 medium, 2 easy.
- Layer legend — A: scRNA preprocessing floor (PBMC); B: scRNA-seq downstream workflows; C: spatial transcriptomics (Visium/Visium-HD); D: multi-omics integration (multiome); E: bulk RNA-seq & metabolomics; F: trajectory & RNA velocity; G: 16S microbiome; L: single-cell foundation models (scLLM). Modality is derived from the task's data type (notebooks/05_task_taxonomy_heatmap.ipynb TAXONOMY); bulk RNA and metabolomics both live in layer E.
- Each task's prompt states the scientific GOAL only and never names an ov.X.Y API, so every harness (with or without OmicVerse) composes its own solution; the bench grades the result. Fixtures resolve under <OVBENCH_ROOT>/fixtures/ at run time; the 'oracle' precomputed artifact (not shown) defines numeric tolerances.
- Grading checks are programmatic (bench/grader.py, ~50 typed handlers): they assert presence of specific obs/var/obsm/uns/layers keys (with alias groups), shape ranges, value ranges, and biology/numerics — e.g. clustering ARI vs a known cell-type label, marker-overlap vs an oracle, multi-method consensus (≥2 doublet/DE/annotation methods), kBET/LISI

batch metrics, pseudotime root-anchoring and correlation. A task passes a check only if its biological/numerical criterion is met, not merely if a key exists.

- The 'Expected output' and 'Grading checks' columns are faithful summaries of each task's prompt and check list (generated by a per-layer agent pass over bench/tasks.py + bench/grader.py); the authoritative, exhaustive definitions remain bench/tasks.py.

### *Supplementary Table 14 | OmicBench task taxonomy (exported from the taxonomy notebook)*

All 40 OmicBench tasks mapped to data modality, detailed analysis category, analysis family (15, S4), capability class (6, S3 / Fig. 2g) and biological domain. Table 12b/12c are the coverage grids behind Extended Data Fig. 8 and Fig. 2g.

#### Table 14a | Per-task taxonomy

| Task ID | Capability class (Fig. 2g) | Analysis family (Fig. 2/EDF8) | Detailed category | Data modality | Biological domain | Layer | Difficulty |
|---|---|---|---|---|---|---|---|
| A01_qc_filter | Preprocessing & Annotation | Preprocessing & QC | Quality Control | scRNA-seq | Immunology | A | medium |
| B01_full_pipeline | Preprocessing & Annotation | Preprocessing & QC | End-to-end Pipeline | scRNA-seq | Immunology | B | hard |
| A02_normalize_log | Preprocessing & Annotation | Preprocessing & QC | Normalization | scRNA-seq | Immunology | A | easy |
| A03_hvg | Preprocessing & Annotation | Preprocessing & QC | Feature Selection (HVG) | scRNA-seq | Immunology | A | easy |
| A04_pca_neighbors_umap | Preprocessing & Annotation | Preprocessing & QC | Dimensionality Reduction | scRNA-seq | Immunology | A | medium |
| B04_multi_method_batch_correction | Preprocessing & Annotation | Preprocessing & QC | Batch Correction | scRNA-seq | Immunology | B | hard |
| B02_subset_recluster | Preprocessing & Annotation | Clustering | Clustering | scRNA-seq | Immunology | B | hard |
| A05_leiden_clustering | Preprocessing & Annotation | Clustering | Clustering | scRNA-seq | Immunology | A | hard |
| B05_multi_method_annotation | Preprocessing & Annotation | Cell-Type Annotation | Cell-Type Annotation | scRNA-seq | Immunology | B | hard |
| B06_reference_annotation | Preprocessing & Annotation | Cell-Type Annotation | Cell-Type Annotation | scRNA-seq | Immunology | B | medium |
| G01_16s_diversity | Microbiome & Foundation Model | Diversity & Ordination | Alpha / Beta Diversity | Microbiome 16S | Microbiology | G | medium |
| G03_16s_ordination_pcoa | Microbiome & Foundation Model | Diversity & Ordination | Ordination (PCoA) | Microbiome 16S | Microbiology | G | medium |
| G05_phylogenetic_diversity | Microbiome & Foundation Model | Diversity & Ordination | Phylogenetic Diversity | Microbiome 16S | Microbiology | G | hard |
| G04_16s_taxonomic_collapse | Microbiome & Foundation Model | Taxonomic Profiling | Taxonomic Profiling | Microbiome 16S | Microbiology | G | medium |
| L01_foundation_model_cell_embedding | Microbiome & Foundation Model | Foundation Model Application | FM Cell Embedding | FM (scLLM) | Immunology | L | hard |
| L02_foundation_model_celltype_annotation | Microbiome & Foundation Model | Foundation Model Application | FM Cell-Type Annotation | FM (scLLM) | Immunology | L | hard |
| L03_foundation_model_zero_shot_inference | Microbiome & Foundation Model | Foundation Model Application | FM Zero-shot Inference | FM (scLLM) | Immunology | L | hard |
| L04_foundation_model_perturbation_response | Microbiome & Foundation Model | Foundation Model Application | FM Perturbation Response | FM (scLLM) | Immunology | L | hard |
| F01_multi_mode_velocity | Trajectory & Integration | Trajectory & Velocity | RNA Velocity | scRNA-seq | Endocrinology | F | hard |
| F02_multi_method_trajectory | Trajectory & Integration | Trajectory & Velocity | Trajectory / Pseudotime | scRNA-seq | Endocrinology | F | hard |
| F03_paga_pseudotime | Trajectory & Integration | Trajectory & Velocity | Trajectory / Pseudotime | scRNA-seq | Endocrinology | F | hard |
| C04_spatial_trajectory_stt | Trajectory & Integration | Trajectory & Velocity | Spatial Trajectory | Spatial | Neurology | C | hard |
| F04_multi_backend_pseudotime | Trajectory & Integration | Trajectory & Velocity | Trajectory / Pseudotime | scRNA-seq | Endocrinology | F | hard |

| | | | | | | | |
|---|---|---|---|---|---|---|---|
| D01_multiome_joint_embedding | Trajectory & Integration | Multi-omic Integration | Multi-omic Integration | Multi-omics | Immunology | D | hard |
| D02_peak_gene_linking | Trajectory & Integration | Multi-omic Integration | Peak-Gene Linking | Multi-omics | Immunology | D | hard |
| E06_pathway_enrichment | Pathway & Networks | Pathway & Signature | Pathway Enrichment | Bulk RNA | Neurology | E | medium |
| B08_aucell_pathway_scoring | Pathway & Networks | Pathway & Signature | Signature Scoring | scRNA-seq | Immunology | B | medium |
| B09_cnmf_gene_programs | Pathway & Networks | Pathway & Signature | Gene Programs (NMF) | scRNA-seq | Immunology | B | hard |
| E04_bulk_wgcna | Pathway & Networks | Regulatory & Networks | Co-expression Network | Bulk RNA | Neurology | E | hard |
| E07_ppi_network | Pathway & Networks | Regulatory & Networks | PPI Network | Bulk RNA | Neurology | E | medium |
| B10_gene_regulatory_network | Pathway & Networks | Regulatory & Networks | Gene Regulatory Network | scRNA-seq | Endocrinology | B | hard |
| B11_scrna_cell_cell_communication | Pathway & Networks | Cell-Cell Communication | Cell-Cell Communication | scRNA-seq | Immunology | B | hard |
| G02_multi_method_da | Differential & Statistical | Differential Analysis | Differential Abundance | Microbiome 16S | Microbiology | G | hard |
| B03_diff_expression | Differential & Statistical | Differential Analysis | Differential Expression | scRNA-seq | Immunology | B | hard |
| E03_bulk_de_multi_method | Differential & Statistical | Differential Analysis | Differential Expression | Bulk RNA | Neurology | E | hard |
| E05_bulk_metabol_multivariate | Differential & Statistical | Multivariate Stats | Multivariate Stats | Metabolomics | Oncology & Metabolic | E | hard |
| C01_visium_spatial_domain | Spatial Methods | Spatial Analysis | Spatial Domain | Spatial | Neurology | C | hard |
| C02_visium_hd_svg | Spatial Methods | Spatial Analysis | Spatially Variable Genes | Spatial | Others | C | hard |
| E01_bulk_deconvolution | Spatial Methods | Cell Deconvolution | Bulk Deconvolution | Bulk RNA | Neurology | E | hard |
| C05_spatial_celltype_deconvolution | Spatial Methods | Cell Deconvolution | Spatial Deconvolution | Spatial | Immunology | C | hard |

### Table 14b | Analysis family × biological domain (Extended Data Fig. 8)

| Analysis family (Fig. 2/EDF8) | Immunology | Neurology | Endocrinology | Oncology & Metabolic | Microbiology | Others | Total |
|---|---|---|---|---|---|---|---|
| Preprocessing & QC | 6 | 0 | 0 | 0 | 0 | 0 | 6 |
| Clustering | 2 | 0 | 0 | 0 | 0 | 0 | 2 |
| Cell-Type Annotation | 2 | 0 | 0 | 0 | 0 | 0 | 2 |
| Diversity & Ordination | 0 | 0 | 0 | 0 | 3 | 0 | 3 |
| Taxonomic Profiling | 0 | 0 | 0 | 0 | 1 | 0 | 1 |
| Foundation Model Application | 4 | 0 | 0 | 0 | 0 | 0 | 4 |
| Trajectory & Velocity | 0 | 1 | 4 | 0 | 0 | 0 | 5 |
| Multi-omic Integration | 2 | 0 | 0 | 0 | 0 | 0 | 2 |
| Pathway & Signature | 2 | 1 | 0 | 0 | 0 | 0 | 3 |
| Regulatory & Networks | 0 | 2 | 1 | 0 | 0 | 0 | 3 |
| Cell-Cell Communication | 1 | 0 | 0 | 0 | 0 | 0 | 1 |
| Differential Analysis | 1 | 1 | 0 | 0 | 1 | 0 | 3 |
| Multivariate Stats | 0 | 0 | 0 | 1 | 0 | 0 | 1 |
| Spatial Analysis | 0 | 1 | 0 | 0 | 0 | 1 | 2 |
| Cell Deconvolution | 1 | 1 | 0 | 0 | 0 | 0 | 2 |

### Table 14c | Capability class × data modality (Fig. 2g)

| Capability class (Fig. 2g) | scRNA-seq | Spatial | Multi-omics | Bulk RNA | Metabolomics | Microbiome 16S | FM (scLLM) | Total |
|---|---|---|---|---|---|---|---|---|
| Preprocessing & Annotation | 10 | 0 | 0 | 0 | 0 | 0 | 0 | 10 |
| Microbiome & Foundation Model | 0 | 0 | 0 | 0 | 0 | 4 | 4 | 8 |

| | | | | | | | | |
|---|---|---|---|---|---|---|---|---|
| Trajectory & Integration | 4 | 1 | 2 | 0 | 0 | 0 | 0 | 7 |
| Pathway & Networks | 4 | 0 | 0 | 3 | 0 | 0 | 0 | 7 |
| Differential & Statistical | 1 | 0 | 0 | 1 | 1 | 1 | 0 | 4 |
| Spatial Methods | 0 | 3 | 0 | 1 | 0 | 0 | 0 | 4 |

#### Notes

- Taxonomy exported verbatim from notebooks/05_task_taxonomy_heatmap.ipynb (TAXONOMY + FAMILY_TO_GROUP). Each of the 40 OmicBench tasks maps to one data modality, one detailed analysis category, one analysis family (15 fine-grained groups, S4 / Extended Data Fig. 8 y-axis), one capability class (6 coarse groups, S3 / Fig. 2g), and one biological domain (6 groups, derived from the task's tissue/disease fixture).
- The 6 capability classes group the 15 families: Preprocessing & Annotation = {Preprocessing & QC, Clustering, Cell-Type Annotation}; Differential & Statistical = {Differential Analysis, Multivariate Stats}; Pathway & Networks = {Pathway & Signature, Regulatory & Networks, Cell-Cell Communication}; Trajectory & Integration = {Trajectory & Velocity, Multi-omic Integration}; Spatial Methods = {Spatial Analysis, Cell Deconvolution}; Microbiome & Foundation Model = {Diversity & Ordination, Taxonomic Profiling, Foundation Model Application}.
- Tasks per capability class: Preprocessing & Annotation 10, Microbiome & Foundation Model 8, Trajectory & Integration 7, Pathway & Networks 7, Spatial Methods 4, Differential & Statistical 4 (=40). Tasks per biological domain: Immunology 21, Neurology 7, Endocrinology 5, Microbiology 5, Oncology & Metabolic 1, Others 1 (=40).
- Table 12b is the analysis-family x biological-domain coverage grid underlying Extended Data Fig. 8 (cell counts; the figure renders these as check marks). Table 12c is the coarse capability-class x data-modality grid underlying Fig. 2g. Domain is the disease/tissue axis; modality is the data-type axis (OmicBench's primary organizing structure).
- Biological domain is assigned from each task's fixture tissue/disease context (e.g. PBMC -> Immunology, pancreas endocrinogenesis -> Endocrinology, brain/Visium -> Neurology, 16S gut -> Microbiology); it is a coverage descriptor, not a grading axis.

### *Supplementary Table 15 | OmicBench Mini-SWE-Agent baseline vs +OmicVerse Pass@1 (Fig. 2j)*

Per-model Pass@1 under the Mini-SWE-Agent harness, baseline vs +OmicVerse, with numerator/denominator (passed cells / total graded (task×seed) cells). 7 models × 2 arms = 14 arms; sorted by uplift descending. Recomputed from grade records.

| Model | Provider | Open weights | Baseline Pass@1 | Baseline % | +OmicVerse Pass@1 | +OmicVerse % | Δ Pass@1 (pp) | Tasks | Seeds |
|---|---|---|---|---|---|---|---|---|---|
| Qwen3.6-35B-A3B | Alibaba | yes | 17/38 | 44.7 | 30/38 | 78.9 | **34.2** | 38 | 1 |
| GLM-5.1 | Zhipu | no | 76/112 | 67.9 | 98/112 | 87.5 | **19.6** | 38 | 3 |
| GPT-5.5 | OpenAI | no | 82/114 | 71.9 | 103/113 | 91.2 | **19.2** | 38 | 3 |
| DeepSeek-v4-pro | DeepSeek | yes | 81/114 | 71.1 | 102/114 | 89.5 | **18.4** | 38 | 3 |
| Gemini-3.1-flash-lite | Google | no | 71/112 | 63.4 | 89/112 | 79.5 | **16.1** | 38 | 3 |
| DeepSeek-v4-flash | DeepSeek | yes | 75/103 | 72.8 | 88/101 | 87.1 | **14.3** | 38 | 3 |
| MiniMax-M2.7 | MiniMax | no | 88/114 | 77.2 | 91/114 | 79.8 | **2.6** | 38 | 3 |

#### Summary

- Adding OmicVerse to the same Mini-SWE-Agent harness raised Pass@1 for every model family, with the largest gains in weaker/smaller models: Qwen3.6-35B-A3B +34.2 pp (44.7→78.9), GLM-5.1 +19.6 (67.9→87.5), GPT-5.5 +19.2 (71.9→91.2), DeepSeek-v4-pro +18.4 (71.1→89.5), Gemini-3.1-flash-lite +16.1 (63.4→79.5), DeepSeek-v4-flash +14.3 (72.8→87.1) and MiniMax-M2.7 +2.6 (77.2→79.8). The smallest open-weights model gains most, consistent with OmicVerse supplying domain capability the base model lacks rather than amplifying an already strong model.

#### Method

- Harness: Mini-SWE-Agent, a deliberately minimal controlled agent loop, run on OmicBench. Each model is evaluated in two arms — baseline (Mini-SWE-Agent alone) and +OmicVerse (same harness with the OmicVerse function registry / skills added) — so any difference is attributable to OmicVerse, not the harness or model.
- Pass@1(model, arm) = passed cells / total graded cells, where a cell is one (task, seed) and a task passes only if all of its programmatic grading checks pass (bench/grader.py). Arm is read from the grades.csv `system` suffix (_baseline / _omicverse). Values recomputed directly from results/*/grades.csv; no agent trajectories are read.
- Denominators: the canonical sweep ran 38 OmicBench tasks under Mini-SWE-Agent across 3 seeds (=114 cells/arm); a few cells per arm are absent from grades.csv because the agent produced no gradeable output (adapter error) and are excluded from the denominator, so totals range 101-114. Qwen3.6-35B-A3B (locally served, open weights) was run at a single seed (38 cells/arm).
- Ablation arms in results/ (omicverse_no_registry, omicverse_doc_rag) are NOT part of these 14 arms and are excluded here; they isolate the registry vs RAG contribution in a separate analysis.

#### Caveats

- Pass@1 denominators are not identical across arms/models (101-114 for the 3-seed models, 38 for single-seed Qwen), because adapter-error cells are dropped; the percentages are over gradeable cells only, so a model with more dropped cells is scored on slightly fewer tasks.
- Qwen3.6-35B-A3B is a single seed, so its Pass@1 (and its headline +34.2 pp gain) has no seed-level variance estimate and is higher-variance than the 3-seed models.
- Grading is programmatic per-task checks (all-checks-pass = task pass); it is binary Pass@1, not a partial-credit rubric mean, so it is stricter than the per-check capability scores used elsewhere in the figure.

### *Supplementary Table 16 | OmicBench task-level pass/fail matrix by model, seed and layer (Fig. 2l)*

Per-model majority-seed task passes by OmicBench layer, baseline vs +OmicVerse (Δ shaded). The exact (model, arm, task, seed) pass/fail matrix (1,411 cells) and the 38-task majority-seed grid are provided in the spreadsheet (sheets 'Raw matrix' and 'Maj-seed task grid').

#### Per-model layer × arm passes (majority-seed)

| Model | Layer | n tasks | Baseline passed | +OmicVerse passed | Δ tasks |
|---|---|---|---|---|---|
| GPT-5.5 | A: scRNA preprocessing | 5 | 5 | 5 | 0 |
| GPT-5.5 | B: scRNA workflows | 10 | 7 | 9 | 2 |
| GPT-5.5 | C: spatial | 4 | 2 | 3 | 1 |
| GPT-5.5 | E: bulk RNA-seq | 6 | 3 | 6 | 3 |
| GPT-5.5 | F: velocity / trajectory | 4 | 4 | 4 | 0 |
| GPT-5.5 | G: 16S microbiome | 5 | 5 | 5 | 0 |
| GPT-5.5 | L: foundation model | 4 | 2 | 3 | 1 |
| DeepSeek-v4-pro | A: scRNA preprocessing | 5 | 5 | 4 | -1 |
| DeepSeek-v4-pro | B: scRNA workflows | 10 | 6 | 8 | 2 |
| DeepSeek-v4-pro | C: spatial | 4 | 1 | 3 | 2 |
| DeepSeek-v4-pro | E: bulk RNA-seq | 6 | 5 | 6 | 1 |
| DeepSeek-v4-pro | F: velocity / trajectory | 4 | 4 | 4 | 0 |
| DeepSeek-v4-pro | G: 16S microbiome | 5 | 4 | 5 | 1 |
| DeepSeek-v4-pro | L: foundation model | 4 | 2 | 4 | 2 |
| DeepSeek-v4-flash | A: scRNA preprocessing | 5 | 5 | 5 | 0 |
| DeepSeek-v4-flash | B: scRNA workflows | 10 | 6 | 9 | 3 |
| DeepSeek-v4-flash | C: spatial | 4 | 2 | 3 | 1 |
| DeepSeek-v4-flash | E: bulk RNA-seq | 6 | 4 | 6 | 2 |
| DeepSeek-v4-flash | F: velocity / trajectory | 4 | 4 | 4 | 0 |
| DeepSeek-v4-flash | G: 16S microbiome | 5 | 5 | 5 | 0 |
| DeepSeek-v4-flash | L: foundation model | 4 | 2 | 3 | 1 |
| Gemini-3.1-flash-lite | A: scRNA preprocessing | 5 | 4 | 5 | 1 |
| Gemini-3.1-flash-lite | B: scRNA workflows | 10 | 7 | 8 | 1 |
| Gemini-3.1-flash-lite | C: spatial | 4 | 1 | 2 | 1 |
| Gemini-3.1-flash-lite | E: bulk RNA-seq | 6 | 3 | 5 | 2 |

| | | | | | |
|---|---|---|---|---|---|
| Gemini-3.1-flash-lite | F: velocity / trajectory | 4 | 4 | 4 | 0 |
| Gemini-3.1-flash-lite | G: 16S microbiome | 5 | 4 | 5 | 1 |
| Gemini-3.1-flash-lite | L: foundation model | 4 | 2 | 2 | 0 |
| GLM-5.1 | A: scRNA preprocessing | 5 | 5 | 5 | 0 |
| GLM-5.1 | B: scRNA workflows | 10 | 5 | 9 | 4 |
| GLM-5.1 | C: spatial | 4 | 2 | 4 | 2 |
| GLM-5.1 | E: bulk RNA-seq | 6 | 4 | 6 | 2 |
| GLM-5.1 | F: velocity / trajectory | 4 | 3 | 4 | 1 |
| GLM-5.1 | G: 16S microbiome | 5 | 5 | 5 | 0 |
| GLM-5.1 | L: foundation model | 4 | 1 | 3 | 2 |
| MiniMax-M2.7 | A: scRNA preprocessing | 5 | 5 | 5 | 0 |
| MiniMax-M2.7 | B: scRNA workflows | 10 | 6 | 7 | 1 |
| MiniMax-M2.7 | C: spatial | 4 | 2 | 2 | 0 |
| MiniMax-M2.7 | E: bulk RNA-seq | 6 | 5 | 5 | 0 |
| MiniMax-M2.7 | F: velocity / trajectory | 4 | 4 | 4 | 0 |
| MiniMax-M2.7 | G: 16S microbiome | 5 | 5 | 5 | 0 |
| MiniMax-M2.7 | L: foundation model | 4 | 4 | 2 | -2 |
| Qwen3.6-35B-A3B | A: scRNA preprocessing | 5 | 3 | 5 | 2 |
| Qwen3.6-35B-A3B | B: scRNA workflows | 10 | 4 | 8 | 4 |
| Qwen3.6-35B-A3B | C: spatial | 4 | 1 | 2 | 1 |
| Qwen3.6-35B-A3B | E: bulk RNA-seq | 6 | 2 | 4 | 2 |
| Qwen3.6-35B-A3B | F: velocity / trajectory | 4 | 3 | 4 | 1 |
| Qwen3.6-35B-A3B | G: 16S microbiome | 5 | 4 | 5 | 1 |
| Qwen3.6-35B-A3B | L: foundation model | 4 | 0 | 2 | 2 |

#### Interpretation

**S3:** Under +OmicVerse, majority-seed task passes increase across multiple layers, not one task family: gains appear in scRNA preprocessing (A), scRNA workflows (B), bulk RNA-seq (E), velocity/trajectory (F), 16S microbiome (G) and foundation-model (L) layers across the model panel (per-model layer breakdown in 14c).

**S4:** GLM-5.1 improves from 25/38 to 36/38 majority-seed task passes with OmicVerse, and Qwen3.6-35B-A3B from 17/38 to 30/38 (the largest gain). Note MiniMax-M2.7 is the only arm with a small NEGATIVE majority-seed delta (31->30), even though its cell-level Pass@1 rises slightly (Table 13) — majority-seed collapse and cell-level mean can disagree at the margin on a near-flat model.

#### Method & caveats

- 14a (raw matrix) is the exact task-level pass/fail record: one row per (model, arm, task, seed) with layer and difficulty, taken verbatim from results/*/grades.csv (1,411 graded cells). A task passes a cell only if ALL its programmatic grading checks pass.
- Seeds: six models ran 3 seeds (0,1,2); Qwen3.6-35B-A3B (locally served, open weights) ran a single seed. The 38-task eval set covers layers A,B,C,E,F,G,L (layer D / multiome's 2 tasks were not run under this minimal-agent sweep, so 40-2=38).
- 14b/14c collapse seeds to a single per-task pass by MAJORITY vote (pass if >=50% of that arm's seeds passed), matching the Fig. 2l heat-map convention. The per-model majority-seed task totals are: GPT-5.5 28->35, DeepSeek-v4-pro 27->34, DeepSeek-v4-flash 28->35, Gemini-3.1-flash-lite 25->31, GLM-5.1 25->36, MiniMax-M2.7 31->30, Qwen3.6-35B-A3B 17->30.
- Layer legend: A scRNA preprocessing, B scRNA workflows, C spatial, E bulk RNA-seq, F velocity/trajectory, G 16S microbiome, L foundation model.
- Majority-seed collapse (14b/14c) is a 3-seed vote for six models but a single seed for Qwen (no vote), so Qwen's per-task pass is one run and higher-variance.
- Cell-level Pass@1 (Table 13) and majority-seed task pass (this table) are different aggregations of the same raw matrix (14a) and need not rank models identically near ties (e.g. MiniMax).
- Pass/fail is binary all-checks-must-pass per task; it does not capture partial progress, which the per-check capability scores elsewhere in the figure do.

### *Supplementary Table 17 | Residual failure taxonomy (OmicBench +OmicVerse arm)*

Every task that still failed after OmicVerse augmentation (majority-seed fail, 35 cells), with the bench failure_mode, the atomic checks that failed, and the dominant cause category. Cause shaded. Full failed-check / secondary-cause / seed columns in the spreadsheet.

| Model | Failed task | Layer | Failure mode | Primary cause | Failed checks | Notes (cause) |
|---|---|---|---|---|---|---|
| DeepSeek-v4-pro | A01_qc_filter | A: scRNA preprocessing | wrong_tool_choice | model reasoning | cells_after_qc: filtered count 5406 below required minimum 5500; multi_doublet_methods applied successfully | Filtering thresholds or doublet-detection parameters were set too stringently, removing 94 cells below the threshold despite passing 5 other checks. |
| DeepSeek-v4-flash | B08_aucell_pathway_scoring | B: scRNA workflows | wrong_workflow_order | model reasoning | tb_celltype_specificity (only 1 column peaks in expected family; need ≥2) | Score columns were generated correctly but cell-type specificity pattern was incomplete, with only b_cell showing strong peak instead of both t_cell and b_cell. |
| DeepSeek-v4-pro | B05_multi_method_annotation | B: scRNA workflows | exceeded_turns | runtime execution | multi_annotation: zero cell-type annotation columns produced; no output generated despite completing run | Agent reached turn limit before completing multi-method annotation pipeline and producing any cell-type columns. |
| DeepSeek-v4-pro | B08_aucell_pathway_scoring | B: scRNA workflows | wrong_workflow_order | model reasoning | tb_celltype_specificity: only 1 of expected 2+ cell types showed statistically significant peak in score columns; score columns created but lack biological specificity | Pathway scoring method or workflow application order failed to generate sufficient cell-type-specific signal; only b_cell peaked when both b_cell and t_cell were required. |
| GLM-5.1 | B08_aucell_pathway_scoring | B: scRNA workflows | wrong_workflow_order | model reasoning | tb_celltype_specificity — only 1 column peaked in expected cell-type family (need ≥2) | Pathway scores were computed but lacked sufficient cell-type specificity; the workflow did not ensure distinct cell-type separation in scoring distributions. |
| GPT-5.5 | B08_aucell_pathway_scoring | B: scRNA workflows | wrong_workflow_order | model reasoning | tb_celltype_specificity — only 1 peak in expected score family (need ≥2); score columns matched but insufficient statistical separation | Model generated score columns matching the regex but failed to ensure sufficient cell-type specificity across scores; only one cell type showed significant peak (z_gap=6.80), indicating the workflow order or scoring logic was incorrect. |
| Gemini-3.1-flash-lite | B08_aucell_pathway_scoring | B: scRNA workflows | wrong_workflow_order | model reasoning | tb_celltype_specificity: only 1 column peaked in expected family (need ≥2); score_columns_regex and score_count_matching passed, confirming 2 matching columns were found but not properly integrated | Agent created score columns but failed to ensure they properly capture cell-type-specific signals across the expected family; scoring method or signature selection was incomplete. |
| Gemini-3.1-flash-lite | B11_scrna_cell_cell_communication | B: scRNA workflows | wrong_workflow_order | tool grounding | ccc_real_call: real CCC library invocation (cellphonedb/liana/commot/ov.single) produced no markers in tool output; ccc_table check passed, indicating the key was found but no real library call was executed | Agent found or created a CCC results key but did not actually invoke the real cell-cell communication library; hallucinated results instead of calling cellphonedb, liana, or OmicVerse tools. |
| MiniMax-M2.7 | B08_aucell_pathway_scoring | B: scRNA workflows | wrong_workflow_order | model reasoning | score_columns_regex, score_count_matching, tb_celltype_specificity — no scoring columns found in obs | The workflow executed but produced no AUCell score columns at all, indicating the agent chose the wrong analytical method or skipped the scoring step. |
| MiniMax-M2.7 | B10_gene_regulatory_network | B: scRNA workflows | wrong_workflow_order | model reasoning | regulon_obsm, regulon_uns, n_regulons — no regulon/TF-module keys in obsm or uns | The workflow ran but failed to produce any regulon or TF-module analytical outputs, indicating incorrect method selection or wrong workflow order. |

| | | | | | | |
|---|---|---|---|---|---|---|
| MiniMax-M2.7 | B11_scrna_cell_cell_communication | B: scRNA workflows | wrong_workflow_order | data handling | ccc_table — uns missing cell–cell communication table; ccc_real_call passed (cellphonedb_v5 invoked) | The CCC library was invoked correctly but the resulting interaction table was not saved to the expected location in uns. |
| Qwen3.6-35B-A3B | B10_gene_regulatory_network | B: scRNA workflows | exceeded_turns | runtime execution | No final adata produced; no atomic checks failed because the run exhausted turns before any checks could evaluate. | The model exceeded its turn budget and failed to complete the analysis, producing no final AnnData object, indicating a timeout or step-limit exhaustion rather than a logical error. |
| Qwen3.6-35B-A3B | B11_scrna_cell_cell_communication | B: scRNA workflows | exceeded_turns | runtime execution | No final adata produced; no atomic checks failed because the run exhausted turns before completion. | The model exceeded its turn budget before completing the cell-cell communication analysis, resulting in no final AnnData object. |
| DeepSeek-v4-flash | C04_spatial_trajectory_stt | C: spatial | wrong_tool_choice | tool grounding | stt_real_call (no markers in tool output) | The spatial-transition-tensor library was never actually invoked; no import or function-call markers for STT appeared in the agent's output. |
| DeepSeek-v4-pro | C01_visium_spatial_domain | C: spatial | wrong_tool_choice | model reasoning | spatial_silhouette: score -0.036 below 0.05 threshold; ari_vs_ground_truth: 0.204 indicating poor cluster coherence; 5 other checks passed | Spatial clustering algorithm choice or parameter settings produced clusters with negative silhouette score and weak concordance with ground truth. |
| GPT-5.5 | C04_spatial_trajectory_stt | C: spatial | wrong_tool_choice | tool grounding | stt_real_call — no markers in tool output; spatial-transition-tensor library was never actually invoked | The spatial-transition-tensor tool was not genuinely called; despite finding the expected uns key 'spatial_trajectory_transition', no actual STT library markers appeared in the agent output. |
| Gemini-3.1-flash-lite | C01_visium_spatial_domain | C: spatial | wrong_tool_choice | model reasoning | spatial_silhouette: spatial silhouette(spatial_domain \| spatial) = -0.013 < 0.05; ari_vs_ground_truth: ARI = 0.213, both metrics indicate weak clustering quality | Agent chose a spatial clustering method that produced poor separation and low agreement with ground truth, suggesting wrong algorithm selection or parameter tuning. |
| Gemini-3.1-flash-lite | C04_spatial_trajectory_stt | C: spatial | wrong_tool_choice | tool grounding | stt_real_call: real STT (spatial-transition-tensor) invocation produced no markers in tool output | Agent did not invoke the spatial-transition-tensor library; no markers for the real tool call were found in output despite creating transition_matrix key. |
| MiniMax-M2.7 | C04_spatial_trajectory_stt | C: spatial | wrong_tool_choice | tool grounding | stt_real_call — no markers in tool output, real spatial-transition-tensor invocation never occurred | The spatial-transition-tensor library was never actually invoked; the agent either hallucinated the tool call or failed to ground it in the real library. |
| MiniMax-M2.7 | C05_spatial_celltype_deconvolution | C: spatial | wrong_tool_choice | data handling | fractions_present — deconvolution fractions do not sum to 1; deconv_real_call passed | The deconvolution library was called correctly but the output fractions failed the normalization constraint, indicating incorrect post-processing or normalization logic. |
| Qwen3.6-35B-A3B | C04_spatial_trajectory_stt | C: spatial | silent_none | runtime execution | No final adata produced; no atomic checks indicate a silent failure with no output generated. | The model silently produced no final AnnData object, suggesting the run crashed or terminated without generating any results or diagnostic output. |
| Qwen3.6-35B-A3B | C05_spatial_celltype_deconvolution | C: spatial | silent_none | runtime execution | No final adata produced; no atomic checks indicate a silent failure with no results. | The model silently failed to produce a final AnnData object, suggesting a crash or unhandled error during execution. |
| Gemini-3.1-flash-lite | E07_ppi_network | E: bulk RNA-seq | wrong_tool_choice | tool grounding | ppi_real_call: real STRINGdb / PPI library invocation produced no markers in tool output | Agent did not invoke the STRINGdb or PPI library; created ppi_network and hub_genes keys but no actual tool call was executed. |

| | | | | | | |
|---|---|---|---|---|---|---|
| MiniMax-M2.7 | E07_ppi_network | E: bulk RNA-seq | wrong_tool_choice | tool grounding | ppi_real_call — no markers in tool output, real STRINGdb/PPI library invocation never occurred | The PPI/STRINGdb library was never actually invoked; the agent failed to ground the tool call in the real implementation. |
| Qwen3.6-35B-A3B | E04_bulk_wgcna | E: bulk RNA-seq | silent_none | runtime execution | No final adata produced; no atomic checks indicate a silent failure. | The model silently produced no final AnnData object, indicating a crash or silent termination before completion. |
| Qwen3.6-35B-A3B | E06_pathway_enrichment | E: bulk RNA-seq | exceeded_turns | model reasoning | rank_present and enrichment_table failed: neither 'rank_genes_groups' nor enrichment-related keys present in uns. | The model exceeded its turn budget and never produced the required rank-genes or enrichment analysis results, failing to construct the expected uns keys. |
| DeepSeek-v4-flash | L03_foundation_model_zero_shot_inference | L: foundation model | wrong_tool_choice | model reasoning | fm_inference_artifact (no key matches expected pattern in layers/obsm) | The geneformer library was successfully imported and called, but the agent's method did not produce the expected inference output artifact (imputed/denoised/corrected/inferred/reconstructed layer or obsm key). |
| GLM-5.1 | L01_foundation_model_cell_embedding | L: foundation model | wrong_tool_choice | model reasoning | celltype_silhouette_fm — silhouette score = -0.018 < 0.05 threshold, indicating poor cell-type separation despite successful tool invocation | Foundation model was correctly invoked and produced an embedding (X_scgpt with 512 dims), but the embedding lacked biological signal and failed to separate cell types. |
| GPT-5.5 | L03_foundation_model_zero_shot_inference | L: foundation model | wrong_tool_choice | tool grounding | fm_real_call — no markers in tool output; foundation-model library (geneformer) was never actually invoked despite producing correct artifact key | The foundation-model inference tool was not genuinely invoked; although the correct obsm artifact key 'X_geneformer_corrected' was present, no real geneformer library markers appeared in the output. |
| Gemini-3.1-flash-lite | L02_foundation_model_celltype_annotation | L: foundation model | exceeded_turns | runtime execution | fine_tune_evidence: no fine-tune progress markers in trajectory | Agent exceeded turn budget before completing foundation model fine-tuning; no evidence of fine-tuning progress indicates incomplete workflow execution due to timeout. |
| Gemini-3.1-flash-lite | L03_foundation_model_zero_shot_inference | L: foundation model | wrong_tool_choice | data handling | fm_inference_artifact: no key matches foundation-model inference patterns in layers or obsm; fm_real_call matched ov.llm.SCLLMManager call but artifact missing indicates real tool was called but did not produce expected output structure | Agent invoked the foundation model library but the inferred/corrected embeddings were not stored in the expected obsm or layers keys; output structure or key naming did not match rubric expectations. |
| MiniMax-M2.7 | L02_foundation_model_celltype_annotation | L: foundation model | exceeded_turns | runtime execution | fine_tune_evidence — no fine-tuning progress markers found in trajectory | The run exceeded the turn budget before fine-tuning could complete, preventing any progress markers from being recorded. |
| MiniMax-M2.7 | L03_foundation_model_zero_shot_inference | L: foundation model | wrong_tool_choice | data handling | fm_inference_artifact — no key matching inference/denoising/correction patterns in layers/obsm; fm_real_call passed | The foundation-model library was invoked successfully but the inferred output was not stored under the expected key naming scheme in obsm/layers. |
| Qwen3.6-35B-A3B | L03_foundation_model_zero_shot_inference | L: foundation model | exceeded_turns | model reasoning | fm_inference_artifact, fm_inference_summary, and fm_real_call all failed: no foundation-model inference outputs in layers/obsm, no summary keys, and no real FM library invocation markers detected. | The model exceeded its turn budget without actually invoking the foundation model or producing any inference artifacts, suggesting incorrect method selection or workflow. |
| Qwen3.6-35B-A3B | L04_foundation_model_perturbation_response | L: foundation model | exceeded_turns | model reasoning | perturb_layer, perturb_uns, perturb_response_obs, and perturbation_real_call | The model exceeded its turn budget without invoking any perturbation-prediction |

| | | | | | | |
|---|---|---|---|---|---|---|
| | | | | | all failed: no perturbation predictions in layers/uns/obs and no real perturbation-prediction library invocation detected. | tools or generating the required response predictions, indicating wrong tool choice or workflow method. |

#### Cause × layer

| Primary cause | A | B | C | E | L | Total |
|---|---|---|---|---|---|---|
| tool grounding | 0 | 1 | 4 | 2 | 1 | 8 |
| model reasoning | 1 | 7 | 2 | 1 | 4 | 15 |
| data handling | 0 | 1 | 1 | 0 | 2 | 4 |
| runtime execution | 0 | 3 | 2 | 1 | 2 | 8 |

#### Interpretation (S7)

- OmicVerse improves execution capability but does not eliminate all model- or modality-specific limitations: after augmentation, spatial (C) and foundation-model (L) tasks remain the hardest for several model arms, and the dominant residual cause is model reasoning (15 of 35), i.e. the agent still fails to invoke or correctly orchestrate the specialised tool rather than crashing — a routing/grounding gap, not a runtime one.

#### Method & caveats

- Scope: every (model, task) cell that STILL failed under +OmicVerse by majority-seed vote (35 residual failures across the 7 models). Source: results/*/grades.csv; failure_mode is the bench's own 7-class tag, 'Failed checks' and 'Notes' are the grader's atomic-check diagnostics.
- Cause category (one of: tool grounding, model reasoning, data handling, runtime execution) is the dominant reason the task failed, assigned per failure from the failure_mode + which atomic checks failed + the grader notes: tool grounding = the real domain tool/library was never invoked (a *_real_call check failed); model reasoning = wrong method/workflow/order or expected analytical outputs never produced; data handling = data-integrity/object-plumbing checks failed; runtime execution = code crash or turn/time budget exceeded.
- Cause breakdown: model reasoning 15, runtime execution 8, tool grounding 8, data handling 4.
- Residual failures concentrate in the harder modalities: layer C (spatial) and layer L (foundation model) account for most, with layer B (scRNA workflows) covering the multi-step tool-grounding tasks (CCC, GRN, AUCell). Layers A (preprocessing) and G (16S) are nearly saturated after augmentation.
- Cause is the single dominant category; many failures are multi-factor (a secondary cause is recorded where one clearly contributed). The assignment is grounded in the grader's atomic-check diagnostics, not in re-reading every raw trajectory (trajectory files were consulted only where the notes were ambiguous).
- Residual = majority-seed fail in the +OmicVerse arm; for Qwen3.6-35B-A3B (single seed) it is a single run, so its residual list is higher-variance than the 3-seed models.
- Failure_mode tags come from bench/failure_taxonomy.py (a heuristic 7-class tagger on the run result); the four-way cause grouping here is a coarser, manuscript-facing re-classification of those tags plus the failed-check evidence.

### *Supplementary Table 18 | OmicBench cost model: provider, status, pricing and cache-adjusted cost (Fig. 2k)*

Per model arm: provider, open/local/closed status, hardware, size, token pricing and the cache-adjusted cost per task (baseline vs +OmicVerse), with Pass@1. Status shaded. The cache-adjustment formula and pricing assumptions are in the notes.

| Model | Provider | Status | Hardware setting | Model size | Input $/1M | Cache $/1M | Output $/1M | Baseline cost/task (cache-adj, USD) | +OmicVerse cost/task (cache-adj, USD) | Baseline Pass@1 (%) | +OmicVerse Pass@1 (%) |
|---|---|---|---|---|---|---|---|---|---|---|---|
| GPT-5.5 | OpenAI | Closed | Hosted API (OpenAI) | Not disclosed | 5.0 | 0.5 | 30.0 | 0.1618 | 0.2592 | 71.9 | 91.2 |
| DeepSeek-v4-pro | DeepSeek | Open weights | Hosted API (DeepSeek) | Not disclosed (MoE) | 1.74 | 0.0174 | 3.48 | 0.1009 | 0.1415 | 71.1 | 89.5 |

| | | | | | | | | | | | |
|---|---|---|---|---|---|---|---|---|---|---|---|
| DeepSeek-v4-flash | DeepSeek | Open weights | Hosted API (DeepSeek) | Not disclosed (MoE, smaller) | 0.14 | 0.0028 | 0.28 | 0.0104 | 0.0098 | 72.8 | 87.1 |
| Gemini-3.1-flash-lite | Google | Closed | Hosted API (Google) | Not disclosed | 0.1 | 0.025 | 0.4 | 0.0035 | 0.0098 | 63.4 | 79.5 |
| GLM-5.1 | Zhipu | Open weights | Hosted API (Zhipu) | Not disclosed | 0.6 | 0.06 | 2.4 | 0.0435 | 0.0544 | 67.9 | 87.5 |
| MiniMax-M2.7 | MiniMax | Open weights | Hosted API (MiniMax) | Not disclosed (MoE) | 0.3 | 0.03 | 1.2 | 0.0284 | 0.0358 | 77.2 | 79.8 |
| Qwen3.6-35B-A3B | Alibaba | Open weights · local | Local — ollama, single node (GPU) | 35B total / ~3B active (A3B MoE), 256k ctx | 0.15 | 0.15 | 1.0 | 0.0256 | 0.0623 | 44.7 | 78.9 |

#### Cost-adjustment calculation

- Cache-adjusted cost calculation (analysis/bench_cost.py): an OmicBench task is a long Mini-SWE-Agent loop in which every LLM call resends the growing conversation, so cumulative input reaches 1-2M tokens/task of which ~85-95% is an unchanged cached prefix. Per cell, the per-call input-token counts are sorted by timestamp and only the POSITIVE increments are billed as fresh (uncached) input at the input price; the remaining (cached) input is billed at the provider's cache-read price; output tokens at the output price. Formula per cell: cost = (fresh_in·p_in + cached_in·p_cache + out·p_out) / 1e6; the per-task figure is the mean over that arm's cells.
- This assumes a warm cache between calls, so the cache-adjusted cost is a best-case lower bound; the true cost sits between it and the naive cost (input_tokens·p_in, no caching), which is also tabulated. Per-call usage is read from each assistant message's extra.response.usage in the mini-swe-agent trajectory.
- Token prices (input / cache-read / output USD per 1M) are public list prices, approx. 2026-05 (analysis/bench_cost.py:PRICING). Cache-read price is set to ~10% of input where the provider documents standard prompt caching, near-zero for DeepSeek (token cache ≈ free), and equal to input where no cache discount is documented (Gemini-flash-lite, Qwen) — so for those the cache-adjusted cost reduces toward naive.
- Open weights = the provider publishes downloadable weights (DeepSeek, Zhipu/GLM, MiniMax, Alibaba/Qwen); Closed = API-only (OpenAI, Google). 'Local' marks the arm actually served locally in this sweep: Qwen3.6-35B-A3B ran on local ollama at $0 real cost; its tabulated cost is an if-served-on-OpenRouter reference for apples-to-apples comparison with the API arms.
- Frontier-model parameter counts are not publicly disclosed and are marked 'Not disclosed' rather than estimated; only Qwen3.6-35B-A3B has a published size (35B total, ~3B active mixture-of-experts, 256k context).

#### Interpretation (S1)

- In the cost-accuracy plane (Fig. 2k), +OmicVerse moves nearly every arm UPWARD (higher Pass@1) at similar per-task cost, so open/local arms approach the closed frontier: e.g. DeepSeek-v4-pro reaches 89.5% at a cache-adjusted $0.14/task and Qwen3.6-35B-A3B 78.9% at $0.06/task (reference) / $0 local, versus closed GPT-5.5 at 91.2% but $0.26/task — a ~2-4x cost gap for a few points of accuracy. The cheapest open arms (DeepSeek-v4-flash $0.010, Gemini-flash-lite $0.010) sit near the low-cost Pareto edge.

#### Caveats

- Costs are cache-adjusted best-case (warm-cache) lower bounds; absolute USD depends on volatile list prices and assumed cache-read discounts (per-provider, see notes), so cross-provider ratios are more robust than absolute values.
- Qwen's real run cost was $0 (local ollama); the non-zero figure is a hypothetical OpenRouter price for comparison, and Qwen ran a single seed.
- GPT-5.5's measured cache-hit is lower (66-74%) than the other arms (~90%), so its cache-adjusted cost is closer to naive; this reflects its trajectory/caching behavior in this sweep, not a pricing difference.
- Open/closed status reflects whether weights are published, not the serving path used here (most open-weights arms were still called via a hosted API); only Qwen was served locally.

### *Supplementary Table 19 | OmicBench native six-dimensional capability scores (Fig. 2m)*

Per model and per native dimension, baseline vs +OmicVerse. Each dimension score is the pass-rate over its atomic rubric checks pooled across tasks; overall = mean of the six. Table 17c gives the number of atomic checks per dimension.

#### Table 19a | Per-model × per-dimension scores (both arms)

| Model | Arm | Tool grounding | Data integrity | Object plumbing | Biological plausibility | Quantitative quality | Methodological robustness | Overall (mean of 6) |
|---|---|---|---|---|---|---|---|---|
| DeepSeek-v4-pro | baseline | 40.0 | 100.0 | 97.4 | 87.2 | 89.5 | 100.0 | 85.7 |
| DeepSeek-v4-pro | +OmicVerse | 96.7 | 100.0 | 99.0 | 94.9 | 94.9 | 92.9 | 96.4 |
| GLM-5.1 | baseline | 28.6 | 96.2 | 92.8 | 87.2 | 91.5 | 97.6 | 82.3 |
| GLM-5.1 | +OmicVerse | 96.4 | 100.0 | 99.5 | 89.7 | 95.7 | 92.9 | 95.7 |
| GPT-5.5 | baseline | 16.7 | 100.0 | 100.0 | 94.9 | 97.4 | 95.2 | 84.0 |
| GPT-5.5 | +OmicVerse | 79.3 | 100.0 | 100.0 | 94.9 | 99.1 | 97.6 | 95.2 |
| DeepSeek-v4-flash | baseline | 35.7 | 96.0 | 96.0 | 94.3 | 94.3 | 100.0 | 86.1 |
| DeepSeek-v4-flash | +OmicVerse | 84.6 | 96.0 | 97.2 | 94.3 | 97.2 | 100.0 | 94.9 |
| Gemini-3.1-flash-lite-preview | baseline | 21.4 | 96.2 | 96.4 | 91.9 | 87.8 | 85.7 | 79.9 |
| Gemini-3.1-flash-lite-preview | +OmicVerse | 63.0 | 100.0 | 96.9 | 94.7 | 95.7 | 100.0 | 91.7 |
| MiniMax-M2.7 | baseline | 50.0 | 96.3 | 96.5 | 92.3 | 96.6 | 95.2 | 87.8 |
| MiniMax-M2.7 | +OmicVerse | 73.3 | 96.3 | 92.9 | 92.3 | 95.7 | 97.6 | 91.4 |
| Qwen3.6-35B-A3B-256k | baseline | 16.7 | 87.5 | 82.0 | 69.2 | 70.6 | 76.9 | 67.2 |
| Qwen3.6-35B-A3B-256k | +OmicVerse | 66.7 | 88.9 | 90.7 | 100.0 | 97.2 | 100.0 | 90.6 |

#### Table 19b | Baseline → +OmicVerse uplift (Δ)

| Model | Overall baseline | Overall +OmicVerse | Overall Δ | Δ Tool grounding | Δ Data integrity | Δ Object plumbing | Δ Biological plausibility | Δ Quantitative quality | Δ Methodological robustness |
|---|---|---|---|---|---|---|---|---|---|
| DeepSeek-v4-pro | 85.7 | 96.4 | 10.7 | 56.7 | 0.0 | 1.6 | 7.7 | 5.4 | -7.1 |
| GLM-5.1 | 82.3 | 95.7 | 13.4 | 67.9 | 3.8 | 6.7 | 2.6 | 4.3 | -4.8 |
| GPT-5.5 | 84.0 | 95.2 | 11.1 | 62.6 | 0.0 | 0.0 | 0.0 | 1.7 | 2.4 |
| DeepSeek-v4-flash | 86.1 | 94.9 | 8.8 | 48.9 | 0.0 | 1.2 | 0.0 | 2.9 | 0.0 |
| Gemini-3.1-flash-lite-preview | 79.9 | 91.7 | 11.8 | 41.5 | 3.8 | 0.5 | 2.8 | 7.9 | 14.3 |
| MiniMax-M2.7 | 87.8 | 91.4 | 3.6 | 23.3 | 0.0 | -3.5 | 0.0 | -0.9 | 2.4 |
| Qwen3.6-35B-A3B-256k | 67.2 | 90.6 | 23.4 | 50.0 | 1.4 | 8.7 | 30.8 | 26.6 | 23.1 |

#### Table 19c | Dimension key (atomic checks per dimension)

| Dimension | What it measures | # atomic checks |
|---|---|---|
| Tool grounding | The real domain algorithm was actually invoked, not a stub / hallucinated call | 9 |
| Data integrity | Stored values in the correct scale / layer / dtype (counts preserved, X normalized, shapes sane) | 8 |
| Object plumbing | Artifact placed in the right AnnData slot (obs/var/obsm/uns/layers key present) | 55 |
| Biological plausibility | The result respects known biology (canonical markers, lineage splits, root anchoring) | 12 |
| Quantitative quality | A metric / count / statistical test meets its threshold (ARI, silhouette, kBET/LISI, counts) | 33 |
| Methodological robustness | Multi-method comparison & agreement (>=2 methods, consensus, stability) | 11 |
| TOTAL | all atomic rubric checks observed | 128 |

#### Interpretation (S3)

OmicVerse lifts the native overall (mean of six dimensions) for every model: DeepSeek-v4-pro 85.7->96.4 (+10.7), GLM-5.1 82.3->95.7 (+13.4), GPT-5.5 84.0->95.2 (+11.1), Gemini-3.1-flash-lite 79.9->91.7 (+11.8), MiniMax-M2.7 87.8->91.4 (+3.6), Qwen3.6-35B-A3B 67.2->90.6 (+23.4); DeepSeek-v4-flash 86.1->94.9 (+8.8).

#### Interpretation (S4)

The largest native-profile gain is Qwen3.6-35B-A3B (+23.4, weakest baseline 67.2) and the smallest is MiniMax-M2.7 (+3.6, strongest baseline 87.8): ecosystem grounding helps most when the base model's intrinsic omics execution is weakest (uplift correlates negatively with baseline capability).

#### Interpretation (S5)

The uplift is broad across the intermediate competencies, not just final correctness: the biggest per-dimension gains are typically in tool grounding (invoking the real algorithm) and methodological robustness (multi-method agreement) — the dimensions OmicVerse's registry/skills most directly supply — so augmentation improves the analytical process, not only the answer.

#### Method & caveats

- Native six-dimensional capability scoring (analysis/radar_native.py): every atomic rubric check in OmicBench is mapped to one of six dimensions — tool grounding, data integrity, object plumbing, biological plausibility, quantitative quality, methodological robustness — by a fixed key->dimension table (audit: analysis/native_dim_map.csv). No LLM judge; the same binary check outcomes that drive Pass@1 are reused.
- A (model, arm, dimension) score = passed / total over that dimension's rubric items, POOLED across every task cell and seed (so the denominator per cell is the count of that dimension's checks summed over all tasks). The per-model overall = the mean of the six dimension scores (equal weight per dimension), reproducing the Fig. 2m radar and the manuscript S3 figures.
- Atomic-check counts per dimension (distinct check keys observed across the run): Tool grounding 9, Data integrity 8, Object plumbing 55, Biological plausibility 12, Quantitative quality 33, Methodological robustness 11 (128 distinct checks total). Object plumbing and quantitative quality hold the most checks; data integrity and tool grounding the fewest, so a dimension's score reflects a different number of underlying checks (Table 17c).
- Dimensions are EQUAL-weighted in the overall, so a high-check dimension (quantitative quality) and a low-check one (data integrity) count the same toward the radar mean; per-check pass counts are pooled within each dimension before averaging.
- Scores are pooled per-check pass rates (not per-task), so a dimension dominated by easy presence checks (object plumbing) saturates high; compare dimensions using Table 17c's check counts.
- Equal-weight mean over six dimensions differs from binary Pass@1 (Table 13) and from the per-task majority-seed view (Table 14); a model can have a high native profile yet still fail whole tasks (all-checks-must-pass).
- Single fresh run per model (Qwen single seed); pooling across seeds means a 3-seed model contributes ~3x the check evaluations of Qwen to its own denominator.

### *Supplementary Table 20 | OmicVerse ablation: baseline / no-registry / doc-RAG / full (Extended Data Fig. 18)*

Four arms separating documentation access (doc-RAG), named-tool access (no-registry) and structured registry-grounded execution (full OmicVerse), for GPT-5.5 and DeepSeek-v4-flash, on the common seed over 38 tasks. Pass@1 + atomic-check rate + per-dimension check rates. Arm shaded; task-level matrix in the spreadsheet.

| Model | Arm | Pass@1 | Pass@1 (%) | Atomic checks | Atomic-check rate (%) | Tool grounding | Data integrity | Object plumbing | Biological plausibility | Quantitative quality | Methodological robustness |
|---|---|---|---|---|---|---|---|---|---|---|---|
| GPT-5.5 | Baseline | 28/38 | 73.7 | 141/151 | 93.4 | 10.0 | 100.0 | 100.0 | 100.0 | 97.4 | 100.0 |
| GPT-5.5 | OmicVerse no-registry | 24/38 | 63.2 | 137/151 | 90.7 | 10.0 | 100.0 | 100.0 | 84.6 | 97.4 | 85.7 |
| GPT-5.5 | Doc-RAG | 29/38 | 76.3 | 142/151 | 94.0 | 50.0 | 100.0 | 100.0 | 100.0 | 97.4 | 78.6 |
| GPT-5.5 | Full OmicVerse | 37/38 | 97.4 | 150/151 | 99.3 | 90.0 | 100.0 | 100.0 | 100.0 | 100.0 | 100.0 |
| DeepSeek-v4-flash | Baseline | 27/38 | 71.1 | 133/145 | 91.7 | 30.0 | 88.9 | 95.2 | 100.0 | 97.2 | 100.0 |

| | | | | | | | | | | | |
|---|---|---|---|---|---|---|---|---|---|---|---|
| DeepSeek-v4-flash | OmicVerse no-registry | 32/38 | 84.2 | 145/151 | 96.0 | 60.0 | 100.0 | 100.0 | 92.3 | 97.4 | 100.0 |
| DeepSeek-v4-flash | Doc-RAG | 27/38 | 71.1 | 140/151 | 92.7 | 80.0 | 100.0 | 97.0 | 92.3 | 92.3 | 78.6 |
| DeepSeek-v4-flash | Full OmicVerse | 35/38 | 92.1 | 148/151 | 98.0 | 90.0 | 100.0 | 98.5 | 100.0 | 97.4 | 100.0 |

#### Interpretation (S1–S3)

- Full OmicVerse is the best arm on BOTH Pass@1 and atomic-check rate for BOTH models, and beats the doc-RAG and no-registry variants: GPT-5.5 Pass@1 baseline 73.7% -> no-registry 63.2% -> doc-RAG 76.3% -> full 97.4%; DeepSeek-v4-flash 71.1% -> 84.2% (no-reg) -> 71.1% (doc-RAG) -> 92.1% (full). Documentation access (doc-RAG) and named-tool access (no-registry) each recover only part of the gain — and for GPT no-registry is even BELOW baseline — so the structured registry/contract execution layer, not more text context or tool names alone, drives the benchmark gain (supports S3).

#### Method & caveats

- Four arms separate three candidate mechanisms (S2): (1) Doc-RAG = the base agent given retrieved OmicVerse documentation text (documentation access); (2) OmicVerse no-registry = OmicVerse tools exposed by name but WITHOUT the structured function registry / contract layer (named-tool access); (3) Full OmicVerse = the complete registry-grounded execution ecosystem; vs (0) Baseline = Mini-SWE-Agent alone.
- All four arms are compared on the COMMON seed (seed 0) over the 38 OmicBench tasks, because the ablation arms (no-registry, doc-RAG) were run at a single seed; the baseline and full arms are therefore also restricted to seed 0 here, so their numbers differ slightly from the 3-seed values in Supplementary Tables 13/17.
- Pass@1 = passed tasks / 38 (all rubric checks must pass). Atomic-check rate = passed atomic rubric checks / total checks evaluated (the finer-grained signal that pass/fail hides). Per-dimension columns map each atomic check to a native capability dimension (analysis/radar_native.py).
- Total atomic checks per arm ≈ 145-151 (it varies slightly because a few tasks emit conditional checks depending on what the agent produced).
- Single-seed comparison (seed 0 only), so per-arm Pass@1 has no seed-variance estimate and the baseline/full numbers here are not identical to the multi-seed Tables 13/17.
- no-registry can fall below baseline (GPT-5.5: 63.2% vs 73.7%): exposing OmicVerse tool names without the registry/contract grounding can mislead a model into calling tools it cannot use correctly — itself evidence that the registry layer, not tool access, is the active ingredient.
- Atomic-check rate saturates high (>90%) even at baseline because many checks are easy object-plumbing/presence checks; the discriminating movement is concentrated in tool-grounding and quantitative checks (see per-dimension columns).

### *Supplementary Table 21 | OmicVerse reconstruction packages (github.com/omicverse)*

Complete list of the standalone R/Bioconductor-to-Python reconstruction packages published under the github.com/omicverse organization, with original R source, released Python (PyPI) package name, pipeline scope, release status, PyPI/GitHub links and the OmicVerse integration point. Rows grouped by domain.

| Reconstructed package | Original R source | Source ecosystem | Python package (PyPI) | Pipeline scope | Release status | PyPI link | GitHub link | OmicVerse integration point |
|---|---|---|---|---|---|---|---|---|
| **1. Bulk RNA-seq / expression** | | | | | | | | |
| py-edgeR | edgeR (Robinson, McCarthy & Smyth, Bioinformatics 2010) | Bioconductor | pyedger | bulk RNA-seq differential expression (negative-binomial GLM / quasi-likelihood F-test for count data) | PyPI v0.1.0 (2026-05-20) | https://pypi.org/project/pyedger/ | https://github.com/omicverse/py-edgeR | ov.bulk.pyDEG.deg_analysis(method='edger') |
| py-limma | limma (Ritchie et al., Nucleic Acids Research 2015; eBayes from Smyth 2004) | Bioconductor | pylimma | bulk DE — linear models & empirical-Bayes moderated differential expression (voom/lmFit/eBayes/topTable) | PyPI v0.1.0 (2026-05-01) | https://pypi.org/project/pylimma/ | https://github.com/omicverse/py-limma | ov.bulk.pyDEG (limma/limma-voom backend) |

| | | | | | | | | |
|---|---|---|---|---|---|---|---|---|
| py-mfuzz | Mfuzz (Futschik & Carlisle 2005; Kumar & Futschik 2007); fuzzy c-means core ported from e1071 cmeans | Bioconductor | pymfuzz | bulk/time-series gene expression - fuzzy c-means soft clustering of temporal trajectories | PyPI v0.1.2 (2026-05-20) | https://pypi.org/project/pymfuzz/ | https://github.com/omicverse/py-mfuzz | ov.bulk.temporal_clusters |
| **10. Metabolomics / lipidomics** | | | | | | | | |
| py-lipidr | lipidr (Mohamed, Molendijk & Hill, J. Proteome Res. 2020, 19(7):2890-2897) | Bioconductor | pylipidr | lipidomics — normalization, differential analysis, Lipid Set Enrichment Analysis (LSEA) | PyPI v0.1.0 (2026-05-20) | https://pypi.org/project/pylipidr/ | https://github.com/omicverse/py-lipidr | ov.metabol.read_skyline; ov.metabol.summarize_transitions; ov.metabol.normalize_pqn … |
| **11. Spatial transcriptomics** | | | | | | | | |
| py-SpaceTrooper | SpaceTrooper (Righelli, Banzi, Marchionni et al., 2024; github.com/drighelli/SpaceTrooper) | Bioconductor | py-spacetrooper | spatial QC (image-based spatial transcriptomics: CosMx/MERFISH/Xenium per-cell QC + GLM QC score) | PyPI v0.1.1 (2026-05-28) | https://pypi.org/project/py-spacetrooper/ | https://github.com/omicverse/py-SpaceTrooper | standalone (not wired into ov.*) |
| py-spanorm | SpaNorm (Bhuva DD, Salim A, Mohamed A; github.com/bhuvad/SpaNorm) | Bioconductor | py-spanorm | spatial transcriptomics normalisation (spatially-aware GLM normalisation + SVG calling + GLM-PCA) | PyPI v0.1.0 (2026-05-26) | https://pypi.org/project/py-spanorm/ | https://github.com/omicverse/py-spanorm | standalone (not wired into ov.*) |
| py-spotsweeper | SpotSweeper (github.com/MicTott/SpotSweeper, v1.5.0) | Bioconductor | py-spotsweeper | spatial QC (spatially-aware outlier/artifact detection for spatial transcriptomics: local variance, MAD local outliers, artifact + Visium outlier flagging) | PyPI v0.1.0 (2026-05-26) | https://pypi.org/project/py-spotsweeper/ | https://github.com/omicverse/py-spotsweeper | standalone (not wired into ov.*) |
| **12. Statistical genetics / GWAS** | | | | | | | | |
| py-coloc | coloc (Giambartolomei et al., PLoS Genet. 2014; Wallace 2020/2021; Wakefield 2009) — coloc 5.2.3 | CRAN | pycoloc | GWAS/genetics — Bayesian colocalisation of two genetic traits (Approximate Bayes Factors / SuSiE) | PyPI v0.1.0 (2026-05-20) | https://pypi.org/project/pycoloc/ | https://github.com/omicverse/py-coloc | ov.genetics.colocalize; ov.genetics.finemap_abf; ov.genetics.coloc_sensitivity |
| py-ldsc | LDSC / LD Score Regression (Bulik-Sullivan et al. 2015; Finucane et al. 2015, 2018) | Broad/R | pyldsc | GWAS/genetics — LD score regression, SNP heritability, genetic correlation, partitioned h2, LDSC-SEG | PyPI v0.1.0 (2026-05-20) | https://pypi.org/project/pyldsc/ | https://github.com/omicverse/py-ldsc | ov.genetics.heritability; ov.genetics.genetic_correlation; ov.genetics.partitioned_heritability … |
| py-matrixeqtl | MatrixEQTL (Shabalin, Bioinformatics 2012, 28(10):1353-1358) | CRAN | pymatrixeqtl | GWAS/genetics - fast cis/trans eQTL mapping | PyPI v0.1.0 (2026-05-20) | https://pypi.org/project/pymatrixeqtl/ | https://github.com/omicverse/py-matrixeqtl | ov.genetics.eqtl_map |
| py-scdrs | scDRS (Zhang, Hou et al. & Price, Nature Genetics 2022, 54:1572-1580) | R (other) / Python-original (scDRS is a Python+R tool from the Price lab; GWAS-genetics) | pyscdrs | GWAS/genetics — single-cell disease relevance scoring (cell-level association of GWAS gene sets) | PyPI v0.1.0 (2026-05-20) | https://pypi.org/project/pyscdrs/ | https://github.com/omicverse/py-scdrs | ov.genetics.disease_relevance_score; ov.genetics.score_downstream; ov.genetics.scdrs_cell_type_plot |
| py-susie | susieR (Wang, Sarkar, Carbonetto & Stephens, JRSS-B 2020; Zou et al., PLoS Genetics 2022) | CRAN | pysusie | GWAS/genetics — Bayesian statistical fine-mapping (SuSiE / susie_rss credible sets, PIPs) | PyPI v0.1.0 (2026-05-20) | https://pypi.org/project/pysusie/ | https://github.com/omicverse/py-susie | ov.genetics.finemap; ov.genetics.get_credible_sets; ov.genetics.get_pip … |
| py-twas | MetaXcan / S-PrediXcan / PrediXcan (Barbeira et al., Nature Communications 2018; hakyimlab/MetaXcan) | R (other) | pytwas | GWAS/genetics — transcriptome-wide association study (TWAS): S-PrediXcan, S-MultiXcan, PrediXcan | PyPI v0.1.0 (2026-05-20) | https://pypi.org/project/pytwas/ | https://github.com/omicverse/py-twas | ov.genetics.twas; ov.genetics.load_twas_model; ov.genetics.make_twas_covariance … |
| py-twosamplemr | MendelianRandomization (Yavorska & Burgess, Int. J. Epidemiol. 2017) + TwoSampleMR (Hemani et al., eLife 2018) | CRAN | pytwosamplemr | GWAS/genetics — two-sample Mendelian randomization (IVW/Egger/median/mode/maxlik/cML, harmonisation, diagnostics) | PyPI v0.1.0 (2026-05-20) | https://pypi.org/project/pytwosamplemr/ | https://github.com/omicverse/py-twosamplemr | ov.genetics.mendelian_randomization; ov.genetics.harmonize; ov.genetics.mr_heterogeneity … |
| **13. Hi-C / epigenome** | | | | | | | | |
| py-dcHiC | dcHiC (ay-lab/dcHiC) | R (other) | (see repo) | Hi-C differential compartment analysis (A/B compartment calling + differential compartments) | GitHub-only (standalone repo) | — | https://github.com/omicverse/py-dcHiC | standalone (not wired into ov.*) |
| rust-bandnorm | BandNorm (Shen, Zheng, Keleş 2022; sshen82/BandNorm) | R (other) | bandnorm-rs | single-cell Hi-C — scGAD scoring + band-normalisation for cell-type discovery/embedding | PyPI v0.1.0 (2026-04-26) | https://pypi.org/project/bandnorm-rs/ | https://github.com/omicverse/rust-bandnorm | standalone (not wired into ov.*) |
| rust-scHiCluster | scHiCluster (Zhou et al., PNAS 2019) — single-cell Hi-C contact-matrix | R (other) | schicluster-rs | single-cell Hi-C contact-matrix imputation (RWR + convolution); Hi-C | PyPI v0.1.1 (2026-04-26) | https://pypi.org/project/schicluster-rs/ | https://github.com/omicverse/rust-scHiCluster | standalone (not wired into ov.*) |

| | | | | | | | | |
|---|---|---|---|---|---|---|---|---|
| | imputation (Gaussian convolution + random-walk-with-restart); upstream Python at github.com/zhoujt1994/scHiCluster | | | | | | | |
| **14. Microbiome / amplicon** | | | | | | | | |
| py-dada2 | DADA2 (Callahan et al., Nature Methods 2016) | Bioconductor | pydada2 | amplicon microbiome: exact amplicon sequence variant (ASV) inference from amplicon sequencing | PyPI v0.1.0 (2026-04-19) | https://pypi.org/project/pydada2/ | https://github.com/omicverse/py-dada2 | ov.alignment.dada2_pipeline; ov.alignment.dada2.filter_and_trim; ov.alignment.dada2.learn_errors … |
| **15. Clustering & visualization utilities** | | | | | | | | |
| py-funkyheatmap | funkyheatmap 0.5.2 (R; used for Saelens et al. 2019 dynbenchmark figures) | R/dynverse | pyfunkyheatmap | visualization — dynbenchmark-style data-frame heatmaps (funky rectangles/circles/bars/pies) for benchmark/multi-metric tables | PyPI v0.1.8 (2026-05-25) | https://pypi.org/project/pyfunkyheatmap/ | https://github.com/omicverse/py-funkyheatmap | ov.pl.funky_heatmap |
| py-mclustR | mclust (Scrucca, Fop, Murphy, Raftery 2016) | CRAN | pymclustr | Gaussian mixture model clustering (BIC model selection, 14 covariance parameterizations) for single-cell | PyPI v0.2.1 (2026-05-25) | https://pypi.org/project/pymclustr/ | https://github.com/omicverse/py-mclustR | ov.utils.cluster (method='pymclustR') |
| **2. Single-cell trajectory & dynamics** | | | | | | | | |
| py-condiments | condiments (Roux de Bezieux et al., Nature Communications 2024) | R (other) | pycondiments | trajectory: differential trajectory analysis across conditions in scRNA-seq | PyPI v0.1.0 (2026-05-24) | https://pypi.org/project/pycondiments/ | https://github.com/omicverse/py-condiments | standalone (not wired into ov.*) |
| py-CytoTRACE | CytoTRACE (Gulati et al., Science 2020) | R (other) | pycytotrace-bio | trajectory: gene-count-based stemness / developmental-potential pseudotime for scRNA-seq | PyPI v0.1.0 (2026-05-25) | https://pypi.org/project/pycytotrace-bio/ | https://github.com/omicverse/py-CytoTRACE | ov.single.TrajInfer (method='cytotrace') |
| py-destiny | destiny (Angerer et al., Bioinformatics 2016; theislab/destiny) | Bioconductor | pydestiny-bio | trajectory inference — diffusion maps + diffusion pseudotime (DPT) for scRNA-seq | PyPI v0.2.0 (2026-05-25) | https://pypi.org/project/pydestiny-bio/ | https://github.com/omicverse/py-destiny | ov.single Traj method='destiny'; ov.single Traj method='urd' |
| py-dynbenchmark | dynbenchmark (Saelens et al., Nat Biotechnol 2019) | R/dynverse | pydynbenchmark | trajectory inference benchmarking (dyneval metrics + synthetic generators + harness) | PyPI v0.1.0 (2026-05-25) | https://pypi.org/project/pydynbenchmark/ | https://github.com/omicverse/py-dynbenchmark | standalone (not wired into ov.*) |
| py-dyneval | dyneval (Saelens et al., Nat Biotechnol 2019) | R/dynverse | pydyneval-bio | trajectory inference benchmark metrics | PyPI v0.1.0 (2026-05-25) | https://pypi.org/project/pydyneval-bio/ | https://github.com/omicverse/py-dyneval | standalone (not wired into ov.*) |
| py-dynwrap | dynwrap (dynverse; Saelens et al. 2019) | R/dynverse | pydynwrap-bio | trajectory inference data wrapper / benchmarking (geodesic distances, milestone networks for dyneval/dynbenchmark) | PyPI v0.1.0 (2026-05-25) | https://pypi.org/project/pydynwrap-bio/ | https://github.com/omicverse/py-dynwrap | standalone (not wired into ov.*) |
| py-MARVEL | MARVEL (Wen et al., NAR Genomics Bioinformatics 2023) - single-cell alternative splicing | R (other) | marvel-python | single-cell alternative splicing (PSI quantification, differential splicing, isoform switching) | PyPI v0.1.1 (2026-05-26) | https://pypi.org/project/marvel-python/ | https://github.com/omicverse/py-MARVEL | standalone (not wired into ov.*) |
| py-monocle2 | Monocle 2 (Qiu et al., Nature Methods 2017) | Bioconductor | (see repo) | single-cell trajectory inference (DDRTree pseudotime) | GitHub-only (vendored in external/) | — | https://github.com/omicverse/py-monocle2 | ov.single.Monocle |
| py-Monocle3 | monocle3 (Cao et al., Nature 2019; Trapnell et al., Nat Biotechnol 2014) | Bioconductor | pymonocle3-bio | single-cell trajectory inference (principal-graph pseudotime) | PyPI v0.1.0 (2026-05-25) | https://pypi.org/project/pymonocle3-bio/ | https://github.com/omicverse/py-Monocle3 | ov.single.TrajInfer |
| py-SCORPIUS | SCORPIUS (Cannoodt et al., bioRxiv 2016) | R-dynverse (rcannood/SCORPIUS) | pyscorpius | trajectory — linear trajectory inference for scRNA-seq via MDS + Hastie-Stuetzle principal curves | PyPI v0.1.0 (2026-05-24) | https://pypi.org/project/pyscorpius/ | https://github.com/omicverse/py-SCORPIUS | ov.single.TrajInfer(method='scorpius') |
| py-Slingshot | slingshot (Street et al., BMC Genomics 2018) | Bioconductor | (see repo) | trajectory — cell-lineage (MST/principal-curve) + pseudotime inference for scRNA-seq | GitHub-only (vendored in omicverse) | — | https://github.com/omicverse/py-Slingshot | ov.single.TrajInfer(method='slingshot') |
| py-tradeSeq | tradeSeq (Van den Berge et al., Nature Communications 11:1201, 2020) | Bioconductor | pytradeseq | Trajectory — trajectory-based differential expression for scRNA-seq via NB-GAM (associationTest, startVsEndTest, patternTest) | PyPI v0.1.0 (2026-05-24) | https://pypi.org/project/pytradeseq/ | https://github.com/omicverse/py-tradeSeq | standalone (not wired into ov.*) |

| | | | | | | | | |
|---|---|---|---|---|---|---|---|---|
| py-TSCAN | TSCAN (Ji & Ji, Nucleic Acids Research 2016) | Bioconductor | pytscan | Trajectory — mclust+MST+spline pseudotime reconstruction for scRNA-seq | PyPI v0.1.0 (2026-04-25) | https://pypi.org/project/pytscan/ | https://github.com/omicverse/py-TSCAN | ov.single.TrajInfer (inference(method='TSCAN')) |
| py-URD | URD (Farrell et al., Science 2018; farrellja/URD) | R (other) | pyurd-bio | trajectory — branching pseudotime (flood-based) + random-walk lineage attribution for scRNA-seq | PyPI v0.2.0 (2026-05-25) | https://pypi.org/project/pyurd-bio/ | https://github.com/omicverse/py-URD | ov.single.TrajInfer (method='urd') |
| **3. Single-cell integration / networks** | | | | | | | | |
| py-Augur | Augur (Skinnider et al., Nature Communications 2021) | R (other) | pyaugur | single-cell cell-type prioritization (AUC-based ranking of perturbation-responsive cell types) | PyPI v0.1.0 (2026-05-26) | https://pypi.org/project/pyaugur/ | https://github.com/omicverse/py-Augur | standalone (not wired into ov.*) |
| py-cca | Seurat RunCCA (Stuart, Butler, Hoffman, Hafemeister et al., Cell 2019) | R (other) | pyccasc | single-cell batch integration (canonical correlation analysis) | PyPI v0.1.0 (2026-04-20) | https://pypi.org/project/pyccasc/ | https://github.com/omicverse/py-cca | ov.single.batch_correction (methods='cca') |
| py-hdWGCNA | hdWGCNA (Morabito et al., Cell Reports Methods 2023) | R (other) | py-hdwgcna | single-cell co-expression network analysis (WGCNA on scRNA-seq) | PyPI v0.1.0 (2026-04-30) | https://pypi.org/project/py-hdwgcna/ | https://github.com/omicverse/py-hdWGCNA | standalone (not wired into ov.*) |
| py-SigXTalk | SigXTalk — deep-learning (HGNN) framework dissecting pathway crosstalk from cell-cell communication (no first-author/year given in README) | R (other) | py-sigxtalk | single-cell — cell-cell communication / signaling pathway crosstalk (PRS) inference via hypergraph neural network | PyPI v0.1.2 (2026-05-27) | https://pypi.org/project/py-sigxtalk/ | https://github.com/omicverse/py-SigXTalk | standalone (not wired into ov.*) |
| rust-NMF | NMF (Gaujoux & Seoighe 2010; Renaud Gaujoux's R 'NMF' package) — algorithms: brunet (Brunet et al. PNAS 2004), lee (Lee & Seung 2001), offset (Badea 2008), nsNMF (Pascual-Montano 2006), snmf/r & snmf/l (Kim & Park 2007); plus modern HALS (Cichocki & Phan 2009), E-HALS (Ang & Gillis 2019), dnmf/RcppML (DeBruine 2024) | CRAN | nmf-rs | single-cell gene-program / matrix factorization (NMF) for scRNA-seq; dimensionality reduction & module identification | PyPI v0.1.0 (2026-05-08) | https://pypi.org/project/nmf-rs/ | https://github.com/omicverse/rust-NMF | ov.single.NMF |
| **4. Single-cell metabolism** | | | | | | | | |
| py-scmetabolism | scMetabolism (Wu et al., Cancer Discovery 2021) | R (other) — Y. Wu / Z. Zhang lab R package (not Bioconductor/CRAN core); single-cell metabolism | py-scmetabolism | single-cell metabolism — metabolic pathway-activity scoring (AUCell/VISION/ssGSEA/GSVA over KEGG/REACTOME) | PyPI v0.1.0 (2026-04-24) | https://pypi.org/project/py-scmetabolism/ | https://github.com/omicverse/py-scmetabolism | ov.single.Metabolism (method='scmetabolism') |
| **5. Single-cell differential abundance** | | | | | | | | |
| py-alakazam | alakazam (Gupta et al., Bioinformatics 2015) | R/Immcantation | pyalakazam | immune repertoire (AIRR-seq: Hill diversity, gene usage, clonal abundance, CDR3 AA properties, BCR lineage) | PyPI v0.1.1 (2026-05-21) | https://pypi.org/project/pyalakazam/ | https://github.com/omicverse/py-alakazam | ov.airr.hill_diversity; ov.airr.aa_properties; ov.airr.clonal_abundance … |
| py-miloR | Milo / miloR (Dann et al., Nature Biotechnology 2022) | Bioconductor | (see repo) | single-cell differential abundance testing on kNN neighbourhoods | GitHub-only (vendored in omicverse) | — | https://github.com/omicverse/py-miloR | ov.single.Milo |
| py-Statial | Statial (Ameen, Iyengar, Qin, Ghazanfar, Patrick, 2022; github.com/SydneyBioX/Statial) | Bioconductor | pystatial | spatial proteomics / single-cell spatial (cell-state changes vs spatial associations: Kontextual L-function, distances, abundances, state changes) | PyPI v0.1.2 (2026-05-27) | https://pypi.org/project/pystatial/ | https://github.com/omicverse/py-Statial | standalone (not wired into ov.*) |
| **6. Doublet & ambient-RNA QC** | | | | | | | | |
| py-decontx | DecontX / decontX (Yang et al., Genome Biology 2020; campbio/decontX, celda Bioconductor) | Bioconductor | pydecontx | single-cell RNA-seq ambient/cross-contaminating RNA decontamination | PyPI v0.1.0 (2026-05-22) | https://pypi.org/project/pydecontx/ | https://github.com/omicverse/py-decontx | ov.pp.ambient.remove_ambient(method='decontx'); ov.pp.ambient.estimate_contamination(method='decontx') |
| py-DoubletFinder | DoubletFinder (McGinnis et al., Cell Systems 2019) | R (other) | pydoubletfinder | doublet detection (scRNA-seq) | PyPI v0.1.0 (2026-04-18) | https://pypi.org/project/pydoubletfinder/ | https://github.com/omicverse/py-DoubletFinder | ov.pp.qc; ov.pp.qc_cpu_gpu_mixed |

| | | | | | | | | |
|---|---|---|---|---|---|---|---|---|
| py-fastcar | FastCAR (Gocht/Berg et al., BMC Genomics 2023) | R (other) | pyfastcar | single-cell ambient-RNA removal / QC (deterministic per-gene ambient-RNA correction for droplet scRNA-seq) | PyPI v0.1.0 (2026-05-22) | https://pypi.org/project/pyfastcar/ | https://github.com/omicverse/py-fastcar | ov.pp.ambient.remove_ambient(method='fastcar') |
| py-sccdc | scCDC (Wang et al., Genome Biology 2024) | R (other) | pysccdc | single-cell ambient-RNA decontamination (entropy-based gene-specific GCG detection and correction for scRNA-seq/snRNA-seq) | PyPI v0.1.0 (2026-05-22) | https://pypi.org/project/pysccdc/ | https://github.com/omicverse/py-sccdc | ov.pp.ambient.remove_ambient(method='sccdc') |
| py-scDblFinder | scDblFinder (Germain et al., F1000Research 2022) | Bioconductor | pyscdblfinder | doublet detection (classifier-based: artificial doublets -> cxds -> kNN features -> xgboost iterative scoring -> thresholding) for scRNA-seq | PyPI v0.1.0 (2026-04-18) | https://pypi.org/project/pyscdblfinder/ | https://github.com/omicverse/py-scDblFinder | ov.pp.qc (doublets_method='scdblfinder'); ov.pp.scdblfinder |
| py-soupx | SoupX (Young & Behjati, GigaScience 2020, 9(12):giaa151) | R (other) | pysoupx | single-cell QC — ambient ('soup') RNA contamination removal from droplet scRNA-seq | PyPI v0.1.0 (2026-05-22) | https://pypi.org/project/pysoupx/ | https://github.com/omicverse/py-soupx | ov.pp.remove_ambient(method='soupx'); ov.pp.estimate_contamination |
| **7. Single-cell CNV** | | | | | | | | |
| py-CopyKAT | CopyKAT (Gao et al., Nat Biotechnol 2021) | R (other) | pycopykat | single-cell CNV: aneuploid/diploid classification and bin-level CNA calling from scRNA-seq | GitHub-only (vendored / optional) | — | https://github.com/omicverse/py-CopyKAT | ov.single.CNV |
| py-inferCNV | inferCNV (Broad Institute / Tickle, Tirosh et al.) | Broad/R | (see repo) | single-cell CNV inference from scRNA-seq | GitHub-only (optional runtime import) | — | https://github.com/omicverse/py-inferCNV | ov.single.CNV |
| **8. Immune repertoire (AIRR)** | | | | | | | | |
| py-dowser | dowser (Hoehn, Pybus & Kleinstein, PLoS Comput. Biol. 2022) | R/Immcantation | pydowser | immune repertoire / B-cell receptor phylogenetics (AIRR-seq) | PyPI v0.1.1 (2026-05-21) | https://pypi.org/project/pydowser/ | https://github.com/omicverse/py-dowser | ov.airr.reconstruct_germlines; ov.airr.lineage_trees; ov.airr.lineage_tests |
| py-gliph | turboGliph 0.99.2 (R wrapper of GLIPH, Glanville et al. Nature 2017, and GLIPH2, Huang et al. Nat. Biotechnol. 2020) | R (other) | pygliph | immune repertoire / TCR CDR3 specificity-group clustering | PyPI v0.1.0 (2026-05-21) | https://pypi.org/project/pygliph/ | https://github.com/omicverse/py-gliph | ov.airr.specificity_groups |
| py-immunarch | immunarch 0.10.3 (Nazarov & ImmunoMind team) | CRAN | pyimmunarch | immune repertoire / bulk TCR-BCR (AIRR-seq) analytics: diversity, clonality, overlap, gene usage, public clonotypes | PyPI v0.1.1 (2026-05-21) | https://pypi.org/project/pyimmunarch/ | https://github.com/omicverse/py-immunarch | ov.airr.repertoire_diversity; ov.airr.repertoire_overlap_bulk; ov.airr.gene_usage_bulk … |
| py-scoper | scoper (Immcantation, Kleinstein Lab, Yale) — Spectral Clustering for clOne Partitioning | R/Immcantation (CRAN scoper) | pyscoper | immune repertoire — B-cell clonal clustering/partitioning of AIRR-seq BCR data (identical / hierarchical / spectral) | PyPI v0.1.0 (2026-05-21) | https://pypi.org/project/pyscoper/ | https://github.com/omicverse/py-scoper | ov.airr.clonal_clustering |
| py-shazam | shazam (Gupta, Vander Heiden et al., Bioinformatics 2015; Kleinstein Lab, Yale) — upstream CRAN shazam 1.3.2 | R/Immcantation (CRAN) | pyshazam | immune repertoire — B-cell somatic hypermutation (SHM) targeting & BASELINe antigen-driven selection analysis | PyPI v0.1.0 (2026-05-21) | https://pypi.org/project/pyshazam/ | https://github.com/omicverse/py-shazam | ov.airr.distance_threshold; ov.airr.clonal_clustering; ov.airr.mutation_analysis … |
| py-tigger | tigger (Immcantation / Kleinstein Lab, Yale) — tigger 1.1.3 | R/Immcantation | pytigger | Immune repertoire (AIRR-Seq) — immunoglobulin novel-allele discovery & V genotyping | PyPI v0.1.1 (2026-05-21) | https://pypi.org/project/pytigger/ | https://github.com/omicverse/py-tigger | ov.airr.find_novel_alleles; ov.airr.infer_genotype |
| **9. Proteomics / mass-spec** | | | | | | | | |
| py-DEqMS | DEqMS (Zhu et al., Molecular & Cellular Proteomics 2020) | Bioconductor | pydeqms | proteomics differential expression (peptide-count-aware moderated t-tests for mass-spec) | PyPI v0.1.1 (2026-05-20) | https://pypi.org/project/pydeqms/ | https://github.com/omicverse/py-DEqMS | ov.protein.de(method='deqms') |
| py-imputeLCMD | imputeLCMD (Lazar et al., J Proteome Res 2016) | Bioconductor | pyimputelcmd | proteomics — left-censored missing-value imputation for label-free LC-MS/MS | PyPI v0.1.1 (2026-05-20) | https://pypi.org/project/pyimputelcmd/ | https://github.com/omicverse/py-imputeLCMD | ov.protein.pp.impute |
| py-MSstats | MSstats (Choi et al., Bioinformatics 2014) | Bioconductor | pymsstats | mass-spec proteomics (DDA/DIA): summarization, group comparison, sample-size design | PyPI v0.3.0 (2026-05-20) | https://pypi.org/project/pymsstats/ | https://github.com/omicverse/py-MSstats | ov.protein.summarize; ov.protein.sample_size; ov.protein.contrast_matrix |
| py-OlinkAnalyze | OlinkAnalyze 3.8.2 (Olink Proteomics AB) | CRAN | pyolinkanalyze | proteomics (Olink NPX QC, bridge/subset | PyPI v0.2.1 (2026-05-20) | https://pypi.org/project/pyolinkanalyze/ | https://github.com/omicverse/py-OlinkAnalyze | standalone (not wired into ov.*) |

| | | | | | | | | |
|---|---|---|---|---|---|---|---|---|
| | | | | normalization, per-protein differential expression) | | | | |
| py-proDA | proDA (Ahlmann-Eltze & Anders 2020) | Bioconductor | pyproda | proteomics (MNAR probabilistic differential expression for label-free mass-spec; censored-ML dropout model) | PyPI v0.1.1 (2026-05-20) | https://pypi.org/project/pyproda/ | https://github.com/omicverse/py-proDA | ov.protein.de (method='proda') |

#### Summary

- Total reconstruction packages (github.com/omicverse): 62 — 59 pure-Python py- + 3 Rust rust-.
- Released on PyPI: 56 (pip-installable). GitHub-only: 6 — py-CopyKAT, py-dcHiC, py-inferCNV, py-miloR, py-monocle2, py-Slingshot (vendored into omicverse or optional runtime imports; not published as standalone PyPI dists).
- Wired into a public ov. integration point: 45; standalone (no ov. entry yet): 17.
- PyPI naming convention: repo py-X → dist pyx (lowercased, de-hyphenated). Exceptions: py-CytoTRACE→pycytotrace-bio, py-destiny→pydestiny-bio, py-URD→pyurd-bio, py-Monocle3→pymonocle3-bio, py-dyneval→pydyneval-bio, py-dynwrap→pydynwrap-bio, py-cca→pyccasc, py-MARVEL→marvel-python; some keep the hyphen (py-hdwgcna, py-scmetabolism, py-sigxtalk, py-spanorm, py-spotsweeper, py-spacetrooper). Rust dists use <tool>-rs (nmf-rs, bandnorm-rs, schicluster-rs).
- Release cadence: the bulk of the catalog was published to PyPI in April–May 2026 (v0.1–v0.3).

### *Supplementary Table 22 | OmicVerse ecosystem architecture: shared entry points*

How reconstructed-R packages (github.com/omicverse py-*/rust-*) and existing native-Python packages both plug into shared OmicVerse entry points over a unified AnnData/MuData object model (the IBE substrate).

| Shared entry point (ov.*) | Scope | Shared object | Dispatch mechanism | Reconstructed-R backends (← original R tool) | Existing-Python backends (← origin) |
|---|---|---|---|---|---|
| ov.pp.qc(doublets_method=…) | QC / doublet detection for scRNA-seq | AnnData (cells × genes) | doublets_method= if/elif → conditional import of _scrublet/_sccomposite/_doubletfinder/_scdblfinder | py-DoubletFinder (←DoubletFinder, McGinnis 2019); py-scDblFinder (←scDblFinder, Germain 2022) | scrublet (←vendored scanpy fork); sccomposite (←native-Python composite detector) |
| ov.pp.ambient.remove_ambient | Ambient/soup-RNA decontamination (droplet scRNA-seq) | AnnData (raw counts; decontaminated .X/layer) | method= → _DISPATCH dict → backend runner | py-soupx (←SoupX, Young & Behjati 2020); py-decontx (←DecontX, Yang 2020); py-fastcar (←FastCAR, Gocht 2023); py-sccdc (←scCDC, Wang 2024) | cellbender (←deep-learning, not wired); scar (←deep-learning, not wired) |
| ov.pp.ambient.estimate_contamination | Report-only ambient-RNA fraction estimation (counts untouched) | AnnData (counts untouched); returns pandas.Series | method= restricted to _NATIVE_METHODS (no DL wrappers) | py-soupx (←SoupX, Young & Behjati 2020); py-decontx (←DecontX, Yang 2020); py-fastcar (←FastCAR, Gocht 2023); py-sccdc (←scCDC, Wang 2024) | — |
| ov.single.TrajInfer(method=…) | Single-cell trajectory inference / pseudotime | AnnData (X + obsm embeddings) | method= in .inference() → inline import of backend wrapper (11 algorithms) | py-Slingshot (←slingshot, Street 2018); py-SCORPIUS (←SCORPIUS, Cannoodt 2016); py-TSCAN (←TSCAN, Ji & Ji 2016); py-destiny (←destiny, Angerer 2016); py-URD (←URD, Farrell 2018); py-Monocle3 (←monocle3, Cao 2019); py-CytoTRACE (←CytoTRACE, Gulati 2020) | palantir (←vendored diffusion-map/pseudotime); VIA (←vendored graph/topology); sctour (←PyPI deep-learning TNODE); StaVIA (←omicverse spatial VIA extension) |
| ov.single.Velo | RNA velocity / directional state transitions | AnnData (spliced/unspliced or velocity layers) | method= in .cal_velocity() | — | dynamo (←PyPI, Qiu 2022); scvelo (←PyPI, Bergen 2020); latentvelo (←vendored, Boileau 2024); graphvelo (←vendored, Gao 2023); regvelo (←PyPI, Huang 2024) |
| ov.single.batch_correction | Batch-effect correction / integration | AnnData (integrated embeddings → obsm) | methods= string → _BATCH_OBSM key routing; kwargs via _split_kwargs_by_signature() | py-cca/pyccasc (←Seurat::RunCCA, Stuart 2019) | harmonypy (←vendored harmony); scanorama (←vendored); cellanova (←vendored); scanpy (←pp.combat); scvi-tools (←scVI/scANVI/totalVI); scArches (←scPoli); concord-sc (←PyPI contrastive) |

| | | | | | |
|---|---|---|---|---|---|
| ov.single.CNV(method=…) | Single-cell copy-number variation inference | AnnData (obsm['X_cnv'], obs cnv_prediction/score, uns['cnv']) | method= at class init → _run_copykat()/_run_infercnv() | py-CopyKAT (←CopyKAT, Gao 2021); py-inferCNV (←inferCNV, Tickle/Tirosh) | — |
| ov.single.Metabolism(method=…) | Single-cell metabolic-state inference + metabolite CCC | AnnData (obsm['X_metabolism'], uns['metabolism']) | method= at class init (3 values) | py-scmetabolism (←scMetabolism, Wu 2021) | scfea (←vendored native-Python GNN flux); mebocost (←vendored metabolite-sensor CCC); Compass (←precomputed CPLEX/Gurobi output loader) |
| ov.single.grn(method=…) | Gene regulatory network inference | AnnData (cells × genes, optional layer) | method= selects grnboost2/genie3/regdiffusion | — | arboreto (←PyPI, GRNBoost2/GENIE3); regdiffusion (←PyPI neural diffusion) |
| ov.single.run_liana / cellphonedb_v5 / pyCEFCON | Cell-cell communication (LR inference, crosstalk) | AnnData → unified comm AnnData via to_comm_adata() (means/pvalues layers) | Three independent entry points (not unified); to_comm_adata() standardizes results | py-SigXTalk (←SigXTalk HGNN crosstalk; PyPI, not wired) | liana (←PyPI saezlab); cellphonedb (←PyPI); CEFCON (←vendored, Wang 2023); mebocost (←vendored); pyscenic (←vendored SCENIC port); commot (←vendored); flowsig (←vendored); nocd (←vendored, optional torch) |
| ov.single.NMF / cNMF / Hotspot | Single-cell gene programs / matrix factorization | AnnData (cells × genes) | Separate class entry points (no unified dispatcher); within NMF, method= selects rust algorithm variant | rust-NMF/nmf-rs (←NMF, Gaujoux & Seoighe 2010; brunet/lee/offset/nsNMF/snmf/hals/ehals/dnmf) | cnmf (←PyPI/vendored, Kotliar 2019); hotspot (←PyPI/vendored, DeTomaso & Yosef 2021) |
| ov.single.MetaCell(method=…) | Metacell construction (7 pluggable backends) | AnnData (obs metacell_id/conf, obsm X_metacell, uns) | method= → BACKEND_REGISTRY lookup → backend class → unified MetaCell wrapper | — | SEACells (←vendored, Persad 2023); MetaQ (←vendored, Li 2025); supercell (←vendored, Bilous 2022); geosketch (←PyPI, Hie 2019); metacells/MC2 (←PyPI, Ben-Kiki 2022) |
| ov.bulk.pyDEG | Bulk RNA-seq differential expression | pandas.DataFrame (genes × samples) | method= in deg_analysis() (5 methods) | py-edgeR (←edgeR, Robinson 2010); py-limma (←limma, Ritchie 2015) | pydeseq2 (←PyPI DESeq2); plus inmoose (edgepy GLM-LRT) / scipy (ttest) |
| ov.bulk.pyWGCNA | Bulk co-expression network / module detection | pandas.DataFrame (samples × genes) | Direct class instantiation (single unified impl; no method=) | PyWGCNA (←WGCNA, Langfelder & Horvath 2008; vendored) | — |
| ov.protein.de | Proteomics differential expression (LC-MS/MS & Olink) | AnnData (samples × proteins; log2 abundance) | method= string selector (8 values) | py-DEqMS (←DEqMS, Zhu 2020); py-proDA (←proDA, Ahlmann-Eltze & Anders 2020) | statsmodels (←PyPI; olink_lmer LMM); plus limma (vendored in pydeqms), scipy.stats (wilcoxon/welch_t/anova/kruskal) |
| ov.protein.impute | Proteomics missing-value imputation (LCMD / MAR-MNAR) | AnnData (samples × proteins; NaN-marked) | method= string selector (12 values; auto runs model_selector) | py-imputeLCMD (←imputeLCMD, Lazar 2016) | — (half_min/min implemented in NumPy) |
| ov.airr | Immune repertoire (single-cell & bulk TCR/BCR) | AnnData (airr obs schema) / pandas.DataFrame (AIRR) / pyimmunarch.ImmunData | method= per-function + direct function wrapping; lazy backend imports | pyalakazam (←alakazam, Gupta 2015); pyimmunarch (←immunarch); pyscoper (←scoper); pyshazam (←shazam, Gupta 2015); pytigger (←tigger); pydowser (←dowser, Hoehn 2022); pygliph (←turboGliph/GLIPH2) | scirpy (←reimplemented in-house single-cell TCR/BCR I/O + metrics) |
| ov.space.clusters | Spatial domain clustering / embedding | AnnData (obsm['spatial'] + X) | methods= list-of-strings loop; per-method kwargs | SpatialDE (←SpatialDE, Svensson 2018; for SVG); NaiveDE (←NaiveDE, Svensson 2019) | STAGATE_pyG (←native PyTorch GAT); GraphST (←vendored contrastive); CAST (←vendored); BINARY (←vendored autoencoder); Banksy (←pip external) |
| ov.space.Deconvolution(method=…) | Spatial deconvolution / cell-type mapping | AnnData (spots × genes; reference cells × genes) | method= if/elif chain (5 methods) | — | Tangram (←PyPI tangram-sc); cell2location (←vendored, BayraktarLab); FlashDeconv (←PyPI, Feng); Starfysh (←vendored, SrivatsanLab); rctd-py (←PyPI binding, separately maintained) |
| ov.genetics.* | Statistical genetics / GWAS suite (eQTL, fine-mapping, coloc, MR, scDRS, LDSC, TWAS, spatial GWAS) | Heterogeneous: GWAS summary stats (DataFrame), genotypes (matrices), AnnData (scDRS only) | method= per-task dispatcher; deferred _require_*() imports at call-time | py-matrixeqtl (←MatrixEQTL, Shabalin 2012); py-susie (←susieR, Wang 2020); py-coloc (←coloc, Giambartolomei 2014); py-twosamplemr (←MendelianRandomization/ TwoSampleMR, Burgess/Hemani); py-scdrs (←scDRS, Zhang 2022); py-ldsc (←LDSC, Bulik-Sullivan 2015); py-twas (←MetaXcan/S-PrediXcan, Barbeira 2018) | gsmap (←vendored native-Python spatial GWAS mapping) |
| ov.llm.SCLLMManager / ModelFactory | Single-cell foundation models (scLLMs) | AnnData (cells × genes + obs/obsm/varm) | model_type= → _models registry → model class (predict(task=…)) | — | scgpt, geneformer, scfoundation, uce, cellplm (←third-party/HF); scmulan, tosica, popv (←vendored); + 16 stub-registered models (scbert/genecompass/nicheformer/…) |

| | | | | | |
|---|---|---|---|---|---|
| ov.epi | Epigenomics (scATAC, bulk ATAC/ChIP/CUT&RUN, chromVAR, Hi-C) | AnnData (cells × tiles/peaks/bins) | Lazy pass-through to epione via __getattr__ + delegate wrappers (no method= in ov.epi) | rust-scHiCluster (←scHiCluster, Zhou 2019); rust-bandnorm (←BandNorm, Shen 2022); py-dcHiC (←dcHiC, ay-lab) | snapatac2 (←PyPI scATAC); cooler (←PyPI Hi-C I/O); MOODS (←PyPI motif scan); pysam (←PyPI BAM/SAM I/O) |

#### Architecture narrative (the IBE substrate)

A single object model as the shared substrate. Nearly every OmicVerse entry point reads from and writes to a unified AnnData (cells × genes / samples × features) — extended to MuData for multimodal data. Backends agree on a fixed input/output contract within each module: pseudotime in adata.obs, embeddings in adata.obsm (e.g. X_palantir, X_pca_harmony, X_metacell, X_cnv), result matrices in adata.obsm/layers, and method metadata in adata.uns. This common schema is what makes downstream, backend-agnostic tooling possible — e.g. CopyKAT and inferCNV both populate obsm['X_cnv']/obs['cnv_prediction'], so a single ov.pl.cnv_heatmap plots either; CCC backends are normalized into one communication AnnData (means/pvalues layers) via to_comm_adata(). The bulk and statistical-genetics modules are the deliberate exceptions: ov.bulk.pyDEG/pyWGCNA operate on pandas.DataFrame count/expression matrices, and ov.genetics.* is intentionally heterogeneous (GWAS summary stats as DataFrames, genotypes as matrices, AnnData only for scDRS) because forcing those tasks into AnnData would be unnatural.

One dispatch pattern over one namespace. The dominant idiom is a method= (or methods=/model_type=/doublets_method=) string argument on a single class or function that routes to a backend behind an identical call signature. Routing is implemented as if/elif chains (Deconvolution, qc), explicit dispatch dictionaries (ambient._DISPATCH, MetaCell.BACKEND_REGISTRY, batch_correction._BATCH_OBSM), or inline per-branch imports (TrajInfer.inference). Because the dispatcher mediates pre/post-processing and schema writing, a reconstructed-R port (py-Slingshot) and a native-Python tool (palantir) are invoked indistinguishably — only the method= string differs. A minority of areas deliberately keep parallel class entry points rather than a single dispatcher: NMF/cNMF/Hotspot are fundamentally different workflows, and the CCC layer (run_liana/cellphonedb_v5/pyCEFCON) keeps three independent entry points unified only at the result-formatting stage.

Reconstructed-R-dominated vs existing-Python-dominated entry points. The reconstruction strategy concentrates where the mature methodology has historically lived in R/Bioconductor: bulk DE (py-edgeR, py-limma), proteomics (py-DEqMS, py-proDA, py-imputeLCMD), immune repertoire (seven Immcantation/CRAN ports under ov.airr), statistical genetics (seven ports under ov.genetics), ambient-RNA and doublet QC (py-soupx/py-decontx/py-fastcar/py-sccdc, py-DoubletFinder/py-scDblFinder), CNV (py-CopyKAT/py-inferCNV), and co-expression networks (PyWGCNA). By contrast, fields that were Python-first are wired almost entirely to existing-Python tools: RNA velocity (dynamo/scvelo/regvelo/…), spatial deconvolution (Tangram/cell2location/Starfysh/…), GRN/CCC (arboreto/liana/cellphonedb/pyscenic/commot), metacells (SEACells/MetaQ/supercell/…), and single-cell foundation models (ov.llm, ~21 native deep-learning models, zero R ports).

Boundary-blending and the IBE feature. The architecturally distinctive entry points are those that place reconstructed-R ports and existing-Python tools behind the same dispatcher — TrajInfer (7 R ports + 4 Python), batch_correction (1 R + 9 Python), pyDEG (2 R + 3 Python), ov.protein.de (2 R + statsmodels/scipy), ov.airr (7 R + in-house scirpy), ov.genetics (7 R + gsmap), ov.space.clusters (SpatialDE/NaiveDE + 5 Python), ov.epi (3 rust/R + 4 Python), and the QC/ambient dispatchers. A few sit at the category boundary by design: StaVIA is an omicverse spatial extension of the reconstructed py-VIA; CEFCON is itself an R-reconstruction grouped among Python CCC backends; rust-NMF and the rust Hi-C tools are Rust reimplementations of R packages.

Lazy loading and optional dependencies. The whole ecosystem is held together by deferred imports. Backends are imported only inside the dispatch branch that needs them (TrajInfer), behind _require_*() helpers (ov.genetics), via __getattr__ pass-through (ov.epi → epione), or lazily within ov.airr so import omicverse.airr succeeds with no optional extras installed. This keeps the top-level package importable with a minimal core while heavyweight or solver-bound tools (CellBender/scAR, scvi-tools, CPLEX/Gurobi-backed Compass, CLIP-scale foundation-model checkpoints) remain opt-in extras. Graceful fallbacks reinforce robustness (e.g. scdblfinder falls back to scrublet when py-scDblFinder is absent), and not-yet-wired tools (CellBender, scAR, py-SigXTalk, the 16 stub LLMs) are registered but raise informative errors or remain standalone until their runtimes are wired.

#### Summary counts

- Shared entry points: 22 (merging the two ov.pp.ambient functions and the parallel-class groups as listed; 20 module rows in the table).
- Distinct reconstructed-R backends mapped: 38 — py-DoubletFinder, py-scDblFinder, py-soupx, py-decontx, py-fastcar, py-sccdc, py-Slingshot, py-SCORPIUS, py-TSCAN, py-destiny, py-URD, py-Monocle3, py-CytoTRACE, py-cca/pyccasc, py-CopyKAT, py-inferCNV, py-scmetabolism, py-SigXTalk, rust-NMF, py-edgeR, py-limma, PyWGCNA, py-DEqMS, py-proDA, py-imputeLCMD, pyalakazam, pyimmunarch, pyscoper, pyshazam, pytigger, pydowser, pygliph, SpatialDE, NaiveDE, py-matrixeqtl, py-susie, py-coloc,

py-twosamplemr, py-scdrs, py-ldsc, py-twas, rust-scHiCluster, rust-bandnorm, py-dcHiC. (≈44 if SpatialDE/NaiveDE and the rust ports are all counted; ~38 strict py-* R reconstructions.)

- Distinct existing-Python backends mapped: ~70+ across modules (e.g. palantir, VIA, sctour, StaVIA, dynamo, scvelo, latentvelo, graphvelo, regvelo, harmonypy, scanorama, cellanova, scanpy, scvi-tools, scArches, concord-sc, scrublet, sccomposite, cellbender, scar, scfea, mebocost, Compass, arboreto, regdiffusion, liana, cellphonedb, CEFCON, pyscenic, commot, flowsig, nocd, cnmf, hotspot, SEACells, MetaQ, supercell, geosketch, metacells, pydeseq2, statsmodels, scirpy, STAGATE_pyG, GraphST, CAST, BINARY, Banksy, Tangram, cell2location, FlashDeconv, Starfysh, rctd-py, gsmap, snapatac2, cooler, MOODS, pysam, ~21 ov.llm foundation models).
- Entry points that blend BOTH classes (the key architectural feature): 9 — ov.pp.qc, ov.pp.ambient.remove_ambient, ov.single.TrajInfer, ov.single.batch_correction, ov.single.Metabolism, ov.bulk.pyDEG, ov.protein.de, ov.airr, ov.space.clusters, ov.genetics, ov.epi (10–11 if the single-class CCC grouping with py-SigXTalk is included).
- Single-class entry points: reconstructed-R-only → 4 (ov.pp.ambient.estimate_contamination, ov.single.CNV, ov.bulk.pyWGCNA, ov.protein.impute); existing-Python-only → 7 (ov.single.Velo, ov.single.grn, ov.single.MetaCell, ov.single.NMF/cNMF/Hotspot, ov.space.Deconvolution, ov.llm, plus the CCC run_liana/cellphonedb_v5/pyCEFCON cluster).

### *Supplementary Table 23 | Parity taxonomy (output class → metric → threshold → example → failure interpretation)*

Pre-registered parity tests are matched to the class of the output they validate (Fig. 3a,c). For each output class: comparison principle, metric, acceptance threshold, an example reconstructed package with its observed value, and how a parity failure should be interpreted.

| Output class | Comparison principle (invariance) | Parity metric | Acceptance threshold | Example package (observed value) | Failure interpretation |
|---|---|---|---|---|---|
| Dense numeric matrix / vector (deterministic) | Numerical identity — a deterministic kernel has no free invariance, so outputs must match bit-for-bit up to floating-point round-off | max\|Δ\| (absolute / relative error); Pearson r on flattened values | Bit-equivalence: max\|Δ\| < 1e-10–1e-12 for deterministic kernels; else abs err < 1e-6 / r = 1.0 | py-matrixeqtl (β max\|diff\| < 2e-15, p < 2e-14); py-coloc (PP max abs diff < 3e-16); py-MSstats (LogIntensities 5e-14); py-soupx / py-fastcar (bit-for-bit); py-inferCNV (per-step < 1e-10) | Exceeding tolerance on a deterministic kernel is a genuine numerical/algorithmic bug (wrong formula, indexing, normalization order) — it cannot be blamed on randomness and must be fixed before release. |
| Test statistics & p-values (DE / association) | Rank-of-significance + top-hit overlap — absolute p-values are noise-sensitive near 0, so the conclusion (which features are significant) matters more than the exact value | Pearson/Spearman on −log10 p or effect size; Jaccard / % agreement of top-N hits | Spearman ≥ 0.70–0.84; top-N (e.g. top-500) 100% or top-50 Jaccard ≥ 0.7 | py-DEqMS (sca.t/sca.P r = 1.0, top-500 100% agreement); py-tradeSeq (associationTest Spearman 0.84, top-50 Jaccard 0.67); py-edgeR / py-limma (stated parity) | Loss of top-hit overlap means the two implementations reach different biological conclusions and is disqualifying; mild divergence in the insignificant tail (high p) is tolerable and expected. |
| Low-dimensional embedding (PCA / MDS / diffusion / CCA) | Invariance-aware — embeddings are defined up to rotation, sign-flip and scaling, so compare subspaces, not coordinates | Procrustes correlation; column-span / projector Frobenius distance; per-component canonical correlation; eigenvalue Pearson | Procrustes ≥ 0.80; projector distance < 5e-3; singular-value rel err < 1e-5; eigenvalue Pearson ≥ 0.95 | py-destiny (eigenvectors Procrustes 0.916, eigenvalues Pearson 0.967); py-cca (singular-value rel err < 1e-5, projector dist < 5e-3); py-SCORPIUS (low-dim Procrustes 0.999) | Divergence beyond a rigid transform means the manifold geometry differs → neighbor graphs, clusters and trajectories built on it may shift. A raw coordinate mismatch alone is NOT a failure (rotation/sign are expected). |
| Clustering / discrete group labels | Label-permutation-invariant — cluster IDs are arbitrary, only the partition matters | Adjusted Rand Index (ARI); AMI / NMI; Cohen's κ | ARI ≥ 0.90 (strict: ≥ 0.95) | py-CopyKAT (aneuploid/diploid ARI 0.988, κ 0.994); py-scoper (clonal ARI > 0.95); py-mclustR (classification 1.0000); py-mfuzz (hard-assignment ARI ~0.99) | Low ARI means objects are grouped differently — a genuine structural disagreement, since label identity is permuted out. A relabeled-but-identical partition correctly scores ARI = 1. |
| Pseudotime / 1-D ordering | Rank- or correlation-based — absolute pseudotime scale and offset are arbitrary, only the ordering matters | Spearman (rank) / Pearson on pseudotime | Spearman ≥ 0.80 (typically achieved ≥ 0.95) | py-Slingshot (Spearman ≥ 0.99); py-URD (Spearman 0.985 / Pearson 0.992); py-TSCAN (Pearson 1.000000); py-monocle2 (≥ 0.99) | Rank divergence means cells are ordered differently along the trajectory → different developmental interpretation. Differences in absolute pseudotime values (scale/offset) are not failures. |
| Tree / graph topology (lineages, phylogenies) | Topology-aware — compare structure, not node coordinates or labels | Robinson-Foulds distance; maximum-parsimony score; graph isomorphism; HIM distance; F1 on branch assignment | Parsimony score exact / RF small / isomorphic (\|Δ\| = 0) | py-dowser (getTrees parsimony exact; RF small for ambiguous clones); py-dyneval (him / edge_flip / isomorphic / F1_branches \|Δ\| = 0); py-URD (branch attribution) | Topology divergence means the inferred branching/lineage structure differs → changed biological relationships. Small RF on genuinely ambiguous clones is tolerable; large structural divergence is not. |
| Discrete feature sets (credible sets, alleles, ASVs, motifs, compartments) | Set-membership equality / overlap | Exact set equality; Jaccard index; membership agreement % | Exact match (Jaccard = 1.0) for deterministic; Jaccard ≥ 0.7 for thresholded selection | py-susie (credible sets identical); py-tigger (novel-allele table & genotype match R exactly); py-dada2 (ASV-level identity); py-dcHiC | A different recovered set = different downstream targets (different causal SNPs, novel alleles, ASVs). Set disagreement maps directly to |

| | | | | | |
|---|---|---|---|---|---|
| | | | | (differential compartments top-50 Jaccard 0.96) | a different scientific output and is interpretable per-element. |
| Stochastic / RNG-bound outputs (bootstrap, MCMC, random init) | Distributional equivalence — exact reproduction is impossible when RNG streams differ, so compare distributions / point estimates, not elements | Kolmogorov–Smirnov test on sampled values; Pearson on point estimates with bootstrap SE within Monte-Carlo tolerance; CI overlap | KS p > 0.01; point estimate exact + SE within ~10% (Monte-Carlo error) | py-imputeLCMD (QRILC/MinProb KS p > 0.01; MinDet/ZERO bit-exact); py-alakazam (alphaDiversity bootstrap r > 0.999, CIs within a few %); py-twosamplemr (closed-form bit-exact; bootstrap SE within MC error) | Only a difference in the distribution (not the seed) indicates a wrong stochastic model. A per-element mismatch under a different RNG stream is expected and is NOT a failure — the parity test deliberately targets the distribution / closed-form estimate. |

#### How the taxonomy is applied (S1–S3 protocol)

- S1 — reference-grounded specification. For each target package the R reference implementation plus shared test fixtures (canonical example datasets bundled with the R package, e.g. coloc_test_data, dowser ExampleAirr, MiSeq SOP) define expected behaviour. The reconstruction is validated against running R outputs, not a loose reimplementation from the paper text or high-level docs (Fig. 3a).

- S2 — equivalence agent. An agent iteratively translates the R behaviour into a candidate Python package and compares Python outputs against the R reference under pre-registered parity tests committed before implementation (tests/test_r_parity.py / test_exact_match.py in each repo).

- S3 — output-class-matched tests. Each parity test is selected from the taxonomy above according to the class of the output it checks — a single package typically registers several (e.g. py-destiny checks an embedding by Procrustes, eigenvalues by Pearson, and pseudotime by rank correlation simultaneously). A package passes only when every registered class-appropriate test clears its threshold.

#### Summary

- 8 output classes in the parity taxonomy, spanning the four named in S3 (exact matrices, embeddings, clustering, pseudotime) plus four observed in practice (test statistics/p-values, tree/graph topology, discrete feature sets, stochastic/RNG-bound outputs).
- Metric families: absolute/relative numerical error & Pearson (matrices); rank correlation + top-N overlap (statistics, pseudotime); invariance-aware geometry (Procrustes, subspace distance — embeddings); permutation-invariant agreement (ARI/AMI — clustering); topology distances (RF, parsimony, isomorphism — trees); set equality/Jaccard (discrete sets); KS / bootstrap-CI (stochastic).
- Threshold philosophy: deterministic kernels are held to bit-equivalence (max|Δ| < 1e-10); everything defined up to an invariance is held to a saturating similarity gate (correlation/ARI/Procrustes ≥ 0.8–1.0); stochastic outputs are held to distributional equivalence.
- Failure interpretation key: a failed deterministic-matrix test is a code bug; a failed invariance-class test (embedding/clustering/pseudotime/topology/set) is a genuine scientific disagreement, not a relabeling artifact; a failed stochastic test is meaningful only at the distribution level, never per-element.

### *Supplementary Table 24 | Per-package acceleration evolution statistics*

Closed-loop acceleration agent (policy→action→environment→reward, Fig. 3b): proposes admissibility-proven rewrites — (E) exact identity, (B) bounded ε-approximation, (C) class-containment (Fig. 3c) — and accepts only those that are faster while clearing the equivalence gate. Full ITERATION_LOG telemetry is published for 7 packages.

#### Part A | Per-package evolution statistics

| Package | Baseline parity (class) | Iterations | Accepted | Rejected (by reason) | Admissibility classes | Final speedup | Final parity | Documented limitations |
|---|---|---|---|---|---|---|---|---|
| py-Statial | 24 parity tests (smoke+structural) | 5 | 5 | 0 (—) | E | ≈1× (correctness port; Class A translation-only) | 24/24 pass | Pure correctness port — iters 2–4 were bug fixes (NaN handling, kNN self-match, type cast), no acceleration attempted |
| py-TSCAN | Pearson 1.000000 (pseudotime/ordinal) | 2 | 1 | 1 (1 REJECT_SLOW) | E | 1.18× vs baseline (≈28× vs R via translation) | Pearson 1.000000 (pass) | Playbook exhausted: dataset too small (131 cells, G≤9) for |

| | | | | | | | | |
|---|---|---|---|---|---|---|---|---|
| | | | | | | | | cache-XtX/Woodbury/sparse rewrites; Class A |
| py-spanorm | correlation 0.974 | 4 | 3 | 0 (—) | E | 7.5× | correlation unchanged (pass) | none documented |
| py-spotsweeper | 3/5 parity tests failing | 5 | 4 | 0 (—) | E,B | 3–19× | 5/5 pass (F1=1.0, ARI=1.0) | residual IRLS (R MASS::rlm) + cross-language kNN tie-breaking differences; manifest relaxed to deterministic-bounded 1e-5 |
| py-Augur | parity 0.9839 (AUC correlation) | 6 | 4 | 2 (1 REJECT_GATE, 1 REJECT_SLOW) | E,B | 2.15× final (3.02× peak @iter5) | 0.9999 (pass) | none documented |
| py-SigXTalk | total 1268 s (Pearson parity gate 0.95) | 6 | 5 | 0 (—) | E,B | 15.85× | Pearson 0.9912 (pass) | n_estimators 10 vs R 500 (7× larger SE); ε≈0.0088 bounded-approx within 0.95 gate |
| py-SpaceTrooper | Pearson 0.7399 (FAILS gate 0.90) | 6 | 5 | 1 (1 REJECT_GATE) | E,B | 102× | Pearson 0.9891 (pass) | max_abs_error 0.194; residual λ difference (sklearn LogisticRegression CV vs R cv.glmnet λ-sequence/CV strategy) |
| Total (7 packages) | — | 34 | 27 | 4 (2 GATE, 2 SLOW) | E (all 7), B (4) | up to 102× | 7/7 pass | — |

### Part B | Accepted vs rejected rewrites

| Package (iter) | Rewrite action | Admissibility class | Parity before→after | Runtime before→after | Decision |
|---|---|---|---|---|---|
| py-SpaceTrooper (iter 5) | Fixed λ=7.9 matching R glmnet (grid-searched C=0.1265) | (B) bounded-ε | 0.951 → 0.989 | baseline → 102× faster | ACCEPT |
| py-SigXTalk (iter 4) | Random-Forest n_estimators 500→10 | (B) bounded-ε (ε≈0.0088) | 0.9912 → 0.9912 | 3.45× → 15.85× | ACCEPT |
| py-SigXTalk (iter 3) | Vectorize Spearman via precomputed ranks + np.corrcoef | (E) exact | 0.9853 → 0.9912 | corr 80 s → corr 2.37 s | ACCEPT |
| py-Augur (iter 5) | Custom _FastRandomForest bypassing sklearn param validation | (E) exact | 0.9977 → 0.9980 | baseline → 3.02× | ACCEPT |
| py-spanorm (iter 2) | Vectorized NB log-PMF (gammaln+xlogy+xlog1py) | (E) exact | 0.974 → 0.974 | 1.0× → 6.2× | ACCEPT |
| py-TSCAN (iter 1) | np.linalg.svd(compute_uv=False) on first SVD | (E) exact | 1.000000 → 1.000000 | baseline → 1.18× | ACCEPT |
| py-Augur (iter 1) | Sorted sliding-window loess (assume 1-D kNN contiguous) | attempted (E) — invalid | 0.9839 → 0.7530 | — → — | REJECT_GATE (parity below threshold) |
| py-SpaceTrooper (iter 0) | sklearn LogisticRegression default (lbfgs solver) | — | — → 0.7399 | — → — | REJECT_GATE (0.7399 < 0.90 gate) |
| py-Augur (iter 3) | Parallel tree building n_jobs=-1 | (E) exact | 0.9977 → 0.9977 | 60 s → 170.7 s | REJECT_SLOW (joblib overhead on 600 cells) |
| py-TSCAN (iter 2) | scipy.linalg.svd lapack_driver='gesdd' | (E) exact | 1.000000 → 1.000000 | prev → 0.34× (slower) | REJECT_SLOW (wrapper overhead at 131×16776) |

### Part C | Final speedups (broader catalogue, no per-iteration telemetry)

| Package | Final speedup vs R | Backend |
|---|---|---|
| py-CopyKAT | median 23.4x faster on 8 cores (per-sample 23.5x / 46.8x / 82.4x on 1k/3k/7k-cell ladder) | pure-Python (NumPy/SciPy/Numba) |
| py-DEqMS | 4.78x (4.8x); 88x including Rscript startup | pure-Python/numpy (vectorised NumPy/SciP |
| py-MARVEL | 3.23x (plate full replay), 2.96x (droplet replay) | pure-Python/numpy |
| py-MSstats | 2.5x-3.9x (2.53x pipeline; 3.91x incl. Rscript startup; groupComparison 6.20x) | pure-Python/numpy (NumPy/SciPy/statsmode |
| py-hdWGCNA | 3.1x total (44.62s R vs 14.43s Python); per-step up to 17.4x (TestSoftPowers), 13.8x (ModuleConnectivity), 9.8x (MetacellsByGroups), 5.6x (ConstructNetwork) | pure-Python/numpy (AnnData-native; MiniB |
| py-immunarch | ~60x (Python pipeline vs R pipeline, includes Rscript startup) | pure-Python/numpy (numpy, scipy, pandas, |
| py-inferCNV | 25-192x py-vs-R | pure-Python/numpy + numba JIT |
| py-lipidr | ~8x | pure-Python/numpy/scipy (reuses pygoslin |
| py-monocle2 | 30-100x faster than R (102x on pancreas 3,696 cells, 'fast' mode) | pure-Python/numpy (NumPy/SciPy; fast DDR |
| py-proDA | ~equal inner-loop (Py ~3.4s vs R proDA pipeline ~3.3s); ~2.7x end-to-end vs R total ~9.1s (avoids Rscript startup) | pure-Python/numpy (scipy.optimize.L-BFGS |
| py-scmetabolism | 3-45x faster than R (VISION 45.7x, GSVA 43.5x, AUCell 3.5x, ssGSEA 3.0x) | pure-Python/numpy (optimized algorithms; |
| py-tradeSeq | 5x faster than R fitGAM | pure-Python (statsmodels.gam, vs R mgcv) |
| rust-NMF | 76x (lee, bit-eq) up to ~215-280x (hals+NNDSVD); ~250x (dnmf+NNDSVD), 16 threads | Rust (PyO3/maturin) + rayon + matrixmult |

| | | |
|---|---|---|
| rust-bandnorm | ~24x (10-30x depending on dataset) | Rust (PyO3/maturin) |
| rust-scHiCluster | ~9.6x on long chromosomes (chr1), ~2.7x end-to-end per cell; +3.1x from multiprocess parallelism | Rust (PyO3/maturin); steps 2-9 in Rust, |

#### Summary

- Packages with full iteration telemetry: 7 (ITERATION_LOG.md). Total acceleration iterations: 34 → 27 accepted, 4 rejected (2 REJECT_GATE = parity dropped below threshold; 2 REJECT_SLOW = no wall-clock gain). No REJECT_INADMISSIBLE recorded (inadmissible actions are filtered before execution).
- Admissibility classes observed: (E) exact identity dominates (all 7 packages); (B) bounded ε-approximation in 4 (Augur loess it=2, SigXTalk RF 500→10, SpaceTrooper fixed-λ, spotsweeper IRLS). (C) class-containment is in the playbook (MST⊆Delaunay) but not exercised in these 7 logged ports.
- Speedup range (logged packages): ≈1× (py-Statial / py-TSCAN — correctness-only / dataset-too-small 'Class A' ports) to 102× (py-SpaceTrooper). Broader catalogue (Part C) ranges to ~280× (rust-NMF) and ~192× (py-inferCNV).
- Two recurring rejection patterns: (i) an assumed exact identity that is actually false (py-Augur iter 1: 1-D kNN contiguity assumption → parity 0.984→0.753 → REJECT_GATE); (ii) a parallelisation/library swap whose overhead exceeds the benefit at the fixture size (py-Augur iter 3 joblib; py-TSCAN iter 2 scipy gesdd → REJECT_SLOW).
- Coverage limitation: structured policy/action/reward telemetry is currently published for 7 of the ~62 reconstructed packages; a complete Results section would require running the rebuildr loop with logging enabled across the full catalogue. Part C documents the available final speedups for the remainder.

### *Supplementary Table 25 | Per-package reconstruction landscape*

One row per reconstructed/targeted R package in the Fig. 3d landscape: methodology family, source ecosystem, scope, version, completeness and release status. 58 components across 5 ecosystems and 24 families.

| R package | Methodology family | Source ecosystem | Pipeline scope | Version | Completeness | Release status | GitHub |
|---|---|---|---|---|---|---|---|
| **Bioconductor (22)** | | | | | | | |
| MARVEL | Alternative splicing | Bioconductor | single-cell alternative splicing (PSI quantification, differential splicing, isoform switching) | 0.1.1 | Full | PyPI v0.1.1 (2026-05-26) | https://github.com/omicverse/py-MARVEL |
| inferCNV | CNV / aneuploidy | Bioconductor | single-cell CNV inference from scRNA-seq | — | Full | GitHub-only (optional runtime import) | https://github.com/omicverse/py-inferCNV |
| proDA | Censored / probabilistic DE | Bioconductor | proteomics (MNAR probabilistic differential expression for label-free mass-spec; censored-ML dropout model) | 0.1.1 | Full | PyPI v0.1.1 (2026-05-20) | https://github.com/omicverse/py-proDA |
| destiny | Diffusion / random-walk trajectory | Bioconductor | trajectory inference — diffusion maps + diffusion pseudotime (DPT) for scRNA-seq | 0.2.0 | Full | PyPI v0.2.0 (2026-05-25) | https://github.com/omicverse/py-destiny |
| DEqMS | Empirical-Bayes moderated t | Bioconductor | proteomics differential expression (peptide-count-aware moderated t-tests for mass-spec) | 0.1.1 | Full | PyPI v0.1.1 (2026-05-20) | https://github.com/omicverse/py-DEqMS |
| edgeR | GLM / linear models | Bioconductor | bulk RNA-seq differential expression (negative-binomial GLM / quasi-likelihood F-test for count data) | 0.1.0 | Core subset (GLM/QL-F path) | PyPI v0.1.0 (2026-05-20) | https://github.com/omicverse/py-edgeR |
| limma | GLM / linear models | Bioconductor | bulk DE — linear models & empirical-Bayes moderated differential expression (voom/lmFit/eBayes/topTable) | 0.1.0 | Core subset (lmFit/eBayes/voom) | PyPI v0.1.0 (2026-05-01) | https://github.com/omicverse/py-limma |
| miloR | GLM / linear models | Bioconductor | single-cell differential abundance testing on kNN neighbourhoods | — | Full | GitHub-only (vendored in omicverse) | https://github.com/omicverse/py-miloR |

| | | | | | | | |
|---|---|---|---|---|---|---|---|
| MSstats | GLM / linear models | Bioconductor | mass-spec proteomics (DDA/DIA): summarization, group comparison, sample-size design | 0.3.0 | Full | PyPI v0.3.0 (2026-05-20) | https://github.com/omicverse/py-MSstats |
| Slingshot | Graph-based trajectory | Bioconductor | trajectory — cell-lineage (MST/principal-curve) + pseudotime inference for scRNA-seq | — | Full | GitHub-only (vendored in omicverse) | https://github.com/omicverse/py-Slingshot |
| TSCAN | Graph-based trajectory | Bioconductor | Trajectory — mclust+MST+spline pseudotime reconstruction for scRNA-seq | 0.1.0 | Full | PyPI v0.1.0 (2026-04-25) | https://github.com/omicverse/py-TSCAN |
| DADA2 | Iterative denoising / ambient cleanup | Bioconductor | amplicon microbiome: exact amplicon sequence variant (ASV) inference from amplicon sequencing | 0.1.0 | Full | PyPI v0.1.0 (2026-04-19) | https://github.com/omicverse/py-dada2 |
| decontX | Mixture / model-based clustering | Bioconductor | single-cell RNA-seq ambient/cross-contaminating RNA decontamination | 0.1.0 | Full | PyPI v0.1.0 (2026-05-22) | https://github.com/omicverse/py-decontx |
| Mfuzz | Mixture / model-based clustering | Bioconductor | bulk/time-series gene expression - fuzzy c-means soft clustering of temporal trajectories | 0.1.2 | Full | PyPI v0.1.2 (2026-05-20) | https://github.com/omicverse/py-mfuzz |
| lipidr | Pathway / scoring | Bioconductor | lipidomics — normalization, differential analysis, Lipid Set Enrichment Analysis (LSEA) | 0.1.0 | Full | PyPI v0.1.0 (2026-05-20) | https://github.com/omicverse/py-lipidr |
| scDblFinder | Random-forest classification | Bioconductor | doublet detection (classifier-based: artificial doublets -> cxds -> kNN features -> xgboost iterative scoring -> thresholding) for scRNA-seq | 0.1.0 | Full | PyPI v0.1.0 (2026-04-18) | https://github.com/omicverse/py-scDblFinder |
| SpaceTrooper | Spatial analysis & QC | Bioconductor | spatial QC (image-based spatial transcriptomics: CosMx/MERFISH/Xenium per-cell QC + GLM QC score) | 0.1.1 | Full | PyPI v0.1.1 (2026-05-28) | https://github.com/omicverse/py-SpaceTrooper |
| SpaNorm | Spatial analysis & QC | Bioconductor | spatial transcriptomics normalisation (spatially-aware GLM normalisation + SVG calling + GLM-PCA) | 0.1.0 | Full | PyPI v0.1.0 (2026-05-26) | https://github.com/omicverse/py-spanorm |
| SpotSweeper | Spatial analysis & QC | Bioconductor | spatial QC (spatially-aware outlier/artifact detection for spatial transcriptomics: local variance, MAD local outliers, artifact + Visium outlier flagging) | 0.1.0 | Full | PyPI v0.1.0 (2026-05-26) | https://github.com/omicverse/py-spotsweeper |
| Statial | Spatial analysis & QC | Bioconductor | spatial proteomics / single-cell spatial (cell-state changes vs spatial associations: Kontextual L-function, distances, abundances, state changes) | 0.1.2 | Full | PyPI v0.1.2 (2026-05-27) | https://github.com/omicverse/py-Statial |
| condiments | Trajectory DE / benchmarking | Bioconductor | trajectory: differential trajectory analysis across conditions in scRNA-seq | 0.1.0 | Partial (topologyTest simplified in v0.1) | PyPI v0.1.0 (2026-05-24) | https://github.com/omicverse/py-condiments |
| tradeSeq | Trajectory DE / benchmarking | Bioconductor | Trajectory — trajectory-based differential expression for scRNA-seq via NB-GAM (associationTest, startVsEndTest, patternTest) | 0.1.0 | Partial (associationTest full; startVsEndTest approx) | PyPI v0.1.0 (2026-05-24) | https://github.com/omicverse/py-tradeSeq |
| **CRAN (8)** | | | | | | | |
| coloc | Bayesian fine-mapping / coloc | CRAN | GWAS/genetics — Bayesian colocalisation of two genetic traits | 0.1.0 | Full | PyPI v0.1.0 (2026-05-20) | https://github.com/omicverse/py-coloc |

| | | | | | | | |
|---|---|---|---|---|---|---|---|
| | | | (Approximate Bayes Factors / SuSiE) | | | | |
| SuSiE | Bayesian fine-mapping / coloc | CRAN | GWAS/genetics — Bayesian statistical fine-mapping (SuSiE / susie_rss credible sets, PIPs) | 0.1.0 | Full | PyPI v0.1.0 (2026-05-20) | https://github.com/omicverse/py-susie |
| imputeLCMD | Imputation | CRAN | proteomics — left-censored missing-value imputation for label-free LC-MS/MS | 0.1.1 | Full | PyPI v0.1.1 (2026-05-20) | https://github.com/omicverse/py-imputeLCMD |
| SoupX | Iterative denoising / ambient cleanup | CRAN | single-cell QC — ambient ('soup') RNA contamination removal from droplet scRNA-seq | 0.1.0 | Full | PyPI v0.1.0 (2026-05-22) | https://github.com/omicverse/py-soupx |
| OlinkAnalyze | Mixed-effects models | CRAN | proteomics (Olink NPX QC, bridge/subset normalization, per-protein differential expression) | 0.2.1 | Full | PyPI v0.2.1 (2026-05-20) | https://github.com/omicverse/py-OlinkAnalyze |
| mclust | Mixture / model-based clustering | CRAN | Gaussian mixture model clustering (BIC model selection, 14 covariance parameterizations) for single-cell | 0.2.1 | Full | PyPI v0.2.1 (2026-05-25) | https://github.com/omicverse/py-mclustR |
| immunarch | Repertoire & B-cell phylogeny | CRAN | immune repertoire / bulk TCR-BCR (AIRR-seq) analytics: diversity, clonality, overlap, gene usage, public clonotypes | 0.1.1 | Full | PyPI v0.1.1 (2026-05-21) | https://github.com/omicverse/py-immunarch |
| MatrixEQTL | eQTL / MR / TWAS | CRAN | GWAS/genetics - fast cis/trans eQTL mapping | 0.1.0 | Full | PyPI v0.1.0 (2026-05-20) | https://github.com/omicverse/py-matrixeqtl |
| **Immunogenomics (Immcantation) (5)** | | | | | | | |
| alakazam | Repertoire & B-cell phylogeny | Immcantation | immune repertoire (AIRR-seq: Hill diversity, gene usage, clonal abundance, CDR3 AA properties, BCR lineage) | 0.1.1 | Full | PyPI v0.1.1 (2026-05-21) | https://github.com/omicverse/py-alakazam |
| dowser | Repertoire & B-cell phylogeny | Immcantation | immune repertoire / B-cell receptor phylogenetics (AIRR-seq) | 0.1.1 | Full | PyPI v0.1.1 (2026-05-21) | https://github.com/omicverse/py-dowser |
| scoper | Repertoire & B-cell phylogeny | Immcantation | immune repertoire — B-cell clonal clustering/partitioning of AIRR-seq BCR data (identical / hierarchical / spectral) | 0.1.0 | Full | PyPI v0.1.0 (2026-05-21) | https://github.com/omicverse/py-scoper |
| shazam | Repertoire & B-cell phylogeny | Immcantation | immune repertoire — B-cell somatic hypermutation (SHM) targeting & BASELINe antigen-driven selection analysis | 0.1.0 | Full | PyPI v0.1.0 (2026-05-21) | https://github.com/omicverse/py-shazam |
| tigger | Repertoire & B-cell phylogeny | Immcantation | Immune repertoire (AIRR-Seq) — immunoglobulin novel-allele discovery & V genotyping | 0.1.1 | Full | PyPI v0.1.1 (2026-05-21) | https://github.com/omicverse/py-tigger |
| **dynverse (4)** | | | | | | | |
| SCORPIUS | Graph-based trajectory | dynverse | trajectory — linear trajectory inference for scRNA-seq via MDS + Hastie-Stuetzle principal curves | 0.1.0 | Full | PyPI v0.1.0 (2026-05-24) | https://github.com/omicverse/py-SCORPIUS |
| dynbenchmark | Trajectory DE / benchmarking | dynverse | trajectory inference benchmarking (dyneval metrics + synthetic generators + harness) | 0.1.0 | Partial (metrics+generators harness; no parity gate) | PyPI v0.1.0 (2026-05-25) | https://github.com/omicverse/py-dynbenchmark |
| dyneval | Trajectory DE / benchmarking | dynverse | trajectory inference benchmark metrics | 0.1.0 | Partial (8/14 metric IDs ported) | PyPI v0.1.0 (2026-05-25) | https://github.com/omicverse/py-dyneval |
| dynwrap | Trajectory DE / benchmarking | dynverse | trajectory inference data wrapper / benchmarking (geodesic distances, milestone networks for dyneval/dynbenchmark) | 0.1.0 | Full | PyPI v0.1.0 (2026-05-25) | https://github.com/omicverse/py-dynwrap |

| Academic GitHub (19) | | | | | | | |
|---|---|---|---|---|---|---|---|
| CopyKAT | CNV / aneuploidy | Academic GitHub | single-cell CNV: aneuploid/diploid classification and bin-level CNA calling from scRNA-seq | — | Full | GitHub-only (vendored / optional) | https://github.com/omicverse/py-CopyKAT |
| SigXTalk | Cell–cell communication | Academic GitHub | single-cell — cell-cell communication / signaling pathway crosstalk (PRS) inference via hypergraph neural network | 0.1.2 | Core (PRS crosstalk; HGNN) | PyPI v0.1.2 (2026-05-27) | https://github.com/omicverse/py-SigXTalk |
| CytoTRACE | Diffusion / random-walk trajectory | Academic GitHub | trajectory: gene-count-based stemness / developmental-potential pseudotime for scRNA-seq | 0.1.0 | Full | PyPI v0.1.0 (2026-05-25) | https://github.com/omicverse/py-CytoTRACE |
| URD | Diffusion / random-walk trajectory | Academic GitHub | trajectory — branching pseudotime (flood-based) + random-walk lineage attribution for scRNA-seq | 0.2.0 | Full | PyPI v0.2.0 (2026-05-25) | https://github.com/omicverse/py-URD |
| Monocle 2 | Graph-based trajectory | Academic GitHub | single-cell trajectory inference (DDRTree pseudotime) | — | Full | GitHub-only (vendored in external/) | https://github.com/omicverse/py-monocle2 |
| Monocle 3 | Graph-based trajectory | Academic GitHub | single-cell trajectory inference (principal-graph pseudotime) | 0.1.0 | Wrapper (re-exports Monocle3-python) | PyPI v0.1.0 (2026-05-25) | https://github.com/omicverse/py-Monocle3 |
| dcHiC | Hi-C / 3D genome | Academic GitHub | Hi-C differential compartment analysis (A/B compartment calling + differential compartments) | — | Full | GitHub-only (standalone repo) | https://github.com/omicverse/py-dcHiC |
| FastCAR | Iterative denoising / ambient cleanup | Academic GitHub | single-cell ambient-RNA removal / QC (deterministic per-gene ambient-RNA correction for droplet scRNA-seq) | 0.1.0 | Full | PyPI v0.1.0 (2026-05-22) | https://github.com/omicverse/py-fastcar |
| scCDC | Iterative denoising / ambient cleanup | Academic GitHub | single-cell ambient-RNA decontamination (entropy-based gene-specific GCG detection and correction for scRNA-seq/snRNA-seq) | 0.1.0 | Full | PyPI v0.1.0 (2026-05-22) | https://github.com/omicverse/py-sccdc |
| LDSC | LD score regression / heritability | Academic GitHub | GWAS/genetics — LD score regression, SNP heritability, genetic correlation, partitioned h2, LDSC-SEG | 0.1.0 | Full | PyPI v0.1.0 (2026-05-20) | https://github.com/omicverse/py-ldsc |
| Seurat CCA | Multivariate integration | Academic GitHub | single-cell batch integration (canonical correlation analysis) | 0.1.0 | Full | PyPI v0.1.0 (2026-04-20) | https://github.com/omicverse/py-cca |
| scMetabolism | Pathway / scoring | Academic GitHub | single-cell metabolism — metabolic pathway-activity scoring (AUCell/VISION/ssGSEA/GSVA over KEGG/REACTOME) | 0.1.0 | Full | PyPI v0.1.0 (2026-04-24) | https://github.com/omicverse/py-scmetabolism |
| scDRS | Polygenic scoring | Academic GitHub | GWAS/genetics — single-cell disease relevance scoring (cell-level association of GWAS gene sets) | 0.1.0 | Full | PyPI v0.1.0 (2026-05-20) | https://github.com/omicverse/py-scdrs |
| Augur | Random-forest classification | Academic GitHub | single-cell cell-type prioritization (AUC-based ranking of perturbation-responsive cell types) | 0.1.0 | Full | PyPI v0.1.0 (2026-05-26) | https://github.com/omicverse/py-Augur |
| DoubletFinder | Random-forest classification | Academic GitHub | doublet detection (scRNA-seq) | 0.1.0 | Full | PyPI v0.1.0 (2026-04-18) | https://github.com/omicverse/py-DoubletFinder |
| GLIPH | Repertoire & B-cell phylogeny | Academic GitHub | immune repertoire / TCR CDR3 specificity-group clustering | 0.1.0 | Full | PyPI v0.1.0 (2026-05-21) | https://github.com/omicverse/py-gliph |
| hdWGCNA | Topological overlap network | Academic GitHub | single-cell co-expression network analysis | 0.1.0 | Full | PyPI v0.1.0 (2026-04-30) | https://github.com/omicverse/py-hdWGCNA |

| | | | | | | | |
|---|---|---|---|---|---|---|---|
| | | | (WGCNA on scRNA-seq) | | | | |
| S-PrediXcan | eQTL / MR / TWAS | Academic GitHub | GWAS/genetics — transcriptome-wide association study (TWAS): S-PrediXcan, S-MultiXcan, PrediXcan | 0.1.0 | Full | PyPI v0.1.0 (2026-05-20) | https://github.com/omicverse/py-twas |
| TwoSampleMR | eQTL / MR / TWAS | Academic GitHub | GWAS/genetics — two-sample Mendelian randomization (IVW/Egger/median/mode/maxlik/cML, harmonisation, diagnostics) | 0.1.0 | Full | PyPI v0.1.0 (2026-05-20) | https://github.com/omicverse/py-twosampleмr |

#### Summary

- 58 reconstructed/targeted components across 5 source ecosystems and 24 methodology families (Fig. 3d).
- Ecosystem split: 22 Bioconductor, 8 CRAN, 5 immunogenomics, 4 dynverse, 19 academic GitHub.
- Update vs prior snapshot (52→58): added py-SpaceTrooper, py-spotsweeper, py-spanorm, py-Statial (Bioconductor spatial QC/normalization/cell-state), py-dcHiC (academic GitHub, Hi-C 3D-genome) and py-SigXTalk (academic GitHub, cell–cell communication). Bioconductor 18→22, academic GitHub 17→19; CRAN/Immcantation/dynverse unchanged.
- Completeness: most ports are full-function reconstructions; documented exceptions are core-subset ports (py-edgeR, py-limma), a namespace wrapper (py-Monocle3), and partial ports (py-tradeSeq, py-condiments, py-dyneval, py-dynbenchmark).
- Release status: the large majority are published on PyPI (v0.1–v0.3, Apr–May 2026); a handful are github-only (vendored into omicverse or optional runtime imports).

### *Supplementary Table 26 | Per-package reconstruction maturity*

One row per reconstructed/targeted R package in the Fig. 3d landscape: methodology family, source ecosystem, scope, version, completeness and release status. Five distinct maturity statuses per package: ported, validated, optimized, integrated, agent-executable.

| Package | Ecosystem | Methodology family | Ported (completeness) | Validated | Optimized | Integrated (ov.*) | Agent-exec |
|---|---|---|---|---|---|---|---|
| **Bioconductor (22)** | | | | | | | |
| MARVEL | Bioconductor | Alternative splicing | ✓ (Full) | ✓ | ✓ | — | — |
| inferCNV | Bioconductor | CNV / aneuploidy | ✓ (Full) | ✓ | ✓ | ✓ ov.single.CNV | ◐ |
| proDA | Bioconductor | Censored / probabilistic DE | ✓ (Full) | ✓ | ✓ | ✓ ov.protein.de (method='proda') | ✓ |
| destiny | Bioconductor | Diffusion / random-walk trajectory | ✓ (Full) | ✓ | — | ✓ ov.single Traj method='destiny'; ov.single Traj method='urd' | ✓ |
| DEqMS | Bioconductor | Empirical-Bayes moderated t | ✓ (Full) | ✓ | ✓ | ✓ ov.protein.de(method ='deqms') | ✓ |
| edgeR | Bioconductor | GLM / linear models | ✓ (Core subset (GLM/QL-F path)) | ◐ | — | ✓ ov.bulk.pyDEG.deg_analysis(method='edger' ) | ✓ |
| limma | Bioconductor | GLM / linear models | ✓ (Core subset (lmFit/eBayes/voom) ) | ◐ | — | ✓ ov.bulk.pyDEG (limma/limma-voom backend) | ✓ |
| miloR | Bioconductor | GLM / linear models | ✓ (Full) | ◐ | — | ✓ ov.single.Milo | ✓ |
| MSstats | Bioconductor | GLM / linear models | ✓ (Full) | ✓ | ✓ | ✓ ov.protein.summarize; ov.protein.sample_size … | ✓ |
| Slingshot | Bioconductor | Graph-based trajectory | ✓ (Full) | ✓ | — | ✓ ov.single.TrajInfer(method='slingshot') | ✓ |
| TSCAN | Bioconductor | Graph-based trajectory | ✓ (Full) | ✓ | ✓ | ✓ ov.single.TrajInfer (inference(method='TSCAN')) | ✓ |
| DADA2 | Bioconductor | Iterative denoising / ambient cleanup | ✓ (Full) | ✓ | — | ✓ ov.alignment.dada2_pipeline; ov.alignment.dada2.filter_and_trim … | ✓ |

| | | | | | | | |
|---|---|---|---|---|---|---|---|
| decontX | Bioconductor | Mixture / model-based clustering | ✓ (Full) | ✓ | — | ✓ ov.pp.ambient.remove_ambient(method='decontx'); ov.pp.ambient.estimate_contamination(method='decontx') | ✓ |
| Mfuzz | Bioconductor | Mixture / model-based clustering | ✓ (Full) | ✓ | — | ✓ ov.bulk.temporal_clusters | ✓ |
| lipidr | Bioconductor | Pathway / scoring | ✓ (Full) | ◐ | ✓ | ✓ ov.metabol.read_skyline; ov.metabol.summarize_transitions … | ✓ |
| scDblFinder | Bioconductor | Random-forest classification | ✓ (Full) | ◐ | — | ✓ ov.pp.qc (doublets_method='scdblfinder'); ov.pp.scdblfinder | ✓ |
| SpaceTrooper | Bioconductor | Spatial analysis & QC | ✓ (Full) | ✓ | ✓ | — | — |
| SpaNorm | Bioconductor | Spatial analysis & QC | ✓ (Full) | ✓ | ✓ | — | — |
| SpotSweeper | Bioconductor | Spatial analysis & QC | ✓ (Full) | ✓ | ✓ | — | — |
| Statial | Bioconductor | Spatial analysis & QC | ✓ (Full) | ✓ | ✓ | — | — |
| condiments | Bioconductor | Trajectory DE / benchmarking | ✓ (Partial (topologyTest simplified in v0.1)) | ✓ | — | — | — |
| tradeSeq | Bioconductor | Trajectory DE / benchmarking | ✓ (Partial (associationTest full; startVsEndTest approx)) | ✓ | ✓ | — | — |
| **CRAN (8)** | | | | | | | |
| coloc | CRAN | Bayesian fine-mapping / coloc | ✓ (Full) | ✓ | — | ✓ ov.genetics.colocalize; ov.genetics.finemap_abf … | ✓ |
| SuSiE | CRAN | Bayesian fine-mapping / coloc | ✓ (Full) | ◐ | — | ✓ ov.genetics.finemap; ov.genetics.get_credible_sets … | ✓ |
| imputeLCMD | CRAN | Imputation | ✓ (Full) | ✓ | — | ✓ ov.protein.pp.impute | ✓ |
| SoupX | CRAN | Iterative denoising / ambient cleanup | ✓ (Full) | ✓ | — | ✓ ov.pp.remove_ambient (method='soupx'); ov.pp.estimate_contamination | ✓ |
| OlinkAnalyze | CRAN | Mixed-effects models | ✓ (Full) | ✓ | — | — | — |
| mclust | CRAN | Mixture / model-based clustering | ✓ (Full) | ✓ | — | ✓ ov.utils.cluster (method='pymclustR') | ◐ |
| immunarch | CRAN | Repertoire & B-cell phylogeny | ✓ (Full) | ✓ | ✓ | ✓ ov.airr.repertoire_diversity; ov.airr.repertoire_overlap_bulk … | ✓ |
| MatrixEQTL | CRAN | eQTL / MR / TWAS | ✓ (Full) | ✓ | — | ✓ ov.genetics.eqtl_map | ✓ |
| **Immunogenomics (Immcantation) (5)** | | | | | | | |
| alakazam | Immcantation | Repertoire & B-cell phylogeny | ✓ (Full) | ◐ | — | ✓ ov.airr.hill_diversity; ov.airr.aa_properties … | ✓ |
| dowser | Immcantation | Repertoire & B-cell phylogeny | ✓ (Full) | ✓ | — | ✓ ov.airr.reconstruct_germlines; ov.airr.lineage_trees … | ✓ |
| scoper | Immcantation | Repertoire & B-cell phylogeny | ✓ (Full) | ◐ | — | ✓ ov.airr.clonal_clustering | ✓ |
| shazam | Immcantation | Repertoire & B-cell phylogeny | ✓ (Full) | ◐ | — | ✓ ov.airr.distance_threshold; ov.airr.clonal_clustering … | ✓ |
| tigger | Immcantation | Repertoire & B-cell phylogeny | ✓ (Full) | ✓ | — | ✓ ov.airr.find_novel_alleles; ov.airr.infer_genotype | ✓ |
| **dynverse (4)** | | | | | | | |
| SCORPIUS | dynverse | Graph-based trajectory | ✓ (Full) | ✓ | — | ✓ ov.single.TrajInfer(method='scorpius') | ✓ |
| dynbenchmark | dynverse | Trajectory DE / benchmarking | ✓ (Partial (metrics+generators harness; no parity gate)) | ◐ | — | — | — |

| | | | | | | | |
|---|---|---|---|---|---|---|---|
| dyneval | dynverse | Trajectory DE / benchmarking | ✓ (Partial (8/14 metric IDs ported)) | ✓ | — | — | — |
| dynwrap | dynverse | Trajectory DE / benchmarking | ✓ (Full) | ✓ | — | — | — |
| **Academic GitHub (19)** | | | | | | | |
| CopyKAT | Academic GitHub | CNV / aneuploidy | ✓ (Full) | ✓ | ✓ | ✓ ov.single.CNV | ✓ |
| SigXTalk | Academic GitHub | Cell–cell communication | ✓ (Core (PRS crosstalk; HGNN)) | ✓ | ✓ | — | — |
| CytoTRACE | Academic GitHub | Diffusion / random-walk trajectory | ✓ (Full) | ◐ | — | ✓ ov.single.TrajInfer (method='cytotrace') | ✓ |
| URD | Academic GitHub | Diffusion / random-walk trajectory | ✓ (Full) | ✓ | — | ✓ ov.single.TrajInfer (method='urd') | ✓ |
| Monocle 2 | Academic GitHub | Graph-based trajectory | ✓ (Full) | ✓ | ✓ | ✓ ov.single.Monocle | ✓ |
| Monocle 3 | Academic GitHub | Graph-based trajectory | ✓ (Wrapper (re-exports Monocle3-python)) | ◐ | — | ✓ ov.single.TrajInfer | ✓ |
| dcHiC | Academic GitHub | Hi-C / 3D genome | ✓ (Full) | ✓ | — | — | — |
| FastCAR | Academic GitHub | Iterative denoising / ambient cleanup | ✓ (Full) | ✓ | — | ✓ ov.pp.ambient.remove_ambient(method='fastcar') | ✓ |
| scCDC | Academic GitHub | Iterative denoising / ambient cleanup | ✓ (Full) | ✓ | — | ✓ ov.pp.ambient.remove_ambient (method='sccdc') | ✓ |
| LDSC | Academic GitHub | LD score regression / heritability | ✓ (Full) | ✓ | — | ✓ ov.genetics.heritability; ov.genetics.genetic_correlation … | ✓ |
| Seurat CCA | Academic GitHub | Multivariate integration | ✓ (Full) | ✓ | — | ✓ ov.single.batch_correction (methods='cca') | ◐ |
| scMetabolism | Academic GitHub | Pathway / scoring | ✓ (Full) | ✓ | ✓ | ✓ ov.single.Metabolism(method='scmetabolism') | ✓ |
| scDRS | Academic GitHub | Polygenic scoring | ✓ (Full) | ✓ | — | ✓ ov.genetics.disease_relevance_score; ov.genetics.score_downstream … | ✓ |
| Augur | Academic GitHub | Random-forest classification | ✓ (Full) | ✓ | ✓ | — | — |
| DoubletFinder | Academic GitHub | Random-forest classification | ✓ (Full) | ✓ | — | ✓ ov.pp.qc; ov.pp.qc_cpu_gpu_mixed | ✓ |
| GLIPH | Academic GitHub | Repertoire & B-cell phylogeny | ✓ (Full) | ✓ | — | ✓ ov.airr.specificity_groups | ✓ |
| hdWGCNA | Academic GitHub | Topological overlap network | ✓ (Full) | ✓ | ✓ | — | — |
| S-PrediXcan | Academic GitHub | eQTL / MR / TWAS | ✓ (Full) | ✓ | — | ✓ ov.genetics.twas; ov.genetics.load_twas_model … | ✓ |
| TwoSampleMR | Academic GitHub | eQTL / MR / TWAS | ✓ (Full) | ✓ | — | ✓ ov.genetics.mendelian_randomization; ov.genetics.harmonize … | ✓ |

#### Summary

- Ported: 58/58 (all). Completeness: most Full; core-subset (py-edgeR, py-limma), wrapper (py-Monocle3), partial (py-tradeSeq, py-condiments, py-dyneval, py-dynbenchmark).
- Validated: 46 with a quantified parity metric, 12 parity-stated-only, 0 unvalidated.
- Optimized: 19 (7 with full acceleration-loop ITERATION_LOG, the rest with a benchmarked speedup); 39 translation-only / not yet benchmarked.
- Integrated into OmicVerse: 43 have a public ov.* entry point; 15 are standalone repos not yet wired in (spatial-QC quartet, Hi-C trio, dynverse, hdWGCNA, Augur, MARVEL, SigXTalk, OlinkAnalyze, etc.).
- Agent-executable: 40 are registered + shipped (callable out-of-the-box by the agent/MCP); 3 are registered but require an optional pip install (py-cca, py-CopyKAT, py-inferCNV); 15 are not integrated.
- The five statuses are NOT nested: a package can be integrated without being optimized (e.g. genetics ports are integrated + agent-executable but not acceleration-looped), and optimized without being integrated (the spatial-QC ports have ITERATION_LOG telemetry but no ov.* entry yet). This is exactly why they must be reported as distinct columns.

- Scope = the 58 Fig. 3d py-* ports. The 3 Rust ports (rust-NMF, rust-bandnorm, rust-scHiCluster) are optimized-by-construction and integrated for NMF only; the 2 external-only vendored ports (supercell, scTenifold) are integrated but lack standalone repos — both excluded here to match the Fig. 3d landscape.
- 'Agent-executable' reflects the @register_function registry (788 registered functions across the OmicVerse modules) exposed to ov.Agent and the MCP server; a package is only counted ✓ if its registered wrapper can run without an undeclared optional dependency.

#### *Supplementary Table 27 | Per-package trajectory benchmark datasets*

One row per reconstructed/targeted R package in the Fig. 3d landscape: methodology family, source ecosystem, scope, version, completeness and release status. 14 synthetic trajectory datasets: source, topology, cells, genes, gold-standard structure (Fig. 3e).

| Dataset | Source (simulator family) | Generative model | Topology class | Gold-standard trajectory structure | # Cells | # Genes | # Branches |
|---|---|---|---|---|---|---|---|
| **Dyntoy (6)** | | | | | | | |
| dyntoy_bifurcation | Dyntoy | Dyntoy-style (Poisson counts) | bifurcation | Single bifurcation: 1 trunk → 2 branches (3 segments, 1 branch point) | 300 | 100 | 3 |
| dyntoy_convergence | Dyntoy | Dyntoy-style (Poisson counts) | convergence | Convergence: 2 branches → 1 common late state (3 segments, 1 merge point) | 300 | 100 | 3 |
| dyntoy_cycle | Dyntoy | Dyntoy-style (Poisson counts) | cycle | Closed cycle, no defined start/end (1 loop) | 300 | 100 | 1 |
| dyntoy_linear | Dyntoy | Dyntoy-style (Poisson counts) | linear | Linear path, no branch point (1 segment) | 300 | 100 | 1 |
| dyntoy_multifurcation | Dyntoy | Dyntoy-style (Poisson counts) | multifurcation | Multifurcation: 1 source → 4 branches (5 segments, 1 branch point) | 400 | 120 | 5 |
| dyntoy_tree | Dyntoy | Dyntoy-style (Poisson counts) | tree | Multifurcating tree (5 segments, multiple branch points) | 400 | 120 | 5 |
| **Splatter (3)** | | | | | | | |
| splatter_bifurcation | Splatter | Splatter-style (negative-binomial counts) | bifurcation | Single bifurcation: 1 trunk → 2 branches (3 segments, 1 branch point) | 300 | 100 | 3 |
| splatter_linear | Splatter | Splatter-style (negative-binomial counts) | linear | Linear path, no branch point (1 segment) | 300 | 100 | 1 |
| splatter_tree | Splatter | Splatter-style (negative-binomial counts) | tree | Multifurcating tree (5 segments, multiple branch points) | 400 | 120 | 5 |
| **Prosstt (3)** | | | | | | | |
| prosstt_bifurcation | Prosstt | PROSSTT-style (Gaussian expression) | bifurcation | Single bifurcation: 1 trunk → 2 branches (3 segments, 1 branch point) | 300 | 100 | 3 |
| prosstt_linear | Prosstt | PROSSTT-style (Gaussian expression) | linear | Linear path, no branch point (1 segment) | 300 | 100 | 1 |
| prosstt_tree | Prosstt | PROSSTT-style (Gaussian expression) | tree | Multifurcating tree (5 segments, multiple branch points) | 400 | 120 | 5 |
| **Dyngen (2)** | | | | | | | |
| dyngen_bifurcation | Dyngen | dyngen-style (Poisson, larger trees) | bifurcation | Single bifurcation: 1 trunk → 2 branches (3 segments, 1 branch point) | 500 | 150 | 3 |
| dyngen_tree | Dyngen | dyngen-style (Poisson, larger trees) | tree | Multifurcating tree (5 segments, multiple branch points) | 600 | 150 | 5 |

#### Summary

- 14 datasets across 4 simulator families (Dyntoy 6, Splatter 3, Prosstt 3, Dyngen 2) and 6 topology classes (bifurcation 4, convergence 1, cycle 1, linear 3, multifurcation 1, tree 4).
- Cell counts: 300–600 cells per dataset. Gene counts: 100–150 genes.
- Generative models: Poisson (Dyntoy), negative-binomial (Splatter), Gaussian (PROSSTT), Poisson with larger trees (dyngen) — chosen to span the noise-model × topology grid of Saelens et al. 2019.
- Gold standard: each dataset carries a known milestone/branch network (linear, single bifurcation, multifurcating tree, cycle, multifurcation, convergence), against which Topology (HIM distance), Branch-assignment, Cell-positions and Feature metrics are scored.
- Methods benchmarked (13): 6 R references (CytoTRACE, SCORPIUS, Slingshot, TSCAN, URD, destiny) + 7 Python ports (the six R-port equivalents + Monocle3-py).

### *Supplementary Table 28 | Trajectory benchmark — full method × dataset × metric matrix*

13 methods × 14 datasets × 4 metrics (Topology / Branch_assignment / Cell_positions / Features, each in [0,1]). Overall = unweighted mean.

#### Part A | Per-method summary (ranked) + R-vs-Python paired

| Rank | Method | Language | Overall | Topology | Branch_assignment | Cell_positions | Features | vs R counterpart (Δ overall) |
|---|---|---|---|---|---|---|---|---|
| 1 | Slingshot | Python | 0.485 | 0.286 | 0.559 | 0.631 | 0.464 | py 0.485 − R 0.455 = +0.030 (py≥R) |
| 2 | Slingshot | R | 0.455 | 0.286 | 0.493 | 0.594 | 0.448 | (see py row) |
| 3 | URD | Python | 0.439 | 0.23 | 0.436 | 0.437 | 0.654 | py 0.439 − R 0.348 = +0.092 (py≥R) |
| 4 | destiny | R | 0.431 | 0.272 | 0.455 | 0.563 | 0.433 | (see py row) |
| 5 | SCORPIUS | R | 0.417 | 0.285 | 0.476 | 0.584 | 0.324 | (see py row) |
| 6 | destiny | Python | 0.403 | 0.23 | 0.436 | 0.568 | 0.377 | py 0.403 − R 0.431 = -0.028 |
| 7 | TSCAN | R | 0.4 | 0.004 | 0.864 | 0.434 | 0.299 | (see py row) |
| 8 | SCORPIUS | Python | 0.398 | 0.285 | 0.476 | 0.565 | 0.266 | py 0.398 − R 0.417 = -0.019 |
| 9 | TSCAN | Python | 0.393 | 0.004 | 0.827 | 0.44 | 0.3 | py 0.393 − R 0.400 = -0.008 |
| 10 | URD | R | 0.348 | 0.23 | 0.436 | 0.514 | 0.211 | (see py row) |
| 11 | CytoTRACE | Python | 0.241 | 0.23 | 0.436 | 0.084 | 0.213 | py 0.241 − R 0.189 = +0.052 (py≥R) |
| 12 | Monocle3 | Python | 0.19 | 0.285 | 0.476 | 0.0 | 0.0 | (no R/py pair) |
| 13 | CytoTRACE | R | 0.189 | 0.166 | 0.389 | 0.101 | 0.1 | (see py row) |

#### Part B | Full method × dataset × metric matrix (182 rows)

| Method | Language | Dataset | Source | Topology class | Topology | Branch_assignment | Cell_positions | Features | Mean | Runtime_s |
|---|---|---|---|---|---|---|---|---|---|---|
| Slingshot (py) | Py | dyngen_bifurcation | Dyngen | bifurcation | 0.0007 | 0.3347 | 0.6135 | 0.0 | 0.2372 | 0.0525 |
| Slingshot (py) | Py | dyngen_tree | Dyngen | tree | 0.0011 | 0.4878 | 0.5251 | 0.375 | 0.3473 | 0.1154 |
| Slingshot (py) | Py | dyntoy_bifurcation | Dyntoy | bifurcation | 0.0007 | 0.3333 | 0.6294 | 0.0 | 0.2408 | 0.0803 |
| Slingshot (py) | Py | dyntoy_convergence | Dyntoy | convergence | 0.0007 | 0.3333 | 0.3995 | 0.7784 | 0.378 | 0.0438 |
| Slingshot (py) | Py | dyntoy_cycle | Dyntoy | cycle | 1.0 | 1.0 | 0.5586 | 0.7873 | 0.8365 | 0.6785 |
| Slingshot (py) | Py | dyntoy_linear | Dyntoy | linear | 1.0 | 1.0 | 0.9999 | 1.0 | 1.0 | 0.3189 |
| Slingshot (py) | Py | dyntoy_multifurcation | Dyntoy | multifurcation | -0.0017 | 0.2 | 0.3798 | 0.0 | 0.1445 | 0.0471 |
| Slingshot (py) | Py | dyntoy_tree | Dyntoy | tree | 0.0011 | 0.4878 | 0.488 | 0.6585 | 0.4089 | 0.0766 |
| Slingshot (py) | Py | prosstt_bifurcation | Prosstt | bifurcation | 0.0007 | 0.3333 | 0.6191 | 0.0 | 0.2383 | 0.028 |
| Slingshot (py) | Py | prosstt_linear | Prosstt | linear | 1.0 | 1.0 | 1.0 | 0.9959 | 0.999 | 0.7995 |
| Slingshot (py) | Py | prosstt_tree | Prosstt | tree | 0.0011 | 0.4878 | 0.488 | 0.7193 | 0.4241 | 0.0705 |

| | | | | | | | | | | |
|---|---|---|---|---|---|---|---|---|---|---|
| Slingshot (py) | Py | splatter_bifurcation | Splatter | bifurcation | 0.0007 | 0.3333 | 0.6313 | 0.0 | 0.2413 | 0.032 |
| Slingshot (py) | Py | splatter_linear | Splatter | linear | 1.0 | 1.0 | 0.9999 | 0.9967 | 0.9991 | 1.27 |
| Slingshot (py) | Py | splatter_tree | Splatter | tree | 0.0011 | 0.4878 | 0.5007 | 0.1911 | 0.2952 | 0.071 |
| Slingshot | R | dyngen_bifurcation | Dyngen | bifurcation | 0.0007 | 0.3347 | 0.5635 | 0.0 | 0.2247 | 6.7 |
| Slingshot | R | dyngen_tree | Dyngen | tree | -0.0017 | 0.2 | 0.3519 | 0.2068 | 0.1892 | 7.0839 |
| Slingshot | R | dyntoy_bifurcation | Dyntoy | bifurcation | 0.0007 | 0.3333 | 0.5998 | 0.0 | 0.2335 | 6.7237 |
| Slingshot | R | dyntoy_convergence | Dyntoy | convergence | 0.0007 | 0.3333 | 0.4507 | 0.9538 | 0.4346 | 7.5705 |
| Slingshot | R | dyntoy_cycle | Dyntoy | cycle | 1.0 | 1.0 | 0.5616 | 0.9033 | 0.8662 | 7.2281 |
| Slingshot | R | dyntoy_linear | Dyntoy | linear | 1.0 | 1.0 | 1.0 | 1.0 | 1.0 | 7.126 |
| Slingshot | R | dyntoy_multifurcation | Dyntoy | multifurcation | -0.0017 | 0.2 | 0.3814 | 0.0 | 0.1449 | 7.6226 |
| Slingshot | R | dyntoy_tree | Dyntoy | tree | -0.0017 | 0.2 | 0.3578 | 0.4812 | 0.2593 | 7.5043 |
| Slingshot | R | prosstt_bifurcation | Prosstt | bifurcation | 0.0007 | 0.3333 | 0.5897 | 0.0 | 0.2309 | 7.1819 |
| Slingshot | R | prosstt_linear | Prosstt | linear | 1.0 | 1.0 | 1.0 | 1.0 | 1.0 | 6.7254 |
| Slingshot | R | prosstt_tree | Prosstt | tree | -0.0017 | 0.2 | 0.3743 | 0.4838 | 0.2641 | 7.5549 |
| Slingshot | R | splatter_bifurcation | Splatter | bifurcation | 0.0007 | 0.3333 | 0.6003 | 0.0 | 0.2336 | 7.2653 |
| Slingshot | R | splatter_linear | Splatter | linear | 1.0 | 1.0 | 1.0 | 1.0 | 1.0 | 7.0023 |
| Slingshot | R | splatter_tree | Splatter | tree | 0.0011 | 0.4279 | 0.478 | 0.2484 | 0.2889 | 7.0812 |
| URD (py) | Py | dyngen_bifurcation | Dyngen | bifurcation | 0.0007 | 0.3347 | 0.3094 | 0.9491 | 0.3985 | 0.2295 |
| URD (py) | Py | dyngen_tree | Dyngen | tree | -0.0017 | 0.2 | 0.24 | 0.2629 | 0.1753 | 0.2015 |
| URD (py) | Py | dyntoy_bifurcation | Dyntoy | bifurcation | 0.0007 | 0.3333 | 0.3148 | 0.9621 | 0.4027 | 0.1428 |
| URD (py) | Py | dyntoy_convergence | Dyntoy | convergence | 0.0007 | 0.3333 | 0.3683 | 0.9309 | 0.4083 | 0.0632 |
| URD (py) | Py | dyntoy_cycle | Dyntoy | cycle | nan | nan | nan | nan | nan | 0.0629 |
| URD (py) | Py | dyntoy_linear | Dyntoy | linear | 1.0 | 1.0 | 0.9051 | 0.9374 | 0.9606 | 0.0645 |
| URD (py) | Py | dyntoy_multifurcation | Dyntoy | multifurcation | -0.0017 | 0.2 | 0.1122 | 0.7143 | 0.2562 | 0.2092 |
| URD (py) | Py | dyntoy_tree | Dyntoy | tree | -0.0017 | 0.2 | 0.2766 | 0.8217 | 0.3241 | 0.3548 |
| URD (py) | Py | prosstt_bifurcation | Prosstt | bifurcation | 0.0007 | 0.3333 | 0.4054 | 0.0 | 0.1849 | 0.0814 |
| URD (py) | Py | prosstt_linear | Prosstt | linear | 1.0 | 1.0 | 0.9201 | 0.4324 | 0.8381 | 0.2411 |
| URD (py) | Py | prosstt_tree | Prosstt | tree | -0.0017 | 0.2 | 0.3178 | 0.7713 | 0.3218 | 0.1509 |
| URD (py) | Py | splatter_bifurcation | Splatter | bifurcation | 0.0007 | 0.3333 | 0.3786 | 0.9362 | 0.4122 | 0.0796 |
| URD (py) | Py | splatter_linear | Splatter | linear | 1.0 | 1.0 | 0.847 | 0.7859 | 0.9082 | 0.159 |
| URD (py) | Py | splatter_tree | Splatter | tree | -0.0017 | 0.2 | 0.2898 | 0.0 | 0.122 | 0.2925 |
| destiny | R | dyngen_bifurcation | Dyngen | bifurcation | 0.0007 | 0.3347 | 0.4824 | 0.739 | 0.3892 | 9.4078 |
| destiny | R | dyngen_tree | Dyngen | tree | -0.0017 | 0.2 | 0.3521 | 0.0713 | 0.1554 | 9.7993 |
| destiny | R | dyntoy_bifurcation | Dyntoy | bifurcation | 0.0007 | 0.3333 | 0.5493 | 0.0 | 0.2208 | 9.176 |
| destiny | R | dyntoy_convergence | Dyntoy | convergence | nan | nan | nan | nan | nan | 9.0534 |
| destiny | R | dyntoy_cycle | Dyntoy | cycle | nan | nan | nan | nan | nan | 9.302 |
| destiny | R | dyntoy_linear | Dyntoy | linear | 1.0 | 1.0 | 0.9291 | 0.9443 | 0.9683 | 9.6196 |
| destiny | R | dyntoy_multifurcation | Dyntoy | multifurcation | -0.0017 | 0.2 | 0.4174 | 0.0 | 0.1539 | 18.2078 |
| destiny | R | dyntoy_tree | Dyntoy | tree | -0.0017 | 0.2 | 0.3737 | 0.5503 | 0.2806 | 11.4507 |
| destiny | R | prosstt_bifurcation | Prosstt | bifurcation | nan | nan | nan | nan | nan | 9.2494 |
| destiny | R | prosstt_linear | Prosstt | linear | 1.0 | 1.0 | 0.9528 | 0.9252 | 0.9695 | 9.6136 |
| destiny | R | prosstt_tree | Prosstt | tree | -0.0017 | 0.2 | 0.3134 | 0.7175 | 0.3073 | 12.1819 |
| destiny | R | splatter_bifurcation | Splatter | bifurcation | 0.0007 | 0.3333 | 0.5799 | 0.0 | 0.2285 | 9.9383 |
| destiny | R | splatter_linear | Splatter | linear | 1.0 | 1.0 | 0.9144 | 0.8125 | 0.9317 | 9.1599 |
| destiny | R | splatter_tree | Splatter | tree | -0.0017 | 0.2 | 0.3337 | 0.0 | 0.133 | 10.0307 |
| SCORPIUS | R | dyngen_bifurcation | Dyngen | bifurcation | 0.0007 | 0.3347 | 0.5781 | 0.0 | 0.2284 | 4.1781 |
| SCORPIUS | R | dyngen_tree | Dyngen | tree | -0.0017 | 0.2 | 0.4245 | 0.0 | 0.1557 | 5.9038 |
| SCORPIUS | R | dyntoy_bifurcation | Dyntoy | bifurcation | 0.0007 | 0.3333 | 0.5753 | 0.0 | 0.2273 | 4.0887 |
| SCORPIUS | R | dyntoy_convergence | Dyntoy | convergence | 0.0007 | 0.3333 | 0.5809 | 0.9341 | 0.4622 | 4.399 |
| SCORPIUS | R | dyntoy_cycle | Dyntoy | cycle | 1.0 | 1.0 | 0.5564 | 0.9708 | 0.8818 | 4.3276 |
| SCORPIUS | R | dyntoy_linear | Dyntoy | linear | 1.0 | 1.0 | 0.8863 | 0.9444 | 0.9577 | 6.1878 |
| SCORPIUS | R | dyntoy_multifurcation | Dyntoy | multifurcation | -0.0017 | 0.2 | 0.441 | 0.0 | 0.1598 | 4.4855 |
| SCORPIUS | R | dyntoy_tree | Dyntoy | tree | -0.0017 | 0.2 | 0.3975 | 0.0234 | 0.1548 | 4.8998 |
| SCORPIUS | R | prosstt_bifurcation | Prosstt | bifurcation | 0.0007 | 0.3333 | 0.6271 | 0.0 | 0.2403 | 8.2323 |
| SCORPIUS | R | prosstt_linear | Prosstt | linear | 1.0 | 1.0 | 0.9333 | 0.8096 | 0.9357 | 7.2241 |
| SCORPIUS | R | prosstt_tree | Prosstt | tree | -0.0017 | 0.2 | 0.4464 | 0.0093 | 0.1635 | 4.0562 |
| SCORPIUS | R | splatter_bifurcation | Splatter | bifurcation | 0.0007 | 0.3333 | 0.5611 | 0.0 | 0.2238 | 4.3701 |
| SCORPIUS | R | splatter_linear | Splatter | linear | 1.0 | 1.0 | 0.7931 | 0.842 | 0.9088 | 4.3457 |
| SCORPIUS | R | splatter_tree | Splatter | tree | -0.0017 | 0.2 | 0.3757 | 0.0 | 0.1435 | 4.0558 |
| destiny (py) | Py | dyngen_bifurcation | Dyngen | bifurcation | 0.0007 | 0.3347 | 0.5575 | 0.7457 | 0.4096 | 0.0627 |
| destiny (py) | Py | dyngen_tree | Dyngen | tree | -0.0017 | 0.2 | 0.3823 | 0.0715 | 0.163 | 0.0965 |
| destiny (py) | Py | dyntoy_bifurcation | Dyntoy | bifurcation | 0.0007 | 0.3333 | 0.5543 | 0.0 | 0.2221 | 0.0522 |
| destiny (py) | Py | dyntoy_convergence | Dyntoy | convergence | 0.0007 | 0.3333 | 0.4287 | 0.9352 | 0.4245 | 0.0741 |
| destiny (py) | Py | dyntoy_cycle | Dyntoy | cycle | nan | nan | nan | nan | nan | 0.0551 |

| | | | | | | | | | | |
|---|---|---|---|---|---|---|---|---|---|---|
| destiny (py) | Py | dyntoy_linear | Dyntoy | linear | 1.0 | 1.0 | 0.9423 | 0.8813 | 0.9559 | 0.0982 |
| destiny (py) | Py | dyntoy_multifurcation | Dyntoy | multifurcation | -0.0017 | 0.2 | 0.417 | 0.0 | 0.1538 | 0.022 |
| destiny (py) | Py | dyntoy_tree | Dyntoy | tree | -0.0017 | 0.2 | 0.3637 | 0.525 | 0.2717 | 0.018 |
| destiny (py) | Py | prosstt_bifurcation | Prosstt | bifurcation | 0.0007 | 0.3333 | 0.5136 | 0.0 | 0.2119 | 0.0099 |
| destiny (py) | Py | prosstt_linear | Prosstt | linear | 1.0 | 1.0 | 0.9572 | 0.7606 | 0.9294 | 0.2432 |
| destiny (py) | Py | prosstt_tree | Prosstt | tree | -0.0017 | 0.2 | 0.4381 | 0.0658 | 0.1755 | 0.1563 |
| destiny (py) | Py | splatter_bifurcation | Splatter | bifurcation | 0.0007 | 0.3333 | 0.5856 | 0.0 | 0.2299 | 0.1126 |
| destiny (py) | Py | splatter_linear | Splatter | linear | 1.0 | 1.0 | 0.8772 | 0.9156 | 0.9482 | 0.0137 |
| destiny (py) | Py | splatter_tree | Splatter | tree | -0.0017 | 0.2 | 0.3604 | 0.0 | 0.1397 | 0.036 |
| TSCAN | R | dyngen_bifurcation | Dyngen | bifurcation | 0.005 | 1.0 | 0.5233 | 0.0 | 0.3821 | 12.1314 |
| TSCAN | R | dyngen_tree | Dyngen | tree | 0.005 | 0.9967 | 0.2979 | 0.1486 | 0.362 | 20.2278 |
| TSCAN | R | dyntoy_bifurcation | Dyntoy | bifurcation | 0.005 | 1.0 | 0.5013 | 0.0 | 0.3766 | 7.4434 |
| TSCAN | R | dyntoy_convergence | Dyntoy | convergence | 0.0039 | 0.7059 | 0.4228 | 0.9351 | 0.5169 | 8.4568 |
| TSCAN | R | dyntoy_cycle | Dyntoy | cycle | nan | nan | nan | nan | nan | 7.0815 |
| TSCAN | R | dyntoy_linear | Dyntoy | linear | 0.0027 | 0.5946 | 0.6411 | 0.9676 | 0.5515 | 7.9696 |
| TSCAN | R | dyntoy_multifurcation | Dyntoy | multifurcation | 0.0047 | 0.8331 | 0.2819 | 0.0 | 0.2799 | 8.8685 |
| TSCAN | R | dyntoy_tree | Dyntoy | tree | 0.005 | 1.0 | 0.2927 | 0.0 | 0.3244 | 9.0318 |
| TSCAN | R | prosstt_bifurcation | Prosstt | bifurcation | 0.005 | 0.9934 | 0.5079 | 0.0 | 0.3766 | 7.8046 |
| TSCAN | R | prosstt_linear | Prosstt | linear | 0.0027 | 0.5522 | 0.6568 | 0.5834 | 0.4488 | 7.5321 |
| TSCAN | R | prosstt_tree | Prosstt | tree | 0.005 | 0.995 | 0.2474 | 0.32 | 0.3919 | 8.6602 |
| TSCAN | R | splatter_bifurcation | Splatter | bifurcation | 0.005 | 0.9934 | 0.5043 | 0.0 | 0.3757 | 7.971 |
| TSCAN | R | splatter_linear | Splatter | linear | 0.0027 | 0.5856 | 0.5776 | 0.9306 | 0.5241 | 7.7905 |
| TSCAN | R | splatter_tree | Splatter | tree | 0.005 | 0.9852 | 0.1896 | 0.0 | 0.2949 | 15.3088 |
| SCORPIUS (py) | Py | dyngen_bifurcation | Dyngen | bifurcation | 0.0007 | 0.3347 | 0.5865 | 0.0 | 0.2305 | 0.8829 |
| SCORPIUS (py) | Py | dyngen_tree | Dyngen | tree | -0.0017 | 0.2 | 0.3627 | 0.0453 | 0.1516 | 0.5627 |
| SCORPIUS (py) | Py | dyntoy_bifurcation | Dyntoy | bifurcation | 0.0007 | 0.3333 | 0.5872 | 0.0 | 0.2303 | 0.48 |
| SCORPIUS (py) | Py | dyntoy_convergence | Dyntoy | convergence | 0.0007 | 0.3333 | 0.5794 | 0.9384 | 0.463 | 1.0793 |
| SCORPIUS (py) | Py | dyntoy_cycle | Dyntoy | cycle | 1.0 | 1.0 | 0.5959 | 0.2345 | 0.7076 | 0.9846 |
| SCORPIUS (py) | Py | dyntoy_linear | Dyntoy | linear | 1.0 | 1.0 | 0.8922 | 0.841 | 0.9333 | 0.6052 |
| SCORPIUS (py) | Py | dyntoy_multifurcation | Dyntoy | multifurcation | -0.0017 | 0.2 | 0.4124 | 0.0 | 0.1527 | 1.014 |
| SCORPIUS (py) | Py | dyntoy_tree | Dyntoy | tree | -0.0017 | 0.2 | 0.3423 | 0.0 | 0.1352 | 1.0642 |
| SCORPIUS (py) | Py | prosstt_bifurcation | Prosstt | bifurcation | 0.0007 | 0.3333 | 0.5889 | 0.0 | 0.2307 | 0.8526 |
| SCORPIUS (py) | Py | prosstt_linear | Prosstt | linear | 1.0 | 1.0 | 0.8934 | 0.7143 | 0.9019 | 0.7182 |
| SCORPIUS (py) | Py | prosstt_tree | Prosstt | tree | -0.0017 | 0.2 | 0.3756 | 0.6755 | 0.3123 | 0.7903 |
| SCORPIUS (py) | Py | splatter_bifurcation | Splatter | bifurcation | 0.0007 | 0.3333 | 0.5782 | 0.0 | 0.2281 | 1.9734 |
| SCORPIUS (py) | Py | splatter_linear | Splatter | linear | 1.0 | 1.0 | 0.7746 | 0.2733 | 0.762 | 2.1835 |
| SCORPIUS (py) | Py | splatter_tree | Splatter | tree | -0.0017 | 0.2 | 0.3391 | 0.0 | 0.1343 | 0.9099 |
| TSCAN (py) | Py | dyngen_bifurcation | Dyngen | bifurcation | 0.005 | 1.0 | 0.5354 | 0.0 | 0.3851 | 0.3363 |
| TSCAN (py) | Py | dyngen_tree | Dyngen | tree | 0.005 | 0.9901 | 0.3106 | 0.1047 | 0.3526 | 0.4164 |
| TSCAN (py) | Py | dyntoy_bifurcation | Dyntoy | bifurcation | 0.005 | 1.0 | 0.4992 | 0.0 | 0.376 | 0.4567 |
| TSCAN (py) | Py | dyntoy_convergence | Dyntoy | convergence | 0.005 | 0.8302 | 0.2836 | 0.9273 | 0.5115 | 0.1918 |
| TSCAN (py) | Py | dyntoy_cycle | Dyntoy | cycle | -0.0017 | 0.2462 | 0.367 | 0.0494 | 0.1652 | 0.1592 |
| TSCAN (py) | Py | dyntoy_linear | Dyntoy | linear | 0.0027 | 0.5161 | 0.673 | 0.9917 | 0.5459 | 5.1191 |
| TSCAN (py) | Py | dyntoy_multifurcation | Dyntoy | multifurcation | 0.005 | 0.9901 | 0.2862 | 0.0 | 0.3203 | 0.224 |
| TSCAN (py) | Py | dyntoy_tree | Dyntoy | tree | 0.005 | 0.971 | 0.2737 | 0.0 | 0.3124 | 0.204 |
| TSCAN (py) | Py | prosstt_bifurcation | Prosstt | bifurcation | 0.005 | 0.9934 | 0.4623 | 0.0 | 0.3651 | 0.127 |
| TSCAN (py) | Py | prosstt_linear | Prosstt | linear | 0.0027 | 0.5522 | 0.7458 | 0.6919 | 0.4982 | 0.2245 |
| TSCAN (py) | Py | prosstt_tree | Prosstt | tree | 0.005 | 0.995 | 0.3189 | 0.5441 | 0.4657 | 0.1958 |
| TSCAN (py) | Py | splatter_bifurcation | Splatter | bifurcation | 0.005 | 0.9868 | 0.5254 | 0.0 | 0.3793 | 0.1842 |
| TSCAN (py) | Py | splatter_linear | Splatter | linear | 0.0027 | 0.537 | 0.7341 | 0.8899 | 0.5409 | 0.285 |
| TSCAN (py) | Py | splatter_tree | Splatter | tree | 0.005 | 0.966 | 0.1456 | 0.0 | 0.2791 | 0.2387 |
| URD | R | dyngen_bifurcation | Dyngen | bifurcation | 0.0007 | 0.3347 | 0.556 | 0.0 | 0.2228 | 11.1879 |
| URD | R | dyngen_tree | Dyngen | tree | -0.0017 | 0.2 | 0.3525 | 0.0 | 0.1377 | 13.0146 |
| URD | R | dyntoy_bifurcation | Dyntoy | bifurcation | 0.0007 | 0.3333 | 0.5605 | 0.0 | 0.2236 | 10.6559 |
| URD | R | dyntoy_convergence | Dyntoy | convergence | 0.0007 | 0.3333 | 0.4758 | 0.9356 | 0.4363 | 10.0232 |
| URD | R | dyntoy_cycle | Dyntoy | cycle | nan | nan | nan | nan | nan | 9.0568 |
| URD | R | dyntoy_linear | Dyntoy | linear | 1.0 | 1.0 | 0.854 | 0.0 | 0.7135 | 9.8563 |
| URD | R | dyntoy_multifurcation | Dyntoy | multifurcation | -0.0017 | 0.2 | 0.3496 | 0.0 | 0.137 | 13.4923 |
| URD | R | dyntoy_tree | Dyntoy | tree | -0.0017 | 0.2 | 0.3641 | 0.1268 | 0.1723 | 14.8646 |
| URD | R | prosstt_bifurcation | Prosstt | bifurcation | 0.0007 | 0.3333 | 0.2177 | 0.0 | 0.1379 | 9.6859 |
| URD | R | prosstt_linear | Prosstt | linear | 1.0 | 1.0 | 0.915 | 0.4917 | 0.8517 | 13.2975 |
| URD | R | prosstt_tree | Prosstt | tree | -0.0017 | 0.2 | 0.321 | 0.3736 | 0.2232 | 10.0919 |
| URD | R | splatter_bifurcation | Splatter | bifurcation | 0.0007 | 0.3333 | 0.5724 | 0.0 | 0.2266 | 9.6881 |
| URD | R | splatter_linear | Splatter | linear | 1.0 | 1.0 | 0.7889 | 0.8112 | 0.9 | 12.1561 |
| URD | R | splatter_tree | Splatter | tree | -0.0017 | 0.2 | 0.3487 | 0.0 | 0.1367 | 12.247 |

| | | | | | | | | | | |
|---|---|---|---|---|---|---|---|---|---|---|
| CytoTRACE (py) | Py | dyngen_bifurcation | Dyngen | bifurcation | 0.0007 | 0.3347 | 0.0913 | 0.0 | 0.1067 | 0.2501 |
| CytoTRACE (py) | Py | dyngen_tree | Dyngen | tree | -0.0017 | 0.2 | 0.0286 | 0.0 | 0.0567 | 0.4288 |
| CytoTRACE (py) | Py | dyntoy_bifurcation | Dyntoy | bifurcation | 0.0007 | 0.3333 | 0.0395 | 0.0 | 0.0934 | 0.5278 |
| CytoTRACE (py) | Py | dyntoy_convergence | Dyntoy | convergence | 0.0007 | 0.3333 | 0.0854 | 0.7762 | 0.2989 | 0.1747 |
| CytoTRACE (py) | Py | dyntoy_cycle | Dyntoy | cycle | nan | nan | nan | nan | nan | 0.0146 |
| CytoTRACE (py) | Py | dyntoy_linear | Dyntoy | linear | 1.0 | 1.0 | 0.0968 | 0.6758 | 0.6932 | 0.0712 |
| CytoTRACE (py) | Py | dyntoy_multifurcation | Dyntoy | multifurcation | -0.0017 | 0.2 | 0.0073 | 0.3865 | 0.148 | 0.5821 |
| CytoTRACE (py) | Py | dyntoy_tree | Dyntoy | tree | -0.0017 | 0.2 | 0.0179 | 0.1569 | 0.0933 | 0.9417 |
| CytoTRACE (py) | Py | prosstt_bifurcation | Prosstt | bifurcation | 0.0007 | 0.3333 | 0.1108 | 0.0 | 0.1112 | 0.1654 |
| CytoTRACE (py) | Py | prosstt_linear | Prosstt | linear | 1.0 | 1.0 | 0.2818 | 0.3761 | 0.6645 | 0.1064 |
| CytoTRACE (py) | Py | prosstt_tree | Prosstt | tree | -0.0017 | 0.2 | 0.176 | 0.392 | 0.1916 | 0.2108 |
| CytoTRACE (py) | Py | splatter_bifurcation | Splatter | bifurcation | 0.0007 | 0.3333 | 0.0032 | 0.0 | 0.0843 | 0.1415 |
| CytoTRACE (py) | Py | splatter_linear | Splatter | linear | 1.0 | 1.0 | 0.075 | 0.0 | 0.5188 | 0.1519 |
| CytoTRACE (py) | Py | splatter_tree | Splatter | tree | -0.0017 | 0.2 | 0.0801 | 0.0 | 0.0696 | 0.19 |
| Monocle3 (py) | Py | dyngen_bifurcation | Dyngen | bifurcation | 0.0007 | 0.3347 | 0.0 | 0.0 | 0.0838 | 44.0741 |
| Monocle3 (py) | Py | dyngen_tree | Dyngen | tree | -0.0017 | 0.2 | 0.0 | 0.0 | 0.0496 | 20.1668 |
| Monocle3 (py) | Py | dyntoy_bifurcation | Dyntoy | bifurcation | 0.0007 | 0.3333 | 0.0 | 0.0 | 0.0835 | 54.9096 |
| Monocle3 (py) | Py | dyntoy_convergence | Dyntoy | convergence | 0.0007 | 0.3333 | 0.0 | 0.0 | 0.0835 | 46.7716 |
| Monocle3 (py) | Py | dyntoy_cycle | Dyntoy | cycle | 1.0 | 1.0 | 0.0 | 0.0 | 0.5 | 39.1786 |
| Monocle3 (py) | Py | dyntoy_linear | Dyntoy | linear | 1.0 | 1.0 | 0.0 | 0.0 | 0.5 | 39.7495 |
| Monocle3 (py) | Py | dyntoy_multifurcation | Dyntoy | multifurcation | -0.0017 | 0.2 | 0.0 | 0.0 | 0.0496 | 103.0708 |
| Monocle3 (py) | Py | dyntoy_tree | Dyntoy | tree | -0.0017 | 0.2 | 0.0 | 0.0 | 0.0496 | 108.2128 |
| Monocle3 (py) | Py | prosstt_bifurcation | Prosstt | bifurcation | 0.0007 | 0.3333 | 0.0 | 0.0 | 0.0835 | 56.0388 |
| Monocle3 (py) | Py | prosstt_linear | Prosstt | linear | 1.0 | 1.0 | 0.0 | 0.0 | 0.5 | 13.2691 |
| Monocle3 (py) | Py | prosstt_tree | Prosstt | tree | -0.0017 | 0.2 | 0.0 | 0.0 | 0.0496 | 31.481 |
| Monocle3 (py) | Py | splatter_bifurcation | Splatter | bifurcation | 0.0007 | 0.3333 | 0.0 | 0.0 | 0.0835 | 60.9895 |
| Monocle3 (py) | Py | splatter_linear | Splatter | linear | 1.0 | 1.0 | 0.0 | 0.0 | 0.5 | 44.4199 |
| Monocle3 (py) | Py | splatter_tree | Splatter | tree | -0.0017 | 0.2 | 0.0 | 0.0 | 0.0496 | 18.0316 |
| CytoTRACE | R | dyngen_bifurcation | Dyngen | bifurcation | 0.0007 | 0.3347 | 0.0737 | 0.0 | 0.1023 | 4.0007 |
| CytoTRACE | R | dyngen_tree | Dyngen | tree | -0.0017 | 0.2 | 0.0515 | 0.0 | 0.0624 | 4.3486 |
| CytoTRACE | R | dyntoy_bifurcation | Dyntoy | bifurcation | 0.0007 | 0.3333 | 0.0436 | 0.0 | 0.0944 | 4.3009 |
| CytoTRACE | R | dyntoy_convergence | Dyntoy | convergence | 0.0007 | 0.3333 | 0.0466 | 0.0 | 0.0952 | 4.8175 |
| CytoTRACE | R | dyntoy_cycle | Dyntoy | cycle | nan | nan | nan | nan | nan | 4.3849 |
| CytoTRACE | R | dyntoy_linear | Dyntoy | linear | 1.0 | 1.0 | 0.0879 | 0.286 | 0.5935 | 4.417 |
| CytoTRACE | R | dyntoy_multifurcation | Dyntoy | multifurcation | -0.0017 | 0.2 | 0.039 | 0.0 | 0.0593 | 4.1397 |
| CytoTRACE | R | dyntoy_tree | Dyntoy | tree | -0.0017 | 0.2 | 0.0527 | 0.3453 | 0.1491 | 4.67 |
| CytoTRACE | R | prosstt_bifurcation | Prosstt | bifurcation | 0.0007 | 0.3333 | 0.4016 | 0.0 | 0.1839 | 7.0564 |
| CytoTRACE | R | prosstt_linear | Prosstt | linear | nan | nan | nan | nan | nan | 4.3046 |
| CytoTRACE | R | prosstt_tree | Prosstt | tree | -0.0017 | 0.2 | 0.1607 | 0.2019 | 0.1402 | 5.2977 |
| CytoTRACE | R | splatter_bifurcation | Splatter | bifurcation | 0.0007 | 0.3333 | 0.1307 | 0.3634 | 0.207 | 5.1299 |
| CytoTRACE | R | splatter_linear | Splatter | linear | 1.0 | 1.0 | 0.0338 | 0.0 | 0.5084 | 9.9969 |
| CytoTRACE | R | splatter_tree | Splatter | tree | -0.0017 | 0.2 | 0.0904 | 0.0 | 0.0722 | 4.038 |

#### Summary

- 13 methods × 14 datasets × 4 metrics = 182 scored (method×dataset) rows.
- Python ≥ R overall in 3/6 paired comparisons: py-Slingshot (0.485 vs 0.455), py-URD (0.439 vs 0.348), py-CytoTRACE (0.241 vs 0.189); R ahead for destiny (0.431 vs 0.403), SCORPIUS (0.417 vs 0.398), TSCAN (0.400 vs 0.393). Monocle3 = Python port only (0.190).
- Topology = Hamming–Ipsen–Mikhailov distance between gold and predicted milestone graphs; Branch_assignment / Cell_positions / Features per the pydyneval suite; all in [0,1], higher = better; Overall = unweighted mean.
- Dataset source/topology/dimensions are detailed in Supplementary Table 4_7.

### *Supplementary Table 29 | Trajectory benchmark — runtime & memory*

Per-method runtime, R-vs-Python speed-up (median 22.6×), re-measured Python replicate variance (warm-up + 3 reps) and tracemalloc peak memory, by method × dataset × language.

#### Part A | Per-method runtime, speed-up, variance, memory

| Method | R_runtime_s_n1 | Py_runtime_s_sweep_n1 | Speedup_R_over_Py | Py_remeas_mean_s | Py_remeas_SD_s | Py_rep_CV_pct | Py_peak_mem_MB |
|---|---|---|---|---|---|---|---|
| destiny | 10.442 | 0.075 | 139.2× | 0.013 | 0.0 | 1.0 | 9.8 |
| URD | 11.38 | 0.167 | 68.3× | 0.083 | 0.001 | 0.6 | 9.8 |
| Slingshot | 7.169 | 0.263 | 27.2× | 0.097 | 0.002 | 2.1 | 2.3 |
| CytoTRACE | 5.064 | 0.283 | 17.9× | 0.116 | 0.005 | 4.2 | 5.3 |
| TSCAN | 9.734 | 0.597 | 16.3× | 0.362 | 0.042 | 11.7 | 38.2 |
| SCORPIUS | 5.054 | 1.007 | 5.0× | 0.898 | 0.394 | 43.9 | 5.7 |
| Monocle3 | — | 48.597 | — | 3.408 | 0.377 | 11.1 | 265.5 |

#### Part B | Full matrix (method × dataset × language)

| Method | Language | Dataset | Cells | Genes | Runtime_s | Rep_SD_s | Peak_mem_MB | Note |
|---|---|---|---|---|---|---|---|---|
| destiny | R | dyngen_bifurcation | 500 | 150 | 9.408 | — | — | |
| destiny | R | dyngen_tree | 600 | 150 | 9.799 | — | — | |
| destiny | R | dyntoy_bifurcation | 300 | 100 | 9.176 | — | — | |
| destiny | R | dyntoy_convergence | 300 | 100 | 9.053 | — | — | ERR |
| destiny | R | dyntoy_cycle | 300 | 100 | 9.302 | — | — | ERR |
| destiny | R | dyntoy_linear | 300 | 100 | 9.62 | — | — | |
| destiny | R | dyntoy_multifurcation | 400 | 120 | 18.208 | — | — | |
| destiny | R | dyntoy_tree | 400 | 120 | 11.451 | — | — | |
| destiny | R | prosstt_bifurcation | 300 | 100 | 9.249 | — | — | ERR |
| destiny | R | prosstt_linear | 300 | 100 | 9.614 | — | — | |
| destiny | R | prosstt_tree | 400 | 120 | 12.182 | — | — | |
| destiny | R | splatter_bifurcation | 300 | 100 | 9.938 | — | — | |
| destiny | R | splatter_linear | 300 | 100 | 9.16 | — | — | |
| destiny | R | splatter_tree | 400 | 120 | 10.031 | — | — | |
| destiny | Py | dyngen_bifurcation | 500 | 150 | 0.0279 | 0.0002 | 16.94 | n=3 |
| destiny | Py | dyngen_tree | 600 | 150 | 0.0371 | 0.0003 | 24.21 | n=3 |
| destiny | Py | dyntoy_bifurcation | 300 | 100 | 0.0074 | 0.0001 | 6.17 | n=3 |
| destiny | Py | dyntoy_convergence | 300 | 100 | 0.0077 | 0.0001 | 6.17 | n=3 |
| destiny | Py | dyntoy_cycle | 300 | 100 | — | — | — | ERR: ArpackError: ARPACK error -9999: Could not build an Arnoldi factorization. IPARA |
| destiny | Py | dyntoy_linear | 300 | 100 | 0.0075 | 0.0001 | 6.17 | n=3 |
| destiny | Py | dyntoy_multifurcation | 400 | 120 | 0.0142 | 0.0 | 10.87 | n=3 |
| destiny | Py | dyntoy_tree | 400 | 120 | 0.014 | 0.0001 | 10.87 | n=3 |
| destiny | Py | prosstt_bifurcation | 300 | 100 | 0.0072 | 0.0001 | 6.17 | n=3 |
| destiny | Py | prosstt_linear | 300 | 100 | 0.0075 | 0.0001 | 6.17 | n=3 |
| destiny | Py | prosstt_tree | 400 | 120 | 0.0139 | 0.0001 | 10.87 | n=3 |
| destiny | Py | splatter_bifurcation | 300 | 100 | 0.0076 | 0.0001 | 6.17 | n=3 |
| destiny | Py | splatter_linear | 300 | 100 | 0.0076 | 0.0001 | 6.17 | n=3 |
| destiny | Py | splatter_tree | 400 | 120 | 0.0136 | 0.0001 | 10.87 | n=3 |
| URD | R | dyngen_bifurcation | 500 | 150 | 11.188 | — | — | |
| URD | R | dyngen_tree | 600 | 150 | 13.015 | — | — | |
| URD | R | dyntoy_bifurcation | 300 | 100 | 10.656 | — | — | |
| URD | R | dyntoy_convergence | 300 | 100 | 10.023 | — | — | |
| URD | R | dyntoy_cycle | 300 | 100 | 9.057 | — | — | ERR |
| URD | R | dyntoy_linear | 300 | 100 | 9.856 | — | — | |
| URD | R | dyntoy_multifurcation | 400 | 120 | 13.492 | — | — | |
| URD | R | dyntoy_tree | 400 | 120 | 14.865 | — | — | |
| URD | R | prosstt_bifurcation | 300 | 100 | 9.686 | — | — | |
| URD | R | prosstt_linear | 300 | 100 | 13.298 | — | — | |
| URD | R | prosstt_tree | 400 | 120 | 10.092 | — | — | |
| URD | R | splatter_bifurcation | 300 | 100 | 9.688 | — | — | |
| URD | R | splatter_linear | 300 | 100 | 12.156 | — | — | |
| URD | R | splatter_tree | 400 | 120 | 12.247 | — | — | |
| URD | Py | dyngen_bifurcation | 500 | 150 | 0.1281 | 0.0003 | 16.94 | n=3 |
| URD | Py | dyngen_tree | 600 | 150 | 0.1823 | 0.0004 | 24.21 | n=3 |
| URD | Py | dyntoy_bifurcation | 300 | 100 | 0.0582 | 0.0002 | 6.17 | n=3 |

| | | | | | | | | |
|---|---|---|---|---|---|---|---|---|
| URD | Py | dyntoy_convergence | 300 | 100 | 0.0505 | 0.0017 | 6.17 | n=3 |
| URD | Py | dyntoy_cycle | 300 | 100 | — | — | — | ERR: ArpackError: ARPACK error -9999: Could not build an Arnoldi factorization. IPARA |
| URD | Py | dyntoy_linear | 300 | 100 | 0.0518 | 0.0 | 6.17 | n=3 |
| URD | Py | dyntoy_multifurcation | 400 | 120 | 0.0701 | 0.0002 | 10.87 | n=3 |
| URD | Py | dyntoy_tree | 400 | 120 | 0.0934 | 0.0 | 10.87 | n=3 |
| URD | Py | prosstt_bifurcation | 300 | 100 | 0.0663 | 0.0015 | 6.17 | n=3 |
| URD | Py | prosstt_linear | 300 | 100 | 0.0596 | 0.0 | 6.17 | n=3 |
| URD | Py | prosstt_tree | 400 | 120 | 0.1225 | 0.0001 | 10.87 | n=3 |
| URD | Py | splatter_bifurcation | 300 | 100 | 0.0494 | 0.0002 | 6.17 | n=3 |
| URD | Py | splatter_linear | 300 | 100 | 0.0475 | 0.0002 | 6.17 | n=3 |
| URD | Py | splatter_tree | 400 | 120 | 0.0988 | 0.0004 | 10.87 | n=3 |
| Slingshot | R | dyngen_bifurcation | 500 | 150 | 6.7 | — | — | |
| Slingshot | R | dyngen_tree | 600 | 150 | 7.084 | — | — | |
| Slingshot | R | dyntoy_bifurcation | 300 | 100 | 6.724 | — | — | |
| Slingshot | R | dyntoy_convergence | 300 | 100 | 7.571 | — | — | |
| Slingshot | R | dyntoy_cycle | 300 | 100 | 7.228 | — | — | |
| Slingshot | R | dyntoy_linear | 300 | 100 | 7.126 | — | — | |
| Slingshot | R | dyntoy_multifurcation | 400 | 120 | 7.623 | — | — | |
| Slingshot | R | dyntoy_tree | 400 | 120 | 7.504 | — | — | |
| Slingshot | R | prosstt_bifurcation | 300 | 100 | 7.182 | — | — | |
| Slingshot | R | prosstt_linear | 300 | 100 | 6.725 | — | — | |
| Slingshot | R | prosstt_tree | 400 | 120 | 7.555 | — | — | |
| Slingshot | R | splatter_bifurcation | 300 | 100 | 7.265 | — | — | |
| Slingshot | R | splatter_linear | 300 | 100 | 7.002 | — | — | |
| Slingshot | R | splatter_tree | 400 | 120 | 7.081 | — | — | |
| Slingshot | Py | dyngen_bifurcation | 500 | 150 | 0.0438 | 0.0001 | 4.1 | n=3 |
| Slingshot | Py | dyngen_tree | 600 | 150 | 0.1023 | 0.0001 | 5.9 | n=3 |
| Slingshot | Py | dyntoy_bifurcation | 300 | 100 | 0.025 | 0.0002 | 1.51 | n=3 |
| Slingshot | Py | dyntoy_convergence | 300 | 100 | 0.0239 | 0.0001 | 1.51 | n=3 |
| Slingshot | Py | dyntoy_cycle | 300 | 100 | 0.2375 | 0.0184 | 1.54 | n=3 |
| Slingshot | Py | dyntoy_linear | 300 | 100 | 0.2305 | 0.0113 | 1.54 | n=3 |
| Slingshot | Py | dyntoy_multifurcation | 400 | 120 | 0.0371 | 0.0003 | 2.64 | n=3 |
| Slingshot | Py | dyntoy_tree | 400 | 120 | 0.0646 | 0.0005 | 2.66 | n=3 |
| Slingshot | Py | prosstt_bifurcation | 300 | 100 | 0.0244 | 0.0 | 1.51 | n=3 |
| Slingshot | Py | prosstt_linear | 300 | 100 | 0.233 | 0.0137 | 1.53 | n=3 |
| Slingshot | Py | prosstt_tree | 400 | 120 | 0.062 | 0.0002 | 2.66 | n=3 |
| Slingshot | Py | splatter_bifurcation | 300 | 100 | 0.0251 | 0.0001 | 1.51 | n=3 |
| Slingshot | Py | splatter_linear | 300 | 100 | 0.1815 | 0.0113 | 1.53 | n=3 |
| Slingshot | Py | splatter_tree | 400 | 120 | 0.0629 | 0.0001 | 2.66 | n=3 |
| CytoTRACE | R | dyngen_bifurcation | 500 | 150 | 4.001 | — | — | |
| CytoTRACE | R | dyngen_tree | 600 | 150 | 4.349 | — | — | |
| CytoTRACE | R | dyntoy_bifurcation | 300 | 100 | 4.301 | — | — | |
| CytoTRACE | R | dyntoy_convergence | 300 | 100 | 4.817 | — | — | |
| CytoTRACE | R | dyntoy_cycle | 300 | 100 | 4.385 | — | — | ERR |
| CytoTRACE | R | dyntoy_linear | 300 | 100 | 4.417 | — | — | |
| CytoTRACE | R | dyntoy_multifurcation | 400 | 120 | 4.14 | — | — | |
| CytoTRACE | R | dyntoy_tree | 400 | 120 | 4.67 | — | — | |
| CytoTRACE | R | prosstt_bifurcation | 300 | 100 | 7.056 | — | — | |
| CytoTRACE | R | prosstt_linear | 300 | 100 | 4.305 | — | — | ERR |
| CytoTRACE | R | prosstt_tree | 400 | 120 | 5.298 | — | — | |
| CytoTRACE | R | splatter_bifurcation | 300 | 100 | 5.13 | — | — | |
| CytoTRACE | R | splatter_linear | 300 | 100 | 9.997 | — | — | |
| CytoTRACE | R | splatter_tree | 400 | 120 | 4.038 | — | — | |
| CytoTRACE | Py | dyngen_bifurcation | 500 | 150 | 0.2003 | 0.0004 | 9.1 | n=3 |
| CytoTRACE | Py | dyngen_tree | 600 | 150 | 0.4732 | 0.2312 | 12.36 | n=3 |
| CytoTRACE | Py | dyntoy_bifurcation | 300 | 100 | 0.0491 | 0.0002 | 3.43 | n=3 |
| CytoTRACE | Py | dyntoy_convergence | 300 | 100 | 0.053 | 0.0003 | 3.43 | n=3 |
| CytoTRACE | Py | dyntoy_cycle | 300 | 100 | — | — | — | ERR: ValueError: array must not contain infs or NaNs |
| CytoTRACE | Py | dyntoy_linear | 300 | 100 | 0.0586 | 0.0001 | 3.43 | n=3 |

| | | | | | | | | |
|---|---|---|---|---|---|---|---|---|
| CytoTRACE | Py | dyntoy_multifurcation | 400 | 120 | 0.1288 | 0.0003 | 5.85 | n=3 |
| CytoTRACE | Py | dyntoy_tree | 400 | 120 | 0.1055 | 0.0006 | 5.85 | n=3 |
| CytoTRACE | Py | prosstt_bifurcation | 300 | 100 | 0.0494 | 0.0001 | 3.43 | n=3 |
| CytoTRACE | Py | prosstt_linear | 300 | 100 | 0.0593 | 0.0002 | 3.43 | n=3 |
| CytoTRACE | Py | prosstt_tree | 400 | 120 | 0.1081 | 0.0002 | 5.85 | n=3 |
| CytoTRACE | Py | splatter_bifurcation | 300 | 100 | 0.0518 | 0.0003 | 3.43 | n=3 |
| CytoTRACE | Py | splatter_linear | 300 | 100 | 0.0559 | 0.0001 | 3.43 | n=3 |
| CytoTRACE | Py | splatter_tree | 400 | 120 | 0.1111 | 0.0018 | 5.85 | n=3 |
| TSCAN | R | dyngen_bifurcation | 500 | 150 | 12.131 | — | — | |
| TSCAN | R | dyngen_tree | 600 | 150 | 20.228 | — | — | |
| TSCAN | R | dyntoy_bifurcation | 300 | 100 | 7.443 | — | — | |
| TSCAN | R | dyntoy_convergence | 300 | 100 | 8.457 | — | — | |
| TSCAN | R | dyntoy_cycle | 300 | 100 | 7.081 | — | — | ERR |
| TSCAN | R | dyntoy_linear | 300 | 100 | 7.97 | — | — | |
| TSCAN | R | dyntoy_multifurcation | 400 | 120 | 8.869 | — | — | |
| TSCAN | R | dyntoy_tree | 400 | 120 | 9.032 | — | — | |
| TSCAN | R | prosstt_bifurcation | 300 | 100 | 7.805 | — | — | |
| TSCAN | R | prosstt_linear | 300 | 100 | 7.532 | — | — | |
| TSCAN | R | prosstt_tree | 400 | 120 | 8.66 | — | — | |
| TSCAN | R | splatter_bifurcation | 300 | 100 | 7.971 | — | — | |
| TSCAN | R | splatter_linear | 300 | 100 | 7.791 | — | — | |
| TSCAN | R | splatter_tree | 400 | 120 | 15.309 | — | — | |
| TSCAN | Py | dyngen_bifurcation | 500 | 150 | 0.3648 | 0.0244 | 64.41 | n=3 |
| TSCAN | Py | dyngen_tree | 600 | 150 | 0.4062 | 0.0009 | 82.55 | n=3 |
| TSCAN | Py | dyntoy_bifurcation | 300 | 100 | 0.1865 | 0.0388 | 25.26 | n=3 |
| TSCAN | Py | dyntoy_convergence | 300 | 100 | 0.1895 | 0.0018 | 25.26 | n=3 |
| TSCAN | Py | dyntoy_cycle | 300 | 100 | 0.2581 | 0.0186 | 25.26 | n=3 |
| TSCAN | Py | dyntoy_linear | 300 | 100 | 1.7579 | 1.0522 | 17.31 | n=3 |
| TSCAN | Py | dyntoy_multifurcation | 400 | 120 | 0.2259 | 0.0019 | 52.33 | n=3 |
| TSCAN | Py | dyntoy_tree | 400 | 120 | 0.2101 | 0.0028 | 52.33 | n=3 |
| TSCAN | Py | prosstt_bifurcation | 300 | 100 | 0.127 | 0.0007 | 25.26 | n=3 |
| TSCAN | Py | prosstt_linear | 300 | 100 | 0.2246 | 0.0053 | 17.31 | n=3 |
| TSCAN | Py | prosstt_tree | 400 | 120 | 0.1903 | 0.003 | 52.33 | n=3 |
| TSCAN | Py | splatter_bifurcation | 300 | 100 | 0.2111 | 0.0378 | 25.26 | n=3 |
| TSCAN | Py | splatter_linear | 300 | 100 | 0.4647 | 0.1667 | 17.31 | n=3 |
| TSCAN | Py | splatter_tree | 400 | 120 | 0.2519 | 0.0193 | 52.33 | n=3 |
| SCORPIUS | R | dyngen_bifurcation | 500 | 150 | 4.178 | — | — | |
| SCORPIUS | R | dyngen_tree | 600 | 150 | 5.904 | — | — | |
| SCORPIUS | R | dyntoy_bifurcation | 300 | 100 | 4.089 | — | — | |
| SCORPIUS | R | dyntoy_convergence | 300 | 100 | 4.399 | — | — | |
| SCORPIUS | R | dyntoy_cycle | 300 | 100 | 4.328 | — | — | |
| SCORPIUS | R | dyntoy_linear | 300 | 100 | 6.188 | — | — | |
| SCORPIUS | R | dyntoy_multifurcation | 400 | 120 | 4.485 | — | — | |
| SCORPIUS | R | dyntoy_tree | 400 | 120 | 4.9 | — | — | |
| SCORPIUS | R | prosstt_bifurcation | 300 | 100 | 8.232 | — | — | |
| SCORPIUS | R | prosstt_linear | 300 | 100 | 7.224 | — | — | |
| SCORPIUS | R | prosstt_tree | 400 | 120 | 4.056 | — | — | |
| SCORPIUS | R | splatter_bifurcation | 300 | 100 | 4.37 | — | — | |
| SCORPIUS | R | splatter_linear | 300 | 100 | 4.346 | — | — | |
| SCORPIUS | R | splatter_tree | 400 | 120 | 4.056 | — | — | |
| SCORPIUS | Py | dyngen_bifurcation | 500 | 150 | 0.7323 | 0.1404 | 10.07 | n=3 |
| SCORPIUS | Py | dyngen_tree | 600 | 150 | 0.5704 | 0.1268 | 14.48 | n=3 |
| SCORPIUS | Py | dyntoy_bifurcation | 300 | 100 | 1.0158 | 0.823 | 3.65 | n=3 |
| SCORPIUS | Py | dyntoy_convergence | 300 | 100 | 0.7318 | 0.4761 | 3.65 | n=3 |
| SCORPIUS | Py | dyntoy_cycle | 300 | 100 | 1.3669 | 0.0243 | 3.65 | n=3 |
| SCORPIUS | Py | dyntoy_linear | 300 | 100 | 1.3861 | 0.0859 | 3.65 | n=3 |
| SCORPIUS | Py | dyntoy_multifurcation | 400 | 120 | 1.0174 | 0.6048 | 6.46 | n=3 |
| SCORPIUS | Py | dyntoy_tree | 400 | 120 | 1.5089 | 0.7886 | 6.46 | n=3 |
| SCORPIUS | Py | prosstt_bifurcation | 300 | 100 | 0.5461 | 0.1102 | 3.65 | n=3 |
| SCORPIUS | Py | prosstt_linear | 300 | 100 | 0.4462 | 0.1154 | 3.65 | n=3 |
| SCORPIUS | Py | prosstt_tree | 400 | 120 | 0.731 | 0.4653 | 6.46 | n=3 |
| SCORPIUS | Py | splatter_bifurcation | 300 | 100 | 0.772 | 0.5652 | 3.65 | n=3 |
| SCORPIUS | Py | splatter_linear | 300 | 100 | 0.7305 | 0.3498 | 3.65 | n=3 |

| | | | | | | | | |
|---|---|---|---|---|---|---|---|---|
| SCORPIUS | Py | splatter_tree | 400 | 120 | 1.0189 | 0.7755 | 6.46 | n=3 |
| Monocle3 | Py | dyngen_bifurcation | 500 | 150 | 3.2725 | 0.0466 | 264.58 | n=3 |
| Monocle3 | Py | dyngen_tree | 600 | 150 | 3.488 | 0.1054 | 264.85 | n=3 |
| Monocle3 | Py | dyntoy_bifurcation | 300 | 100 | 5.1932 | 1.6201 | 263.8 | n=3 |
| Monocle3 | Py | dyntoy_convergence | 300 | 100 | 4.0253 | 1.3096 | 263.93 | n=3 |
| Monocle3 | Py | dyntoy_cycle | 300 | 100 | 2.7524 | 0.348 | 273.96 | n=3 |
| Monocle3 | Py | dyntoy_linear | 300 | 100 | 3.0633 | 0.1433 | 273.98 | n=3 |
| Monocle3 | Py | dyntoy_multifurcatio n | 400 | 120 | 3.2879 | 0.3048 | 264.17 | n=3 |
| Monocle3 | Py | dyntoy_tree | 400 | 120 | 3.0284 | 0.075 | 264.17 | n=3 |
| Monocle3 | Py | prosstt_bifurcation | 300 | 100 | 2.7892 | 0.0345 | 263.8 | n=3 |
| Monocle3 | Py | prosstt_linear | 300 | 100 | 2.8479 | 0.1304 | 263.78 | n=3 |
| Monocle3 | Py | prosstt_tree | 400 | 120 | 3.1689 | 0.0773 | 264.19 | n=3 |
| Monocle3 | Py | splatter_bifurcation | 300 | 100 | 3.2534 | 0.6419 | 263.8 | n=3 |
| Monocle3 | Py | splatter_linear | 300 | 100 | 2.7838 | 0.096 | 263.8 | n=3 |
| Monocle3 | Py | splatter_tree | 400 | 120 | 4.7549 | 1.2549 | 264.17 | n=3 |

#### Part C | Benchmark configuration

| Item | Value |
|---|---|
| Hardware | Intel Xeon Platinum 8462Y+ · 64 logical cores · 1081 GB RAM |
| OS | Linux 3.10.0 (x86_64, glibc 2.17) |
| Environment | conda omicdev (/scratch/users/steorra/env/omicdev) |
| Python | 3.10.20 — numpy 2.2.6, scipy 1.15.3, scikit-learn 1.7.2, pandas 2.3.3 |
| R | 4.5.3 (same env; Rscript driver) |
| Timer | time.perf_counter() (re-measurement); original sweep time.time() |
| Warm-up | 1 untimed warm-up discarded before timed reps (re-measurement); original sweep none |
| Replicates | Python n=3 after warm-up (this work); R + original sweep n=1 |
| Memory metric | tracemalloc peak Python-heap per run (Python ports only); excludes C/BLAS off-heap; R not measured |
| Failures | Python re-measurement 95/98 OK (3 errored); original sweep 10/182 errored. |

#### Summary

- Every Python port faster than its R reference; median speed-up 22.6× (destiny 139×, URD 68×, Slingshot 27×, CytoTRACE 18×, TSCAN 16×, SCORPIUS 5×) — reproduces Fig. 3f / S2.
- Replicate variance (Python, re-measured): rep-CV 0.6% (URD) to ~44% (SCORPIUS); fast deterministic ports (destiny/URD/Slingshot) CV ≤ 2%.
- Peak memory (Python, re-measured): 2.3 MB (Slingshot) to 265 MB (Monocle3-py); native ports < 40 MB.
- Dataset-size dependence: runtime grows with cells (300→600) and genes (100→150); largest dataset dyngen_tree (600×150) dominates per-method means.
- Caveats: original Fig. 3f sweep = single time.time() run, no memory; this table adds warm-up+3-rep timing and tracemalloc peak memory for Python ports only. R variance/memory + off-heap (BLAS) memory remain unmeasured.

### *Supplementary Table 30 | Per-package evolution-log audit*

One row per reconstructed/targeted R package in the Fig. 3d landscape: methodology family, source ecosystem, scope, version, completeness and release status. Package-wide evolution-log audit: iterations, rejected, documented limitations, final parity (14 packages).

| Package | Scope | Iterations | Rejected | Documented limitations | Final parity | Key rejected / documented steps |
|---|---|---|---|---|---|---|
| py-tradeSeq | Trajectory — trajectory-based differential expression for scRNA-seq via NB-GAM (associationTest, startVsEndTest, patternTest) | 12 | 3 | 1 | 0.84 | First associationTest run: Spearman 0.40 on -log10(p); Discover statsmodels param naming: x0_s0, x0_s1, … not smooth_*; startVsEndTest only Spearman 0.45 — document as known limitation |

| | | | | | | |
|---|---|---|---|---|---|---|
| py-SCORPIUS | trajectory — linear trajectory inference for scRNA-seq via MDS + Hastie-Stuetzle principal curves | 11 | 1 | 0 | 0.989 | UnivariateSpline principal curve degenerates to zeros |
| py-condiments | trajectory: differential trajectory analysis across conditions in scRNA-seq | 9 | 0 | 1 | 1.0 | topologyTest discrepancy: R=0 sig, Py=1 sig — document |
| py-destiny | trajectory inference — diffusion maps + diffusion pseudotime (DPT) for scRNA-seq | 9 | 0 | 0 | 0.916 | — |
| py-URD | trajectory — branching pseudotime (flood-based) + random-walk lineage attribution for scRNA-seq | 9 | 2 | 0 | 0.985 | First parity run: 0 non-NA cells in pseudotime — flood stalled; Diagnose: destiny transition matrix is too sparse for single-cell root |
| py-TSCAN | Trajectory — mclust+MST+spline pseudotime reconstruction for scRNA-seq | 9 | 1 | 1 | 1.0 | Cluster ARI only 0.85 — investigate mclust divergence; difftest documented divergence: pygam ≠ mgcv at small df |
| py-Slingshot | trajectory — cell-lineage (MST/principal-curve) + pseudotime inference for scRNA-seq | 8 | 0 | 2 | 0.997 | LOWESS warning: invalid value encountered in divide; Document lineage count mismatch: R=3, Py=2 |
| py-Monocle3 | single-cell trajectory inference (principal-graph pseudotime) | 8 | 0 | 0 | 1.0 | — |
| py-CytoTRACE | trajectory: gene-count-based stemness / developmental-potential pseudotime for scRNA-seq | 7 | 1 | 1 | 0.93 | First smoke test failure: cytotrace negatively correlated with cell index; R-side parity attempt: HiClimR + ncdf4 fail to install |
| py-dynbenchmark | trajectory inference benchmarking (dyneval metrics + synthetic generators + harness) | 7 | 0 | 0 | 0.797 | — |
| py-spotsweeper | spatial QC (spatially-aware outlier/artifact detection for spatial transcriptomics: local variance, MAD local outliers, artifact + Visium outlier flagging) | 5 | 0 | 0 | F1=1.0 / ARI=1.0 | — |
| py-Statial | spatial proteomics / single-cell spatial (cell-state changes vs spatial associations: Kontextual L-function, distances, abundances, state changes) | 5 | 0 | 0 | machine-precision (Kontextual 5% rel) | — |
| py-dynwrap | trajectory inference data wrapper / benchmarking (geodesic distances, milestone networks for dyneval/dynbenchmark) | 4 | 0 | 0 | bit-equivalent geodesics | — |
| py-dyneval | trajectory inference benchmark metrics | 3 | 0 | 0 | corr 0.98 (8/14 metrics) | — |
| Total (14 packages) | — | 106 | 8 | 6 | 14/14 pass | — |

#### Summary

- 14 reconstructed packages carry a published evolution log (evolution.ipynb): the trajectory-inference family (SCORPIUS, tradeSeq, condiments, destiny, URD, Slingshot, Monocle3, TSCAN, CytoTRACE) + the dynverse benchmarking trio (dynwrap, dyneval, dynbenchmark) + spatial QC (spotsweeper, Statial).
- 106 total iterations recorded; 8 rejected and 6 documented-limitation steps — i.e. failures and caveats are logged, not just successes. Every logged package ended above its parity threshold (14/14 pass).
- Fig. 3g (py-Slingshot): 8 iterations, 0 rejected, 2 documented limitations (LOWESS warning on a sparse curve end; R emits 3 lineages vs Python 2 — neither affects pseudotime parity), final parity 0.997, overall benchmark score 0.485 > 0.455 (R), 27× faster.
- Fig. 3h (py-URD): 9 iterations, 2 rejected (flood stalled → 0 cells reached → diagnosed as the R transition matrix being too sparse for a single-root flood), then the accepted iter-3 fix (16-cell root cluster + min_cells_flooded=1) drove parity 0 → 0.985 in one step; final overall 0.439 > 0.348 (R), 68× faster.
- Most rejections per package: py-tradeSeq (3 rejected — NB-GAM convergence/identifiability fixes) and py-URD (2). The audit shows the same accept/reject/document discipline across unrelated algorithm families (GAMs, random walks, principal curves, diffusion maps, spatial QC).

- Coverage caveat: evolution.ipynb logs are published for these 14 of ~62 reconstructed packages; the acceleration-specific ITERATION_LOG.md audit (7 packages) is reported separately in Supplementary Table 4_4. The remaining packages report a final parity (Supplementary Table 2) but no per-iteration reconstruction log yet — closing that gap across the catalogue is the recommended next step.
- Supplementary Notebooks: the full py-Slingshot and py-URD evolution.ipynb (per-iteration code, parity, runtime, rationale) are provided as supplementary notebooks (supplementary_notebooks/py-Slingshot_evolution.ipynb, py-URD_evolution.ipynb).

### *Supplementary Table 31 | PseudotimeFate terminal macrostates (pancreatic endocrine)*

3,696 cells × 3,000 HVGs; Palantir pseudotime; PseudotimeFate (n_macrostates=10, residency_threshold=0.60, late_pt_quantile=0.70). Four mature endocrine terminals: Beta, Delta, Epsilon, Alpha.

#### Part A | Terminal macrostates

| Endocrine terminal | Macrostate ID | Self-residency | Dominant cluster | Cluster purity | Pseudotime rank | # cells | Terminal cell cluster |
|---|---|---|---|---|---|---|---|
| Beta | ms9 | 0.9266 | Beta | 0.395 | 0.788 | 1422 | Beta |
| Delta | ms2 | 0.9017 | Delta | 1.0 | 0.813 | 24 | Delta |
| Epsilon | ms4 | 0.8169 | Epsilon | 0.905 | 0.856 | 21 | Epsilon |
| Alpha | ms7 | 0.7524 | Alpha | 0.977 | 0.867 | 88 | Alpha |

#### Part B | All macrostates + terminal-cell selection criteria

| Macrostate ID | Self-residency | Dominant cluster | Cluster purity | Mean pt-rank | # cells | Terminal? | Selection outcome |
|---|---|---|---|---|---|---|---|
| ms9 | 0.9266 | Beta | 0.395 | 0.788 | 1422 | Yes | TERMINAL |
| ms0 | 0.9255 | Ngn3 high EP | 0.85 | 0.409 | 20 | No | excluded — pt-rank<0.7 |
| ms8 | 0.9021 | Ngn3 high EP | 0.421 | 0.349 | 1472 | No | excluded — pt-rank<0.7 |
| ms2 | 0.9017 | Delta | 1.0 | 0.813 | 24 | Yes | TERMINAL |
| ms5 | 0.8768 | Ductal | 0.859 | 0.381 | 156 | No | excluded — pt-rank<0.7 |
| ms3 | 0.834 | Ductal | 0.795 | 0.168 | 361 | No | excluded — initial state |
| ms6 | 0.8223 | Ductal | 0.689 | 0.335 | 103 | No | excluded — pt-rank<0.7 |
| ms4 | 0.8169 | Epsilon | 0.905 | 0.856 | 21 | Yes | TERMINAL |
| ms7 | 0.7524 | Alpha | 0.977 | 0.867 | 88 | Yes | TERMINAL |
| ms1 | 0.7369 | Beta | 1.0 | 0.981 | 29 | No | excluded — cluster dedup (vs ms9) |

#### Configuration

| Parameter | Value |
|---|---|
| n_macrostates | 10 |
| residency_threshold | 0.6 |
| late_pt_quantile | 0.7 |
| groupby | clusters |
| pseudotime_key | palantir_pseudotime |
| dataset | 3696 cells × 3000 HVGs |
| clusters | Alpha, Beta, Delta, Ductal, Epsilon, Ngn3 high EP, Ngn3 low EP, Pre-endocrine |
| initial state | ms3 (lowest mean pseudotime) |

### *Supplementary Table 32 | Lineage driver genes — pancreatic endocrine fates*

Pearson correlation (gene expression vs fate probability) with Fisher-z 95% CI and BH-adjusted q, top 20 per lineage. Headline markers Gcg/Pdx1/Ins2/Sst/Ghrl reproduce the main-text values exactly.

| Lineage | Rank | Gene | Correlation | 95% CI | Adjusted P (q) | Canonical marker |
|---|---|---|---|---|---|---|
| Alpha | 1 | Gcg | 0.593 | [0.571, 0.613] | 0 (<ε) | ✓ |
| Alpha | 2 | Irx1 | 0.532 | [0.509, 0.555] | 0 (<ε) | ✓ |
| Alpha | 3 | Irx2 | 0.465 | [0.439, 0.490] | 0 (<ε) | ✓ |
| Alpha | 4 | Ctxn2 | 0.386 | [0.358, 0.413] | 0 (<ε) | |
| Alpha | 5 | Peg10 | 0.382 | [0.354, 0.409] | 0 (<ε) | |
| Alpha | 6 | Wnk3 | 0.375 | [0.347, 0.403] | 0 (<ε) | |
| Alpha | 7 | Rgs4 | 0.360 | [0.331, 0.387] | 0 (<ε) | |
| Alpha | 8 | Smarca1 | 0.347 | [0.319, 0.375] | 0 (<ε) | |
| Alpha | 9 | Dock11 | 0.341 | [0.312, 0.369] | 0 (<ε) | |
| Alpha | 10 | Edn3 | 0.324 | [0.295, 0.353] | 0 (<ε) | |
| Alpha | 11 | Etv1 | 0.323 | [0.294, 0.352] | 0 (<ε) | |
| Alpha | 12 | Tmsb15l | 0.321 | [0.292, 0.350] | 0 (<ε) | |
| Alpha | 13 | Tmsb15b2 | 0.317 | [0.288, 0.346] | 0 (<ε) | |
| Alpha | 14 | Pla2g1b | 0.307 | [0.278, 0.336] | 0 (<ε) | |
| Alpha | 15 | Arx | 0.301 | [0.271, 0.330] | 0 (<ε) | ✓ |
| Alpha | 16 | Gast | 0.296 | [0.266, 0.325] | 0 (<ε) | |
| Alpha | 17 | Asb4 | 0.290 | [0.260, 0.319] | 0 (<ε) | |
| Alpha | 18 | Sphkap | 0.284 | [0.254, 0.314] | 0 (<ε) | |
| Alpha | 19 | Tmem27 | 0.275 | [0.245, 0.304] | 0 (<ε) | |
| Alpha | 20 | Gria2 | 0.269 | [0.239, 0.298] | 0 (<ε) | |
| Beta | 1 | Pdx1 | 0.498 | [0.474, 0.522] | 0 (<ε) | ✓ |
| Beta | 2 | Ins2 | 0.491 | [0.466, 0.515] | 0 (<ε) | ✓ |
| Beta | 3 | Mnx1 | 0.486 | [0.461, 0.511] | 0 (<ε) | ✓ |
| Beta | 4 | Gng12 | 0.486 | [0.461, 0.510] | 0 (<ε) | |
| Beta | 5 | Nkx6-1 | 0.479 | [0.454, 0.504] | 0 (<ε) | ✓ |
| Beta | 6 | Sytl4 | 0.474 | [0.449, 0.499] | 0 (<ε) | |
| Beta | 7 | Nnat | 0.459 | [0.433, 0.484] | 0 (<ε) | |
| Beta | 8 | Adra2a | 0.451 | [0.425, 0.477] | 0 (<ε) | |
| Beta | 9 | Gm27033 | 0.417 | [0.390, 0.443] | 0 (<ε) | |
| Beta | 10 | Npepl1 | 0.410 | [0.383, 0.437] | 0 (<ε) | |
| Beta | 11 | Cryba2 | 0.382 | [0.354, 0.409] | 0 (<ε) | |
| Beta | 12 | Mafb | 0.377 | [0.349, 0.405] | 0 (<ε) | |
| Beta | 13 | Mapt | 0.374 | [0.346, 0.401] | 0 (<ε) | |
| Beta | 14 | C2cd4b | 0.368 | [0.340, 0.396] | 0 (<ε) | |
| Beta | 15 | Chgb | 0.362 | [0.334, 0.390] | 0 (<ε) | |
| Beta | 16 | Ins1 | 0.358 | [0.329, 0.385] | 0 (<ε) | ✓ |
| Beta | 17 | Frzb | 0.351 | [0.322, 0.379] | 0 (<ε) | |
| Beta | 18 | Mlxipl | 0.342 | [0.313, 0.370] | 0 (<ε) | |
| Beta | 19 | Ero1lb | 0.340 | [0.311, 0.368] | 0 (<ε) | |
| Beta | 20 | G6pc2 | 0.338 | [0.310, 0.367] | 0 (<ε) | |
| Delta | 1 | Sst | 0.658 | [0.640, 0.676] | 0 (<ε) | ✓ |
| Delta | 2 | Hhex | 0.355 | [0.327, 0.383] | 0 (<ε) | ✓ |
| Delta | 3 | Arg1 | 0.343 | [0.314, 0.371] | 0 (<ε) | |

| | | | | | | |
|---|---|---|---|---|---|---|
| Delta | 4 | Mest | 0.296 | [0.266, 0.325] | 0 (<ε) | |
| Delta | 5 | Mef2c | 0.295 | [0.265, 0.324] | 0 (<ε) | |
| Delta | 6 | Ptprz1 | 0.291 | [0.261, 0.320] | 0 (<ε) | |
| Delta | 7 | Cd24a | 0.274 | [0.244, 0.303] | 0 (<ε) | |
| Delta | 8 | Nrsn1 | 0.267 | [0.237, 0.297] | 0 (<ε) | |
| Delta | 9 | Cfap61 | 0.264 | [0.233, 0.293] | 0 (<ε) | |
| Delta | 10 | Spock3 | 0.255 | [0.225, 0.285] | 0 (<ε) | |
| Delta | 11 | Th | 0.252 | [0.222, 0.282] | 0 (<ε) | |
| Delta | 12 | Fam159b | 0.249 | [0.219, 0.279] | 0 (<ε) | |
| Delta | 13 | Pdyn | 0.218 | [0.187, 0.249] | 0 (<ε) | |
| Delta | 14 | Frzb | 0.216 | [0.185, 0.246] | 0 (<ε) | |
| Delta | 15 | Gap43 | 0.214 | [0.183, 0.245] | 0 (<ε) | |
| Delta | 16 | Scn3a | 0.213 | [0.182, 0.244] | 0 (<ε) | |
| Delta | 17 | Igfbp5 | 0.212 | [0.181, 0.243] | 0 (<ε) | |
| Delta | 18 | Kctd8 | 0.210 | [0.179, 0.241] | 0 (<ε) | |
| Delta | 19 | Bmp3 | 0.209 | [0.178, 0.240] | 0 (<ε) | |
| Delta | 20 | Fxyd7 | 0.208 | [0.177, 0.239] | 0 (<ε) | |
| Epsilon | 1 | Ghrl | 0.523 | [0.499, 0.546] | 0 (<ε) | ✓ |
| Epsilon | 2 | Mboat4 | 0.463 | [0.438, 0.488] | 0 (<ε) | ✓ |
| Epsilon | 3 | Mdk | 0.437 | [0.411, 0.463] | 0 (<ε) | |
| Epsilon | 4 | Serpina1c | 0.410 | [0.383, 0.436] | 0 (<ε) | |
| Epsilon | 5 | Irs4 | 0.401 | [0.373, 0.427] | 0 (<ε) | |
| Epsilon | 6 | Cdkn1a | 0.391 | [0.363, 0.418] | 0 (<ε) | |
| Epsilon | 7 | Anpep | 0.371 | [0.343, 0.398] | 0 (<ε) | |
| Epsilon | 8 | Gm11837 | 0.361 | [0.333, 0.389] | 0 (<ε) | |
| Epsilon | 9 | Serpina1e | 0.339 | [0.310, 0.367] | 0 (<ε) | |
| Epsilon | 10 | Upk3bl | 0.313 | [0.284, 0.342] | 0 (<ε) | |
| Epsilon | 11 | St8sia2 | 0.313 | [0.284, 0.342] | 0 (<ε) | |
| Epsilon | 12 | Arg1 | 0.303 | [0.273, 0.332] | 0 (<ε) | |
| Epsilon | 13 | Mfap4 | 0.302 | [0.272, 0.331] | 0 (<ε) | |
| Epsilon | 14 | Cldn6 | 0.297 | [0.267, 0.326] | 0 (<ε) | |
| Epsilon | 15 | Ppp1r14a | 0.295 | [0.265, 0.324] | 0 (<ε) | |
| Epsilon | 16 | Ifitm2 | 0.292 | [0.262, 0.321] | 0 (<ε) | |
| Epsilon | 17 | Lrrtm3 | 0.292 | [0.262, 0.321] | 0 (<ε) | |
| Epsilon | 18 | Cd24a | 0.291 | [0.261, 0.320] | 0 (<ε) | |
| Epsilon | 19 | Serpinh1 | 0.290 | [0.260, 0.319] | 0 (<ε) | |
| Epsilon | 20 | Igfbpl1 | 0.289 | [0.260, 0.319] | 0 (<ε) | |

#### *Supplementary Table 33*

Source datasets and per-(organ × age) reference assembly for the whole-body spatial AD-risk analysis. 809,284 cells across 126 organ–age combinations, projected onto the 6-week whole-body Array-seq spatial reference (GSE248904) via CMap. Each sample feeds gsMap spatial-LDSC to produce per-spot AD heritability.

#### 1 | Source datasets (5)

| rank | source | paper | accession | assay | organs_covered | ages_offered | preprocessing_status |
|---|---|---|---|---|---|---|---|
| **1** | PanSci | Zhang et al. Nature 2024 | GSE247719 | sci-RNA-seq3 (PanSci) | Kidney, Lung, Heart, Liver, Muscle, Stomach, BAT, iWAT, gWAT, Ileum, Colon, Pancreas, Skin, Thymus | 3, 6, 12, 16, 23 months | Curated cellType + Lineage + Sub_cell_type annotations from source |
| **2** | EasySci | Sziraki et al. Nature Aging 2023 | GSE212606 | EasySci (sci-RNA-seq3 variant, brain-optimised) | Brain (whole) | 3, 6, 21 months | Main_cell_type + Sub_cluster from source |
| **3** | TMS | Schaum et al. Nature 2020 (Tabula Muris Senis) | figshare:8273102 (TMS) | Smart-seq2 (FACS) + 10x Chromium (Droplet) | 23 organs (BAT, Bone Marrow, Brain Myeloid/NonMyeloid, Heart, Kidney, Liver, Lung, Pancreas, Spleen, Skin, Muscle, Stomach, Colon, Thymus, Trachea, …) | 1, 3, 18, 21, 24, 30 months | cell_ontology_class from source (Tabula Muris cell ontology) |
| **4** | MCA2 | Han et al. Cell 2022 (Mouse Cell Atlas 2.0) | GSE153562 | Microwell-seq (MCA2) | Stomach, Ileum, Spleen, Thymus (mid-late life gaps for TMS organs) | 6, 12, 18, 21, 24 months (per-organ subset) | leiden clusters with marker-based manual annotation |
| **5** | Hammond_microglia | Olah et al. Nature Communications 2020 | GSE179358 | 10x Chromium (microglia FACS-sorted, brain) | Brain (microglia only) | 6, 12, 18 months | cellType=microglial cell (single-cell-type focus) |

#### 2 | Per-source contribution

| source | samples | distinct_organs | total_cells | age_range | pct_of_atlas |
|---|---|---|---|---|---|
| **PanSci** | 47 | 10 | 454620 | 3–23 mo | 56.18 |
| **TMS** | 65 | 13 | 207717 | 1–30 mo | 25.67 |
| **MCA2** | 12 | 6 | 110309 | 6–24 mo | 13.63 |
| **EasySci** | 3 | 1 | 26640 | 3–21 mo | 3.29 |
| **Hammond_microglia** | 4 | 1 | 9998 | 6–24 mo | 1.24 |
| **Total** | 126 | 15 | 809284 | 1–30 mo | 100.0 |

#### 3 | Full per-(organ × age × source) ledger

*131 rows; 5 are multi-source Brain samples where ≥1 dataset is co-mapped to the same spatial reference.*

| organ | age_months | source | sample_id | n_cells | source_accession | preprocess_status | mapping_target |
|---|---|---|---|---|---|---|---|
| **BAT** | 3 | PanSci | BAT_03_months | 10000 | GSE247719 | Curated cellType + Lineage + Sub_cell_type | Brown_Fat_CTRL1 (GSE248904) |

| | | | | | | | |
|---|---|---|---|---|---|---|---|
| | | | | | | annotations from source | |
| **BAT** | 3 | TMS | BAT_3m | 713 | figshare:8273102 (TMS) | cell_ontology_class from source (Tabula Muris cell ontology) | Brown_Fat_CTRL1 (GSE248904) |
| **BAT** | 6 | PanSci | BAT_06_months | 10000 | GSE247719 | Curated cellType + Lineage + Sub_cell_type annotations from source | Brown_Fat_CTRL1 (GSE248904) |
| **BAT** | 12 | PanSci | BAT_12_months | 10000 | GSE247719 | Curated cellType + Lineage + Sub_cell_type annotations from source | Brown_Fat_CTRL1 (GSE248904) |
| **BAT** | 16 | PanSci | BAT_16_months | 10000 | GSE247719 | Curated cellType + Lineage + Sub_cell_type annotations from source | Brown_Fat_CTRL1 (GSE248904) |
| **BAT** | 18 | TMS | BAT_18m | 662 | figshare:8273102 (TMS) | cell_ontology_class from source (Tabula Muris cell ontology) | Brown_Fat_CTRL1 (GSE248904) |
| **BAT** | 21 | TMS | BAT_21m | 794 | figshare:8273102 (TMS) | cell_ontology_class from source (Tabula Muris cell ontology) | Brown_Fat_CTRL1 (GSE248904) |
| **BAT** | 23 | PanSci | BAT_23_months | 10000 | GSE247719 | Curated cellType + Lineage + Sub_cell_type annotations from source | Brown_Fat_CTRL1 (GSE248904) |
| **BAT** | 24 | TMS | BAT_24m | 848 | figshare:8273102 (TMS) | cell_ontology_class from source (Tabula Muris cell ontology) | Brown_Fat_CTRL1 (GSE248904) |
| **BAT** | 30 | TMS | BAT_30m | 2015 | figshare:8273102 (TMS) | cell_ontology_class from source (Tabula Muris cell ontology) | Brown_Fat_CTRL1 (GSE248904) |
| **Bone_Marrow** | 1 | TMS | Bone_Marrow_1m | 3027 | figshare:8273102 (TMS) | cell_ontology_class from source (Tabula Muris cell ontology) | Bone_Marrow_CTRL1 (GSE248904) |
| **Bone_Marrow** | 3 | TMS | Bone_Marrow_3m | 3490 | figshare:8273102 (TMS) | cell_ontology_class from source (Tabula Muris cell ontology) | Bone_Marrow_CTRL1 (GSE248904) |
| **Bone_Marrow** | 18 | TMS | Bone_Marrow_18m | 6712 | figshare:8273102 (TMS) | cell_ontology_class from source (Tabula | Bone_Marrow_CTRL1 (GSE248904) |

| | | | | | | | |
|---|---|---|---|---|---|---|---|
| | | | | | | Muris cell ontology) | |
| **Bone_Marrow** | 21 | TMS | Bone_Marrow_21m | 5216 | figshare:8273102 (TMS) | cell_ontology_class from source (Tabula Muris cell ontology) | Bone_Marrow_CTRL1 (GSE248904) |
| **Bone_Marrow** | 24 | TMS | Bone_Marrow_24m | 8279 | figshare:8273102 (TMS) | cell_ontology_class from source (Tabula Muris cell ontology) | Bone_Marrow_CTRL1 (GSE248904) |
| **Bone_Marrow** | 30 | TMS | Bone_Marrow_30m | 10000 | figshare:8273102 (TMS) | cell_ontology_class from source (Tabula Muris cell ontology) | Bone_Marrow_CTRL1 (GSE248904) |
| **Brain** | 3 | EasySci | Brain_03_months | 8875 | GSE212606 | Main_cell_type + Sub_cluster from source | Brain_CTRL1 (GSE248904) |
| **Brain** | 3 | PanSci | Brain_3m | 4405 | GSE247719 | Curated cellType + Lineage + Sub_cell_type annotations from source | Brain_CTRL1 (GSE248904) |
| **Brain** | 3 | TMS | Brain_3m | 4643 | figshare:8273102 (TMS) | cell_ontology_class from source (Tabula Muris cell ontology) | Brain_CTRL1 (GSE248904) |
| **Brain** | 6 | EasySci | Brain_06_months | 8892 | GSE212606 | Main_cell_type + Sub_cluster from source | Brain_CTRL1 (GSE248904) |
| **Brain** | 6 | Hammond_microglia | Brain_6m | 2528 | GSE179358 | cellType=microglial cell (single-cell-type focus) | Brain_CTRL1 (GSE248904) |
| **Brain** | 6 | PanSci | Brain_6m | 4478 | GSE247719 | Curated cellType + Lineage + Sub_cell_type annotations from source | Brain_CTRL1 (GSE248904) |
| **Brain** | 12 | Hammond_microglia | Brain_12m | 2197 | GSE179358 | cellType=microglial cell (single-cell-type focus) | Brain_CTRL1 (GSE248904) |
| **Brain** | 12 | MCA2 | Brain_12m | 4242 | GSE153562 | leiden clusters with marker-based manual annotation | Brain_CTRL1 (GSE248904) |
| **Brain** | 18 | Hammond_microglia | Brain_18m | 2614 | GSE179358 | cellType=microglial cell (single-cell-type focus) | Brain_CTRL1 (GSE248904) |
| **Brain** | 18 | TMS | Brain_18m | 4022 | figshare:8273102 (TMS) | cell_ontology_class from source (Tabula Muris cell ontology) | Brain_CTRL1 (GSE248904) |
| **Brain** | 21 | EasySci | Brain_21_months | 8873 | GSE212606 | Main_cell_type + Sub_cluster from source | Brain_CTRL1 (GSE248904) |

| | | | | | | | |
|---|---|---|---|---|---|---|---|
| **Brain** | 21 | TMS | Brain_21m | 11824 | figshare:8273102 (TMS) | cell_ontology_class from source (Tabula Muris cell ontology) | Brain_CTRL1 (GSE248904) |
| **Brain** | 24 | Hammond_microglia | Brain_24m | 2659 | GSE179358 | cellType=microglial cell (single-cell-type focus) | Brain_CTRL1 (GSE248904) |
| **Brain** | 24 | TMS | Brain_24m | 2677 | figshare:8273102 (TMS) | cell_ontology_class from source (Tabula Muris cell ontology) | Brain_CTRL1 (GSE248904) |
| **Colon** | 3 | PanSci | Colon_03_months | 10000 | GSE247719 | Curated cellType + Lineage + Sub_cell_type annotations from source | Colon_CTRL1 (GSE248904) |
| **Colon** | 3 | TMS | Colon_3m | 3987 | figshare:8273102 (TMS) | cell_ontology_class from source (Tabula Muris cell ontology) | Colon_CTRL1 (GSE248904) |
| **Colon** | 6 | PanSci | Colon_06_months | 10000 | GSE247719 | Curated cellType + Lineage + Sub_cell_type annotations from source | Colon_CTRL1 (GSE248904) |
| **Colon** | 12 | PanSci | Colon_12_months | 10000 | GSE247719 | Curated cellType + Lineage + Sub_cell_type annotations from source | Colon_CTRL1 (GSE248904) |
| **Colon** | 16 | PanSci | Colon_16_months | 10000 | GSE247719 | Curated cellType + Lineage + Sub_cell_type annotations from source | Colon_CTRL1 (GSE248904) |
| **Colon** | 18 | TMS | Colon_18m | 2369 | figshare:8273102 (TMS) | cell_ontology_class from source (Tabula Muris cell ontology) | Colon_CTRL1 (GSE248904) |
| **Colon** | 21 | MCA2 | Colon_21m | 10000 | GSE153562 | leiden clusters with marker-based manual annotation | Colon_CTRL1 (GSE248904) |
| **Colon** | 23 | PanSci | Colon_23_months | 10000 | GSE247719 | Curated cellType + Lineage + Sub_cell_type annotations from source | Colon_CTRL1 (GSE248904) |
| **Colon** | 24 | TMS | Colon_24m | 1955 | figshare:8273102 (TMS) | cell_ontology_class from source (Tabula Muris cell ontology) | Colon_CTRL1 (GSE248904) |

| | | | | | | | |
|---|---|---|---|---|---|---|---|
| **Colon** | 30 | TMS | Colon_30m | 1887 | figshare:8273102 (TMS) | cell_ontology_class from source (Tabula Muris cell ontology) | Colon_CTRL1 (GSE248904) |
| **Heart** | 1 | TMS | Heart_1m | 946 | figshare:8273102 (TMS) | cell_ontology_class from source (Tabula Muris cell ontology) | Heart_CTRL1 (GSE248904) |
| **Heart** | 3 | PanSci | Heart_03_months | 10000 | GSE247719 | Curated cellType + Lineage + Sub_cell_type annotations from source | Heart_CTRL1 (GSE248904) |
| **Heart** | 3 | TMS | Heart_3m | 534 | figshare:8273102 (TMS) | cell_ontology_class from source (Tabula Muris cell ontology) | Heart_CTRL1 (GSE248904) |
| **Heart** | 6 | PanSci | Heart_06_months | 10000 | GSE247719 | Curated cellType + Lineage + Sub_cell_type annotations from source | Heart_CTRL1 (GSE248904) |
| **Heart** | 12 | PanSci | Heart_12_months | 10000 | GSE247719 | Curated cellType + Lineage + Sub_cell_type annotations from source | Heart_CTRL1 (GSE248904) |
| **Heart** | 16 | PanSci | Heart_16_months | 10000 | GSE247719 | Curated cellType + Lineage + Sub_cell_type annotations from source | Heart_CTRL1 (GSE248904) |
| **Heart** | 18 | TMS | Heart_18m | 1143 | figshare:8273102 (TMS) | cell_ontology_class from source (Tabula Muris cell ontology) | Heart_CTRL1 (GSE248904) |
| **Heart** | 21 | TMS | Heart_21m | 918 | figshare:8273102 (TMS) | cell_ontology_class from source (Tabula Muris cell ontology) | Heart_CTRL1 (GSE248904) |
| **Heart** | 23 | PanSci | Heart_23_months | 10000 | GSE247719 | Curated cellType + Lineage + Sub_cell_type annotations from source | Heart_CTRL1 (GSE248904) |
| **Heart** | 24 | TMS | Heart_24m | 768 | figshare:8273102 (TMS) | cell_ontology_class from source (Tabula Muris cell ontology) | Heart_CTRL1 (GSE248904) |
| **Heart** | 30 | TMS | Heart_30m | 4304 | figshare:8273102 (TMS) | cell_ontology_class from source (Tabula Muris cell ontology) | Heart_CTRL1 (GSE248904) |

| | | | | | | | |
|---|---|---|---|---|---|---|---|
| **Ileum** | 3 | PanSci | Ileum_03_months | 9994 | GSE247719 | Curated cellType + Lineage + Sub_cell_type annotations from source | Small_Intestine_CTRL1 (GSE248904) |
| **Ileum** | 6 | PanSci | Ileum_06_months | 9994 | GSE247719 | Curated cellType + Lineage + Sub_cell_type annotations from source | Small_Intestine_CTRL1 (GSE248904) |
| **Ileum** | 12 | MCA2 | Ileum_12m | 9981 | GSE153562 | leiden clusters with marker-based manual annotation | Small_Intestine_CTRL1 (GSE248904) |
| **Ileum** | 12 | PanSci | Ileum_12_months | 9996 | GSE247719 | Curated cellType + Lineage + Sub_cell_type annotations from source | Small_Intestine_CTRL1 (GSE248904) |
| **Ileum** | 16 | PanSci | Ileum_16_months | 9995 | GSE247719 | Curated cellType + Lineage + Sub_cell_type annotations from source | Small_Intestine_CTRL1 (GSE248904) |
| **Ileum** | 18 | MCA2 | Ileum_18m | 9983 | GSE153562 | leiden clusters with marker-based manual annotation | Small_Intestine_CTRL1 (GSE248904) |
| **Ileum** | 21 | MCA2 | Ileum_21m | 9985 | GSE153562 | leiden clusters with marker-based manual annotation | Small_Intestine_CTRL1 (GSE248904) |
| **Ileum** | 23 | PanSci | Ileum_23_months | 9998 | GSE247719 | Curated cellType + Lineage + Sub_cell_type annotations from source | Small_Intestine_CTRL1 (GSE248904) |
| **Ileum** | 24 | MCA2 | Ileum_24m | 9986 | GSE153562 | leiden clusters with marker-based manual annotation | Small_Intestine_CTRL1 (GSE248904) |
| **Kidney** | 1 | TMS | Kidney_1m | 2422 | figshare:8273102 (TMS) | cell_ontology_class from source (Tabula Muris cell ontology) | Kidney_CTRL1 (GSE248904) |
| **Kidney** | 3 | PanSci | Kidney_03_months | 9802 | GSE247719 | Curated cellType + Lineage + Sub_cell_type annotations from source | Kidney_CTRL1 (GSE248904) |
| **Kidney** | 3 | TMS | Kidney_3m | 2254 | figshare:8273102 (TMS) | cell_ontology_class from source (Tabula | Kidney_CTRL1 (GSE248904) |

| | | | | | | | |
|---|---|---|---|---|---|---|---|
| | | | | | | Muris cell ontology) | |
| **Kidney** | 6 | PanSci | Kidney_06_months | 9773 | GSE247719 | Curated cellType + Lineage + Sub_cell_type annotations from source | Kidney_CTRL1 (GSE248904) |
| **Kidney** | 12 | PanSci | Kidney_12_months | 9795 | GSE247719 | Curated cellType + Lineage + Sub_cell_type annotations from source | Kidney_CTRL1 (GSE248904) |
| **Kidney** | 16 | PanSci | Kidney_16_months | 9800 | GSE247719 | Curated cellType + Lineage + Sub_cell_type annotations from source | Kidney_CTRL1 (GSE248904) |
| **Kidney** | 18 | TMS | Kidney_18m | 3040 | figshare:8273102 (TMS) | cell_ontology_class from source (Tabula Muris cell ontology) | Kidney_CTRL1 (GSE248904) |
| **Kidney** | 21 | TMS | Kidney_21m | 2204 | figshare:8273102 (TMS) | cell_ontology_class from source (Tabula Muris cell ontology) | Kidney_CTRL1 (GSE248904) |
| **Kidney** | 23 | PanSci | Kidney_23_months | 9795 | GSE247719 | Curated cellType + Lineage + Sub_cell_type annotations from source | Kidney_CTRL1 (GSE248904) |
| **Kidney** | 24 | TMS | Kidney_24m | 5508 | figshare:8273102 (TMS) | cell_ontology_class from source (Tabula Muris cell ontology) | Kidney_CTRL1 (GSE248904) |
| **Kidney** | 30 | TMS | Kidney_30m | 5511 | figshare:8273102 (TMS) | cell_ontology_class from source (Tabula Muris cell ontology) | Kidney_CTRL1 (GSE248904) |
| **Liver** | 1 | TMS | Liver_1m | 2785 | figshare:8273102 (TMS) | cell_ontology_class from source (Tabula Muris cell ontology) | Liver_CTRL1 (GSE248904) |
| **Liver** | 3 | PanSci | Liver_03_months | 9981 | GSE247719 | Curated cellType + Lineage + Sub_cell_type annotations from source | Liver_CTRL1 (GSE248904) |
| **Liver** | 3 | TMS | Liver_3m | 729 | figshare:8273102 (TMS) | cell_ontology_class from source (Tabula Muris cell ontology) | Liver_CTRL1 (GSE248904) |
| **Liver** | 6 | PanSci | Liver_06_months | 9973 | GSE247719 | Curated cellType + Lineage + | Liver_CTRL1 (GSE248904) |

| | | | | | | | |
|---|---|---|---|---|---|---|---|
| | | | | | | Sub_cell_type annotations from source | |
| **Liver** | 12 | PanSci | Liver_12_months | 9969 | GSE247719 | Curated cellType + Lineage + Sub_cell_type annotations from source | Liver_CTRL1 (GSE248904) |
| **Liver** | 16 | PanSci | Liver_16_months | 9969 | GSE247719 | Curated cellType + Lineage + Sub_cell_type annotations from source | Liver_CTRL1 (GSE248904) |
| **Liver** | 18 | TMS | Liver_18m | 1176 | figshare:8273102 (TMS) | cell_ontology_class from source (Tabula Muris cell ontology) | Liver_CTRL1 (GSE248904) |
| **Liver** | 21 | TMS | Liver_21m | 367 | figshare:8273102 (TMS) | cell_ontology_class from source (Tabula Muris cell ontology) | Liver_CTRL1 (GSE248904) |
| **Liver** | 23 | PanSci | Liver_23_months | 9983 | GSE247719 | Curated cellType + Lineage + Sub_cell_type annotations from source | Liver_CTRL1 (GSE248904) |
| **Liver** | 24 | TMS | Liver_24m | 945 | figshare:8273102 (TMS) | cell_ontology_class from source (Tabula Muris cell ontology) | Liver_CTRL1 (GSE248904) |
| **Liver** | 30 | TMS | Liver_30m | 2272 | figshare:8273102 (TMS) | cell_ontology_class from source (Tabula Muris cell ontology) | Liver_CTRL1 (GSE248904) |
| **Lung** | 1 | TMS | Lung_1m | 2507 | figshare:8273102 (TMS) | cell_ontology_class from source (Tabula Muris cell ontology) | Lung_CTRL1 (GSE248904) |
| **Lung** | 3 | PanSci | Lung_03_months | 9996 | GSE247719 | Curated cellType + Lineage + Sub_cell_type annotations from source | Lung_CTRL1 (GSE248904) |
| **Lung** | 3 | TMS | Lung_3m | 3610 | figshare:8273102 (TMS) | cell_ontology_class from source (Tabula Muris cell ontology) | Lung_CTRL1 (GSE248904) |
| **Lung** | 6 | PanSci | Lung_06_months | 9991 | GSE247719 | Curated cellType + Lineage + Sub_cell_type annotations from source | Lung_CTRL1 (GSE248904) |
| **Lung** | 12 | PanSci | Lung_12_months | 9995 | GSE247719 | Curated cellType + | Lung_CTRL1 (GSE248904) |

| | | | | | | | |
|---|---|---|---|---|---|---|---|
| | | | | | | Lineage + Sub_cell_type annotations from source | |
| **Lung** | 16 | PanSci | Lung_16_months | 9996 | GSE247719 | Curated cellType + Lineage + Sub_cell_type annotations from source | Lung_CTRL1 (GSE248904) |
| **Lung** | 18 | TMS | Lung_18m | 4376 | figshare:8273102 (TMS) | cell_ontology_class from source (Tabula Muris cell ontology) | Lung_CTRL1 (GSE248904) |
| **Lung** | 21 | TMS | Lung_21m | 2950 | figshare:8273102 (TMS) | cell_ontology_class from source (Tabula Muris cell ontology) | Lung_CTRL1 (GSE248904) |
| **Lung** | 23 | PanSci | Lung_23_months | 9997 | GSE247719 | Curated cellType + Lineage + Sub_cell_type annotations from source | Lung_CTRL1 (GSE248904) |
| **Lung** | 24 | TMS | Lung_24m | 1870 | figshare:8273102 (TMS) | cell_ontology_class from source (Tabula Muris cell ontology) | Lung_CTRL1 (GSE248904) |
| **Lung** | 30 | TMS | Lung_30m | 9994 | figshare:8273102 (TMS) | cell_ontology_class from source (Tabula Muris cell ontology) | Lung_CTRL1 (GSE248904) |
| **Muscle** | 3 | PanSci | Muscle_03_months | 9995 | GSE247719 | Curated cellType + Lineage + Sub_cell_type annotations from source | Muscle_CTRL1 (GSE248904) |
| **Muscle** | 3 | TMS | Muscle_3m | 1102 | figshare:8273102 (TMS) | cell_ontology_class from source (Tabula Muris cell ontology) | Muscle_CTRL1 (GSE248904) |
| **Muscle** | 6 | PanSci | Muscle_06_months | 9999 | GSE247719 | Curated cellType + Lineage + Sub_cell_type annotations from source | Muscle_CTRL1 (GSE248904) |
| **Muscle** | 12 | PanSci | Muscle_12_months | 9995 | GSE247719 | Curated cellType + Lineage + Sub_cell_type annotations from source | Muscle_CTRL1 (GSE248904) |
| **Muscle** | 16 | PanSci | Muscle_16_months | 9997 | GSE247719 | Curated cellType + Lineage + Sub_cell_type annotations from source | Muscle_CTRL1 (GSE248904) |

| | | | | | | | |
|---|---|---|---|---|---|---|---|
| **Muscle** | 18 | TMS | Muscle_18m | 1521 | figshare:8273102 (TMS) | cell_ontology_class from source (Tabula Muris cell ontology) | Muscle_CTRL1 (GSE248904) |
| **Muscle** | 21 | TMS | Muscle_21m | 3528 | figshare:8273102 (TMS) | cell_ontology_class from source (Tabula Muris cell ontology) | Muscle_CTRL1 (GSE248904) |
| **Muscle** | 23 | PanSci | Muscle_23_months | 9995 | GSE247719 | Curated cellType + Lineage + Sub_cell_type annotations from source | Muscle_CTRL1 (GSE248904) |
| **Muscle** | 24 | TMS | Muscle_24m | 1231 | figshare:8273102 (TMS) | cell_ontology_class from source (Tabula Muris cell ontology) | Muscle_CTRL1 (GSE248904) |
| **Muscle** | 30 | TMS | Muscle_30m | 3284 | figshare:8273102 (TMS) | cell_ontology_class from source (Tabula Muris cell ontology) | Muscle_CTRL1 (GSE248904) |
| **Pancreas** | 3 | TMS | Pancreas_3m | 1588 | figshare:8273102 (TMS) | cell_ontology_class from source (Tabula Muris cell ontology) | Pancreas_CTRL1 (GSE248904) |
| **Pancreas** | 18 | TMS | Pancreas_18m | 2071 | figshare:8273102 (TMS) | cell_ontology_class from source (Tabula Muris cell ontology) | Pancreas_CTRL1 (GSE248904) |
| **Pancreas** | 21 | TMS | Pancreas_21m | 1437 | figshare:8273102 (TMS) | cell_ontology_class from source (Tabula Muris cell ontology) | Pancreas_CTRL1 (GSE248904) |
| **Pancreas** | 30 | TMS | Pancreas_30m | 2642 | figshare:8273102 (TMS) | cell_ontology_class from source (Tabula Muris cell ontology) | Pancreas_CTRL1 (GSE248904) |
| **Skin** | 3 | TMS | Skin_3m | 2346 | figshare:8273102 (TMS) | cell_ontology_class from source (Tabula Muris cell ontology) | Skin_CTRL1 (GSE248904) |
| **Skin** | 21 | TMS | Skin_21m | 4352 | figshare:8273102 (TMS) | cell_ontology_class from source (Tabula Muris cell ontology) | Skin_CTRL1 (GSE248904) |
| **Skin** | 24 | TMS | Skin_24m | 1122 | figshare:8273102 (TMS) | cell_ontology_class from source (Tabula Muris cell ontology) | Skin_CTRL1 (GSE248904) |
| **Spleen** | 1 | TMS | Spleen_1m | 2801 | figshare:8273102 (TMS) | cell_ontology_class from source (Tabula | Spleen_CTRL1 (GSE248904) |

| | | | | | | | |
|---|---|---|---|---|---|---|---|
| | | | | | | Muris cell ontology) | |
| **Spleen** | 3 | TMS | Spleen_3m | 6480 | figshare:8273102 (TMS) | cell_ontology_class from source (Tabula Muris cell ontology) | Spleen_CTRL1 (GSE248904) |
| **Spleen** | 6 | MCA2 | Spleen_6m | 9412 | GSE153562 | leiden clusters with marker-based manual annotation | Spleen_CTRL1 (GSE248904) |
| **Spleen** | 12 | MCA2 | Spleen_12m | 9569 | GSE153562 | leiden clusters with marker-based manual annotation | Spleen_CTRL1 (GSE248904) |
| **Spleen** | 18 | TMS | Spleen_18m | 5929 | figshare:8273102 (TMS) | cell_ontology_class from source (Tabula Muris cell ontology) | Spleen_CTRL1 (GSE248904) |
| **Spleen** | 21 | TMS | Spleen_21m | 6186 | figshare:8273102 (TMS) | cell_ontology_class from source (Tabula Muris cell ontology) | Spleen_CTRL1 (GSE248904) |
| **Spleen** | 24 | TMS | Spleen_24m | 3954 | figshare:8273102 (TMS) | cell_ontology_class from source (Tabula Muris cell ontology) | Spleen_CTRL1 (GSE248904) |
| **Spleen** | 30 | TMS | Spleen_30m | 8615 | figshare:8273102 (TMS) | cell_ontology_class from source (Tabula Muris cell ontology) | Spleen_CTRL1 (GSE248904) |
| **Stomach** | 3 | PanSci | Stomach_03_months | 9440 | GSE247719 | Curated cellType + Lineage + Sub_cell_type annotations from source | Stomach_CTRL1 (GSE248904) |
| **Stomach** | 6 | PanSci | Stomach_06_months | 9361 | GSE247719 | Curated cellType + Lineage + Sub_cell_type annotations from source | Stomach_CTRL1 (GSE248904) |
| **Stomach** | 12 | MCA2 | Stomach_12m | 8218 | GSE153562 | leiden clusters with marker-based manual annotation | Stomach_CTRL1 (GSE248904) |
| **Stomach** | 12 | PanSci | Stomach_12_months | 9457 | GSE247719 | Curated cellType + Lineage + Sub_cell_type annotations from source | Stomach_CTRL1 (GSE248904) |
| **Stomach** | 16 | PanSci | Stomach_16_months | 9353 | GSE247719 | Curated cellType + Lineage + Sub_cell_type | Stomach_CTRL1 (GSE248904) |

| | | | | | | | |
|---|---|---|---|---|---|---|---|
| | | | | | | annotations from source | |
| **Stomach** | 18 | MCA2 | Stomach_18m | 9552 | GSE153562 | leiden clusters with marker-based manual annotation | Stomach_CTRL1 (GSE248904) |
| **Stomach** | 23 | PanSci | Stomach_23_months | 9353 | GSE247719 | Curated cellType + Lineage + Sub_cell_type annotations from source | Stomach_CTRL1 (GSE248904) |
| **Stomach** | 24 | MCA2 | Stomach_24m | 9381 | GSE153562 | leiden clusters with marker-based manual annotation | Stomach_CTRL1 (GSE248904) |
| **Thymus** | 3 | TMS | Thymus_3m | 1430 | figshare:8273102 (TMS) | cell_ontology_class from source (Tabula Muris cell ontology) | Thymus_CTRL1 (GSE248904) |
| **Thymus** | 12 | MCA2 | Thymus_12m | 10000 | GSE153562 | leiden clusters with marker-based manual annotation | Thymus_CTRL1 (GSE248904) |
| **Thymus** | 18 | TMS | Thymus_18m | 2040 | figshare:8273102 (TMS) | cell_ontology_class from source (Tabula Muris cell ontology) | Thymus_CTRL1 (GSE248904) |
| **Thymus** | 21 | TMS | Thymus_21m | 2792 | figshare:8273102 (TMS) | cell_ontology_class from source (Tabula Muris cell ontology) | Thymus_CTRL1 (GSE248904) |
| **Thymus** | 24 | TMS | Thymus_24m | 3013 | figshare:8273102 (TMS) | cell_ontology_class from source (Tabula Muris cell ontology) | Thymus_CTRL1 (GSE248904) |

#### Notes

- **CMap projection.** Each single-cell sample is mapped onto the whole-body spatial reference (GSE248904 CTRL replicate 1) using CMap classification + spot assignment. See scripts/preprocess_new_sc.py and scripts/process_remaining_gaps.py. A spot-classification confidence (CMAP_quality) is retained per cell.
- **gsMap input.** Each (organ × age × source) projected h5ad is one input to `gsmap quick_mode`; outputs include per-spot AD -log10(p) and per-annotation Cauchy combination.
- **Multi-source Brain samples.** 5 Brain samples (Brain_3m, Brain_6m, Brain_12m, Brain_18m, Brain_24m) each combine ≥2 sources to maximise cell-type coverage: PanSci/EasySci supply non-microglia neurons; Hammond and TMS add microglia depth; MCA2 fills aged gaps. See Fig. 4c pie-matrix in notebooks/paper_figure.ipynb.
- **Why 15 organs and not 16.** Lymph_Node spatial spots exist in GSE248904 but no aged single-cell dataset covers lymph node directly; lymph node was excluded from the multi-age atlas (still appears in the 6-week whole-body baseline).

- **Why 1–30 mo and not 1–32 mo.** TMS droplet covers up to 30 m; no public single-cell mouse atlas reliably covers 31–32 months for the organs analysed here.
- **Provenance script.** scripts/build_age_provenance_per_source.py rebuilds results/age_sample_provenance_per_source.csv from raw obs metadata + CMap output paths.

##### Summary statistics

- Total single-cell sources: 5
- Total samples (organ × age × source rows): 131
- Distinct (organ × age) combinations: 126
- Distinct organs: 15
- Distinct ages: 10 (1, 3, 6, 12, 16, 18, 21, 23, 24, 30 months)
- Total cells: 809,284
- Largest source: PanSci (454,620 cells, 56.2%)
- Smallest source: Hammond_microglia (9,998 cells)
- Multi-source samples: 5 (all Brain)

#### *Supplementary Table 34*

Per-organ, per-age sample counts, spot counts and median spatial AD genetic-risk scores across the multi-age CMap-projected atlas. Young (≤6 mo) vs Old (≥18 mo) totals match the paragraph reference numbers — 264,283 young spots / 41 samples vs 322,477 old spots / 62 samples — used to construct the Young/Old whole-body risk maps in Fig. 4e.

##### 1 | Overall age-group totals

| age_group | n_samples | n_spots_total | n_spots_pAD_valid | median_logp | q25_logp | q75_logp | frac_genomewide_sig |
|---|---|---|---|---|---|---|---|
| **Young (≤6 mo)** | 41 | 264283 | 261598 | 1.186 | 0.391 | 2.432 | 0.006 |
| **Mid (6–18 mo)** | 23 | 222524 | 221202 | 1.197 | 0.532 | 2.06 | 0.001 |
| **Old (≥18 mo)** | 62 | 322477 | 319643 | 2.115 | 1.012 | 3.439 | 0.008 |

##### 2 | Per-organ × age-group breakdown

| organ | age_group | n_samples | n_spots_total | n_spots_pAD_valid | median_logp | q25_logp | q75_logp | frac_genomewide_sig |
|---|---|---|---|---|---|---|---|---|
| **BAT** | Young (≤6 mo) | 3 | 20713 | 20678 | 1.097 | 0.653 | 1.613 | 0.0 |
| **BAT** | Mid (6–18 mo) | 2 | 20000 | 19977 | 1.056 | 0.614 | 1.597 | 0.0 |
| **BAT** | Old (≥18 mo) | 5 | 14319 | 14223 | 1.278 | 0.706 | 2.212 | 0.01 |
| **Bone_Marrow** | Young (≤6 mo) | 2 | 6517 | 6428 | 3.503 | 2.327 | 4.873 | 0.028 |
| **Bone_Marrow** | Old (≥18 mo) | 4 | 30207 | 30057 | 3.187 | 2.238 | 4.373 | 0.009 |
| **Brain** | Young (≤6 mo) | 4 | 33821 | 33285 | 0.035 | 0.001 | 0.752 | 0.026 |
| **Brain** | Mid (6–18 mo) | 1 | 6439 | 6417 | 0.666 | 0.221 | 5.351 | 0.014 |
| **Brain** | Old (≥18 mo) | 4 | 32669 | 32275 | 1.049 | 0.016 | 4.632 | 0.021 |

| | | | | | | | | |
|---|---|---|---|---|---|---|---|---|
| **Colon** | Young (≤6 mo) | 3 | 23987 | 23826 | 1.322 | 0.518 | 2.338 | 0.0 |
| **Colon** | Mid (6–18 mo) | 2 | 20000 | 19827 | 1.18 | 0.504 | 2.018 | 0.0 |
| **Colon** | Old (≥18 mo) | 5 | 26211 | 26124 | 1.584 | 0.783 | 2.67 | 0.001 |
| **Heart** | Young (≤6 mo) | 4 | 21480 | 21395 | 1.018 | 0.182 | 2.208 | 0.001 |
| **Heart** | Mid (6–18 mo) | 2 | 20000 | 19954 | 1.057 | 0.178 | 2.12 | 0.0 |
| **Heart** | Old (≥18 mo) | 5 | 17133 | 16955 | 1.58 | 0.464 | 2.724 | 0.005 |
| **Ileum** | Young (≤6 mo) | 2 | 19988 | 19839 | 1.742 | 0.749 | 2.61 | 0.0 |
| **Ileum** | Mid (6–18 mo) | 3 | 29972 | 29807 | 1.771 | 1.049 | 2.587 | 0.0 |
| **Ileum** | Old (≥18 mo) | 4 | 39952 | 39831 | 2.01 | 1.425 | 2.645 | 0.0 |
| **Kidney** | Young (≤6 mo) | 4 | 24251 | 23786 | 0.867 | 0.434 | 1.527 | 0.0 |
| **Kidney** | Mid (6–18 mo) | 2 | 19595 | 19260 | 0.731 | 0.341 | 1.234 | 0.0 |
| **Kidney** | Old (≥18 mo) | 5 | 26058 | 25776 | 2.045 | 0.836 | 3.401 | 0.006 |
| **Liver** | Young (≤6 mo) | 4 | 23468 | 23338 | 1.596 | 0.969 | 2.319 | 0.001 |
| **Liver** | Mid (6–18 mo) | 2 | 19938 | 19913 | 1.37 | 0.861 | 1.953 | 0.0 |
| **Liver** | Old (≥18 mo) | 5 | 14743 | 14494 | 1.625 | 0.824 | 3.156 | 0.026 |
| **Lung** | Young (≤6 mo) | 4 | 26104 | 25537 | 1.41 | 0.646 | 3.123 | 0.007 |
| **Lung** | Mid (6–18 mo) | 2 | 19991 | 19850 | 1.12 | 0.497 | 2.998 | 0.001 |
| **Lung** | Old (≥18 mo) | 5 | 29187 | 28500 | 3.176 | 1.773 | 4.608 | 0.017 |
| **Muscle** | Young (≤6 mo) | 3 | 21096 | 20929 | 0.733 | 0.302 | 1.309 | 0.0 |
| **Muscle** | Mid (6–18 mo) | 2 | 19992 | 19807 | 0.651 | 0.255 | 1.169 | 0.0 |
| **Muscle** | Old (≥18 mo) | 5 | 19559 | 19456 | 1.012 | 0.446 | 2.148 | 0.004 |
| **Pancreas** | Young (≤6 mo) | 1 | 1588 | 1588 | 1.597 | 1.017 | 2.28 | 0.003 |
| **Pancreas** | Old (≥18 mo) | 3 | 6150 | 6015 | 1.287 | 0.795 | 2.104 | 0.005 |
| **Skin** | Young (≤6 mo) | 1 | 2346 | 2308 | 2.255 | 1.578 | 2.946 | 0.0 |
| **Skin** | Old (≥18 mo) | 2 | 5474 | 5454 | 2.346 | 1.522 | 3.22 | 0.0 |
| **Spleen** | Young (≤6 mo) | 3 | 18693 | 18530 | 3.703 | 2.859 | 4.531 | 0.01 |
| **Spleen** | Mid (6–18 mo) | 1 | 9569 | 9515 | 2.113 | 1.636 | 2.676 | 0.0 |
| **Spleen** | Old (≥18 mo) | 4 | 24684 | 24490 | 3.723 | 2.942 | 4.655 | 0.012 |
| **Stomach** | Young (≤6 mo) | 2 | 18801 | 18701 | 0.699 | 0.234 | 1.278 | 0.0 |
| **Stomach** | Mid (6–18 mo) | 3 | 27028 | 26875 | 0.934 | 0.394 | 1.629 | 0.0 |
| **Stomach** | Old (≥18 mo) | 3 | 28286 | 28181 | 1.514 | 0.896 | 2.16 | 0.0 |
| **Thymus** | Young (≤6 mo) | 1 | 1430 | 1430 | 2.468 | 1.938 | 3.143 | 0.001 |
| **Thymus** | Mid (6–18 mo) | 1 | 10000 | 10000 | 2.19 | 1.673 | 2.791 | 0.0 |
| **Thymus** | Old (≥18 mo) | 3 | 7845 | 7812 | 3.004 | 2.324 | 3.767 | 0.004 |

#### 3 | Per-organ × age-months (fine-grained)

| organ | age_months | n_samples | n_spots_total | n_spots_pAD_valid | median_logp | q25_logp | q75_logp | frac_genomewide_sig |
|---|---|---|---|---|---|---|---|---|
| **BAT** | 3 | 2 | 10713 | 10683 | 1.186 | 0.734 | 1.707 | 0.001 |

| | | | | | | | | |
|---|---|---|---|---|---|---|---|---|
| **BAT** | 6 | 1 | 10000 | 9995 | 0.996 | 0.58 | 1.496 | 0.0 |
| **BAT** | 12 | 1 | 10000 | 9994 | 1.057 | 0.606 | 1.6 | 0.0 |
| **BAT** | 16 | 1 | 10000 | 9983 | 1.054 | 0.623 | 1.595 | 0.0 |
| **BAT** | 18 | 1 | 662 | 613 | 3.549 | 1.547 | 5.833 | 0.055 |
| **BAT** | 21 | 1 | 794 | 781 | 2.458 | 1.045 | 3.528 | 0.061 |
| **BAT** | 23 | 1 | 10000 | 9998 | 1.123 | 0.642 | 1.734 | 0.0 |
| **BAT** | 24 | 1 | 848 | 844 | 1.604 | 0.916 | 2.513 | 0.001 |
| **BAT** | 30 | 1 | 2015 | 1987 | 2.421 | 0.793 | 3.406 | 0.031 |
| **Bone_Marrow** | 1 | 1 | 3027 | 2988 | 3.328 | 2.239 | 4.574 | 0.011 |
| **Bone_Marrow** | 3 | 1 | 3490 | 3440 | 3.694 | 2.429 | 5.145 | 0.042 |
| **Bone_Marrow** | 18 | 1 | 6712 | 6656 | 3.357 | 2.405 | 4.502 | 0.005 |
| **Bone_Marrow** | 21 | 1 | 5216 | 5181 | 3.098 | 2.223 | 4.388 | 0.011 |
| **Bone_Marrow** | 24 | 1 | 8279 | 8247 | 3.23 | 2.18 | 4.276 | 0.012 |
| **Bone_Marrow** | 30 | 1 | 10000 | 9973 | 3.088 | 2.17 | 4.364 | 0.009 |
| **Brain** | 3 | 2 | 17923 | 17597 | 0.062 | 0.001 | 1.398 | 0.04 |
| **Brain** | 6 | 2 | 15898 | 15688 | 0.023 | 0.002 | 0.318 | 0.01 |
| **Brain** | 12 | 1 | 6439 | 6417 | 0.666 | 0.221 | 5.351 | 0.014 |
| **Brain** | 18 | 1 | 6636 | 6595 | 3.687 | 2.114 | 4.824 | 0.009 |
| **Brain** | 21 | 2 | 20697 | 20440 | 0.056 | 0.003 | 3.503 | 0.029 |
| **Brain** | 24 | 1 | 5336 | 5240 | 3.514 | 2.456 | 4.639 | 0.004 |
| **Colon** | 3 | 2 | 13987 | 13937 | 1.398 | 0.571 | 2.418 | 0.0 |
| **Colon** | 6 | 1 | 10000 | 9889 | 1.239 | 0.456 | 2.23 | 0.0 |
| **Colon** | 12 | 1 | 10000 | 9900 | 1.186 | 0.497 | 2.021 | 0.0 |
| **Colon** | 16 | 1 | 10000 | 9927 | 1.174 | 0.512 | 2.01 | 0.0 |
| **Colon** | 18 | 1 | 2369 | 2369 | 2.378 | 1.136 | 3.297 | 0.0 |
| **Colon** | 21 | 1 | 10000 | 10000 | 1.571 | 0.863 | 2.71 | 0.001 |
| **Colon** | 23 | 1 | 10000 | 9913 | 1.159 | 0.501 | 1.921 | 0.0 |
| **Colon** | 24 | 1 | 1955 | 1955 | 2.619 | 1.651 | 3.592 | 0.001 |
| **Colon** | 30 | 1 | 1887 | 1887 | 2.864 | 1.884 | 3.898 | 0.003 |
| **Heart** | 1 | 1 | 946 | 923 | 2.483 | 1.309 | 3.476 | 0.001 |
| **Heart** | 3 | 2 | 10534 | 10490 | 1.038 | 0.213 | 2.292 | 0.002 |
| **Heart** | 6 | 1 | 10000 | 9982 | 0.88 | 0.13 | 1.974 | 0.0 |
| **Heart** | 12 | 1 | 10000 | 9993 | 0.942 | 0.176 | 2.07 | 0.0 |
| **Heart** | 16 | 1 | 10000 | 9961 | 1.156 | 0.181 | 2.16 | 0.001 |
| **Heart** | 18 | 1 | 1143 | 1099 | 1.942 | 0.435 | 3.519 | 0.013 |
| **Heart** | 21 | 1 | 918 | 853 | 2.273 | 1.074 | 3.375 | 0.038 |
| **Heart** | 23 | 1 | 10000 | 9994 | 1.15 | 0.155 | 2.073 | 0.002 |
| **Heart** | 24 | 1 | 768 | 728 | 2.41 | 0.886 | 3.643 | 0.014 |
| **Heart** | 30 | 1 | 4304 | 4281 | 2.412 | 1.42 | 3.299 | 0.003 |
| **Ileum** | 3 | 1 | 9994 | 9940 | 1.816 | 0.673 | 2.704 | 0.0 |
| **Ileum** | 6 | 1 | 9994 | 9899 | 1.69 | 0.805 | 2.515 | 0.0 |
| **Ileum** | 12 | 2 | 19977 | 19897 | 1.778 | 1.141 | 2.525 | 0.0 |
| **Ileum** | 16 | 1 | 9995 | 9910 | 1.752 | 0.793 | 2.733 | 0.0 |
| **Ileum** | 18 | 1 | 9983 | 9959 | 1.947 | 1.531 | 2.422 | 0.0 |
| **Ileum** | 21 | 1 | 9985 | 9985 | 2.345 | 1.801 | 2.93 | 0.0 |
| **Ileum** | 23 | 1 | 9998 | 9901 | 1.889 | 1.009 | 2.741 | 0.0 |
| **Ileum** | 24 | 1 | 9986 | 9986 | 1.809 | 1.281 | 2.446 | 0.001 |
| **Kidney** | 1 | 1 | 2422 | 2375 | 2.27 | 1.362 | 3.273 | 0.0 |
| **Kidney** | 3 | 2 | 12056 | 11829 | 0.851 | 0.443 | 1.466 | 0.0 |
| **Kidney** | 6 | 1 | 9773 | 9582 | 0.732 | 0.33 | 1.231 | 0.0 |
| **Kidney** | 12 | 1 | 9795 | 9611 | 0.708 | 0.334 | 1.204 | 0.0 |
| **Kidney** | 16 | 1 | 9800 | 9649 | 0.752 | 0.349 | 1.264 | 0.0 |

| | | | | | | | | |
|---|---|---|---|---|---|---|---|---|
| **Kidney** | 18 | 1 | 3040 | 3003 | 2.587 | 1.726 | 3.407 | 0.004 |
| **Kidney** | 21 | 1 | 2204 | 2141 | 2.833 | 1.763 | 3.752 | 0.003 |
| **Kidney** | 23 | 1 | 9795 | 9638 | 0.706 | 0.349 | 1.176 | 0.0 |
| **Kidney** | 24 | 1 | 5508 | 5508 | 3.659 | 2.97 | 4.509 | 0.017 |
| **Kidney** | 30 | 1 | 5511 | 5486 | 2.508 | 1.67 | 3.541 | 0.01 |
| **Liver** | 1 | 1 | 2785 | 2725 | 3.406 | 2.331 | 4.465 | 0.007 |
| **Liver** | 3 | 2 | 10710 | 10669 | 1.389 | 0.809 | 2.042 | 0.0 |
| **Liver** | 6 | 1 | 9973 | 9944 | 1.564 | 0.988 | 2.145 | 0.0 |
| **Liver** | 12 | 1 | 9969 | 9958 | 1.436 | 0.936 | 2.008 | 0.0 |
| **Liver** | 16 | 1 | 9969 | 9955 | 1.299 | 0.794 | 1.903 | 0.0 |
| **Liver** | 18 | 1 | 1176 | 1094 | 3.191 | 1.388 | 4.443 | 0.067 |
| **Liver** | 21 | 1 | 367 | 321 | 3.445 | 2.862 | 4.344 | 0.0 |
| **Liver** | 23 | 1 | 9983 | 9961 | 1.265 | 0.662 | 2.141 | 0.0 |
| **Liver** | 24 | 1 | 945 | 868 | 3.089 | 1.422 | 4.752 | 0.096 |
| **Liver** | 30 | 1 | 2272 | 2250 | 5.225 | 2.27 | 6.373 | 0.096 |
| **Lung** | 1 | 1 | 2507 | 2299 | 3.76 | 2.287 | 5.013 | 0.011 |
| **Lung** | 3 | 2 | 13606 | 13293 | 1.317 | 0.614 | 2.979 | 0.011 |
| **Lung** | 6 | 1 | 9991 | 9945 | 1.161 | 0.578 | 2.663 | 0.001 |
| **Lung** | 12 | 1 | 9995 | 9935 | 1.165 | 0.533 | 2.997 | 0.001 |
| **Lung** | 16 | 1 | 9996 | 9915 | 1.066 | 0.468 | 2.999 | 0.001 |
| **Lung** | 18 | 1 | 4376 | 4242 | 3.132 | 2.094 | 5.07 | 0.024 |
| **Lung** | 21 | 1 | 2950 | 2751 | 3.092 | 2.268 | 4.309 | 0.008 |
| **Lung** | 23 | 1 | 9997 | 9928 | 1.77 | 0.736 | 3.335 | 0.001 |
| **Lung** | 24 | 1 | 1870 | 1743 | 1.74 | 0.886 | 3.829 | 0.018 |
| **Lung** | 30 | 1 | 9994 | 9836 | 4.227 | 3.279 | 5.304 | 0.032 |
| **Muscle** | 3 | 2 | 11097 | 11046 | 0.881 | 0.353 | 1.616 | 0.001 |
| **Muscle** | 6 | 1 | 9999 | 9883 | 0.631 | 0.267 | 1.032 | 0.0 |
| **Muscle** | 12 | 1 | 9995 | 9862 | 0.65 | 0.252 | 1.143 | 0.0 |
| **Muscle** | 16 | 1 | 9997 | 9945 | 0.652 | 0.259 | 1.2 | 0.0 |
| **Muscle** | 18 | 1 | 1521 | 1521 | 2.318 | 1.477 | 3.369 | 0.018 |
| **Muscle** | 21 | 1 | 3528 | 3477 | 1.675 | 1.049 | 2.878 | 0.005 |
| **Muscle** | 23 | 1 | 9995 | 9943 | 0.541 | 0.239 | 1.01 | 0.0 |
| **Muscle** | 24 | 1 | 1231 | 1231 | 1.604 | 1.079 | 2.347 | 0.016 |
| **Muscle** | 30 | 1 | 3284 | 3284 | 2.226 | 0.872 | 3.544 | 0.004 |
| **Pancreas** | 3 | 1 | 1588 | 1588 | 1.597 | 1.017 | 2.28 | 0.003 |
| **Pancreas** | 18 | 1 | 2071 | 2046 | 1.365 | 0.872 | 2.065 | 0.014 |
| **Pancreas** | 21 | 1 | 1437 | 1368 | 1.803 | 1.22 | 2.514 | 0.0 |
| **Pancreas** | 30 | 1 | 2642 | 2601 | 0.994 | 0.622 | 1.764 | 0.0 |
| **Skin** | 3 | 1 | 2346 | 2308 | 2.255 | 1.578 | 2.946 | 0.0 |
| **Skin** | 21 | 1 | 4352 | 4352 | 2.534 | 1.676 | 3.387 | 0.0 |
| **Skin** | 24 | 1 | 1122 | 1102 | 1.751 | 1.17 | 2.379 | 0.0 |
| **Spleen** | 1 | 1 | 2801 | 2734 | 3.813 | 3.017 | 4.601 | 0.007 |
| **Spleen** | 3 | 1 | 6480 | 6407 | 3.971 | 3.21 | 4.711 | 0.008 |
| **Spleen** | 6 | 1 | 9412 | 9389 | 3.443 | 2.614 | 4.349 | 0.011 |
| **Spleen** | 12 | 1 | 9569 | 9515 | 2.113 | 1.636 | 2.676 | 0.0 |
| **Spleen** | 18 | 1 | 5929 | 5855 | 3.846 | 3.075 | 4.623 | 0.004 |
| **Spleen** | 21 | 1 | 6186 | 6123 | 4.103 | 3.321 | 4.912 | 0.012 |
| **Spleen** | 24 | 1 | 3954 | 3919 | 3.424 | 2.665 | 4.392 | 0.018 |
| **Spleen** | 30 | 1 | 8615 | 8593 | 3.492 | 2.784 | 4.458 | 0.015 |
| **Stomach** | 3 | 1 | 9440 | 9387 | 0.758 | 0.27 | 1.351 | 0.0 |
| **Stomach** | 6 | 1 | 9361 | 9314 | 0.635 | 0.207 | 1.212 | 0.0 |
| **Stomach** | 12 | 2 | 17675 | 17567 | 1.092 | 0.48 | 1.81 | 0.0 |
| **Stomach** | 16 | 1 | 9353 | 9308 | 0.718 | 0.275 | 1.243 | 0.0 |
| **Stomach** | 18 | 1 | 9552 | 9552 | 1.892 | 1.424 | 2.457 | 0.0 |
| **Stomach** | 23 | 1 | 9353 | 9266 | 0.67 | 0.242 | 1.362 | 0.0 |
| **Stomach** | 24 | 1 | 9381 | 9363 | 1.643 | 1.231 | 2.221 | 0.0 |
| **Thymus** | 3 | 1 | 1430 | 1430 | 2.468 | 1.938 | 3.143 | 0.001 |
| **Thymus** | 12 | 1 | 10000 | 10000 | 2.19 | 1.673 | 2.791 | 0.0 |

| | | | | | | | | |
|---|---|---|---|---|---|---|---|---|
| **Thymus** | 18 | 1 | 2040 | 2040 | 2.699 | 2.047 | 3.267 | 0.005 |
| **Thymus** | 21 | 1 | 2792 | 2792 | 2.708 | 2.048 | 3.361 | 0.001 |
| **Thymus** | 24 | 1 | 3013 | 2980 | 3.623 | 2.909 | 4.443 | 0.007 |

#### Notes

- **gsMap input.** Each (organ × age × source) CMap-projected h5ad in the multi-age atlas (Supplementary Table Mouse 1) was scored with `gsmap quick_mode` against the Bellenguez 2022 AD GWAS, producing a per-spot S-LDSC $\chi^2$ statistic and p-value.
- **Median log10(p) metric.** Per-organ × per-age 'median_logp' is the spot-level median of -log10(p_AD). Higher = stronger per-spot AD heritability enrichment in that organ-age stratum. q25 and q75 give the 25th/75th percentiles to show the spread.
- **Genome-wide significance fraction.** frac_genomewide_sig = fraction of spots with p_AD < $5\times10^{-8}$. Conservative; complements the median by capturing the tail.
- **Why n_spots_total ≠ n_spots_pAD_valid.** A small number of spots (~1%) fail gsMap S-LDSC (no LDSC score in the chunk they fell into); we list both for transparency. Median and percentiles use only valid spots.
- **Age-group definitions.** Young = age_months ≤ 6 (1, 3, 6 mo); Mid = 6 < age_months < 18 (12, 16 mo); Old = age_months ≥ 18 (18, 21, 23, 24, 30 mo). Young and Old definitions match the Young/Old comparison in Fig. 4e and the paragraph S2 reference numbers (264,283 young spots / 41 samples; 322,477 old spots / 62 samples).
- **Sample-sample variability.** n_samples in each (organ × age) cell is the number of CMap-projected samples (= h5ad files) feeding gsMap. The same organ-age can have >1 sample from different single-cell sources (see Supplementary Table Mouse 1, Section 3).

#### Summary statistics

- Total samples: 126
- Total spots: 809,284
- Spots with valid p_AD: 802,443 (99.15%)
- Distinct organs: 15
- Distinct ages: 10 months ([1, 3, 6, 12, 16, 18, 21, 23, 24, 30])
- Young (≤6 mo): 264,283 spots, 41 samples
- Old (≥18 mo): 322,477 spots, 62 samples
- Old vs Young median log10(p) ratio (overall): 2.12 / 1.19 = 1.78×

### *Supplementary Table 35*

Full GTEx eQTL query output for the colon-nominated Alzheimer's disease candidate genes (Para 5 S2). Per (gene × tissue): number of significant single-tissue cis-eQTLs, best eQTL P value, lead SNP (from SMR), and whether the locus's AD GWAS lead SNP is itself a significant eQTL.

#### 1 | Per-gene aggregate (colon vs brain vs blood vs liver)

| gene | pathway | colon_n_eQTL | brain_n_eQTL | blood_n_eQTL | liver_n_eQTL | colon_best_p | ad_snp_is_colon_eQTL |
|---|---|---|---|---|---|---|---|
| **ADAM10** | APP | 345 | 14 | 27 | 0 | 8.72697e-10 | False |
| **CD2AP** | APP | 200 | 69 | 98 | 0 | 1.55928e-06 | True |
| **BIN1** | Myeloid | 192 | 73 | 250 | 29 | 3.19881e-22 | False |

| | | | | | | | |
|---|---|---|---|---|---|---|---|
| **PICALM** | APP | 93 | 0 | 59 | 0 | 2.4864e-09 | False |
| **APP** | APP | 28 | 3 | 249 | 0 | 5.99202e-07 | False |
| **CR1** | APP | 24 | 38 | 4 | 0 | 4.60457e-07 | True |
| **PSEN1** | APP | 0 | 1 | 250 | 210 | | False |
| **SORL1** | APP | 0 | 0 | 0 | 0 | | False |
| **SPI1** | Myeloid | 0 | 0 | 0 | 0 | | False |
| **TREM2** | Myeloid | 0 | 0 | 0 | 0 | | False |

#### 2 | Full per-(gene × tissue) GTEx query

| gene | pathway | tissue | n_sig_eQTL | n_tested_total | best_eQTL_p | lead_snp | smr_p_eqtl_top | ad_snp_is_eQTL | ad_snp_eQTL_p |
|---|---|---|---|---|---|---|---|---|---|
| **ADAM10** | APP | Brain_Cerebellum | 14 | | 6.74145e-08 | | | False | |
| **ADAM10** | APP | Brain_Cortex | 0 | | | | | False | |
| **ADAM10** | APP | Colon_Sigmoid | 142 | 4295 | 1.54819e-08 | rs36017602 | 1.54819e-08 | False | |
| **ADAM10** | APP | Colon_Transverse | 203 | 4295 | 8.72697e-10 | rs28455654 | 8.72697e-10 | False | |
| **ADAM10** | APP | Liver | 0 | | | | | False | |
| **ADAM10** | APP | Whole_Blood | 27 | | 3.12644e-21 | | | False | |
| **APP** | APP | Brain_Cerebellum | 3 | | 7.25424e-14 | | | False | |
| **APP** | APP | Brain_Cortex | 0 | | | | | False | |
| **APP** | APP | Colon_Sigmoid | 10 | 4592 | 2.58802e-05 | rs1625289 | 2.58802e-05 | False | |
| **APP** | APP | Colon_Transverse | 18 | 4597 | 5.99202e-07 | rs9976487 | 5.99202e-07 | False | |
| **APP** | APP | Liver | 0 | | | | | False | |
| **APP** | APP | Whole_Blood | 249 | | 7.46545e-16 | | | False | |
| **BIN1** | Myeloid | Brain_Cerebellum | 73 | | 3.94785e-21 | | | False | |
| **BIN1** | Myeloid | Brain_Cortex | 0 | | | | | False | |
| **BIN1** | Myeloid | Colon_Sigmoid | 126 | | 3.19881e-22 | | | False | |
| **BIN1** | Myeloid | Colon_Transverse | 66 | | 2.56874e-12 | | | False | |
| **BIN1** | Myeloid | Liver | 29 | | 4.0699e-07 | | | False | |
| **BIN1** | Myeloid | Whole_Blood | 250 | | 3.15642e-35 | | | False | |
| **CD2AP** | APP | Brain_Cerebellum | 66 | | 1.71378e-08 | | | False | |
| **CD2AP** | APP | Brain_Cortex | 3 | | 3.17547e-05 | | | False | |
| **CD2AP** | APP | Colon_Sigmoid | 25 | | 3.18698e-05 | | | False | |
| **CD2AP** | APP | Colon_Transverse | 175 | | 1.55928e-06 | | | True | 3.53302e-06 |
| **CD2AP** | APP | Liver | 0 | | | | | False | |
| **CD2AP** | APP | Whole_Blood | 98 | | 1.48784e-08 | | | False | |
| **CR1** | APP | Brain_Cerebellum | 0 | | | | | False | |
| **CR1** | APP | Brain_Cortex | 38 | | 7.57438e-10 | | | True | 2.20042e-09 |

| | | | | | | | | | |
|---|---|---|---|---|---|---|---|---|---|
| **CR1** | APP | Colon_Sigmoid | 24 | 6111 | 4.60457e-07 | rs679515 | 4.60457e-07 | True | 9.18396e-07 |
| **CR1** | APP | Colon_Transverse | 0 | 6111 | | rs72741284 | 0.00072104 | False | |
| **CR1** | APP | Liver | 0 | | | | | False | |
| **CR1** | APP | Whole_Blood | 4 | | 8.29772e-06 | | | False | |
| **PICALM** | APP | Brain_Cerebellum | 0 | | | | | False | |
| **PICALM** | APP | Brain_Cortex | 0 | | | | | False | |
| **PICALM** | APP | Colon_Sigmoid | 48 | 3672 | 2.4864e-09 | rs7131120 | 2.4864e-09 | False | |
| **PICALM** | APP | Colon_Transverse | 45 | 3672 | 5.37765e-08 | rs597672 | 5.37765e-08 | False | |
| **PICALM** | APP | Liver | 0 | | | | | False | |
| **PICALM** | APP | Whole_Blood | 59 | | 1.38196e-06 | | | False | |
| **PSEN1** | APP | Brain_Cerebellum | 1 | | 2.21868e-05 | | | False | |
| **PSEN1** | APP | Brain_Cortex | 0 | | | | | False | |
| **PSEN1** | APP | Colon_Sigmoid | 0 | 1771 | | rs3742825 | 7.42804e-05 | False | |
| **PSEN1** | APP | Colon_Transverse | 0 | 1771 | | rs11625005 | 0.00175464 | False | |
| **PSEN1** | APP | Liver | 210 | | 6.30566e-08 | | | False | |
| **PSEN1** | APP | Whole_Blood | 250 | | 1.60036e-27 | | | False | |
| **SORL1** | APP | Brain_Cerebellum | 0 | | | | | False | |
| **SORL1** | APP | Brain_Cortex | 0 | | | | | False | |
| **SORL1** | APP | Colon_Sigmoid | 0 | 1569 | | rs923893 | 0.000915174 | False | |
| **SORL1** | APP | Colon_Transverse | 0 | 1569 | | rs139568145 | 0.000174324 | False | |
| **SORL1** | APP | Liver | 0 | | | | | False | |
| **SORL1** | APP | Whole_Blood | 0 | | | | | False | |
| **SPI1** | Myeloid | Brain_Cerebellum | 0 | | | | | False | |
| **SPI1** | Myeloid | Brain_Cortex | 0 | | | | | False | |
| **SPI1** | Myeloid | Colon_Sigmoid | 0 | | | | | False | |
| **SPI1** | Myeloid | Colon_Transverse | 0 | | | | | False | |
| **SPI1** | Myeloid | Liver | 0 | | | | | False | |
| **SPI1** | Myeloid | Whole_Blood | 0 | | | | | False | |
| **TREM2** | Myeloid | Brain_Cerebellum | 0 | | | | | False | |
| **TREM2** | Myeloid | Brain_Cortex | 0 | | | | | False | |
| **TREM2** | Myeloid | Colon_Sigmoid | 0 | | | | | False | |
| **TREM2** | Myeloid | Colon_Transverse | 0 | | | | | False | |
| **TREM2** | Myeloid | Liver | 0 | | | | | False | |

| | | | | | | | | |
|---|---|---|---|---|---|---|---|---|
| **TREM2** | Myeloid | Whole_Blood | 0 | | | | | False |

#### Notes

- **Data source.** Variants queried via GTEx Portal v8 (https://gtexportal.org) per (gene, tissue), filtered to significant single-tissue cis-eQTLs at the GTEx-default FDR. Colon = Colon_Transverse + Colon_Sigmoid; Brain = Brain_Cortex + Brain_Cerebellum; Whole_Blood = blood bulk; Liver = liver bulk.
- **Paragraph S2 verification.** PICALM colon_n_eQTL = 93 (matches paragraph); brain_n_eQTL = 0 (matches 'no detected bulk-brain eQTL'). CD2AP colon_n_eQTL = 200 (matches), ad_snp_is_colon_eQTL = True. CR1 colon_n_eQTL = 24 (matches), ad_snp_is_colon_eQTL = True.
- **Lead SNP source.** lead_snp is the SMR top-eQTL variant from the colocalisation analysis (scripts/21_eqtl_coloc_analysis.ipynb pipeline). It is the variant with the strongest eQTL signal in that gene's window in that tissue. Available for the 6 SMR-tested genes (PICALM, ADAM10, CR1, APP, SORL1, PSEN1) × 2 colon tissues. Non-SMR rows leave lead_snp blank.
- **n_tested_total vs n_sig_eQTL.** n_tested_total = number of variants tested in the locus by SMR (union of GWAS + eQTL variants after MAF/LD filtering). n_sig_eQTL = number of variants reaching GTEx single-tissue eQTL significance for that (gene, tissue). The ratio gives an effective association rate.
- **ad_snp_is_eQTL definition.** True if the Bellenguez 2022 AD GWAS lead SNP at this locus is itself annotated as a significant eQTL for this gene in this tissue. ad_snp_eQTL_p is its eQTL p-value when annotated.
- **Genes without colon eQTL.** SORL1, SPI1, TREM2 show 0 colon_n_eQTL in this GTEx bulk query. For TREM2 and SPI1 this likely reflects cell-type dilution (myeloid-specific expression invisible in bulk colon). For SORL1 it is a true negative in colon. These are honest limitations of the GTEx bulk eQTL framework, not analysis errors.

#### Data sources

| data_source | citation | accession | url | version_snapshot | derived_path_in_repo | notes |
|---|---|---|---|---|---|---|
| **GTEx Portal v8 eQTL query** | GTEx Consortium Science 2020 | v8 | https://gtexportal.org/home/ | queried 2026-04 (snapshot) | results/gtex_eqtl_query_results.csv | Per (gene × tissue): n_eqtl (significant single-tissue cis-eQTLs at GTEx FDR), best_p, and ad_snp_is_eqtl flag for the Bellenguez 2022 AD GWAS lead SNP |
| **Bellenguez 2022 AD GWAS** | Bellenguez et al. Nat Genet 2022 | GCST90027158 | https://www.ebi.ac.uk/gwas/studies/GCST90027158 | 2022-04 (GRCh38) | data/gwas/Bellenguez2022_AD_withN.tsv.gz (gitignored) | Used to identify AD GWAS lead SNPs per locus and to test for eQTL overlap |
| **SMR colocalisation pipeline** | Zhu et al. Nat Genet 2016 (SMR/HEIDI) | - | https://yanglab.westlake.edu.cn/software/smr/ | v1.3.1 | results/smr_colon_AD_results.csv | Provides lead_snp (top-eQTL SNP per gene × tissue), n_tested variants, SMR effect size, HEIDI p. Lead-SNP |

| | | | | | | |
|---|---|---|---|---|---|---|
| | | | | | | column in this table is sourced here. |
| **Pipeline notebook** | This work | - | - | - | notebooks/21_eqtl_coloc_analysis.ipynb (also executed.ipynb) | Runs the GTEx query + SMR for the 10 AD candidate genes |
| **Coloc-ABF + SharePro** | Giambartolomei 2014; Zhang Genome Biol 2023 | - | https://github.com/zhwm/SharePro_coloc | SharePro v5.0.0 | results/coloc_FULL_eqtl_results.csv + results/sharepro/{GENE}_v3_result.sharepro.txt | Used in Fig. 4 main analysis; not directly in this table but cross-references the AD-SNP overlap calls |

### *Supplementary Table 36*

Final per-gene evidence matrix combining GTEx colon eQTL, AD-SNP × eQTL overlap, single-causal coloc-ABF (full summary statistics), multi-causal SharePro, SMR + HEIDI, with explicit missing-analysis flags (Para 5 S5–S6). Highlights the CD2AP follow-up gap called out in S6.

#### 1 | Final evidence matrix (8 candidate genes)

| gene | GTEx_colon_n_eQTL | GTEx_colon_best_p | AD_SNP_is_colon_eQTL | coloc_ABF_best_PP_H4 | coloc_ABF_best_PP_H3 | SharePro_share | SMR_best_p | HEIDI_p_at_best_SMR | passes_coloc_ABF | passes_SharePro | passes_SMR | passes_HEIDI | missing_analyses | verdict |
|---|---|---|---|---|---|---|---|---|---|---|---|---|---|---|
| **CR1** | 24 | 4.60e-07 | Yes | 0.9925 | 0.3416 | 0.9991 | 2.07e-06 | 0.00e+00 | ✓ | ✓ | ✓ | — | (complete) | ✓✓ shared-causal (coloc + SharePro) |
| **PICALM** | 93 | 2.49e-09 | No | 0.0001 | 0.9995 | 0.9977 | 1.34e-03 | 0.00e+00 | — | ✓ | ✓ | — | (complete) | ✓ shared-causal (multi-causal SharePro rescue) |
| **CD2AP** | 200 | 1.56e-06 | Yes | | | | | | — | — | — | — | SharePro, coloc-ABF, SMR/HEIDI | REQUIRES FOLLOW-UP: AD GWAS SNP overlaps a colon eQTL but multi-causal |

| | | | | | | | | | | | | | | |
|---|---|---|---|---|---|---|---|---|---|---|---|---|---|---|
| | | | | | | | | | | | | | | SharePro not yet run (SharePro + coloc-ABF + SMR/HEIDI missing) |
| **ADAM10** | 345 | 8.73e-10 | No | 0.0004 | 0.9996 | 0.0005 | 4.00e-04 | 0.00e+00 | — | — | ✓ | — | (complete) | colon eQTL present but no coloc / SharePro / SMR support |
| **APP** | 28 | 5.99e-07 | No | 0.0982 | 0.9621 | | 1.48e-02 | 1.47e-129 | — | — | — | — | SharePro | REQUIRES FOLLOW-UP: has colon eQTL but missing SharePro |
| **SORL1** | 0 | | No | 0.0514 | 0.1161 | | 4.22e-01 | 2.61e-43 | — | — | — | — | SharePro | no colon eQTL |
| **PSEN1** | 0 | | No | 0.037 | 0.0388 | | 1.49e-02 | 4.28e-17 | — | — | — | — | SharePro | no colon eQTL |
| **BIN1** | 192 | 3.20e-22 | No | 0.0 | 1.0 | | | | — | — | — | — | SharePro, SMR/HEIDI | REQUIRES FOLLOW-UP: has colon eQTL but missing SharePro + SMR/HEIDI |

#### 2 | Per-method raw values (long format)

| gene | method | metric | value | tissue |
|---|---|---|---|---|
| **CR1** | GTEx | colon_n_eQTL | 24 | Colon (Trans+Sigmoid) |
| **CR1** | GTEx | colon_best_eQTL_p | 4.60e-07 | Colon (Trans+Sigmoid) |
| **CR1** | GTEx | AD_SNP_is_colon_eQTL | Yes | Colon (Trans+Sigmoid) |
| **CR1** | coloc-ABF | best_PP_H4 | 0.9925 | Colon (Trans+Sigmoid) |

| | | | | |
|---|---|---|---|---|
| **CR1** | coloc-ABF | best_PP_H3 | 0.3416 | Colon (Trans+Sigmoid) |
| **CR1** | SharePro | share_probability | 0.9991 | Colon (multi-causal) |
| **CR1** | SMR | best_SMR_p | 2.07e-06 | Colon (Trans+Sigmoid) |
| **CR1** | HEIDI | p_at_best_SMR | 0.00e+00 | Colon (Trans+Sigmoid) |
| **PICALM** | GTEx | colon_n_eQTL | 93 | Colon (Trans+Sigmoid) |
| **PICALM** | GTEx | colon_best_eQTL_p | 2.49e-09 | Colon (Trans+Sigmoid) |
| **PICALM** | GTEx | AD_SNP_is_colon_eQTL | No | Colon (Trans+Sigmoid) |
| **PICALM** | coloc-ABF | best_PP_H4 | 0.0001 | Colon (Trans+Sigmoid) |
| **PICALM** | coloc-ABF | best_PP_H3 | 0.9995 | Colon (Trans+Sigmoid) |
| **PICALM** | SharePro | share_probability | 0.9977 | Colon (multi-causal) |
| **PICALM** | SMR | best_SMR_p | 1.34e-03 | Colon (Trans+Sigmoid) |
| **PICALM** | HEIDI | p_at_best_SMR | 0.00e+00 | Colon (Trans+Sigmoid) |
| **CD2AP** | GTEx | colon_n_eQTL | 200 | Colon (Trans+Sigmoid) |
| **CD2AP** | GTEx | colon_best_eQTL_p | 1.56e-06 | Colon (Trans+Sigmoid) |
| **CD2AP** | GTEx | AD_SNP_is_colon_eQTL | Yes | Colon (Trans+Sigmoid) |
| **CD2AP** | coloc-ABF | best_PP_H4 | | Colon (Trans+Sigmoid) |
| **CD2AP** | coloc-ABF | best_PP_H3 | | Colon (Trans+Sigmoid) |
| **CD2AP** | SharePro | share_probability | | Colon (multi-causal) |
| **CD2AP** | SMR | best_SMR_p | | Colon (Trans+Sigmoid) |
| **CD2AP** | HEIDI | p_at_best_SMR | | Colon (Trans+Sigmoid) |
| **ADAM10** | GTEx | colon_n_eQTL | 345 | Colon (Trans+Sigmoid) |
| **ADAM10** | GTEx | colon_best_eQTL_p | 8.73e-10 | Colon (Trans+Sigmoid) |
| **ADAM10** | GTEx | AD_SNP_is_colon_eQTL | No | Colon (Trans+Sigmoid) |
| **ADAM10** | coloc-ABF | best_PP_H4 | 0.0004 | Colon (Trans+Sigmoid) |
| **ADAM10** | coloc-ABF | best_PP_H3 | 0.9996 | Colon (Trans+Sigmoid) |
| **ADAM10** | SharePro | share_probability | 0.0005 | Colon (multi-causal) |
| **ADAM10** | SMR | best_SMR_p | 4.00e-04 | Colon (Trans+Sigmoid) |
| **ADAM10** | HEIDI | p_at_best_SMR | 0.00e+00 | Colon (Trans+Sigmoid) |
| **APP** | GTEx | colon_n_eQTL | 28 | Colon (Trans+Sigmoid) |
| **APP** | GTEx | colon_best_eQTL_p | 5.99e-07 | Colon (Trans+Sigmoid) |
| **APP** | GTEx | AD_SNP_is_colon_eQTL | No | Colon (Trans+Sigmoid) |
| **APP** | coloc-ABF | best_PP_H4 | 0.0982 | Colon (Trans+Sigmoid) |
| **APP** | coloc-ABF | best_PP_H3 | 0.9621 | Colon (Trans+Sigmoid) |
| **APP** | SharePro | share_probability | | Colon (multi-causal) |
| **APP** | SMR | best_SMR_p | 1.48e-02 | Colon (Trans+Sigmoid) |
| **APP** | HEIDI | p_at_best_SMR | 1.47e-129 | Colon (Trans+Sigmoid) |
| **SORL1** | GTEx | colon_n_eQTL | 0 | Colon (Trans+Sigmoid) |
| **SORL1** | GTEx | colon_best_eQTL_p | | Colon (Trans+Sigmoid) |
| **SORL1** | GTEx | AD_SNP_is_colon_eQTL | No | Colon (Trans+Sigmoid) |
| **SORL1** | coloc-ABF | best_PP_H4 | 0.0514 | Colon (Trans+Sigmoid) |
| **SORL1** | coloc-ABF | best_PP_H3 | 0.1161 | Colon (Trans+Sigmoid) |
| **SORL1** | SharePro | share_probability | | Colon (multi-causal) |
| **SORL1** | SMR | best_SMR_p | 4.22e-01 | Colon (Trans+Sigmoid) |
| **SORL1** | HEIDI | p_at_best_SMR | 2.61e-43 | Colon (Trans+Sigmoid) |
| **PSEN1** | GTEx | colon_n_eQTL | 0 | Colon (Trans+Sigmoid) |
| **PSEN1** | GTEx | colon_best_eQTL_p | | Colon (Trans+Sigmoid) |
| **PSEN1** | GTEx | AD_SNP_is_colon_eQTL | No | Colon (Trans+Sigmoid) |
| **PSEN1** | coloc-ABF | best_PP_H4 | 0.037 | Colon (Trans+Sigmoid) |
| **PSEN1** | coloc-ABF | best_PP_H3 | 0.0388 | Colon (Trans+Sigmoid) |
| **PSEN1** | SharePro | share_probability | | Colon (multi-causal) |
| **PSEN1** | SMR | best_SMR_p | 1.49e-02 | Colon (Trans+Sigmoid) |
| **PSEN1** | HEIDI | p_at_best_SMR | 4.28e-17 | Colon (Trans+Sigmoid) |
| **BIN1** | GTEx | colon_n_eQTL | 192 | Colon (Trans+Sigmoid) |
| **BIN1** | GTEx | colon_best_eQTL_p | 3.20e-22 | Colon (Trans+Sigmoid) |
| **BIN1** | GTEx | AD_SNP_is_colon_eQTL | No | Colon (Trans+Sigmoid) |
| **BIN1** | coloc-ABF | best_PP_H4 | 0.0 | Colon (Trans+Sigmoid) |

| | | | | |
|---|---|---|---|---|
| **BIN1** | coloc-ABF | best_PP_H3 | 1.0 | Colon (Trans+Sigmoid) |
| **BIN1** | SharePro | share_probability | | Colon (multi-causal) |
| **BIN1** | SMR | best_SMR_p | | Colon (Trans+Sigmoid) |
| **BIN1** | HEIDI | p_at_best_SMR | | Colon (Trans+Sigmoid) |

#### Notes

- **Thresholds.** coloc-ABF PP.H4 ≥ 0.8 = strong shared-causal; SharePro share ≥ 0.5 = multi-causal shared signal; SMR p < 0.0042 (Bonferroni on 12 tests); HEIDI p > 0.05 = pass (single-variant model consistent).
- **Para 5 S5 verification.** PICALM SharePro share = 0.9977 (matches 0.998 in paragraph). CR1 SharePro share = 0.9991 (matches 0.999 in paragraph). ADAM10 SharePro share = 0.0005 → multi-causal NOT supported (matches paragraph).
- **Para 5 S6 verification.** CD2AP has GTEx colon n_eQTL = 200, AD_SNP_is_colon_eQTL = Yes, but SharePro analysis was not run for CD2AP in this work (file CD2AP_v3_result.sharepro.txt does not exist). Verdict: 'REQUIRES FOLLOW-UP'. This is the gene called out by the paragraph as needing additional multi-causal colocalization work.
- **Why ADAM10 fails SharePro despite high SMR.** ADAM10 has SMR p = 4.0e-04 (Bonferroni-significant) but HEIDI p ≈ 0 (rejects single-variant model). SharePro share = 0.0005 (no shared causal under multi-causal model). Interpretation: ADAM10 has eQTL + GWAS signal but the variants are NOT shared; SMR captures a linkage artifact that HEIDI correctly flags.
- **Final causal-candidate ranking (Colon).** GOLD (coloc + SMR + HEIDI all pass): none. SILVER (coloc OR SharePro shared-causal): CR1 (both methods), PICALM (SharePro rescue). INCOMPLETE: CD2AP (SharePro not yet run). DROP: ADAM10 (multi-causal contradicts shared-causal interpretation). NO COLON EQTL: SORL1, PSEN1 (zero significant colon variants in GTEx).
- **How to extend.** To run SharePro for CD2AP: place GWAS + eQTL summary statistics + LD matrix in results/sharepro/CD2AP_v3_{gwas,eqtl,ld}.txt, then run the SharePro CLI as in `notebooks/21_eqtl_coloc_analysis.ipynb`. Output will be results/sharepro/CD2AP_v3_result.sharepro.txt and this builder will pick it up automatically on re-run.

#### Data sources

| data_source | citation | accession | url | version_snapshot | derived_path_in_repo | notes |
|---|---|---|---|---|---|---|
| **GTEx Portal v8 eQTL query** | GTEx Consortium Science 2020 | v8 | https://gtexportal.org/home/ | queried 2026-04 (snapshot) | results/gtex_eqtl_query_results.csv | Per (gene × tissue) significant cis-eQTL counts + best p + AD-SNP-is-eQTL flag |
| **Bellenguez 2022 AD GWAS** | Bellenguez et al. Nat Genet 2022 | GCST90027158 | https://www.ebi.ac.uk/gwas/studies/GCST90027158 | 2022-04 (GRCh38) | data/gwas/Bellenguez2022_AD_withN.tsv.gz (gitignored) | AD GWAS lead SNPs per locus |
| **coloc-ABF (full summary stats)** | Giambartolomei et al. PLoS Genet 2014 | - | https://github.com/chr1swallace/coloc | coloc v5.x | results/coloc_FULL_eqtl_results.csv | Single-causal coloc using full eQTL summary stats; PP.H0–H4 per (gene × colon tissue) |
| **SharePro multi-causal coloc** | Zhang et al. Genome Biol 2023 | - | https://github.com/zhwm/SharePro_coloc | SharePro v5.0.0 | results/sharepro/{GENE}_v3_result.sharepro.txt | Multi-causal shared-component coloc; produces "share" |

| | | | | | | |
|---|---|---|---|---|---|---|
| | | | | | | probability per credible set |
| **SMR + HEIDI** | Zhu et al. Nat Genet 2016 | - | https://yanglab.westlake.edu.cn/software/smr/ | SMR v1.3.1 | results/smr_colon_AD_results.csv | Summary-data-based Mendelian Randomization with HEIDI heterogeneity test |
| **Pipeline notebook** | This work | - | - | - | notebooks/21_eqtl_coloc_analysis.ipynb (+ _executed) | Orchestrates GTEx query + coloc-ABF + SharePro + SMR for the 8 focal genes |
| **Comparison data: sig-only vs full** | Supplementary Fig (Para 5 S4) | - | - | - | results/coloc_sig_vs_full_comparison.csv | Sister CSV showing PP.H4 drop when switching from sig-only to full summary |
